# A Lipid Atlas of the Human Kidney

**DOI:** 10.1101/2022.04.07.487155

**Authors:** Melissa A. Farrow, Léonore E.M. Tideman, Elizabeth K. Neumann, Nathan Heath Patterson, Lukasz G. Migas, Madeline E. Colley, Jamie L. Allen, Emilio S. Rivera, Carrie E. Romer, Haichun Yang, Maya Brewer, Ellie Pingry, Martin Dufresne, Katerina Djambazova, Kavya Sharman, Angela R.S. Kruse, Danielle B. Gutierrez, Raymond C. Harris, Agnes B. Fogo, Mark P. de Caestecker, Richard M. Caprioli, Raf Van de Plas, Jeffrey M. Spraggins

**Affiliations:** Department of Biochemistry, Vanderbilt University, Nashville, TN, USA 37232; Mass Spectrometry Research Center, Vanderbilt University, Nashville, TN, USA 37232; Delft Center for Systems and Control, Delft University of Technology, 2628 CD Delft, The Netherlands; Department of Cell and Developmental Biology, Vanderbilt University, Nashville, TN, USA 37232; Division of Nephrology and Hypertension, Department of Medicine, Vanderbilt University Medical Center, Nashville, TN USA 37232; Department of Pathology, Microbiology and Immunology, Vanderbilt University Medical Center, Nashville, TN USA 37232; Department of Medicine, Vanderbilt University Medical Center, Nashville, TN, USA 37232; Department of Pharmacology, Vanderbilt University, Nashville, TN, USA 37232; Department of Chemistry, Vanderbilt University, Nashville, TN, USA 37232

## Abstract

Tissue atlases provide foundational knowledge on the cellular organization and molecular distributions across molecular classes and spatial scales. Here, we construct a comprehensive spatio-molecular lipid atlas of the human kidney from 29 donor tissues using integrated multimodal molecular imaging. Our approach leverages high spatial resolution matrix-assisted laser desorption/ionization (MALDI) imaging mass spectrometry (IMS) for untargeted lipid mapping, stained microscopy for histopathological assessment, and tissue segmentation using autofluorescence microscopy. With a combination of unsupervised, supervised, and interpretive machine learning, the atlas provides multivariate lipid profiles of specific multicellular functional tissue units (FTUs) of the nephron, including the glomerulus, proximal tubules, thick ascending limb, distal tubules, and collecting ducts. In total, the atlas consists of tens of thousands of FTUs and millions of mass spectrometry measurements. Detailed patient, clinical, and histopathologic information allowed molecular data to be mined based on these features. As examples, we highlight the discovery of how lipid profiles are altered with sex and differences in body mass index.

## INTRODUCTION

The human kidney is a highly organized organ responsible for filtering waste products from the blood, maintaining ion and fluid balance, releasing hormones that regulate blood pressure, and controlling the production of red blood cells. Filtration and maintenance of ion and water balance are achieved in the nephron, which is vital for maintaining homeostasis. Each human kidney has approximately one million nephrons comprised of distinct multicellular functional tissue units (FTUs) surrounded by a network of capillaries. FTUs of the nephron include the glomerulus, where blood filtration and formation of ultrafiltrate is initiated, as well as a series of tubules that are comprised of proximal convoluted tubules, the loop of Henle, distal convoluted tubules, and collecting ducts. Each renal FTU specializes in balancing ion and nutrient concentration and osmolality of the urine by mechanisms of secretion and absorption.

Fundamental to comprehending the architectural and functional complexity of an organ is determining the organization of cell types and FTUs and cataloging the localized molecular profiles of these tissue features. Model systems, such as cell cultures and organoids, have provided insight into aspects of tissue function, but not necessarily into the spatial anatomical and cellular organization of tissue. Animal models are better suited to yield the latter, but may not fully recapitulate human physiology. Therefore, recently there has been a push to develop cellular atlases of human tissue instead. Nevertheless, such studies have primarily focused on transcriptional data and with limited spatial context^1–3^. To address these gaps, multiple large-scale research consortia have been established to develop and deploy spatial technologies for deep molecular profiling and mapping of the transcriptome, proteome, lipidome, and metabolome of human tissues^4^. Comprehensive spatio-molecular atlases offer a means to generate new hypotheses and advance biomedical research by providing an unprecedented view into tissue at cellular resolution^5–7^. Exploring relationships between cellular and molecular organization of tissues enables the discovery of underlying drivers of functional efficiency, transition to disease, and disease severity in function of key patient factors (*e.g.,* age, sex, race, comorbidities). Recently, the National Institutes of Health and private organizations have funded various atlas efforts to address this grand challenge, such as the Human Biomolecular Atlas Program (HuBMAP)^4,7^, Allen Brain Atlas^8^, Human Cell Atlas (HCA)^9^, BRAIN Initiative^10^, Kidney Precision Medicine Project^11^, and Human Tumor Atlas Network^12^, targeting an array of normal and/or diseased organs.

Integration of multimodal molecular imaging technologies can provide a more rounded, systems biology view of organ function (and dysfunction) across a wide range of molecular classes (e.g., lipids, metabolites, proteins, and RNA) and spatial scales (e.g., whole organs to single cells). The Human Biomolecular Atlas Program is a consortium funded by the National Institutes of Health that is leading research efforts in this area with the goal of creating an open, comprehensive molecular atlas of the human body at cellular resolution^4,7^. Here, as part of HuBMAP, we have developed an FTU-specific lipidomic atlas of the human kidney to characterize the molecular organization of the nephron in normal-appearing tissue. This atlas incorporates untargeted imaging mass spectrometry (IMS) data^13,14^ that is fully integrated with various forms of microscopy^15,16^ to enable the discovery of spatially specific molecular marker candidates for key patient factors such as obesity and sex. Each component of the atlas is publicly available (https://portal.hubmapconsortium.org/). This atlas spans multiple scales of spatial granularity, ranging from large tissue areas including multiple anatomical regions to specific functional tissue units and cell types and from averaged populations to individual subjects, serving as a resource for exploring the biomolecular landscape that informs renal function and disease. To punctuate the potential of this resource, we demonstrate how our integration of IMS and microscopy can be used to reveal spatio-molecular relationships that are difficult to access through other means, and how these unique insights can yield key lipidomic marker candidates for, *e.g.*, obesity for various functional components of the nephron. Although this work is focused on the human kidney, the analytical and computational pipelines we have developed as part of our atlasing efforts have broad utility and are applicable to any organ system or disease type. Together, these developed pipelines provide critical insight into how molecules and cells are organized into multicellular functional components that work in concert to drive macroscopic tissue function.

## RESULTS

We have collected nephrectomies from 29 donors (**Table S1**) and analyzed these tissues with a customized workflow that integrates pathological assessment and image segmentation on the basis of autofluorescence (AF) microscopy with high spatial resolution matrix-assisted laser desorption/ionization (MALDI) imaging mass spectrometry (IMS) and unsupervised and supervised machine learning approaches that exploit the spatial co-registration of the different data types (**Figure 1**). The spatial match achieved by acquisition of microscopy images and IMS data from the same tissue section outperforms what would be possible with measurements from subsequent tissue sections. Whole slide microscopy images were segmented into glomeruli, proximal tubule, thick ascending limb, distal tubule, and collecting duct features, and spatially associated untargeted IMS data were mined to discover FTU-specific lipid profiles. This approach, for the first time, provides insight into the FTU-level organization of the kidney at the molecular level. **Figure 2A.1-3** shows example whole slide microscopy images collected from a selected donor (56-year-old white female). Autofluorescence microscopy (**Figure 2A.1**) was utilized to drive image co-registration and FTU segmentation (**Figure 2A.2**). PAS-stained images (**Figure 2A.3** and **figure S1**) were collected after MALDI IMS acquisition for histological assessment of overall specimen composition (*i.e.,* % cortex vs. medulla), tubular atrophy, glomerular sclerosis, and measures of other key tissue features and pathologies (**table S2**). Autofluorescence images (**figure S2**), acquired prior to MALDI IMS, were used to automatically segment the FTUs of the nephron, including glomeruli, proximal tubules, thick ascending limb, distal tubules, and collecting ducts (**figure S3**) as described previously^17^. MALDI IMS measurement regions were selected to capture a mixture of FTUs and anatomical regions in both positive and negative ion modes (**Figure 2A.4-6**). It is noted that both polarities are collected from each tissue section. Examples of IMS-provided lipid distribution images are shown in **Figure 2B.1-5**. All MALDI IMS data in the atlas were collected with a pixel size of 10 µm which, together with multimodal acquisition from the same tissue section, provided sufficient spatial specificity and accuracy to associate mass spectrometry (MS) signals with specific FTUs *in situ*. For example, the sphingomyelin SM(34:1);2O (*m/z* 687.545) was found to localize to the glomeruli and other surrounding structures (**Figure 2B.1**). The phosphatidylethanolamine PE(36:4) (*m/z* 738.508) localized primarily to the proximal tubules (**Figure 2B.2**), whereas PE(36:1) (*m/z* 744.555) was found with greater abundance in the collecting ducts (**Figure 2B.3**). The substantial variation in distribution between these latter two lipids is particularly notable given that they only differ by the number of double bonds in their fatty acyl tails. Finally, the sulfatide SHexCer(42:1);3O (*m/z* 906.635) was found specifically in the thick ascending limb (**Figure 2B.4**). The overlay of these four lipid species (**Figure 2B.5**) highlights the localized diversity of these molecules and their ability to differentiate FTUs as compared to complimentary microscopy. Besides a spatial view into a selection of the hundreds of lipid species recorded, the atlas can also yield spectrum-wide molecular signatures for each FTU. One approach is demonstrated in **Figure 2B.6**, where cohort-wide average mass spectra specific for each FTU type are provided. Using the automated AF microscopy-based FTU segmentations and the common spatial coordinate system as a guide, MALDI IMS mass spectra (*i.e.,* IMS pixels) specific to each FTU type can be collected at scale from across all tissues in the atlas. These FTU-specific mass spectra are then combined to provide donor cohort-wide average mass spectra for each FTU in both negative (**figure S4 and S5**) and positive (**figure S6 and S7**) ion modes. The negative ion mode average mass spectra for each FTU are also shown in **Figure 2B.6** for a selected mass window (*m/z* 650-900). While subtle changes in ion intensity or in the presence of low abundant molecular species can be difficult to discern in average spectra, the manner in which average spectra diverge between FTUs is more readily apparent when these data are shown as difference spectra (**figure S8-27)**. For example, the difference spectrum comparing glomeruli to proximal tubules in negative ion mode shows variations in lipid ion intensities between these two critical structures of the nephron more clearly (**Figure 2B.7**).

**Figure 1.**
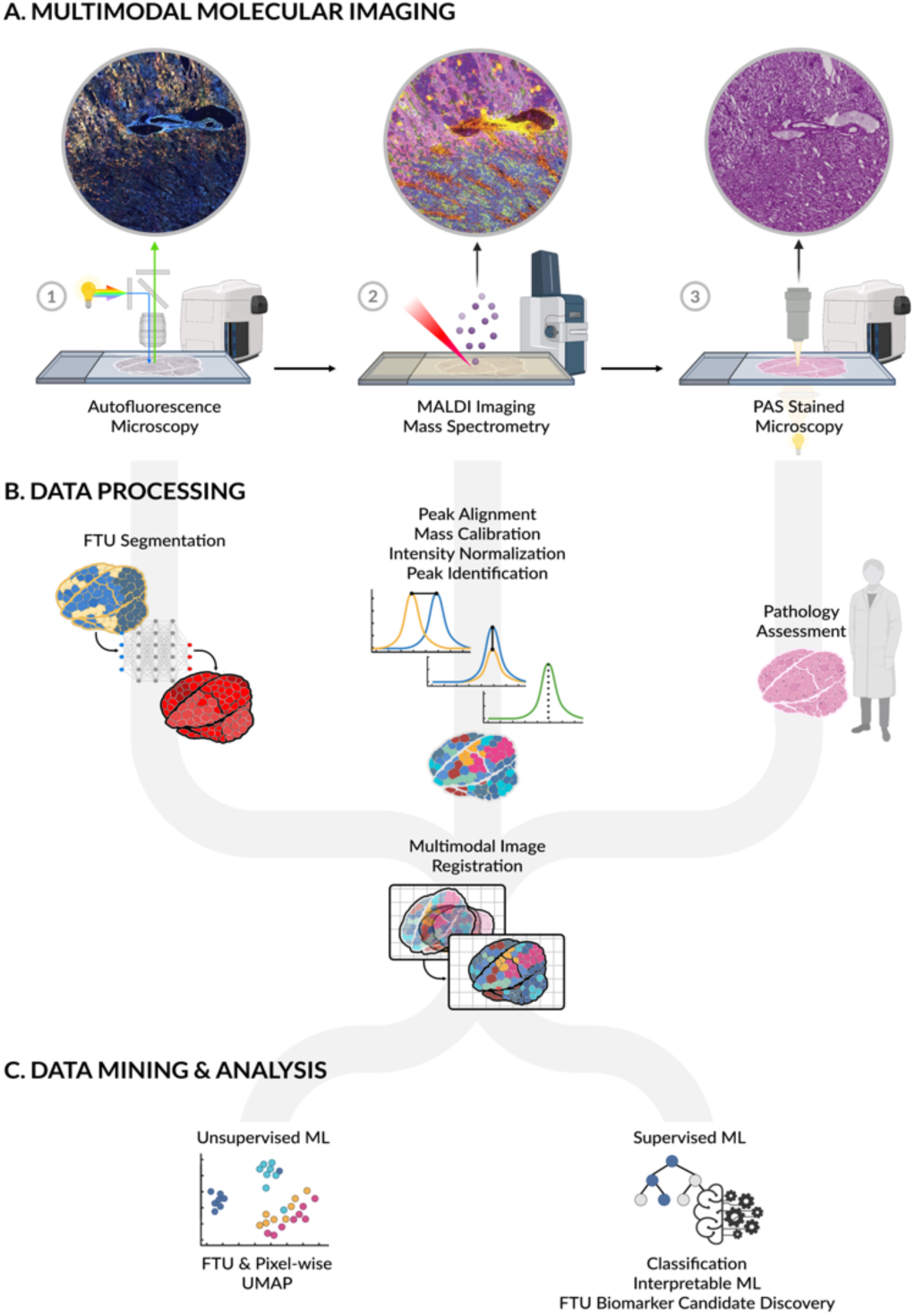
Construction of the FTU lipid atlas of the human kidney. Multimodal molecular imaging data were collected from 28 donor kidney tissues. Each tissue section was subjected to autofluorescence microscopy (**1**), MALDI imaging mass spectrometry (**2**), and PAS-stained microscopy (**3**). Each modality is processed individually, resp. to provide FTU segmentations, to ensure MALDI IMS measurements are comparable and to remove potential batch effects, and to provide histopathological assessment of each tissue. These datasets are then integrated by spatially co-registering them onto the same spatial coordinate system and by performing a combination of unsupervised and cross-modal supervised machine learning analyses. Interpretable machine learning is then used to uncover spatially distinct biomarker candidates for FTUs across the overall data cohort as well as scoped to specific donor metadata such as body mass index or sex.

**Figure 2.**
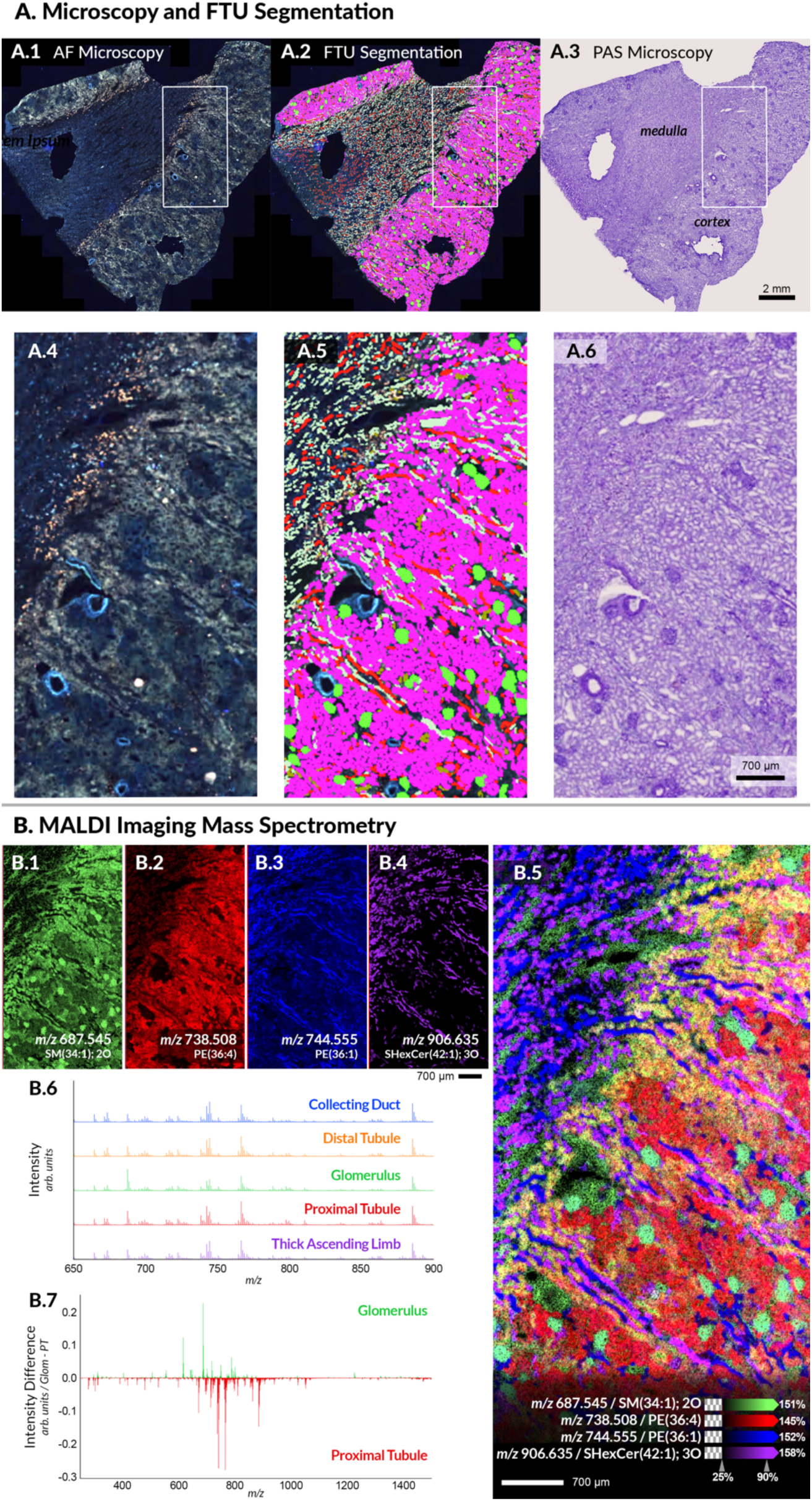
Example of multimodal molecular characterization of human kidney tissue. Whole slide microscopy images from donor VAN0028 (56-year-old white female) were collected using autofluorescence (**A.1 and A.2**) prior to high spatial resolution (10 µm pixel size) MALDI IMS measurement and PAS-stained microscopy (**A.3**) was acquired post-IMS. AF microscopy data was automatically segmented into renal FTUs (**A.2**), including the glomeruli (green), proximal tubules (magenta), thick ascending limb (light green), distal tubules (brown), and collecting ducts (red). MALDI IMS measurement regions (white boxes) were selected to include a mixture of tissue features. The microscopy data for the MALDI IMS measurement region are highlighted in **Panels A.4-6**. Selected individual molecular distribution images from the negative ion mode MALDI IMS measurement and an overlay image are provided in **Panels B.1-4 and B.5**, respectively. The selected ions demonstrate unique localizations within the kidney, without the need for prior labeling, and these are just four of the hundreds of lipids that make up this molecular atlas. The mean mass spectrum associated with each FTU, obtained by averaging FTU-specific IMS-pixels across all donors, displays subtle differences in the lipids detected and their intensities (**B.6**). It is noted that full size versions of the spectra can be found in the supplemental materials. Variations in intensity profiles of FTUs are more evident using difference spectra, as shown in the comparison between the normalized (between 0 and 1 to allow for direct comparison) average spectrum of proximal tubules subtracted from that of the glomeruli (**B.7**).

### Characteristics of the Donor Tissue Cohort

The molecular atlas was constructed with normal portions of fresh-frozen renal cancer nephrectomy tissue (*i.e.,* remnant tissue) collected from 29 human donors (**table S1**). All tissues were collected through the Cooperative Human Tissue Network at Vanderbilt University Medical Center. Warm and cold ischemia times were shorter than 5 min and 30 min, respectively. **Figure 3A** highlights the distributions of sex, age, and body mass index (BMI) of the donor cohort. The cohort includes 14 female and 15 male donors with an age distribution ranging from 20 to 78 years and an average age of 60. The BMI values of the included donors were similar to averages for the population of the United States^18,19^, ranging from 22.3 to 45.5 with an overall average of 30.6, and averages of 31.8 and 29.5 for female and male donors, respectively. It is noted that the cohort used for this study is limited based on tissues available and currently only includes non-Hispanic white individuals. This limitation will be specifically addressed in future versions of the atlas.

**Figure 3.**
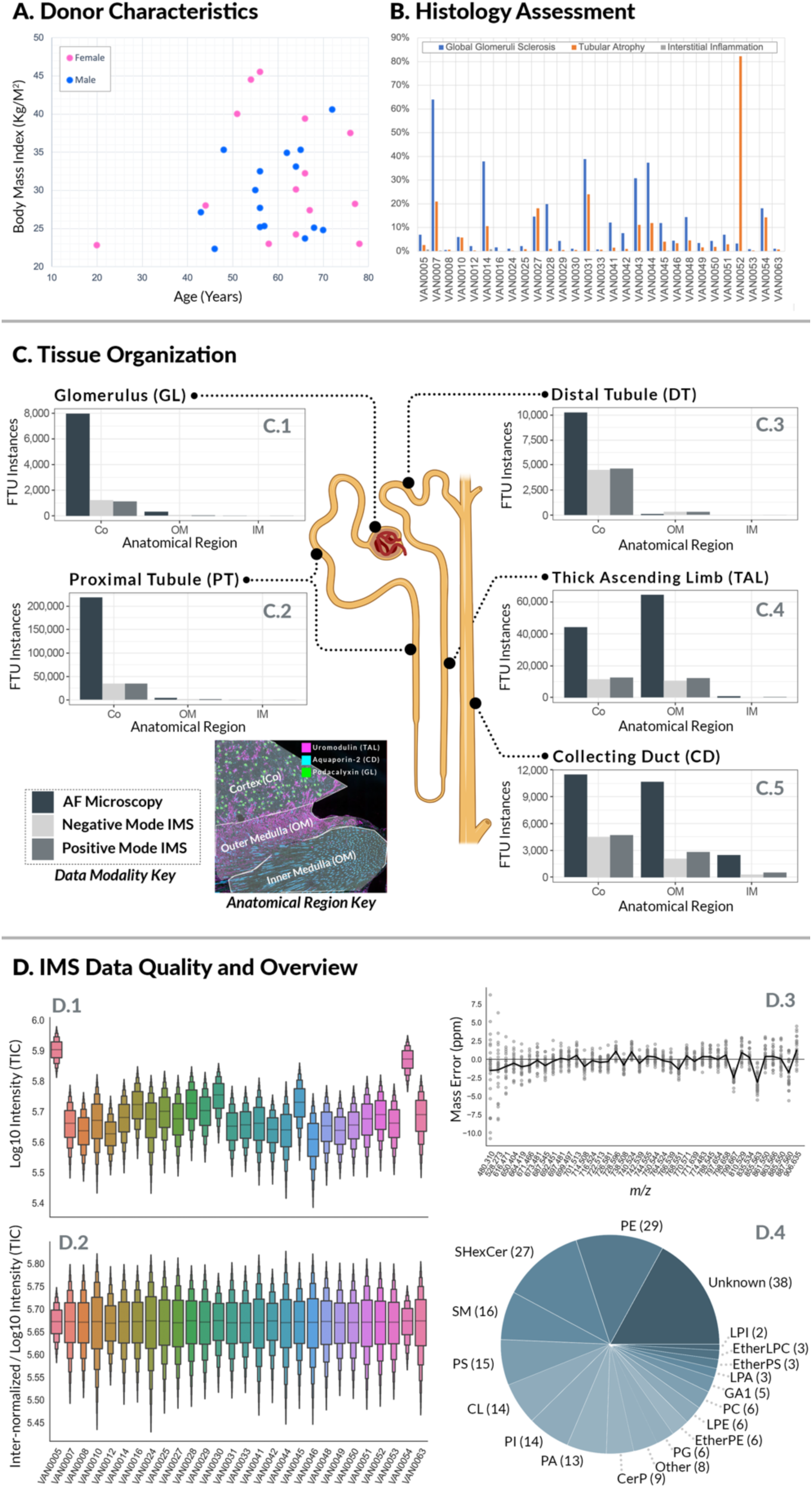
Global Characteristics of the Atlas. The atlas was constructed using tissues from 29 adult human donors, for which key donor characteristics such as age and BMI (**A**) and detailed histopathology assessments (**B**) are available. Selected measures of tissue normalcy are highlighted in the bar plot, including % global glomeruli sclerosis, % interstitial fibrosis tubular atrophy in cortex, and % interstitial inflammation in the cortex. To delineate critical components of the nephron and support mining of MALDI IMS data, autofluorescence microscopy was used to comprehensively segment renal functional tissue units such as the glomerulus (**C.1**), proximal tubule (**C.2**), thick ascending limb of the loop of Henley (**C.4**), distal tubule (**C.3**), and collecting duct (**C.5**). The total number of detected instances for each FTU was quantified within the cortex (Co), outer medulla (OM), and inner medulla (IM) of the kidney across all autofluorescence whole slide images (dark grey) and specific to the MALDI IMS measurement regions in both negative (light grey) and positive ion (medium grey) modes. The example immunofluorescence data shows how markers (*e.g.,* uromodulin-pink, Aquaporin-2-pink, and podocalyxin-green) used to train the AF-based segmentation algorithms were also able to differentiate the broader anatomical zones of the kidney (*i.e.,* Co, OM, and IM). The integrated MALDI IMS data underwent data preprocessing to address non-biological variability, including peak alignment, calibration, and intensity normalization. Boxen plots show the variability of the total ion current (Log10 intensity) for the negative ion mode data following intra-sample normalization (**D.1**) and the consistency after inter-sample normalization (D.2). Panel **D.3** provides the mass error in parts per million (ppm) for selected negative ion mode lipids. The black line represents the mean mass error from all pixels collected from all donor samples, and the grey dots represent the spread of the data for each *m/z*. Following preprocessing, m/z features are annotated, using mass accuracy to compare to LC-MS/MS-based identifications and on-tissue fragmentation. The provided pie chart summarizes the number of annotations for various lipid classes from the negative ion mode data (**D.4**). It is noted that 28 of the 29 samples were analyzed in negative ion mode.

Although it is difficult to obtain tissues from completely normal individuals and most adult tissues display some pathological features, histopathological analysis by expert clinical pathologists determined that all tissues included in the atlas were age-appropriate normal tissues (**table S2**). PAS-stained images show no clear indication of cancer or immune cell infiltration. Tissues were assessed for anatomical composition (i.e., percentage of cortex and medulla) to ensure all key renal FTUs are represented. Tissue normalcy was determined using measures of interstitial fibrosis tubular atrophy, arteriosclerosis, global glomeruli sclerosis, arteriolar hyalinosis, and interstitial inflammation. The frequency of key pathologies determined for each donor tissue is summarized in **Figure 3B**. As expected, the levels of tissue pathologies vary greatly in human subjects, yet our analytical and computational pipelines were still able to uncover lipidomic profiles for specific FTUs that were consistent across the entire atlas donor cohort.

### Functional Tissue Unit Frequency and Distribution

Multimodal analysis of renal tissue provides a means of connecting rich molecular information from imaging mass spectrometry with relevant tissue cell types and structures when integrated with complementary microscopy and the derived functional tissue unit segmentation masks (**figure S3**). By applying the multimodal collection pipeline to 29 donors, FTU segmentation labels and molecular profiles (*i.e.*, mass spectra) on the order of millions of data points are obtained and tied to specific spatial tissue areas with high fidelity. **Figure 3C** shows the number of instances (discrete FTUs) found across the AF microscopy images overall as well as specific to the MALDI IMS measurement regions. Each donor sample has similar numbers of FTU instances per class, but relative counts are influenced by the gross morphology of each section, which was calculated as the percent of cortex or medulla in a section during pathological assessment^17^ (**table S2**). FTU instances were distributed as expected, with glomeruli and proximal tubules almost exclusive to the cortex (**Figure 3C.1 and 3C.2**, respectively), thick ascending limb segments found in the outer medulla and cortex (**Figure 3C.4**), distal tubules in the cortex (**Figure 3C.3**), and collecting ducts found both in medullary and cortical regions (**Figure 3C.5**). In terms of robustness of the atlas and the FTU-specific molecular signatures provided by MALDI IMS, it is critical to note that the analyses highlighted herein are based on thousands of individual FTU instances for each targeted kidney structure, together with millions of mass spectrometry measurements that are co-registered and integrated with them. For example, positive ion mode MALDI IMS measurements from all 29 donors amassed 6,779,166 mass spectra (individual IMS pixels). From this, approximately 68.8% (4,662,712 pixels) were spatially annotated into the five targeted functional tissue units: glomeruli (GL), proximal tubules (PT), thick ascending limb (TAL), distal tubules (DT), and collecting ducts (CD). Similarly, in negative ionization mode, 6,568,017 mass spectra were collected from 29 of the donor samples, with 70.8% assigned to FTUs. The fraction of MALDI IMS data that could be labeled as part of targeted FTUs is as expected. The unlabeled fraction of the data is partially due to how the segmentation models were constructed, namely with an emphasis on avoiding false positives. Given the number of instances available, for integration with MALDI IMS data it is less important to detect every last FTU than it is to be sure that the FTU label assignments are of high confidence. Therefore, the resulting segmentation masks label rather conservatively and only high-confidence FTUs so that only high-quality data are included in subsequent IMS data mining and modeling approaches. Furthermore, the current version of the kidney lipid atlas does not include tissue features such as vasculature and extracellular matrix, which also account for a nontrivial tissue area. More granular delineation of the nephron and the addition of these other key tissue features will be the focus of future extensions of the atlas.

### Imaging Mass Spectrometry Data Quality and Annotations

Ensuring MALDI IMS data quality was essential when constructing the kidney FTU lipid atlas. Extensive quality control procedures and data preprocessing were performed to minimize non-biological variability and to ensure comparability of the data. Peak alignment and calibration were critical for enabling high-confidence annotation of IMS data. IMS peak intensity normalization was performed both on a per-sample basis (*i.e.,* intra-sample normalization) and across the entire data cohort (*i.e.,* inter-sample normalization). Following intra-sample normalization, the sample-level intensity variance across the entire cohort was ±8% (**Figure 3D.1, figure S28**) and ±9% (**figure S31**) for negative and positive ion modes, respectively. This was improved upon further with inter-sample normalization reducing the global intensity variance for negative ion mode to ±2% (**Figure 3D.2, figure S28**) and ±3% (**figure S31)** for positive ion mode. A combination of LC-MS/MS and on-tissue fragmentation (**table S4**) was used to identify lipids from serial tissue sections from each of the donor samples. Although thousands of MS features were detected, the current atlas is built around 212 (negative ion mode) and 211 (positive ion mode) peaks that were either annotated or that remain unidentified at the moment, but that do exhibit a high S/N and that are clearly distinguishable from the isotopic envelopes of other annotated molecular species. All IMS peak annotations were made based on mass accuracy with a threshold of better than 5 parts per million (ppm) in order to link to the theoretical *m/z* of the lipids identified by MS/MS. Plots are provided to highlight the consistency of the data, showing the spread of the mass error for selected negative (**Figure 3D.3, figure S29**) and positive (**figure S30**) ion mode lipids. The MALDI IMS analysis detected a wide range of lipid classes. Negative ion mode data included annotations and images from phosphatidylethanolamines (PE), sulfatides (SHexCer), sphingomyelins (SM), phosphatidylserines (PS), cardiolipins (CL), phosphatidylinositols (PI), and others (**Figure 3D.4, figure S32A**). In positive ion mode, the atlas includes lipid classes such as phosphatidylcholines (PC), sphingomyelins (SM), ether-linked PCs (etherPC), and non-polar lipids such as diglycerides (DG) and monoglycerides (MG), among others (**figure S32B**). Full datasets are available as part of the atlas, and additional features can be added to future analyses if other ions are identified or found to be important.

### Unsupervised Analysis

IMS measurements were acquired in both negative and positive ionization mode, yielding two data cohorts. All mass spectra within a cohort (one mass spectrum per IMS-pixel) have been preprocessed to remove non-biological variation where possible and to enable ion intensity values to be compared across experiments. This preprocessing phase includes m/z alignment, m/z calibration, ion intensity normalization, and peak picking, and it is complemented with donor-specific variation removal by means of reComBat^20^. Each cohort dataset consists of all IMS-pixels (i.e., spectra) collected across donor tissue samples and uses the same set of peak-picked features throughout. Specifically, the negative ionization mode cohort dataset entails 6,568,017 pixels, each reporting a vector of 212 features. The features correspond to peak intensities of a selection of 212 lipid species (**table S4**) that were chosen for identifiability and broad coverage from all peaks reported by the full profile mass spectra acquired.

Initial insight into the variation captured by this dataset was obtained through an unsupervised machine learning approach, Uniform Manifold Approximation and Projection (UMAP)^21^ with a focus on exploring the structure underlying the measurements. This resulted in the dimensionality-reduced representation of the spectral data in **Figure 4**. The latent space representation of chemical variation (irrespective of donor origin) is shown color-coded for donor origin (**Figure 4A**) and for FTU type (**Figure 4C**). Using the cosine distance, this visualization captures primarily overall variation across all 212 measured features. The presence of donor-specific variation is apparent in **Figure 4A**, where certain donor-specific measurement sets tend to cluster away from other measurements. However, **Figure 4C** shows that, besides donor-specific variation, FTU-specific variation is also clearly present in the data. Specifically, subsets of measurements that share FTU membership tend to cluster together regardless of their donor origin (*e.g.,* note proximal tubule concentration in the bottom right corner of **Figure 4C**). **Figures 4A** and **4C** suggest that the measurements collected as part of this atlas effectively capture different types of biological variation. Furthermore, since a rather low-dimensional representation is pursued here, it is unsurprising that the resulting manifold tends to be dominated by the more prominent variabilities captured by the data, which include donor-related and FTU-related chemical variation.

**Figure 4.**
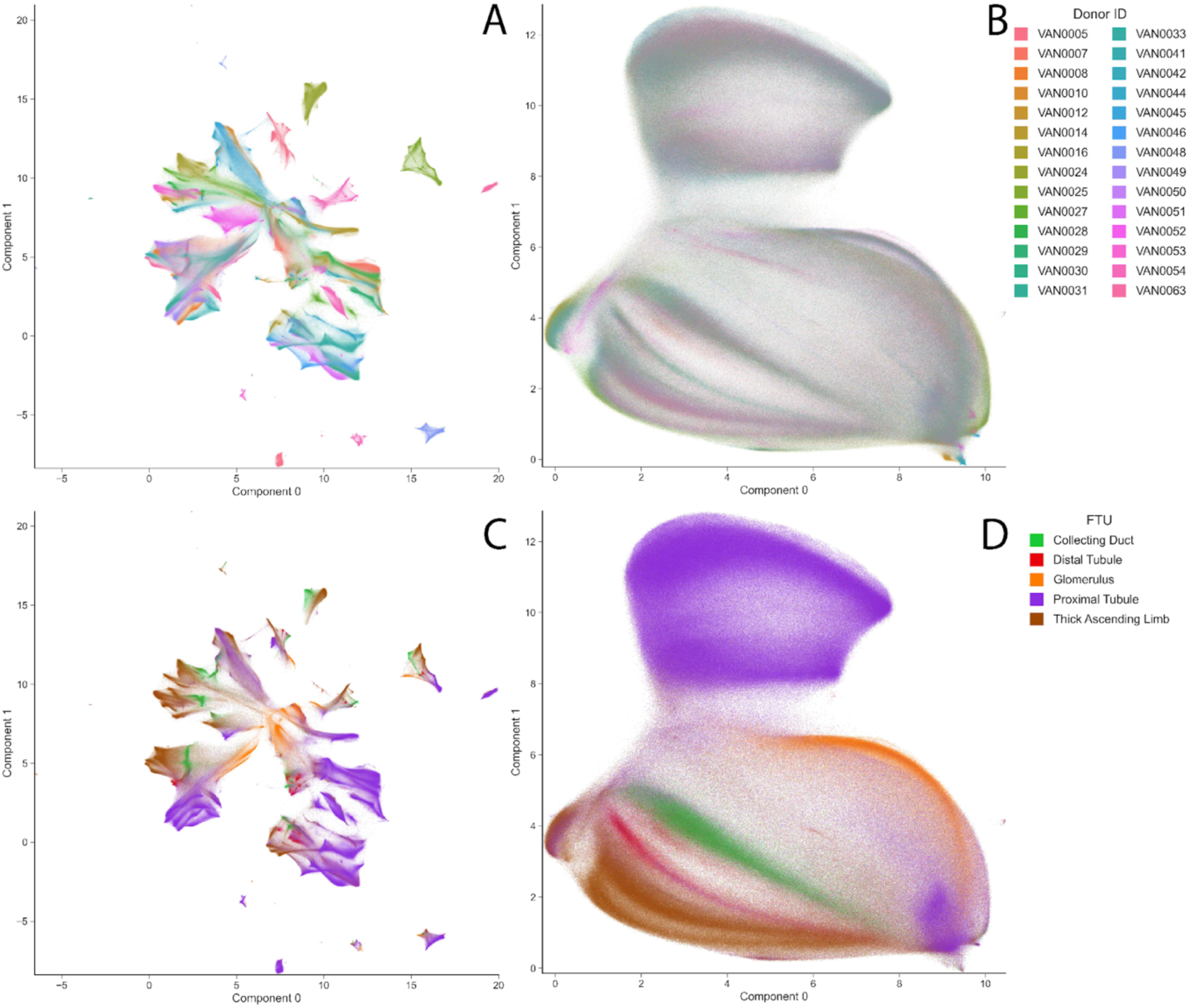
Pixel-level chemical variation MALDI IMS measurements across 28 of the donors analyzed by negative ion mode. Two-dimensional visualization of chemical variation in the negative ionization mode experiment cohort, using UMAP to cast a matrix of 6,568,017 observations (i.e., IMS pixels across 28 donor tissues) by 212 features (i.e., lipid species) into a table of 6,568,017 observations by 2 latent variables, while retaining neighborhood relationships between observations as captured by a cosine distance measure. (**A**) Latent space representation of chemical variation after preprocessing, with pixels color-coded for donor origin. (**C**) Same latent space representation as in panel A, with pixels color-coded for FTU type (as automatically recognized from microscopy). (**B**) Latent space representation of chemical variation after preprocessing and reduction of donor variation by reComBat, with pixels color-coded for donor origin. (**D**) Same latent space representation as in panel B, with pixels color-coded for FTU type (as automatically recognized from microscopy). Note, while reComBat has not been optimized for use on mass spectrometry data, it is applied here to demonstrate that if donor-specific variation can be removed, FTU-related variation becomes more readily discernable in an unsupervised context.

To further our unsupervised exploration towards FTU-specific chemical differentiation, we removed a substantial part of the donor-related variation using the batch normalization method reComBat^20^ prior to conducting a secondary UMAP analysis with identical hyperparameters. Although reComBat has not been optimized for use on mass spectrometry data, it is applied here as a proof-of-concept demonstration of how, in an unsupervised context, FTU-related variation becomes more readily discernable when at least part of the donor-specific variation can be reduced. This effect is apparent in **Figures 4B** and **4D**. While **Figures 4A** and **4B** share the donor-origin color-coding, in panel **A**, certain donor-specific subsets are clearly separated, and in panel **B**’s visualization, it is harder to discern donor-related subsets. This suggests that latent space donor-related variation has been relatively reduced from **Figure 4A** to **4B**. In **Figures 4C** and **4D**, the FTU-type color-coding suggests that, as donor-related variation is reduced, the FTU-related variation in these measurements tends to come more to the foreground in the chemical variation reported by the cohort dataset. The visualization in **Figure 4D** demonstrates that based on the 212 lipid species tracked in this dataset, the 302,142 glomeruli (GL) measurements acquired across all tissue experiments (in green) tend to group together, which suggests that this data captures genuine biological variation across different human donors. This is further supported by 462,935 collecting duct (CD) pixels (in orange), 253,961 distal tubule (DT) pixels (in red), 2,575,639 proximal tubule (PT) pixels (in purple), and 1,049,658 thick ascending limb (TAL) pixels (in brown) clustering together, regardless of donor. Similar groupings can be observed whether the analysis is done on a per-pixel basis (**Figure 4**, **figure S33**) or if a mean spectrum is calculated for each FTU instance and the unsupervised analysis is performed at the FTU-instance-level (**figure S34**). These observations suggest that the data collected as part of this atlas reports several biologically relevant underlying trends. Some of these chemical variations are donor-specific, while other trends are donor-agnostic and hold up across the cohort irrespective of the particular donor sampled, supporting the biological validity of the atlas dataset.

Similar observations on chemical variation can be discerned in the positive ionization mode IMS data (**Figure 5, figure S35,** and **figure S36**). This cohort dataset comprises 6,779,166 pixels, each reporting a vector of 211 features that report the peak intensities of a selection of 211 lipid species (**table S4**). **Figure 5** shows that the chemical variation captured by the data tends to group measurements according to the FTU-type they represent, regardless of the donor origin, and that there chemical differentiation between FTUs. **Figure 5D** depicts the FTU-instance-level grouping of 290,321 GL mean spectra (orange), 524,092 CD mean spectra (green), 271,080 DT mean spectra (red), 2,557,196 PT mean spectra (purple), and 1,020,023 TAL mean spectra (brown). In figures S33-36, we provide colorings for meta-data such as sex and BMI in addition to the FTU-type and donor origin labels shown here. However, those visualizations show less clear separation, which is possibly the result of the two-dimensional representation acquired through UMAP being too low-dimensional to accurately represent and separate all these different variation types in one cohesive 2-D space. While this is a testament to the richness of the atlas cohort datasets, it also highlights some of the limitations that come with open-ended exploration by unsupervised machine learning for this data and it hints at the need for deeper and more targeted analyses to discern how chemical variation relates to particular donor meta-data. To this end, in the next section, a more in-depth and targeted supervised machine learning approach using interpretable ML concepts is utilized in combination with this atlas to uncover potential relationships that are hard to discern through unsupervised means.

**Figure 5.**
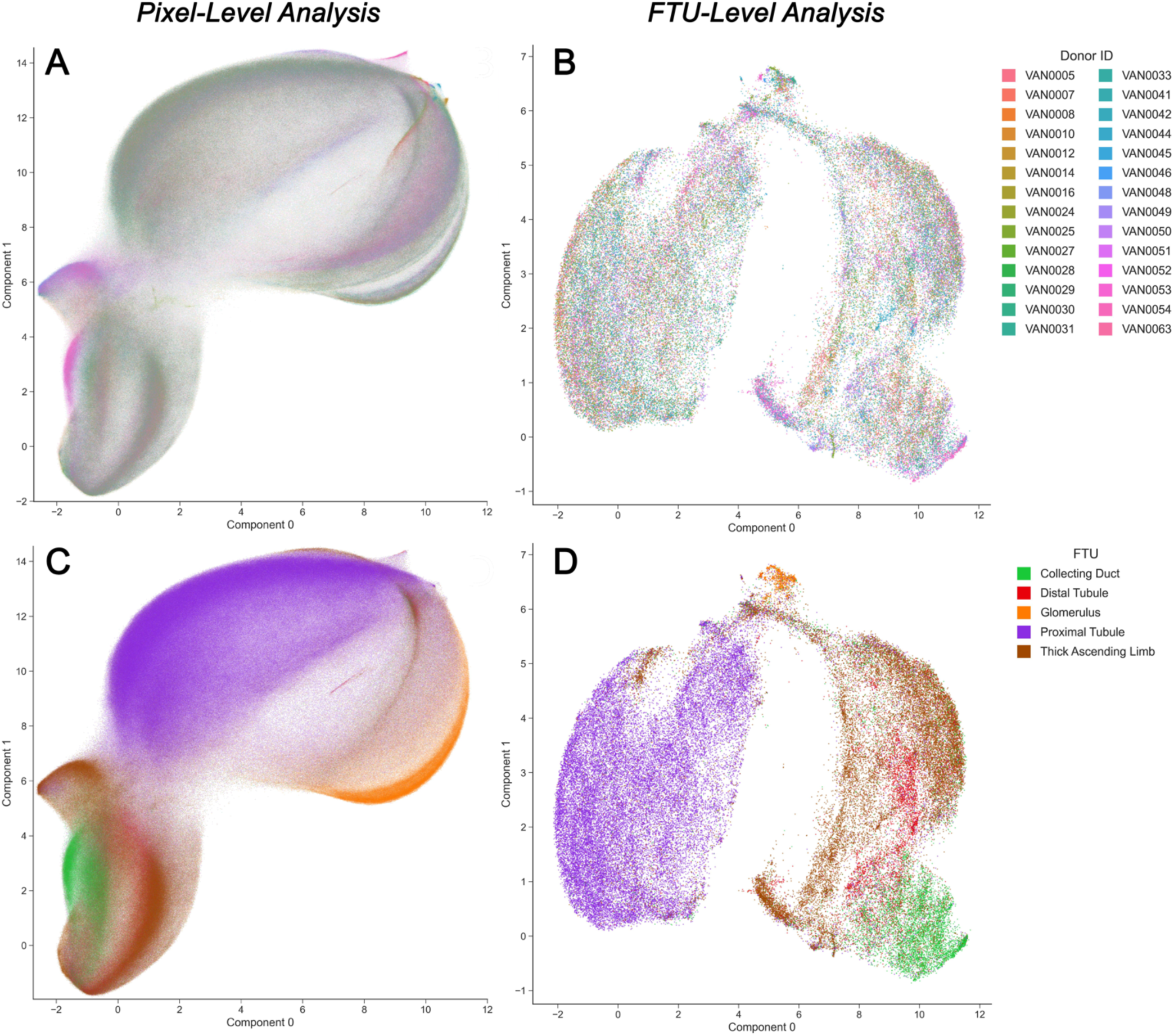
Pixel-level and FTU-instance-level chemical variation in positive ionization mode IMS measurements across 28 donors. Note, for consistency only the 28 samples that had been analyzed by negative ion mode as well were included in this visualization. (**A** and **C**) Two-dimensional visualizations of chemical variation in the positive ionization mode experiment cohort, using UMAP to cast a matrix of 6,779,166 observations (i.e., IMS pixels across 28 donor tissues) by 211 features (*i.e.,* lipid species) into a table of 6,779,166 observations by 2 latent variables, while retaining neighborhood relationships between observations as captured by a cosine distance measure. (FTU-instance-level column) Two-dimensional visualizations of chemical variation in the positive ionization mode experiment cohort, using UMAP to cast a matrix of 75,846 FTU-instances (*i.e.,* mean spectrum per FTU instance found across 28 donor tissues) by 211 features (i.e., lipid species) into a table of 74,959 observations by 2 latent variables, while retaining neighborhood relationships between observations as captured by a cosine distance measure. (**A**) Pixel-level latent space representation of chemical variation after preprocessing and reduction of donor variation by reComBat, with pixels color-coded for donor origin. (**C**) Same pixel-level latent space representation as in panel A, with pixels color-coded for FTU type (as automatically recognized from microscopy). (**B**) FTU-instance-level latent space representation of chemical variation after preprocessing and reduction of donor variation by reComBat, with pixels color-coded for donor origin. (**D**) Same FTU-instance-level latent space representation as in panel B, with pixels color-coded for FTU type (as automatically recognized from microscopy). Note, while reComBat has not been optimized for use on mass spectrometry data, it is applied here to demonstrate that if donor-specific variation can be removed, FTU-related variation becomes more readily discernable in an unsupervised context.

### Supervised Analysis and Spatially Driven Discovery of Biomarker Candidates

While informative, the unsupervised machine learning analysis of the atlas tends to show the dominant types of variation present in the datasets. This is not necessarily the chemical variation most relevant to certain questions of biological interest. For example, investigations into which lipid species are specific to certain FTUs or which species vary with sex or other donor characteristics require a more narrowly focused, in-depth analysis. To this end, we have developed an interpretable supervised machine learning workflow that enables automated discovery of which molecular species, among the hundreds tracked, are potential biomarker candidates for a specific FTU, or which species seem to be informative when differentiating sex or other donor characteristics^22^. Unlike unsupervised methods, supervised machine learning algorithms are guided by a specific prediction objective and they, therefore, tend to disregard data variation that is not relatable to the defined classification task. This ability to focus in on variation that aids in the recognition task at hand and attenuate other types of variation, makes supervised machine learning and specifically the training of classification models an effective means of filtering high-dimensional data down to a more compact set of features relevant to a certain biological question of interest. With a deepening of the molecular mapping of the kidney in mind, the following classification tasks were defined and models were built: recognition (one-versus-all classification) of five FTUs, namely glomeruli, proximal tubules, thick ascending limb, distal tubules, and collecting ducts, with the goal of obtaining lipid marker candidates for each FTU type, binary classification of female versus male donors with the aim of discerning sex-related marker candidates, and binary classification of normal-BMI (22 < BMI < 25) versus high-BMI (BMI > 35) donors directed at suggesting lipid marker candidates that differentiate these two classes. The interpretability method employed, Shapley additive explanations (SHAP)^23^, is used in each classification task to quantitatively estimate, for each of the hundreds of IMS-detected lipid species, the relevance of a particular molecular species for recognizing the target biological class. This provides, for each task, a ranked list of each molecular species’ relative predictive importance for that task, offering an experiment-wide heuristic for a lipid’s biomarker potential in a spatially driven manner. While a lipid’s relevance in a classification task trying to recognize a certain FTU is not the same as that lipid being a genuine biomarker for that FTU, such a relevance heuristic can help suggest which out of hundreds of lipid species have a potential relationship to the FTU and could merit further investigation. The automated process furthermore avoids the human bias and drift involved with performing such a task manually, and it can be performed at scale across a whole cohort of datasets, increasing the robustness of the suggested marker candidate panels. For a given classification task, our SHAP workflow quantifies the importance (as reported by a SHAP value) of each molecular species, reporting its donor cohort-wide relevance (*i.e.*, averaged across all donors), its tissue sample-wide relevance (*i.e.*, specific to a donor, but averaged across that donor’s tissue sample), and its pixel-specific relevance (*i.e.*, specific to a location within a tissue sample). Since the cohort-wide SHAP importance scores report marker candidate potential across many donors, these scores are more robust cues than donor-specific estimates. We therefore use the cohort-wide SHAP scores to extract from the atlas subsets of highly discriminative molecular species that may be biologically relevant marker candidates for each specific FTU, sex, and BMI category. It is noted that the selected comparisons (*i.e.,* FTU, sex, and BMI classifications) are just example comparisons that can be made with this atlas. The assembled and integrated data that provides the foundation of the atlas is a valuable resource that can be mined further to ask other biologically relevant questions and to include additional molecular species detected beyond the 212 (negative ion mode) and 211 (positive ion mode) lipids that have been annotated and examined here.

### Lipid Profiles of Renal Functional Tissue Units

In **figures S37-S46**, for each FTU, the top twenty cohort-wide SHAP importance scores and the corresponding suggested top twenty biomarker candidates for a specific FTU are provided, both in negative and positive ionization modes. While these plots give a cohort-wide overview of a lipid’s relevance to an FTU, an analysis of marker potential down to the donor-specific level is also provided for both modes in the form of bubble plots (**figure S47**-negative ion mode, **figure S48** – positive ion mode). For simplicity, selected ions that were found to be defining molecular features of one or multiple FTUs are represented in a summary bubble plot in **Figure 6**, combining both polarities. The bubble plot data representations provide an effective means of visualizing complex molecular species-FTU relationships in a way that makes trends for specific lipid species across the donor cohort very apparent. Molecular species that are consistent, strong marker candidates for a specific FTU across all donor samples are represented by columns of large bubbles in the chart, and the color of the bubbles indicates whether the molecular marker candidate’s signal intensity is positively (red) or negatively (blue) correlated with the FTU. Ion images for the top ten molecular markers for each FTU for both negative and positive ion modes are provided in **figures S49-S148**.

**Figure 6.**
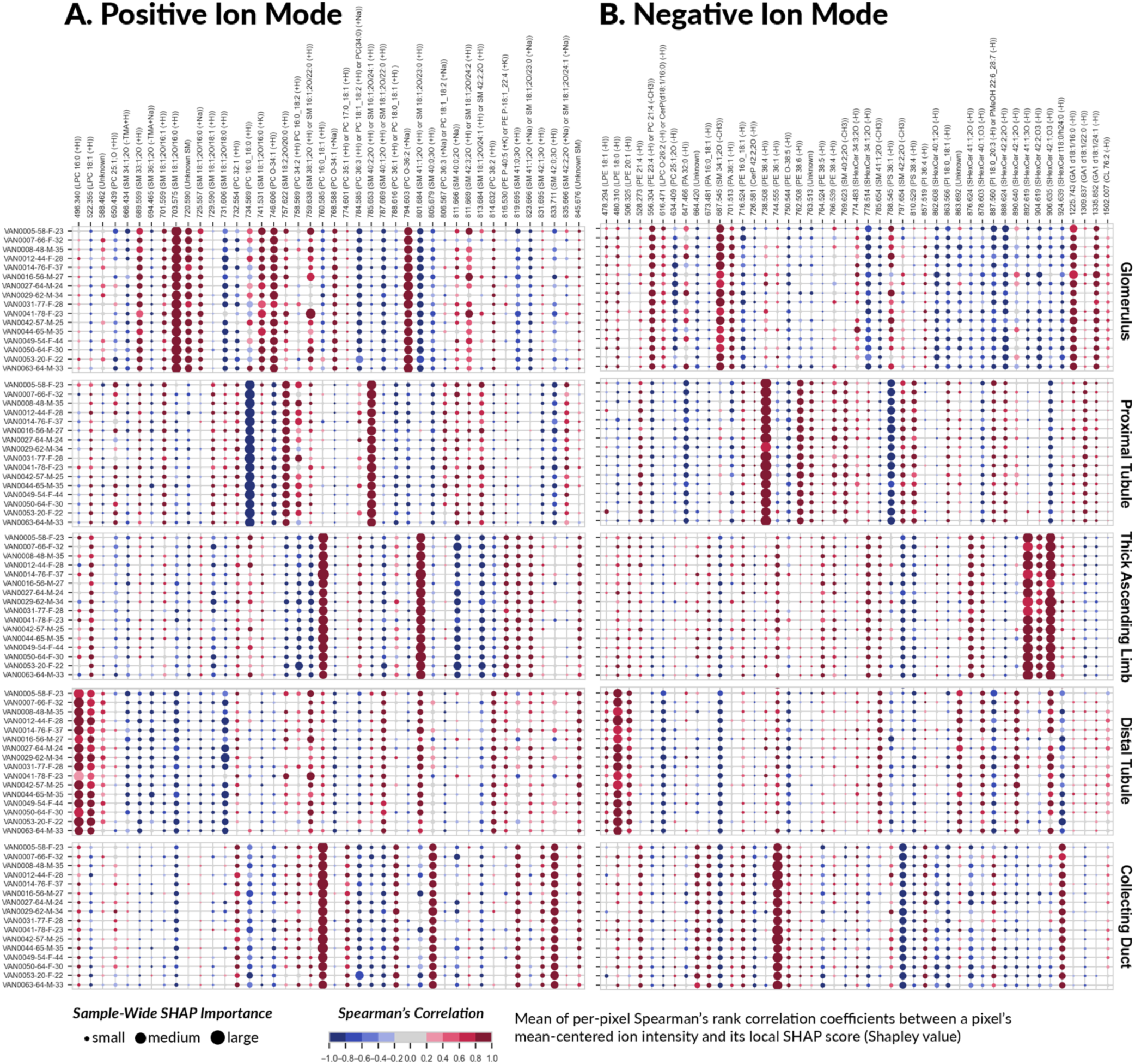
**Summary of biomarker candidates for five functional tissue units (FTUs), namely glomeruli, proximal tubules, distal tubules, collecting ducts, and the thick ascending limb, obtained by applying our SHAP-based workflow to the atlas**. The bubble plot reports both positive ion mode (left) and negative ion mode (right) findings. The columns correspond to a selection of molecular species (in increasing order of mass-to-charge ratios) that are biomarker candidates for one (or multiple) of the five FTUs under study. The rows correspond to different donors, each row labeled with its donor ID number and followed the donor’s age, sex, and BMI. Each bubble marker is informative of the direction (positive or negative correlation) and magnitude (relatively large or small) of a molecular species’ influence on the classification model designed to recognize one of the five FTUs. The marker size represents the magnitude of the molecular species’ influence, as measured by its tissue sample-wide SHAP importance score for a given donor sample. The marker color indicates the direction of the molecular species’ influence, as measured by the Spearman’s rank correlation coefficient between the molecular species’ mean-centered ion intensity values and its local pixel-specific SHAP scores. A positive Spearman’s rank correlation coefficient indicates that a high intensity of the molecular species correlates with the FTU. Conversely, a negative Spearman’s rank correlation coefficient indicates that a low intensity of the molecular species correlates with the FTU.

Tissue atlases depend on comparisons to previous studies to anchor outputs and provide a reliable foundation for new observations. For example, phosphatidic acid PA(36:1) has previously been found to mark glomeruli using IMS (at 30 um pixel size)^24^. Our higher spatial resolution multimodal imaging workflows and SHAP analysis are consistent with this observation, finding the presence of PA(36:1) to be a marker candidate for glomeruli (**Figure 6B, figures S37** and **S47**). We also found multiple gangliosides to be specific to glomeruli, including GA1(d18:1/16:0) (*m/z* 1225.743) and GA1(d18:1/24:1) (*m/z* 1335.852) (**Figure 6B, figures S37, S47, S50,** and **S54**). While GA1 species have not previously been reported to be specifically localized to the glomerulus, products of the GA1 ganglioside biosynthesis pathway, GM1b and GD1a, have been detected as part of the lipid profiles of podocytes^25^, a unique and important cell type contributing to the filtration barrier in the glomerulus. We and others have previously reported the specific localization of the sphingomyelin molecule SM(d34:1;2O) to glomeruli^26^. Here, SM(d34:1;2O) (identified as SM(d18:1;2O/16:0) in positive ion mode) represents the most robust glomerular signature, having the highest SHAP scores in both negative (*m/z* 687.545) (**Figure 6B, figures S37, S47,** and **S49**) and positive ion modes (*m/z* 703.575, H+ adduct) (**Figure 6A, figures S38, S48,** and **S59**). Other positive ion mode adducts of SM(d34:1) (Na+: *m/z* 741.531 and K+: *m/z* 725.557) were detected with correlation to glomeruli (**Figure 6A, figures S38, S48** and **S63**). SHAP analysis also revealed a negative correlation of SM(d34:1;2O) with the thick ascending limb (*m/z* 687.545), distal tubules (*m/z* 666.434, *m/z* 687.545, and *m/z* 703.575) and collecting ducts (*m/z* 687.545 and *m/z* 703.575) (**Figure 6A** and **6B, figures S41** and **S43-S46**). Sphingomyelin species are enriched in glomeruli and proximal tubules (**Figure 6A and B**, **figures S47** and **S48**), but the distinct spatial distributions of SM(d34:1;2O) to glomeruli (**Figure 6A** and **6B** and supplemental **figure S49, S59,** and **S63**) indicates a functionally specific localization.

The proximal tubule, which is the primary site of glucose reabsorption in the nephron, is known to have an unusually rigid apical membrane^27^. Decreased membrane fluidity is commonly associated with a high ratio of sphingomyelin^28^. Our analysis of the lipid atlas showed that the proximal tubule was the most sphingomyelin-enriched FTU, with nine sphingomyelin species detected and identified as having a high positive association with proximal tubules based on cohort-wide SHAP importance scores (negative mode *m/z* 769.623 and *m/z* 797.654, positive mode *m/z* 701.559, *m/z* 729.590, *m/z* 757.622, *m/z* 785.653, *m/z* 811.666, *m/z* 813.684, *m/z* 835.666) (**Figure 6A** and **6B, figures S39, S40, S47, S48, S74, S75, S80, S81, S83, S84, S86, and S88**). Of those, eight were exclusively discriminative for proximal tubules (negative mode *m/z* 769.623 and *m/z* 797.654, positive mode *m/z* 701.559, *m/z* 729.590, *m/z* 785.653, *m/z* 811.666, *m/z* 813.684, *m/z* 835.666). The acyl chain identity for five of the eight sphingomyelin species was revealed to contain long-chain fatty acids (>C18 up to C26 in length) (*m/z* 729.590, *m/z* 757.622, *m/z* 785.653, *m/z* 813.684, *m/z* 835.666), which contribute to decreased membrane fluidity. Membrane fluidity is a critical regulator of glucose uptake in the proximal tubule, as increasing fluidity inhibits glucose reabsorption by sodium-glucose cotransporter-2 (SGLT2)^29^. In addition to membrane fluidity as a regulator of SGLT2, its function is also regulated by protein kinase C (PKC)^30^. While PKC binds membranes enriched in sphingomyelin, it also requires phosphatidylserine (PS) for activation in such environments^31^. In addition to a sphingomyelin rich environment in proximal tubules, we also identified the phosphatidylserine PS(38:4) (*m/z* 810.529) as being a positive marker for proximal tubules (**Figure 6B, figures S39, S47,** and **S73**).

In proximal tubules, a sodium gradient must be established for uptake of sodium, and subsequently glucose, from the lumen across the apical membrane via SGLT2. This gradient is established by the basolateral Na+/K+ ATPase pump^32^. Na+/K+ ATPase activity is known to be modified by neutral phospholipids, particularly polyunsaturated phosphatidylethanolamines (PE). Of the eleven polyunsaturated phosphatidylethanolamines detected in the atlas (*m/z* 528.273, *m/z* 556.304, *m/z* 714.508, *m/z* 722.513, *m/z* 738.508, *m/z* 750.544, *m/z* 762.508, *m/z* 764.524, *m/z* 766.539, *m/z* 778.576, and *m/z* 816.530) PE (21:4) (*m/z* 528.273), PE(36:4) (*m/z* 738.508), PE(38:4) (*m/z* 766.539), and PE(38:6) (*m/z* 762.508) had cohort-wide SHAP importance scores that were highly associated with proximal tubules (**Figure 6B, figures S39, S47**, **S69, S71, and S76**). Of those, PE(38:4) has been shown to directly stimulate Na+/K+ ATPase pump activity^33^, and this lipid was found to be a significant biomarker candidate of proximal tubules. The enrichment of polyunsaturated PEs and the localization of PE(38:4) (*m/z* 766.539) to the proximal tubules (**Figure 2B.2**, **figures S47** and **S76** may be important for the tuning of Na+/K+ ATPase activity to modulate SGLT2 function.

The kidney is rich in sulfatides, a class of glycosphingolipids that contain a ceramide bound to a sulfated carbohydrate. Within the kidney, sulfatidyl hexose ceramides (SHexCer) are strongly associated with the thick ascending limb (**Figure 6B, figures S41** and **S47**). In the thick ascending limb, six of the seven sulfatidyl hexose ceramide species (*m/z* 778.514, *m/z* 876.624, *m/z* 878.603, *m/z* 892.619, *m/z* 904.619, *m/z* 906.635) detected had positive discriminatory values for this FTU (**figure S41**). Furthermore, among the atlas-measured species, the sulfatidyl hexose ceramides detected at *m/z* 876.624, *m/z* 892.619, *m/z* 904.619, and *m/z* 906.635 are the most robustly associated thick ascending limb biomarker candidates detected (**Figure 6B, figures S41, S47, S89-92**). In positive ion mode, of the five species with the highest cohort-wide SHAP importance scores (*m/z* 758.569, *m/z* 760.585, *m/z* 801.684, *m/z* 811.666, and *m/z* 813.684) (**figure S42**) none are unique to thick ascending limb and they were not considered as discriminatory biomarker candidates of this FTU specifically (**Figures 6A** and **S48**). These observations suggest that the sulfatidyl hexose ceramide signature in the thick ascending limb represents a defining feature for this FTU.

The distal tubule is the shortest segment of the nephron. However, it is critical for absorption of divalent cations, including magnesium and calcium, as well as for maintaining sodium and potassium homeostasis^34^. Distinct from the preceding nephron segments, the movement of ions in the distal tubule occurs primarily via transcellular transport. The distal tubule is unique in that it is highly plastic and can fine-tune ion uptake and secretion in response to luminal ion concentration by modulating ion channel expression and activity. Interestingly, the most robust biomarker candidates for distal tubules include lysophosphatidylcholine (*m/z* 496.340 and *m/z* 522.355) in **Figure 6A** (and **supplemental figures S44, S48, S119, S120**) and lysophosphatidylethanolamine (*m/z* 480.310) as highlighted in **Figure 6B** (and **supplemental figures S43**, **S47, and S109**). Lysophospholipids are present at low cellular concentrations. However, both LPC(16:0) (*m/z* 496.340, **supplemental figure S119**) and LPE(18:0) (*m/z* 480.310, **supplemental figure S109**) are discriminative for distal tubules and have the highest cohort-wide SHAP importance scores in positive and negative ion modes, respectively. In addition, LPE(18:1) (*m/z* 478.294) and LPE(20:1) (*m/z* 506.325) are DT biomarker candidates (**figures S47 and S115**). Various sphingomyelins (*m/z* 731.60, *m/z* 801.68, *m/z* 759.63) and sulfatidyl hexose ceramides (*m/z* 906.63, *m/z* 924.63, *m/z* 890.63) also have high cohort-wide SHAP importance scores for this FTU, but none of these are exclusive to distal tubules (**Figures 6A and B**, and **figures S47 and S48**). Prevalence of lysolipids seems to be a defining feature of distal tubule functional tissue units.

The renal filtrate from tubules drains into the collecting ducts, the final stage where nutrients are reabsorbed to adjust final concentration of electrolyte and acid-base components of the resulting urine. The lipid profile of the collecting ducts was found to be enriched in phosphatidylcholines (**Figure 6A, figures S46** and **S48**) and phosphatidylethanolamines (**Figure 6B**, **figures S45** and **S47**), both of which serve as discriminatory markers for collecting ducts as per our SHAP analysis. **Figures S45** and **S46** show that four of the top *m/z* markers for collecting ducts were either phosphatidylcholines (*m/z* 734.569, *m/z* 760.585, *m/z* 788.616) (**figures S139, S142, and S143**) or phosphatidylethanolamines (*m/z* 744.555) (**figure S129**). PE(36:1) (*m/z* 744.555) is the most significant biomarker candidate of collecting ducts detected in negative ion mode (**figure S45**). While not exclusive to collecting ducts, its relative cohort-wide SHAP importance is significantly higher than in distal tubules and the thick ascending limb, where it was determined to be a less prominent positive marker (**Figure 6B, figures S41, S43,** and **S45**). In positive ion mode, PC(16:0_18:1) (*m/z* 760.585) has the highest cohort-wide SHAP importance score for collecting ducts (**Figure 6A, figures S46** and **S48**) but is also positively associated with the thick ascending limb (**Figure 6A, figure S42**). PC(32:1) (*m/z* 732.554), PC(35:1) (*m/z* 774.601), and PE(38:1) (*m/z* 772.586) are nearly exclusive markers of collecting ducts (**Figure 6A and B, figures S37-46, S147-148)**. Three additional phosphatidylcholine (*m/z* 758.569) (**figure S46**) or phosphatidylethanolamine lipids (*m/z* 528.273 and *m/z* 716.524) (**figure S45**) are also positively associated biomarker candidates for collecting ducts (**figures S45** and **S46**), albeit not as robustly. Although none of these three are exclusively discriminatory, the overall number of positively associated phosphatidylcholine and phosphatidylethanolamine biomarker candidates for collecting ducts as determined by our SHAP analysis is twelve total the highest number for any FTU (**figures S45** and **S46**). Collecting ducts contain mitochondria rich cells, which would be consistent with a lipid profile enriched for PC and PE as they are the most abundant lipids in the mitochondrial membrane^35^.

The sulfatidyl hexose ceramide SHexCer(t18:0/h24:0) (*m/z* 924.639) also has strong positive association with the collecting duct, the second highest positively associated biomarker candidate detected in negative ion mode (**Figure 6B**, **figures S45, S47, and S131)**. Interestingly, it is anti-correlated with the preceding distal tubule segment (**Figure 6B, figures S43** and **S47**), suggesting a collecting duct specific function. SHexCer(t18:0/h24:0) (*m/z* 924.639) incorporates a phytosphingosine as the sphingoid base as well as a hydroxy fatty acid. It is of note that it is one of only two phytosphingosine ceramides detected in the dataset and the only one positively correlated with an FTU. SHexCer(t18:0/h24:0) (*m/z* 924.639) has been reported to specifically localize to the collecting duct^36^. The presence of the collecting duct-specific 4-hydroxylated SHexCer(t18:0/h24:0) (*m/z* 924.639) in conjunction with long chain sphingomyelins, SM(40:0;2O) (*m/z* 805.679) and SM(42:0;3O) (*m/z* 833.711), which are two of the top collecting duct biomarker candidates (**Figure 6, figures S45-48, and S140-141**), points to a functional role in membrane structure for this group of lipids. It is speculated that tuning these lipid components of the collecting duct membrane could allow for strict control of NH_3_ and H^+^ transport via a transcellular route exclusively.

### Sex and BMI Comparisons

Using the demographic information provided in the donor metadata of the atlas, we can go beyond the discovery of FTU-level biomarker candidates and explore lipid associations with sex and other important characteristics such as BMI. Binary classification of female versus male donors and binary classification of normal-BMI (22 < BMI < 25) versus high-BMI (BMI > 35) donors, in conjunction with our SHAP-based interpretable machine learning workflow, enables discovery of lipid biomarker candidates that are associated with these donor categories. The cohort-wide SHAP importance scores, highlighting the top biomarker candidates for the classification of female from male donor tissue overall (*i.e.,* not specific to an FTU) for both negative and positive ion modes, are provided in **figures S149** and **S150** and summarized in bubble plots (**Figure 7A** and **7B**). FTU-specific analyses were also performed, differentiating female from male kidney functional tissue units for both polarities (**figures S151-S160**). While our SHAP approach is inherently multivariate, we can also analyze univariate ion intensity distribution comparisons. These are provided as split violin plots for a selection of biomarker candidates for both polarities (positive ion mode – **Figure 7C** and **figure S162**, negative ion mode – **Figure 7D** and **figure S161**). Ion images for the top ten overall sex biomarker candidates for both negative and positive ion modes are provided in **figures S163-S182**.

**Figure 7.**
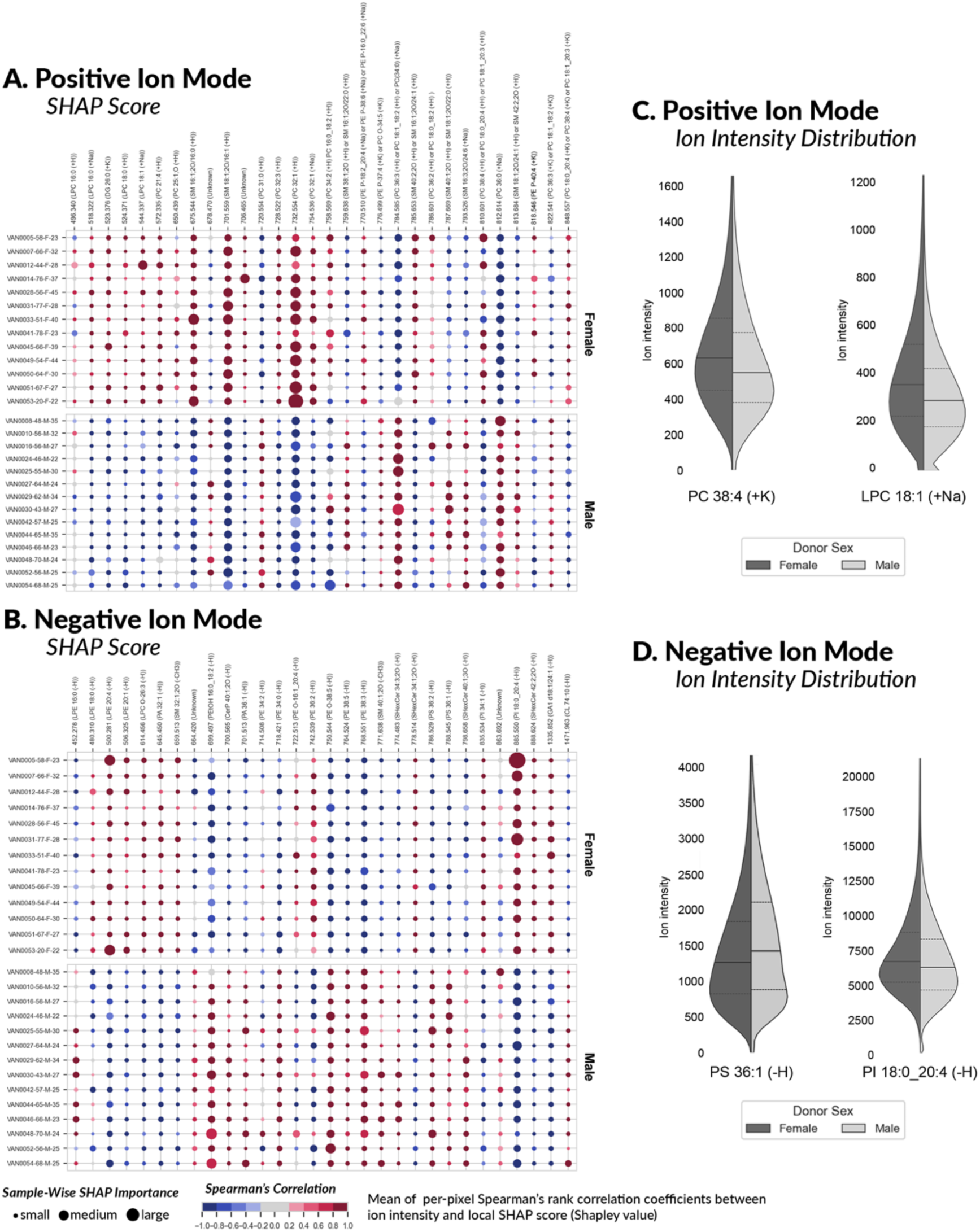
**Summary of biomarker candidates for sex, obtained by applying our SHAP-based workflow to the atlas**. The bubble plots report positive ion mode (A) and negative ion mode (B) findings. The columns correspond to a selection of molecular species (in increasing order of mass-to-charge ratios) that are biomarker candidates for the female sex. The rows correspond to the different donors. Each bubble marker is informative of the direction and magnitude of a molecular species’ influence on the classification model designed to recognize female donor tissue from male donor tissue. A molecular species with a large marker acts as a differentiator between sexes. The marker color indicates the direction of the molecular species’ influence, as measured by the Spearman’s rank correlation coefficient between the molecular species’ mean-centered ion intensity values and its local pixel-specific SHAP scores. Molecular species that are positively correlated with female sex are negatively correlated with male sex, and vice-versa. A positive Spearman’s rank correlation coefficient indicates that a high intensity of the molecular species correlates with either the female sex (A/B top) or the male sex (A/B bottom). Conversely, a negative Spearman’s rank correlation coefficient indicates that a low intensity of the molecular species correlates with the female sex (A/B top) or the male sex (A/B bottom). The split violin plots on the right report the ion intensity distributions of select sex biomarker candidates in positive mode (C) and negative mode (D), approximated using kernel density estimation. The violin plots are cropped at the 99th percentile of the distribution of one of the two sexes (whichever is larger) to facilitate visual comparison. The full line of each violin plot indicates the median of each class’ distribution, whereas the dashed lines indicate its interquartile range.

Stratifying the dataset for females versus males reveals enrichment in phosphatidylcholine and lysophosphatidylcholine in females. There are nine molecular signals, representing phosphatidylcholine species, presenting as positively correlating marker candidates for females (*m/z* 572.335, *m/z* 650.439, *m/z* 728.522, *m/z* 732.554, *m/z* 754.536, *m/z* 758.569, *m/z* 786.601, *m/z* 810.601, *m/z* 848.557), compared to four species positively correlating with males (*m/z* 720.554, *m/z* 784.585, *m/z* 812.614, *m/z* 822.541) (**Figure 7A and figures S150, S173, S175, S176, S178-S180**). For lyso-phosphatidylcholines, none are positively correlated with males, while five (*m/z* 496.340 slightly, *m/z* 518.322, *m/z* 524.371, *m/z* 544.33, *m/z* 614.456) show positively correlating discriminative association to females (**Figures 7A and B**, and **figure S150**), consistent with the phosphatidylcholine pattern. Our analysis suggests that phospholipids containing arachidonic acid LPE(20:4) (*m/z* 500.281), PE(O-16:1_20:4) (*m/z* 722.513), PC(18:0_20:4) (*m/z* 810.601), and PI(18:0_20:4) (*m/z* 885.55), are more (positively correlating) associated with females (**Figures 7A-D, figure S149, S150, S163, and S167**). Out of the molecular species captured by this atlas, PI(18:0_20:4) (*m/z* 885.55) is suggested to be the most robust sex biomarker candidate, positively correlating in females (**Figures 7B** and **7D**).

Figures 7B and **7D** show that phosphatidylserines play a consistent role in differentiating females and males, with our analysis reporting that several PS lipids have a negative Spearman’s rank correlation coefficient, indicating that low intensity of these molecular species correlates with the female sex and high intensity correlates with males. This is further apparent when evaluated by individual FTU (**figures S151, S153, S155, S157,** and **S159**). In a recent study, it was shown that phosphatidylserines comprise 60% of the urine lipidome in males and 45% in females, with PS(36:1) (*m/z* 788.545) being the most abundant urine lipid. In our analysis, PS(36:1) (*m/z* 788.545) was found to be a biomarker candidate for males overall (Figures 7B and **7D, figure S149**) and an even higher-ranked differentiating marker for the thick ascending limb (**figure S155**). PS(36:2) was also found to be higher in male urine, and in our atlas, this phosphatidylserine was also found to be an overall molecular marker for male kidney tissue (Figures 7B**, figures S149 and S171**) with PS(36:2) (*m/z* 786.529) having a high ranking cohort-wide SHAP importance score in proximal tubules (**figure S153**) relative to other FTUs and the overall tissue ranking.

To identify molecular signatures for obesity, 14 donors were selected from the atlas to compare those with a normal BMI range (BMI 20-25, 7 donors) to those with BMI values classified as obese (BMI >35, 7 donors). The cohort-wide SHAP importance scores highlighting the top biomarker candidates for the classification of obese from normal donor tissue overall (*i.e.,* not specific to an FTU) for both negative and positive ion modes are provided in **figures S183** and **S184** and summarized in bubble plots (Figure 8A and **8B**). FTU-specific analysis was also performed, differentiating obese from normal kidney functional tissue units for both polarities (**figures S185-S194**). Intensity distribution comparisons are provided as split violin plots for a selection of obesity biomarker candidates for both positive ion mode (Figure 8C and **figure S195)** and negative ion mode (Figure 8D and **figure S196**). Ion images for the top ten overall obesity biomarker candidates for both negative and positive ion modes are provided in **figures S197-S216**.

**Figure 8.**
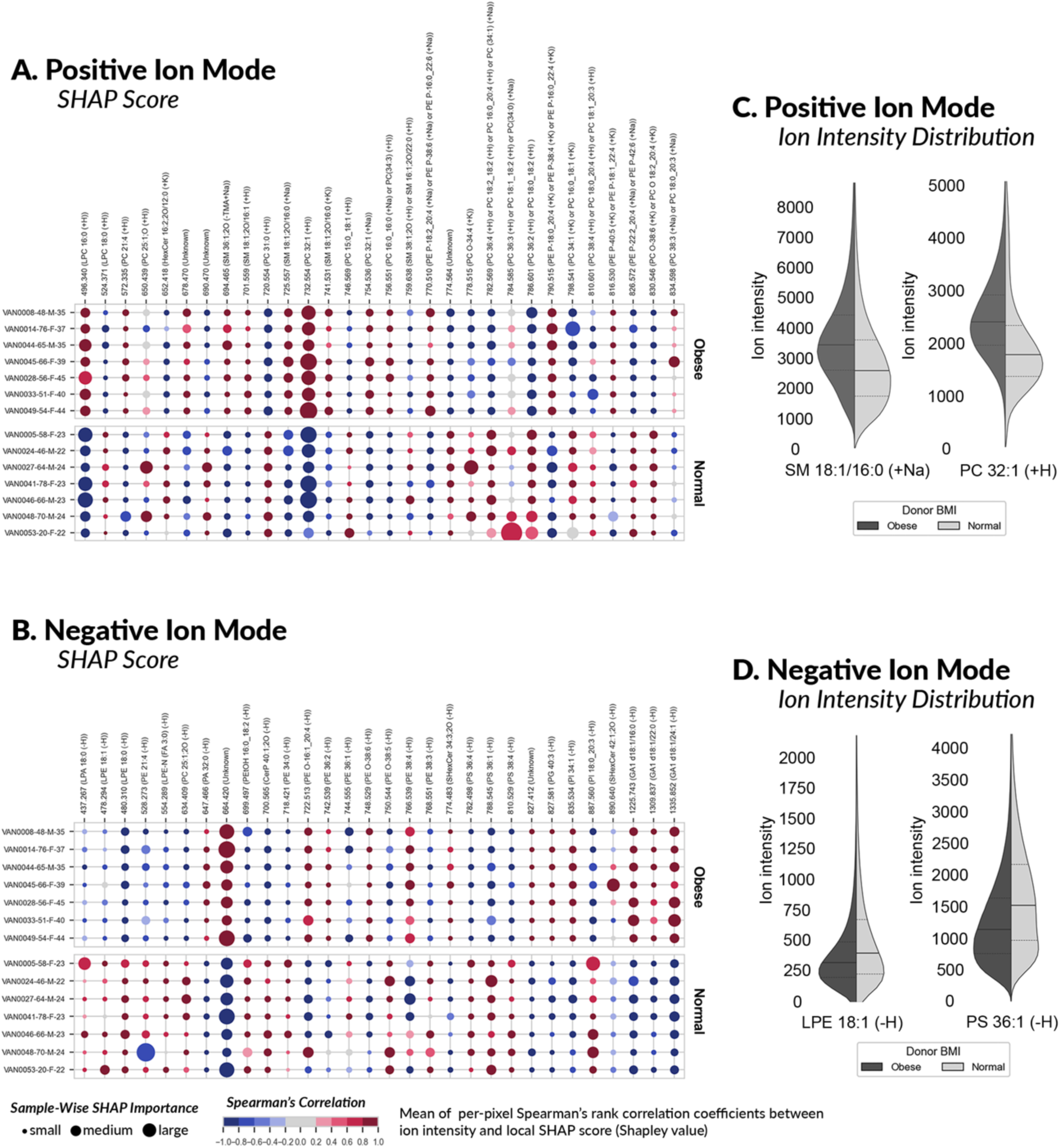
Summary of biomarker candidates for BMI, obtained by applying our SHAP-based workflow to the atlas. The bubble plots report positive ion mode (A) and negative ion mode (B) findings. The columns correspond to a selection of molecular species (in increasing order of mass-to-charge ratios) that are biomarker candidates for the characteristic of obesity. The rows correspond to the different donors. Each bubble marker is informative of the direction and magnitude of a molecular species’ influence on the classification model designed to recognize high-BMI donor tissue from normal-BMI donor tissue. A molecular species with a large marker acts as a differentiator between BMI classes. The marker color indicates the direction of the molecular species’ influence, as measured by the Spearman’s rank correlation coefficient between the molecular species’ mean-centered ion intensity values and its local pixel-specific SHAP scores. Molecular species that are positively correlated with high-BMI are negatively correlated with normal-BMI, and vice-versa. A positive Spearman’s rank correlation coefficient indicates that a high intensity of the molecular species correlates with either high-BMI (A/B top) or normal-BMI (A/B bottom). Conversely, a negative Spearman’s rank correlation coefficient indicates that a low intensity of the molecular species correlates with either high-BMI (A/B bottom) or normal-BMI (A/B bottom). The split violin plots on the right report the ion intensity distributions of select obesity biomarker candidates in positive mode (C) and negative mode (D), approximated using kernel density estimation. The violin plots are cropped at the 99th percentile of the distribution of one of the BMI categories (whichever is larger) to facilitate visual comparison. The full line of each violin plot indicates the median of each class’ distribution, whereas the dashed lines indicate its interquartile range.

Lysophosphatidylethanolamine (LPE) species, including LPE(18:1) (*m/z* 478.284), LPE(18:0) (*m/z* 480.310), and N-acyl-lysophosphatidylethanolamine LPE-N(FA3:0) (*m/z* 554.289), were found to be differentiators and positively correlating to normal donors (Figures 8B and **8D, figures S183** and **S205**). Although many phosphatidylcholines were found to be markers for both normal and obese BMI kidney tissue (Figures 8A and **8C, figures S183** and **S184**), PC(32:1) (*m/z* 732.554) was found to be the greatest discriminator of normal and obese tissues. PC(32:1) (*m/z* 732.554) is a positively correlating discriminative marker candidate for obesity and appears negatively correlated to normal tissue, with the greatest cohort-wide SHAP importance score (Figures 8A**, figure S184** and **S207**). Phosphatidylcholine PC(36:2) (*m/z* 786.601) (most likely PC(18:0_18:2) based on LC-MS/MS data) was found to be one of the most consistent (positively correlating) biomarker candidates for normal BMI, and PC(O-34:4) (*m/z* 778.515) and PC(34:1) (*m/z* 798.541)suggest less prominent but similar behavior (Figure 8A and **figures S184, S209, S211,** and **S214**). In negative ion mode, the ion detected at *m/z* 664.420 was the most discriminant of BMI, reporting the highest absolute cohort-wide SHAP importance score and correlating positively with obesity and negatively with normal BMI (Figure 8B**, figures S183** and **S197**). Although we were unable to confirm the identity of this molecule by MS/MS, based on mass accuracy, this is likely a PAz-PC lipid species (*i.e.,* 1-palmitoyl-2-azelaoyl PC), an oxidized phosphatidylcholine previously observed in kidney disease^37,38^. Obesity represents a chronically inflamed state, and under these conditions, coagulation factors are aberrantly expressed or regulated, promoting increased prothrombotic risk^39,40^. The oxidized phospholipid, PAz-PC, could be an important molecule in the feedback loop that exists between obesity and risk factors for cardiovascular disease as PAz-PC inhibits the tissue factor pathway inhibitor (TFPI)^41^. TFPI inhibits the tissue factor-dependent, or extrinsic coagulation, pathway by directly and indirectly perturbing the activation of the serine protease Factor Xa that interacts with fibrin, a marker of obesity^42^ and linked to cardiovascular disease^43^.

Other marker candidates for obesity included phosphatidylserines and sphingomyelins. Phosphatidylserines PS(36:4) (*m/z* 782.498), PS(36:1) (*m/z* 788.545), and PS(38:4) (*m/z* 810.529) were all (positively correlating) molecular indicators of normal BMI donor tissues (Figure 8B and **figures S183** and **S201**). PS(36:1) (*m/z* 788.545) was also a positively correlating biomarker candidate for glomeruli and a negatively correlating marker for proximal tubules (not considering obesity), although it showed a relatively high degree of variability as an FTU indicator (Figure 6B). When considering the classification of normal versus obese BMI categories, PS(36:1) (*m/z* 788.545) was found to be a more consistent marker candidate positively associated with normal BMI donor tissues (Figures 8B and D and **figures S183** and **S201**). This is an example of how our SHAP analysis of the kidney lipid atlas can differentiate consistent biomarker candidates from those that are rather variable and linked to specific donor comorbidities. The sphingomyelins SM(18:1;2O/16:0) (*m/z* 725.557 and *m/z* 741.531) and SM(18:1;2O/16:1) (*m/z* 701.559) were found to be discriminant for obese donor tissues (Figures 8A and **8C** and **figures S184** and **S213**), correlating positively with high BMI. Given that SM(18:1;2O/16:0) is a robust glomerular biomarker candidate as well (Figure 6A), the specific sphingomyelin profile of glomeruli in obese versus normal donors was analyzed further. SM(18:1;2O/16:0) (*m/z* 725.557) demonstrated the highest cohort-wide SHAP importance score of all measured sphingomyelin species and showed positive correlation to obesity (**Figures S186**). This observation appears consistent with the trend of the percentage of glomerular sclerosis found in the histopathological reports of the PAS-stained tissues. The average glomerular sclerosis for obese donors was 15.96% compared to 7.8% in normal donor tissues (**table S2**). The positive correlation of high SM(18:1;2O/16:0) (*m/z* 725.557) intensity to glomeruli in tissues from donors with obesity (**Figure S186**) and increased glomerular pathology indicates that perturbation of SM(18:1;2O/16:0) (*m/z* 725.557) may be a molecular signature of glomerular disease.

## DISCUSSION

We have constructed a multimodal, high-dimensional atlas of the human kidney, including lipid, multicellular functional tissue unit, and histological imaging data, as well as pathophysiological information for each human subject. It is constructed from millions of mass spectral measurements with spatial annotations delineating more than one hundred thousand functional tissue unit instances across 29 donor whole slide kidney tissue sections. As part of the NIH Human Biomolecular Atlas Program^4^, the data are freely accessible to the greater scientific community (https://portal.hubmapconsortium.org, **table S3**). As an initial demonstration of the atlas’ potential, we highlight lipid differences between specific functional tissue units of the nephron, between subjects with BMIs classified as normal (BMI 20-25) and obese (BMI>35), and differences between men and women. Additional insight into the molecular variance associated with demographics may help explain the functional ramifications resulting in differential severity in diseases such as chronic kidney disease, cancer, and diabetes^44–50^. Lipids, particularly in the context of functional tissue units and cellular neighborhoods, are chronically understudied compared to their transcript and protein counterparts despite their clear involvement in health and disease^51,52^. Many large studies assume that genetic expression is sufficient for understanding disease states. While likely true in some cases, there is a known discordance in gene expression and metabolomic profiles, indicating that further research in this area is required to understand complex physiology. Defining spatially specific molecular profiles is necessary for realizing the promise of personalized medicine and improved therapeutics. As a step towards this goal, our kidney lipid atlas serves to catalog lipid profiles on a multiscale level. Ultimately, multidimensional and multiscale atlas efforts, such as the one discussed here, can provide key findings for a defined, targeted validation, and serve as a resource for others to use to generate new hypotheses and corroborate orthogonal studies. Our multimodal lipid atlas and comprehensive analyses, including both unsupervised and supervised interpretable machine learning approaches, can establish the basal lipid signature of normal renal tissue against which diseased tissue can be compared, providing further insight into pathobiological mechanisms.

### Extraction of biological insight into FTU lipid function

The sphingomyelin SM(18:1;2O/16:0) is highly enriched in glomeruli and requires the podocyte-specific enzyme ceramide synthase 6 (CERS6) for synthesis. Genetic ablation of CERS6 leads to the loss of F-actin fibers and downregulation of synaptopodin and CD2AP (CD2-associated protein)^53^. CD2AP facilitates nephrin binding to the actin cytoskeleton^54^. This complex of CD2AP, synaptopodin, and nephrin tethered to the cytoskeleton localizes to lipid rafts to form the supercomplex that anchors the podocyte slit diaphragm. Lipid raft formation is governed by the interaction of cholesterol with locally available phospholipids. When cholesterol was mixed with sphingomyelin with varying acyl chain lengths, it was shown that sphingomyelins containing 16:0 acyl chains have the highest affinity for cholesterol in ternary PC:SM:cholesterol bilayers, the highest lateral segregation tendency, and the highest thermostability^55^, suggesting that SM(18:1;2O/16:0) is critical for the formation of lipid rafts. Given that SM(18:1;2O/16:0) is specifically correlated with the glomeruli and CERS6 downregulation destabilizes the slit diaphragm, we propose that SM(18:1;2O/16:0) is mechanistically linked to the slit diaphragm by providing a local lipid environment that stabilizes the architecture of, or regulates the formation of, the specific lipid raft microdomain associated with the slit diaphragm.

Long-chain fatty acid-containing phospholipids combined with cholesterol lead to plasma membranes with high rigidity and higher viscosity^28^. While SM(18:1;2O/16:0) was determined to be a strong biomarker candidate for glomeruli (among the species measured in this atlas), a more diverse set of six sphingomyelins were found to exhibit positive marker potential for proximal tubules. This enrichment in sphingomyelins, taken together with the tendency for these proximal tubule-specific sphingomyelins to have very long chain fatty acid tails, driving leaflet interdigitation and a higher propensity for fluid-to-gel phase transition in lipid membranes^56^, is consistent with the rigid apical membrane and lower membrane fluidity of proximal tubules. Increasing the membrane fluidity by applying exogenous insults or under pathological conditions such as diabetes results in the reduction of glucose transport mediated by the sodium-glucose cotransporter SGLT2^27,29,57^, thereby abrogating a key function of the proximal tubule segment of the nephron where the majority of glucose is reabsorbed.

While PKC and Na^+^/K^+^ ATPase are widely expressed, they are important regulators of SGLT2 in the proximal tubule^30,32^. The identification of specific lipids in proximal tubules that can serve as regulatory molecules for these proteins suggests that they could be critical for modifying PKC and Na^+^/K^+^ ATPase activity, which could then fine-tune SGLT2 activity and glucose reabsorption. Of the two phosphatidylserine species that emerged as biomarker candidates in the atlas, the proximal tubule-specific PS(38:4) could be critical for activating PKC in a sphingomyelin-rich environment. While PKC can bind to sphingomyelin-rich membranes with high affinity, this lipid microenvironment inhibits PKC activity. This inhibition is relieved by the binding of phosphatidylserine, resulting in full kinase activity^31^. Similar to PKC activation, Na^+^/K^+^ ATPase is stimulated by specific lipids, including polyunsaturated phosphatidylethanolamines. Proximal tubules had five polyunsaturated PE lipids with (positively correlating) discriminative association, more so than the thick ascending limb and distal tubules, which express higher levels of the sodium pump^58^. This enrichment in specific phosphatidylethanolamines suggests that these may be regulatory molecules that can be leveraged to adjust the sodium gradient by modifying Na^+^/K^+^ ATPase activity in response to glucose absorption cues from the lumen.

Sulfatides in the thick ascending limb present an anionic charge on the plasma membrane, where they are available to modulate membrane ion fluxes by binding cations. They have been proposed to act as counterions to interstitial ammonium (NH ^+^), leading to its accumulation in the thick ascending limb^59^. Additionally, sulfatides have been proposed to be regulators of Na^+^/K^+^ ATPase by binding K^+^ directly or modulating its localization^59^. The ceramide moiety of SHexCer is a bioactive second messenger known to activate phosphatases as well as kinases and modulate several thick ascending limb-specific ion channels. The ceramide target protein phosphatase 1 (PP1)^60^, dephosphorylates sodium–hydrogen antiporter 3 (NHE3), alleviating inhibition of this Na^+^/H^+^ antiporter^61^. If PP1 activity is inhibited, the dephosphorylation of Na^+^/K^+^ ATPase is perturbed, and pump activity is negated^62^. Finally, PP1 can suppress epithelial sodium channel (ENaC) activity and regulate sodium reabsorption. The enrichment of SHexCer in the thick ascending limb and putative roles of both sulfatide and ceramide in regulating ion balance suggest that sulfatidyl hexose ceramides are mechanistically important for thick ascending limb function.

While the nephron as an entire unit is taxed with filtrate reabsorption, the distal tubule functional tissue unit is distinct from preceding nephron segments in how it executes reabsorption. The distal tubule relies more on transcellular transport for a tightly controlled approach to regulating water and ion balance^63^. The ability to adjust the activity of ion channels is critical to the function of distal tubules. Second messengers allow cells to be highly plastic and adapt to environmental cues, such as the change in electrolyte concentration. Reactive oxygen species (ROS) are a class of second messengers that function in signaling cascades that target ion channels and transporters^64^. Lysophosphatidylcholines (LPCs) were recognized early on to be an inducer of ROS^65^. Subsequently, it was discovered that LPC(16:0) is the most potent ROS inducer, followed by LPC(18:0) and LPC(18:1)^66^. In our atlas, LPC(16:0), LPC(18:1), LPE(18:0), LPE(18:1), and LPE(20:1) were all determined to be (positively correlating) biomarker candidates for distal tubules, providing a lipid landscape capable of driving ROS production for modulation of ion channel activity. ROS has been implicated in the upregulation of the distal tubule sodium-chloride cotransporter (NCC)^67^. Also, K^+^ channels in the basolateral membrane can be stimulated by prostaglandin F2α (PGF2a) in a ROS-dependent mechanism^67^. Recently, ROS was shown to modulate the distal tubule-specific Mg^2+^ channel TRPM6^68^. These observations, coupled with finding LPC and LPE molecular markers in distal tubules, point to a role for lipid-induced ROS regulation of ion channels in distal tubules.

The collecting duct is the most distal functional tissue unit of the nephron and is responsible for achieving the final electrolyte, pH, and fluid balance of urine before reaching the calyces and renal pelvis. The collecting duct is composed of principal and intercalated cells. Principal cells express ENaC and Na+/K+ ATPase to regulate sodium reabsorption and potassium secretion. Intercalated cells primarily participate in acid-base secretion and reabsorption and are part of a class of proton-secreting cells known as mitochondria-rich cells^69^. Consistent with the presence of mitochondria-rich cells, phosphatidylcholines and phosphatidylethanolamines are highly enriched in the collecting duct. The outer mitochondrial membrane (OMM) is rich in PC and PE, while the inner mitochondrial membrane (IMM) contains PE^70^. PC is well suited for the smooth OMM as its physical properties make it ideal for forming a smooth bilayer^70^. The IMM lipidome is rich in non-bilayer forming PE to maintain stability and render it impermeable to solutes^70^. Given the critical role these phospholipids play in the structure-function of mitochondria and their selective enrichment in collecting ducts, their detection as strong biomarker candidates in this atlas is likely aligned with the mitochondria-rich intercalated cells of this FTU.

The phytosphingosine ceramide SHexCer(t18:0/h24:0) also has discriminative positive association with the collecting duct. Phytosphingosine ceramides are unique in that they are 4-hydroxylated molecules. These phytoceramides are found in the skin barrier, where it is proposed that they introduce heterogeneity into the membrane and expand the mechanisms of permeability restrictions^71^. While their membrane packing is less ordered, the hydrogen bonding of their head group adds structure to the membrane in a manner that is divergent from other ceramides^72^. Given that the collecting duct must maintain strict permeability regulation, it is intriguing that the same phytoceramide that comprises the skin barrier is detected here. Nakashima et al. also observe SHexCer(t18:0/h24:0) in collecting ducts and propose that its hydroxylation state is critical for regulating the transport of NH_3_ and H^+^ by strengthening lateral interactions of the plasma membrane. Long-chain sphingomyelin species that can stabilize membrane domains^73^ are also found to be associated with the collecting ducts. Taken together, the phytosphingosine ceramide and sphingomyelin species of the collecting duct could create leakproof but flexible membranes for the regulation of ammonia transport.

### Molecular characteristics of sexual dimorphism

Our analysis of the atlas suggests that lipidomic signatures of females are enriched with increased PC and LPC compared to males. This enhanced association of phosphatidylcholines with females could be attributed to phosphatidylethanolamine-*N*-methyltransferase (PEMT) levels, which are regulated by estrogen^74^. Our SHAP-based analysis also found arachidonic acid-containing phospholipids to be marker candidates for female donor tissue. Arachidonic acid is liberated from phospholipids by phospholipases and can be metabolized into prostanoids PGE_2_ and PGI_2_, which act in the kidney to regulate blood pressure. Studies have shown that females produce higher levels of PGE2 and that these levels correlate with lower blood pressure compared to age-matched males^75^. The observation that phosphatidylcholines are key components of the molecular signature of the kidney for females indicates that this important class of lipids should be investigated further related to hormonal regulation of PEMT and the sexual dimorphism of prostanoids and blood pressure regulation.

### Lipid signatures in normal and obese donors

In the comparison of tissues from donors classified as obese versus normal based on BMI values, lysophosphatidylethanolamines were one of the primary lipid classes to show an association with normal BMI donor tissues. This is consistent with previous observations that lysophosphatidylethanolamines are negatively correlated with BMI^76^. While SHAP analyses of FTU and sex detected differences in overall lipid class enrichments, the lipid profiles when using BMI as a classifier showed a pronounced difference in specific PC and SM molecules. This may be a function of sex and FTU being physiologically stable aspects while BMI occurs on a spectrum and is more heterogenous throughout the population. The strongest biomarker for obesity was PC(32:1), which has previously been shown to be associated with BMI^77^. Sphingomyelins also showed differences in their glomerular profiles of obese compared to normal donors. SM(18:1;2O/16:0)’s discriminative association with obesity combined with an elevated detection of glomerular sclerosis in this population suggests a mechanistic link between alterations in sphingomyelin lipids and the development of glomerular disease.

### Future directions and refinements

The kidney lipid atlas that we have constructed provides a publicly available, comprehensive foundation for exploring the relationships between spatio-molecular information, defined functional tissue units of the nephron, histopathological data, and key donor characteristics. Currently, these analyses provide detailed lipid information with functional tissue unit-level specificity. Workflows can be refined to increase the molecular breadth and structural specificity through multimodal and multi-omic assay integration^15^. The ability to connect deep molecular information to specific tissue features with improved granularity, from functional tissue units to specific cell types and eventually single cells, is necessary for building multiscale molecular atlases of human organs associated with normal aging and disease. This goal is now central to the efforts of many large-scale research consortia, such as the Human Biomolecular Atlas Project (HuBMAP)^4,7^, Kidney Precision Medicine Project (KPMP)^11^, Human Tumor Atlas Network^12^, and others. The broader goals of these research efforts and, indeed, the focus of future releases of this kidney lipid atlas is the expansion of our MALDI IMS lipid coverage and integration of additional modalities to capture cellular organization and additional molecular classes more thoroughly. For example, multiplexed immunofluorescence microscopy assays use dozens of well-characterized protein markers to identify distinct cell types and their organization at cellular resolution. On the other hand, transcriptomics and mass spectrometry-based proteomics assays are often collected from bulk samples (no spatial information) or in a profiling mode with limited spatial resolution targeting anatomical features or specific functional tissue units. High spatial resolution approaches to transcriptomic and mass spectrometry-based proteomics are emerging, but with limited molecular coverage. An integrated approach, building on what we have developed as part of the current version of the kidney atlas, will be necessary to refine the atlas in later iterations and leverage the strengths of many individual assays to provide a comprehensive picture of how molecular distributions relate to cellular organization and localized changes in biomolecular pathways in normal and diseased kidney tissue.

## METHODS

### Materials

Acetone, isopentane, tetrahydrofuran (THF), acetonitrile, and methanol were purchased from Fisher Scientific (Pittsburgh, PA, USA). 1,5-Diaminonaphthalene (DAN) and carboxymethylcellulose were purchased from Sigma-Aldrich Chemical Company (St. Louis, MO, USA).

### Sample Preparation

Human kidney tissue was surgically removed during a full nephrectomy and remnant tissue was processed for research purposes by the Cooperative Human Tissue Network at Vanderbilt University Medical Center. Remnant biospecimens were collected in compliance with the Cooperative Human Tissue Network standard protocols and National Cancer Institute’s Best Practices for the procurement of remnant surgical research material.

Participants were consented for remnant tissue collection in accordance with institutional IRB policies. Donor demographics including age, sex, race, weight, height, BMI, and comorbidities are recorded in Table S1. Half of the excised tissue was flash frozen over an isopentane-dry ice slurry, embedded in carboxymethylcellulose, and stored at-80 °C until use. Kidney tissues were cryosectioned to a 10 µm thickness and thaw mounted onto indium tin-oxide (ITO) coated glass slides (Delta Technologies, Loveland, CO, USA) for IMS analysis or regular glass slides for histological staining. Slides were stored at-80 °C and returned to ∼20 °C within a vacuum desiccator prior to further processing. The remaining excised tissue was formalin fixed and paraffin embedded (FFPE). FFPE tissue was sectioned to a 10 µm thickness and stained using periodic acid Schiff staining methods for pathological assessment. In brief, the percentage of cortex, medulla, and other structures was determined within the tissue section. Additionally, a histological assessment is reported in Table S2. Additional method details can be found in these protocol references^78–82^.

### Autofluorescence Microscopy and tissue segmentation

Autofluorescence microscopy images were acquired on each tissue prior to analysis using DAPI, EGFP, and DsRed filters on a AxioScan Z1 slide scanner (Carl Zeiss Microscopy GmbH, Oberkochen, Germany)^83^. After IMS analysis, an additional autofluorescence image was acquired using brightfield and EGFP filters to locate MALDI laser ablation marks for image registration between the marks and their corresponding MALDI IMS pixels. Additional methods can be found here^84^. Segmentation of FTUs based on autofluorescence images were performed as described previously^17^.

### Imaging Mass Spectrometry

Samples for IMS analysis were coated with a 20 mg/mL solution of DAN dissolved in THF using a TM Sprayer M3 (HTX Technologies, LLC, Chapel Hill, NC, USA) yielding a 1.67 mg/cm^2^ coating (0.05 mL/hr, 4 passes, 40 °C spray nozzle). Tissue samples were imaged immediately after matrix deposition^85^. MALDI IMS was performed on a timsTOF fleX mass spectrometer (Bruker Daltonics, Bremen, Germany). The ion images were collected in positive and negative ion modes at 10 µm pixel size with the beam scan set to 6 µm^2^ using 150 laser shots per pixel and 18.6% laser power (30% global attenuator and 62% local laser power) at 10 kHz. Between 75,150 to 400,250 pixels were acquired from each tissue. Data were collected from *m/z* 1500 – 2000 for lipid analysis. Imaging data were visualized using SCiLS Lab Version 2019 (Bruker Daltonics, Bremen, Germany) or custom software described below. Lipids were identified using a combination of mass accuracy (≤ 5 ppm), orthogonal LC-MS experiments, and LIPIDMAPS^86,87^ database searching. Approximately 200 lipids were identified from each patient sample.

### LC-MS/MS

Three to five 10 µm sections were collected per patient into Eppendorf tubes. Samples were transferred into glass vials with 200 µL of methanol. Lipids were extracted using a modified MTBE protocol^88^. 200 µL of cold methanol were added to samples on ice and vortexed for 1 minute. 800 µL of cold MTBE was added and samples were homogenized by sonication in four rounds of 10 minutes each on ice. 200 µL of cold water were then added and samples were allowed to rest for 10 minutes. Samples were centrifuged at 1000 xg for 10 minutes at 4 °C and the upper organic layer was removed and evaporated to dryness. Samples were reconstituted in 500 µL methanol for analysis with 500 ppb Avanti Equisplash internal standards. A Thermo Scientific Q Exactive HF with a Vanquish UHPLC+ and a Bruker TimsTof FleX mass spectrometer with a Waters Premier UHPLC were used for analysis with a Waters Premier reversed-phase C18 BEH column (2.1 mm x 100 mm) at 50 °C. Lipids were separated with a binary gradient of (A) 10 mM ammonium acetate and 0.1% formic acid in water/acetonitrile (4:6) and (B) 10 mM ammonium acetate and 0.1% formic acid in 2-propanol/acetonitrile (9:1) with a flow rate of 250 µL/min for 15 minutes on the Thermo system and 30 minutes on the Bruker system. Ten µL were injected in both negative and positive ion modes. Samples pooled to create a reference sample for QC. Data was analyzed and annotated in MS-DIAL version 4.90^89^.

### MALDI IMS Data Preprocessing

MALDI IMS data was processed using in-house developed software. All datasets were first converted into a custom binary format optimized for storage and speed of analysis of IMS data. Each spectrum was *m/z* aligned using between 6 and 10 alignment peaks that were automatically selected for each dataset. The alignment peaks are selected by their frequency of occurrence throughout the dataset and ensure that several anchor points are available for each spectrum. This is in contrast to using a pre-defined set of peaks that might not be present in every spectrum, hindering the alignment process. The alignment process was accomplished using the Python *msalign* library (v0.2.0)^90,91^. Subsequently, the spectra were calibrated using at least four calibration points in both polarities. Two vectors of normalization factors were calculated for each dataset, where the total ion current (TIC) and outlier-insensitive TIC variant that only includes data lying between the 5th and 95th percentiles of each mass spectrum (5/95% TIC). Normalization aims to counteract noise factors and to project all mass spectra onto a consensus intensity scale so that intensities can be compared between spectra. Since the subsequent analyses compare multiple datasets from different patients, the normalizations of a single dataset have no knowledge of the intensity scales another dataset has, which means that if normalization is performed on a dataset-by-dataset basis, ion intensities in one dataset might not be comparable to ion intensity of another. To counteract this, all normalization vectors were concatenated (TIC and 5/95% TIC were processed separately) into one large vector which was divided by its median factor. This ensures that intensities are comparable between datasets. At this point, rather than creating a copy of the data for both normalization factors, we keep the IMS data in the *m/z* aligned and calibrated state and apply the normalization factors when needed. Following the pre-processing, we calculate the average mass spectrum for each dataset which is subsequently annotated using in-house developed software (see below). The unsupervised and classification workflows (discussed in depth below) require that a common set of features (peaks) is present for each analyzed dataset, hence a set of annotated peaks was used to extract ion images from the processed binary data. The data was extracted into a two-dimensional matrix of shape (N x M) where N is the number of pixels and M is the number of identified peaks. An image is created by summing the intensity of a specified peak with a ± 3-5ppm extraction window. Before further analysis, the image intensity matrix is normalized using one of the available normalization factors.

### MALDI IMS Tentative Identification

The average mass spectrum was scaled between 0 and 1, peak picked, deisotoped and filtered using a signal-to-noise ratio (S/N) threshold of 0.03 (calculated as fraction of the base peak of the mass spectrum). Features below this threshold were excluded from identification process. IMS identification was performed using in-house developed software that associated detected peaks with tentative lipids from the LIPID MAPS Structure Database (LMSD) and custom LC-MS/MS database (discussed above). Parameters for annotating peaks include [M+H]^+^, [M+Na]^+^ and [M+K]^+^ adducts in positive and [M-H]^-^ and [M-CH_3_]^-^ in negative mode and search window of ± 5ppm. To provide more confident identification, IMS identification was subsequently associated with LC-MS/MS identifications.

### MALDI IMS-Microscopy and Microscopy-Microscopy Image Registration

MALDI IMS pixels were registered to microscopy using *IMS MicroLink*, an in-house developed, open-source plugin developed for the napari image viewer. Within *IMS MicroLink*, the theoretical coordinate (.i.e., the x,y integer coordinates) of each IMS pixel was extracted from the IMS metadata and visualized as a an IMS pixel map with randomized intensities and the post-AF image was registered to the IMS pixel map by selecting 5 corresponding fiducials where a fiducial is an IMS pixel in the IMS pixel map image and a laser ablation mark in the post-AF image. This creates an exact registration of the IMS pixel to its origin in microscopy coordinates, the laser ablation mark. After alignment of the postAF image to IMS, the pre-acquisition AF (preAF (IMS)) image is automatically registered to the registered postAF image (PostAF (IMS)) using *wsireg*, an in-house developed, open-source Python package to perform whole slide image registration. The CODEX image is also automatically registered with *wsireg* to the postAF (IMS) image by concatenating transformations from modality to modality. After all registrations and alignments are completed, all microscopy images and IMS are sampled into a common coordinate space.

### Unsupervised Machine Learning

Uniform Manifold Approximation and Projection (UMAP) was used to uncover molecular trends across all datasets. UMAP aims to preserve the global neighborhood structure of the data while producing a low-dimensional embedding. UMAP embedding was performed at pixel level where each pixel across all samples was considered as an observation (6,779,166 in positive and 6,568,017 in negative mode) and at FTU-level where an average profile was obtained for each detected FTU (78,190 in positive and 74,959 in negative mode). In both cases, 211 and 212 features were used in positive and negative modes, respectively.

UMAP projection was performed in Python using the umap-learn (v0.5.2)^21,92^ and scikit-learn (v1.0.2)^93^ libraries. In our analysis, the purpose of UMAP was to identify a low-dimensional projection, hence, the number of target dimensions was limited to 2 (n_components=2). Several other parameters were adjusted, including the distance metric (metric=’cosine’), minimum number of neighbors (n_neighbors=250) and the minimal distance (min_dist=0). The random seed was also specified to allow us to reproduce the results.

Due to the multi-dataset nature of the data, two normalization approaches were explored. First, standard TIC normalization was used which results in inter-normalized data. The second approach was to remove inherent batch effects caused by the fact that each sample came from different patients with vastly different BMI, age or gender. Batch effects were removed using the reComBat algorithm^20^.

### Supervised Machine Learning & Shapely additive explanations

The identification of biomarker candidates was performed computationally following a workflow similar to that presented in [Tideman, 2021]. The first step of the workflow consists in learning a classification model from a set of labeled IMS datasets via supervised machine learning – 28 datasets in negative ion mode and 29 in positive ion mode. The following classification tasks were performed: recognition (one-versus-all classification) of five FTUs (glomeruli, distal tubules, proximal tubules, collecting ducts, thick ascending limb), binary classification of male versus female donors, and binary classification of normal-BMI (22 < BMI < 25) versus high-BMI (BMI > 35) donors. In the case of the FTU recognition task, segmentation by autofluorescence provided pixel-wise labeling of the IMS data. In the case of the sex and BMI classification, each IMS dataset was labeled using its corresponding donor metadata (female or male, high-BMI or low-BMI). We choose eXtreme Gradient Boosting (XGBoost) models [Chen, 2016] for the classification of IMS data for the following reasons: XGBoost can encode non-linear dependencies between molecular species, does not make assumptions about the data distribution, does not depend on measuring distances between high-dimensional mass spectra, and does not require the scaling of ion intensity values. Rather than use one XGBoost model, as per Tideman^22^, we use an ensemble of XGBoost models for each classification task. The reason for using an ensemble of XGBoost models is to ensure that our SHAP-based biomarker candidate discovery workflow is robust to correlated inputs (e.g. molecular species with similar ion images). The problem posed by correlated inputs is that multiple models may fit the data equally well without necessarily being faithful to the true biological relationships. There is therefore a risk that spurious correlations influence the results produced by SHAP on one single model, and that the results vary from one model to the next. Given that our aim is to select a shortlist of biologically relevant candidate biomarkers, we cannot allow for correlation between molecular species to cause inconsistencies. The solution we propose is to stabilize the output of SHAP by applying it to an ensemble of ten XGBoost models. The training process of each of these XGBoost models is initialized using a different random seed, and the models are trained on slightly different training datasets (the split between training and testing datasets is done by sampling without replacement with ten different random seeds). Spurious patterns due to correlated inputs cancel out across the different XGBoost models, whereas the true correlations are reinforced by the ensembling process. The patterns that dominate the decision-making process (and the ensuing SHAP explanations) of the ensemble are therefore likely to be biologically relevant.

The second step of the workflow consists in quantifying the importance of each molecular species with respect to a given classification task using Shapley additive explanations (SHAP)^23^. We apply SHAP to the ensemble of 10 XGBoost models obtained in the previous step. The Shapley value, or local SHAP importance score, of a molecular species for a given mass spectrum is the mean of the Shapley values obtained by applying SHAP to the 10 classification models making up the ensemble. As discussed in Tideman^22^, we have one Shapley value per molecular species and per pixel: the Shapley value of a molecular species measures the magnitude and direction of its effect on the classification of one spatially-localized mass spectrum. We have multiple IMS datasets, each of which corresponds to the renal tissue sample of a different donor. The datasets are labeled according to the donor’s metadata: VAN-donor ID-donor age-donor sex-donor BMI. The global SHAP importance score is a dataset-wide summary measure of a molecular species’ biomarker potential that is computed by taking the mean of the magnitude of its Shapley values across all mass spectra in a given dataset. The global SHAP importance score is a donor-specific measure of a molecular species’ relevance to a given classification task. In the summary bubble plots (Figures 6-8), the size of each marker corresponds to the global SHAP importance score of a given molecular species (column) for a given dataset (row). Finally, the total SHAP score (*SHAP_tot_*) of a molecular species is an experiment-wide summary measure of its biomarker potential: it is computed by taking the mean of the molecular species’ global SHAP scores across all datasets. The molecular species are ranked in descending order of total SHAP score in order to facilitate the selection of a shortlist of molecular species that are likely to be useful biomarkers of a given FTU, sex or BMI.

The next question is whether a biomarker candidate is positively or negatively correlated with a given FTU, sex or BMI. We measure the direction and magnitude of the relationship by computing the Spearman rank-order correlation coefficient ***ρ*** between the mean-centered intensity and the Shapley values of a given molecular species. The Spearman rank-order correlation coefficient ***ρ*** also provides a way of assessing the statistical significance of the relationship: ***ρ*** ranges from-1 to 1, and we consider ***ρ*** to be significant if its magnitude exceeds 0.2. In the summary bubble plots (Figures 6-8), the marker color corresponds to the Spearman rank-order correlation coefficient per molecular species (column) and per dataset (row). We distinguish between the following three scenarios:

- Positive Spearman rank-order correlation coefficient (***ρ*** > 0.2)-red marker: pixels with high (above the mean) ion intensity have positive Shapley values and, conversely, pixels with low (below the mean) ion intensity have negative Shapely values. A pixel’s likelihood of belonging to the target class increases monotonically as the ion intensity of the molecular species increases. A high intensity of the molecular species is indicative of the target characteristic.
- Negative Spearman rank-order correlation coefficient (***ρ*** <-0.2)-blue marker: pixels with low (below the mean) ion intensity have positive Shapley values and, conversely, pixels with high (above the mean) ion intensity have negative Shapley values. A pixel’s likelihood of belonging to the target class decreases monotonically as the ion intensity of the molecular species increases. A low intensity of the molecular species is indicative of the target characteristic.
- Non-significant Spearman rank-order correlation coefficient (−0.2 < ***ρ*** < 0.2)-gray marker: the relationship between the molecular species and the target class is either monotonic but non-significant, or non-monotonic (and therefore not captured by the Spearman rank-order correlation coefficient).

For example, we consider the BMI classification task, where an ensemble of XGBoost classification models is made to differentiate between high-BMI (obese) donors from normal-BMI donors. The characteristic of interest, for which we need biomarkers, is obesity. A molecular species with a high total SHAP score is one whose ion intensity distribution varies from one BMI category to the other, which enables classification. It may therefore be a useful biomarker for obesity and should be investigated further. The next step is to determine whether the molecular species is positively or negatively correlated with our characteristic of interest. If the ion intensity of a given molecular species is higher in high-BMI donors, the relationship between ion intensity and the target characteristic is positive: a high ion intensity of the molecular species is a marker of obesity. Conversely, if it is lower in high-BMI donors, the relationship is negative: a low ion intensity of the molecular species is a marker of obesity.

## ACKNOWLEDGEMENTS

The authors thank the Hubmap data curation and ingest team, particularly Brendan Honick, Phil Blood, and Jonathan Silverstein for their support. Research reported in this publication was supported by the National Institutes of Health (NIH)’s Common Fund, National Institute Of Diabetes And Digestive And Kidney Diseases (NIDDK), and the Office Of The Director (OD) under Award Numbers U54DK120058, U54DK134302, and U01DK133766 (J.M.S., R.M.C., and R.V.), by NIH’s Common Fund, National Eye Institute, and the Office Of The Director (OD) under Award Number U54EY032442 (J.M.S., R.M.C., and R.V.), by NIH’s National Institute Of Allergy And Infectious Diseases (NIAID) under Award Numbers R01AI138581 and R01AI145992 (J.M.S. and R.V.), by NIH’s National Institute On Aging (NIA) under Award Number R01AG078803 (J.M.S. and R.V.), and by the National Science Foundation Major Research Instrument Program CBET – 1828299 (J.M.S. and R.M.C.). The research was furthermore made possible in part by grant numbers 2021-240339 and 2022-309518 (L.G.M. and R.V.) from the Chan Zuckerberg Initiative DAF, an advised fund of Silicon Valley Community Foundation.

The content is solely the responsibility of the authors and does not necessarily represent the official views of the National Institutes of Health or the Chan Zuckerberg Initiative.

## Notes

https://portal.hubmapconsortium.org/

### Competing Interest Statement

The authors have declared no competing interest.

### Summary of Updates

This updated version of our lipid altlas manuscript has been refined to focus on the discovery of lipid profiles of kidney-specific fuctional tissue units.

