## Supplemental figures and tables. for "A Lipid Atlas of the Human Kidney"

Figure S14. The intensity difference spectrum between the negative ion mode average spectra for all pixels associated with distal tubules and thick ascending limb from all donor samples. …………………………..26
Figure S18. The intensity difference spectrum between the positive ion mode average spectra for all pixels associated with collecting ducts and distal tubules from all donor samples………………………………….. 28
Figure S19. The intensity difference spectrum between the positive ion mode average spectra for all pixels associated with collecting ducts and glomeruli from all donor samples………………………………………….. 28
Figure S20. The intensity difference spectrum between the positive ion mode average spectra for all pixels associated with collecting ducts and proximal tubules from all donor samples…………………………… 29
Figure S21. The intensity difference spectrum between the positive ion mode average spectra for all pixels associated with collecting ducts and thick ascending limb from all donor samples………………………… 29
Figure S22. The intensity difference spectrum between the positive ion mode average spectra for all pixels associated with distal tubules and glomeruli from all donor samples……………………………………….. 30
Figure S23. The intensity difference spectrum between the positive ion mode average spectra for all pixels associated with distal tubules and proximal tubules from all donor samples……………………………… 30
Figure S24. The intensity difference spectrum between the positive ion mode average spectra for all pixels associated with distal tubules and thick ascending limb from all donor samples………………………….. 31
Figure S25. The intensity difference spectrum between the positive ion mode average spectra for all pixels associated with glomeruli and proximal tubules from all donor samples…………………………………… 31

Figure S26. The intensity difference spectrum between the positive ion mode average spectra for all pixels associated with glomeruli and thick ascending limb from all donor samples……………………………………. 32
Figure S196. Split violin plots of the ion intensity distributions of the **top obesity biomarker candidates in obese (left) and normal (right) donor data, in negative ion mode**, approximated using kernel density estimation...183
Figure S197. 10 µm MALDI IMS images of an unknown (*m/z* 664.42) detected in negative ion mode from each donor sample....184
Figure S198. 10 µm MALDI IMS images of [GA1(d18:1/24:1)-H]- (*m/z* 1,335.852) detected in negative ion mode from each donor sample....184
Figure S199. 10 µm MALDI IMS images of [PE(38:4)-H]- (*m/z* 766.539) detected in negative ion mode from each donor sample....185
Figure S200. 10 µm MALDI IMS images of an unknown [GA1(d18:1/16:0)-H]- (*m/z* 1,225.743) detected in negative ion mode from each donor sample....185
Figure S201. 10 µm MALDI IMS images of [PS(36:1)-H]- (*m/z* 788.545) detected in negative ion mode from each donor sample....186
Figure S202. 10 µm MALDI IMS images of [PI(18:0_20:3)-H]- or [PMeOH(22:6_28:7)-H]- (*m/z* 887.56) detected in negative ion mode from each donor sample....186
Figure S203. 10 µm MALDI IMS images of [PE(O-16:1_20:4)-H]- (*m/z* 722.513) detected in negative ion mode from each donor sample....187
Figure S204. 10 µm MALDI IMS images of [PE(21:4)-H]- (*m/z* 528.273) detected in negative ion mode from each donor sample....187

Figure S205. 10 μm MALDI IMS images of [LPE(18:0)-H]$^-$ (*m/z* 480.31) detected in negative ion mode from each donor sample....188
Figure S206. 10 μm MALDI IMS images of [PE(O-38:5)-H]$^-$ (*m/z* 750.544) detected in negative ion mode from each donor sample....188
Figure S207. 10 μm MALDI IMS images of [PC(32:1)+H]$^+$ (*m/z* 732.554) detected in positive ion mode from each donor sample....189
Figure S208. 10 μm MALDI IMS images of [LPC(16:0)+H]$^+$ (*m/z* 496.34) detected in positive ion mode from each donor sample....189
Figure S209. 10 μm MALDI IMS images of [PC(36:2)+H]$^+$ or [PC(18:0_18:2)+H]$^+$ (*m/z* 786.601) detected in positive ion mode from each donor sample....190
Figure S210. 10 μm MALDI IMS images of an unknown [PC(36:3)+H]$^+$ or [PC(18:1_18:2)+H]$^+$ or [PC(34:0)+Na]$^+$ (*m/z* 784.585) detected in positive ion mode from each donor sample....190
Figure S211. 10 μm MALDI IMS images of [PC(34:1)+K]$^+$ or [PC(16:0_18:1)+K]$^+$ (*m/z* 798.541) detected in positive ion mode from each donor sample....191
Figure S212. 10 μm MALDI IMS images of [PE(P-38:4)+K]$^+$ or [PE(P-18:0_20:4)+K]$^+$ or [PE(P-16:0_22:4)+K]$^+$ (*m/z* 790.515) detected in positive ion mode from each donor sample....191
Figure S213. 10 μm MALDI IMS images of [SM(18:1,2O/16:0)+Na]$^+$ (*m/z* 725.557) detected in positive ion mode from each donor sample....192
Figure S214. 10 μm MALDI IMS images of [PC(36:4)+H]$^+$ or [PC(16:0_20:4)+H]$^+$ or [PC(18:2_18:2)+H]$^+$ or [PC(34:1)+Na]$^+$ (*m/z* 782.569) detected in positive ion mode from each donor sample....192
Figure S215. 10 μm MALDI IMS images of [PC(31:0)+H]$^+$ (*m/z* 720.554) detected in positive ion mode from each donor sample....193
Figure S216. 10 μm MALDI IMS images of [PC(O-34:4)+K]$^+$ (*m/z* 778.515) detected in positive ion mode from each donor sample....193

**Table S1.** Donor demographics and comorbidities.

| Sample ID | Age | Sex | Race | Height (cm) | Weight (kg) | BMI | Comorbidities |
|---|---|---|---|---|---|---|---|
| VAN0005 | 58 | F | Non-Hispanic White | 160 | 59 | 23 | Renal cell carcinoma |
| VAN0007 | 66 | F | Non-Hispanic White | 158.8 | 81.5 | 32.2 | Renal cell carcinoma, obese |
| VAN0008 | 48 | M | Non-Hispanic White | 193 | 131.5 | 35.3 | Renal cell carcinoma, hypertension, extremely obese, previous urinary and kidney cancer |
| VAN0010 | 56 | M | Non-Hispanic White | 167.6 | 91.2 | 32.5 | Renal cell carcinoma, hypertension, obese, soft tissue sarcoma |
| VAN0012 | 44 | F | Non-Hispanic White | 160 | 71.7 | 28 | Renal cell carcinoma |
| VAN0014 | 76 | F | Non-Hispanic White | 157.4 | 93 | 37.5 | Renal cell carcinoma, hypertension, extremely obese |
| VAN0016 | 56 | M | Non-Hispanic White | 181.6 | 91.4 | 27.7 | Oncocytoma, Medullary Fibroma, prostate cancer |
| VAN0024 | 46 | M | Non-Hispanic White | 175.2 | 68.5 | 22.3 | Renal cell carcinoma, laryngeal/throat cancer |
| VAN0025 | 55 | M | Non-Hispanic White | 181.6 | 98.9 | 30 | Renal cell carcinoma, obese |
| VAN0027 | 64 | F | Non-Hispanic White | 170 | 70 | 24.2 | Renal cell carcinoma, coronary artery disease, hypertension, breast cancer |
| VAN0028 | 56 | F | Non-Hispanic White | 175.2 | 139.6 | 45.5 | Renal cell carcinoma, morbidly obese |
| VAN0029 | 62 | M | Non-Hispanic White | 182.9 | 116.9 | 34.9 | Medullary fibroma, renal cell carcinoma, obese |
| VAN0030 | 43 | M | Non-Hispanic White | 175.3 | 83.4 | 27.1 | Renal cell carcinoma |
| VAN0031 | 77 | F | Non-Hispanic White | 162.6 | 74.6 | 28.2 | Renal cell carcinoma, hypertension, type 2 diabetes |
| VAN0033 | 51 | F | Non-Hispanic White | 165.1 | 108.9 | 40 | Renal cell carcinoma, morbidly obese |
| VAN0041 | 78 | F | Non-Hispanic White | 160 | 59 | 23 | Oncocytoma, benign adrenal gland |
| VAN0042 | 57 | M | Non-Hispanic White | 175.3 | 77.6 | 25.3 | Renal cell carcinoma |
| VAN0043* | 72 | M | Non-Hispanic White | 182.9 | 135.7 | 40.6 | Renal cell carcinoma, hypertension, morbidly obese |
| VAN0044 | 65 | M | Non-Hispanic White | 165.1 | 96.2 | 35.3 | Renal cell carcinoma, hypertension, type 2 diabetes, extremely obese |
| VAN0045 | 66 | F | Non-Hispanic White | 165.1 | 107.5 | 39.4 | Renal cell carcinoma, hypertension, type 2 diabetes, extremely obese |
| VAN0046 | 66 | M | Non-Hispanic White | 177.8 | 74.8 | 23.7 | Renal cell carcinoma |
| VAN0048 | 70 | M | Non-Hispanic White | 172 | 73.5 | 24.8 | Renal cell carcinoma, hypertension |
| VAN0049 | 54 | F | Non-Hispanic White | 160 | 113.9 | 44.5 | Renal cell carcinoma, hypertension, arrhythmia, fibromyalgia, morbidly obese |
| VAN0050 | 64 | F | Non-Hispanic White | 160 | 77.1 | 30.1 | Renal cell carcinoma, hypertension, hyperlipidemia, depression, obese |
| VAN0051 | 67 | F | Non-Hispanic White | 157.5 | 68 | 27.4 | Renal cell carcinoma |
| VAN0052 | 56 | M | Non-Hispanic White | 177.8 | 79.6 | 25.2 | Renal cell carcinoma, hypertension, type 2 diabetes |
| VAN0053 | 20 | F | Non-Hispanic White | 172.7 | 68 | 22.8 | Ewing Sarcoma |
| VAN0054 | 68 | M | Non-Hispanic White | 177.8 | 79.4 | 25.1 | Renal cell carcinoma, hypertension |
| VAN0063 | 64 | M | Non-Hispanic White | 162.6 | 87.5 | 33.1 | Renal cell carcinoma, hypertension, obese |

* VAN0043 was only analyzed by MALDI IMS in negative ion mode.

**Table S2.** Histology Assessment of donor tissue blocks.

| Sample Number | Specimen Composition | | Global Glomeruli Sclerosis | Other Glomerular Lesions | Arteriosclerosis | Arteriolar Hyalinosis | Interstitial Fibrosis Tubular Atrophy in Cortex | Interstitial Inflammation in Cortex |
|---|---|---|---|---|---|---|---|---|
| | Cortex (%) | Medulla (%) | % | Segmental Lesions (%) | 0-3 | 0-3 | % | % |
| VAN0005 | 78.0 | 22.0 | 7.0 | 0 | 3 | 0 | 2.59 | 0.76 |
| VAN0007 | 41.6 | 58.4 | 64.0 | 0 | 3 | 3 | 20.92 | 0.28 |
| VAN0008 | 68.5 | 31.5 | 0.5 | 0 | 2 | 0 | 0.65 | 0.04 |
| VAN0010 | 70.7 | 29.3 | 6.0 | 0 | 3 | 3 | 5.74 | 0.1 |
| VAN0012 | 76.3 | 23.7 | 2.2 | 0 | 1 | 0 | 0.35 | 0.0 |
| VAN0014 | 59.3 | 40.7 | 37.8 | 0 | 3 | 3 | 10.57 | 0.91 |
| VAN0016 | 50.6 | 49.4 | 1.7 | 0 | 2 | 0 | 0.03 | 0.03 |
| VAN0024 | 33.5 | 66.5 | 1.1 | 0 | 1 | 1 | 0.20 | 0 |
| VAN0025 | 76.3 | 23.7 | 2.2 | 0 | 2 | 1 | 0.84 | 0.02 |
| VAN0027 | 57.1 | 42.9 | 14.7 | 0 | 3 | 3 | 18.1 | 0 |
| VAN0028 | 63.0 | 37.0 | 19.9 | 0 | 3 | 1 | 1.00 | 0 |
| VAN0029 | 77.9 | 22.1 | 4.3 | 0 | 3 | 3 | 0.56 | 0 |
| VAN0030 | 67.2 | 32.8 | 1.1 | 0 | 2 | 3 | 0.53 | 0.02 |
| VAN0031 | 83.1 | 16.9 | 38.8 | 0 | 3 | 3 | 24.0 | 0.04 |
| VAN0033 | 54.1 | 45.9 | 0.8 | 0 | 1 | 1 | 0.67 | 0.08 |
| VAN0041 | 57.6 | 42.4 | 12.1 | 0 | 3 | 1 | 1.50 | 0 |
| VAN0042 | 60.9 | 39.1 | 7.7 | 0 | 1 | 0 | 0.91 | 0 |
| VAN0043 | 68.5 | 31.5 | 30.8 | 0 | 1 | 3 | 11.1 | 0 |
| VAN0044 | 72.3 | 27.7 | 37.4 | 0 | 3 | 2 | 11.9 | 0 |
| VAN0045 | 55.5 | 44.5 | 11.8 | 0 | 1 | 0 | 4.0 | 0.2 |
| VAN0046 | 54.2 | 45.8 | 4.4 | 0 | 2 | 3 | 3.3 | 0 |
| VAN0048 | 59.8 | 40.2 | 14.4 | 0 | 3 | 1 | 4.61 | 0.01 |
| VAN0049 | 58.8 | 41.2 | 3.5 | 0 | 3 | 1 | 1.64 | 0.00 |
| VAN0050 | 64.0 | 36.0 | 4.4 | 0 | 2 | 0 | 1.81 | 0.00 |
| VAN0051 | 62.5 | 37.5 | 6.9 | 0 | 3 | 1 | 2.91 | 0 |
| VAN0052 | 67.5 | 32.5 | 3.2 | 6.8 | 1 | 3 | 82.27 | 0.00 |
| VAN0053 | 58.2 | 41.8 | 0.9 | 0 | 1 | 0 | 0.31 | 0.00 |
| VAN0054 | 61.4 | 38.6 | 18.1 | 0 | 3 | 3 | 14.28 | 0 |
| VAN0063 | 67.9 | 32.1 | 1.1 | 0 | 1 | 0 | 0.70 | 0 |

**Table S3.** DOI for all modalities.

| Sample ID | AF DOI | PAS DOI | IMS-neg DOI | IMS-pos DOI |
|---|---|---|---|---|
| VAN0005 | 10.35079/HBM836.DHDQ.639 | 10.35079/HBM396.XMSB.623 | 10.35079/HBM644.HLRW.739 | 10.35079/HBM773.XBNC.394 |
| VAN0007 | 10.35079/HBM966.RLGZ.875 | 10.35079/HBM884.QGXZ.934 | 10.35079/HBM777.STQB.672 | 10.35079/HBM453.RTXK.849 |
| VAN0008 | 10.35079/HBM396.ZHCB.959 | N/A | 10.35079/HBM477.FNGG.257 | 10.35079/HBM477.FNGG.257 |
| VAN0010 | 10.35079/HBM533.ZJKS.654 | N/A | 10.35079/HBM758.SQVJ.597 | 10.35079/HBM886.MPTX.392 |
| VAN0012 | 10.35079/HBM445.VDTN.654 | 10.35079/HBM337.ZVPK.369 | 10.35079/HBM554.PGRJ.942 | 10.35079/HBM759.QJTL.573 |
| VAN0014 | 10.35079/HBM335.SQFN.396 | 10.35079/HBM389.KJJP.875 | 10.35079/HBM265.RKZN.565 | 10.35079/HBM377.MFPF.328 |
| VAN0016 | 10.35079/HBM729.GWPJ.492 | 10.35079/HBM957.LDNV.942 | 10.35079/HBM862.DKNM.597 | 10.35079/HBM543.RWZD.568 |
| VAN0024 | 10.35079/HBM576.NKHD.888 | 10.35079/HBM796.FZWQ.899 | 10.35079/HBM747.PXSM.373 | 10.35079/HBM787.DXFR.498 |
| VAN0025 | 10.35079/HBM383.KJZV.973 | 10.35079/HBM852.NQRZ.779 | 10.35079/HBM493.VRQM.643 | 10.35079/HBM869.RKXX.583 |
| VAN0027 | 10.35079/HBM455.BSJM.532 | 10.35079/HBM338.RFPS.979 | 10.35079/HBM544.VSBJ.293 | 10.35079/HBM799.DZFZ.778 |
| VAN0028 | 10.35079/HBM863.KDGD.244 | 10.35079/HBM745.BCRF.958 | 10.35079/HBM578.WFKM.657 | 10.35079/HBM938.XFGN.695 |
| VAN0029 | 10.35079/HBM722.XVHQ.385 | 10.35079/HBM486.KPCH.364 | 10.35079/HBM749.MMKH.463 | 10.35079/HBM784.HQQK.452 |
| VAN0030 | 10.35079/HBM659.TDXR.629 | N/A | 10.35079/HBM252.SRFF.799 | 10.35079/HBM966.RXFK.673 |
| VAN0031 | 10.35079/HBM495.PCPM.434 | 10.35079/HBM538.XVLP.889 | 10.35079/HBM393.RNFV.949 | 10.35079/HBM568.VXSK.299 |
| VAN0033 | 10.35079/HBM733.VXST.856 | 10.35079/HBM654.GRHB.837 | 10.35079/HBM975.BJRS.622 | 10.35079/HBM348.NCLT.522 |
| VAN0041 | 10.35079/HBM328.KRPF.865 | 10.35079/HBM346.WRZP.637 | 10.35079/HBM337.XRWN.985 | 10.35079/HBM884.RCMT.655 |
| VAN0042 | 10.35079/HBM888.PHHS.299 | 10.35079/HBM964.CZLM.657 | 10.35079/HBM437.BGCD.226 | 10.35079/HBM662.MPBS.458 |
| VAN0043 | 10.35079/HBM272.DKBX.433 | 10.35079/HBM987.NCDW.845 | 10.35079/HBM962.PZXB.965 | 10.35079/HBM856.CDPK.464 |
| VAN0044 | 10.35079/HBM645.FQSQ.837 | 10.35079/HBM838.HGWL.498 | 10.35079/HBM229.MHGL.765 | 10.35079/HBM793.CTLW.723 |
| VAN0045 | 10.35079/HBM992.HCFC.472 | 10.35079/HBM535.BDNV.724 | 10.35079/HBM925.WHKF.243 | 10.35079/HBM459.PHNF.976 |
| VAN0046 | 10.35079/HBM578.WQFJ.765 | 10.35079/HBM593.DVGN.469 | 10.35079/HBM794.JRXQ.455 | 10.35079/HBM393.LXBW.274 |
| VAN0048 | 10.35079/HBM236.DQSD.964 | N/A | 10.35079/HBM632.SGXL.444 | 10.35079/HBM447.SQFP.455 |
| VAN0049 | 10.35079/HBM576.CBTX.826 | 10.35079/HBM738.MXDD.328 | 10.35079/HBM286.ZZNV.536 | 10.35079/HBM856.RFNW.734 |
| VAN0050 | 10.35079/HBM759.JWGJ.636 | 10.35079/HBM837.XGBD.427 | 10.35079/HBM494.CRGT.577 | 10.35079/HBM963.DBXF.369 |
| VAN0051 | 10.35079/HBM855.ZJVP.233 | 10.35079/HBM978.CGGC.988 | 10.35079/HBM349.DTWT.383 | 10.35079/HBM928.HZCW.757 |
| VAN0052 | 10.35079/HBM574.SCQM.248 | 10.35079/HBM594.SRGW.833 | 10.35079/HBM488.HRWV.754 | 10.35079/HBM573.JRSG.822 |
| VAN0053 | 10.35079/HBM278.FBRC.748 | 10.35079/HBM947.VKMP.764 | 10.35079/HBM959.MGHL.844 | 10.35079/HBM556.CGCS.879 |
| VAN0054 | 10.35079/HBM438.VGFH.887 | 10.35079/HBM846.FTPG.677 | 10.35079/HBM472.NWBK.884 | 10.35079/HBM782.SCDT.743 |
| VAN0063 | 10.35079/HBM353.XGRQ.833 | 10.35079/HBM864.MFDZ.983 | 10.35079/HBM946.ZHFC.683 | 10.35079/HBM856.FWBK.447 |

**Table S4.** Lipid Annotations for Selected Species as Determined by Shapley Additive Analysis.

| Negative | IMS m/z | Annotation | LC-MS/MS (y/n) | MS/MS | Dot Product score | MSDIAL Total score | Theoretical m/z | ppm error |
|---|---|---|---|---|---|---|---|---|
| | 478.294 | LPE 18:1 (-H) | Y | | 826 | 1198 | 478.2939 | 0.20907647 |
| | 480.31 | LPE 18:0 (-H) | Y | | 830 | 1259 | 480.3096 | 0.83279618 |
| | 500.281 | LPE 20:4 (-H) | Y | | 684 | 1118 | 500.2783 | 5.396996032 |
| | 506.325 | LPE 20:1 (-H) | Y | | 818 | 995 | 506.3252 | 0.395003053 |
| | 528.273 | PE 21:4 (-H) | N | | | | 528.2732 | 0.378591986 |
| | 554.289 | LPE-N (FA 3:0) (-H) | N | | | | 554.2888 | 0.360822734 |
| | 556.304 | PE 23:4 (-H) or PC 21:4 (-CH3) | N | | | | 556.3045 | 0.8987883434 |
| | 616.471 | LPC O-26:2 (-H) or CerP(d18:1/16:0) (-H) | N | | | | 616.4712 | 0.324427159 |
| | 634.409 | PC 25:1;O (-CH3) | N | | | | 634.409 | 0 |
| | 647.466 | PA 32:0 (-H) | Y | | 725 | 1186 | 647.4657 | 0.463345008 |
| | 664.42 | PAz-PC (-H) | N | SORI | | | | |
| | 673.481 | PA 16:0_18:1 (-H) | Y | | 734 | 1219 | 673.4814 | 0.5939288 |
| | 687.545 | SM 34:1;2O (-CH3) | Y | | 817 | 1216 | 687.5447 | 0.436335267 |
| | 701.513 | PA 36:1 (-H) | N | | | | 701.5127 | 0.427647283 |
| | 716.524 | PE 16:0_18:1 (-H) | Y | | 805 | 1173 | 716.5236 | 0.558250977 |
| | 726.581 | CerP 42:2;2O (-H) | N | | | | 726.5807 | 0.412892883 |
| | 738.508 | PE 36:4 (-H) | Y | | 781 | 1177 | 738.5079 | 0.135408166 |
| | 743.607 | SM 38:1;2O (-CH3) | Y | | 790 | 1245 | 743.6073 | 0.403438751 |
| | 744.555 | PE 36:1 (-H) | Y | | 792 | 1174 | 744.5549 | 0.13430843 |
| | 750.544 | PE O-38:5 (-H) | Y | | 785 | 1254 | 750.5443 | 0.399709917 |
| | 762.508 | PE 38:6 (-H) | Y | | 795 | 1218 | 762.5079 | 0.131146182 |
| | 763.513 | Unknown | N | | | | | |
| | 764.524 | PE 38:5 (-H) | Y | | 833 | 1218 | 764.5236 | 0.523201638 |
| | 766.539 | PE 38:4 (-H) | Y | | 846 | 1229 | 766.5392 | 0.260912945 |
| | 769.623 | SM 40:2;2O (-CH3) | Y | | 781 | 1242 | 769.6229 | 0.129933764 |
| | 772.586 | SHexCer 34:4;2O (-H) | N | | | | | #DIV/0! |
| | 774.483 | SHexCer 34:3;2O (-H) | N | | | | 774.4832 | 0.258236719 |
| | 778.514 | SHexCer 34:1;2O (-H) | Y | | 586 | 1045 | 778.5145 | 0.642248796 |
| | 785.654 | SM 41:1;2O (-CH3) | Y | | 816 | 1221 | 785.6542 | 0.254564922 |
| | 788.545 | PS 36:1 (-H) | Y | | 749 | 1191 | 788.5447 | 0.380447678 |
| | 797.654 | SM 42:2;2O (-CH3) | Y | | 824 | 1251 | 797.6542 | 0.250735218 |
| | 810.529 | PS 38:4 (-H) | Y | | 729 | 1242 | 810.5291 | 0.1233762 |
| | 857.519 | PI 36:4 (-H) | Y | | 755 | 1046 | 857.5186 | 0.466462185 |
| | 862.608 | SHexCer 40:1;2O (-H) | Y | | 695 | 1060 | 862.6084 | 0.463709836 |
| | 863.566 | PI 18:0_18:1 (-H) | Y | | 741 | 1148 | 863.5655 | 0.578994877 |

| | | | | | | | | |
|---|---|---|---|---|---|---|---|---|
| | 863.692 | Unknown | N | | | | | |
| | 876.624 | SHexCer 41:1;2O (-H) | Y | | 659 | 1059 | 876.624 | 0 |
| | 878.603 | SHexCer 40:1;O3 (-H) | Y | | 702 | 1069 | 878.6033 | 0.341451028 |
| | 887.56 | PI 18:0_20:3 (-H) | Y | | 763 | 1204 | 887.5655 | 6.196725763 |
| | 888.624 | SHexCer 42:2;2O (-H) | Y | | 723 | 1063 | 888.624 | 0 |
| | 890.64 | SHexCer 42:1;2O (-H) | Y | | 716 | 1063 | 890.6397 | 0.336836546 |
| | 892.619 | SHexCer 41:1;3O (-H) | Y | | 646 | 1052 | 892.6189 | 0.112029893 |
| | 904.619 | SHexCer 42:2;O3 (-H) | Y | | 700 | 1067 | 904.6189 | 0.110543788 |
| | 906.635 | SHexCer 42:1;O3 (-H) | Y | | 702 | 1062 | 906.6346 | 0.441192075 |
| | 924.639 | SHexCer (SM4) t18:0/h24:0 (-H) | Y | | m/z 96 | | 924.6451 | 6.597125751 |
| | 1040.64 | SM3 d18:1/h22:0 (-H) | Y | | | | 1040.6561 | 15.47100911 |
| | 1225.743 | GA1 d18:1/16:0 (-H) | N | | | | 1225.7427 | 0.244749571 |
| | 1309.837 | GA1 d18:1/22:0 (-H) | N | | | | 1309.8366 | 0.305381603 |
| | 1335.852 | GA1 d18:1/24:1 (-H) | N | | | | 1335.8522 | 0.149717162 |
| | 1502.007 | CL 76:2 (-H) | N | | | | 1502.0119 | 3.262291064 |
| **Positive** | **IMS m/z** | **Annotation** | **LC-MS/MS (y/n)** | **MS/MS** | **Dot Product** | **Total score** | **Theoretical m/z** | **ppm error** |
| | 496.34 | LPC 16:0 (+H) | Y | | 824 | 1254 | 496.3398 | 0.402949753 |
| | 522.355 | LPC 18:1 (+H) | Y | | 831 | 1256 | 522.3554 | 0.765762161 |
| | 524.335 | PE 20:0 (+H) | N | | | | 524.3347 | 0.572153626 |
| | 524.371 | LPC 18:0 (+H) | Y | | 752 | 1246 | 524.3711 | 0.190704636 |
| | 588.462 | Unknown | Y | SORI | | | | |
| | 636.46 | PE 28:0 (+H) | N | | | | 636.4599 | 0.15711909 |
| | 650.439 | PC 25:1;O (+H) | Y | SORI | | | 650.4391 | 0.153742295 |
| | 666.434 | SM 34:1;2O (-TMA+Na) | Y | SORI | | | 666.4833 | 73.97034554 |
| | 689.559 | SM 33:1;2O (+H) | Y | | 774 | 1246 | 689.5592 | 0.290040362 |
| | 694.465 | SM 36:1;2O (-TMA+Na) | Y | SORI | 798 | 1248 | 694.5145 | 71.27281 |
| | 701.559 | SM 18:1;2O/16:1 (+H) | Y | | 806 | 1247 | 701.5592 | 0.285079292 |
| | 703.575 | SM 18:1;2O/16:0 (+H) | Y | | 817 | 1216 | 703.5748 | 0.284262597 |
| | 720.59 | Unknown SM | Y | | | | | #DIV/0! |
| | 725.557 | SM 18:1;2O/16:0 (+Na) | Y | | 817 | 1216 | 725.5568 | 0.27565037 |
| | 729.59 | SM 18:1;2O/18:1 (+H) | Y | | 802 | 1250 | 729.5905 | 0.685315941 |
| | 731.606 | SM 18:1;2O/18:0 (+H) | Y | | 798 | 1248 | 731.6061 | 0.136685574 |
| | 732.554 | PC 32:1 (+H) | Y | | 808 | 915 | 732.5538 | 0.27301749 |
| | 734.569 | PC 16:0_16:0 (+H) | Y | | 804 | 1237 | 734.5694 | 0.544536704 |

| | | | | | | | | |
|---|---|---|---|---|---|---|---|---|
| | 741.531 | SM 18:1;2O/16:0 (+K) | Y | | 817 | 1216 | 741.5307 | 0.40456855 |
| | 746.606 | PC O-34:1 (+H) | Y | | 558 | 964 | 746.6058 | 0.26787898 |
| | 756.551 | PC 16:0_16:0 (+Na) or PC(34:3) (+H) | Y | | 810 | 921 | 756.5538 | 3.70099258 |
| | 757.622 | SM 18:1;2O/16:0 (+H) or SM 18:2;2O/20:0 (+H) | Y | | 817 | 1216 | 757.6218 | 0.263983956 |
| | 758.569 | PC 34:2 (+H) PC 16:0_18:2 (+H) | Y | | 838 | 939 | 758.5694 | 0.527308378 |
| | 759.638 | SM 38:1;2O (+H) or SM 16:1;2O/22:0 (+H) | Y | | 790 | 1245 | 759.6374 | 0.789850526 |
| | 760.585 | PC 16:0_18:1 (+H) | Y | | 587 | 809 | 760.5851 | 0.131477727 |
| | 768.588 | PC O-34:1 (+Na) | Y | | 790 | 985 | 768.5878 | 0.260217505 |
| | 774.601 | PC 35:1 (+H) or PC 17:0_18:1 (+H) | Y | | 804 | 1119 | 774.6007 | 0.387296319 |
| | 784.585 | PC 36:3 (+H) or PC 18:1_18:2 (+H) or PC(34:0) (+Na) | Y | | 803 | 1142 | 784.5851 | 0.1274559 |
| | 785.653 | SM 40:2;2O (+H) or SM 16:1;2O/24:1 (+H) | Y | | 781 | 1242 | 785.6531 | 0.127282639 |
| | 787.669 | SM 40:1;2O (+H) or SM 18:1;2O/22:0 (+H) | Y | | 806 | 1248 | 787.6687 | 0.38087079 |
| | 788.616 | PC 36:1 (+H) or PC 18:0_18:1 (+H) | Y | | 824 | 1120 | 788.6164 | 0.507217451 |
| | 794.603 | PC O-36:2 (+Na) | Y | | 822 | 1092 | 794.6034 | 0.503395782 |
| | 801.684 | SM 41:1;2O (+H) or SM 18:1;2O/23:0 (+H) | Y | | 816 | 1221 | 801.6844 | 0.498949462 |
| | 805.679 | SM 40:0;3O (+H) | Y | | 728 | 1073 | 805.6793 | 0.372356594 |
| | 806.567 | PC 36:3 (+Na) or PC 18:1_18:2 (+Na) | Y | | 803 | 1142 | 806.567 | 0 |
| | 811.666 | SM 40:0;2O (+Na) | N | | | | 811.6663 | 0.369610023 |
| | 811.669 | SM 42:3;2O (+H) or SM 18:1;2O/24:2 (+H) | Y | | 778 | 1247 | 811.6687 | 0.36960893 |
| | 813.684 | SM 18:1;2O/24:1 (+H) or SM 42:2;2O (+H) | Y | | 824 | 1251 | 813.6844 | 0.491591089 |
| | 814.632 | PC 38:2 (+H) | Y | | 780 | 928 | 814.632 | 0 |
| | 815.7 | SM 42:1;2O (+H) or SM 18:1;2O/24:0 (+H) | Y | | 814 | 1241 | 815.7 | 0 |
| | 816.53 | PE P-40:5 (+K) or PE P-18:1_22:4 (+K) | Y | | 468 | 1245 | 816.5303 | 0.367408289 |
| | 819.695 | SM 41:0;3O (+H) | Y | | 717 | 1071 | 819.695 | 0 |
| | 823.666 | SM 41:1;2O (+Na) or SM 18:1;2O/23:0 (+Na) | Y | | 814 | 1241 | 823.6663 | 0.364225172 |
| | 831.695 | SM 42:1;3O (+H) | Y | | 716 | 1180 | 831.695 | 0 |
| | 833.711 | SM 42:0;3O (+H) | Y | | 715 | 1059 | 833.7106 | 0.479782793 |

| | | | | | | | | |
|---|---|---|---|---|---|---|---|---|
| | 835.666 | SM 42:2;2O (+Na) or SM 18:1;2O/24:1 (+Na) | Y | | 824 | 1251 | 835.6663 | 0.358994972 |

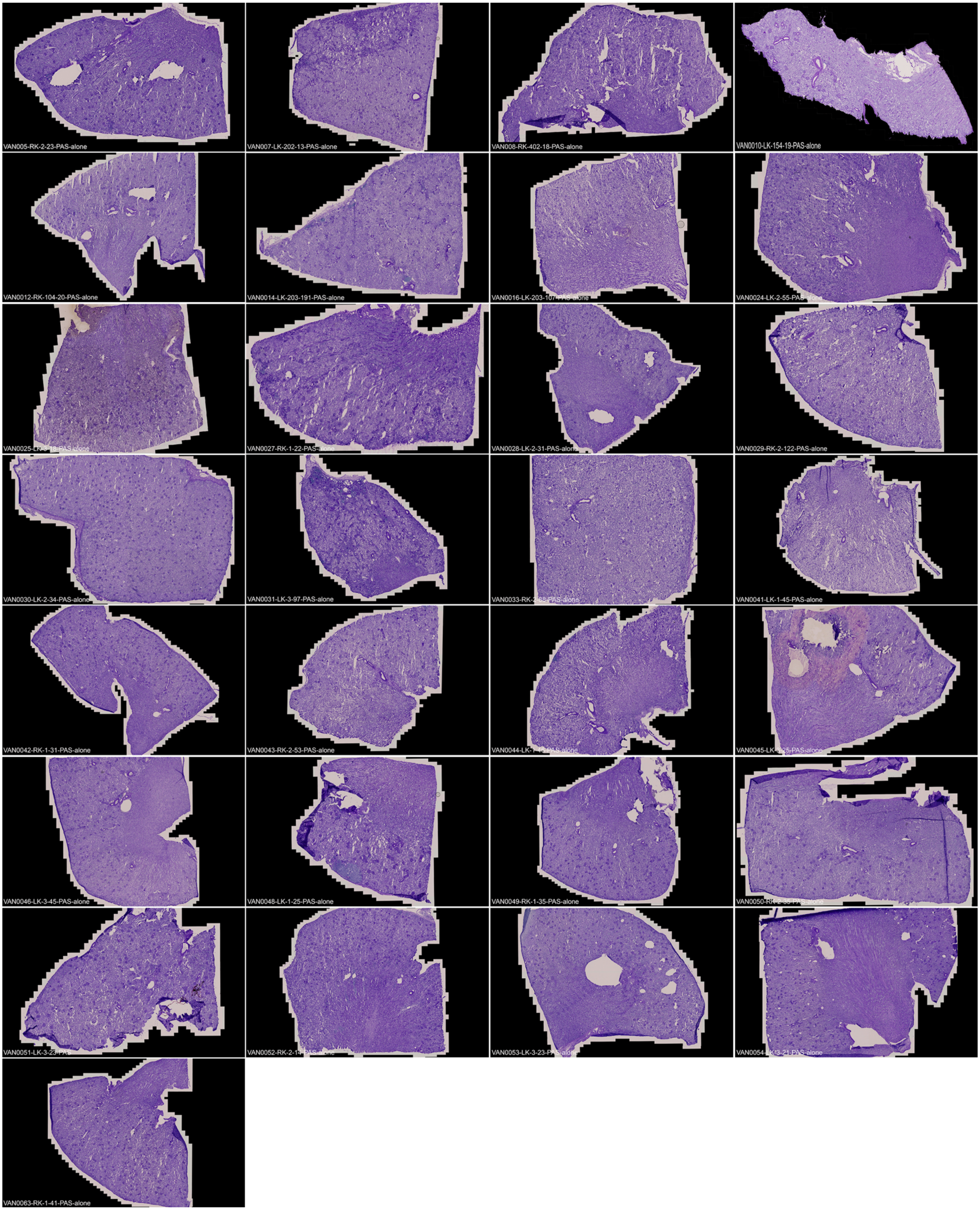

**Figure S1.** PAS Images of all donor samples.

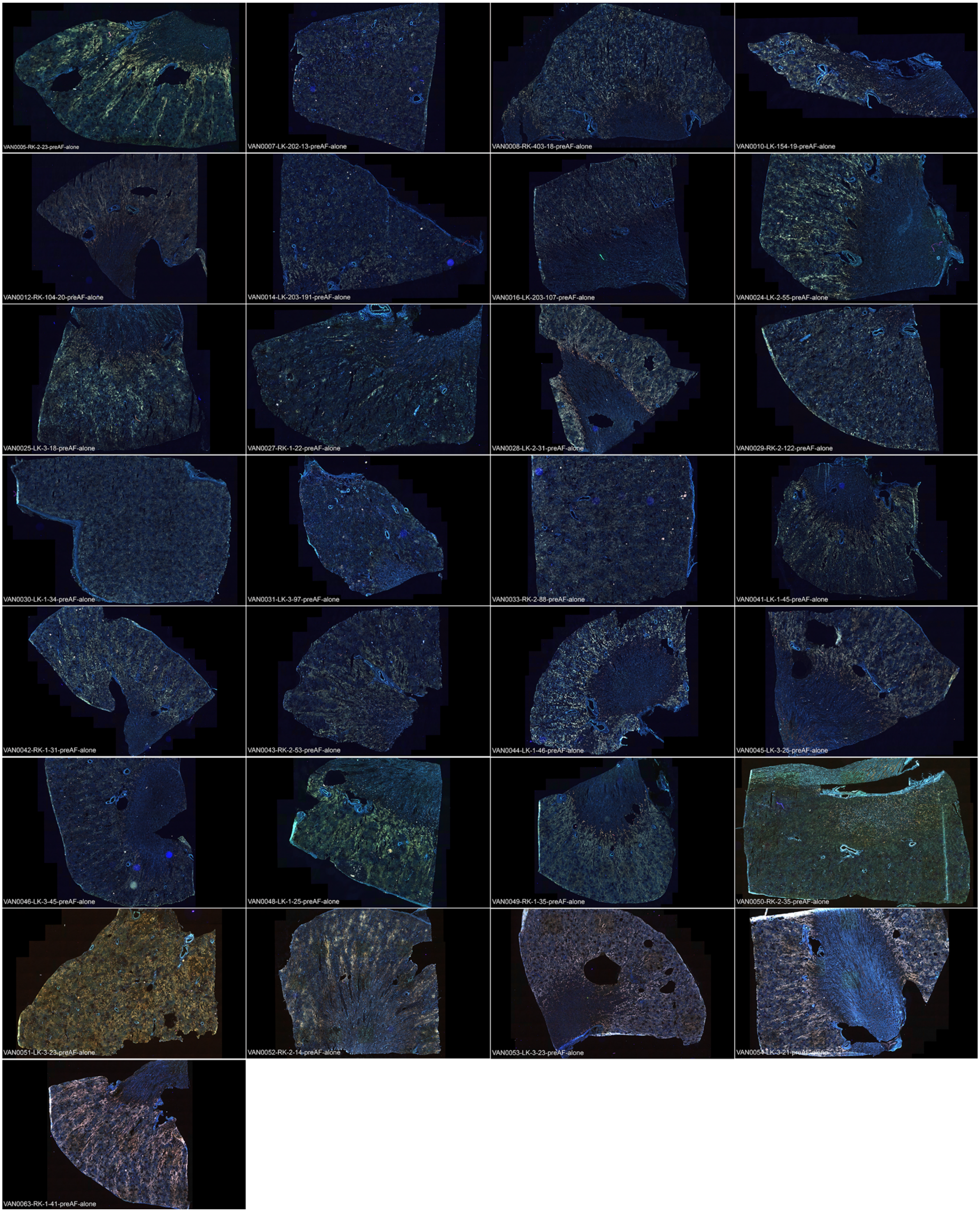

**Figure S2.** Autofluorescence images (pre-IMS-acquisition) of all donor samples.

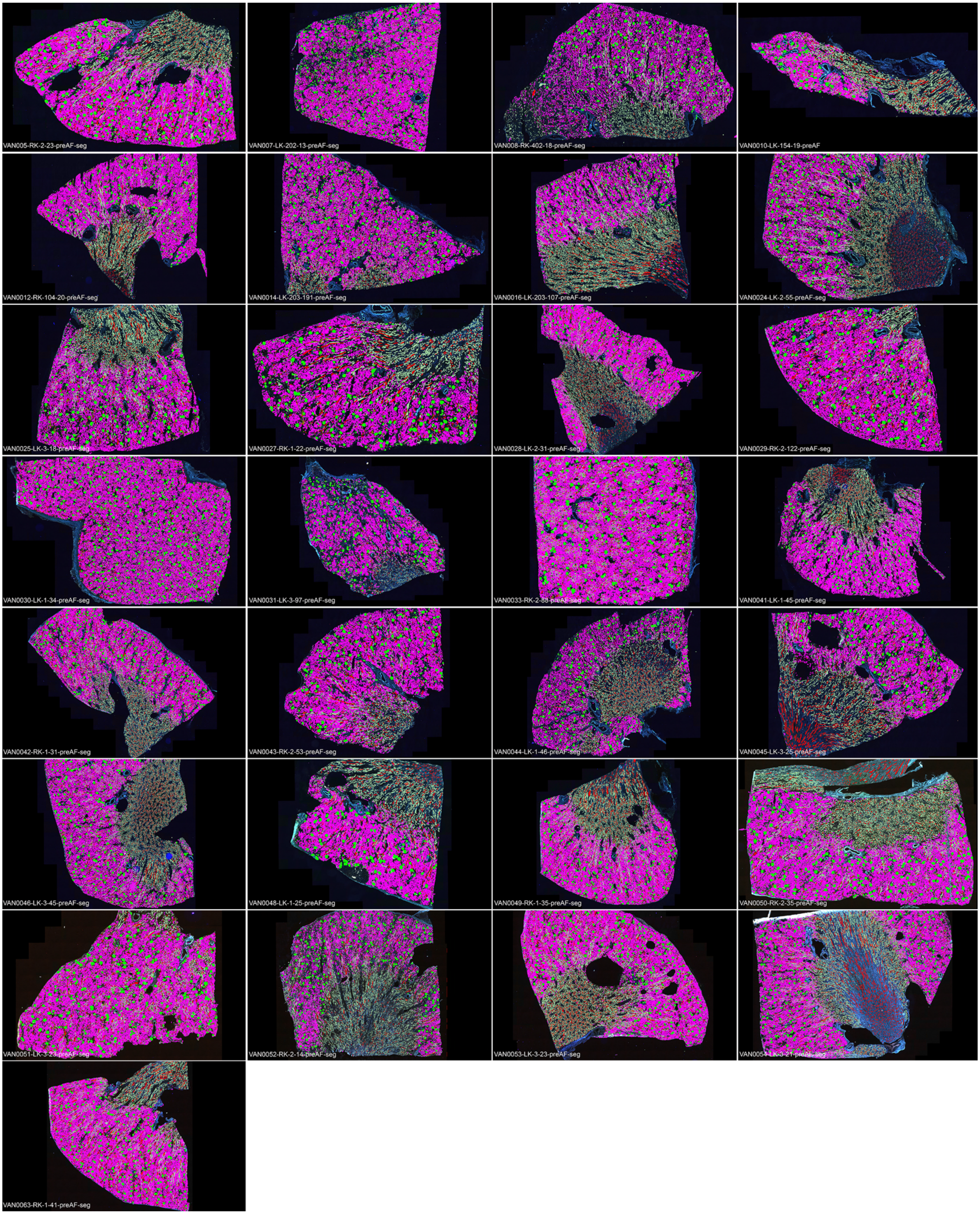

**Figure S3.** Automated FTU segmentation of autofluorescence whole slide human kidney images. FTU segments include glomeruli (green), proximal tubules (magenta), thick ascending limb (light green), distal tubules (brown), and collecting ducts (red).

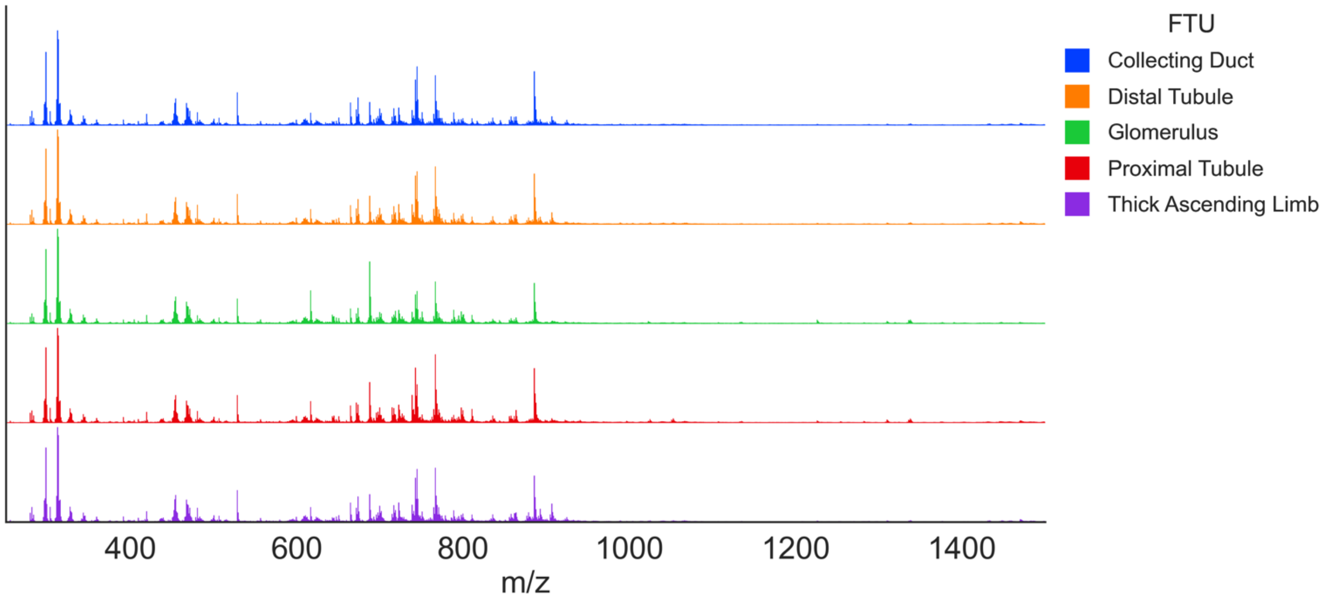

**Figure S4.** The <u>negative ion mode</u> full mass range average spectrum for all pixels associated with each functional tissue unit for all donor samples. The intensity is scaled to the base peak for each spectrum.

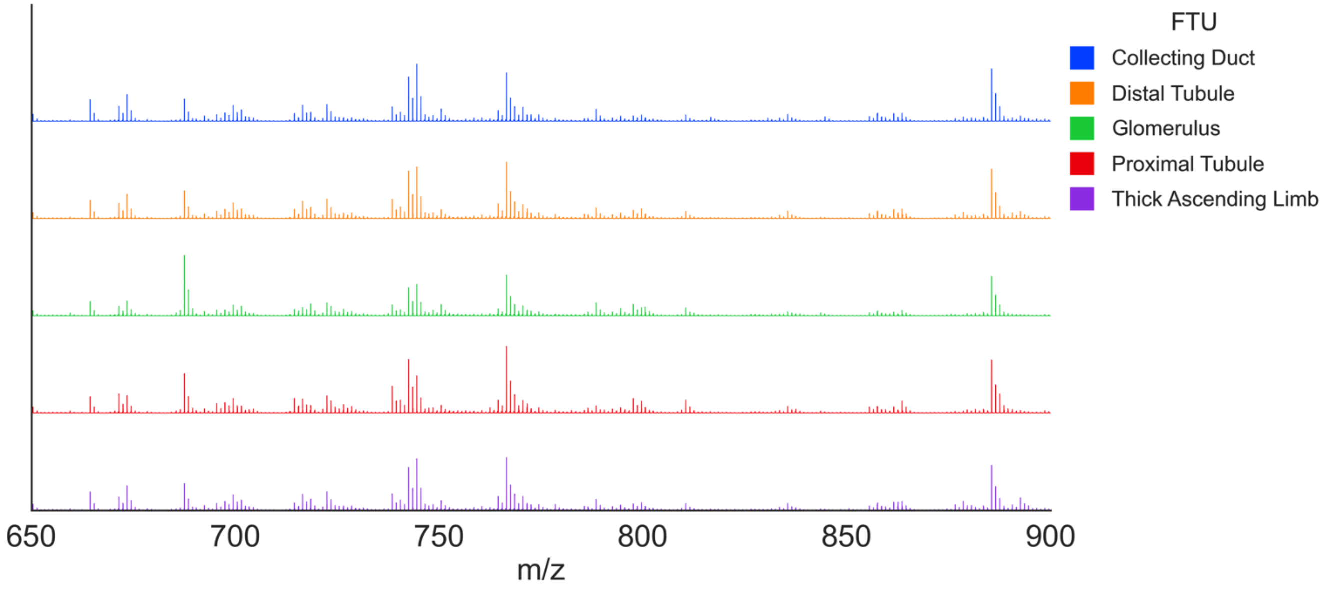

**Figure S5.** The <u>negative ion mode</u> average spectrum from *m/z* 650-900 for all pixels associated with each functional tissue unit for all donor samples. The intensity is scaled to the base peak for each spectrum.

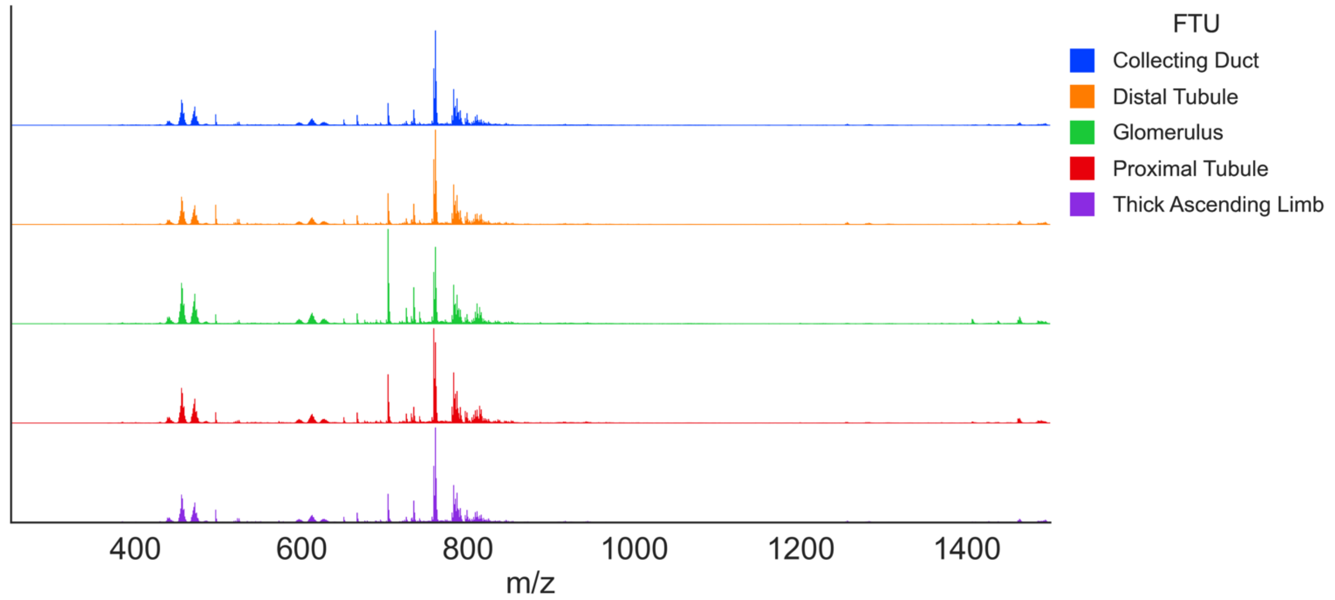

**Figure S6**. The positive ion mode full mass range average spectrum for all pixels associated with each functional tissue unit for all donor samples. The intensity is scaled to the base peak for each spectrum.

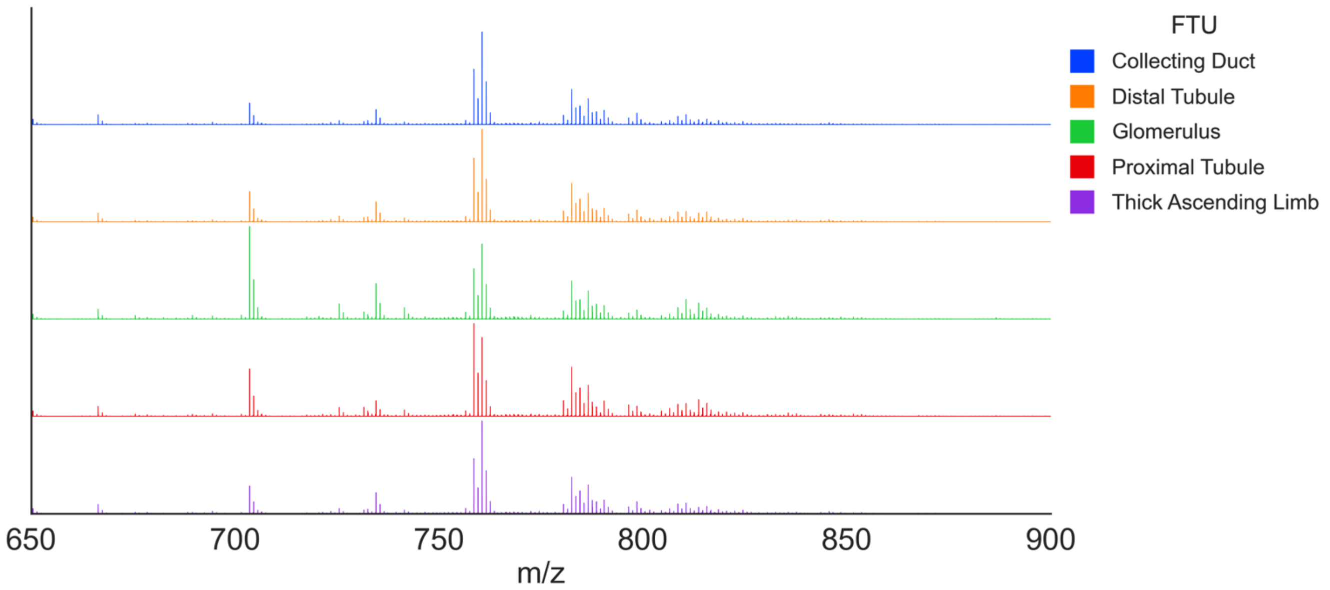

**Figure S7**. The positive ion mode average spectrum from *m/z* 650-900 for all pixels associated with each functional tissue unit for all donor samples. The intensity is scaled to the base peak for each spectrum.

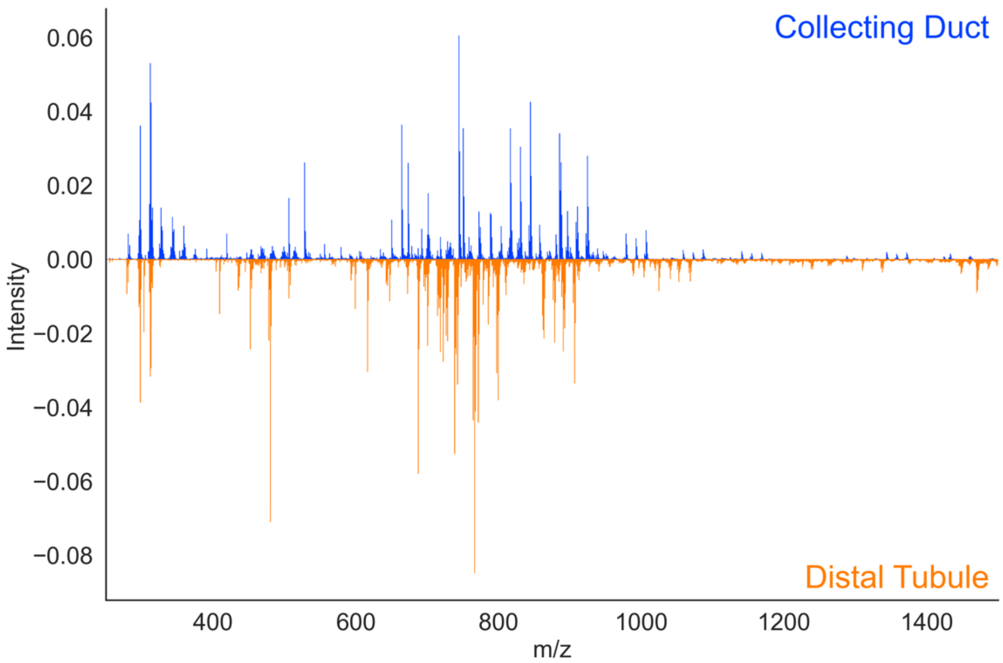

**Figure S8**. The intensity difference spectrum between the <u>negative ion mode</u> average spectra for all pixels associated with collecting ducts (top, blue) and distal tubules (bottom, orange) from all donor samples.

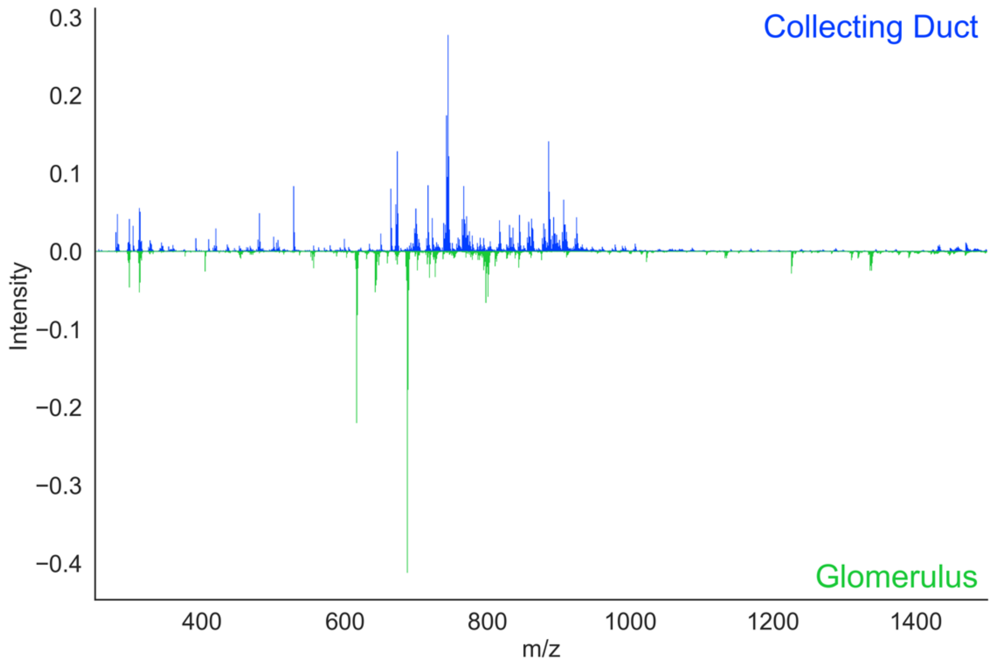

**Figure S9.** The intensity difference spectrum between the <u>negative ion mode</u> average spectra for all pixels associated with collecting ducts (top, blue) and glomeruli (bottom, green) from all donor samples.

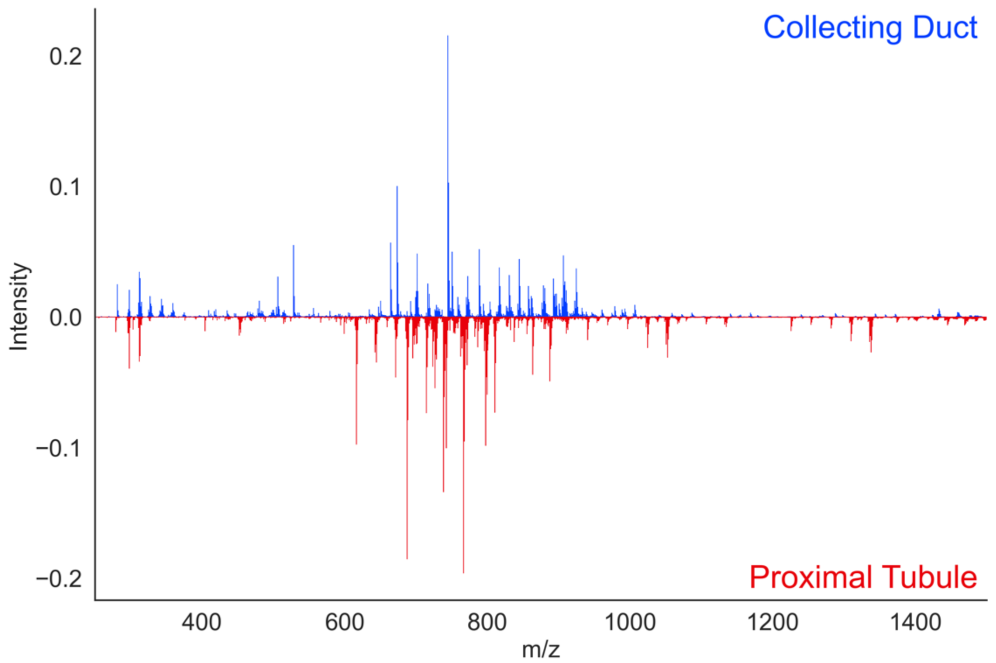

**Figure S10.** The intensity difference spectrum between the <u>negative ion mode</u> average spectra for all pixels associated with collecting ducts (top, blue) and proximal tubules (bottom, red) from all donor samples.

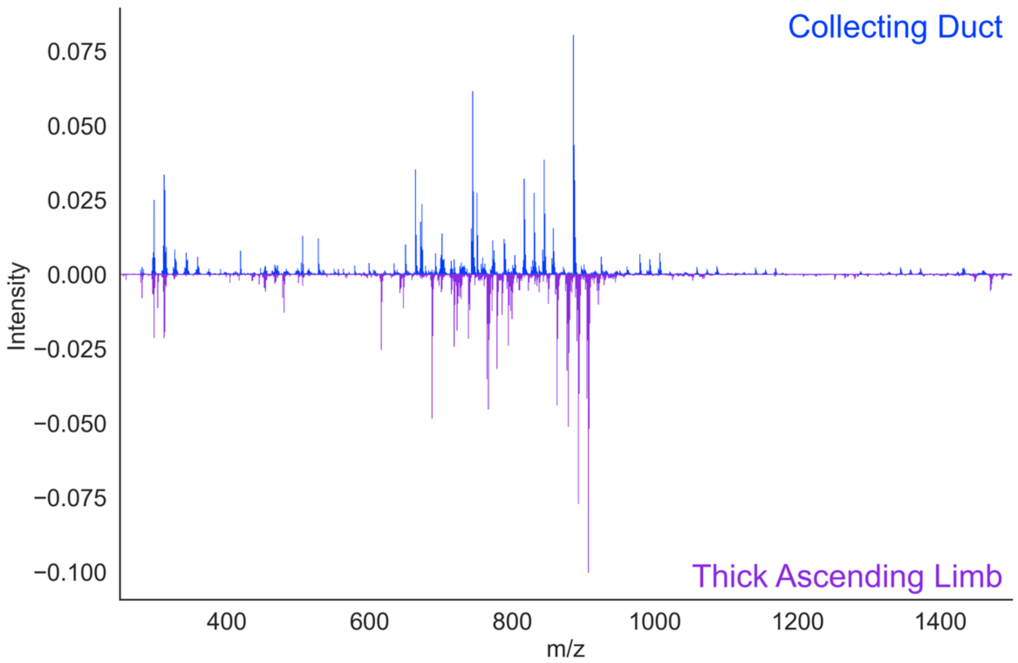

**Figure S11.** The intensity difference spectrum between the <u>negative ion mode</u> average spectra for all pixels associated with collecting ducts (top, blue) and thick ascending limb (bottom, purple) from all donor samples.

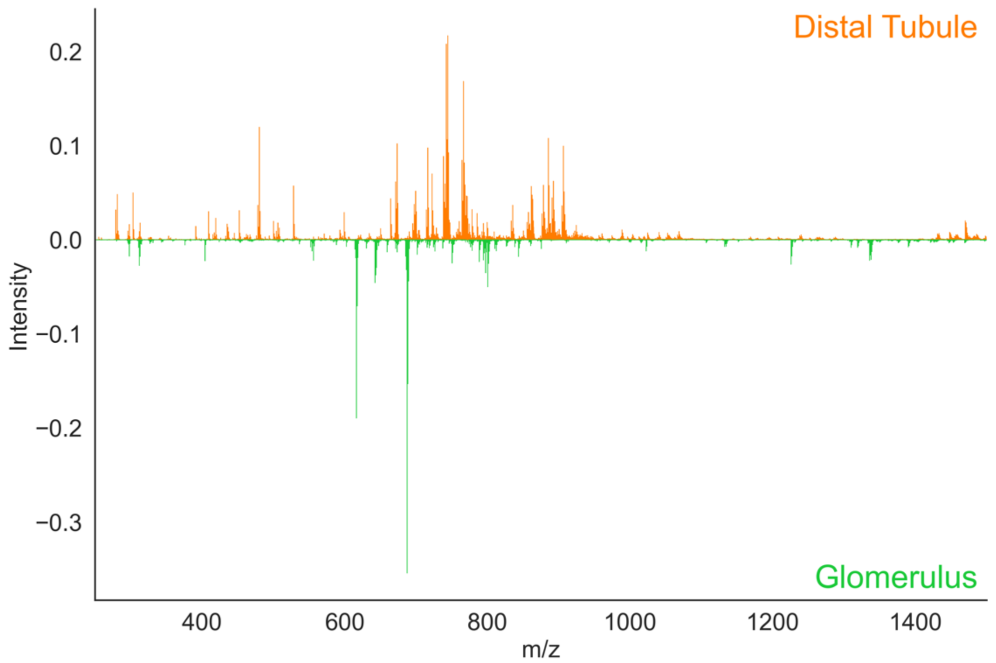

**Figure S12.** The intensity difference spectrum between the negative ion mode average spectra for all pixels associated with distal tubules (top, orange) and glomeruli (bottom, green) from all donor samples.

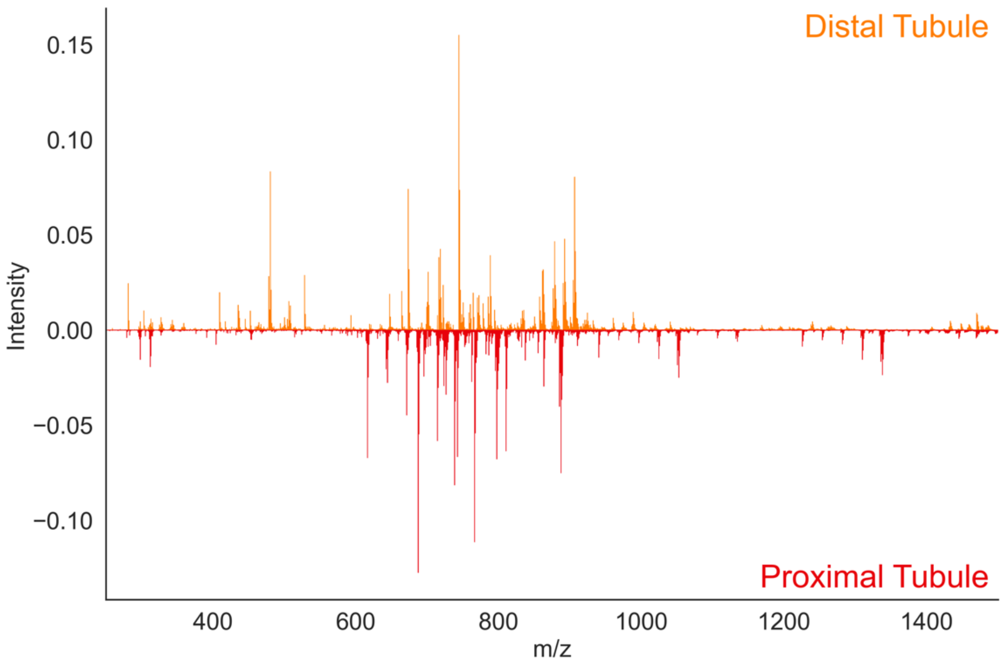

**Figure S13.** The intensity difference spectrum between the negative ion mode average spectra for all pixels associated with distal tubules (top, orange) and proximal tubules (bottom, red) from all donor samples.

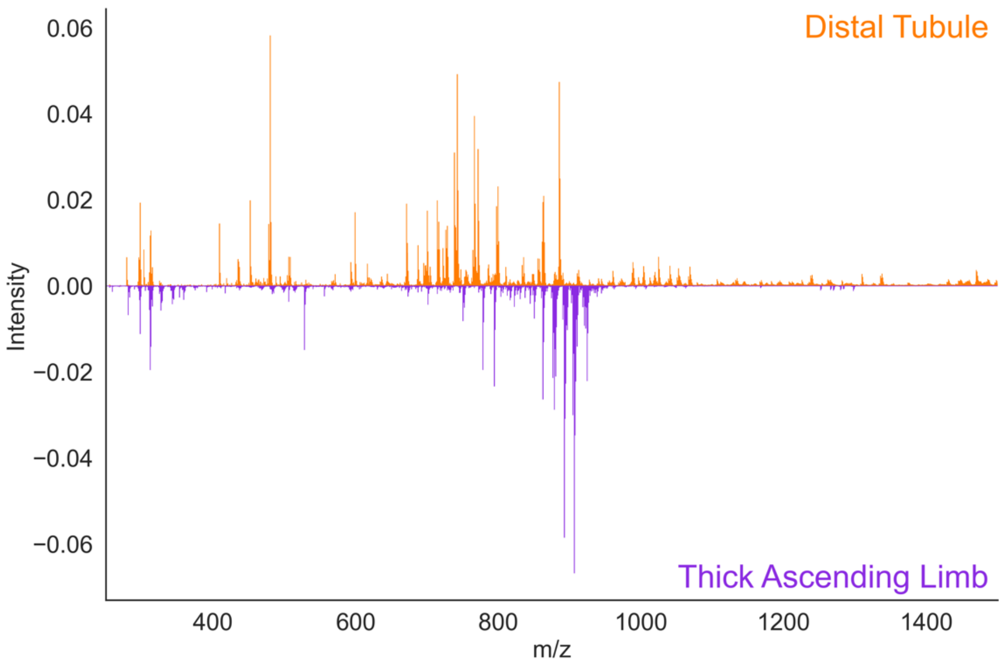

**Figure S14.** The intensity difference spectrum between the negative ion mode average spectra for all pixels associated with distal tubules (top, orange) and thick ascending limb (bottom, purple) from all donor samples.

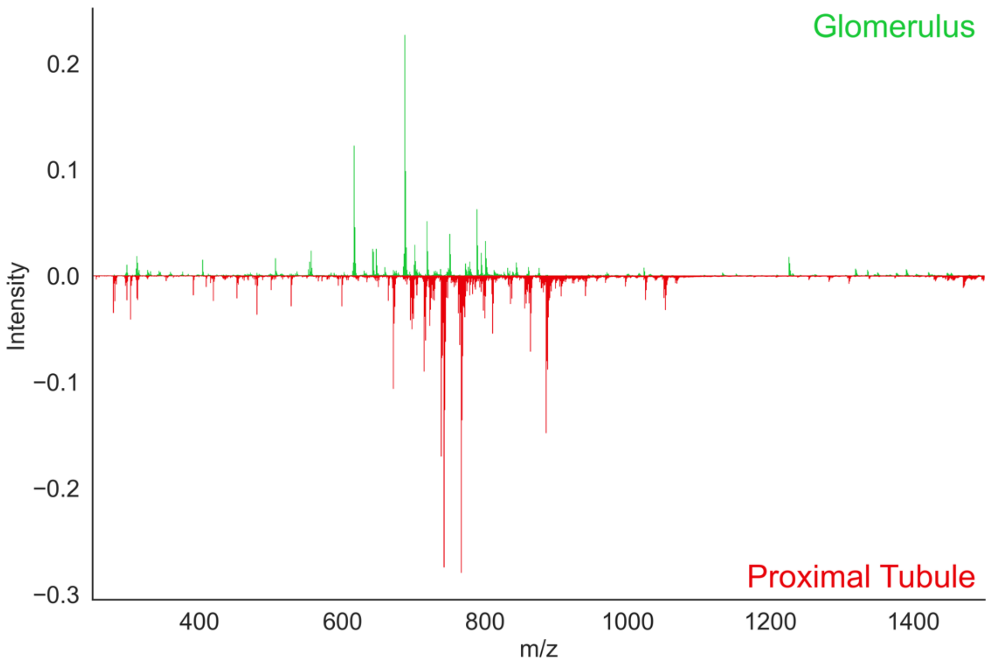

**Figure S15.** The intensity difference spectrum between the negative ion mode average spectra for all pixels associated with glomeruli (top, green) and proximal tubules (bottom, red) from all donor samples.

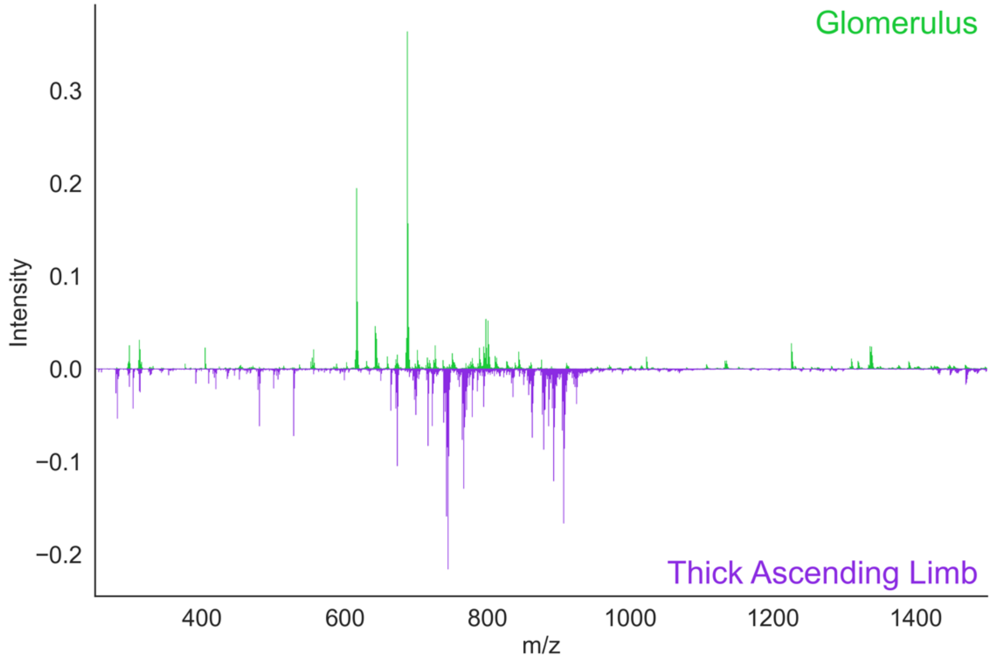

**Figure S16.** The intensity difference spectrum between the negative ion mode average spectra for all pixels associated with glomeruli (top, green) and thick ascending limb (bottom, purple) from all donor samples.

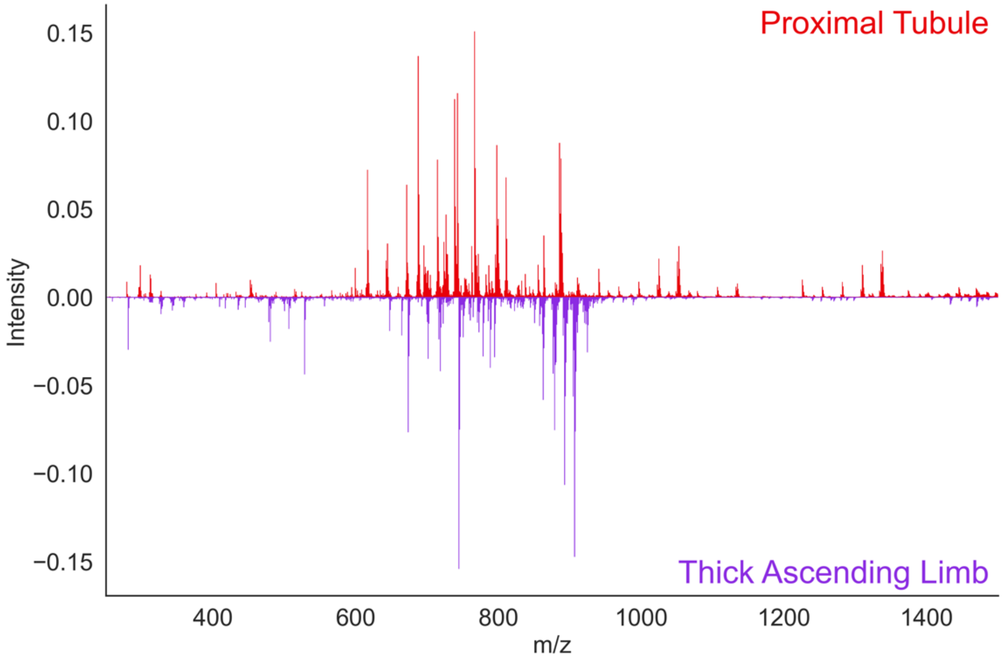

**Figure S17.** The intensity difference spectrum between the negative ion mode average spectra for all pixels associated with proximal tubules (top, red) and thick ascending limb (bottom, purple) from all donor samples.

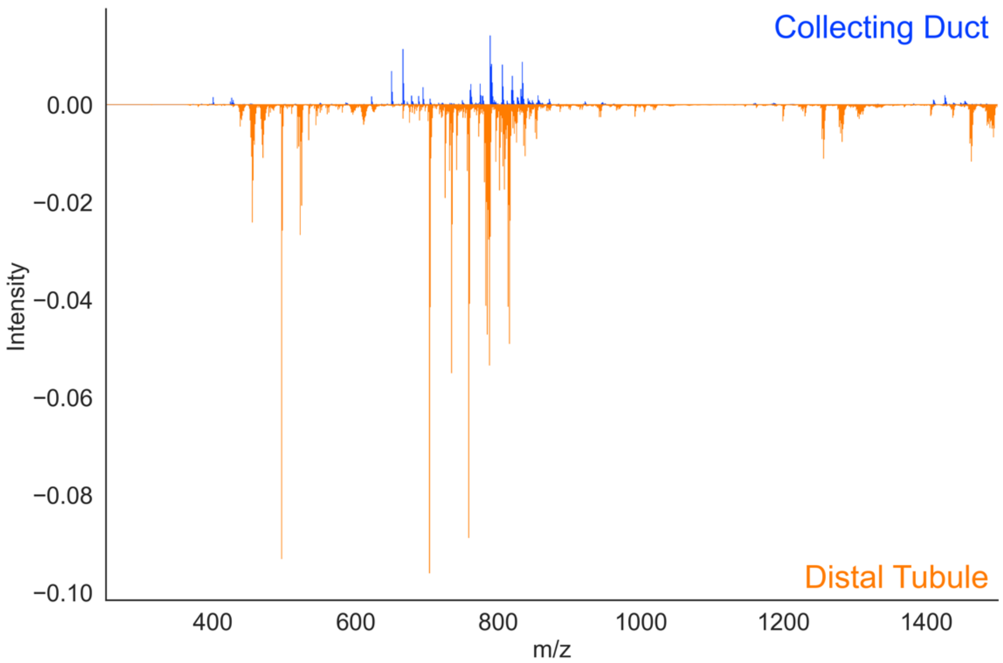

**Figure S18.** The intensity difference spectrum between the positive ion mode average spectra for all pixels associated with collecting ducts (top, blue) and distal tubules (bottom, orange) from all donor samples.

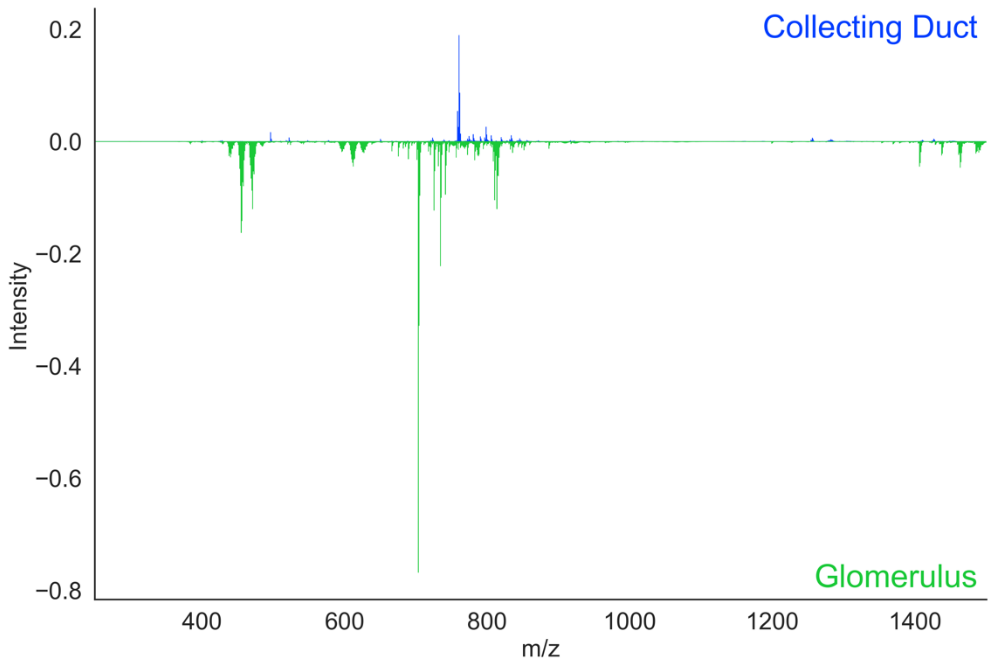

**Figure S19.** The intensity difference spectrum between the positive ion mode average spectra for all pixels associated with collecting ducts (top, blue) and glomeruli (bottom, green) from all donor samples.

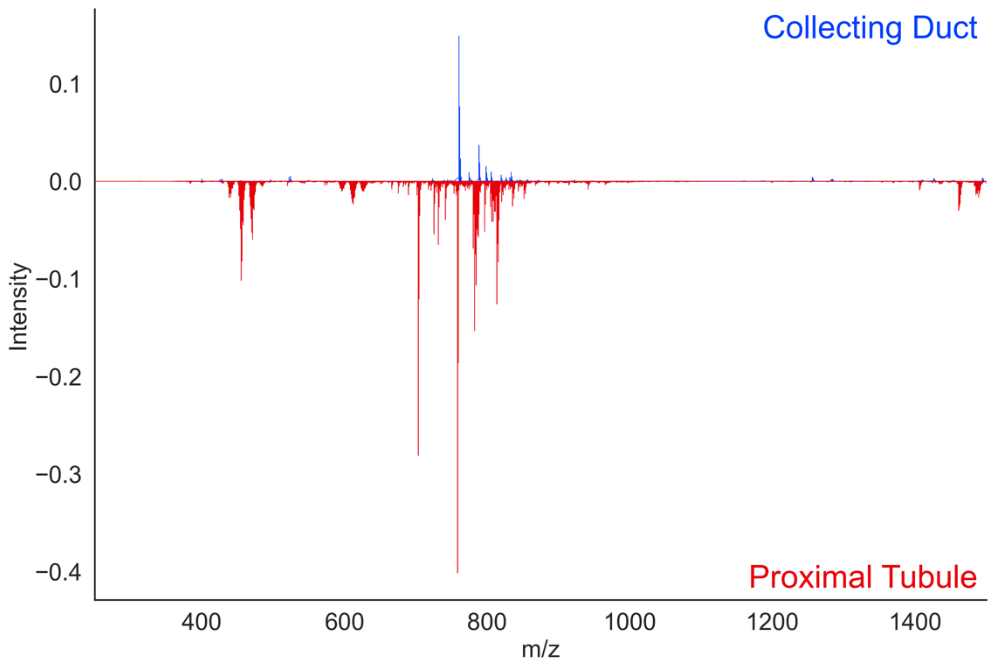

**Figure S20.** The intensity difference spectrum between the positive ion mode average spectra for all pixels associated with collecting ducts (top, blue) and proximal tubules (bottom, red) from all donor samples.

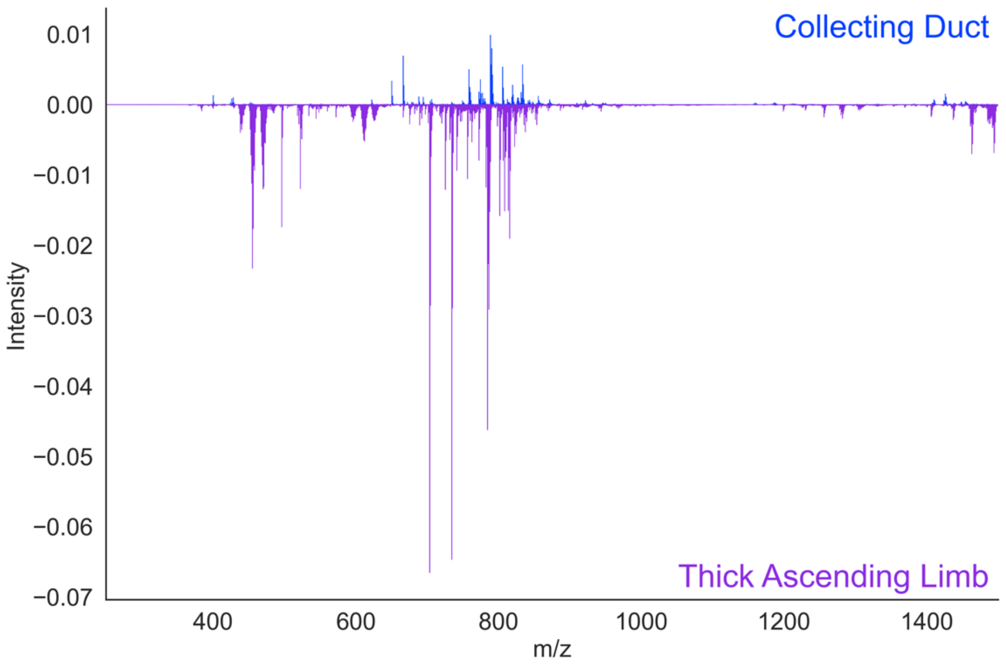

**Figure S21.** The intensity difference spectrum between the positive ion mode average spectra for all pixels associated with collecting ducts (top, blue) and thick ascending limb (bottom, purple) from all donor samples.

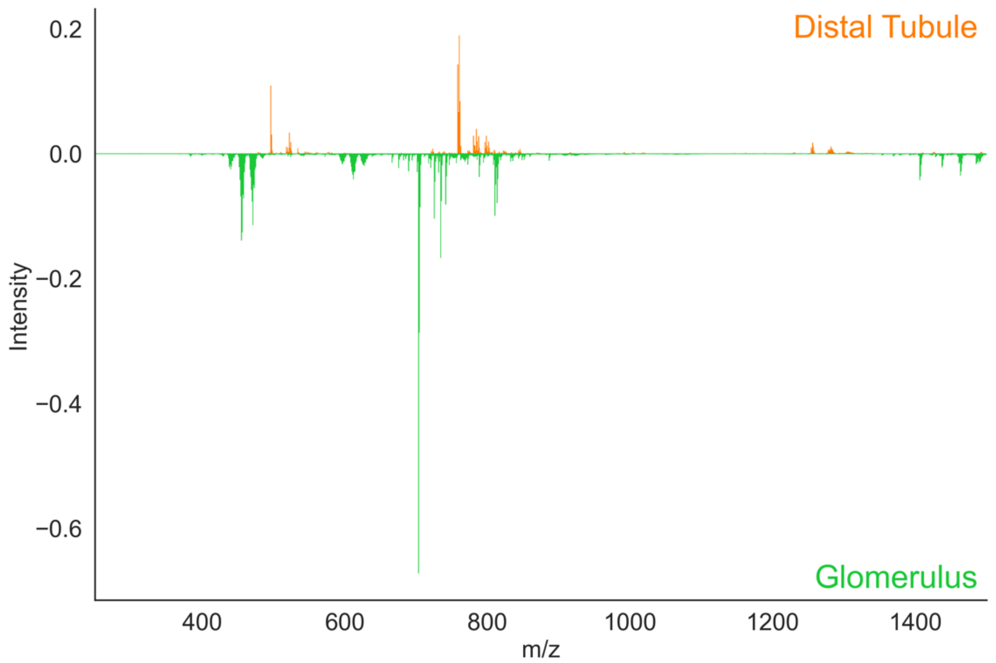

**Figure S22.** The intensity difference spectrum between the positive ion mode average spectra for all pixels associated with distal tubules (top, orange) and glomeruli (bottom, green) from all donor samples.

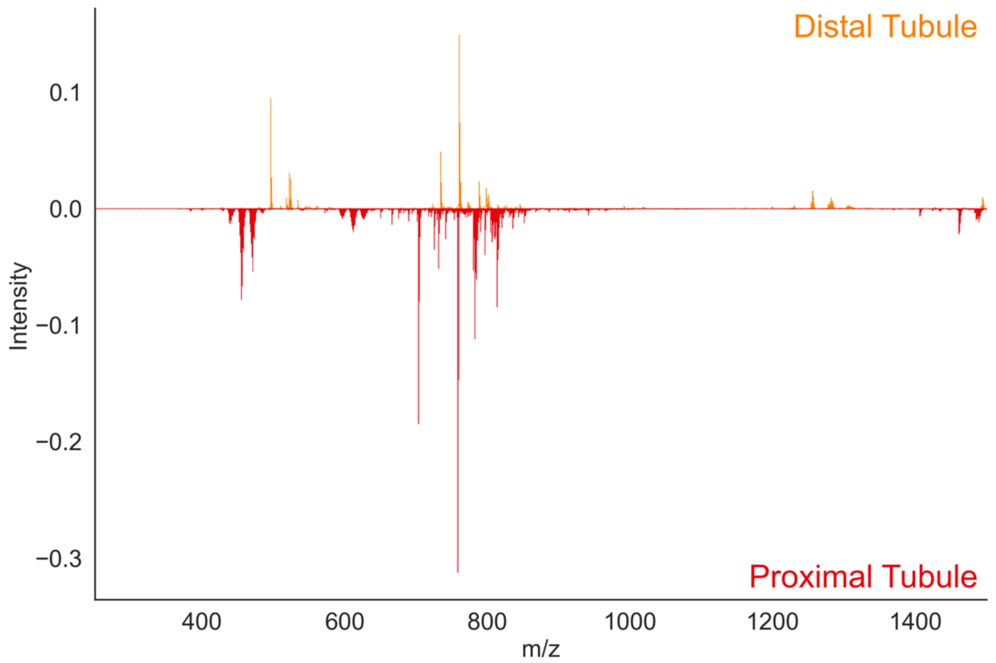

**Figure S23.** The intensity difference spectrum between the positive ion mode average spectra for all pixels associated with distal tubules (top, orange) and proximal tubules (bottom, red) from all donor samples.

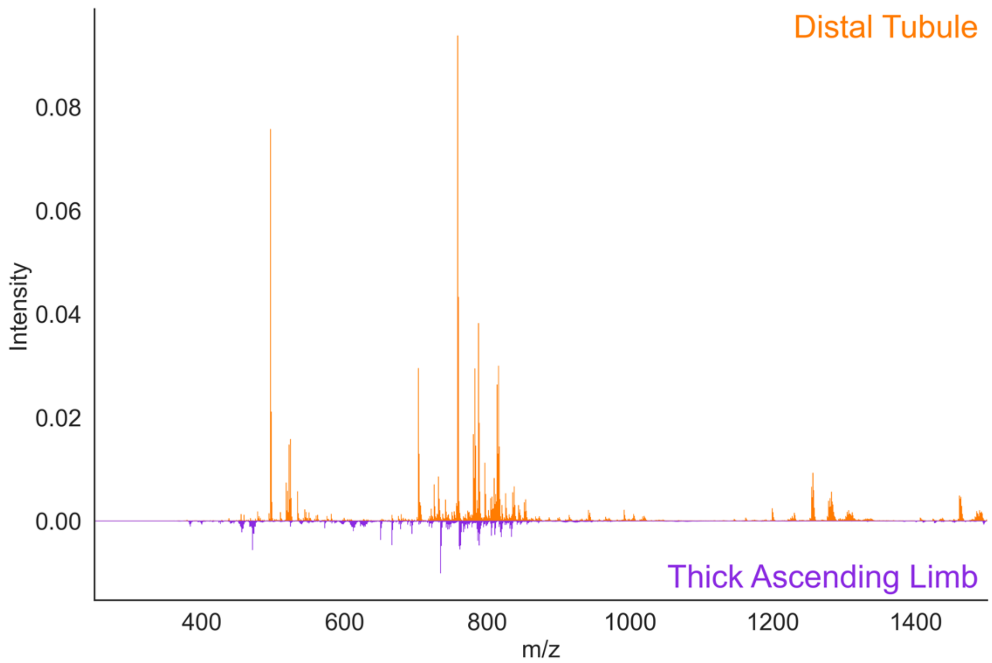

**Figure S24.** The intensity difference spectrum between the positive ion mode average spectra for all pixels associated with distal tubules (top, orange) and thick ascending limb (bottom, purple) from all donor samples.

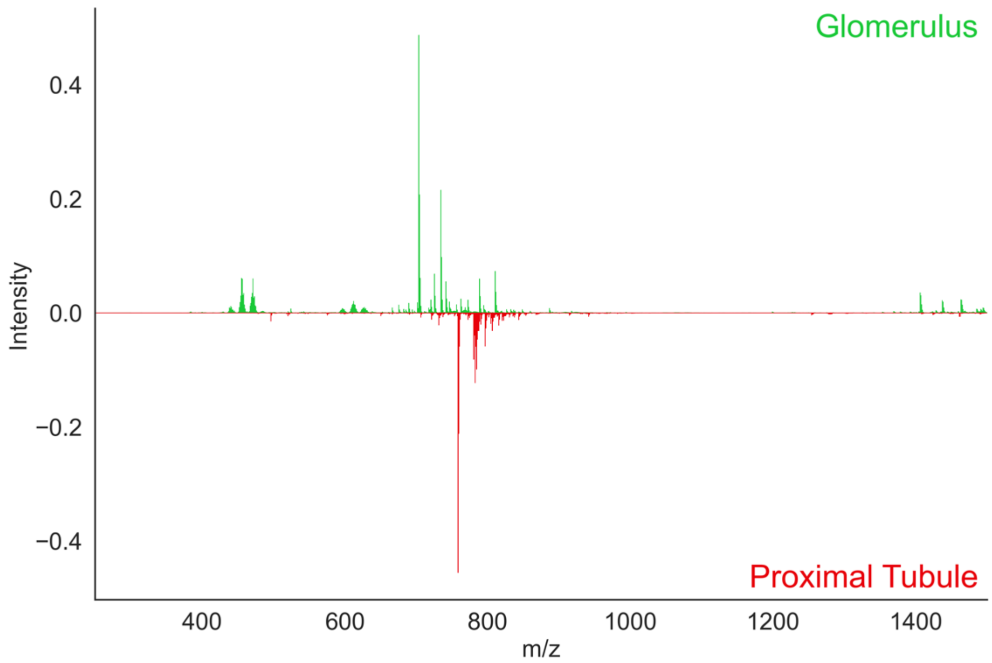

**Figure S25.** The intensity difference spectrum between the positive ion mode average spectra for all pixels associated with glomeruli (top, green) and proximal tubules (bottom, red) from all donor samples.

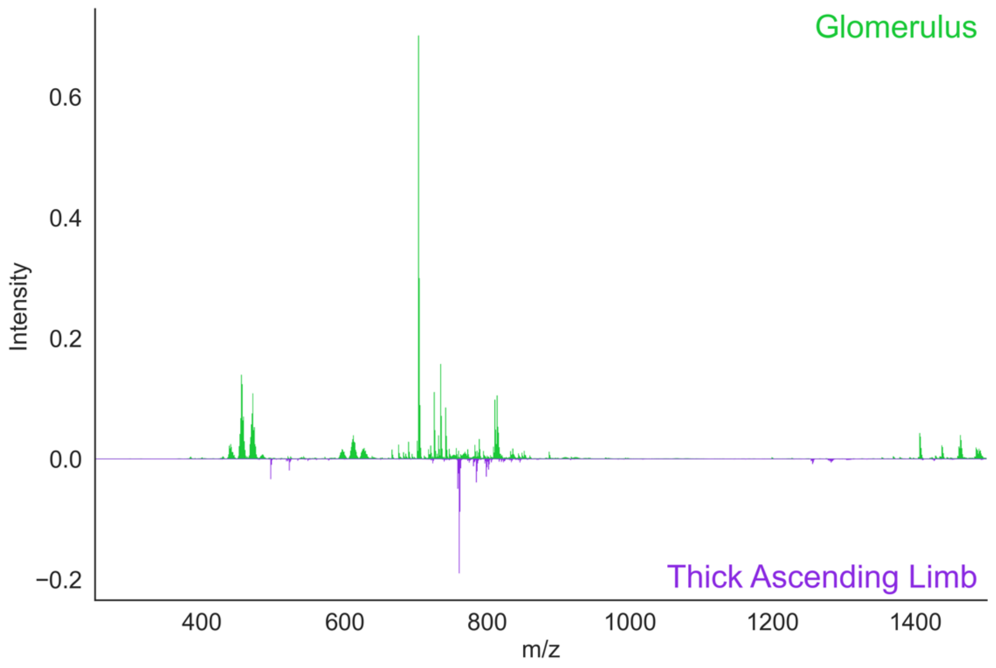

**Figure S26.** The intensity difference spectrum between the <u>positive ion mode</u> average spectra for all pixels associated with glomeruli (top, green) and thick ascending limb (bottom, purple) from all donor samples.

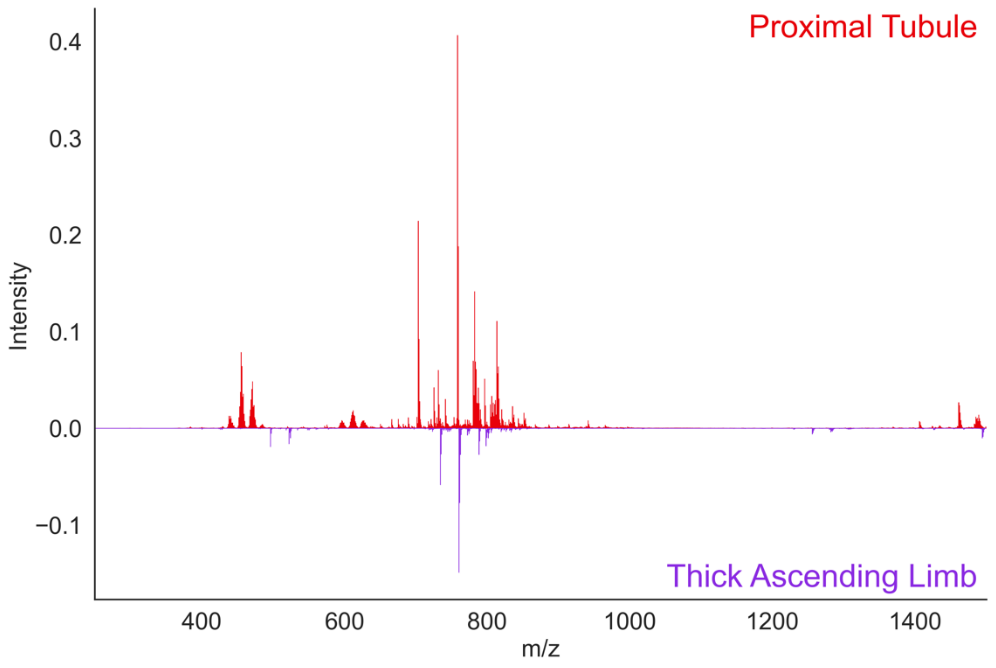

**Figure S27.** The intensity difference spectrum between the <u>positive ion mode</u> average spectra for all pixels associated with proximal tubules (top, red) and thick ascending limb (bottom, purple) from all donor samples.

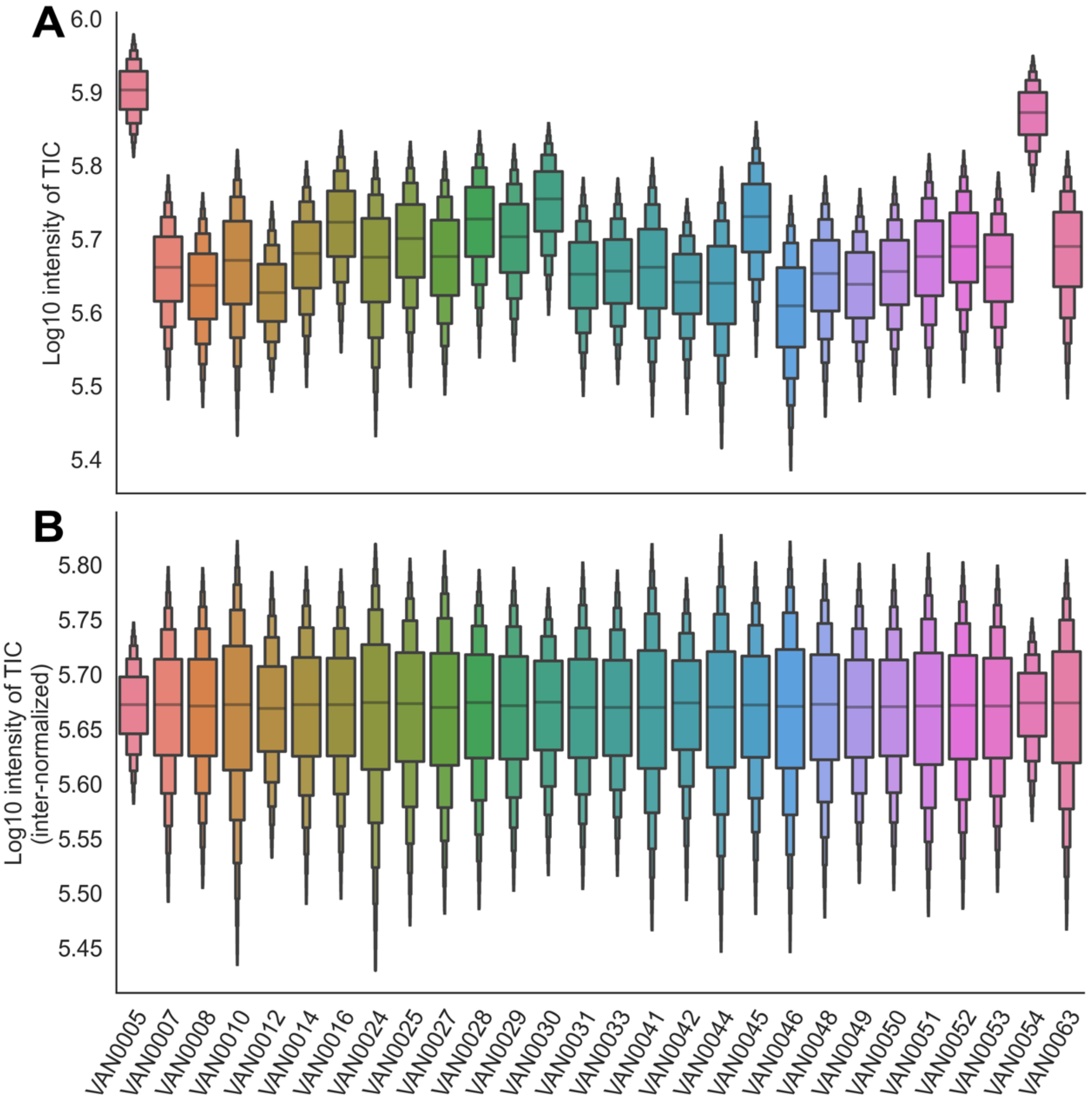

**Figure S28.** The Log10 intensity of the total ion current for all pixels collected from each donor sample in <u>negative ion mode</u>. (A) Data were normalized within each dataset (intra-normalized) and (B) across all datasets (inter-normalized). Inter-normalized data were used for all subsequent analyses. The boxen plots provide a more detailed visualization of the distribution of the log-transformed intensities, offering insights into the median, interquartile range, and the overall spread of the data, thereby highlighting potential outliers or anomalies within each sample set.

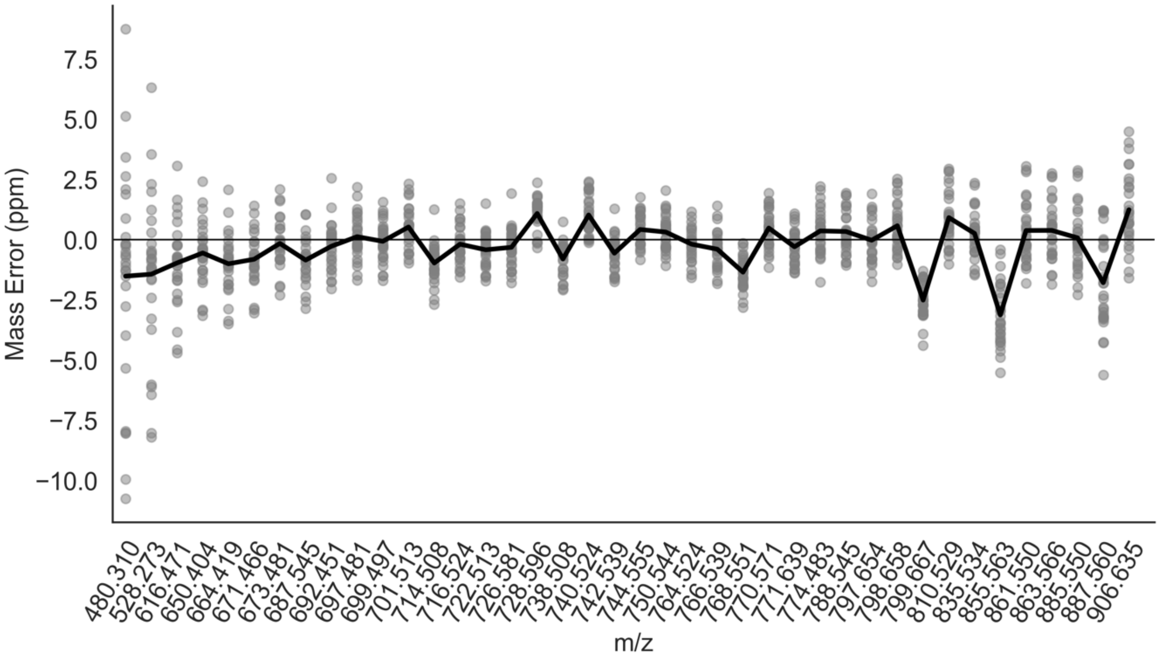

**Figure S29.** The mass error in parts per million (ppm) for selected <u>negative ion mode</u> lipids. The black line represents the mean mass error from all pixels collected from all donor samples, and the grey dots represent the spread of the data for each *m/z*.

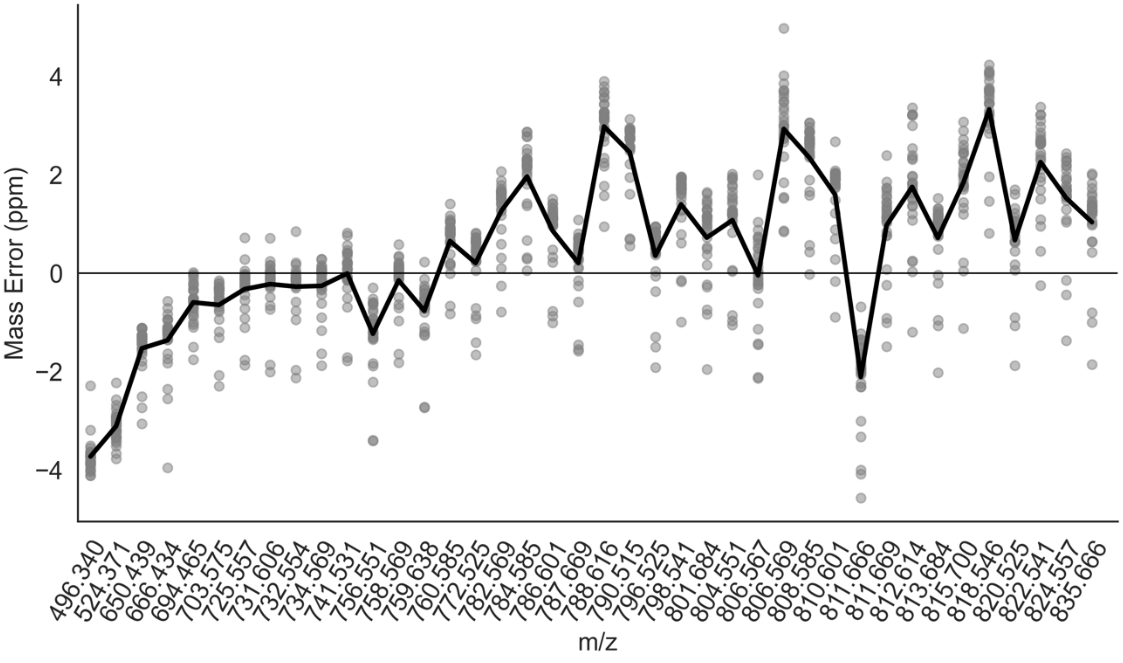

**Figure S30.** The mass error in parts per million (ppm) for selected <u>positive ion mode</u> lipids. The black line represents the mean mass error from all pixels collected from all donor samples, and the grey dots represent the spread of the data for each *m/z*.

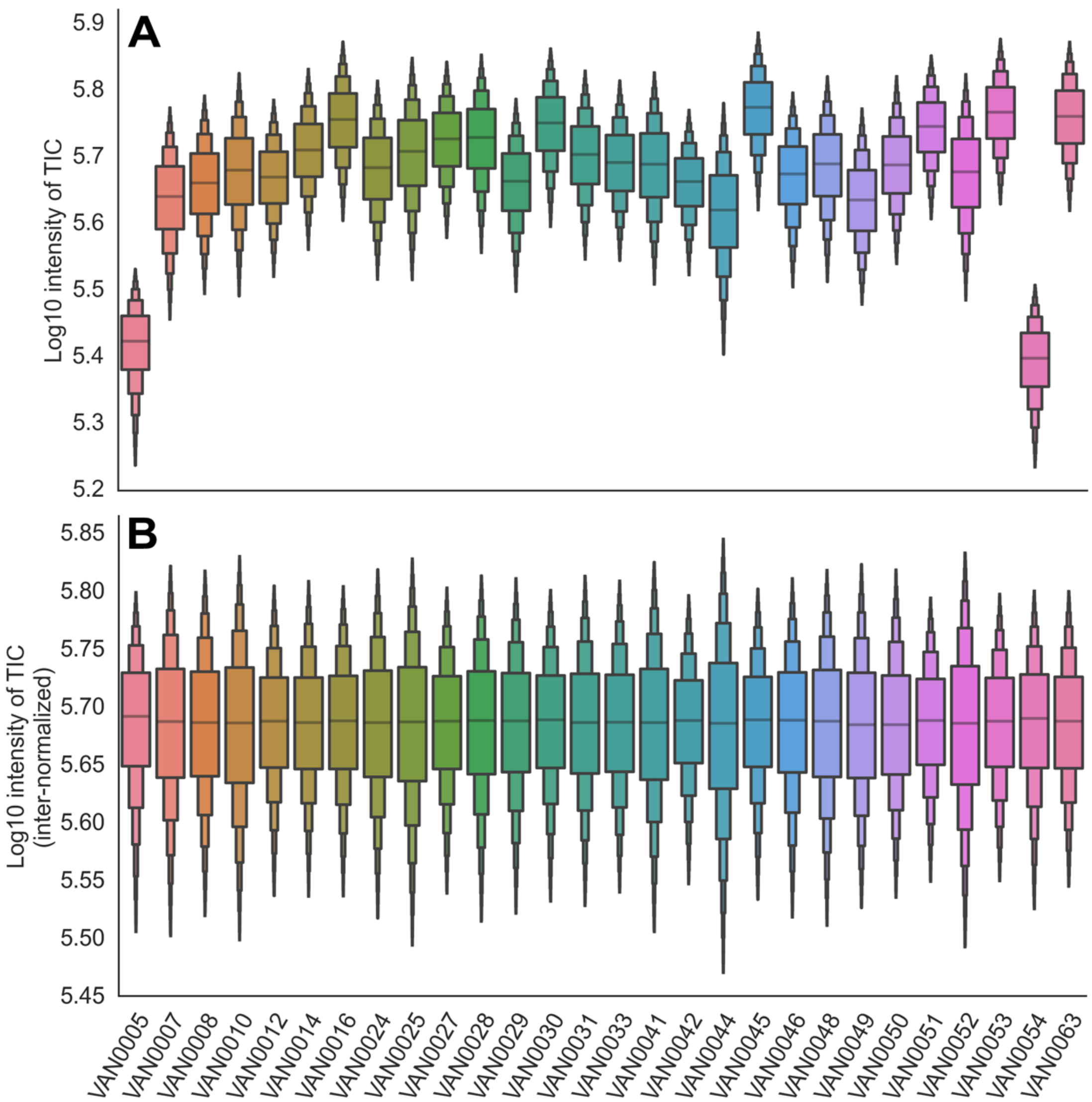

**Figure S31.** The Log10 intensity of the total ion current for all pixels collected from each donor sample in positive ion mode. (A) Data were normalized within each dataset (intra-normalized) and (B) across all datasets (inter-normalized). Inter-normalized data were used for all subsequent analyses. The boxen plots provide a more detailed visualization of the distribution of the log-transformed intensities, offering insights into the median, interquartile range, and the overall spread of the data, thereby highlighting potential outliers or anomalies within each sample set.

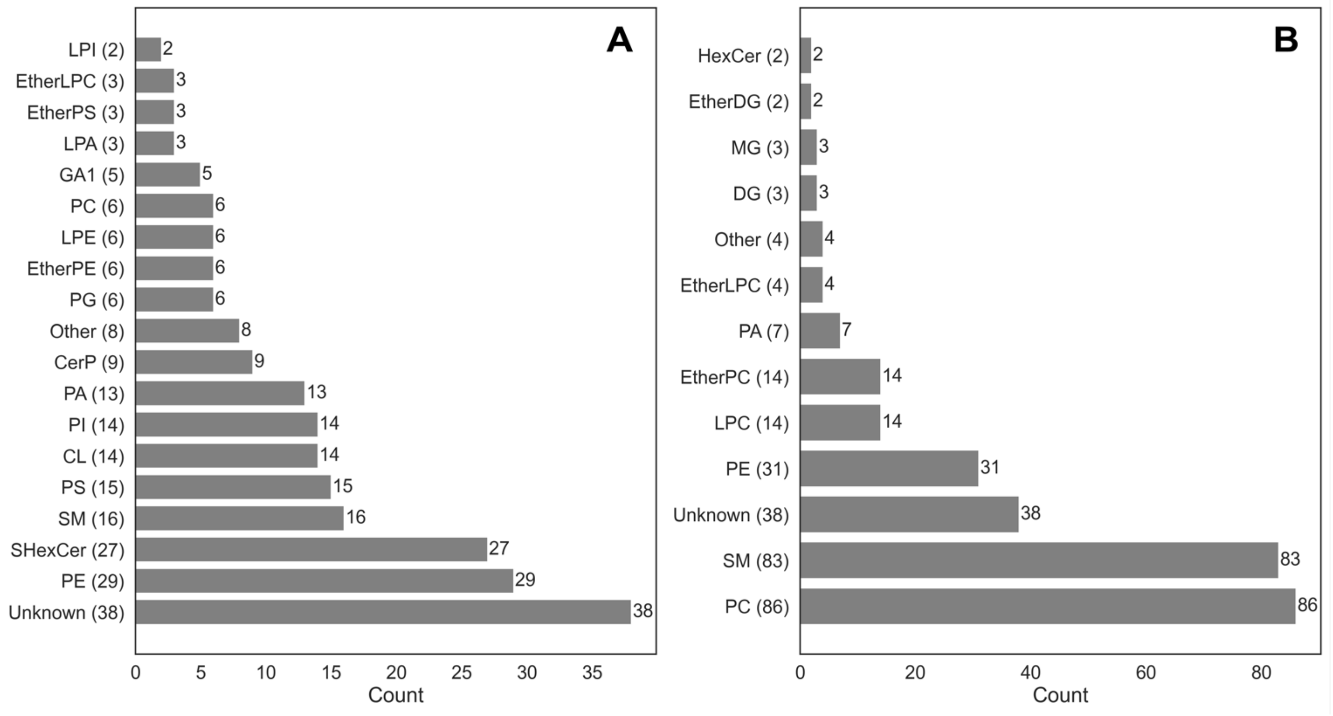

**Figure S32.** The total number of lipids identified from each class that were annotated in the IMS data for (A) negative ion mode and (B) positive ion mode.

**Figure S33.** Pixel-wise UMAP embedding of <u>negative ion mode</u> data, illustrating the effect of batch normalization and categorization based on technical and biological factors. (A) Shows the embedding of IMS data after inter-dataset normalization, colored by donor ID, where separation is observed according to donor, rather than other biological variations. (B) Displays the embedding post inter-dataset and batch normalization, revealing improved integration of IMS data. (C, D) are analogous to (A) and (B), but are distinguished by sex. (E, F) follow the same normalization process, differentiated by BMI Finally, (G, H) are categorized by FTU, with (H) demonstrating clear grouping by different FTUs, indicating that batch normalization has effectively minimized dataset-to-dataset variation, allowing the UMAP embedding to be primarily driven by FTU-related variation.

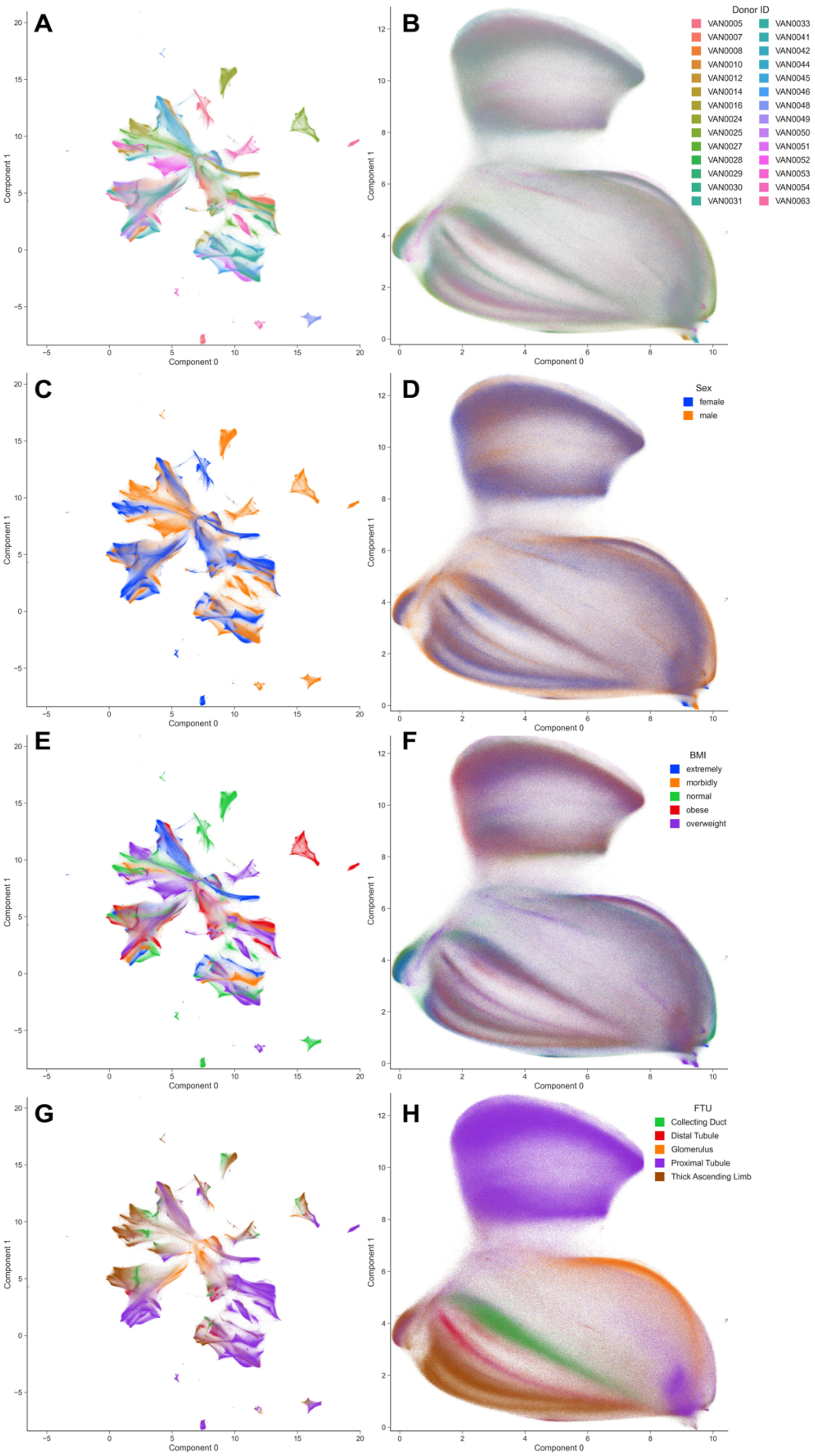

**Figure S34.** FTU-level UMAP negative ion mode where each point represents an aggregated instance of a FTU rather than individual pixels. (A) UMAP embedding before batch normalization, colored by donor ID, illustrating initial data distribution according to donor. (B) UMAP embedding after batch normalization, showing improved data integration across donors. (C) and (D) display UMAP embeddings colored by sex, before and after batch normalization, respectively, highlighting sex-specific patterns. (E) and (F) depict UMAP embeddings colored by BMI categories, before and after batch normalization, visualizing the metabolic variation with respect to body composition. Finally, (G) and (H) present UMAP embeddings colored by FTU, before and after batch normalization, with (H) clearly demonstrating the distinct grouping of different FTUs. This indicates that the batch normalization process successfully reduced the variation between datasets, allowing the UMAP analysis to reflect the diversity of each FTU.

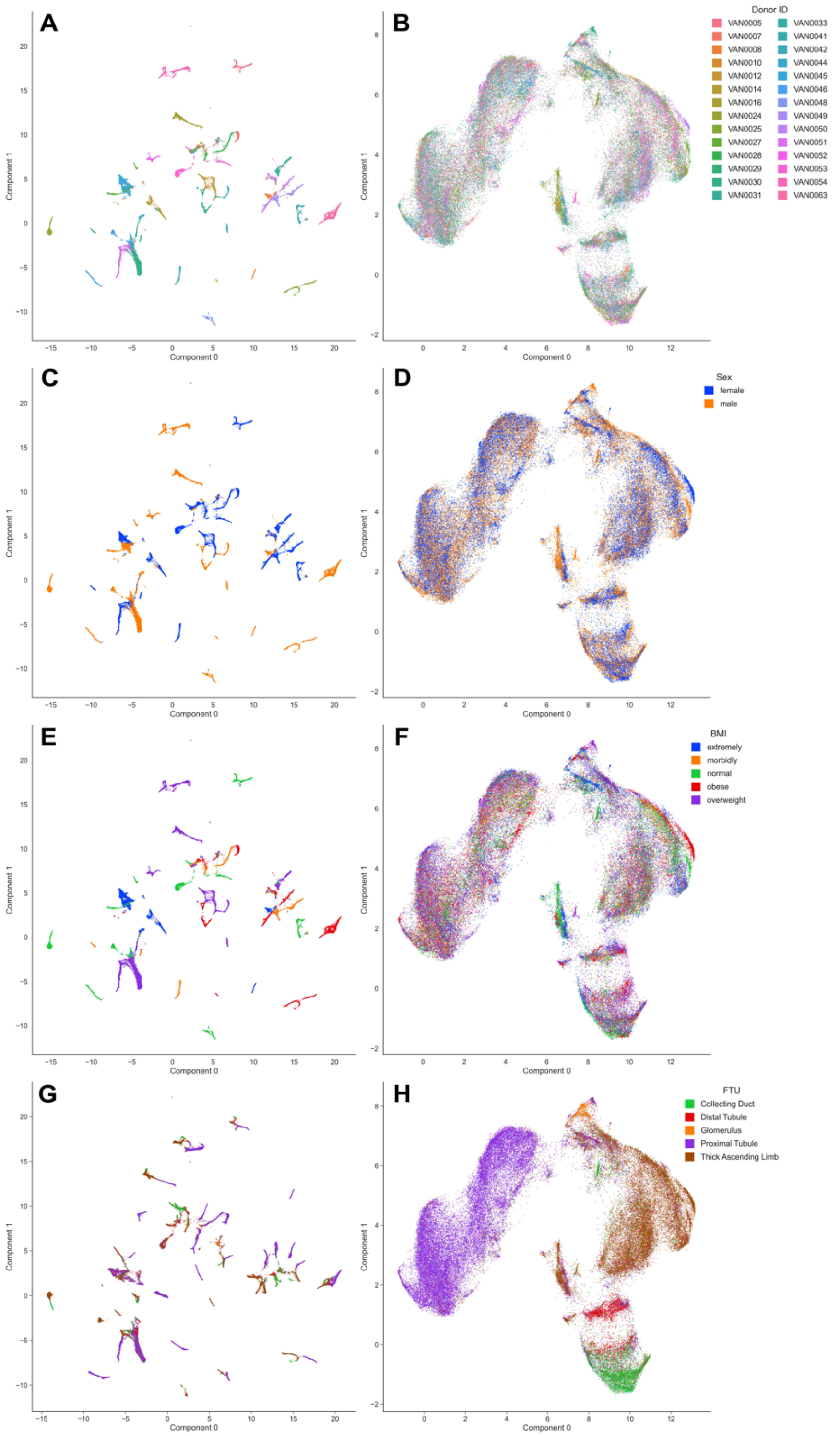

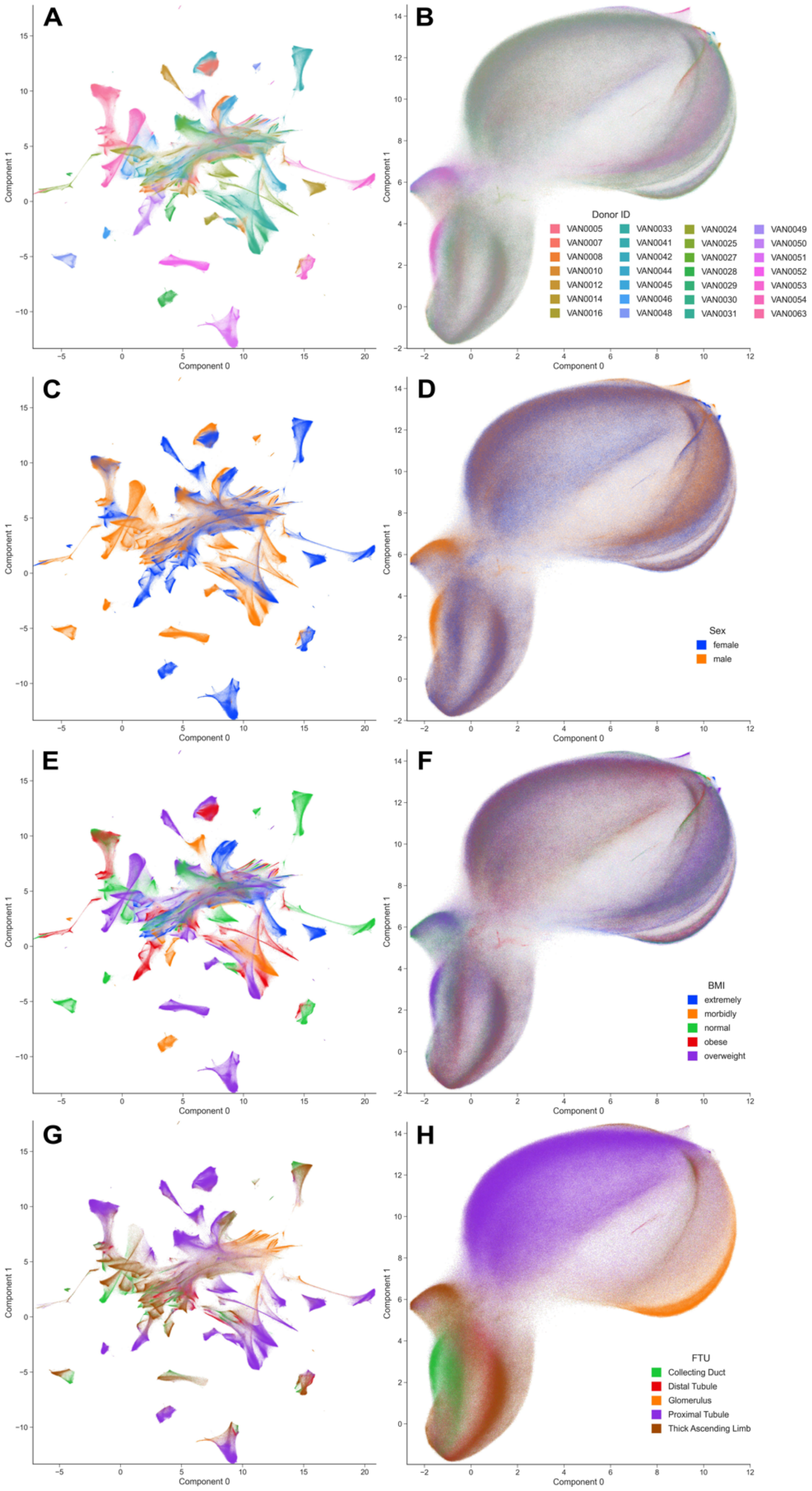

**Figure S35.** Pixel-wise UMAP embedding of positive ion mode data, illustrating the effect of batch normalization and categorization based on technical and biological factors. (A) Shows the embedding of IMS data after inter-dataset normalization, colored by donor ID, where separation is observed according to donor, rather than other biological variations. (B) Displays the embedding post inter-dataset and batch normalization, revealing improved integration of IMS data. (C, D) are analogous to (A) and (B), but are distinguished by sex. (E, F) follow the same normalization process, differentiated by BMI Finally, (G, H) are categorized by FTU, with (H) demonstrating clear grouping by different FTUs, indicating that batch normalization has effectively minimized dataset-to-dataset variation, allowing the UMAP embedding to be primarily driven by FTU-related variation.

**Figure S36.** FTU-level UMAP positive ion mode where each point represents an aggregated instance of a FTU rather than individual pixels. (A) UMAP embedding before batch normalization, colored by donor ID, illustrating initial data distribution according to donor. (B) UMAP embedding after batch normalization, showing improved data integration across donors. (C) and (D) display UMAP embeddings colored by sex, before and after batch normalization, respectively, highlighting sex-specific patterns. (E) and (F) depict UMAP embeddings colored by BMI categories, before and after batch normalization, visualizing the metabolic variation with respect to body composition. Finally, (G) and (H) present UMAP embeddings colored by FTU, before and after batch normalization, with (H) clearly demonstrating the distinct grouping of different FTUs. This indicates that the batch normalization process successfully reduced the variation between datasets, allowing the UMAP analysis to reflect the diversity of each FTU.

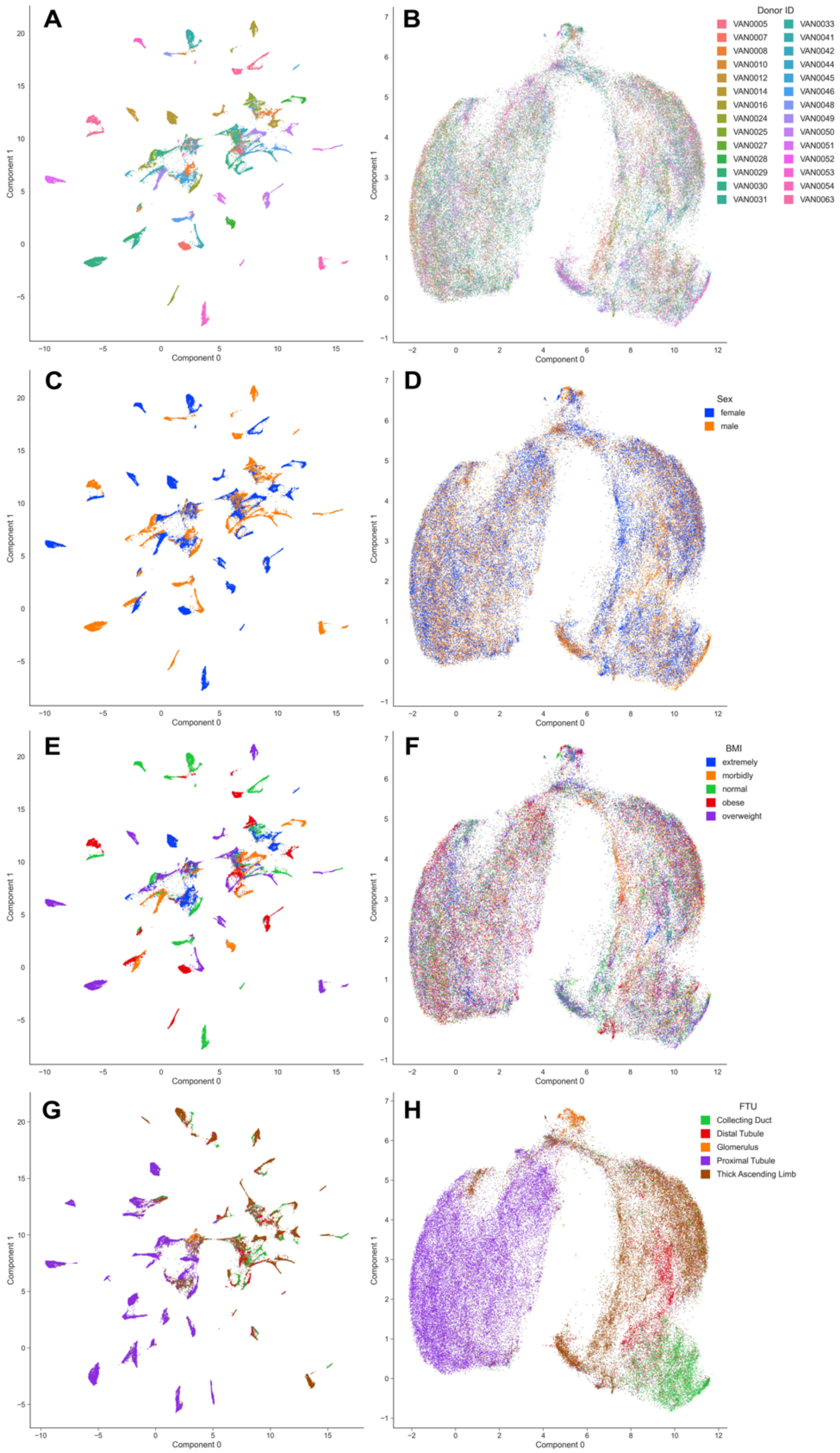

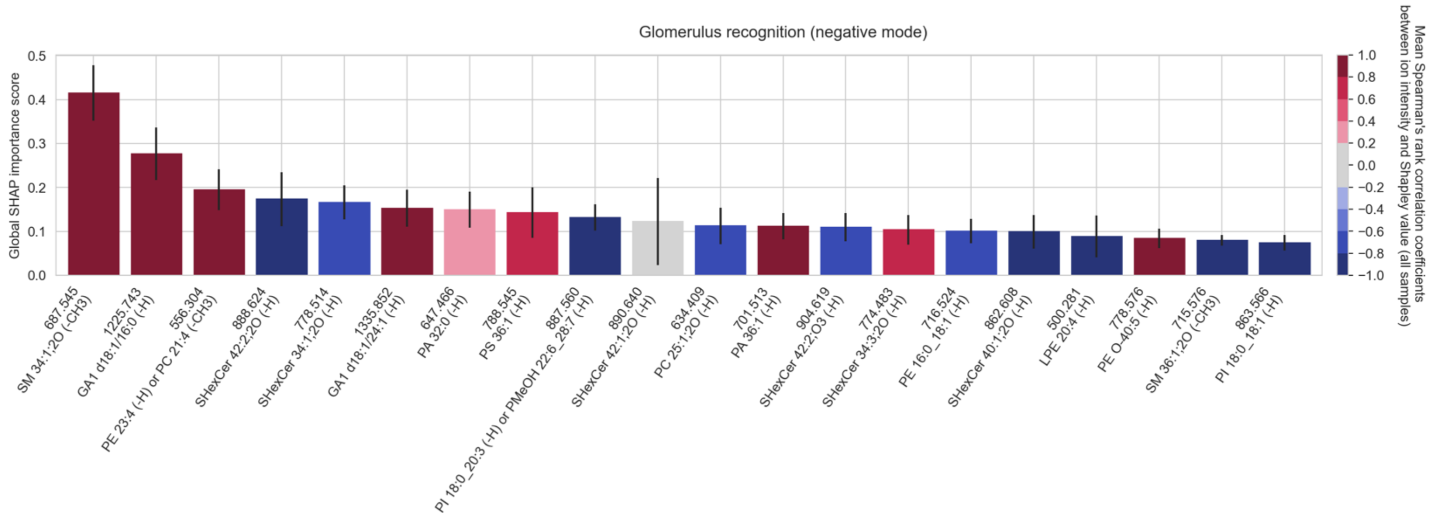

**Figure S37.** The top twenty cohort-wide (absolute) SHAP importance scores reporting the top twenty molecular species among the measured lipid species that enable the recognition (one-versus-all classification) of **<u>glomeruli in negative ion mode</u>**. The error bar is the standard deviation across all donor samples of the tissue sample-wide SHAP importance score for a given molecular species. The color of the bar reports the mean Spearman's rank correlation coefficient (across all donor samples) between the molecular species' mean-centered ion intensity and its Shapley score. A positive Spearman's rank correlation coefficient indicates that a high intensity of the molecular species correlates with the glomeruli. Conversely, a negative Spearman's rank correlation coefficient indicates that a low intensity of the molecular species correlates with the glomeruli.

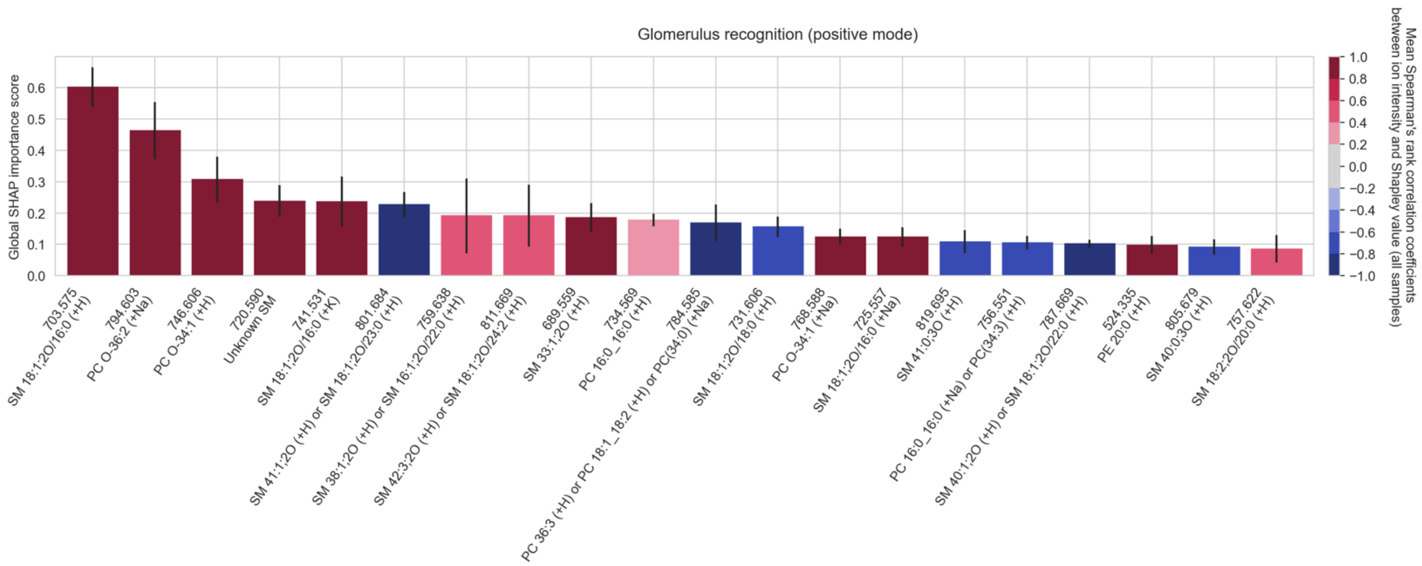

**Figure S38.** The top twenty cohort-wide (absolute) SHAP importance scores reporting the top twenty molecular species among the measured lipid species that enable the recognition (one-versus-all classification) of **<u>glomeruli in positive ion mode</u>**. The error bar is the standard deviation across all donor samples of the tissue sample-wide SHAP importance score for a given molecular species. The color of the bar reports the mean Spearman's rank correlation coefficient (across all donor samples) between the molecular species' mean-centered ion intensity and its Shapley score. A positive Spearman's rank correlation coefficient indicates that a high intensity of the molecular species correlates with the glomeruli. Conversely, a negative Spearman's rank correlation coefficient indicates that a low intensity of the molecular species correlates with the glomeruli.

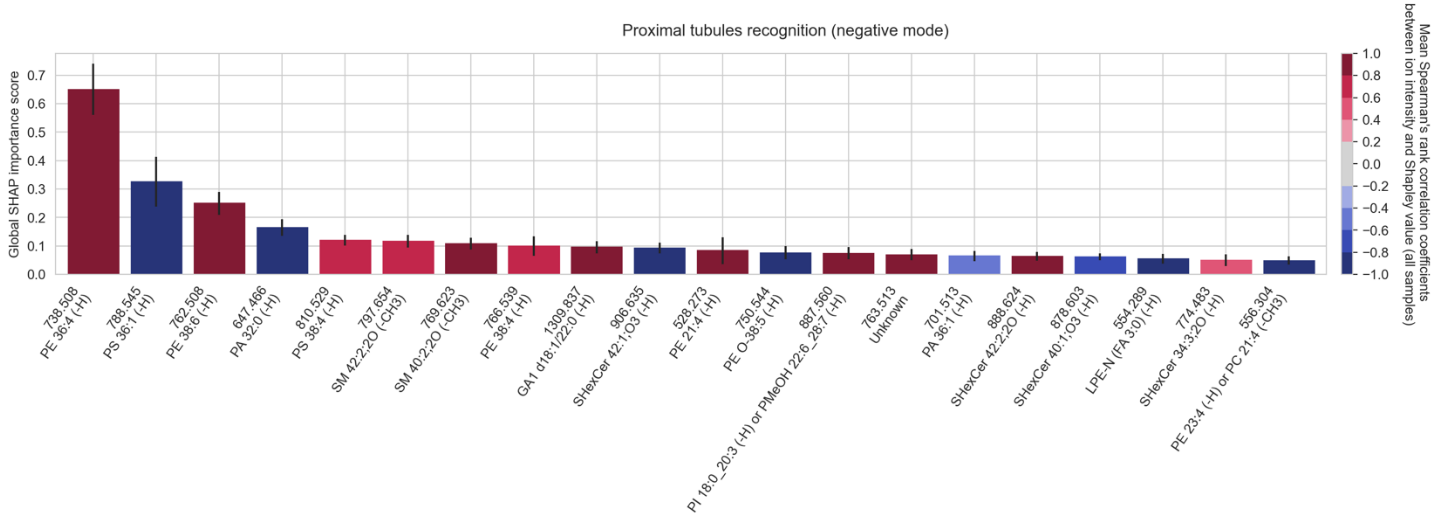

**Figure S39.** The top twenty cohort-wide (absolute) SHAP importance scores reporting the top twenty molecular species among the measured lipid species that enable the recognition (one-versus-all classification) of **proximal tubules in negative ion mode**. The error bar is the standard deviation across all donor samples of the tissue sample-wide SHAP importance score for a given molecular species. The color of the bar reports the mean Spearman's rank correlation coefficient (across all donor samples) between the molecular species' mean-centered ion intensity and its Shapley score. A positive Spearman's rank correlation coefficient indicates that a high intensity of the molecular species correlates with the proximal tubules. Conversely, a negative Spearman's rank correlation coefficient indicates that a low intensity of the molecular species correlates with the proximal tubules.

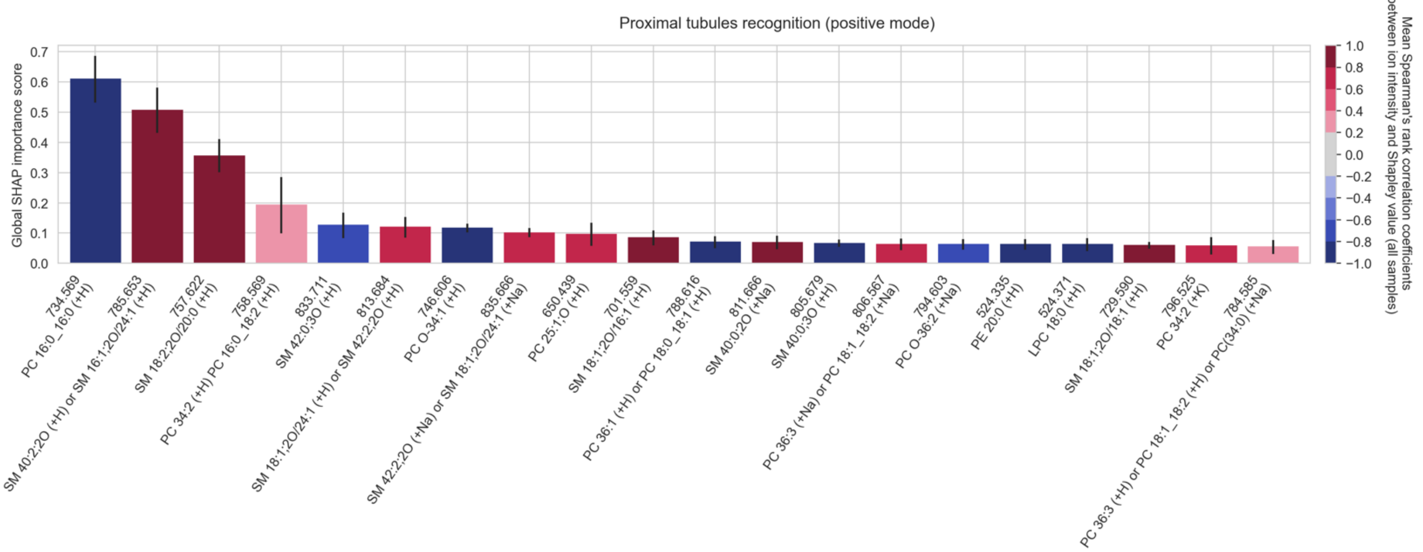

**Figure S40.** The top twenty cohort-wide (absolute) SHAP importance scores reporting the top twenty molecular species among the measured lipid species that enable the recognition (one-versus-all classification) of **proximal tubules in positive ion mode**. The error bar is the standard deviation across all donor samples of the tissue sample-wide SHAP importance score for a given molecular species. The color of the bar reports the mean Spearman's rank correlation coefficient (across all donor samples) between the molecular species' mean-centered ion intensity and its Shapley score. A positive Spearman's rank correlation coefficient indicates that a high intensity of the molecular species correlates with the proximal tubules. Conversely, a negative Spearman's rank correlation coefficient indicates that a low intensity of the molecular species correlates with the proximal tubules.

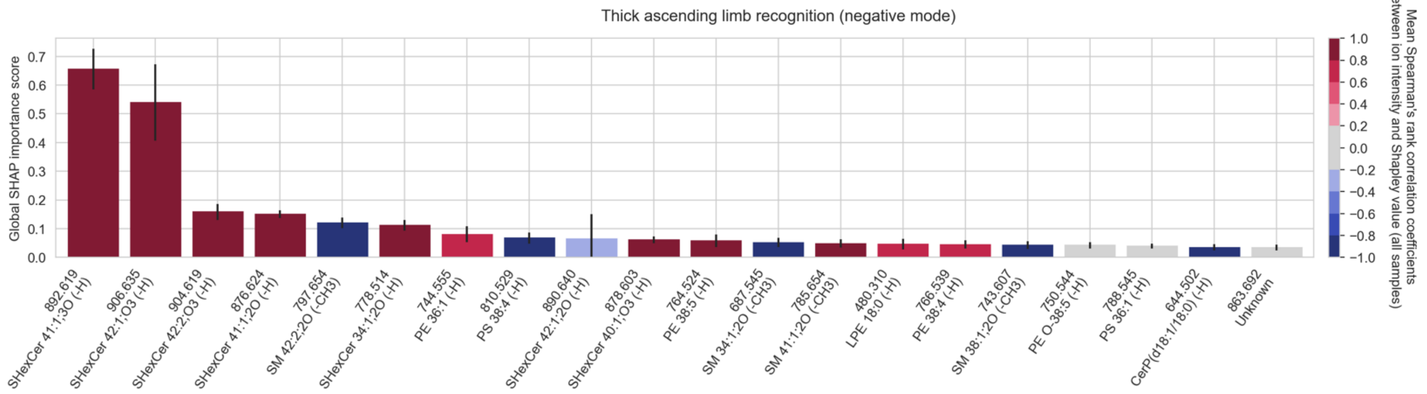

**Figure S41.** The top twenty cohort-wide (absolute) SHAP importance scores reporting the top twenty molecular species among the measured lipid species that enable the recognition (one-versus-all classification) of **thick ascending limb in negative ion mode**. The error bar is the standard deviation across all donor samples of the tissue sample-wide SHAP importance score for a given molecular species. The color of the bar reports the mean Spearman's rank correlation coefficient (across all donor samples) between the molecular species' mean-centered ion intensity and its Shapley score. A positive Spearman's rank correlation coefficient indicates that a high intensity of the molecular species correlates with thick ascending limb. Conversely, a negative Spearman's rank correlation coefficient indicates that a low intensity of the molecular species correlates with thick ascending limb.

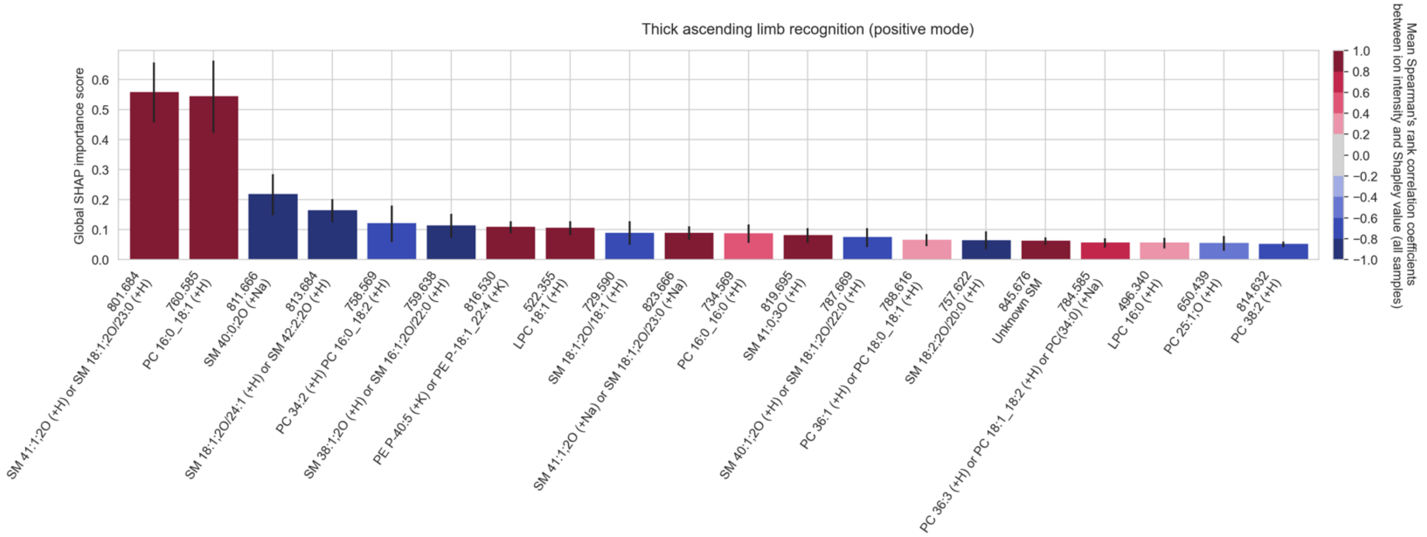

**Figure S42.** The top twenty cohort-wide (absolute) SHAP importance scores reporting the top twenty molecular species among the measured lipid species that enable the recognition (one-versus-all classification) of **thick ascending limb in positive ion mode**. The error bar is the standard deviation across all donor samples of the tissue sample-wide SHAP importance score for a given molecular species. The color of the bar reports the mean Spearman's rank correlation coefficient (across all donor samples) between the molecular species' mean-centered ion intensity and its Shapley score. A positive Spearman's rank correlation coefficient indicates that a high intensity of the molecular species correlates with thick ascending limb. Conversely, a negative Spearman's rank correlation coefficient indicates that a low intensity of the molecular species correlates with thick ascending limb.

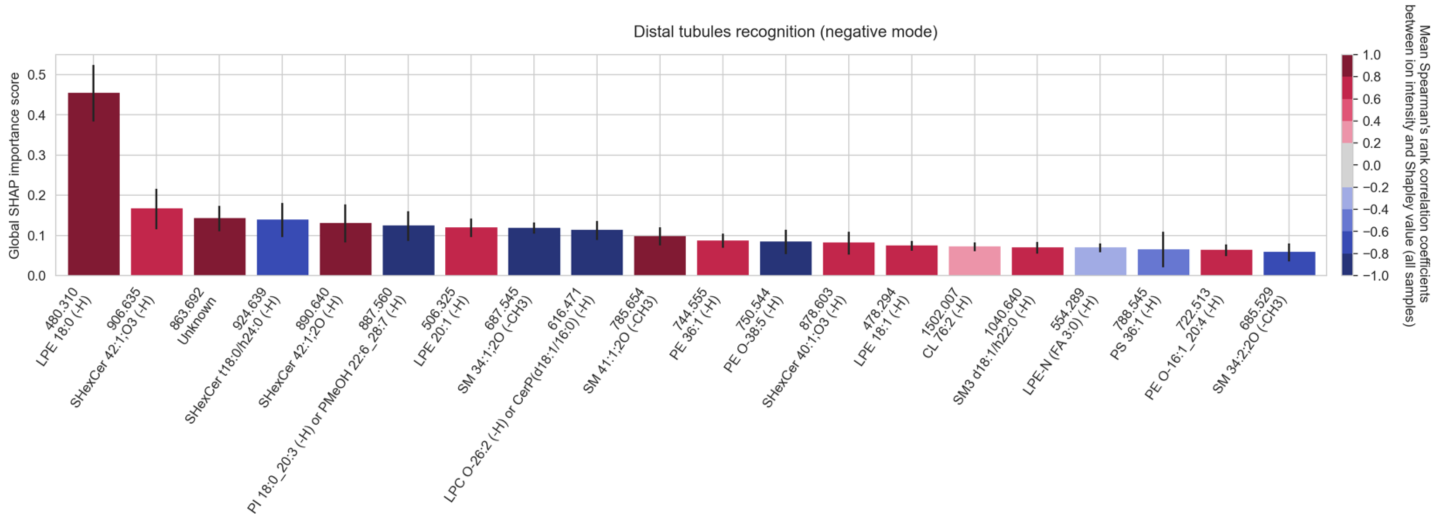

**Figure S43.** The top twenty cohort-wide (absolute) SHAP importance scores reporting the top twenty molecular species among the measured lipid species that enable the recognition (one-versus-all classification) of **distal tubules in negative ion mode**. The error bar is the standard deviation across all donor samples of the tissue sample-wide SHAP importance score for a given molecular species. The color of the bar reports the mean Spearman's rank correlation coefficient (across all donor samples) between the molecular species' mean-centered ion intensity and its Shapley score. A positive Spearman's rank correlation coefficient indicates that a high intensity of the molecular species correlates with the distal tubules. Conversely, a negative Spearman's rank correlation coefficient indicates that a low intensity of the molecular species correlates with the distal tubules.

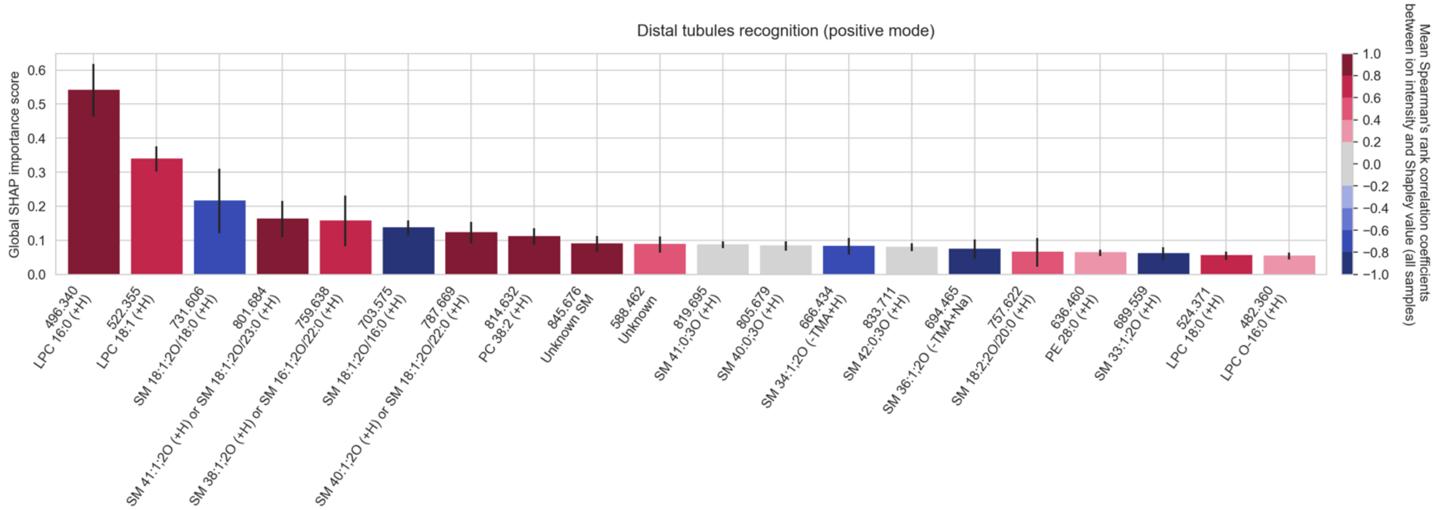

**Figure S44.** The top twenty cohort-wide (absolute) SHAP importance scores reporting the top twenty molecular species among the measured lipid species that enable the recognition (one-versus-all classification) of **distal tubules in positive ion mode**. The error bar is the standard deviation across all donor samples of the tissue sample-wide SHAP importance score for a given molecular species. The color of the bar reports the mean Spearman's rank correlation coefficient (across all donor samples) between the molecular species' mean-centered ion intensity and its Shapley score. A positive Spearman's rank correlation coefficient indicates that a high intensity of the molecular species correlates with the distal tubules. Conversely, a negative Spearman's rank correlation coefficient indicates that a low intensity of the molecular species correlates with the distal tubules.

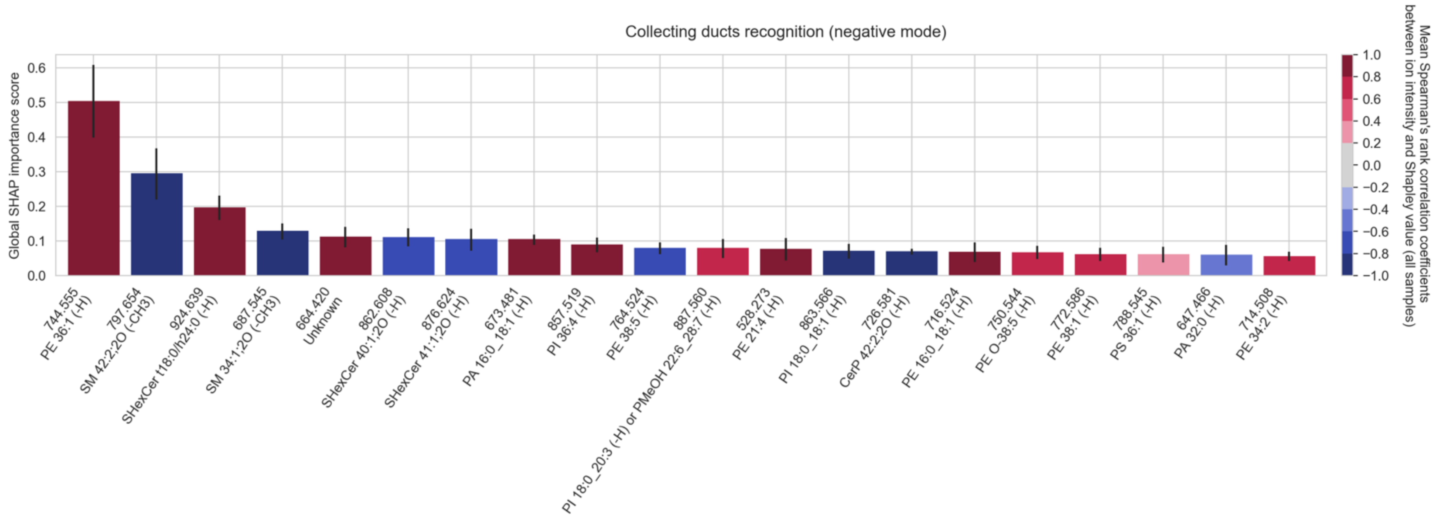

**Figure S45.** The top twenty cohort-wide (absolute) SHAP importance scores reporting the top twenty molecular species among the measured lipid species that enable the recognition (one-versus-all classification) of **collecting ducts in negative ion mode**. The error bar is the standard deviation across all donor samples of the tissue sample-wide SHAP importance score for a given molecular species. The color of the bar reports the mean Spearman's rank correlation coefficient (across all donor samples) between the molecular species' mean-centered ion intensity and its Shapley score. A positive Spearman's rank correlation coefficient indicates that a high intensity of the molecular species correlates with the collecting ducts. Conversely, a negative Spearman's rank correlation coefficient indicates that a low intensity of the molecular species correlates with the collecting ducts.

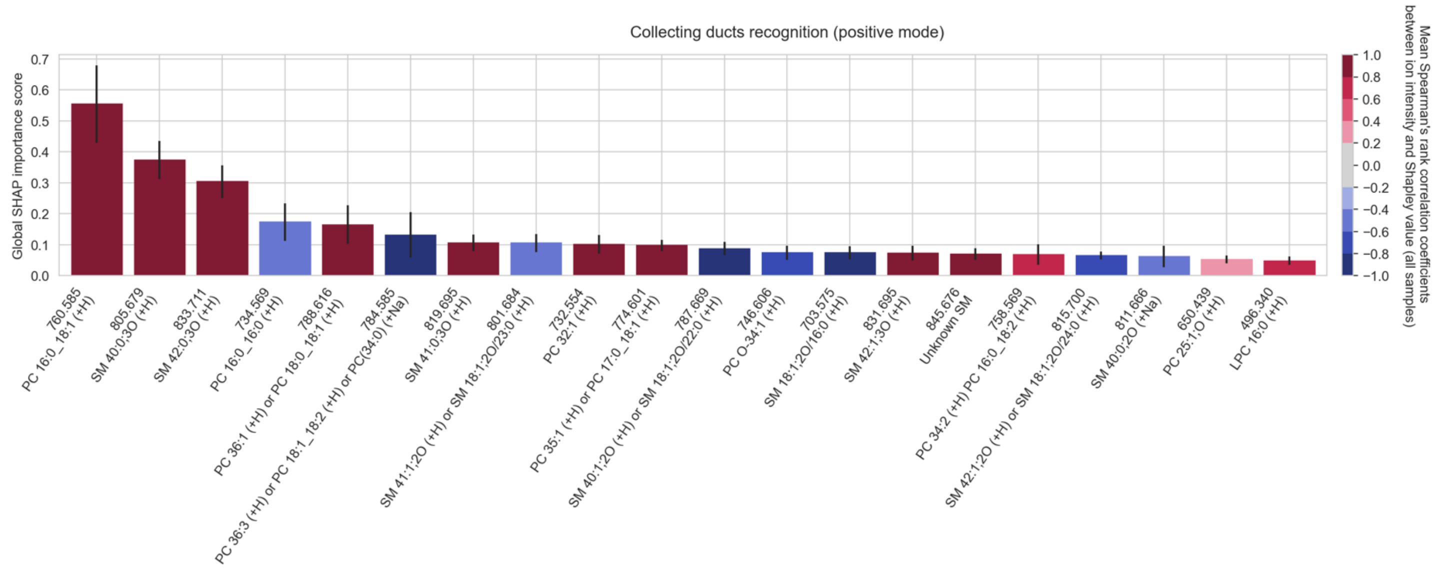

**Figure S46.** The top twenty cohort-wide (absolute) SHAP importance scores reporting the top twenty molecular species among the measured lipid species that enable the recognition (one-versus-all classification) of **collecting ducts in positive ion mode**. The error bar is the standard deviation across all donor samples of the tissue sample-wide SHAP importance score for a given molecular species. The color of the bar reports the mean Spearman's rank correlation coefficient (across all donor samples) between the molecular species' mean-centered ion intensity and its Shapley score. A positive Spearman's rank correlation coefficient indicates that a high intensity of the molecular species correlates with the collecting ducts. Conversely, a negative Spearman's rank correlation coefficient indicates that a low intensity of the molecular species correlates with the collecting ducts.

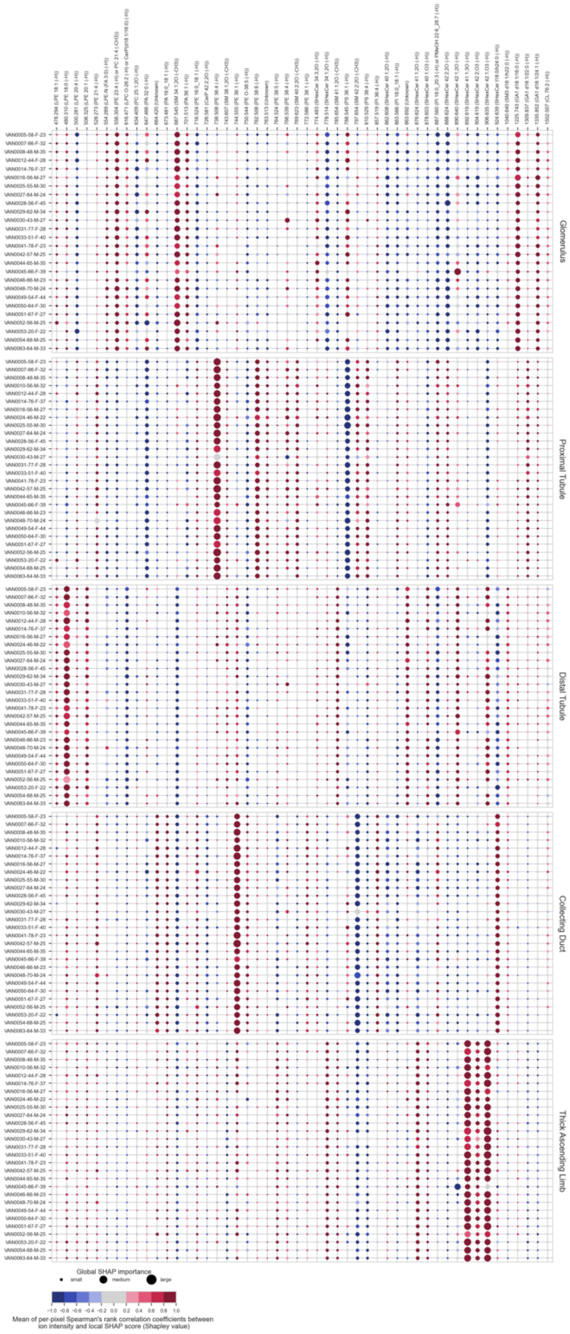

**Figure S47.** Summary of **FTU-specific biomarker candidates in negative ion mode**, obtained by applying our SHAP-based workflow to the atlas. The columns correspond to a selection of molecular species (in increasing order of mass-to-charge ratios) that are biomarker candidates for one (or multiple) of the five FTUs under study. The rows correspond to different donors, each row labeled with its donor ID number and followed by the donor's age, sex, and BMI. Each bubble marker is informative of the direction (positive or negative correlation) and magnitude (relatively large or small) of a molecular species' influence on the classification model designed to recognize one of the five FTUs. The marker size represents the magnitude of the molecular species' influence, as measured by its tissue sample-wide SHAP importance score for a given donor sample. The marker color indicates the direction of the molecular species' influence, as measured by the Spearman's rank correlation coefficient between the molecular species' mean-centered ion intensity values and its local pixel-specific SHAP scores. A positive Spearman's rank correlation coefficient indicates that a high ion intensity of the molecular species correlates with the FTU. Conversely, a negative Spearman's rank correlation coefficient indicates that a low ion intensity of the molecular species correlates with the FTU.

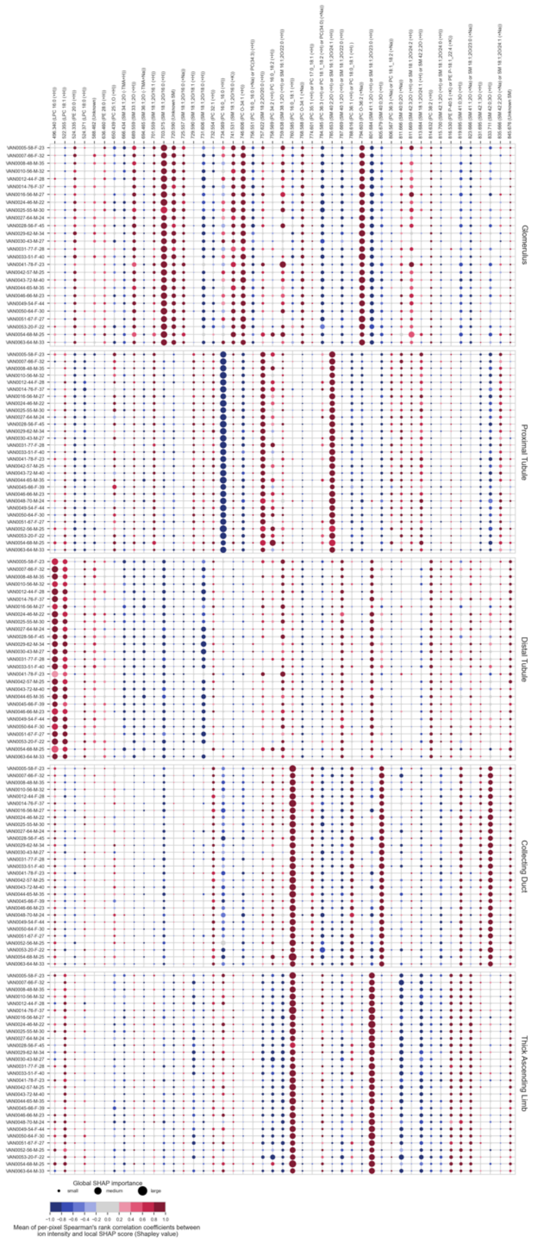

**Figure S48.** Summary of **FTU-specific biomarker candidates in positive ion mode**, obtained by applying our SHAP-based workflow to the atlas. The columns correspond to a selection of molecular species (in increasing order of mass-to-charge ratios) that are biomarker candidates for one (or multiple) of the five FTUs under study. The rows correspond to different donors, each row labeled with its donor ID number and followed by the donor's age, sex, and BMI. Each bubble marker is informative of the direction (positive or negative correlation) and magnitude (relatively large or small) of a molecular species' influence on the classification model designed to recognize one of the five FTUs. The marker size represents the magnitude of the molecular species' influence, as measured by its tissue sample-wide SHAP importance score for a given donor sample. The marker color indicates the direction of the molecular species' influence, as measured by the Spearman's rank correlation coefficient between the molecular species' mean-centered ion intensity values and its local pixel-specific SHAP scores. A positive Spearman's rank correlation coefficient indicates that a high ion intensity of the molecular species correlates with the FTU. Conversely, a negative Spearman's rank correlation coefficient indicates that a low ion intensity of the molecular species correlates with the FTU.

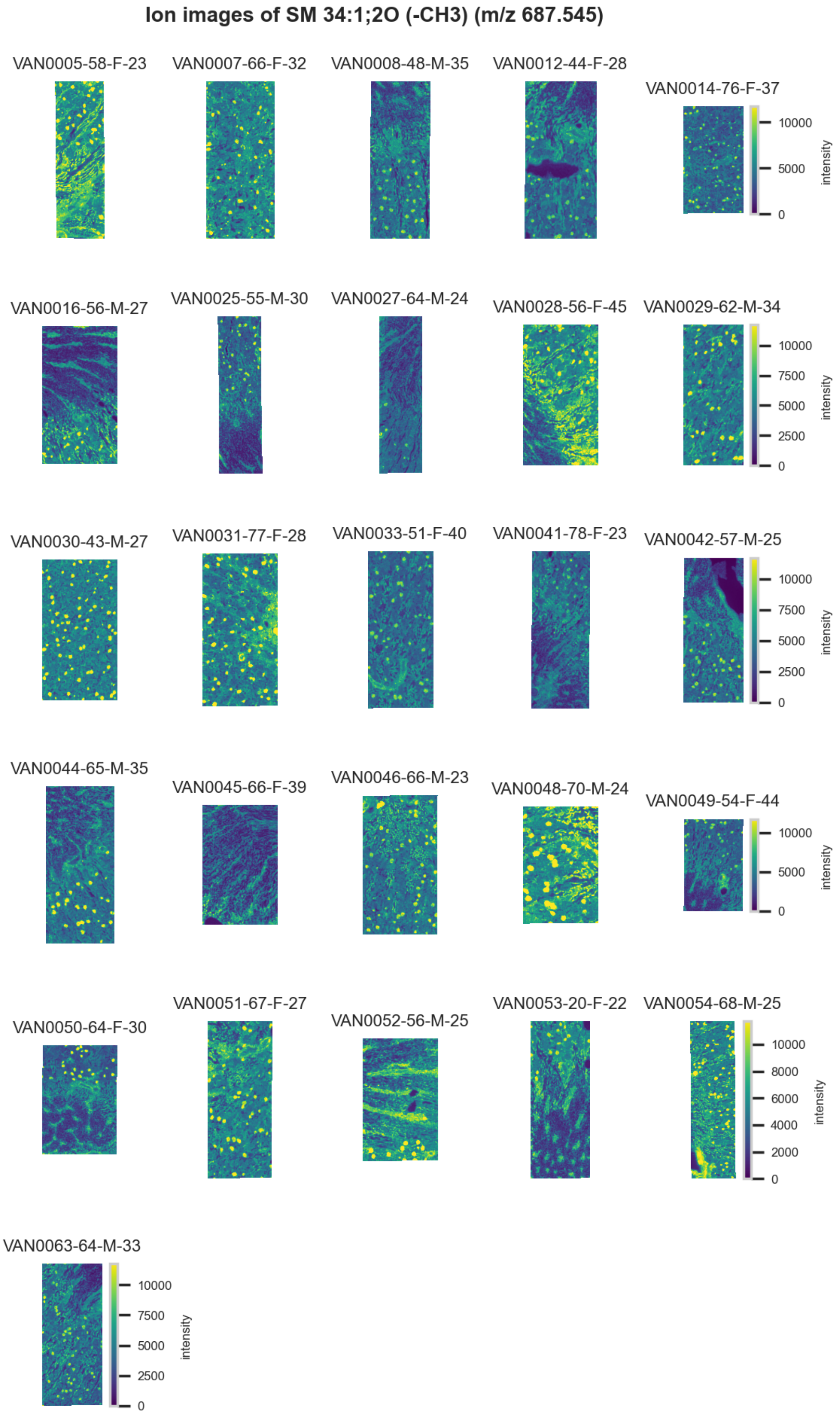

**Figure S49.** 10 μm MALDI IMS images of $[SM(34:1,2O)-CH_3]^-$ (*m/z* 687.545) detected in negative ion mode from each donor sample. Multivariate SHAP analysis determined this to be feature 1 for classification of glomeruli.

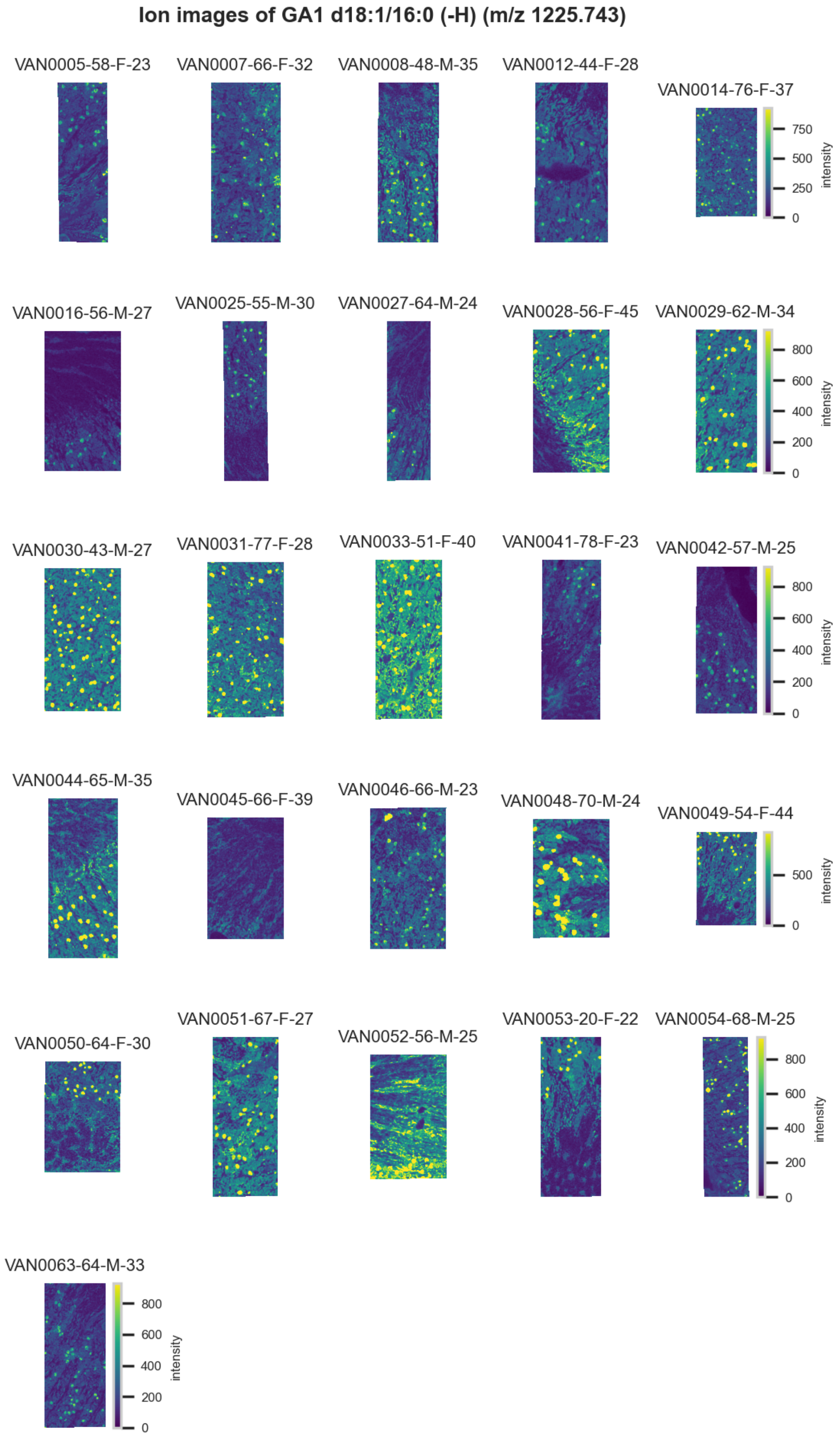

**Figure S50.** 10 µm MALDI IMS images of [GA1(d18:1/16:0)-H]$^-$ (*m/z* 1,225.743) detected in negative ion mode from each donor sample. Multivariate SHAP analysis determined this to be feature 2 for classification of glomeruli.

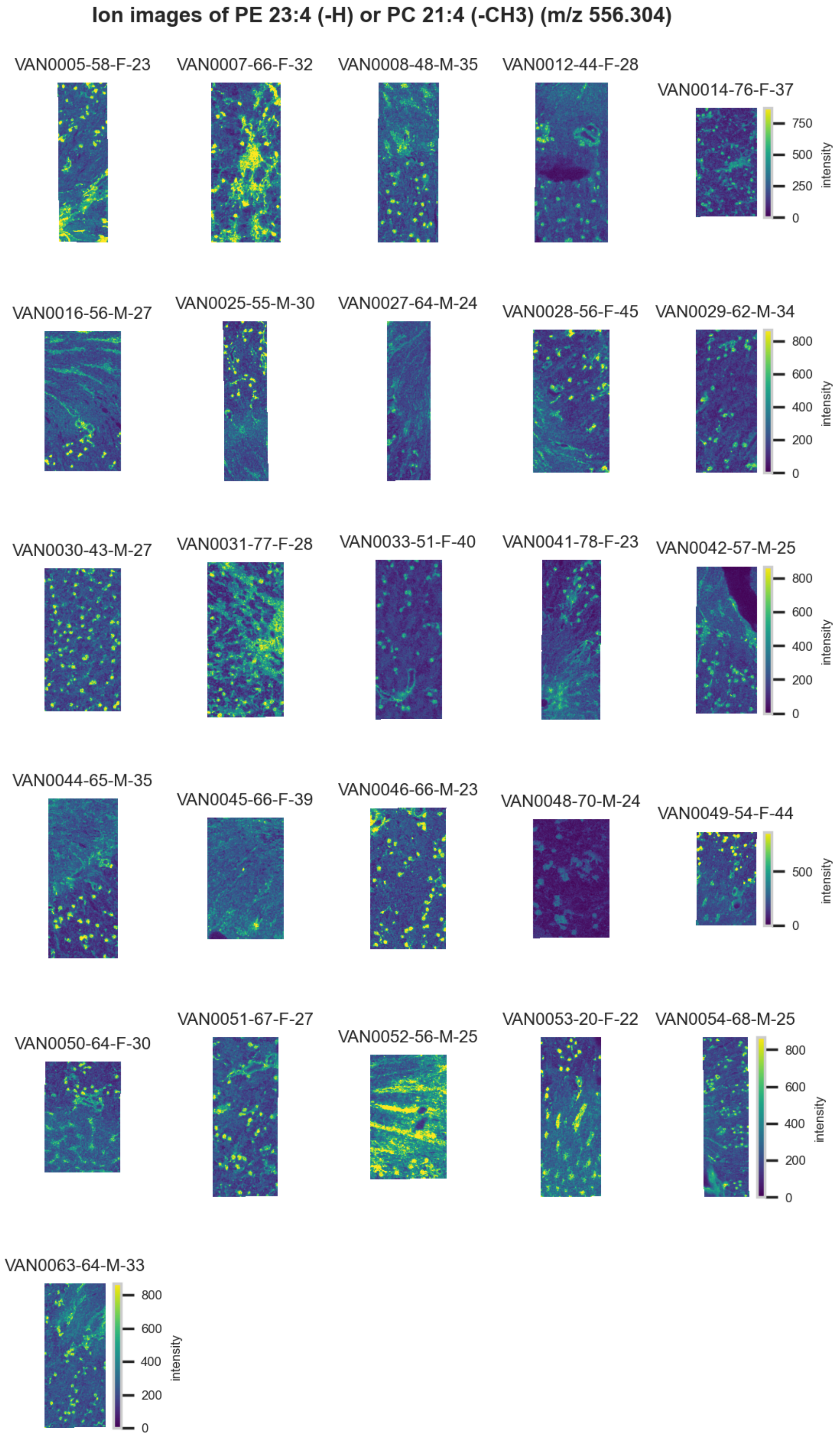

**Figure S51.** 10 µm MALDI IMS images of [PE(23:4)-H]$^-$ or [PC(21:4)-CH3]$^-$ (*m/z* 556.304) detected in <u>negative ion mode</u> from each donor sample. These molecules are isomers and therefore indistinguishable by MS alone. Multivariate SHAP analysis determined this to be feature 3 for classification of glomeruli.

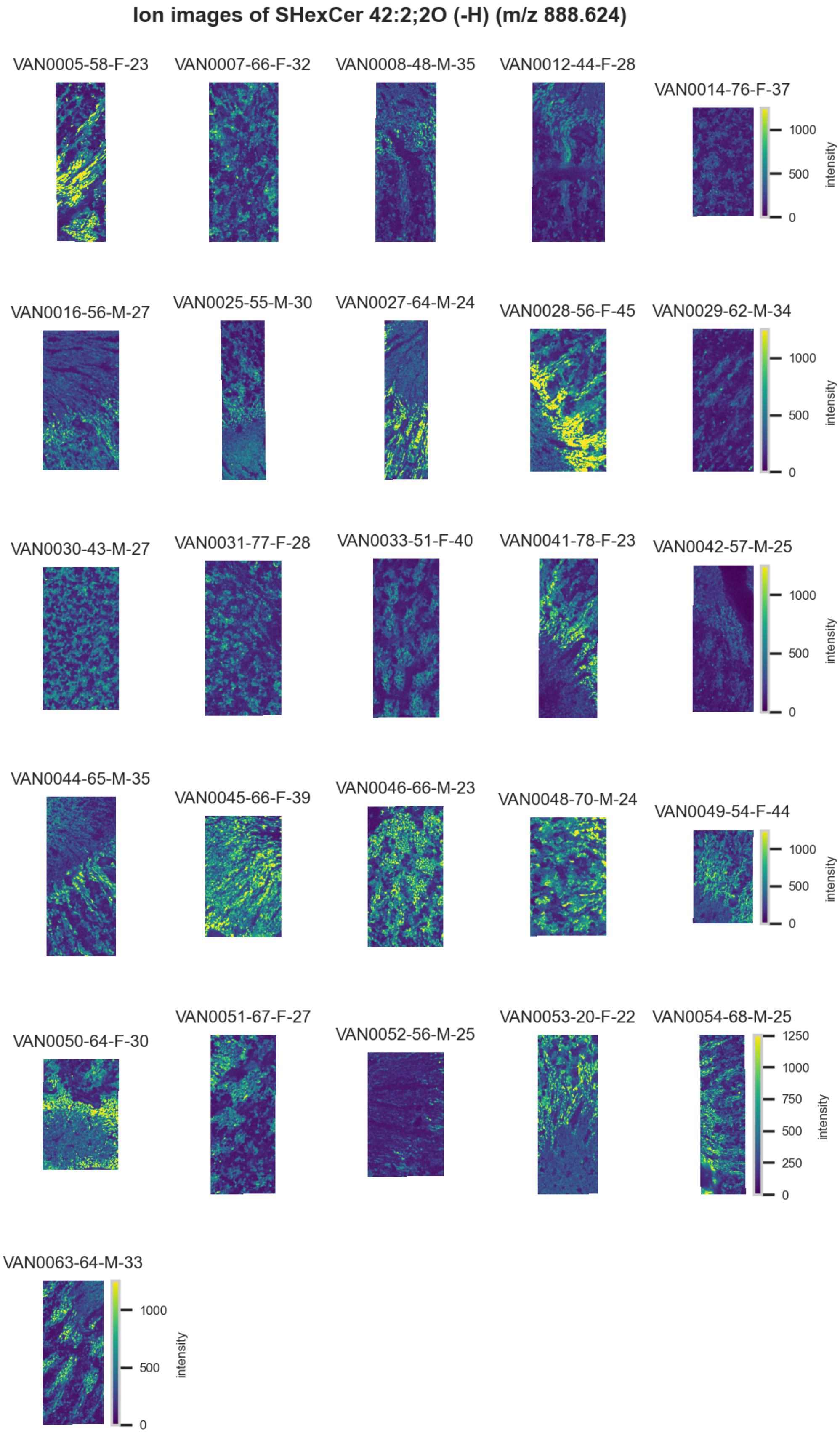

**Figure S52.** 10 µm MALDI IMS images of [SHexCer(42:2,2O)-H]$^-$ (*m/z* 888.624) detected in negative ion mode from each donor sample. Multivariate SHAP analysis determined this to be feature 4 for classification of glomeruli.

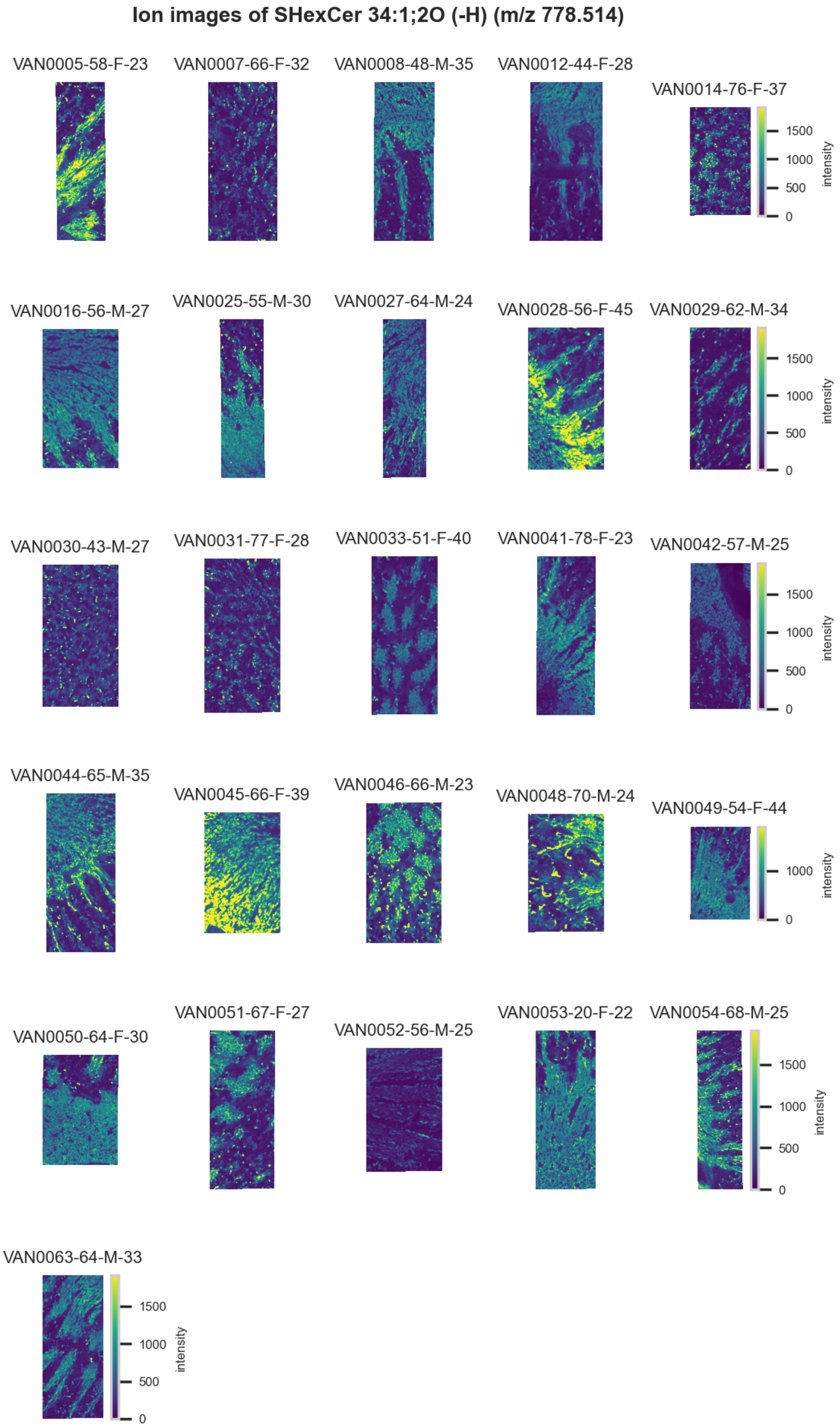

**Figure S53.** 10 µm MALDI IMS images of [SHexCer(34:1,2O)-H]$^{-}$ (*m/z* 778.514) detected in negative ion mode from each donor sample. Multivariate SHAP analysis determined this to be feature 5 for classification of glomeruli.

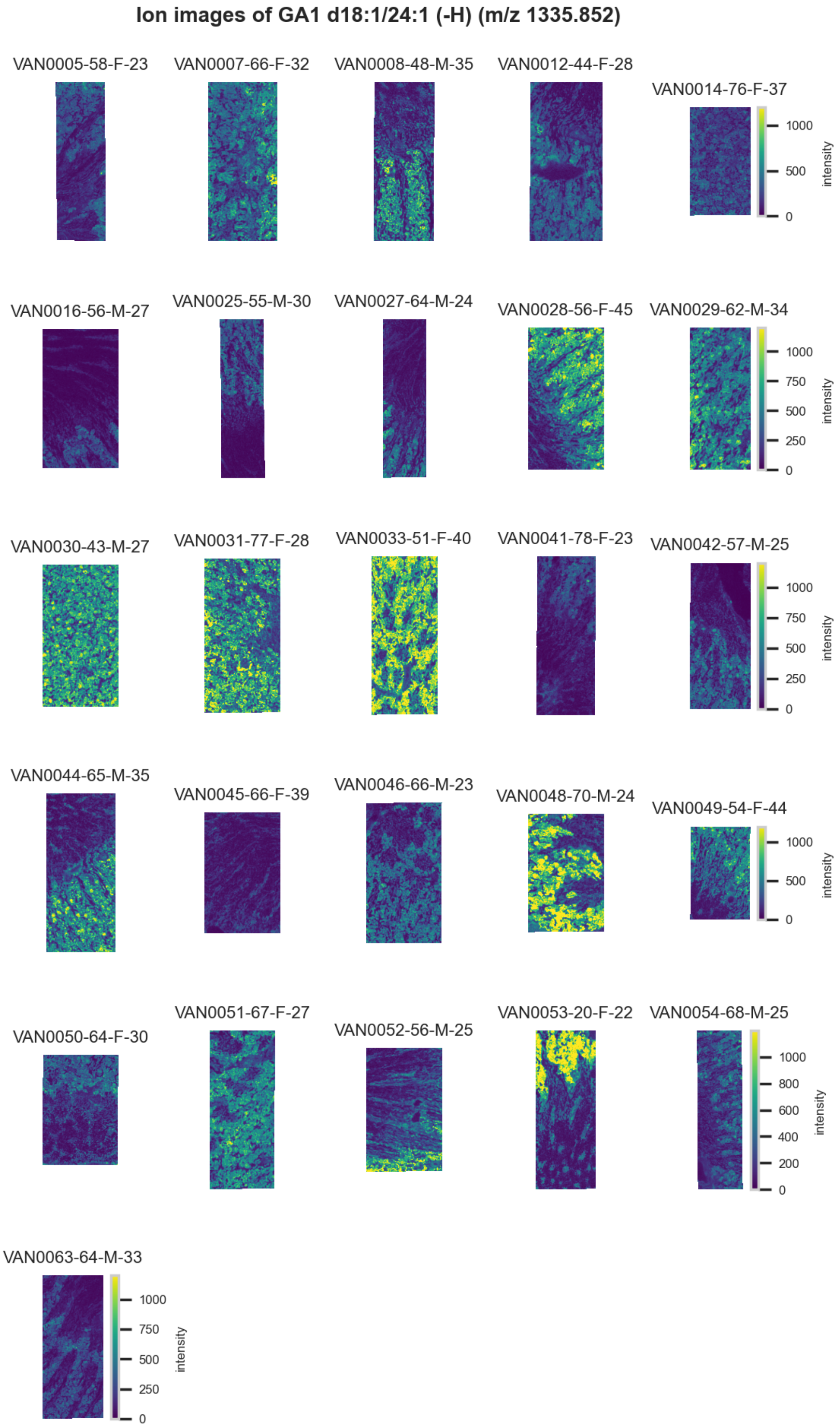

**Figure S54.** 10 µm MALDI IMS images of $[GA1(d18:1/24:1)-H]^-$ (*m/z* 1,335.852) detected in negative ion mode from each donor sample. Multivariate SHAP analysis determined this to be feature 6 for classification of glomeruli.

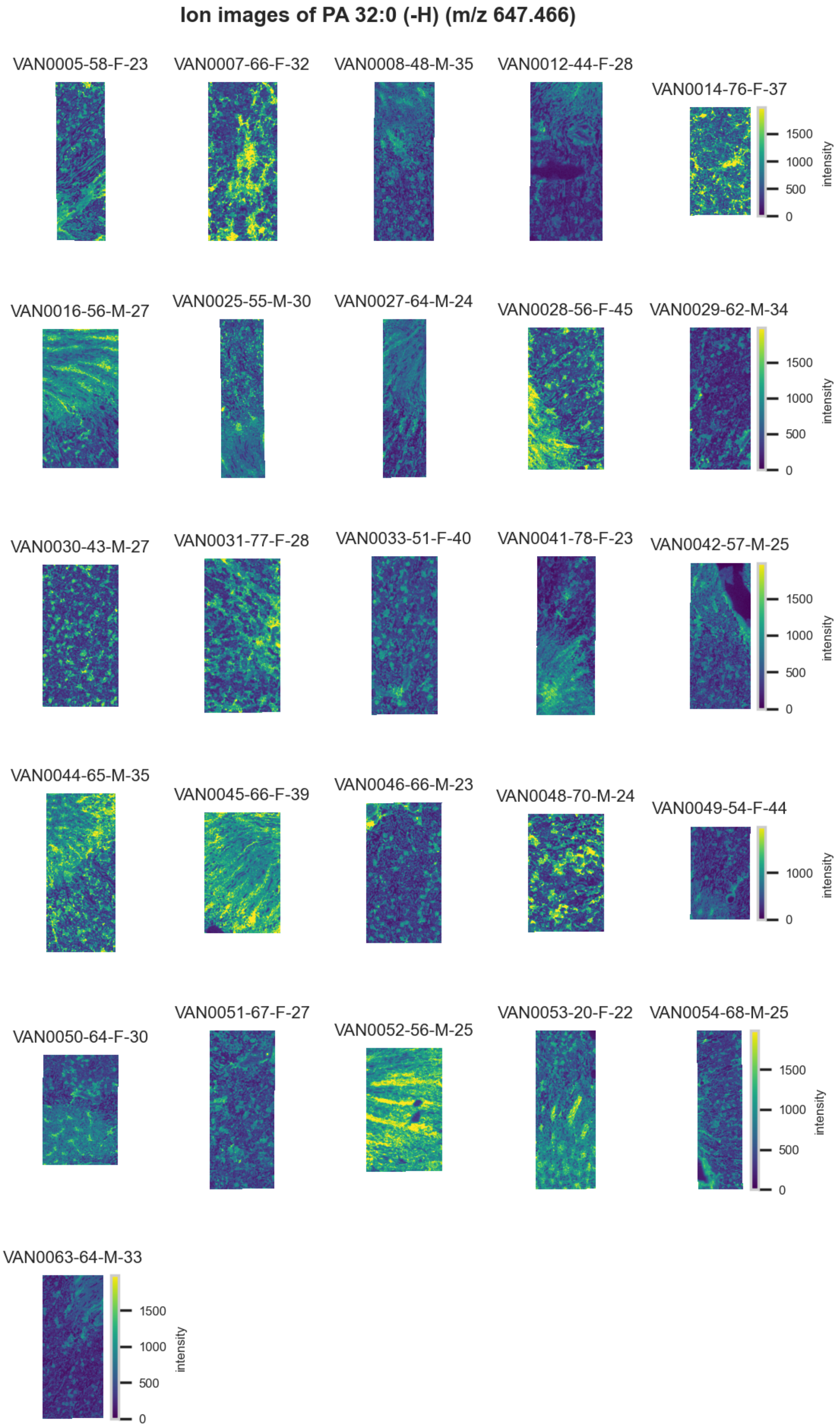

**Figure S55.** 10 µm MALDI IMS images of [PA(32:0)-H]$^-$ (*m/z* 647.466) detected in negative ion mode from each donor sample. Multivariate SHAP analysis determined this to be feature 7 for classification of glomeruli.

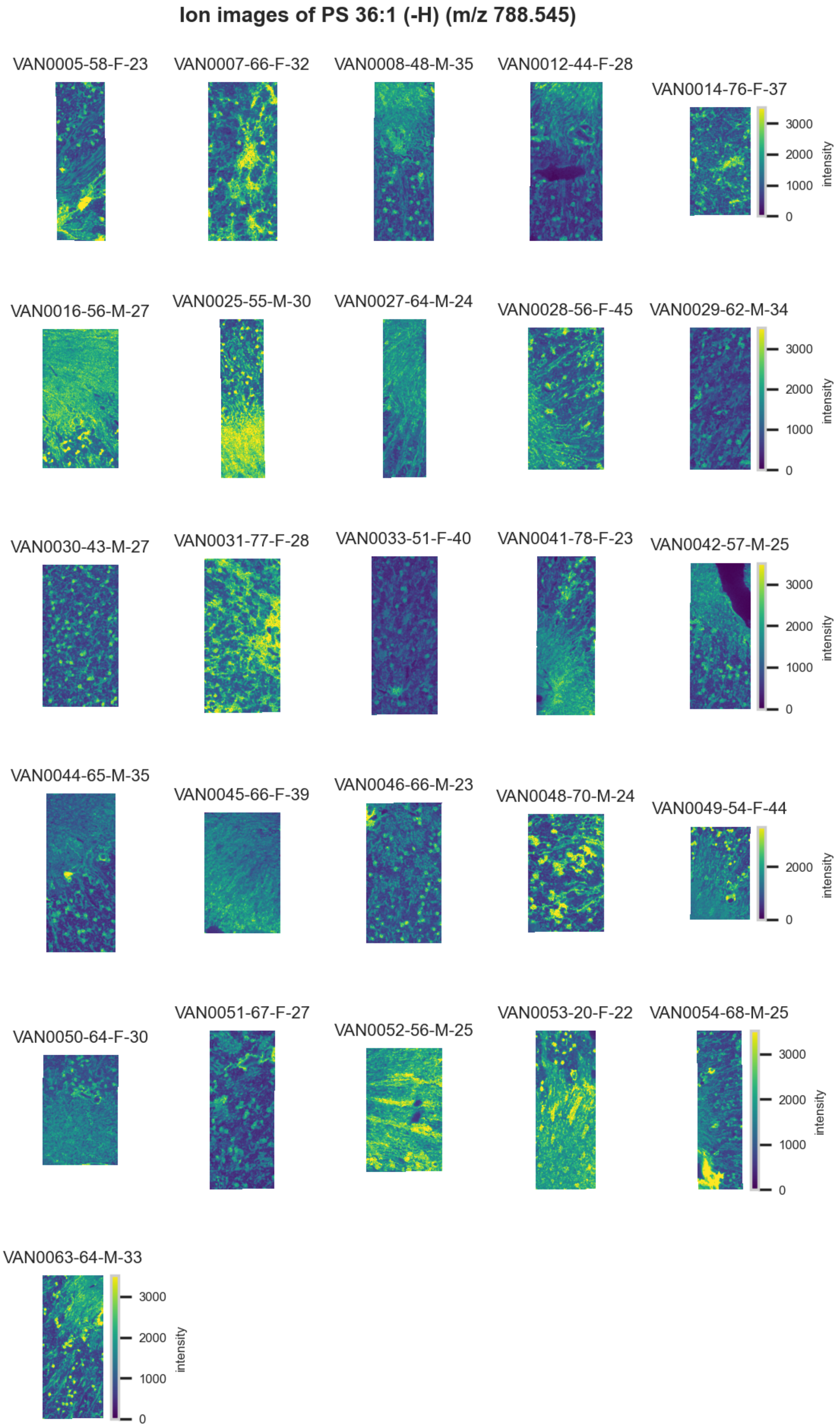

**Figure S56.** 10 µm MALDI IMS images of [PS(36:1)-H]$^-$ (*m/z* 788.545) detected in <u>negative ion mode</u> from each donor sample. Multivariate SHAP analysis determined this to be feature 8 for classification of glomeruli.

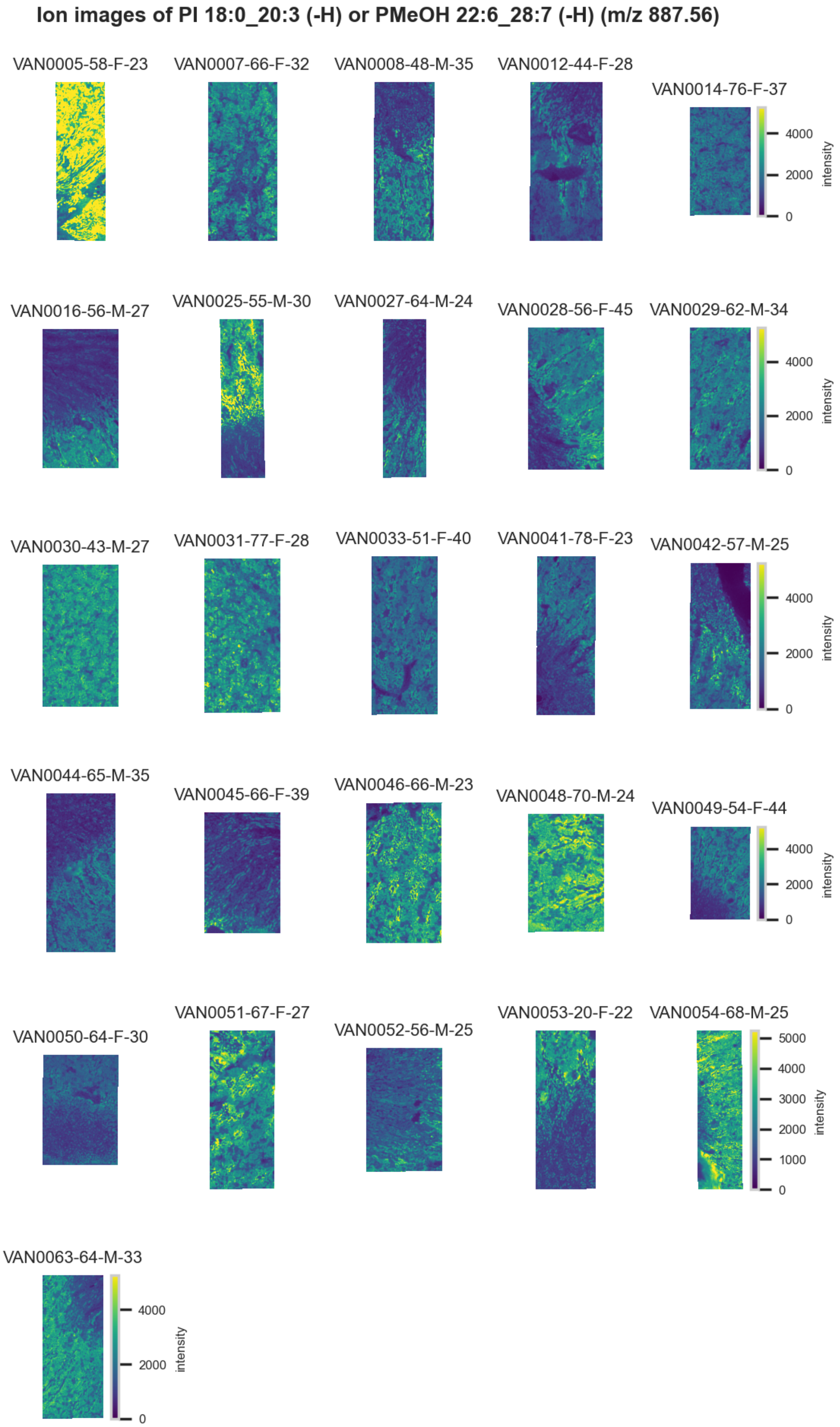

**Figure S57.** 10 µm MALDI IMS images of [PI(18:0_20:3)-H]$^-$ or [PMeOH(22:6_28:7)-H]$^-$ (*m/z* 887.56) detected in negative ion mode from each donor sample. These molecules are isomers and therefore indistinguishable by MS alone. Multivariate SHAP analysis determined this to be feature 9 for classification of glomeruli.

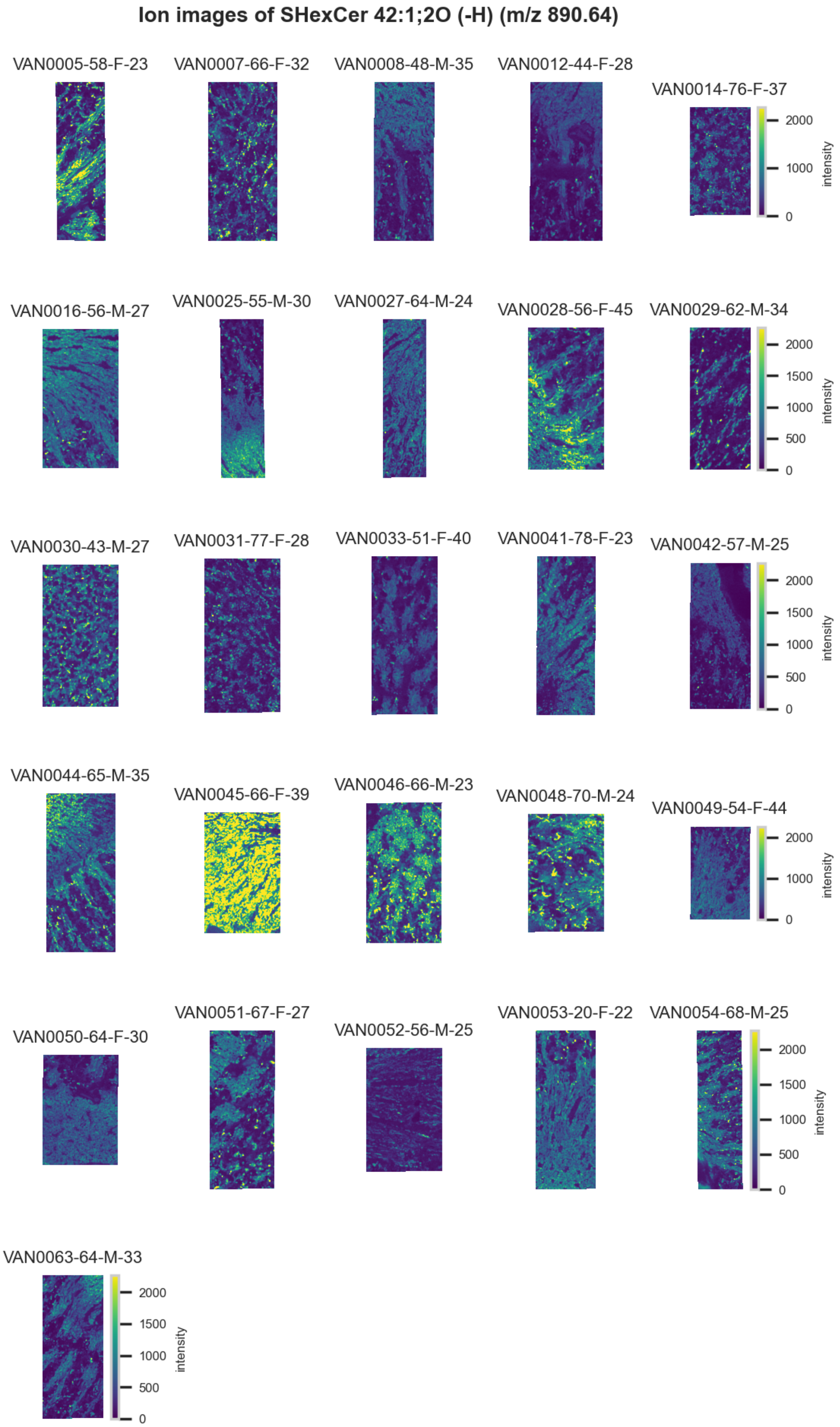

**Figure 58.** 10 µm MALDI IMS images of $[SHexCer(42{:}1{,}2O){-}H]^-$ (*m/z* 890.64) detected in <u>negative ion mode</u> from each donor sample. Multivariate SHAP analysis determined this to be feature 10 for classification of glomeruli.

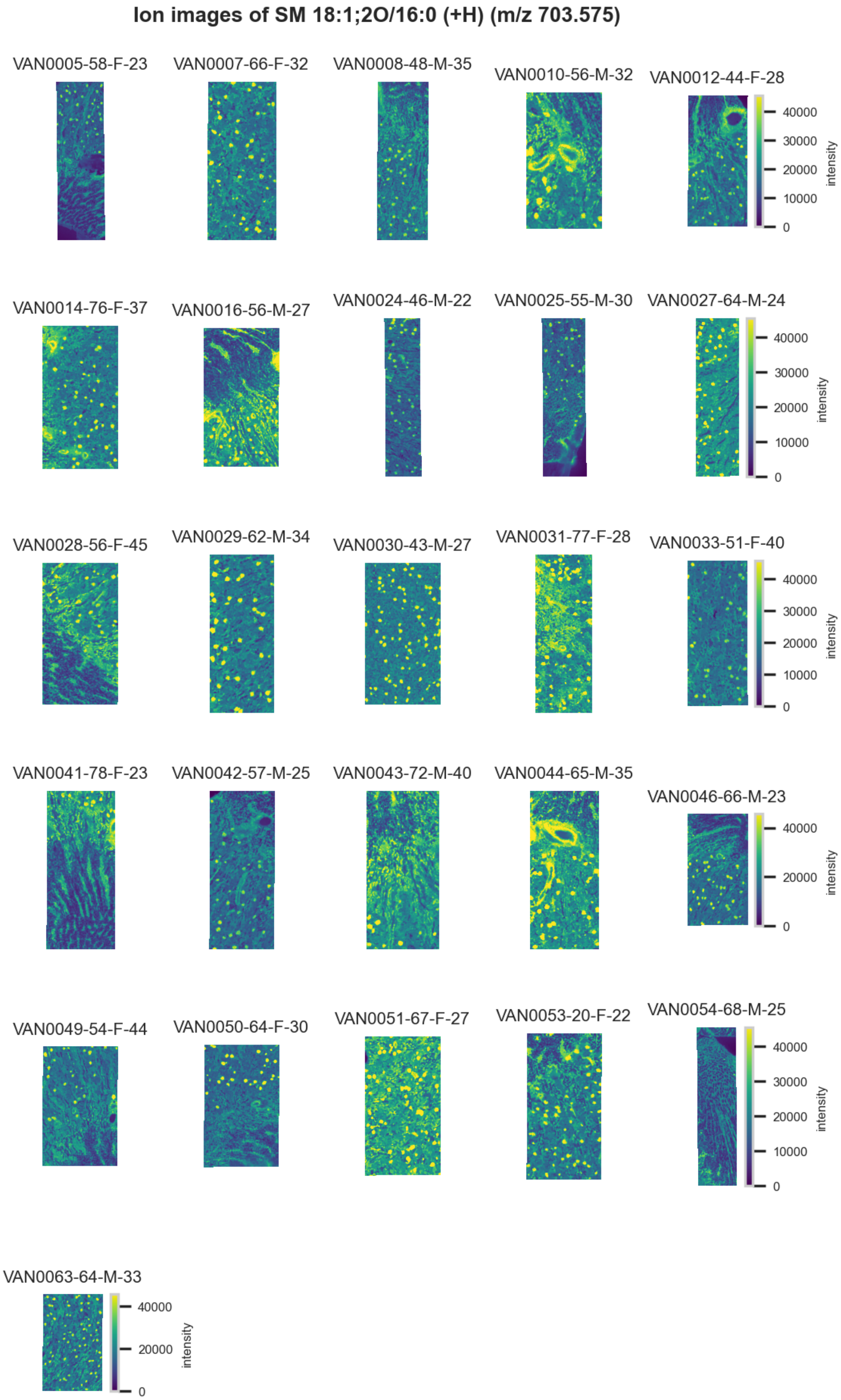

**Figure S59.** 10 μm MALDI IMS images of [SM(18:1,2O:16:0)+H]$^{+}$ (*m/z* 703.575) detected in positive ion mode from each donor sample. Multivariate SHAP analysis determined this to be feature 1 for classification of glomeruli.

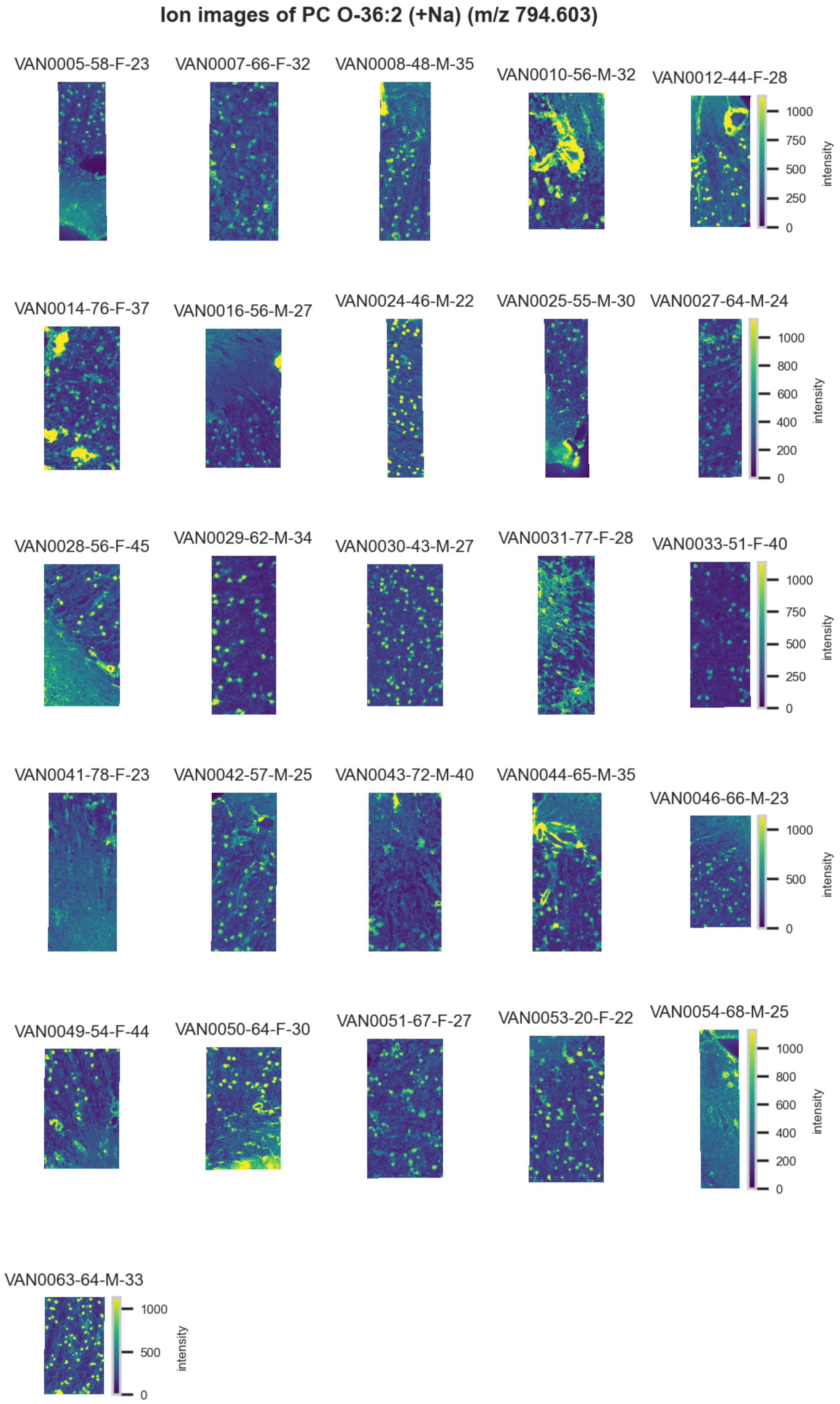

**Figure S60.** 10 μm MALDI IMS images of $[PC(O\text{-}36{:}2)+Na]^+$ (*m/z* 794.603) detected in positive ion mode from each donor sample. Multivariate SHAP analysis determined this to be feature 2 for classification of glomeruli.

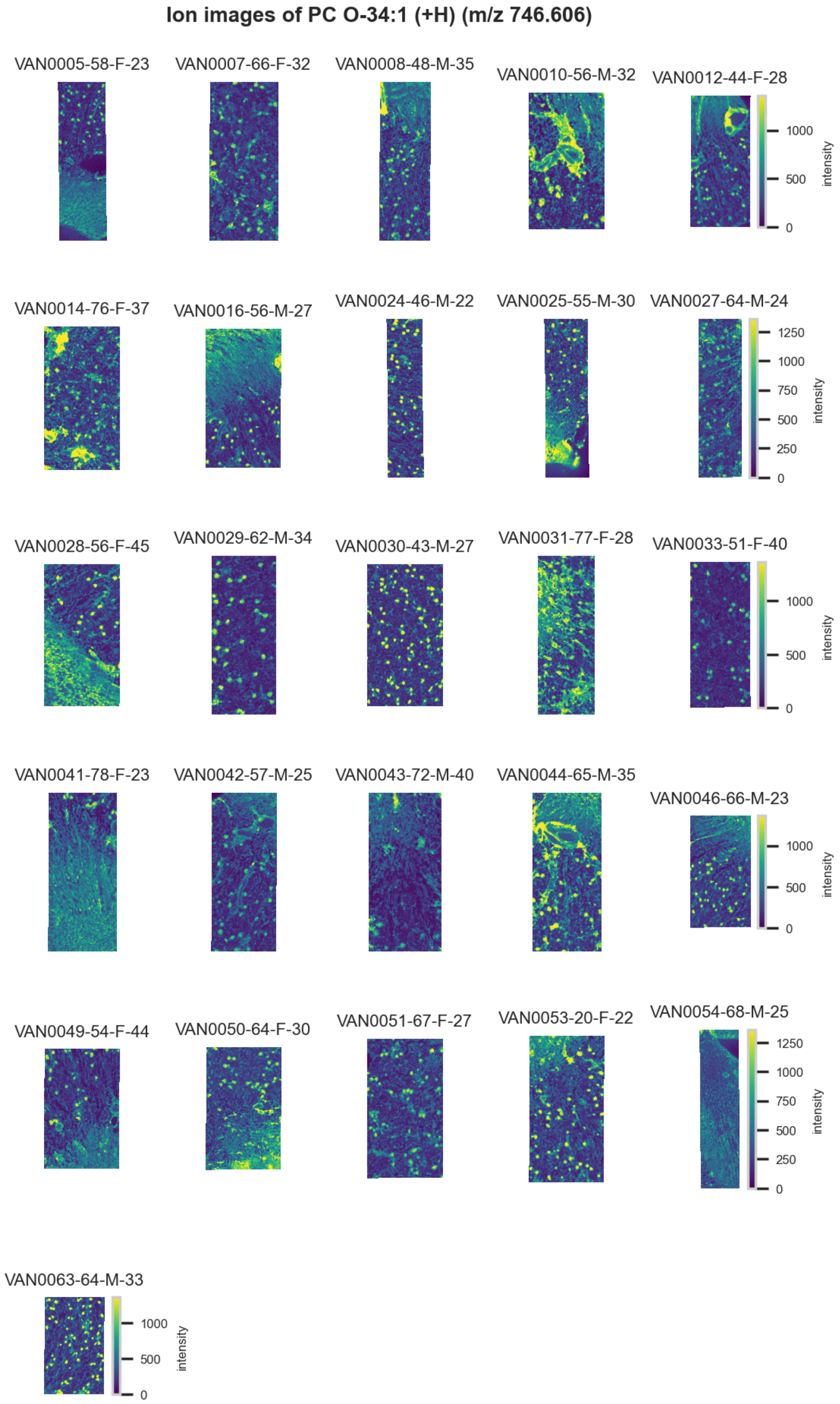

**Figure S61.** 10 µm MALDI IMS images of $[PC(O\text{-}34:1)+H]^+$ (*m/z* 746.606) detected in positive ion mode from each donor sample. Multivariate SHAP analysis determined this to be feature 3 for classification of glomeruli.

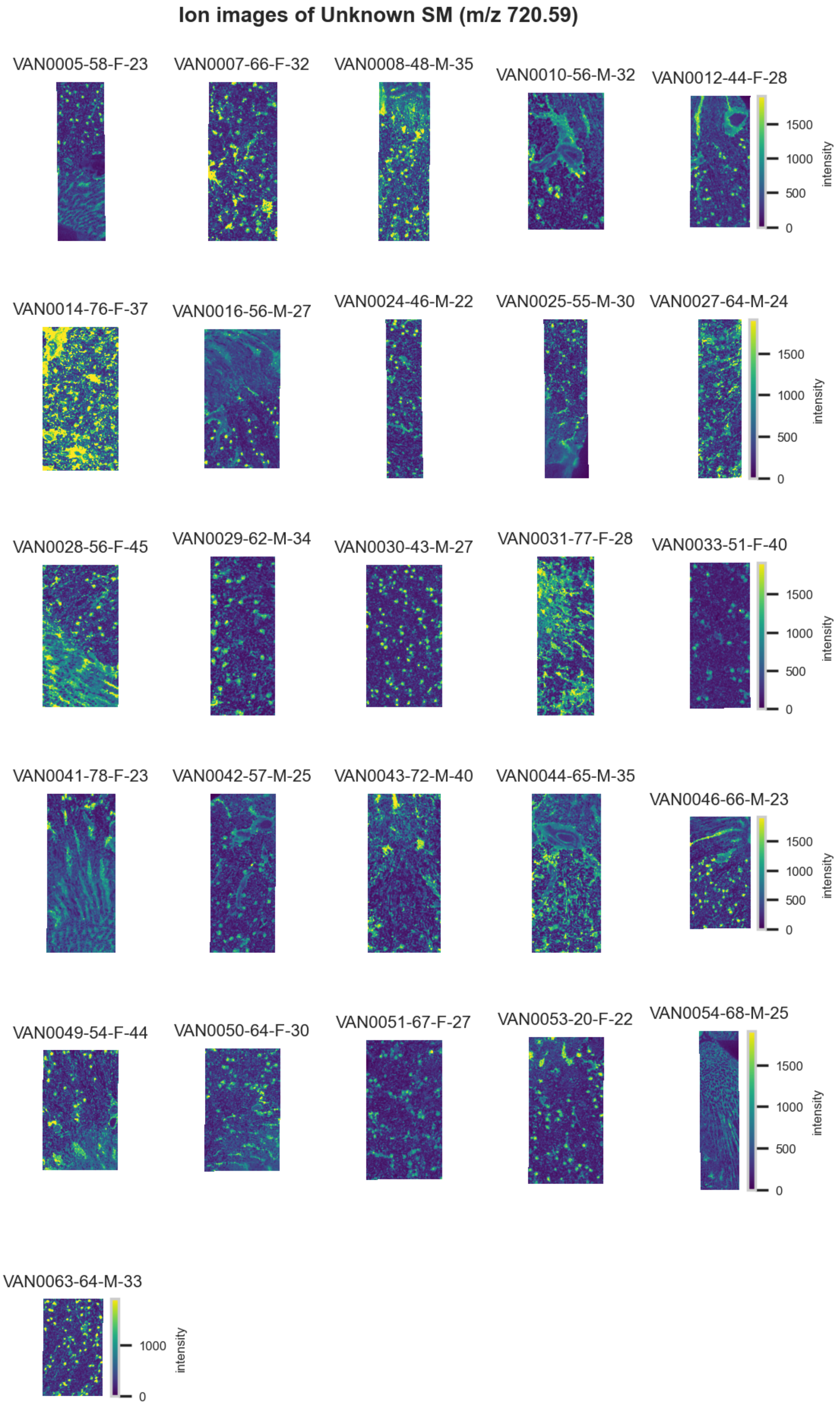

**Figure S62.** 10 µm MALDI IMS images of an unknown SM lipid (*m/z* 720.59) detected in positive ion mode from each donor sample. Multivariate SHAP analysis determined this to be feature 4 for classification of glomeruli.

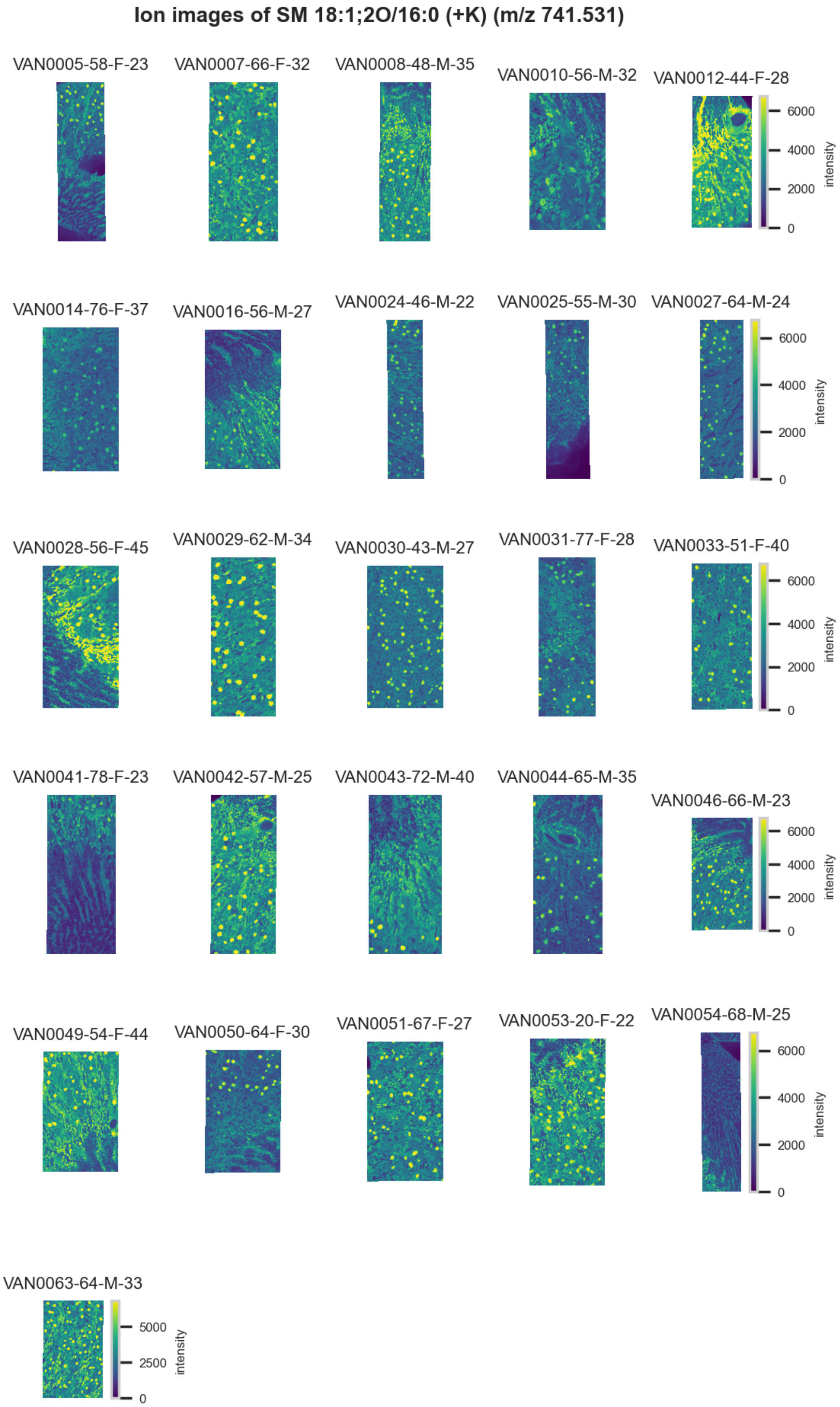

**Figure S63.** 10 µm MALDI IMS images of $[SM(18:1,2O/16:0)+K]^+$ (*m/z* 741.531) detected in positive ion mode from each donor sample. Multivariate SHAP analysis determined this to be feature 5 for classification of glomeruli.

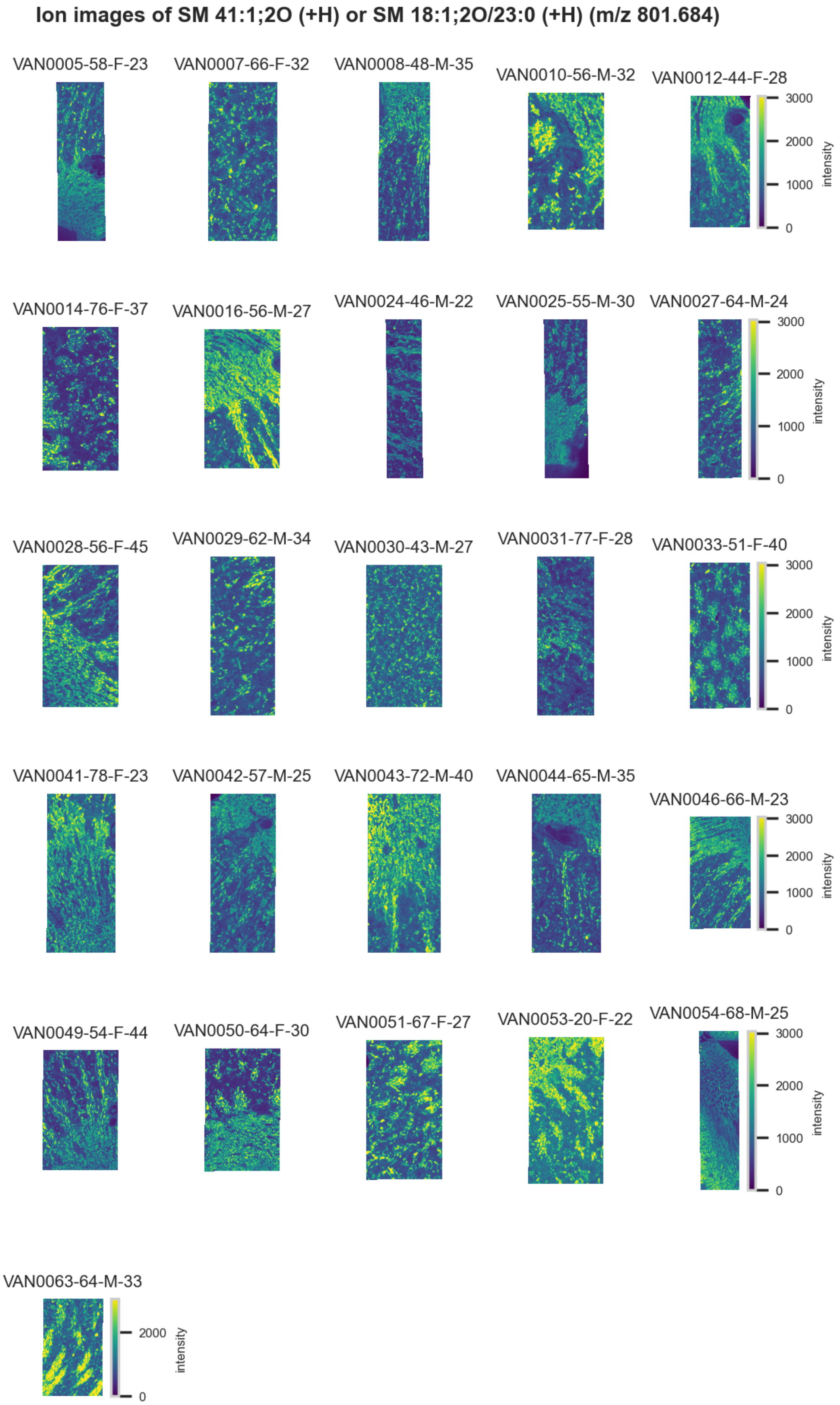

**Figure S64.** 10 µm MALDI IMS images of $[SM(41:1,2O)+H]^+$ or $[SM(18:1,2O/23:0)+H]^+$ (*m/z* 801.684) detected in <u>positive ion mode</u> from each donor sample. These molecules are isomers and therefore indistinguishable by MS alone. Multivariate SHAP analysis determined this to be feature 6 for classification of glomeruli.

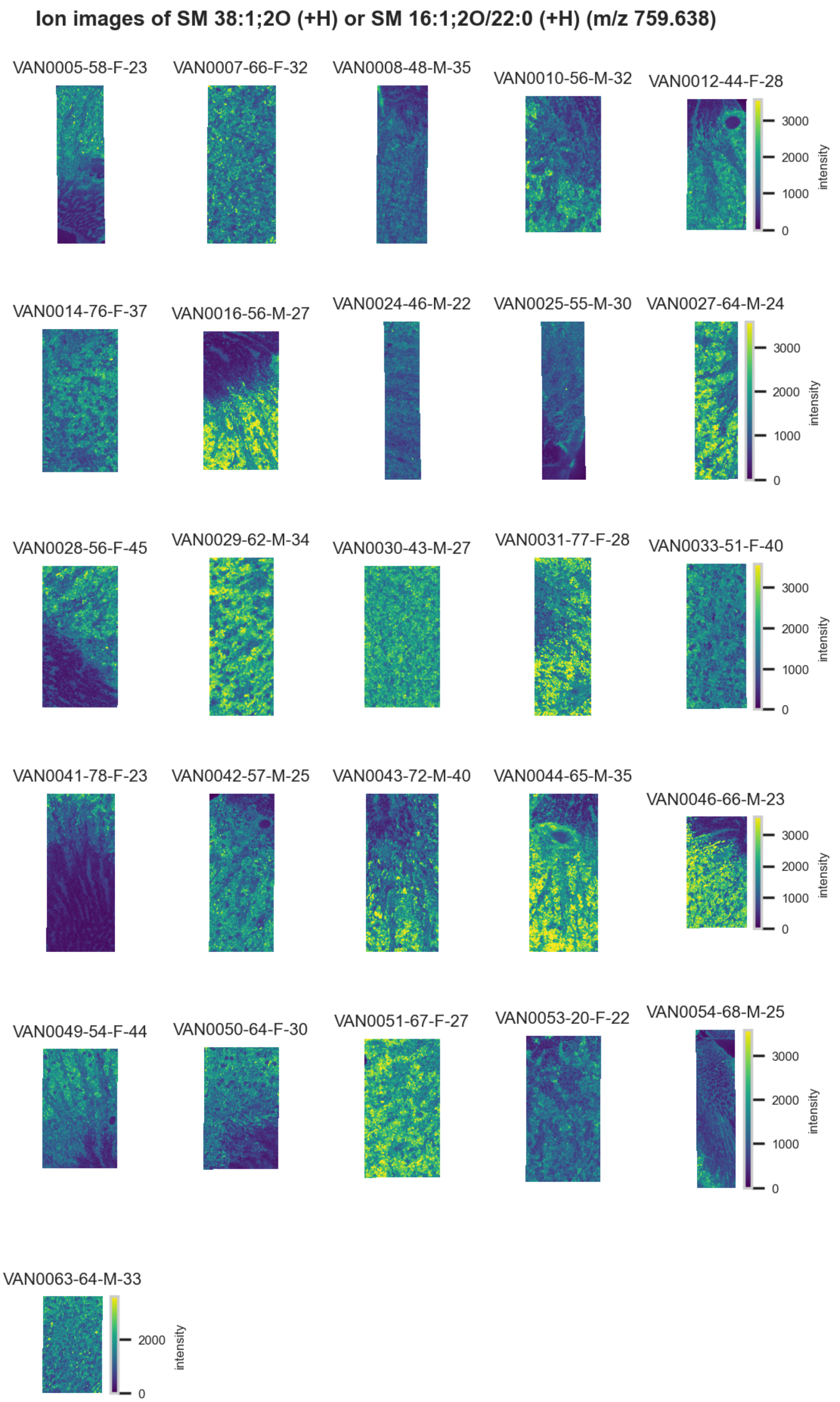

**Figure S65.** 10 µm MALDI IMS images of $[SM(38:1,2O)+H]^+$ or $[SM(16:1,2O/22:2)+H]^+$ (*m/z* 759.638) detected in <u>positive ion mode</u> from each donor sample. These molecules are isomers and therefore indistinguishable by MS alone. Multivariate SHAP analysis determined this to be feature 7 for classification of glomeruli.

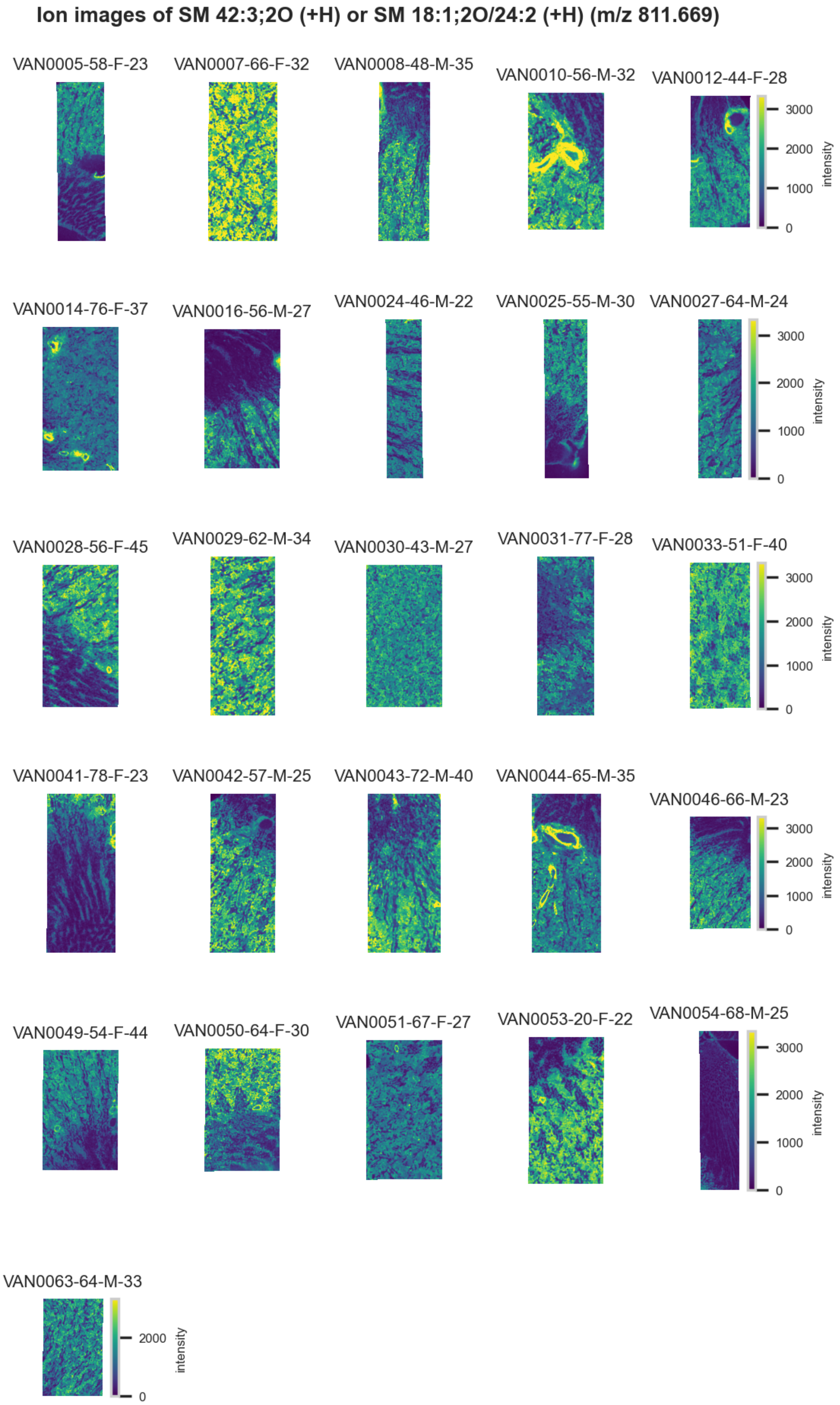

**Figure S66.** 10 μm MALDI IMS images of $[SM(42:3,2O)+H]^+$ or $[SM(18:1,2O/24:2)+H]^+$ (*m/z* 811.669) detected in positive ion mode from each donor sample. These molecules are isomers and therefore indistinguishable by MS alone. Multivariate SHAP analysis determined this to be feature 8 for classification of glomeruli.

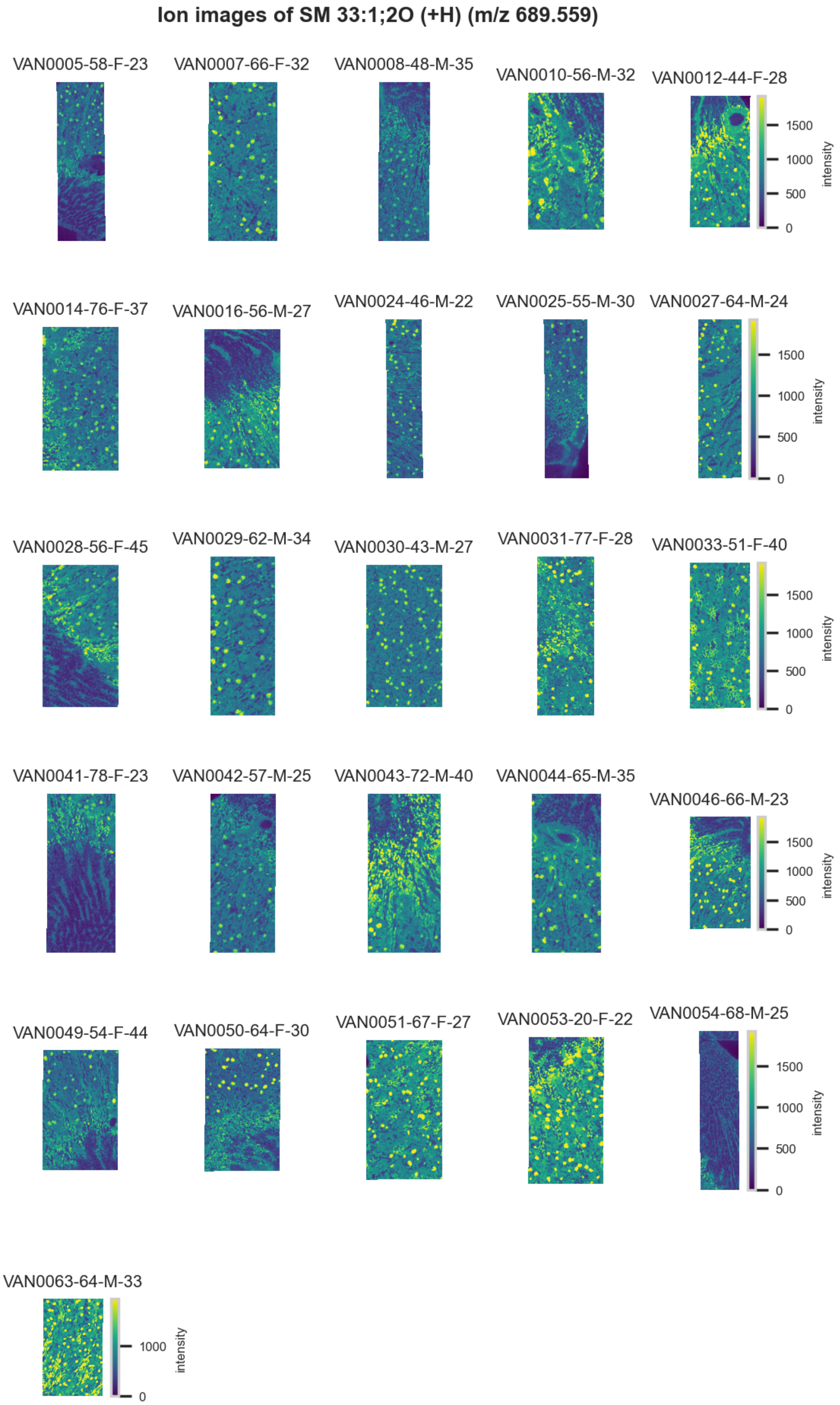

**Figure S67.** 10 µm MALDI IMS images of $[SM(33:1,2O)+H]^+$ (*m/z* 689.559) detected in positive ion mode from each donor sample. Multivariate SHAP analysis determined this to be feature 9 for classification of glomeruli.

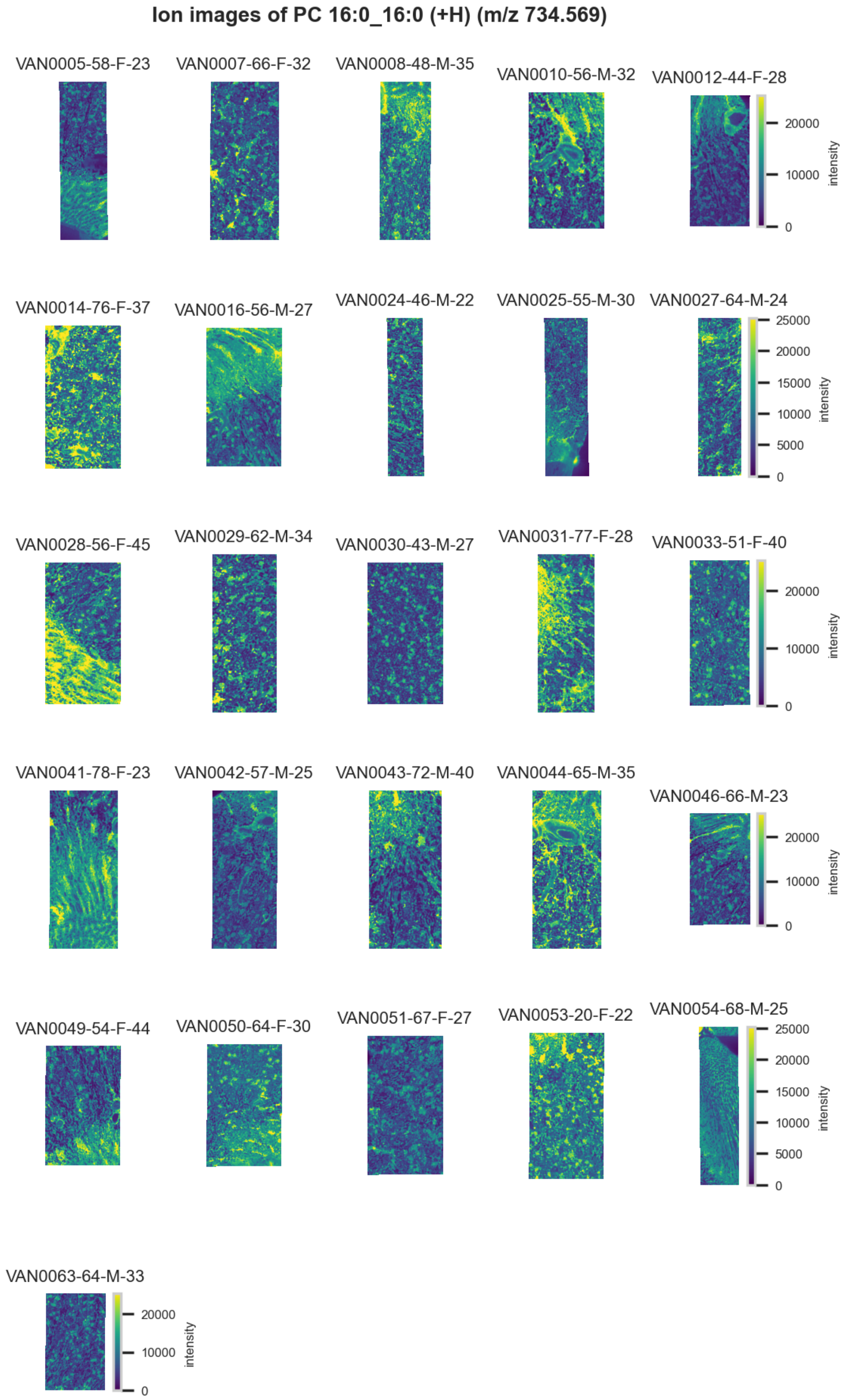

**Figure S68.** 10 µm MALDI IMS images of $[PC(16{:}0\_16{:}0)+H]^+$ (*m/z* 734.569) detected in positive ion mode from each donor sample. Multivariate SHAP analysis determined this to be feature 10 for classification of glomeruli.

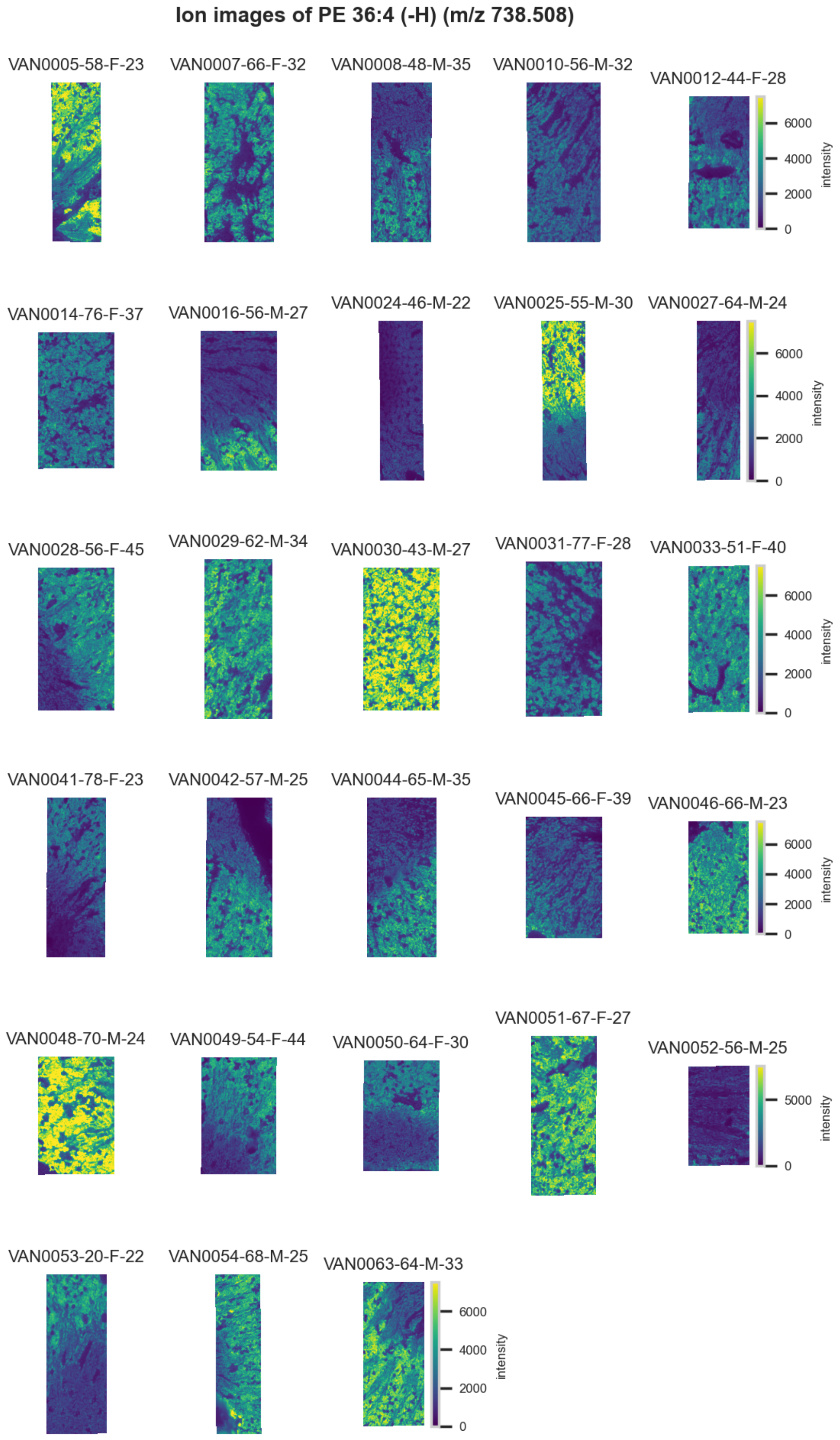

**Figure S69.** 10 µm MALDI IMS images of [PE(36:4)-H]$^-$ (*m/z* 738.508) detected in <u>negative ion mode</u> from each donor sample. Multivariate SHAP analysis determined this to be feature 1 for classification of proximal tubules.

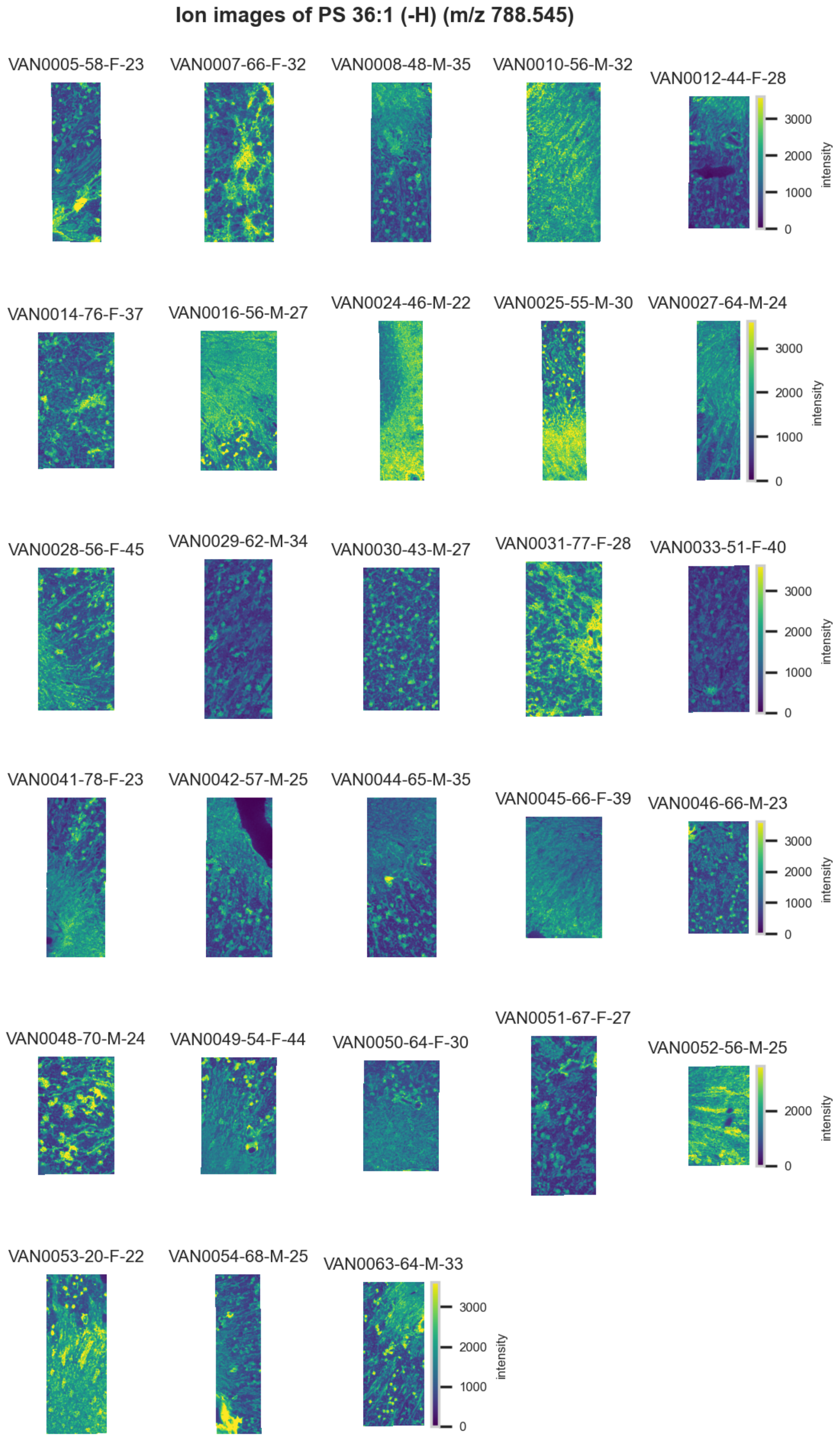

**Figure S70.** 10 µm MALDI IMS images of [PS(36:1)-H]$^-$ (*m/z* 788.545) detected in negative ion mode from each donor sample. Multivariate SHAP analysis determined this to be feature 2 for classification of proximal tubules.

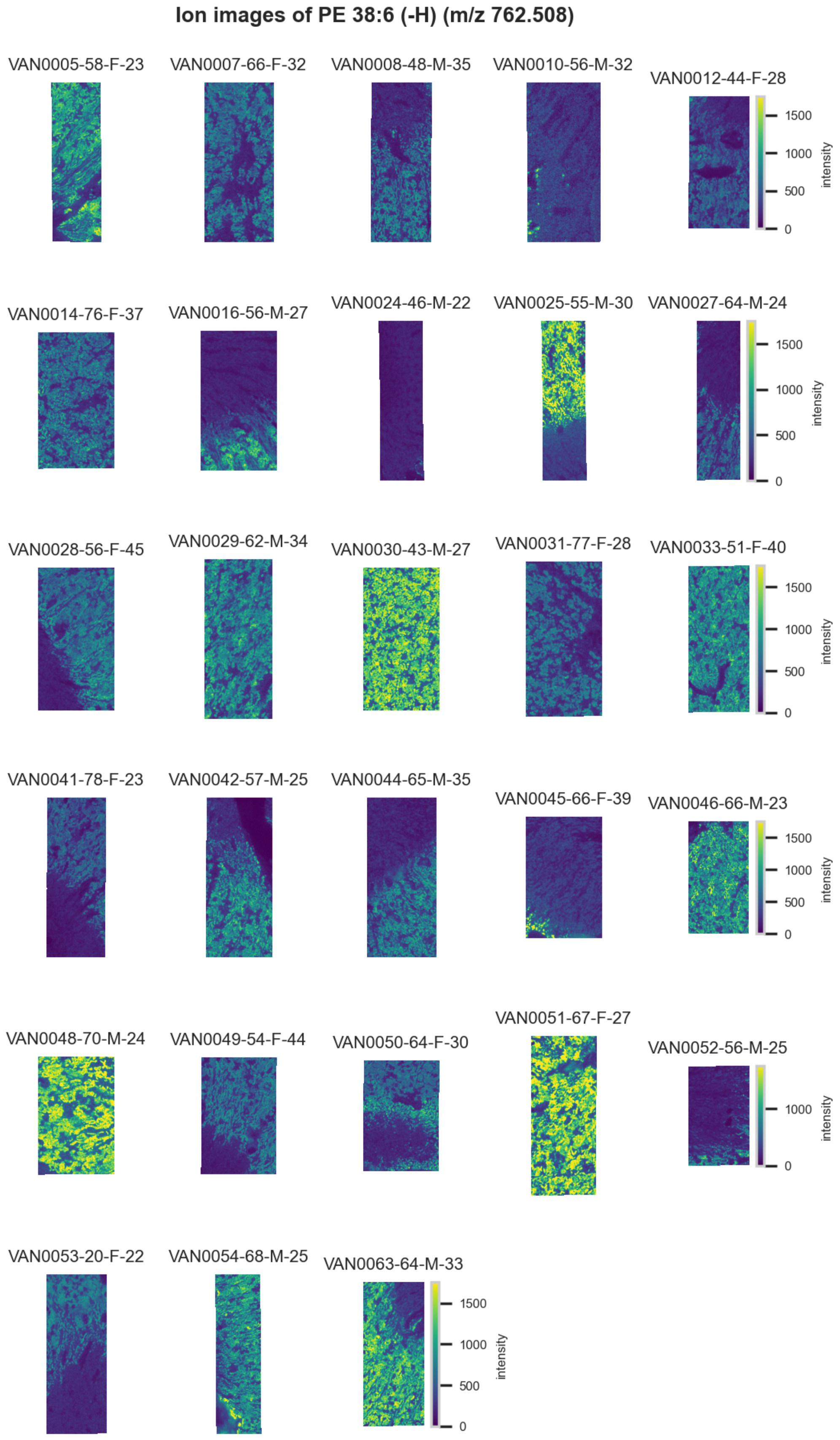

**Figure S71.** 10 µm MALDI IMS images of [PE(38:6)-H]$^-$ (*m/z* 762.508) detected in negative ion mode from each donor sample. Multivariate SHAP analysis determined this to be feature 3 for classification of proximal tubules.

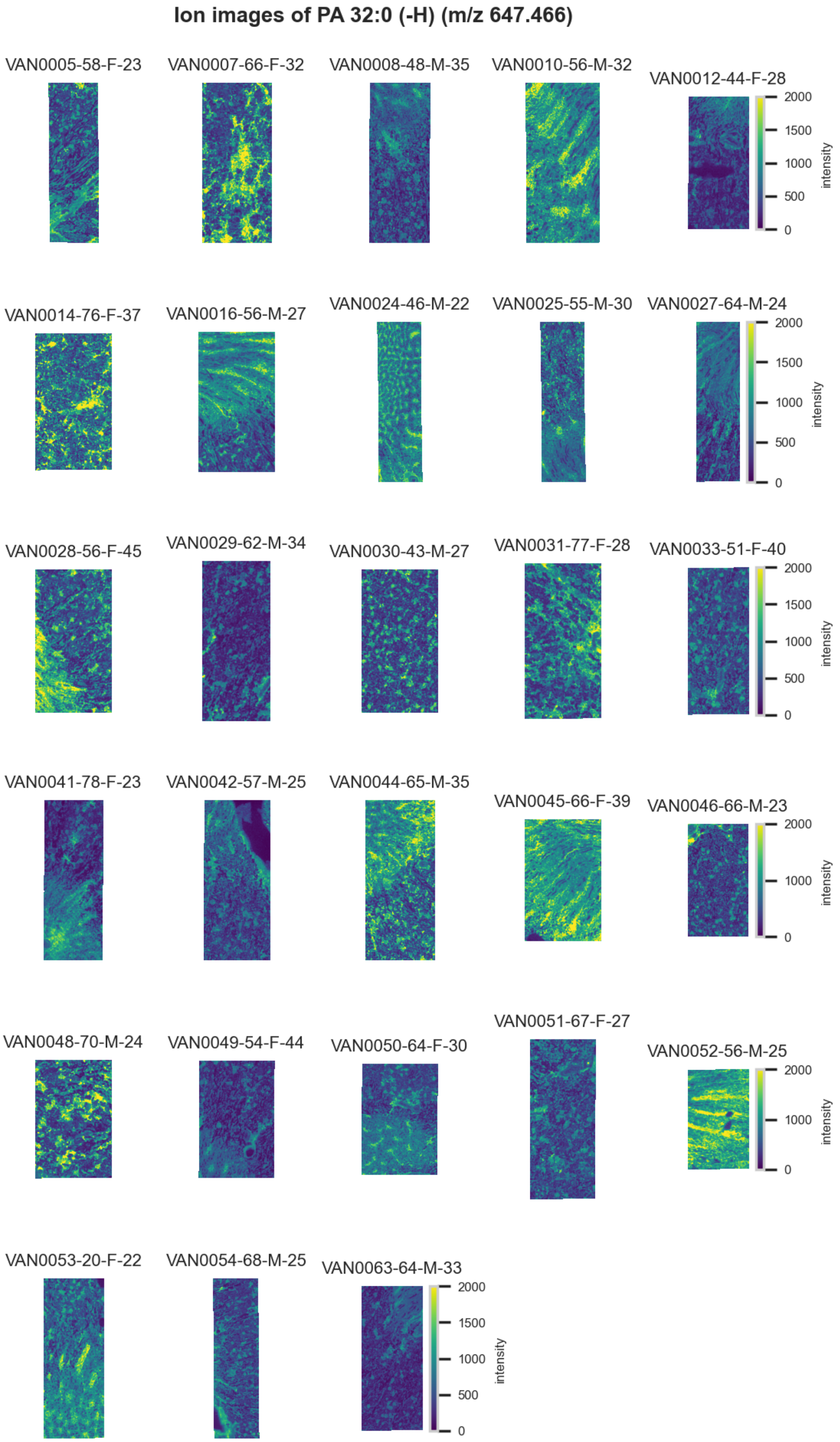

**Figure S72.** 10 µm MALDI IMS images of [PA(32:0)-H]$^-$ (*m/z* 647.466) detected in <u>negative ion mode</u> from each donor sample. Multivariate SHAP analysis determined this to be feature 4 for classification of proximal tubules.

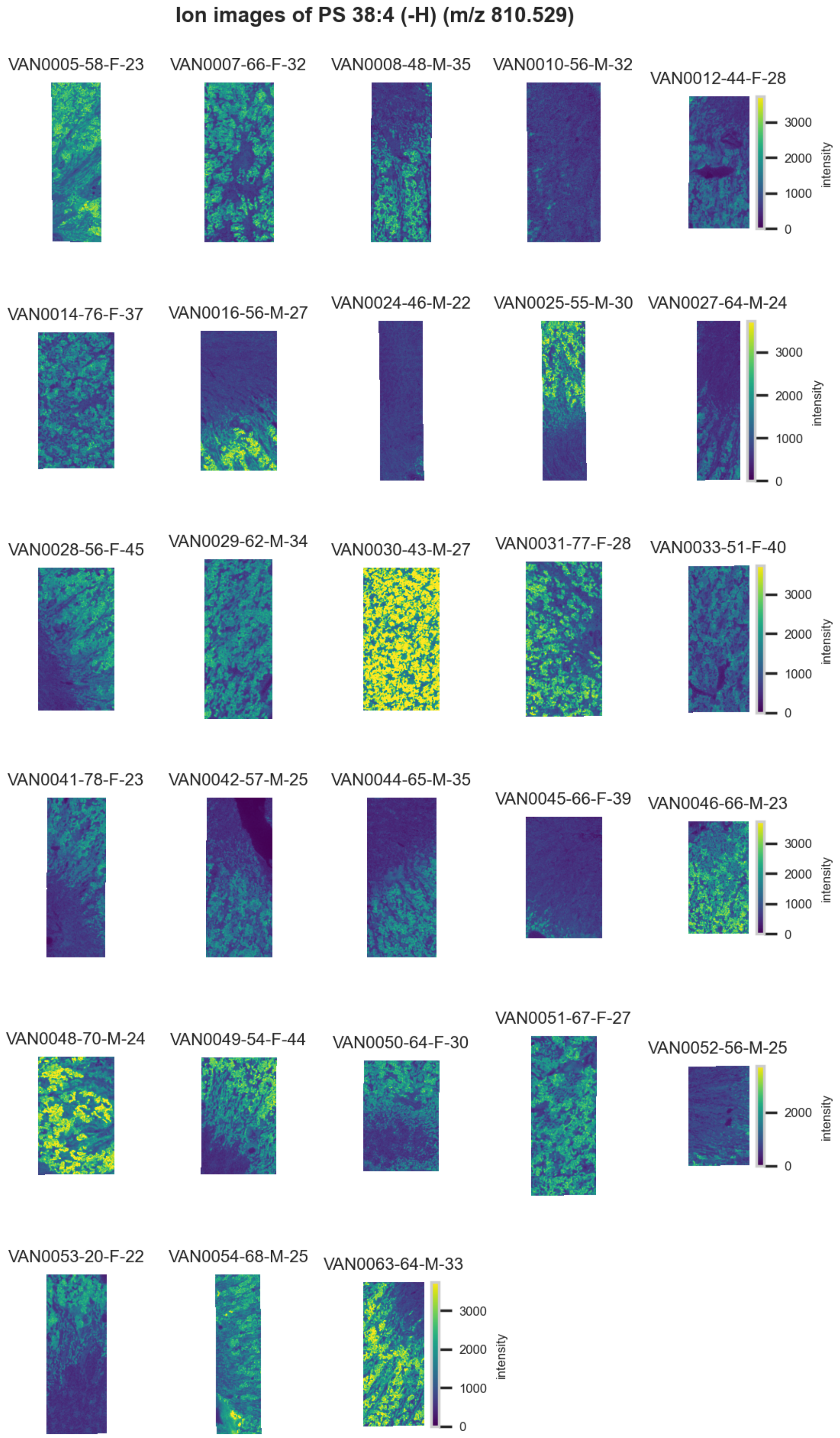

**Figure S73.** 10 µm MALDI IMS images of [PS(38:4)-H]$^-$ (*m/z* 810.529) detected in <u>negative ion mode</u> from each donor sample. Multivariate SHAP analysis determined this to be feature 5 for classification of proximal tubules.

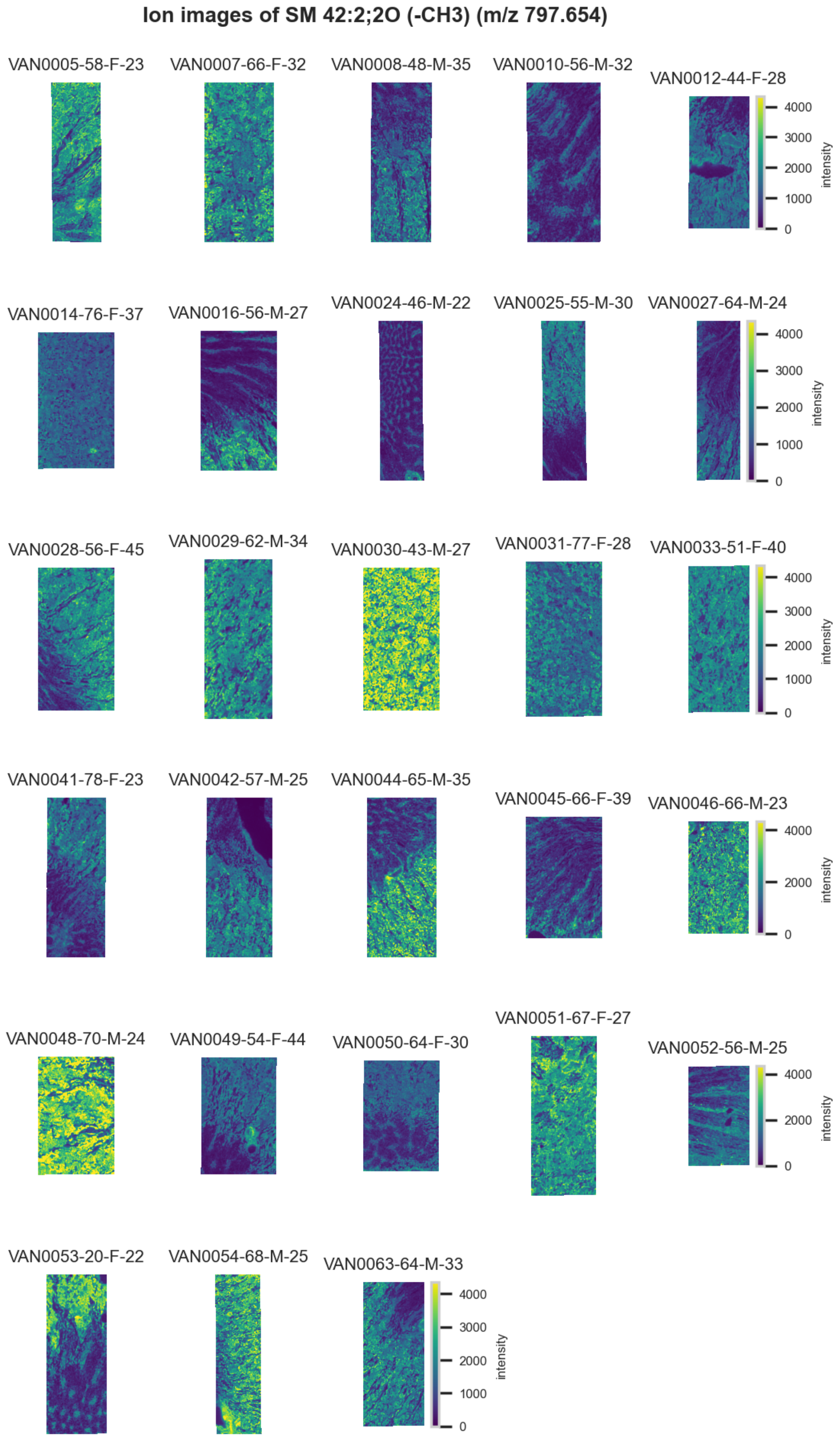

**Figure S74.** 10 μm MALDI IMS images of $[SM(42{:}2{,}2O){-}CH_3]^-$ (*m/z* 797.654) detected in negative ion mode from each donor sample. Multivariate SHAP analysis determined this to be feature 6 for classification of proximal tubules.

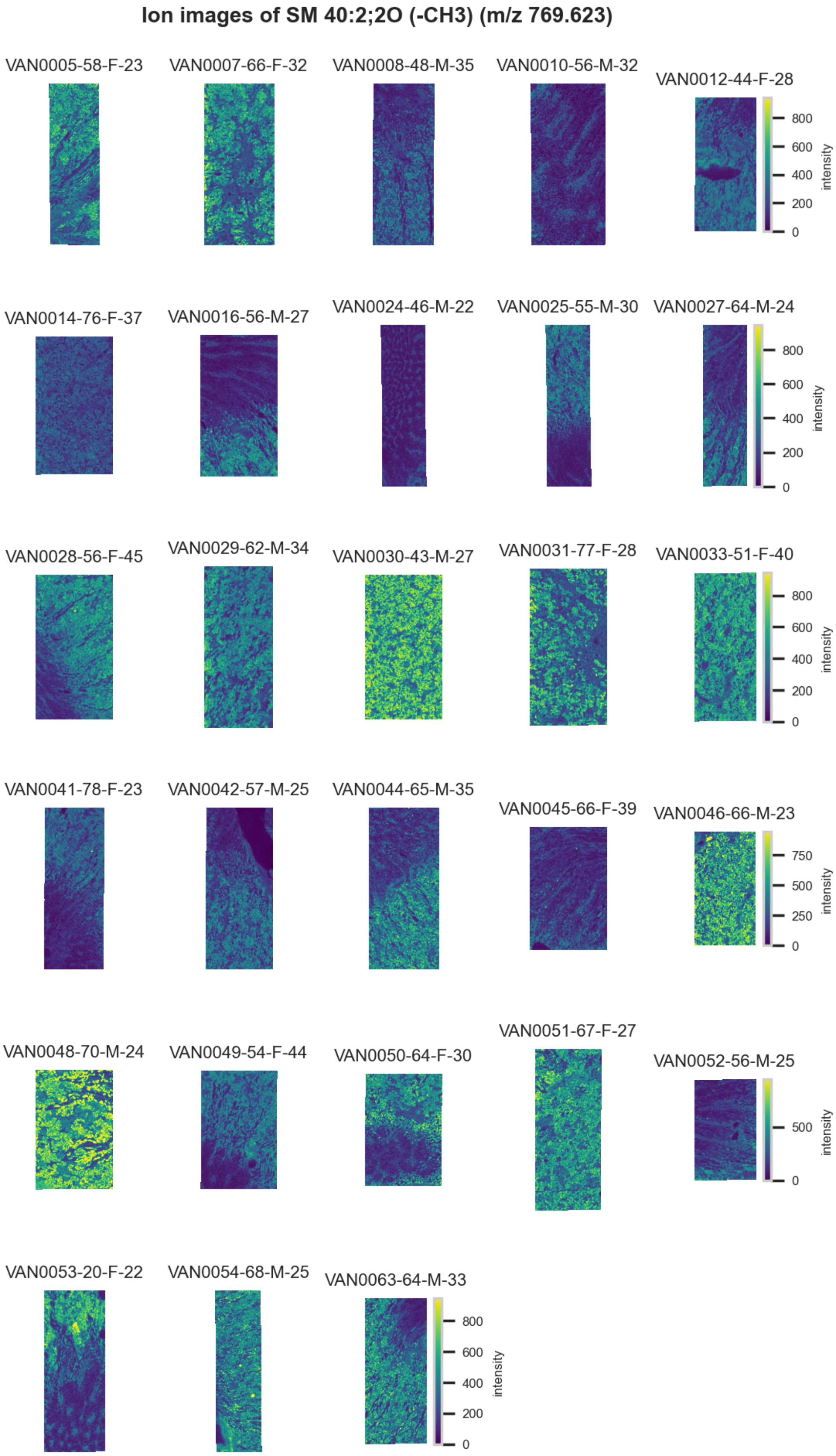

**Figure S75.** 10 µm MALDI IMS images of $[SM(40:2,2O)\text{-}CH_3]^-$ (*m/z* 769.623) detected in negative ion mode from each donor sample. Multivariate SHAP analysis determined this to be feature 7 for classification of proximal tubules.

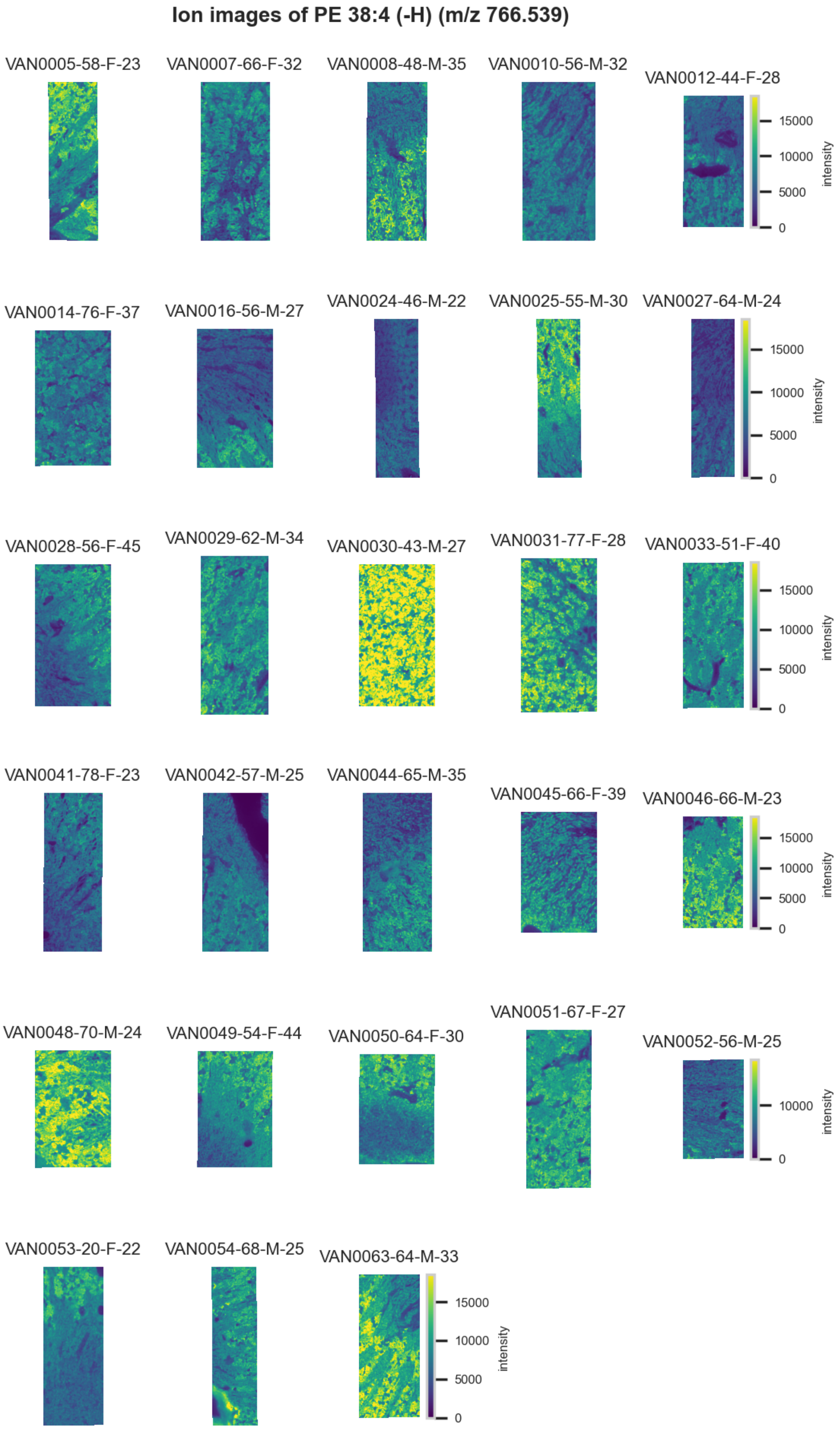

**Figure S76.** 10 µm MALDI IMS images of [PE(38:4)-H]$^-$ (*m/z* 766.539) detected in <u>negative ion mode</u> from each donor sample. Multivariate SHAP analysis determined this to be feature 8 for classification of proximal tubules.

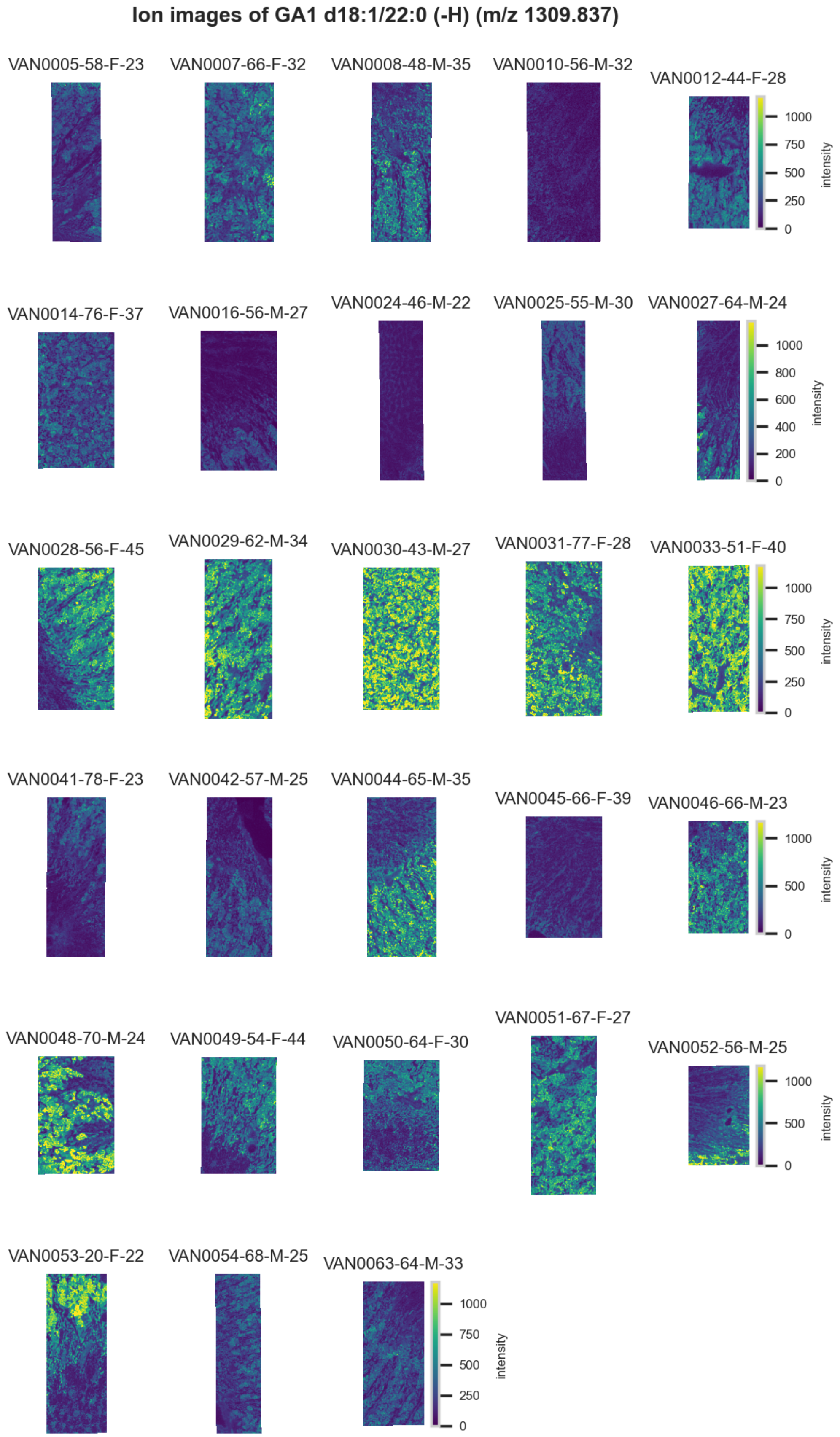

**Figure S77.** 10 µm MALDI IMS images of [GA1(d18:1/22:0)-H]$^{-}$ (*m/z* 1,309.837) detected in negative ion mode from each donor sample. Multivariate SHAP analysis determined this to be feature 9 for classification of proximal tubules.

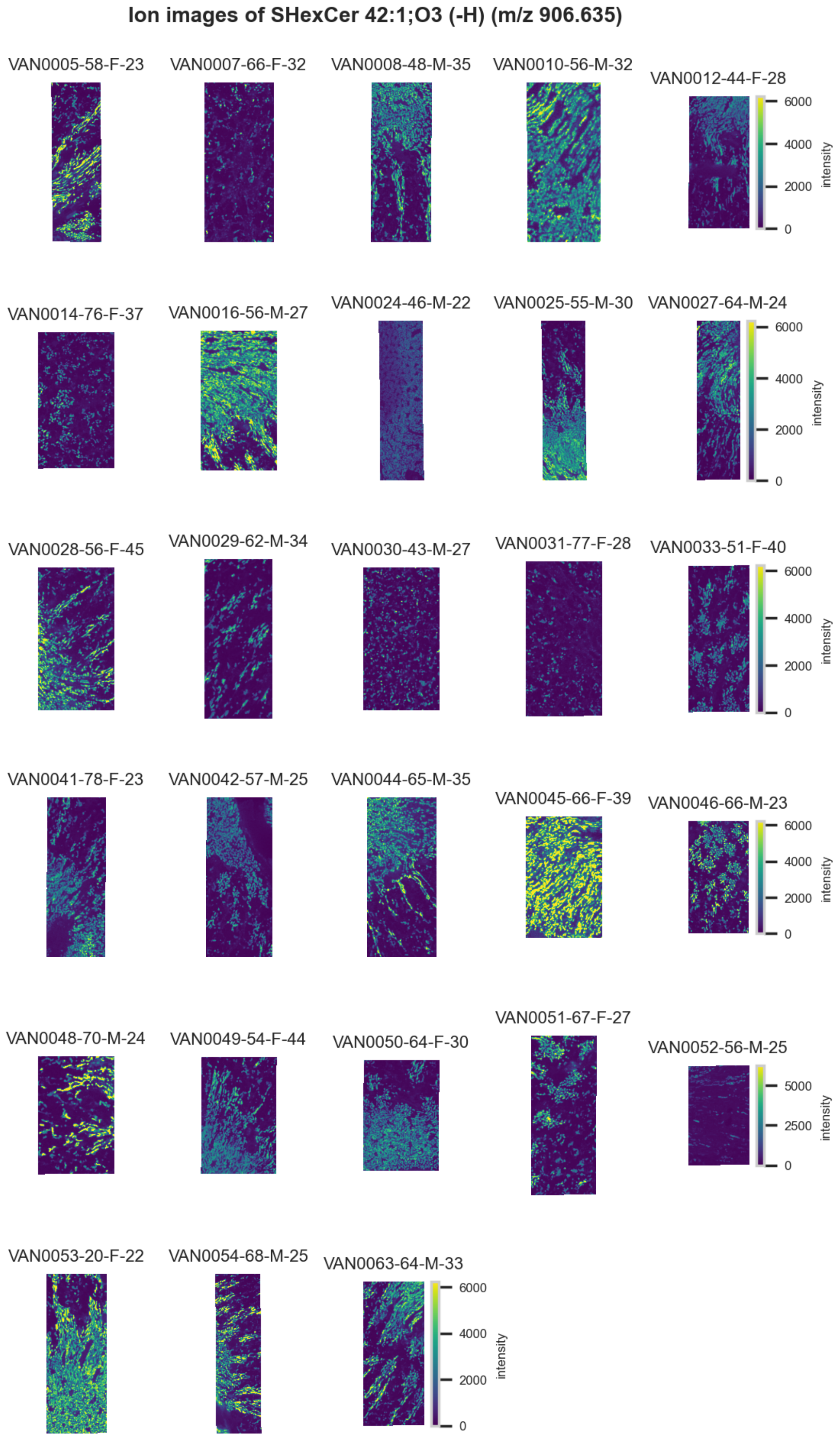

**Figure S78.** 10 μm MALDI IMS images of [SHexCer(42:1,3O)-H]$^-$ (*m/z* 906.635) detected in <u>negative ion mode</u> from each donor sample. Multivariate SHAP analysis determined this to be feature 10 for classification of proximal tubules.

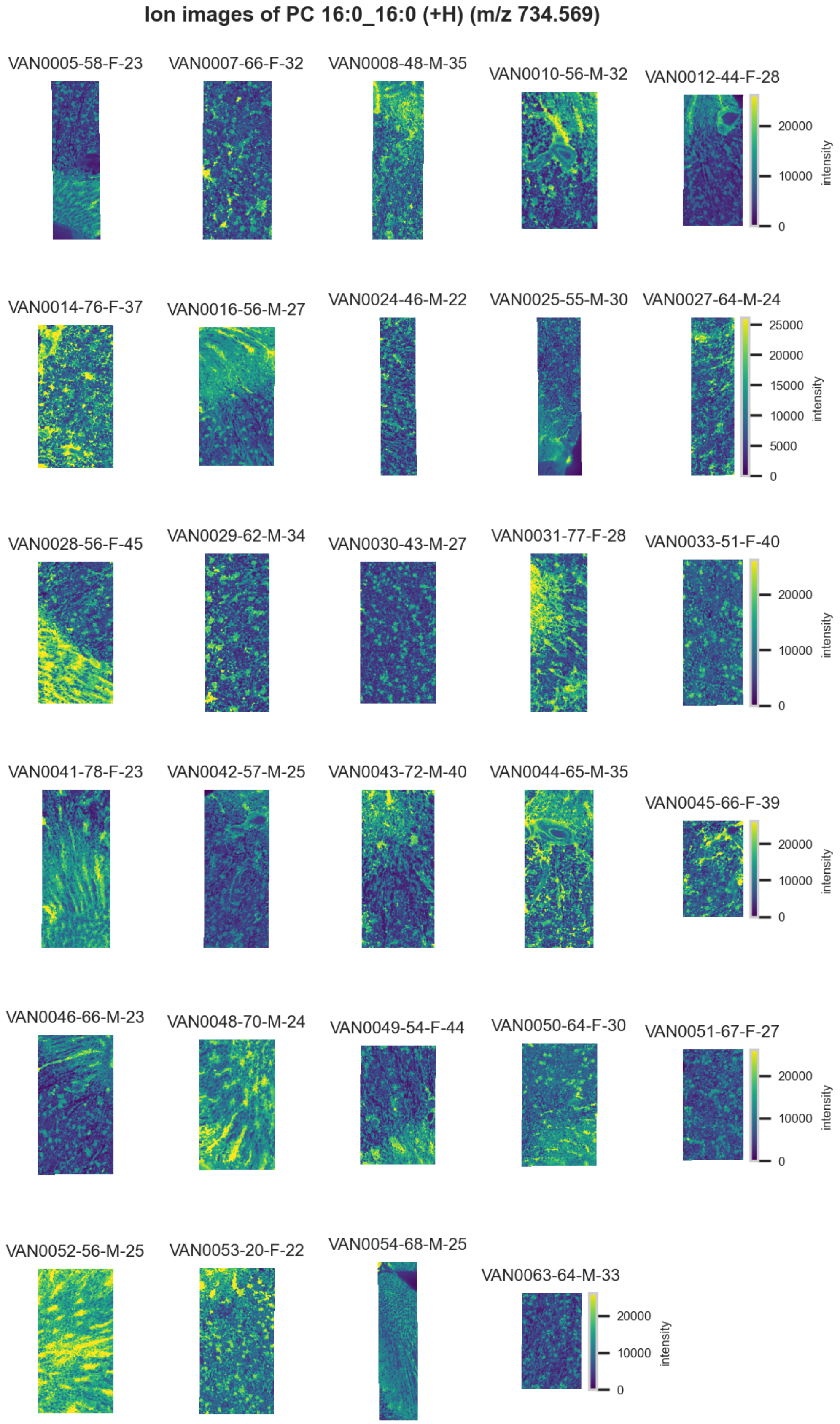

**Figure S79.** 10 µm MALDI IMS images of $[PC(16:0\_16:0)+H]^+$ (*m/z* 734.569) detected in positive ion mode from each donor sample. Multivariate SHAP analysis determined this to be feature 1 for classification of proximal tubules.

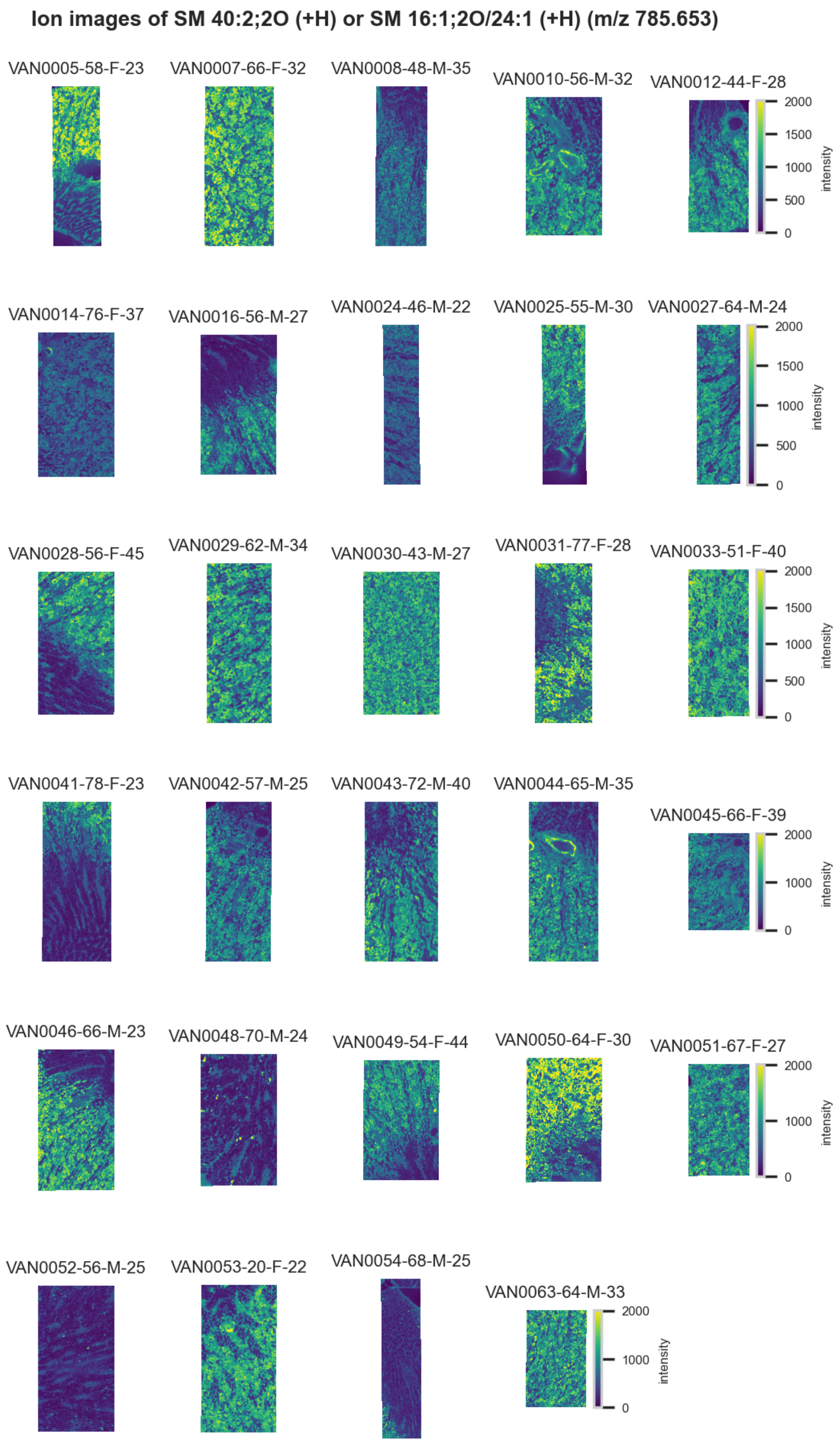

**Figure S80.** 10 µm MALDI IMS images of $[SM(40:2,2O)+H]^+$ or $[SM(16:1,2O/24:1)+H]^+$ (*m/z* 785.653) detected in positive ion mode from each donor sample. These molecules are isomers and therefore indistinguishable by MS alone. Multivariate SHAP analysis determined this to be feature 2 for classification of proximal tubules.

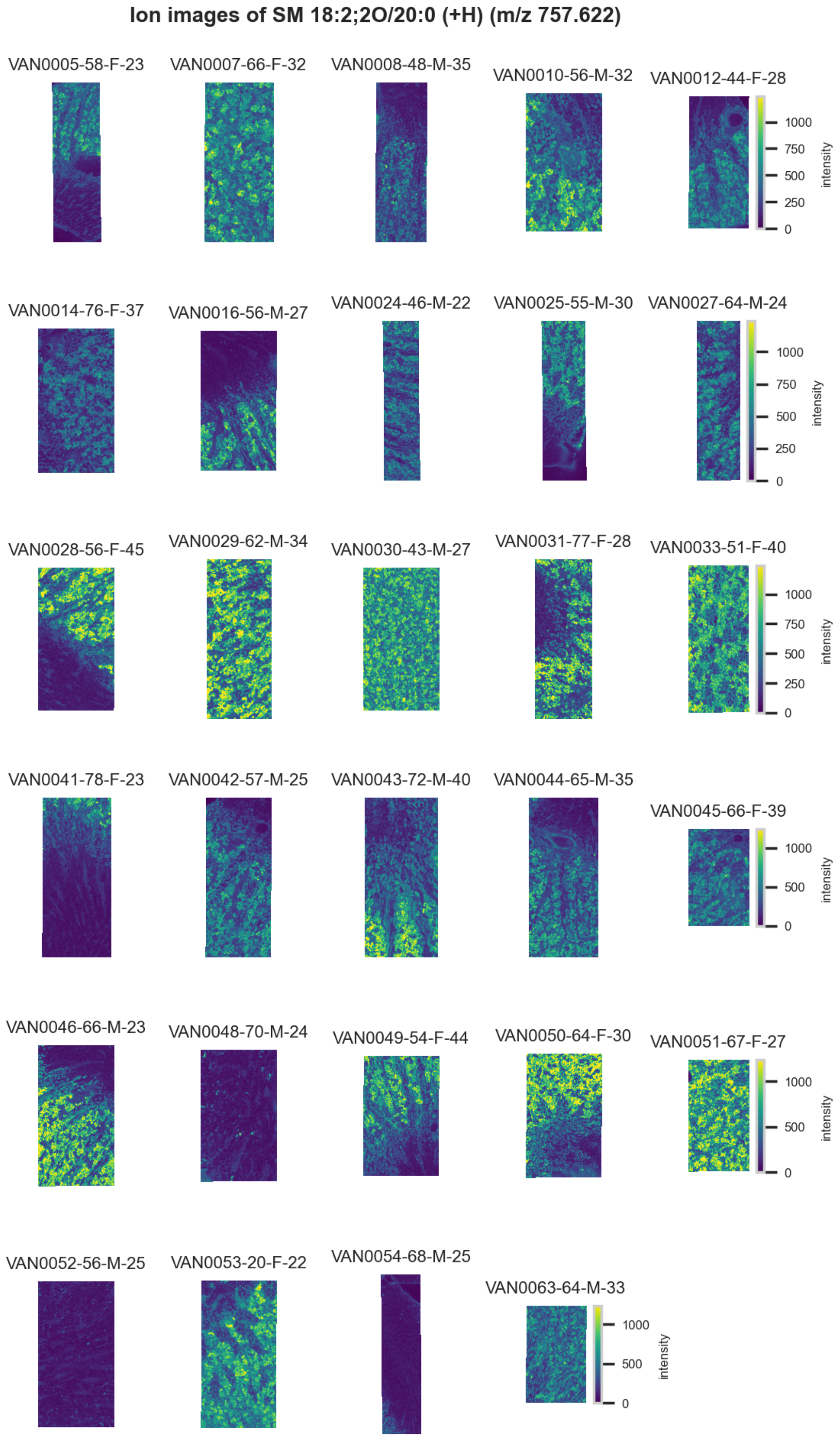

**Figure S81.** 10 µm MALDI IMS images of $[SM(18:2,2O/20:0)+H]^+$ (*m/z* 757.622) detected in positive ion mode from each donor sample. Multivariate SHAP analysis determined this to be feature 3 for classification of proximal tubules.

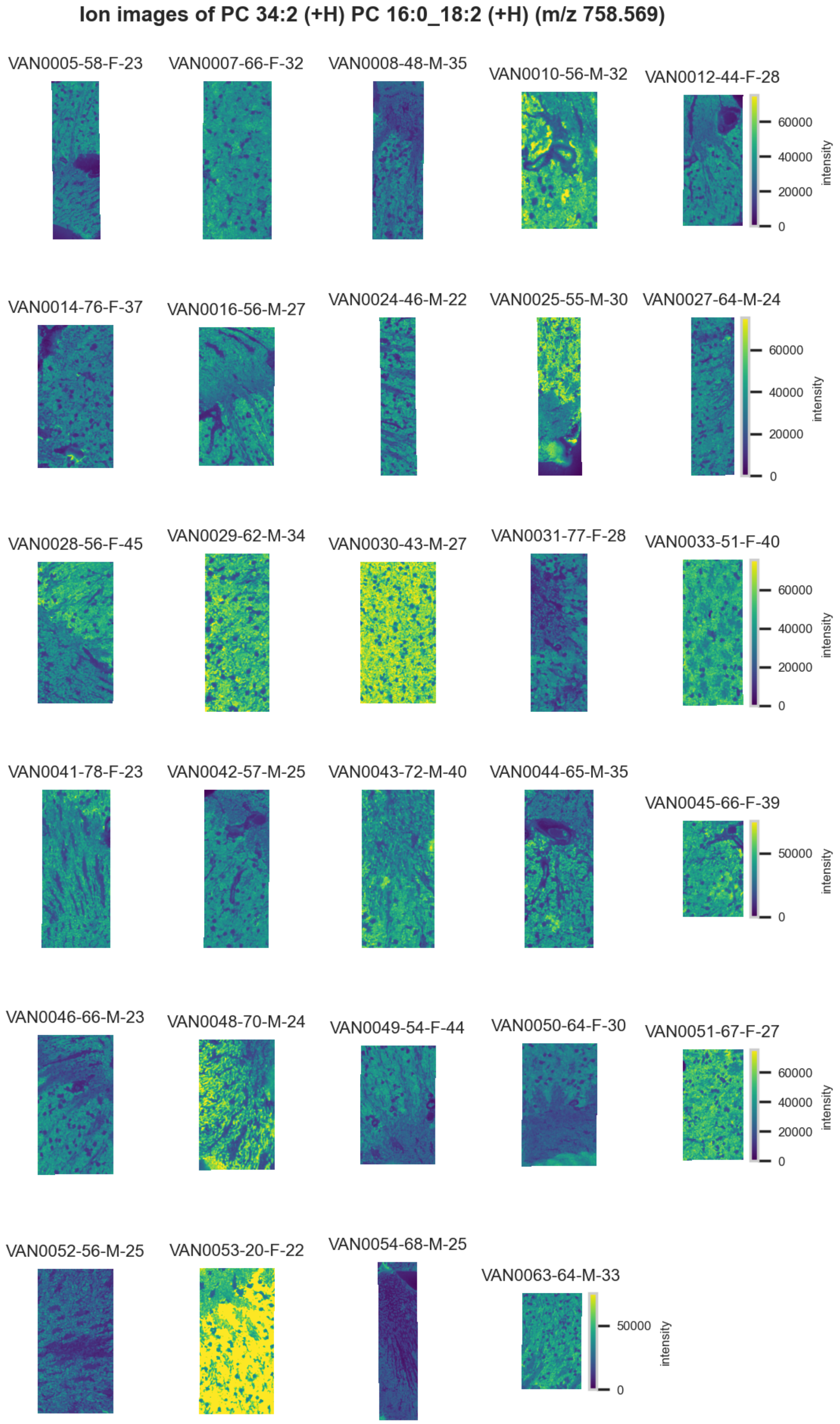

**Figure S82.** 10 µm MALDI IMS images of [PC(16:0_18:2)+H]$^+$ (*m/z* 758.569) detected in <u>positive ion mode</u> from each donor sample. Multivariate SHAP analysis determined this to be feature 4 for classification of proximal tubules.

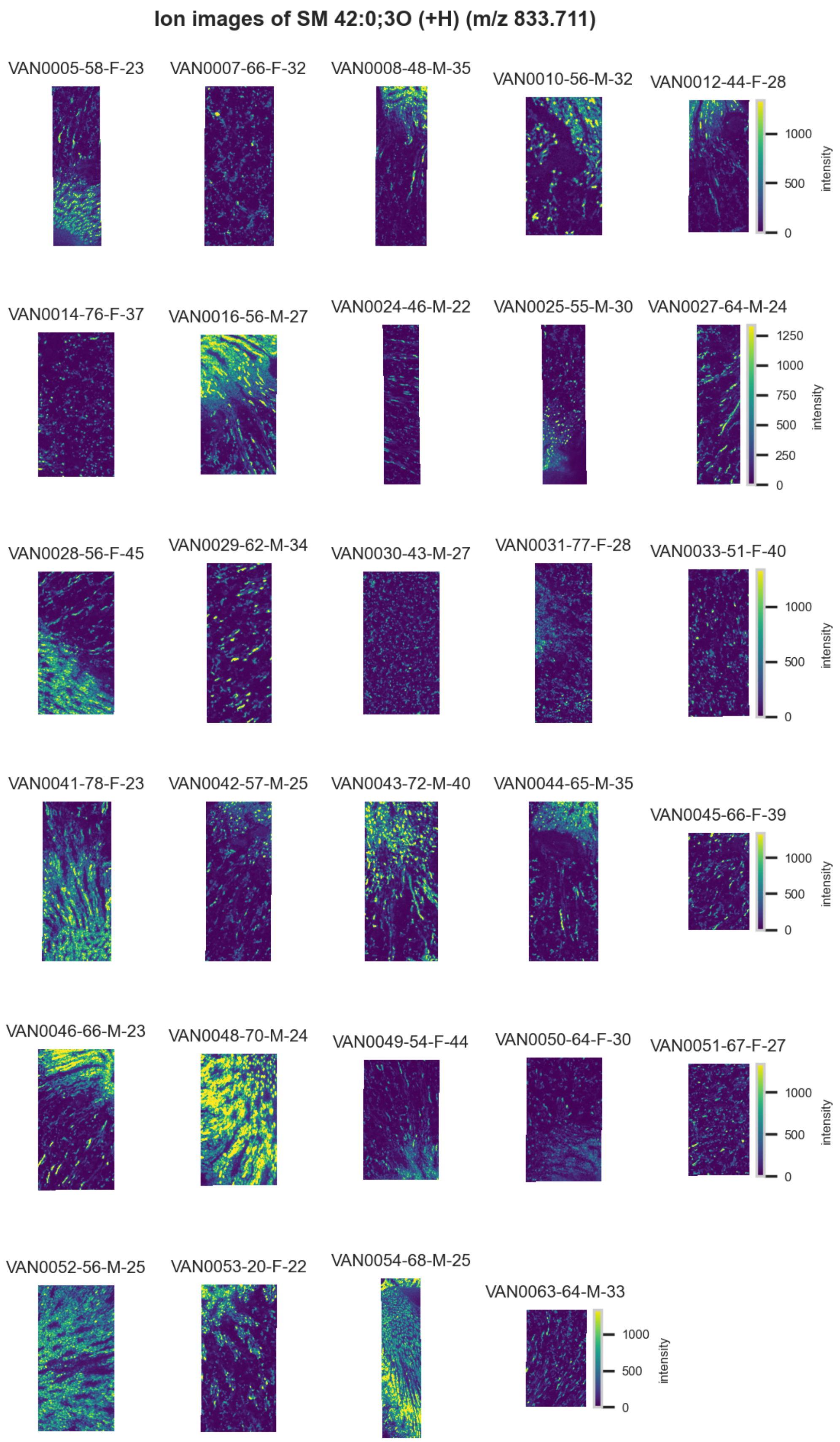

**Figure S83.** 10 µm MALDI IMS images of $[SM(42:0,3O)+H]^+$ (*m/z* 833.711) detected in positive ion mode from each donor sample. Multivariate SHAP analysis determined this to be feature 5 for classification of proximal tubules.

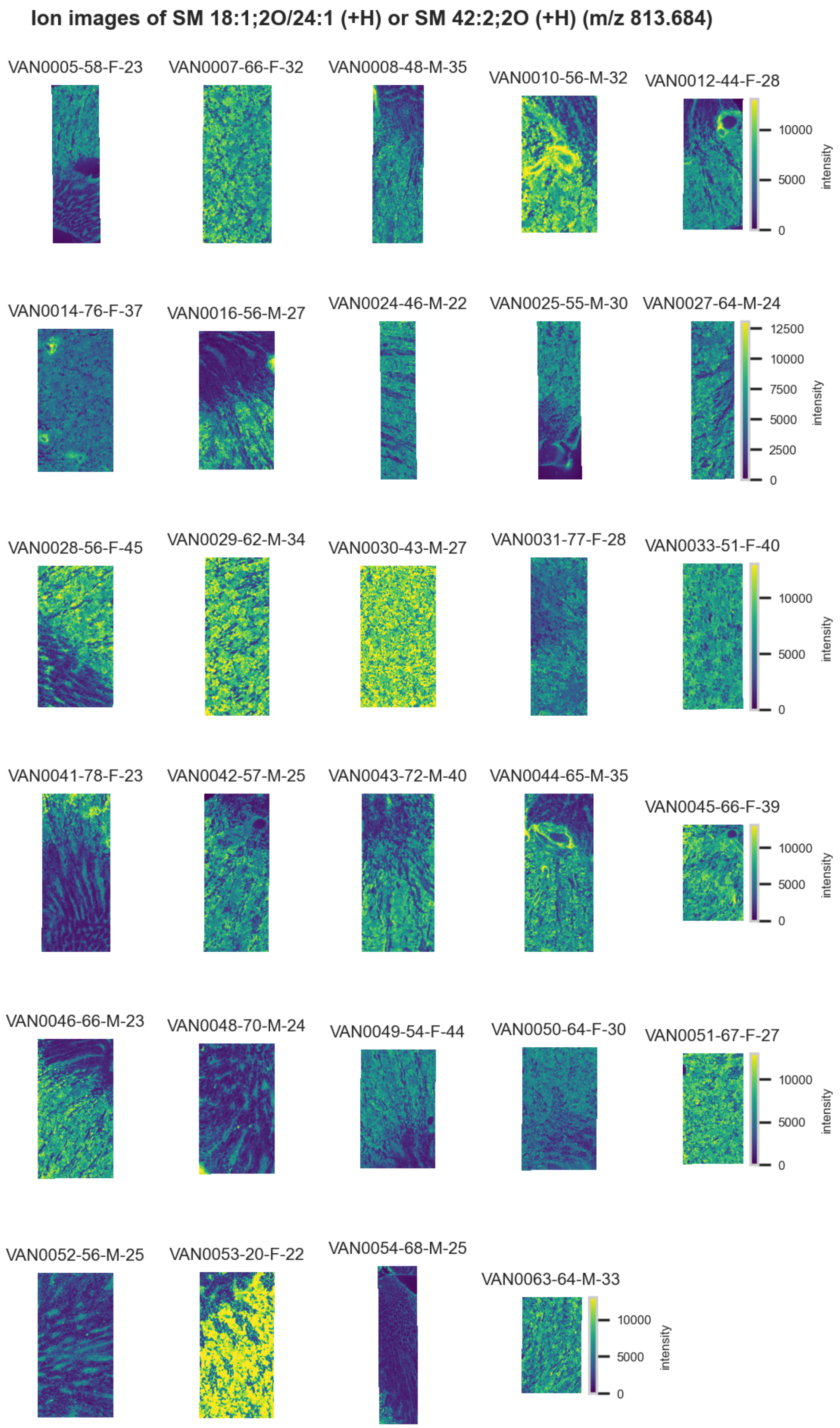

**Figure S84.** 10 µm MALDI IMS images of [SM(42:2,2O)+H]$^+$ or [SM(18:1,2O/24:1)+H]$^+$ (*m/z* 813.684) detected in positive ion mode from each donor sample. These molecules are isomers and therefore indistinguishable by MS alone. Multivariate SHAP analysis determined this to be feature 6 for classification of proximal tubules.

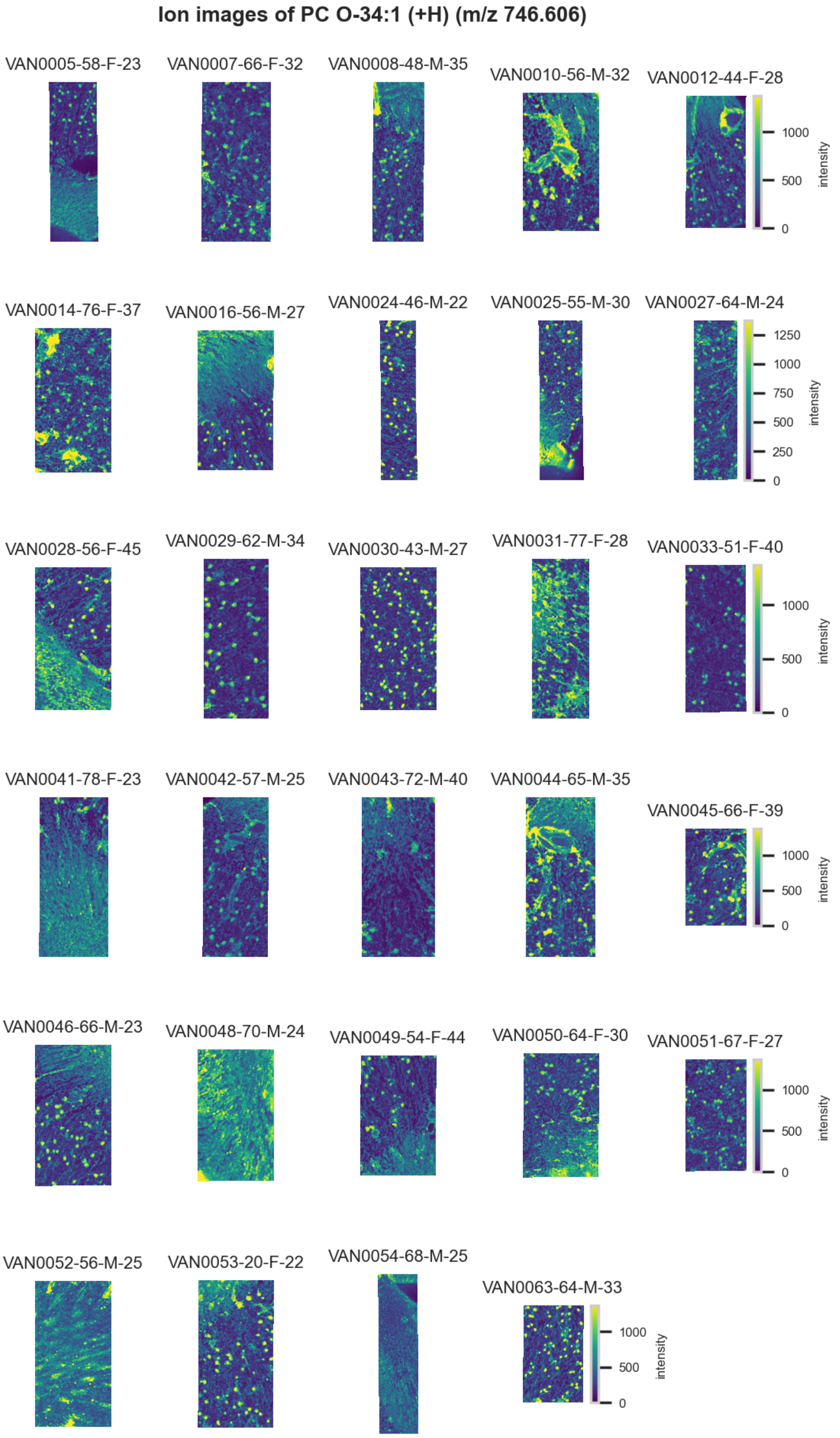

**Figure S85.** 10 µm MALDI IMS images of [PC(O-34:1)+H]$^+$ (*m/z* 746.606) detected in positive ion mode from each donor sample. Multivariate SHAP analysis determined this to be feature 7 for classification of proximal tubules.

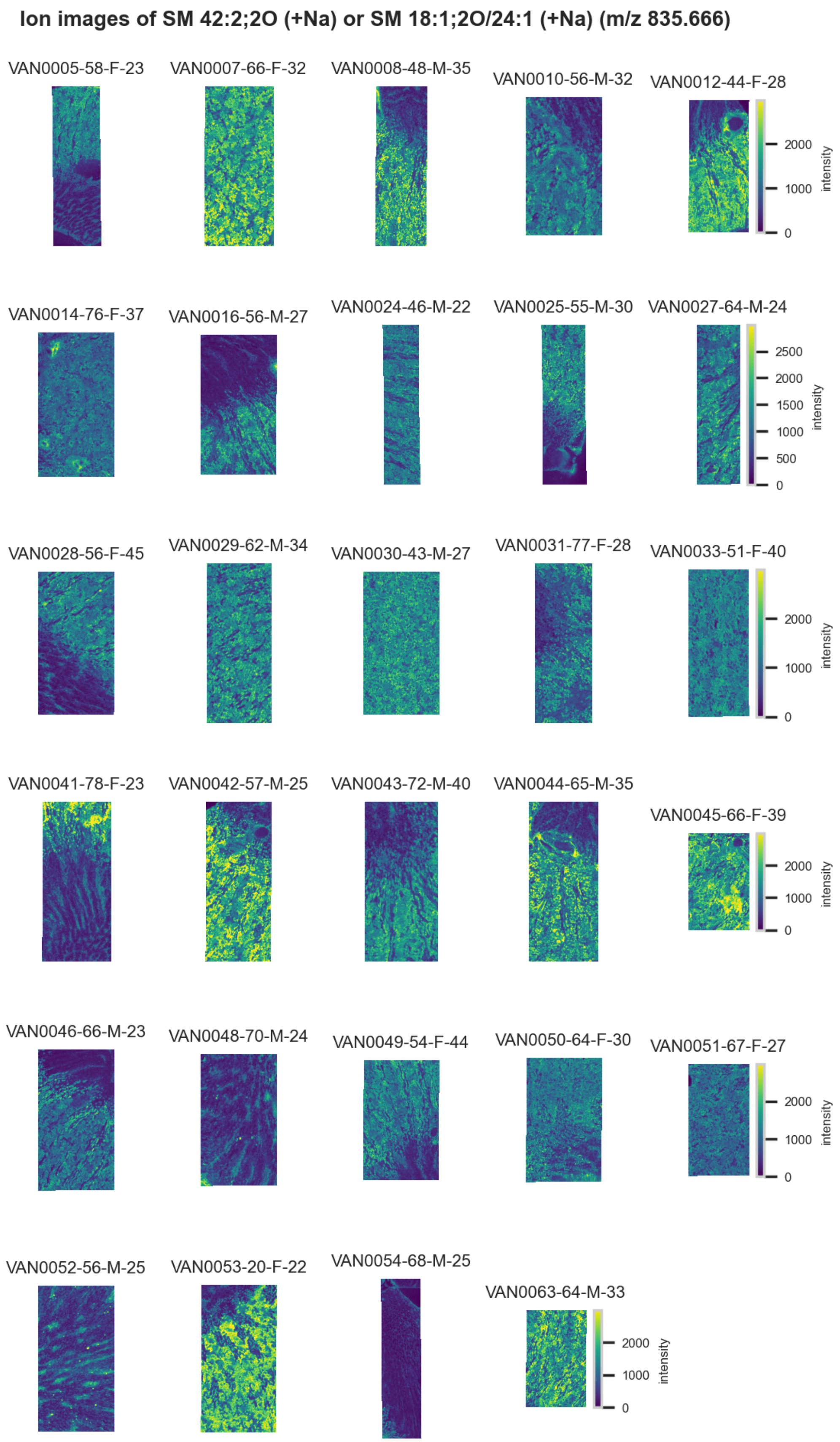

**Figure S86.** 10 µm MALDI IMS images of $[SM(42:2,2O)+Na]^+$ or $[SM(18:1,2O/24:1)+Na]^+$ (*m/z* 835.666) detected in <u>positive ion mode</u> from each donor sample. These molecules are isomers and therefore indistinguishable by MS alone. Multivariate SHAP analysis determined this to be feature 8 for classification of proximal tubules.

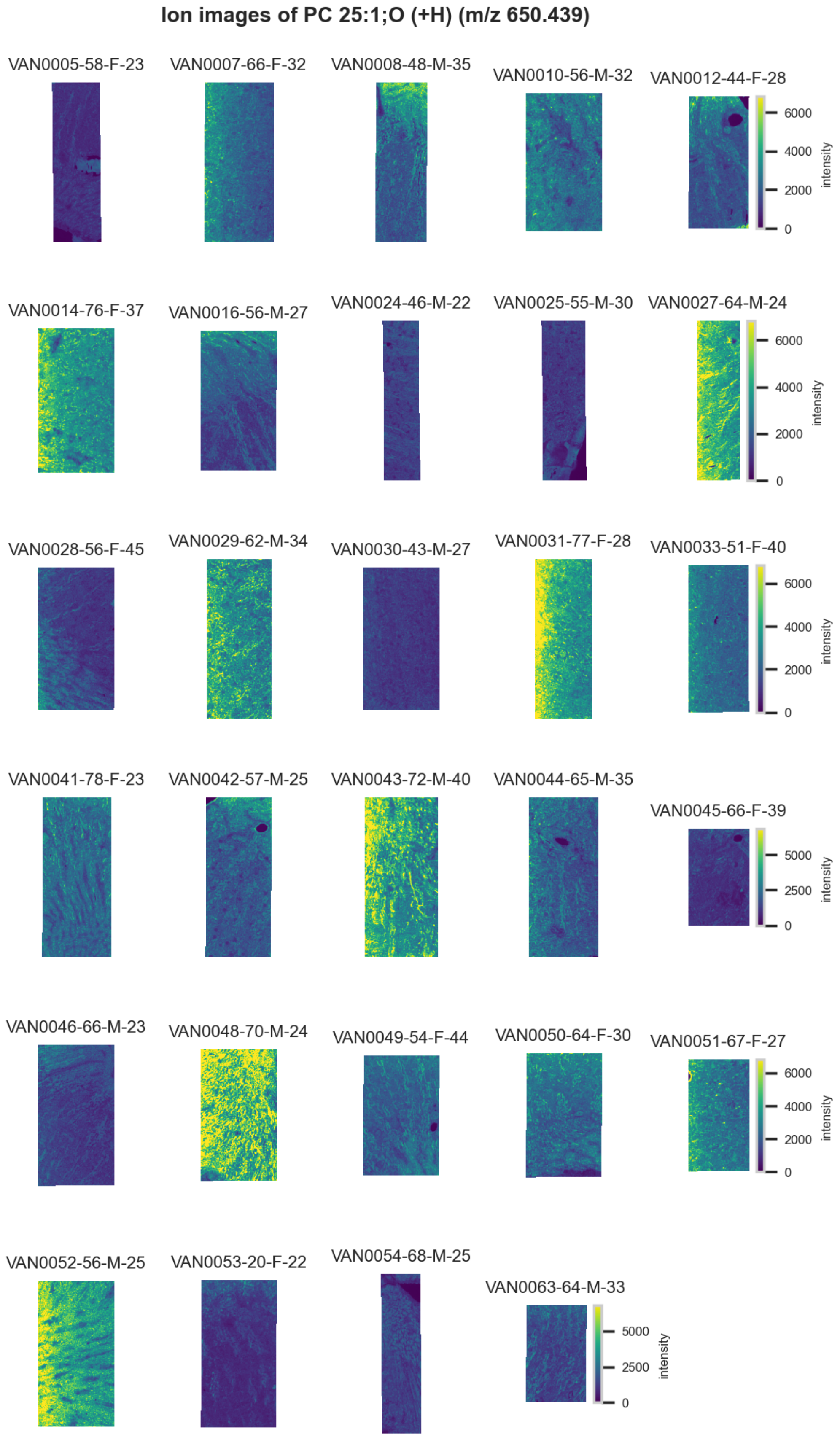

**Figure S87.** 10 µm MALDI IMS images of $[PC(25:1,O)+H]^{+}$ (*m/z* 650.439) detected in <u>positive ion mode</u> from each donor sample. Multivariate SHAP analysis determined this to be feature 9 for classification of proximal tubules.

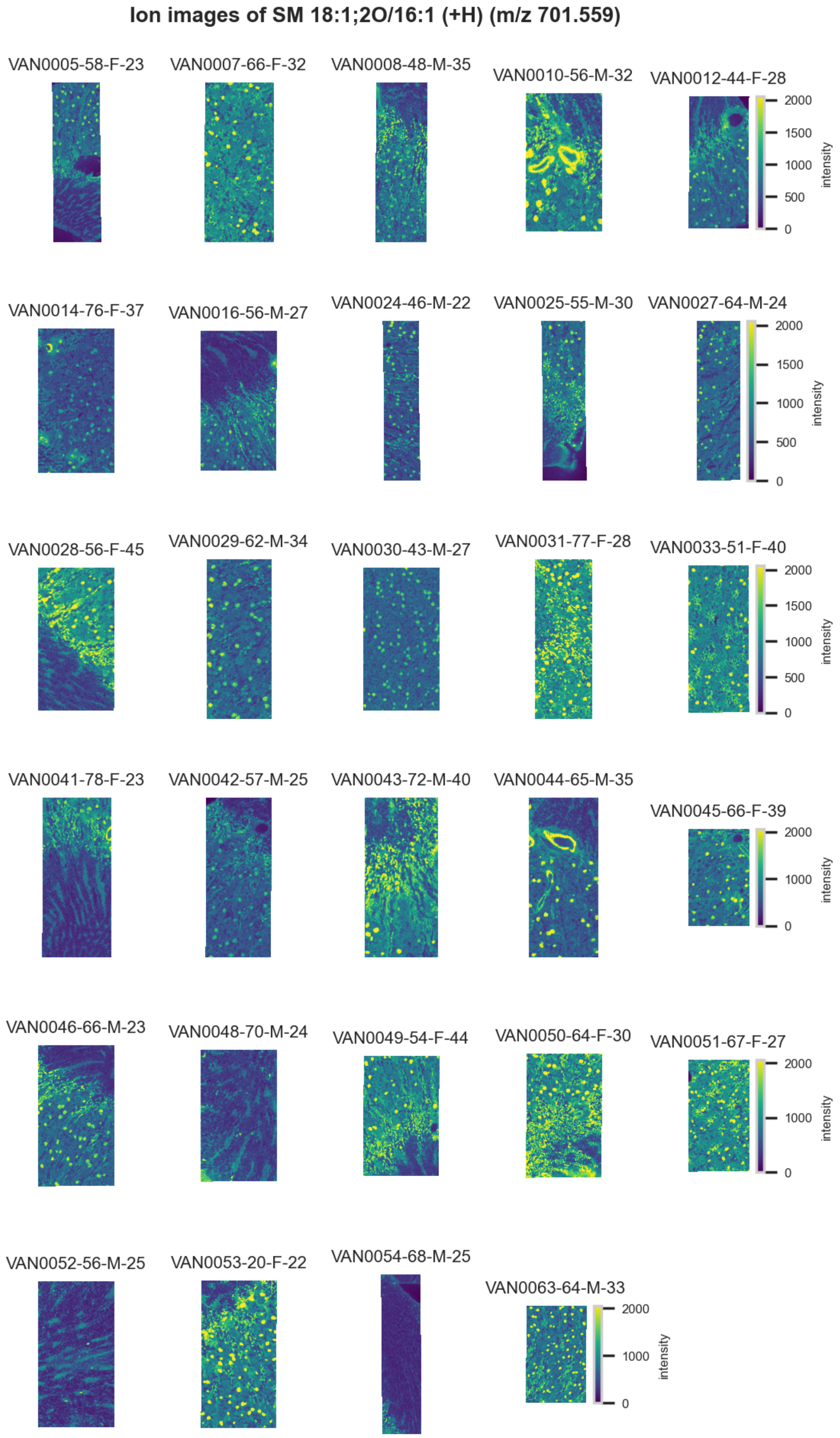

**Figure S88.** 10 µm MALDI IMS images of $[SM(18:1,2O/16:1)+H]^+$ (*m/z* 701.559) detected in positive ion mode from each donor sample. Multivariate SHAP analysis determined this to be feature 10 for classification of proximal tubules.

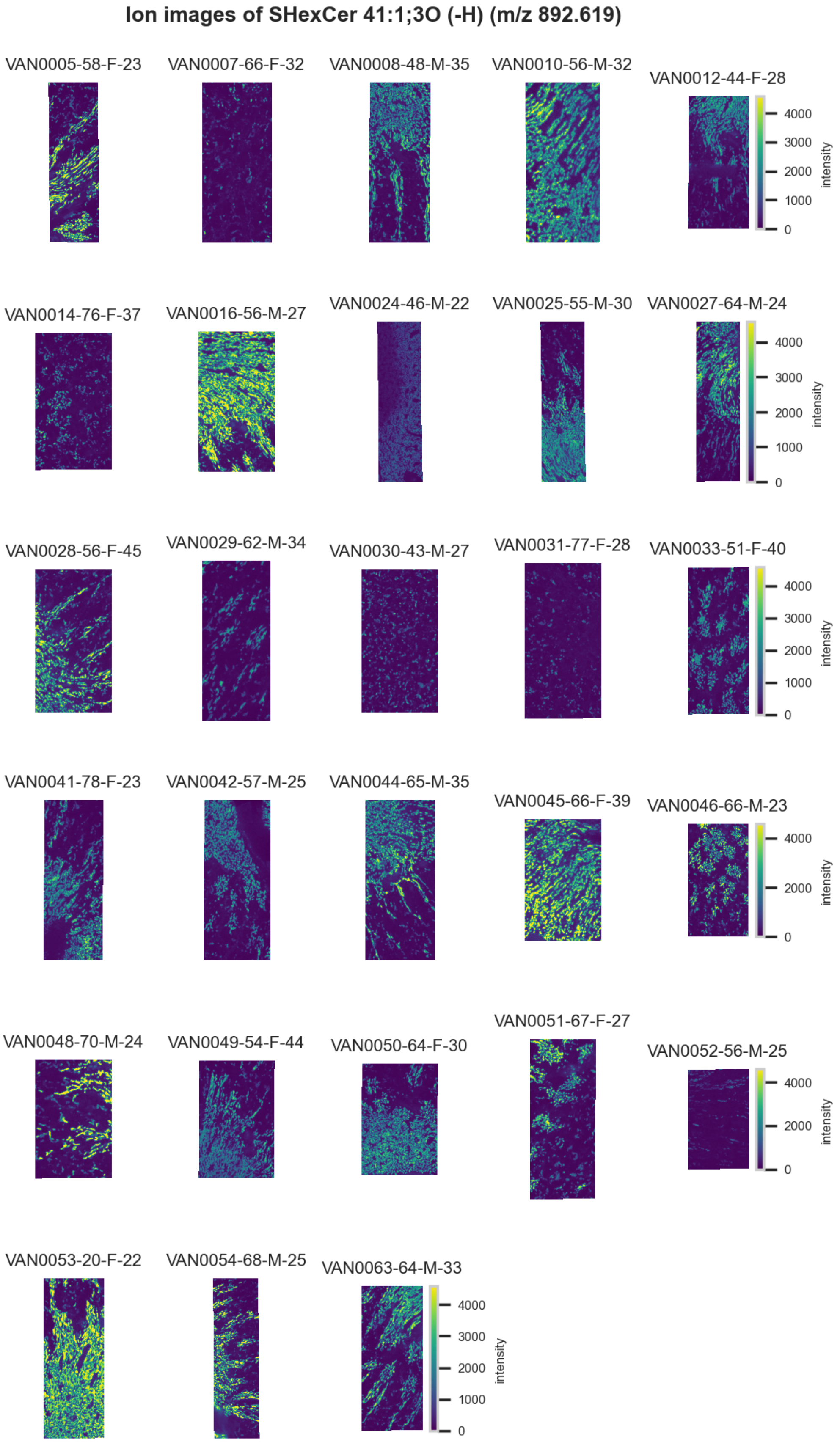

**Figure S89.** 10 µm MALDI IMS images of [SHexCer(41:1,3O)-H]$^-$ (*m/z* 892.619) detected in negative ion mode from each donor sample. Multivariate SHAP analysis determined this to be feature 1 for classification of thick ascending limb.

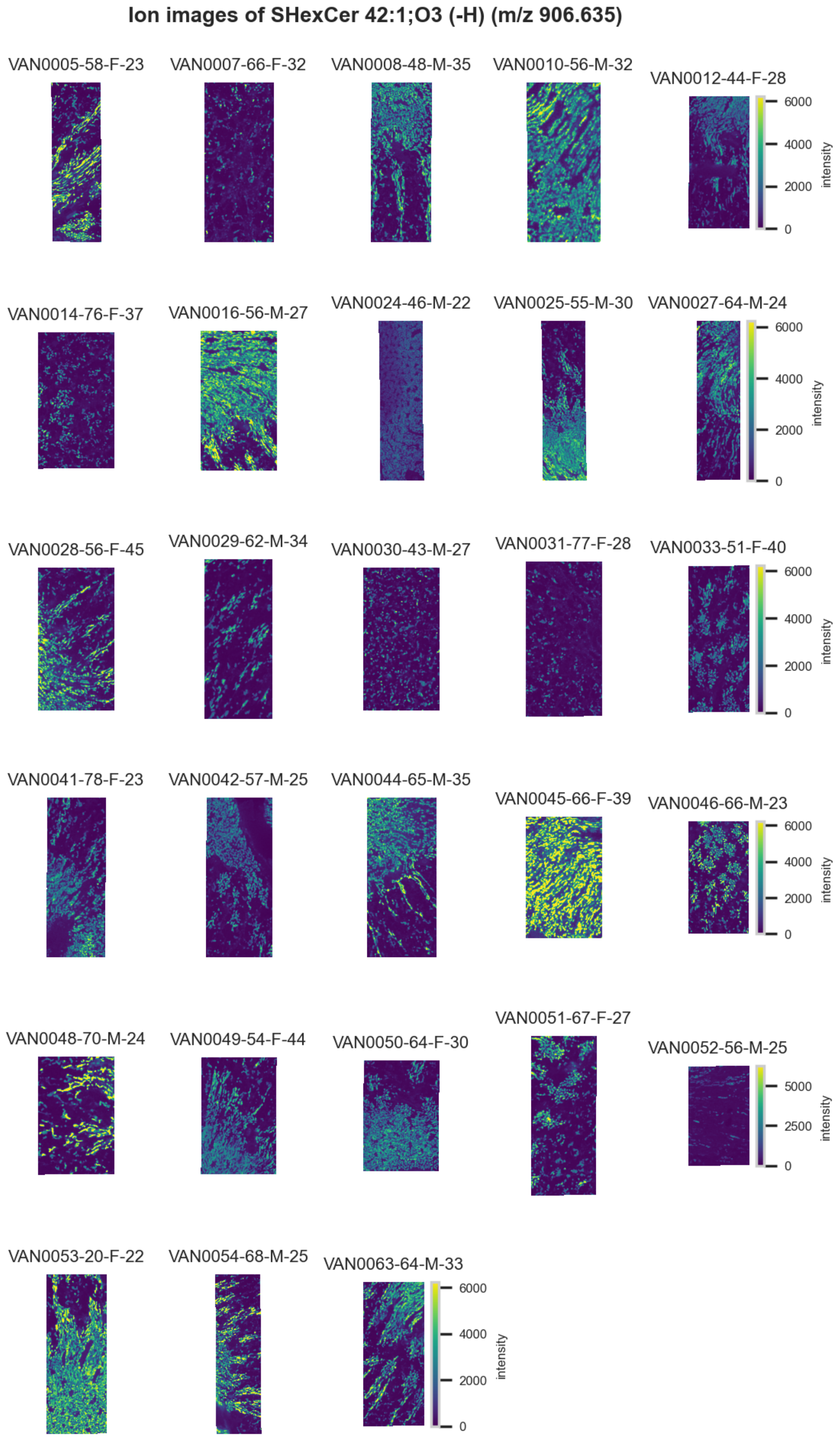

**Figure S90.** 10 μm MALDI IMS images of [SHexCer(42:1,3O)-H]$^-$ (*m/z* 906.635) detected in negative ion mode from each donor sample. Multivariate SHAP analysis determined this to be feature 2 for classification of thick ascending limb.

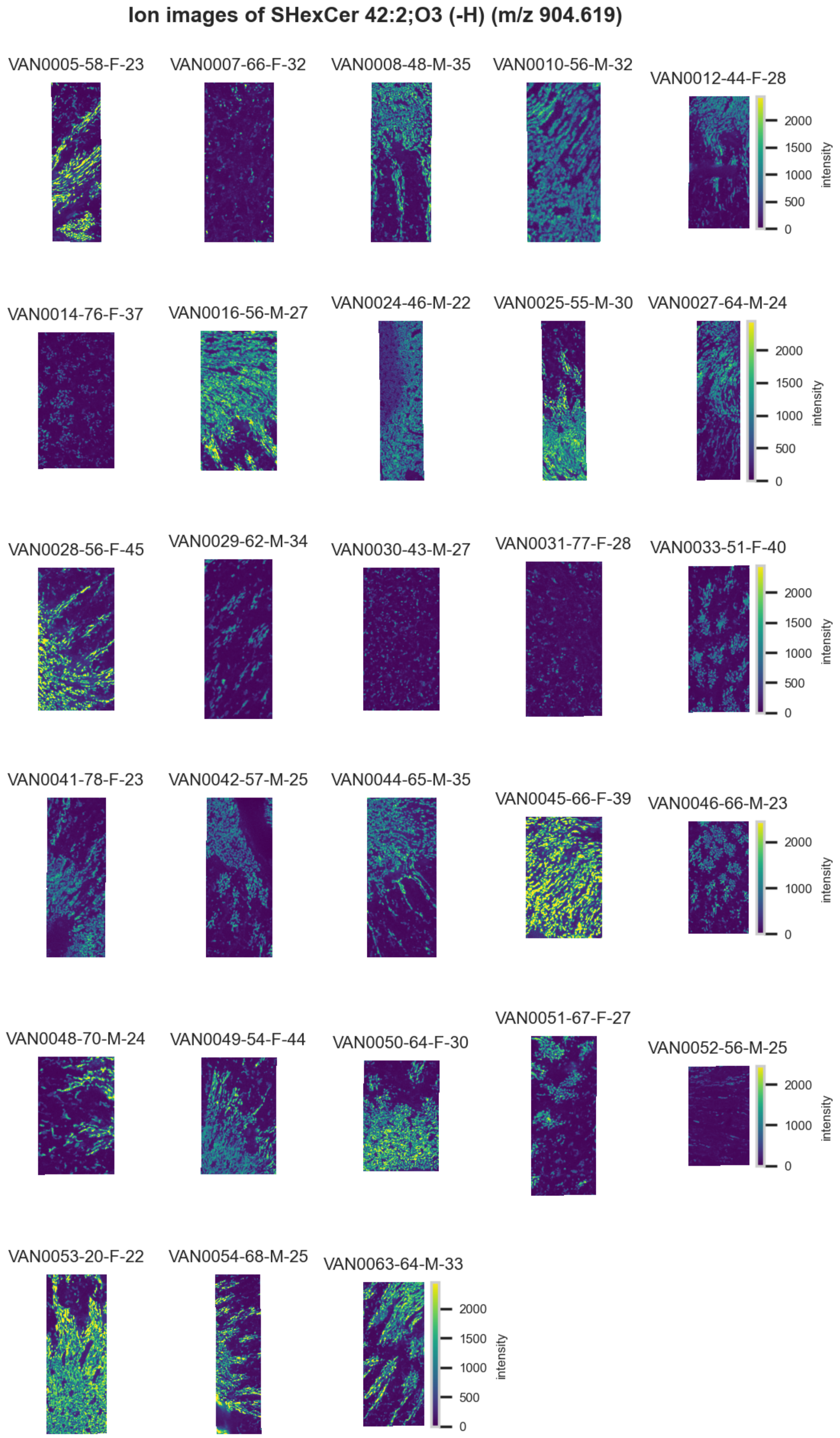

**Figure S91.** 10 µm MALDI IMS images of [SHexCer(42:2,3O)-H]$^-$ (*m/z* 904.619) detected in negative ion mode from each donor sample. Multivariate SHAP analysis determined this to be feature 3 for classification of thick ascending limb.

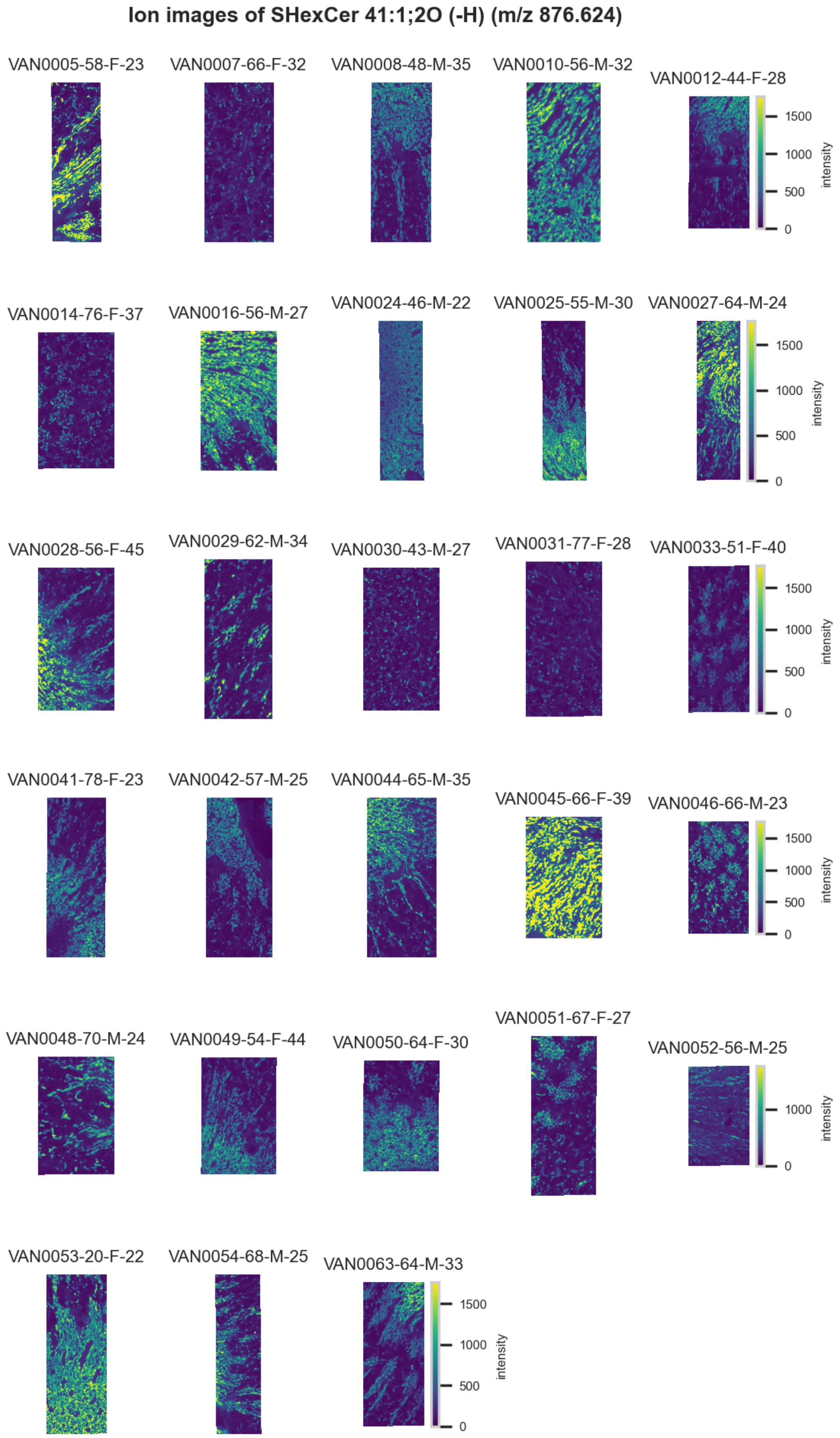

**Figure S92.** 10 μm MALDI IMS images of [SHexCer(41:1,2O)-H]$^-$ (*m/z* 876.624) detected in negative ion mode from each donor sample. Multivariate SHAP analysis determined this to be feature 4 for classification of thick ascending limb.

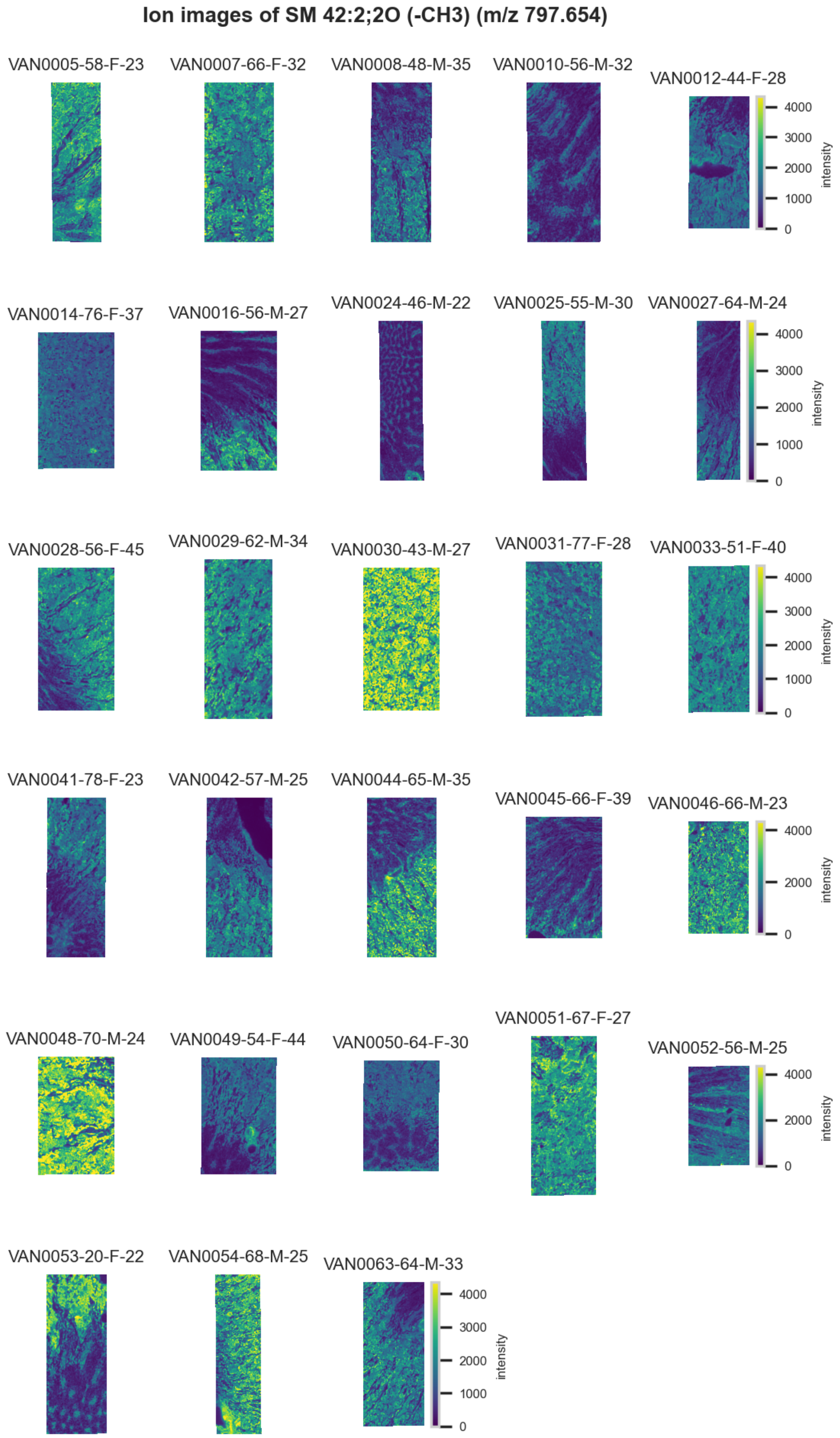

**Figure S93.** 10 µm MALDI IMS images of [SM(42:2,2O)-CH3]$^{-}$ (*m/z* 797.654) detected in negative ion mode from each donor sample. Multivariate SHAP analysis determined this to be feature 5 for classification of thick ascending limb.

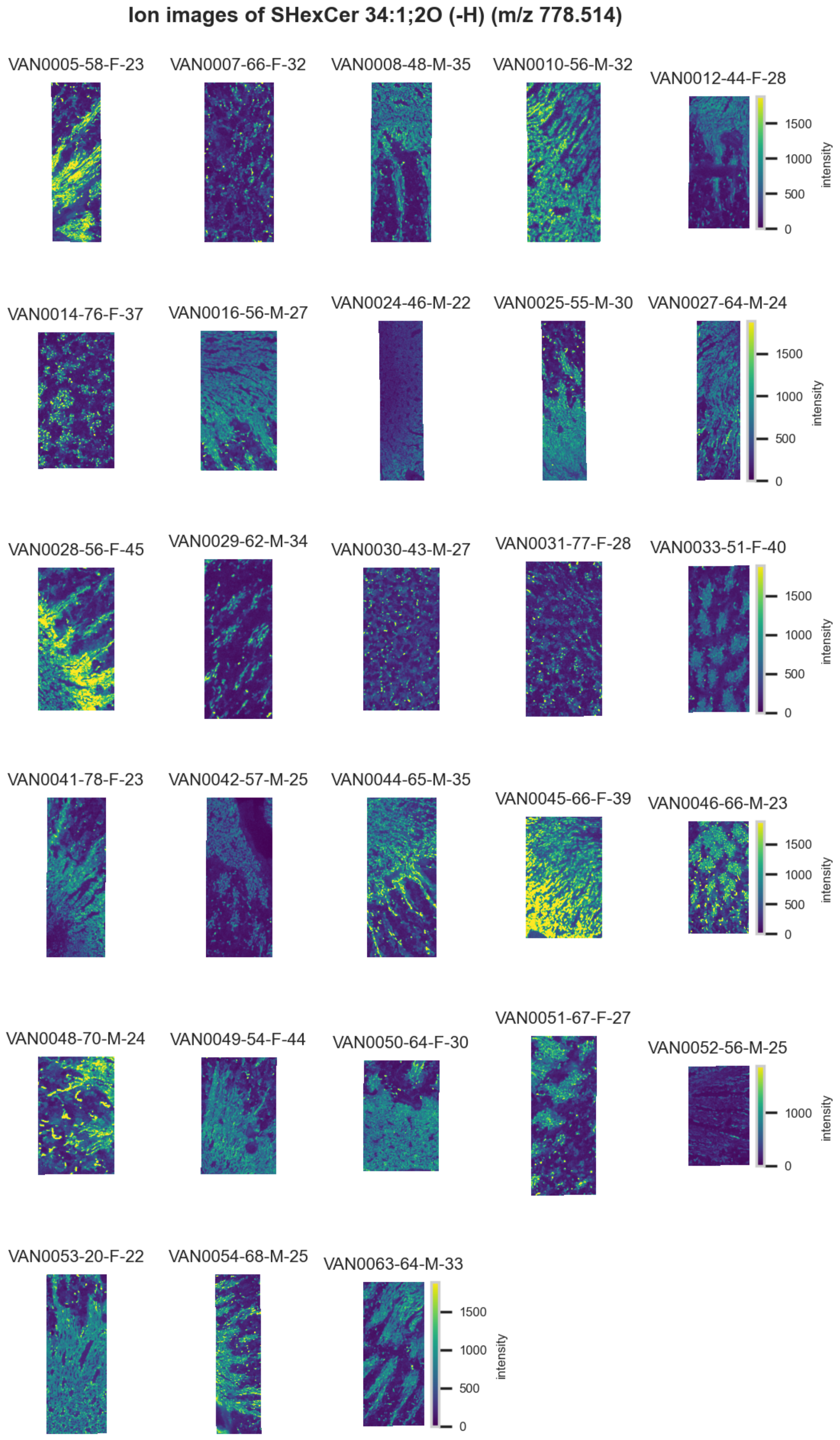

**Figure S94.** 10 µm MALDI IMS images of [SHexCer(34:1,2O)-H]$^{-}$ (*m/z* 778.514) detected in negative ion mode from each donor sample. Multivariate SHAP analysis determined this to be feature 6 for classification of thick ascending limb.

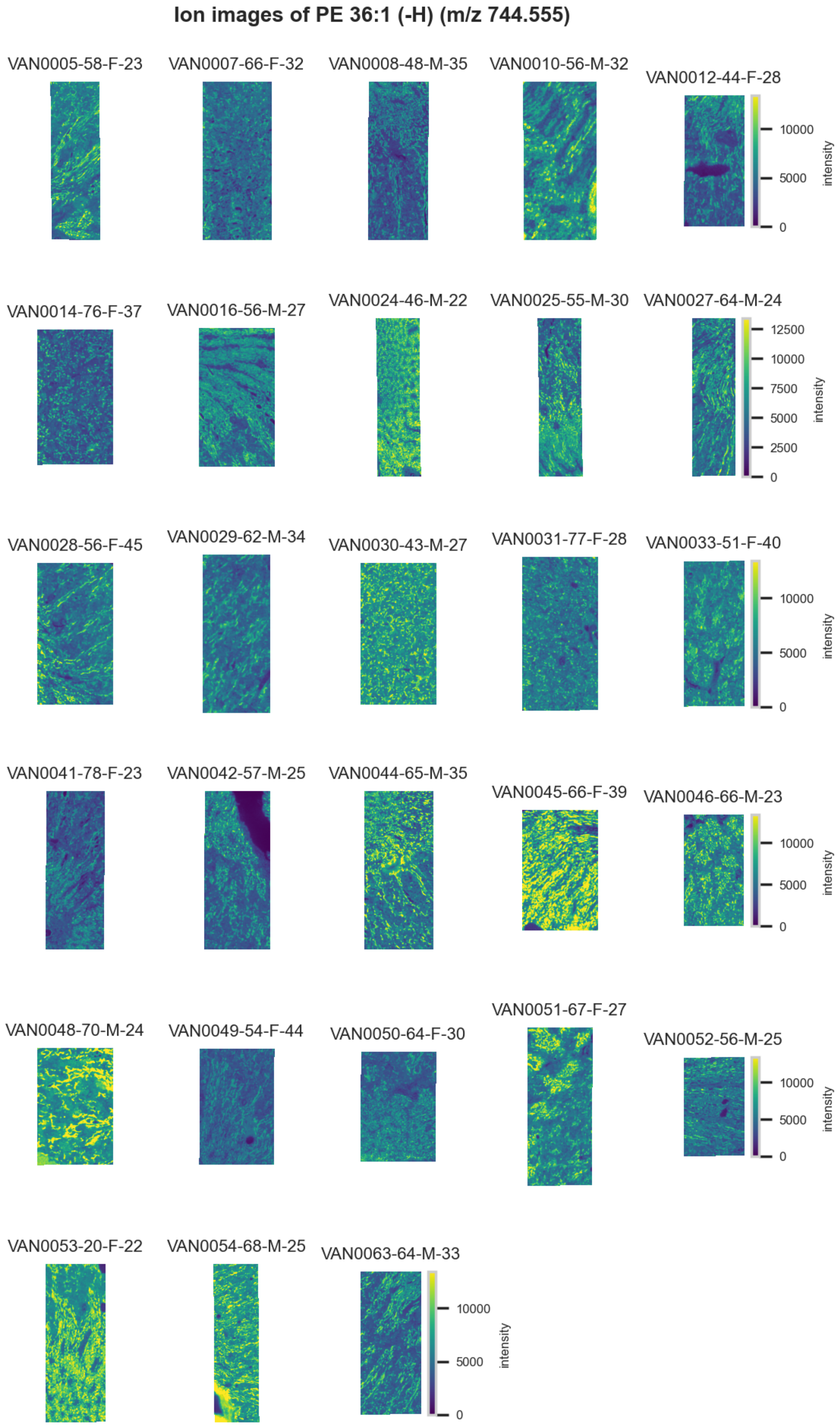

**Figure S95.** 10 µm MALDI IMS images of [PE(36:1)-H]$^-$ (*m/z* 744.555) detected in negative ion mode from each donor sample. Multivariate SHAP analysis determined this to be feature 7 for classification of thick ascending limb.

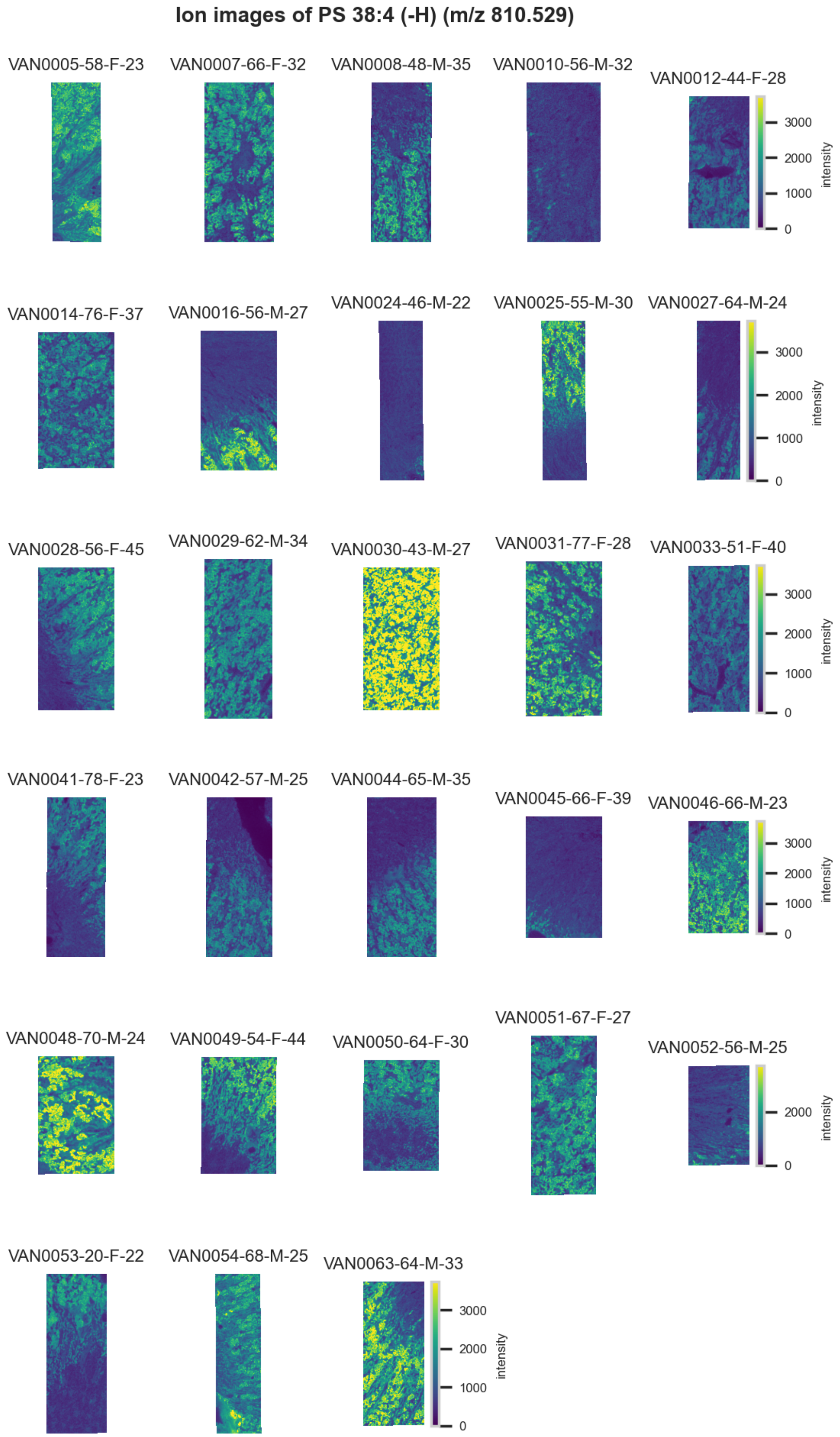

**Figure S96.** 10 µm MALDI IMS images of [PS(38:4)-H]⁻ (*m/z* 810.529) detected in negative ion mode from each donor sample. Multivariate SHAP analysis determined this to be feature 8 for classification of thick ascending limb.

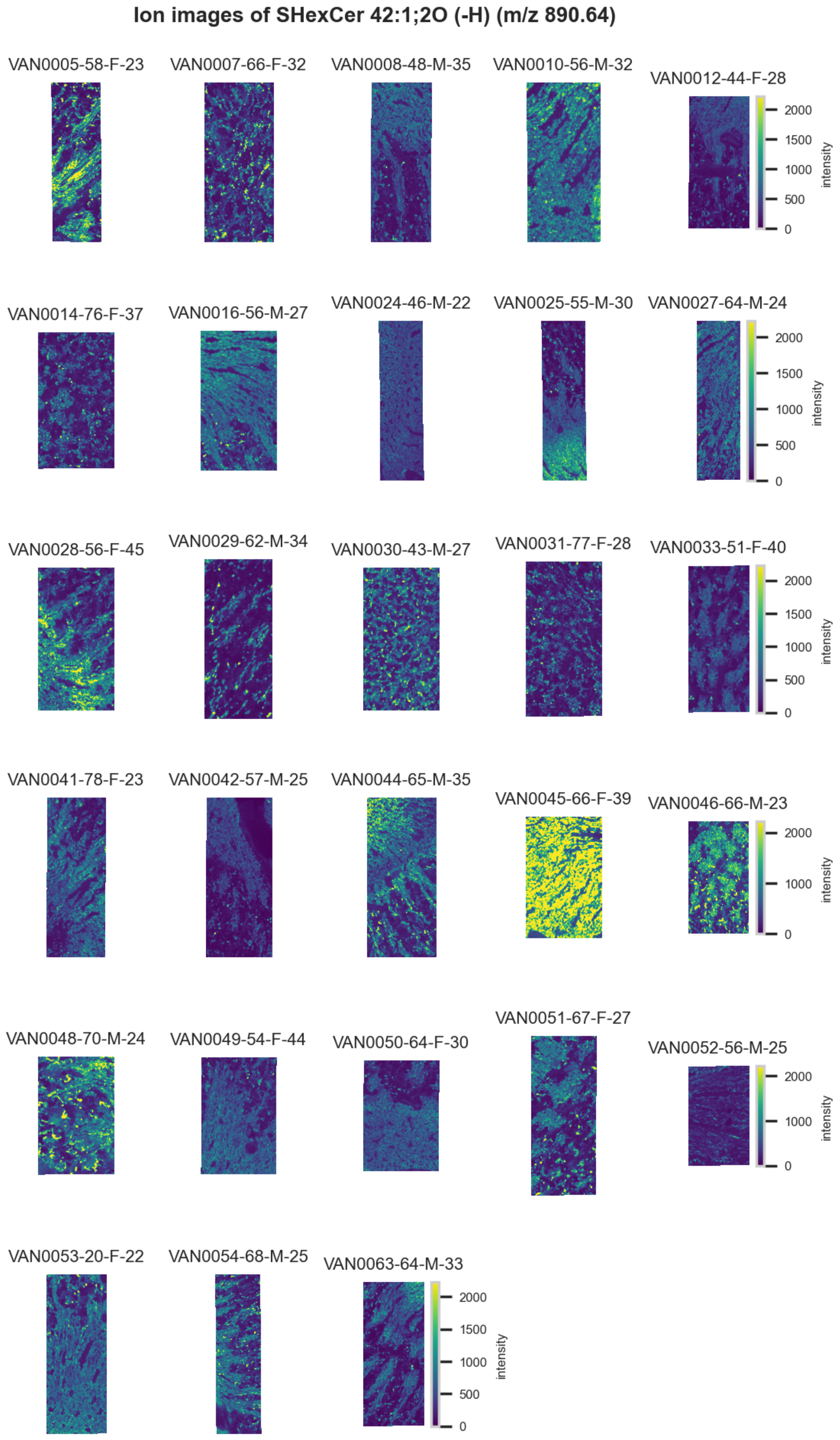

**Figure S97.** 10 µm MALDI IMS images of [SHexCer(42:1,2O)-H]$^-$ (*m/z* 890.64) detected in negative ion mode from each donor sample. Multivariate SHAP analysis determined this to be feature 9 for classification of thick ascending limb.

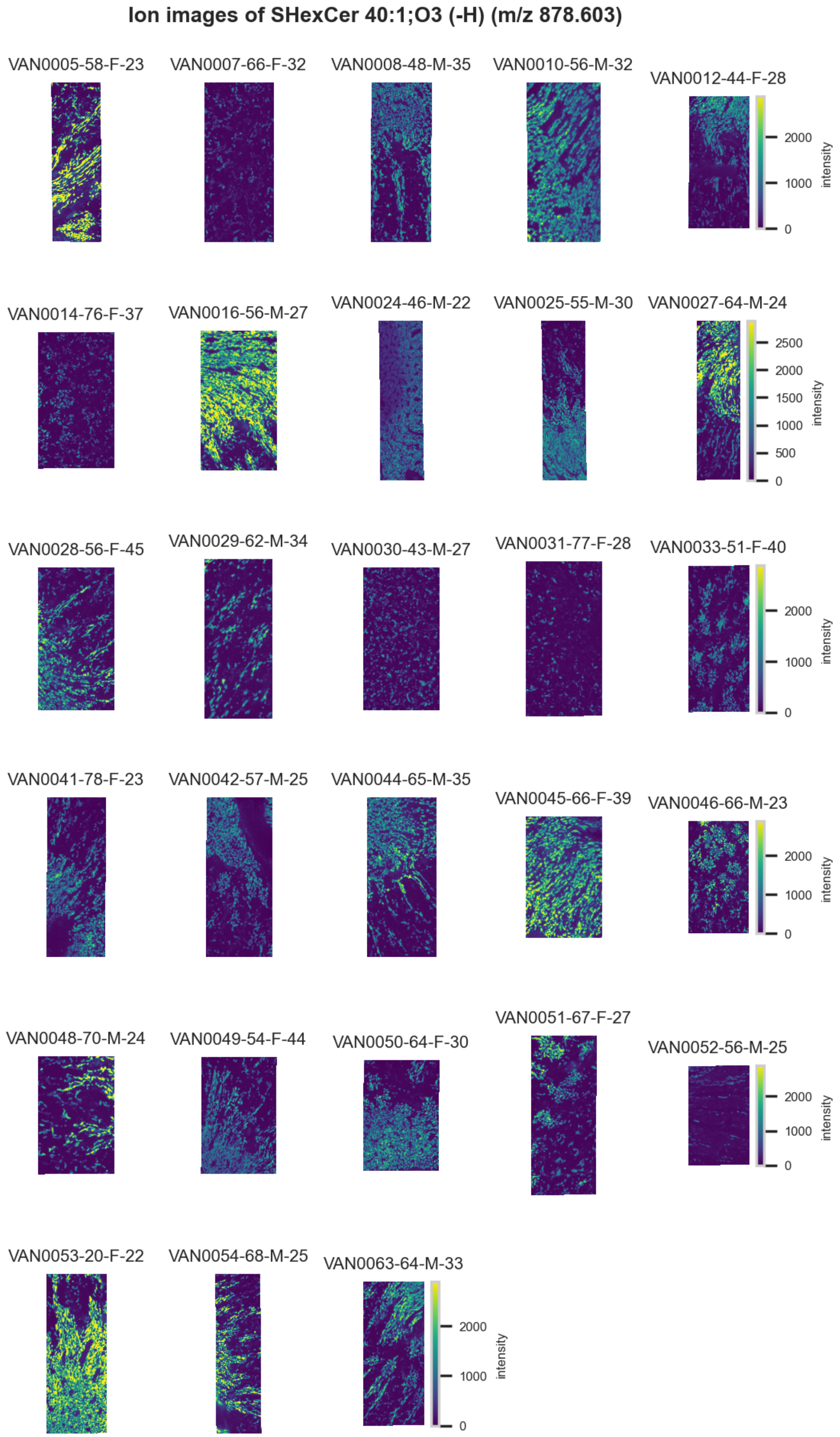

**Figure S98.** 10 µm MALDI IMS images of [SHexCer(40:1,3O)-H]$^-$ (*m/z* 878.603) detected in negative ion mode from each donor sample. Multivariate SHAP analysis determined this to be feature 10 for classification of thick ascending limb.

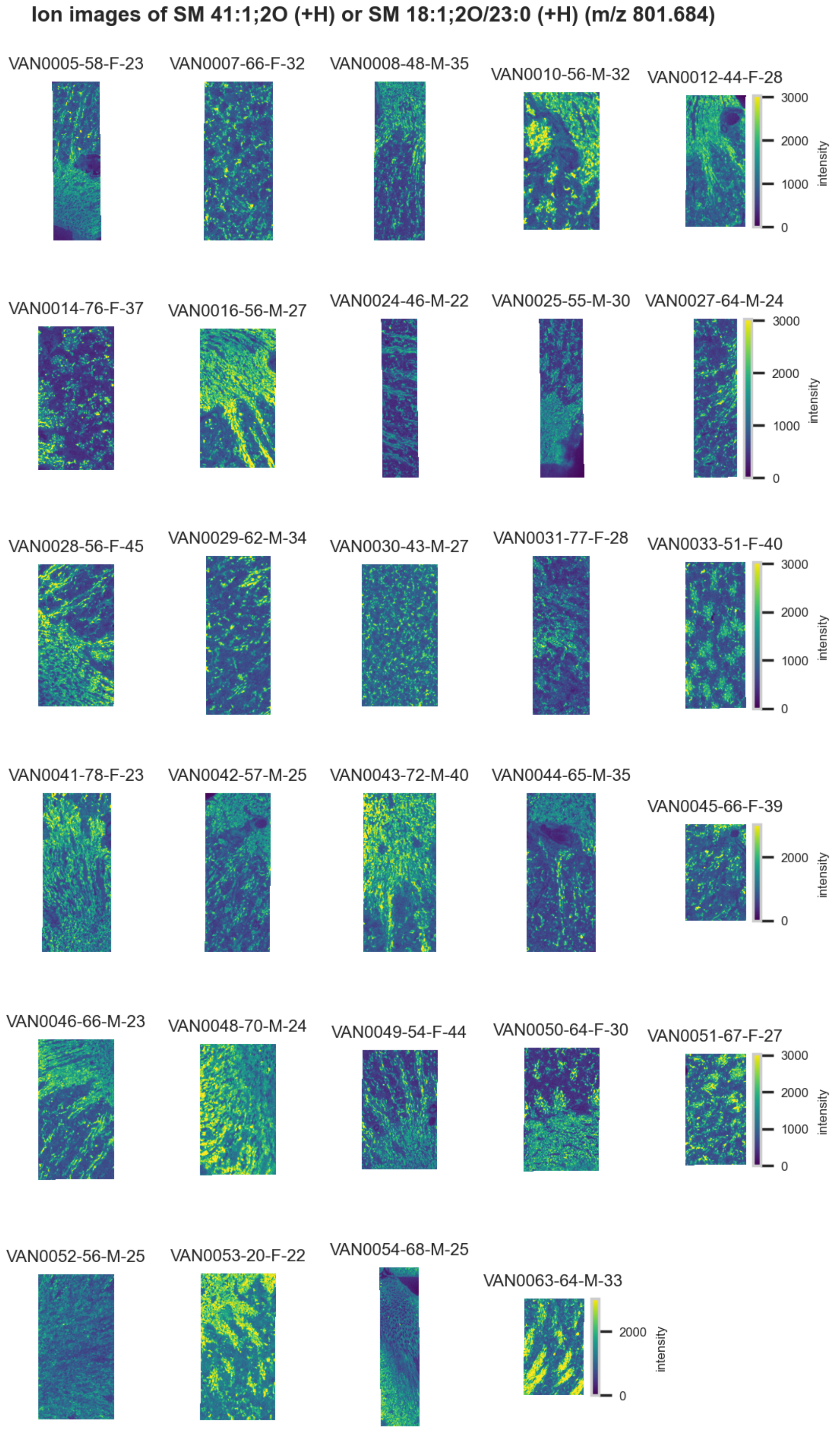

**Figure S99.** 10 µm MALDI IMS images of [SM(41:1,2O)+H]$^+$ or [SM(18:1,2O/23:0)+H]$^+$ (*m/z* 801.684) detected in positive ion mode from each donor sample. These molecules are isomers and therefore indistinguishable by MS alone. Multivariate SHAP analysis determined this to be feature 1 for classification of thick ascending limbs.

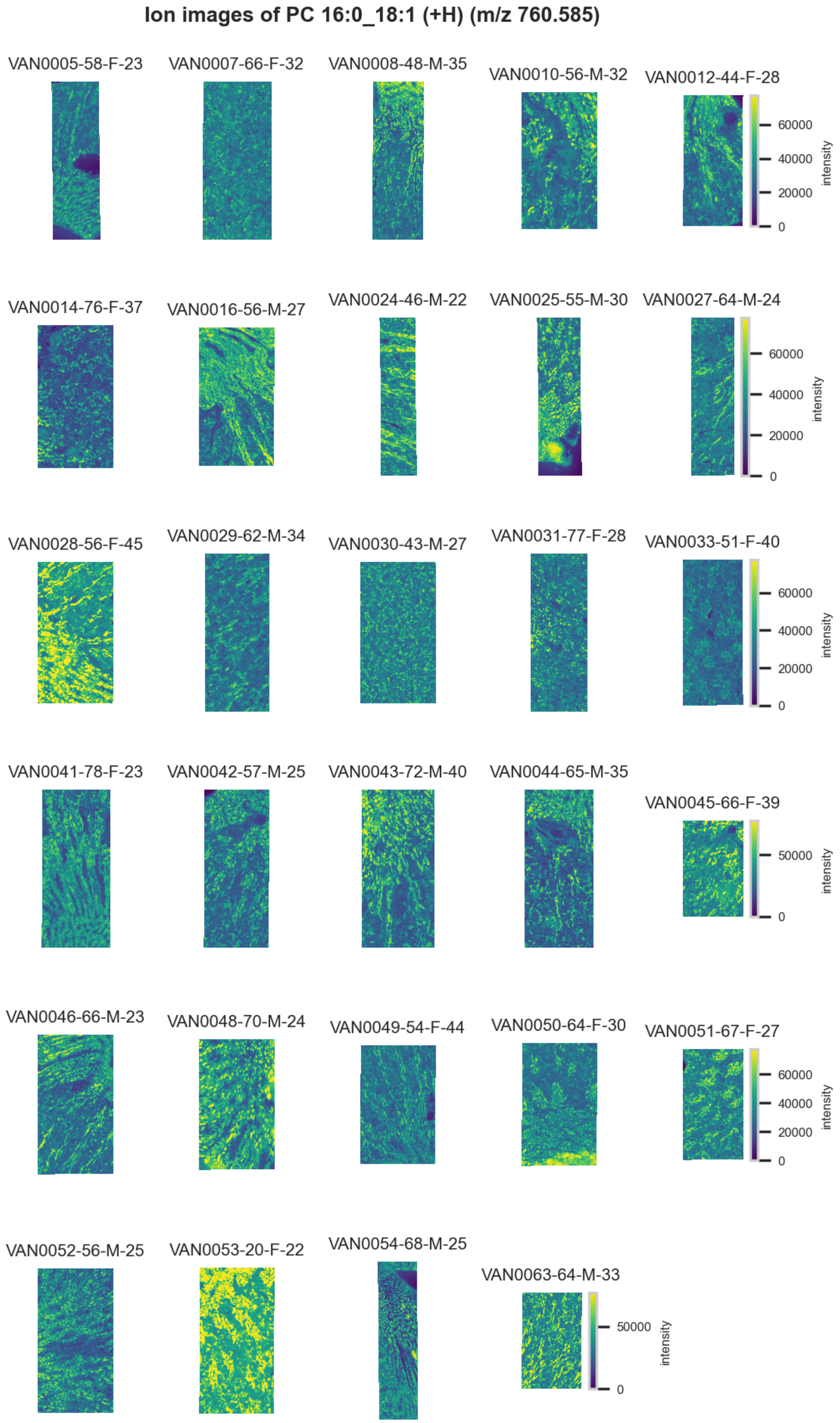

**Figure S100.** 10 µm MALDI IMS images of [PC(16:0_18:1)+H]$^+$ (*m/z* 760.585) detected in positive ion mode from each donor sample. Multivariate SHAP analysis determined this to be feature 2 for classification of thick ascending limbs.

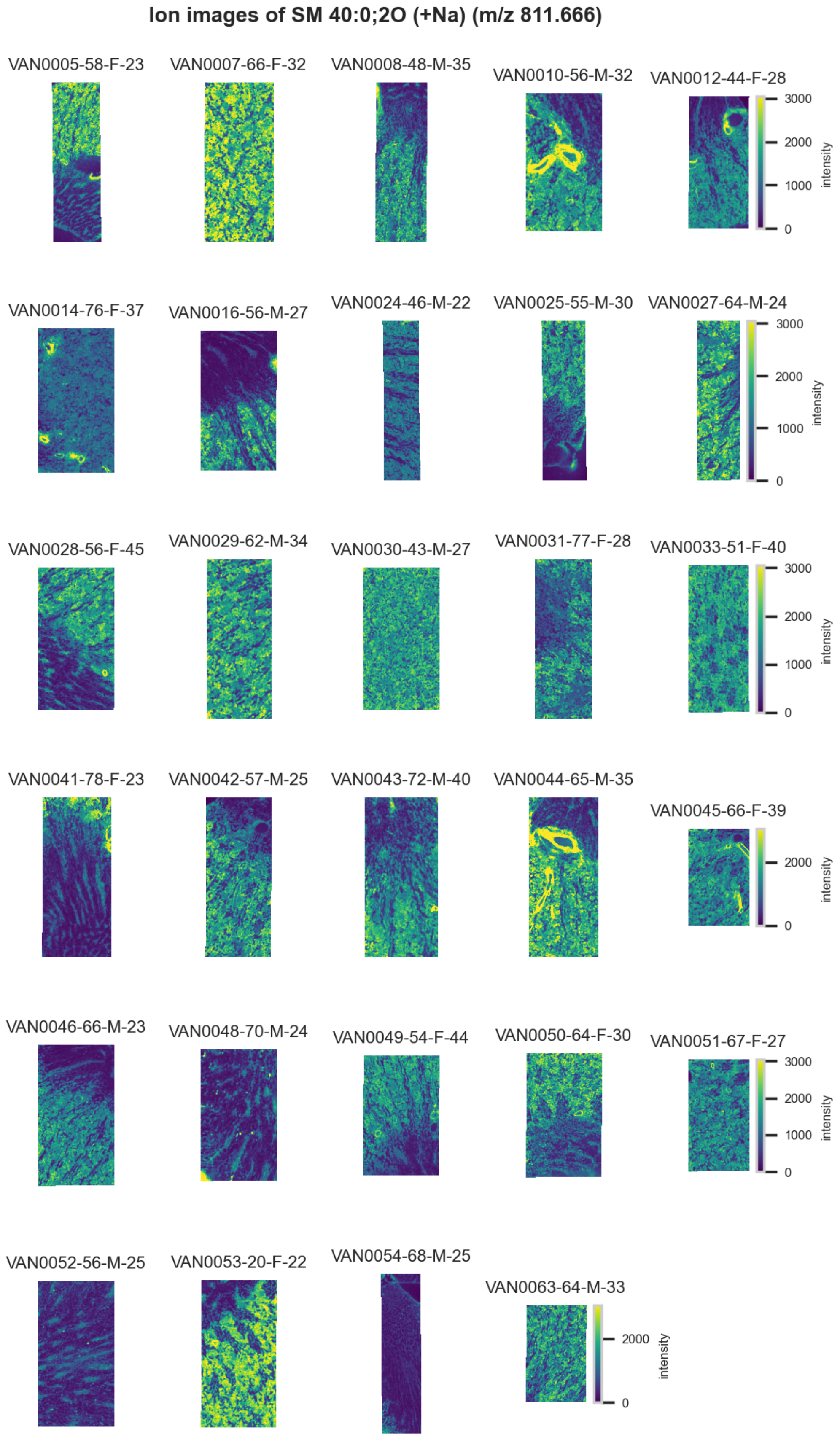

**Figure S101.** 10 µm MALDI IMS images of $[SM(40:0,2O)+Na]^{+}$ (*m/z* 811.666) detected in positive ion mode from each donor sample. Multivariate SHAP analysis determined this to be feature 3 for classification of thick ascending limbs.

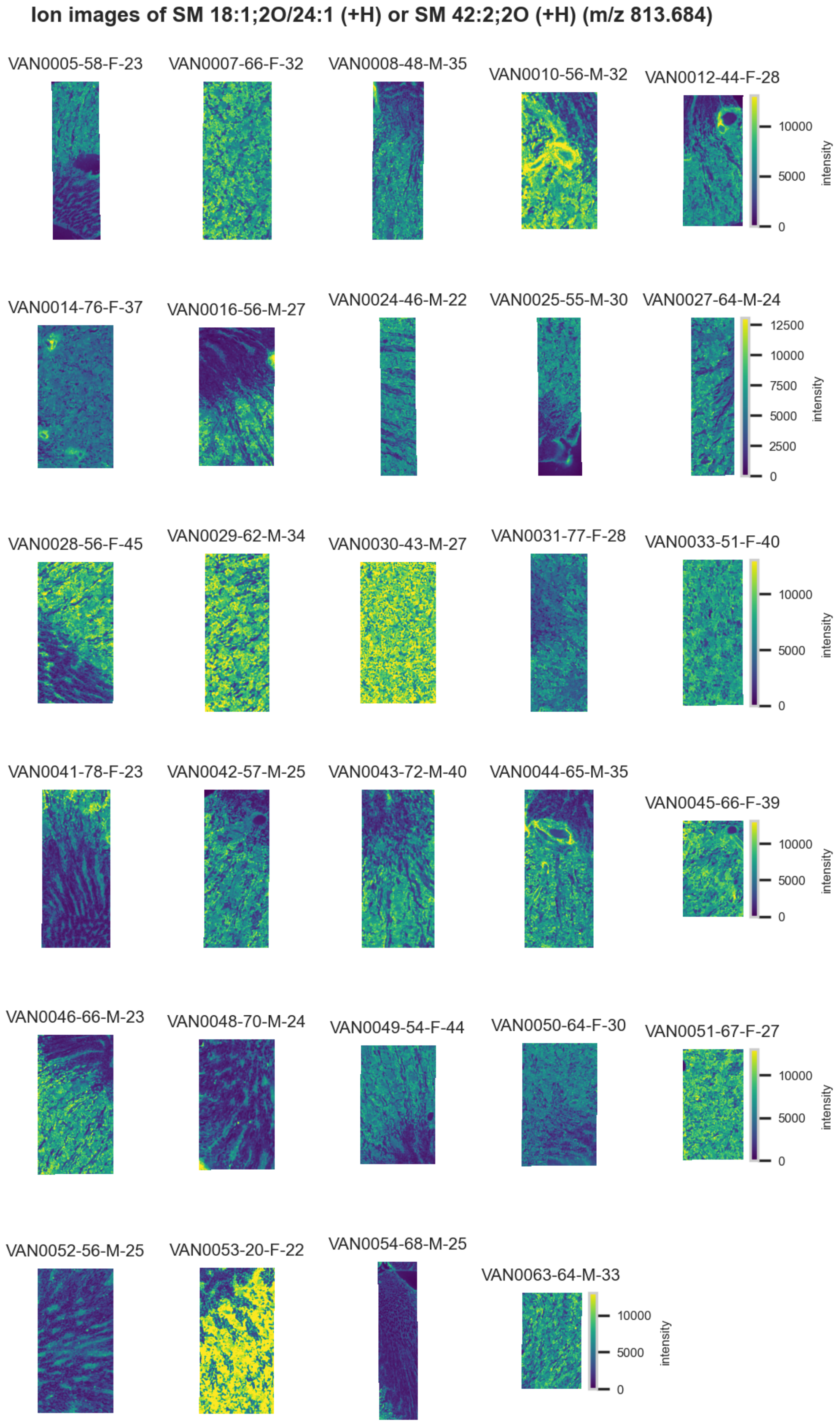

**Figure S102.** 10 µm MALDI IMS images of [SM(42:2,2O)+H]$^+$ or [SM(18:1,2O/24:1)+H]$^+$ (*m/z* 813.684) detected in positive ion mode from each donor sample. These molecules are isomers and therefore indistinguishable by MS alone. Multivariate SHAP analysis determined this to be feature 4 for classification of thick ascending limbs.

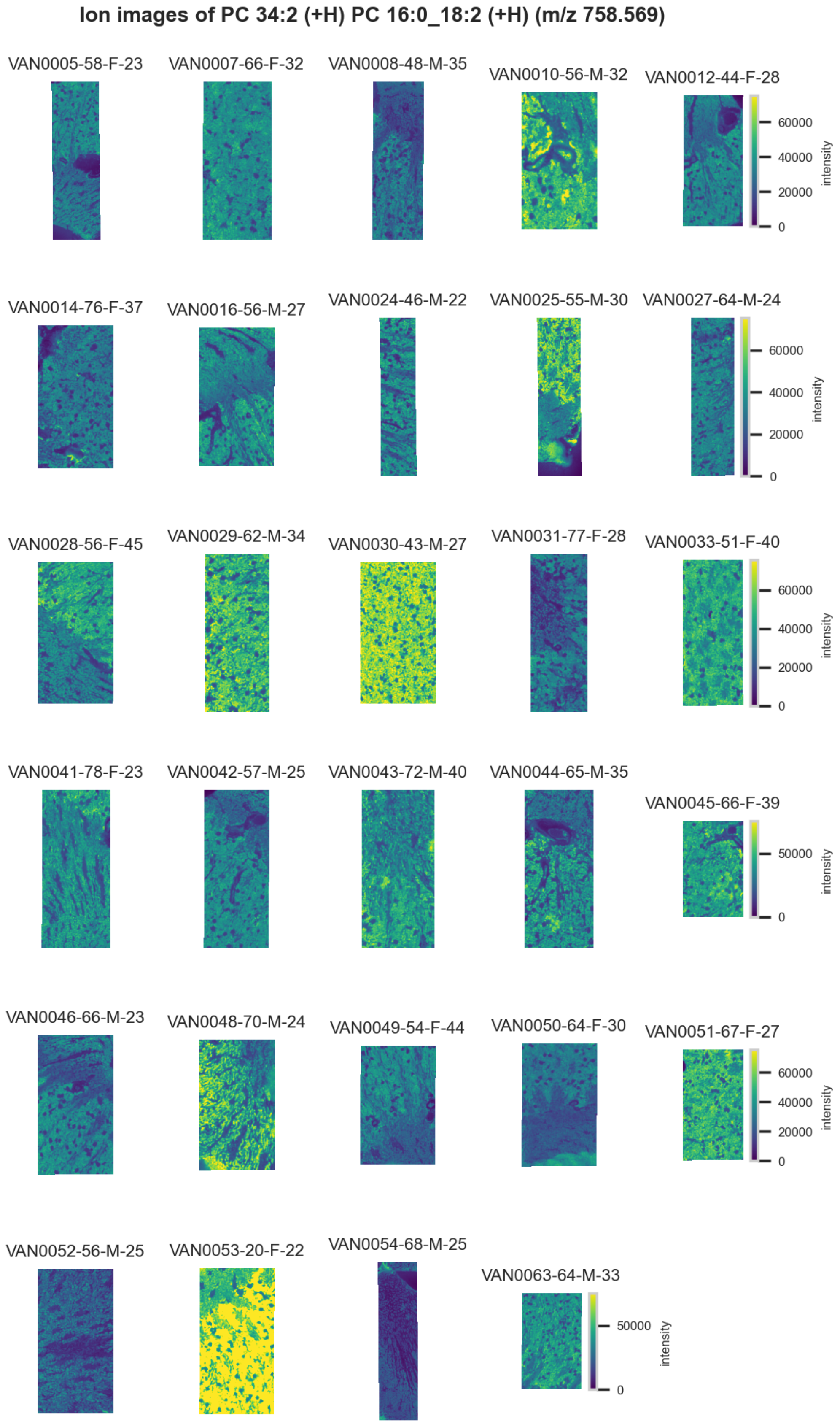

**Figure S103.** 10 µm MALDI IMS images of $[PC(16:0\_18:2)+H]^{+}$ (*m/z* 758.569) detected in positive ion mode from each donor sample. Multivariate SHAP analysis determined this to be feature 5 for classification of thick ascending limbs.

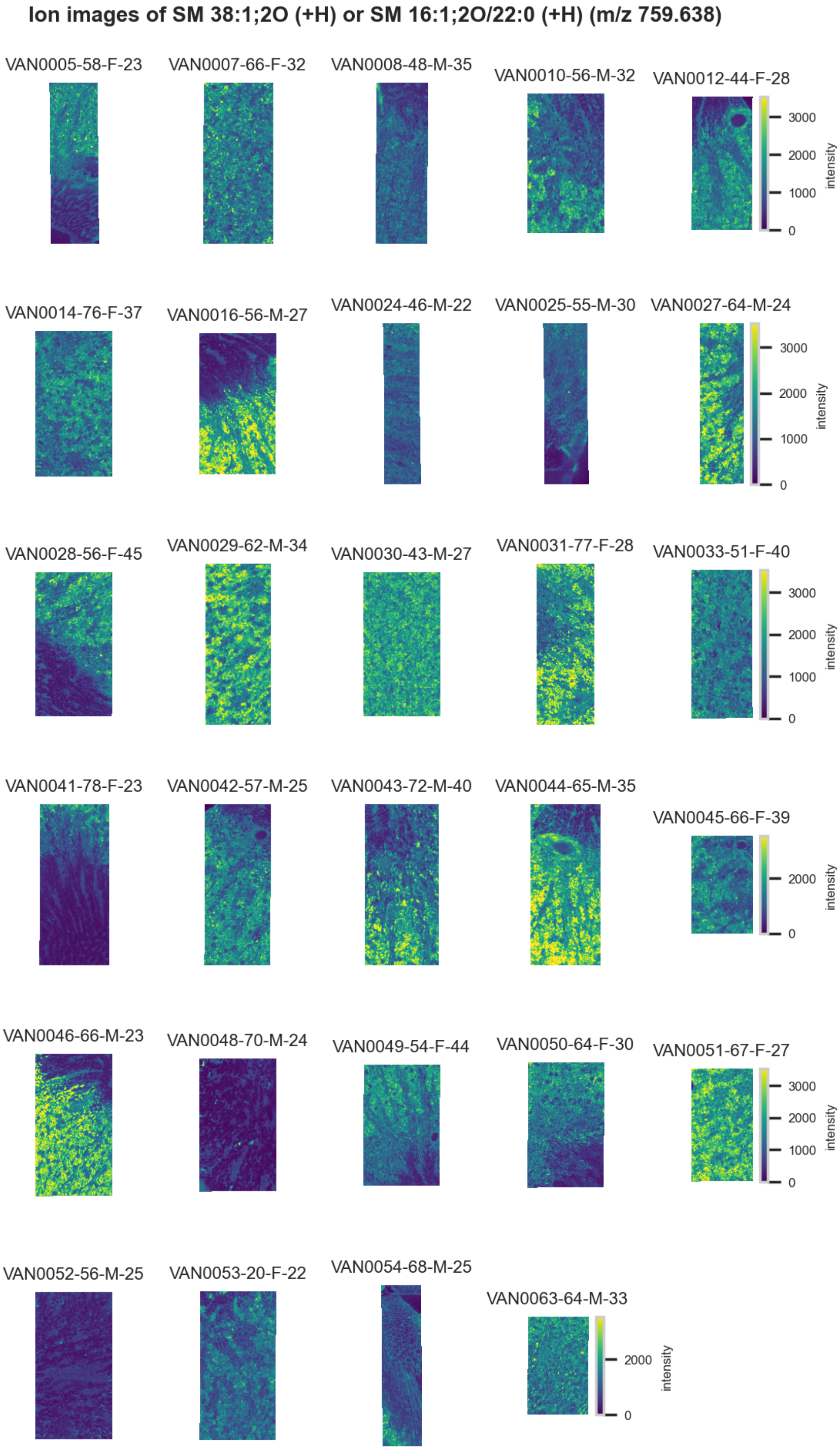

**Figure S104.** 10 µm MALDI IMS images of [SM(38:1,2O)+H]$^+$ or [SM(16:1,2O/22:0)+H]$^+$ (*m/z* 759.638) detected in positive ion mode from each donor sample. These molecules are isomers and therefore indistinguishable by MS alone. Multivariate SHAP analysis determined this to be feature 6 for classification of thick ascending limbs.

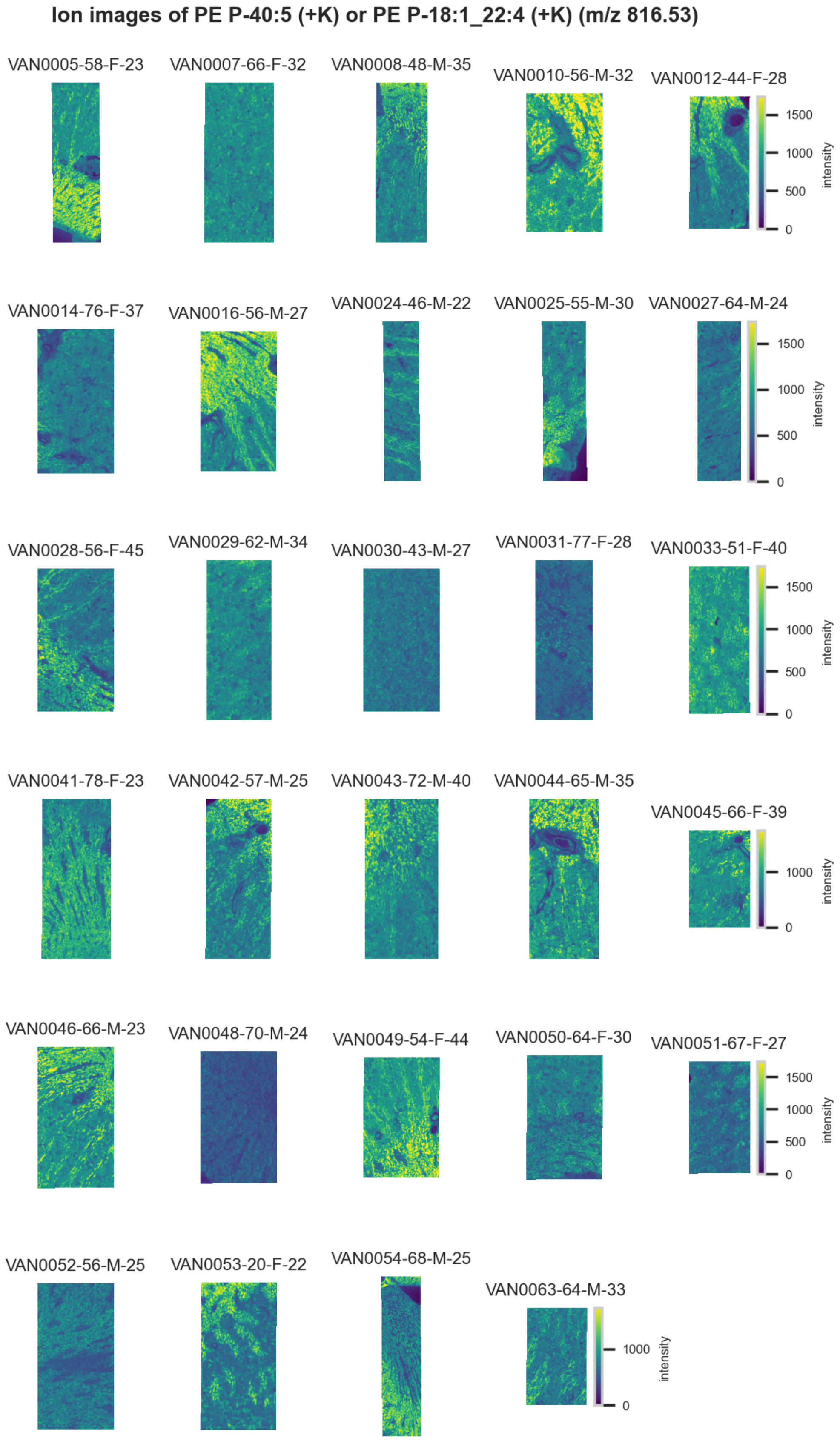

**Figure S105.** 10 µm MALDI IMS images of $[PE(P\text{-}40{:}5)+K]^+$ or $[PE(P\text{-}18{:}1\_22{:}4)+K]^+$ (*m/z* 816.53) detected in positive ion mode from each donor sample. These molecules are isomers and therefore indistinguishable by MS alone. Multivariate SHAP analysis determined this to be feature 7 for classification of thick ascending limbs.

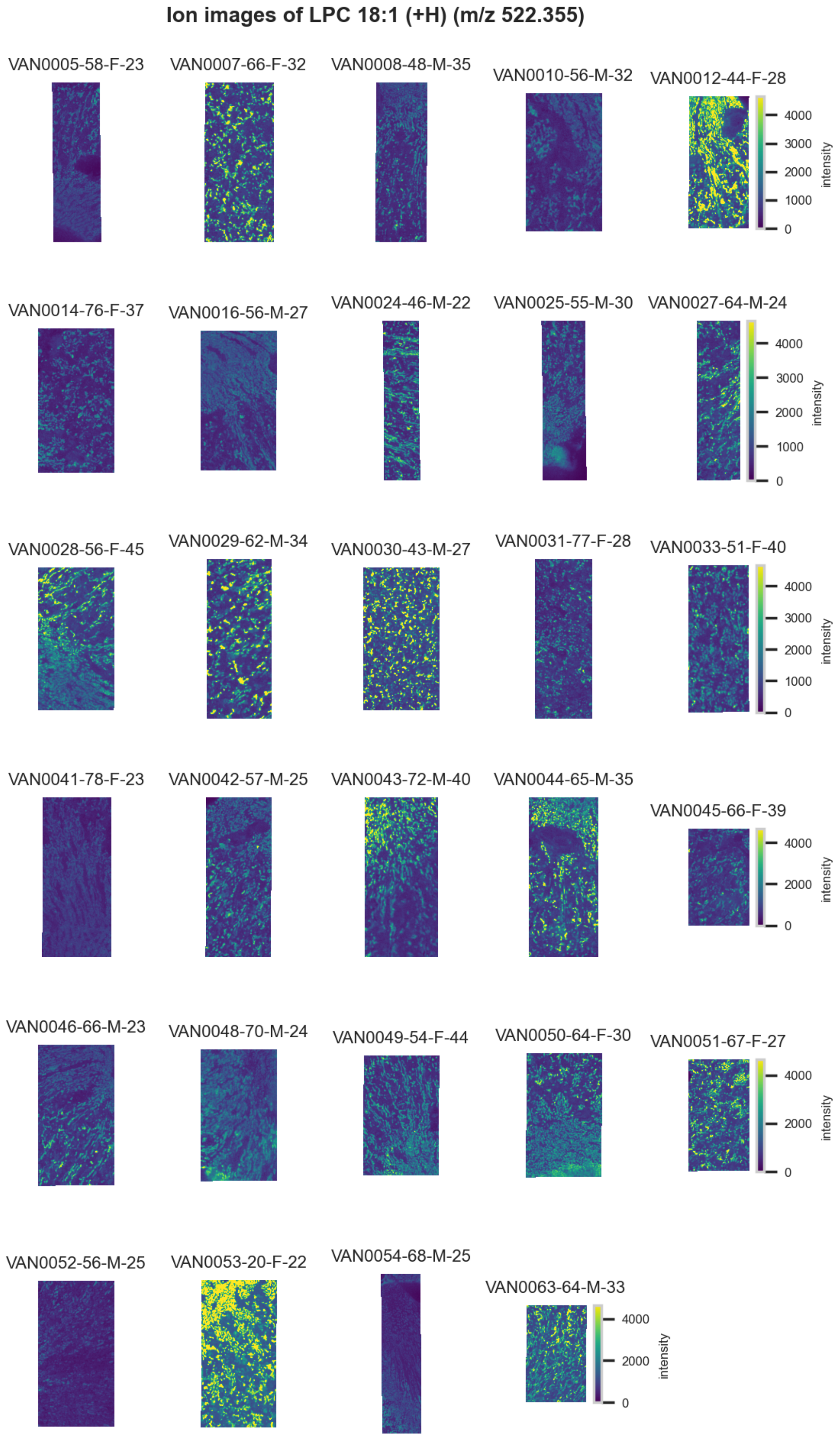

**Figure S106.** 10 µm MALDI IMS images of $[LPC(18:1)+H]^+$ (*m/z* 522.355) detected in positive ion mode from each donor sample. Multivariate SHAP analysis determined this to be feature 8 for classification of thick ascending limbs.

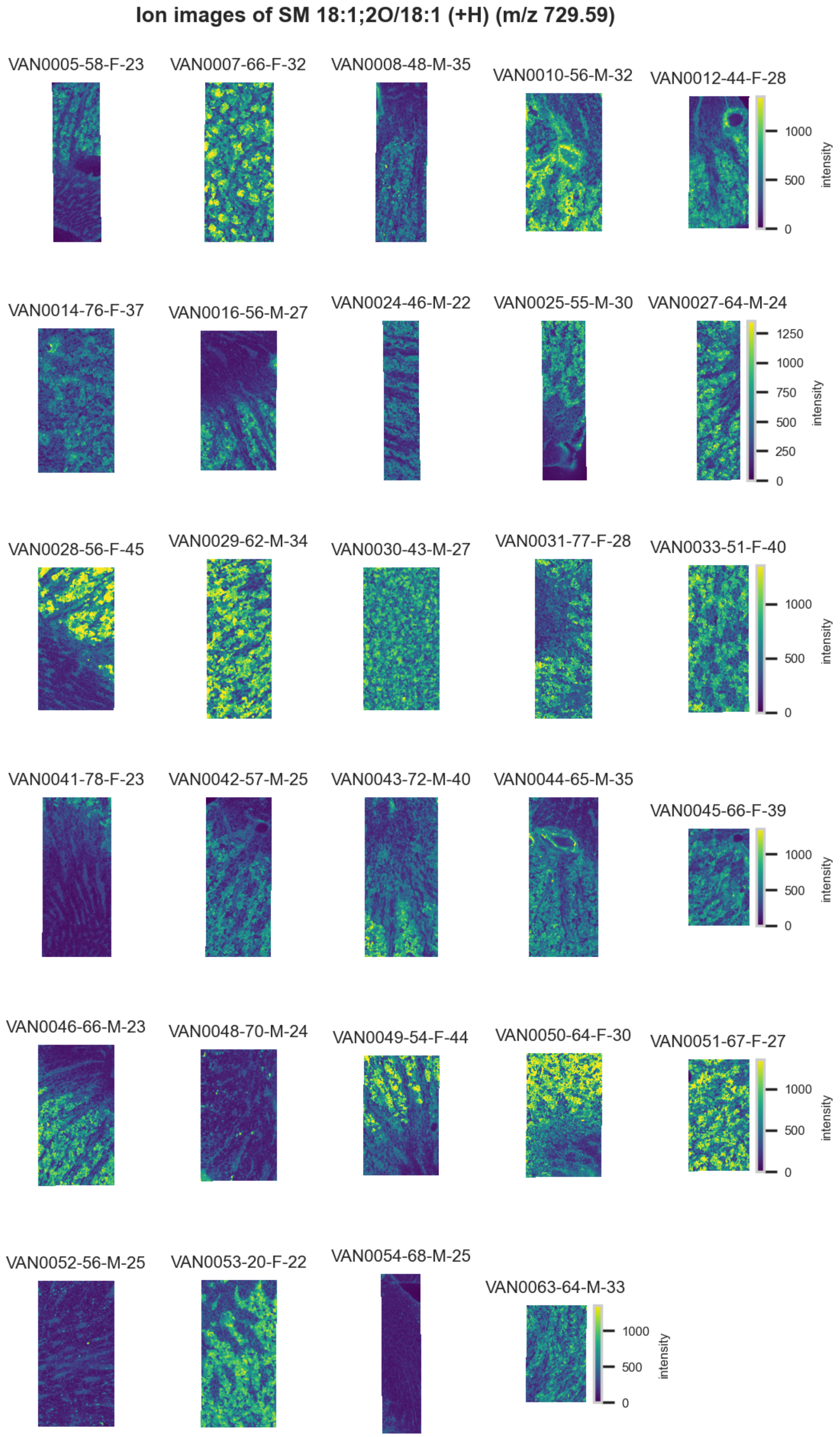

**Figure S107.** 10 µm MALDI IMS images of [SM(18:1,2O/18:1)+H]$^+$ (*m/z* 729.59) detected in <u>positive ion mode</u> from each donor sample. Multivariate SHAP analysis determined this to be feature 9 for classification of thick ascending limbs.

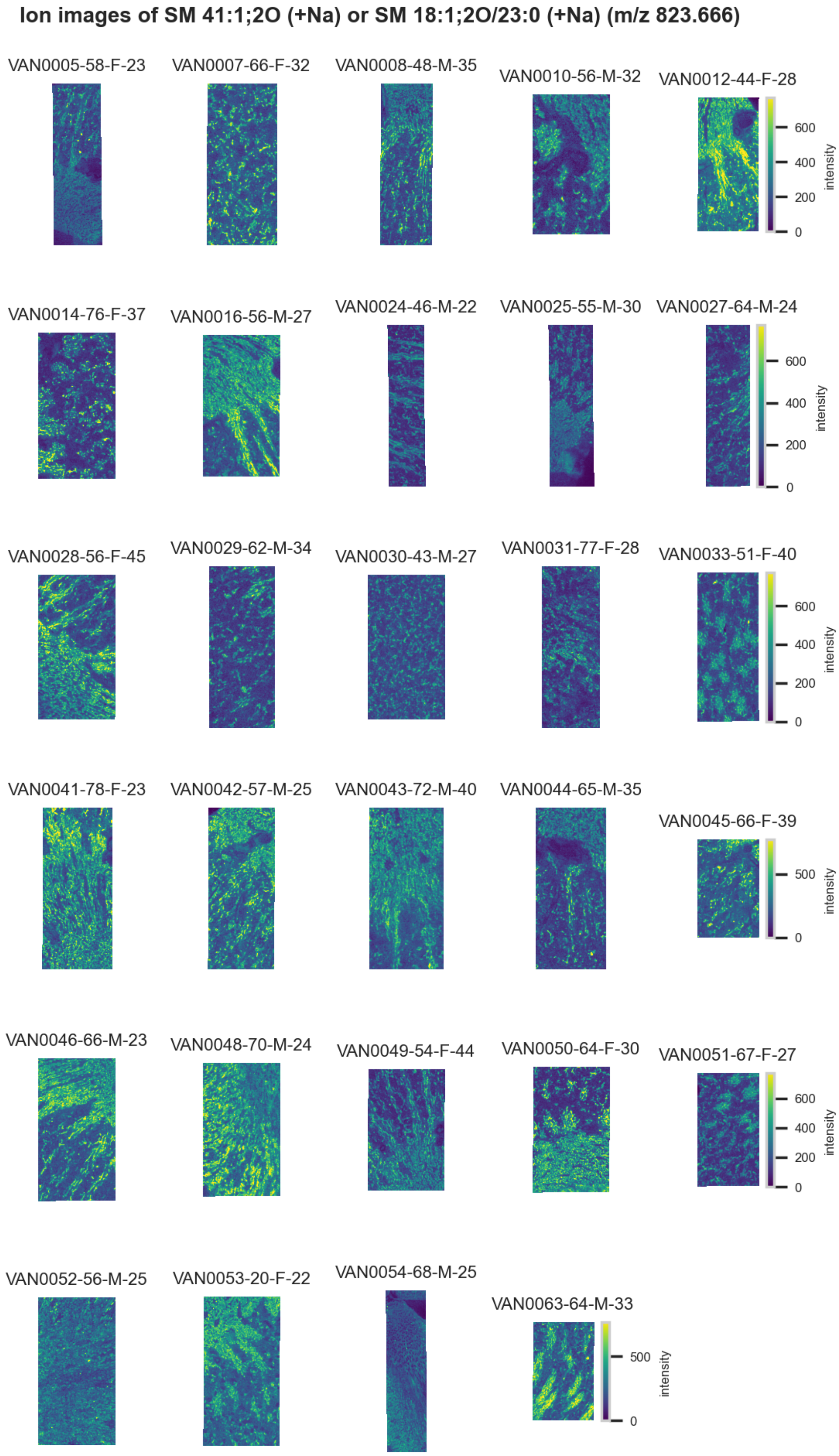

**Figure S108.** 10 µm MALDI IMS images of [SM(41:1,2O)+Na]$^+$ or [SM(18:1,2O/23:0)+Na]$^+$ (*m/z* 823.666) detected in positive ion mode from each donor sample. These molecules are isomers and therefore indistinguishable by MS alone. Multivariate SHAP analysis determined this to be feature 10 for classification of thick ascending limbs.

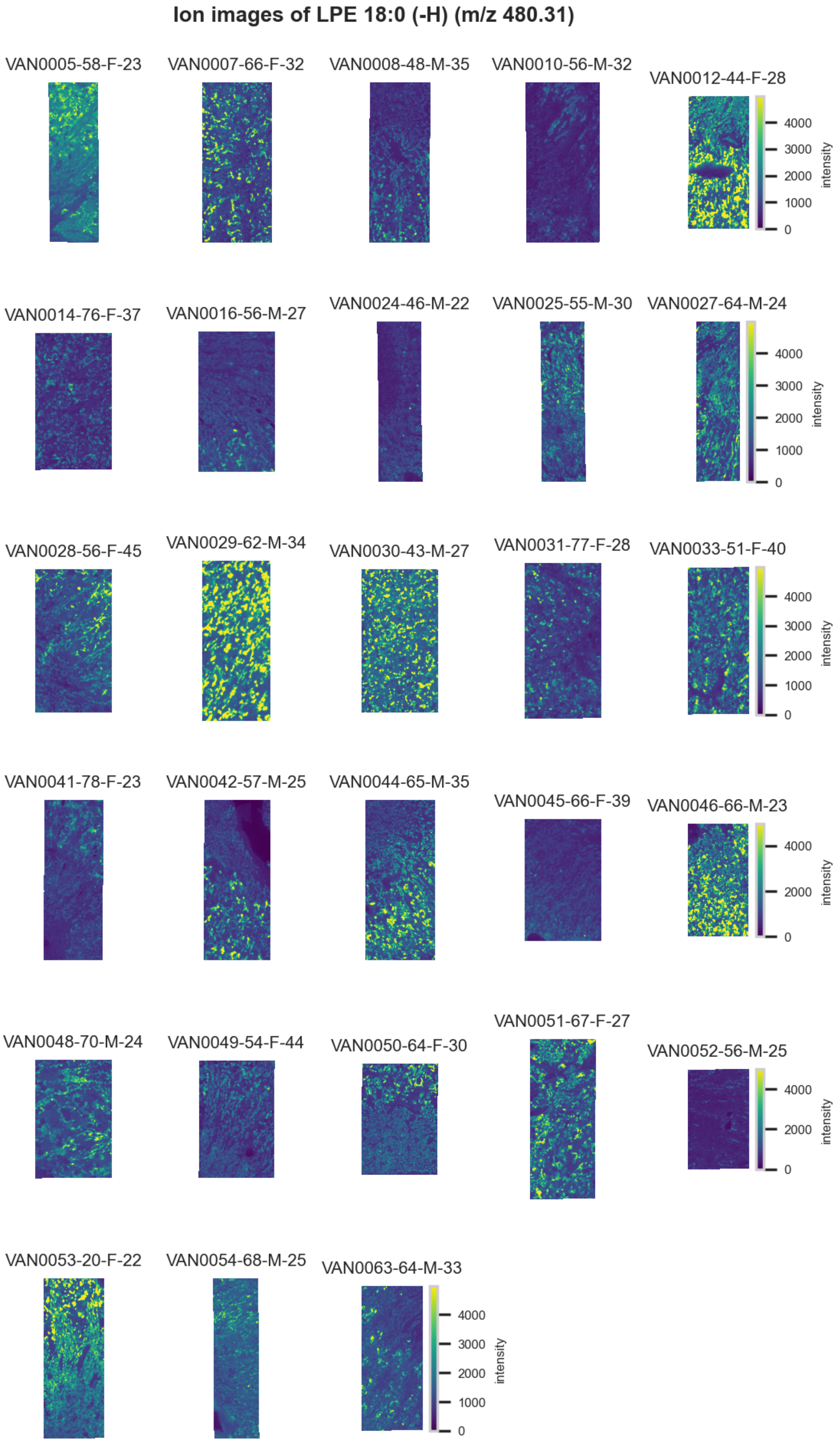

**Figure S109.** 10 µm MALDI IMS images of $[LPE(18:0)\text{-}H]^-$ (*m/z* 480.31) detected in <u>negative ion mode</u> from each donor sample. Multivariate SHAP analysis determined this to be feature 1 for classification of distal tubules.

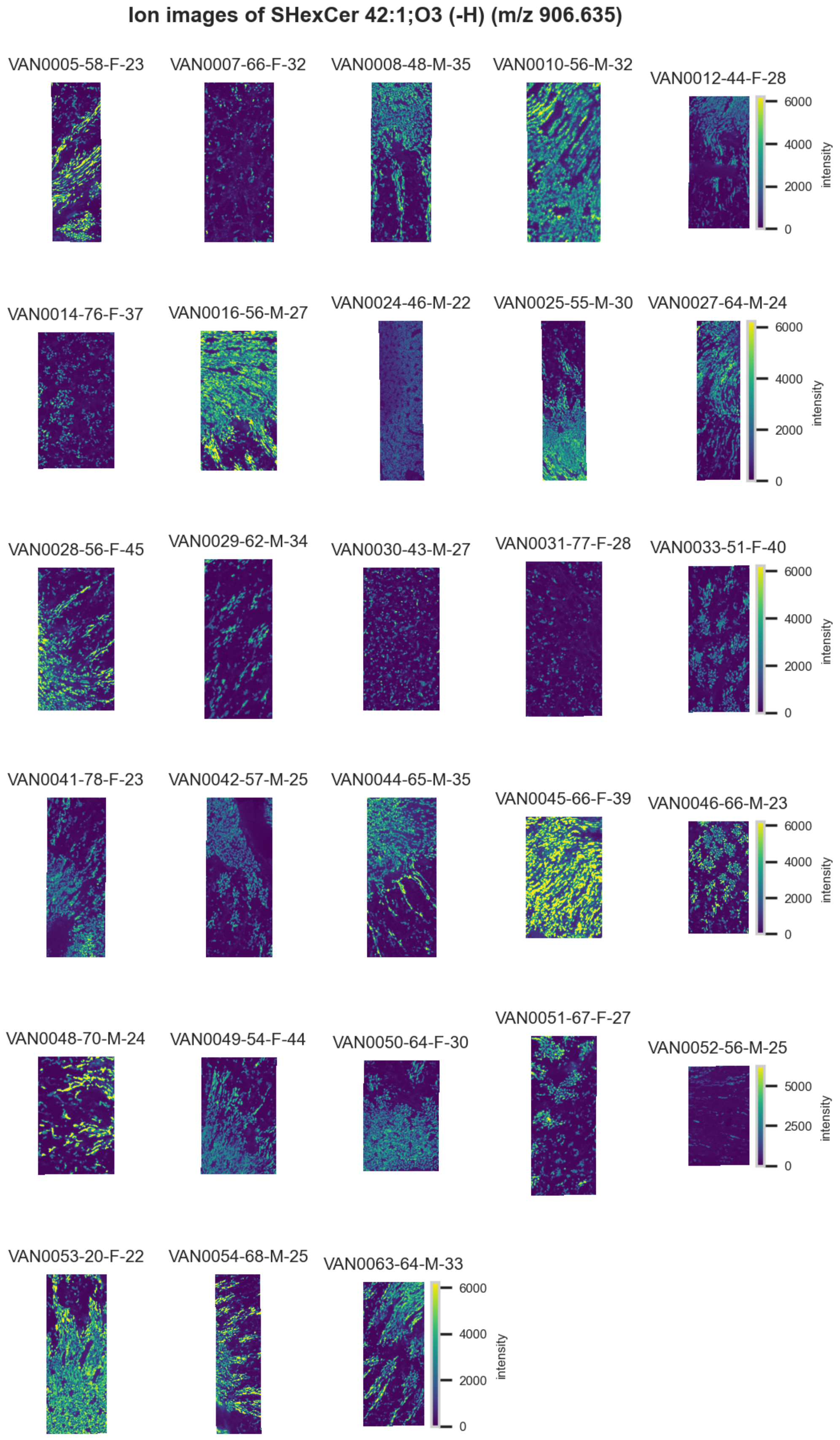

**Figure S110.** 10 µm MALDI IMS images of [SHexCer(42:1,3O)-H]$^-$ (*m/z* 906.635) detected in <u>negative ion mode</u> from each donor sample. Multivariate SHAP analysis determined this to be feature 2 for classification of distal tubules.

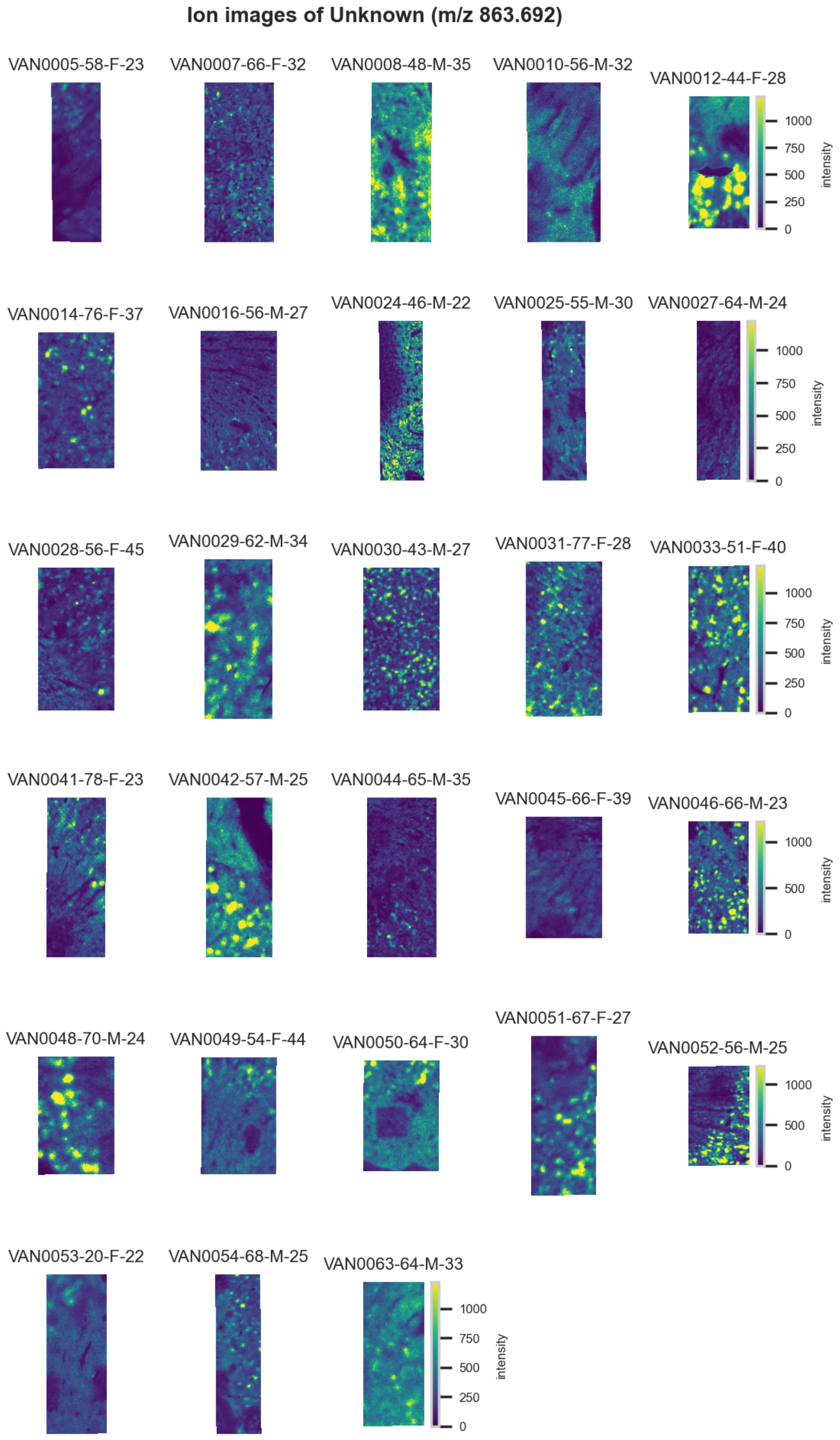

**Figure S111.** 10 µm MALDI IMS images of an unknown (*m/z* 863.692) detected in <u>negative ion mode</u> from each donor sample. Multivariate SHAP analysis determined this to be feature 3 for classification of distal tubules.

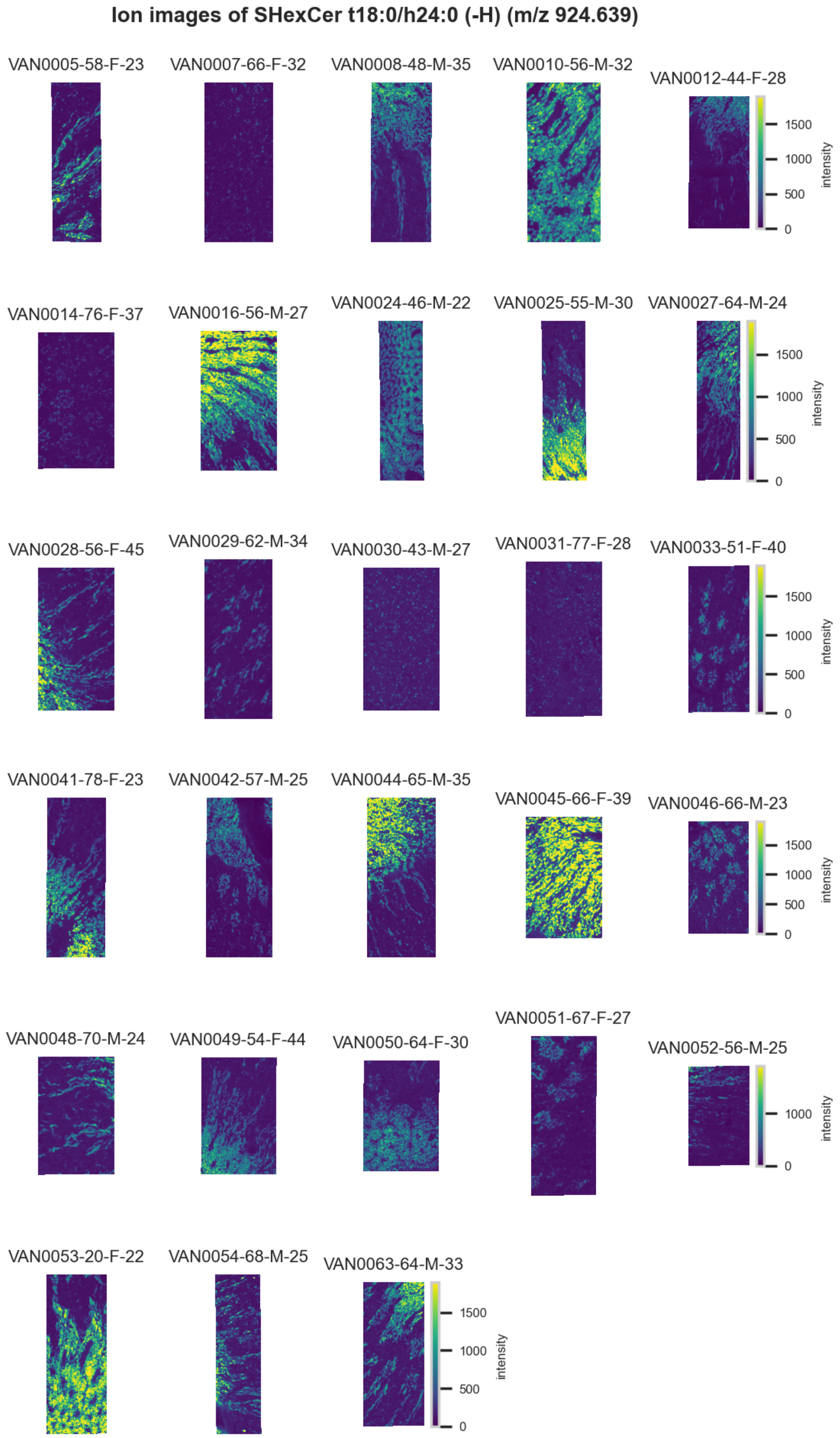

**Figure S112.** 10 µm MALDI IMS images of [SHexCer(t18:0/h24:0)-H]$^-$ (*m/z* 924.639) detected in negative ion mode from each donor sample. Multivariate SHAP analysis determined this to be feature 4 for classification of distal tubules.

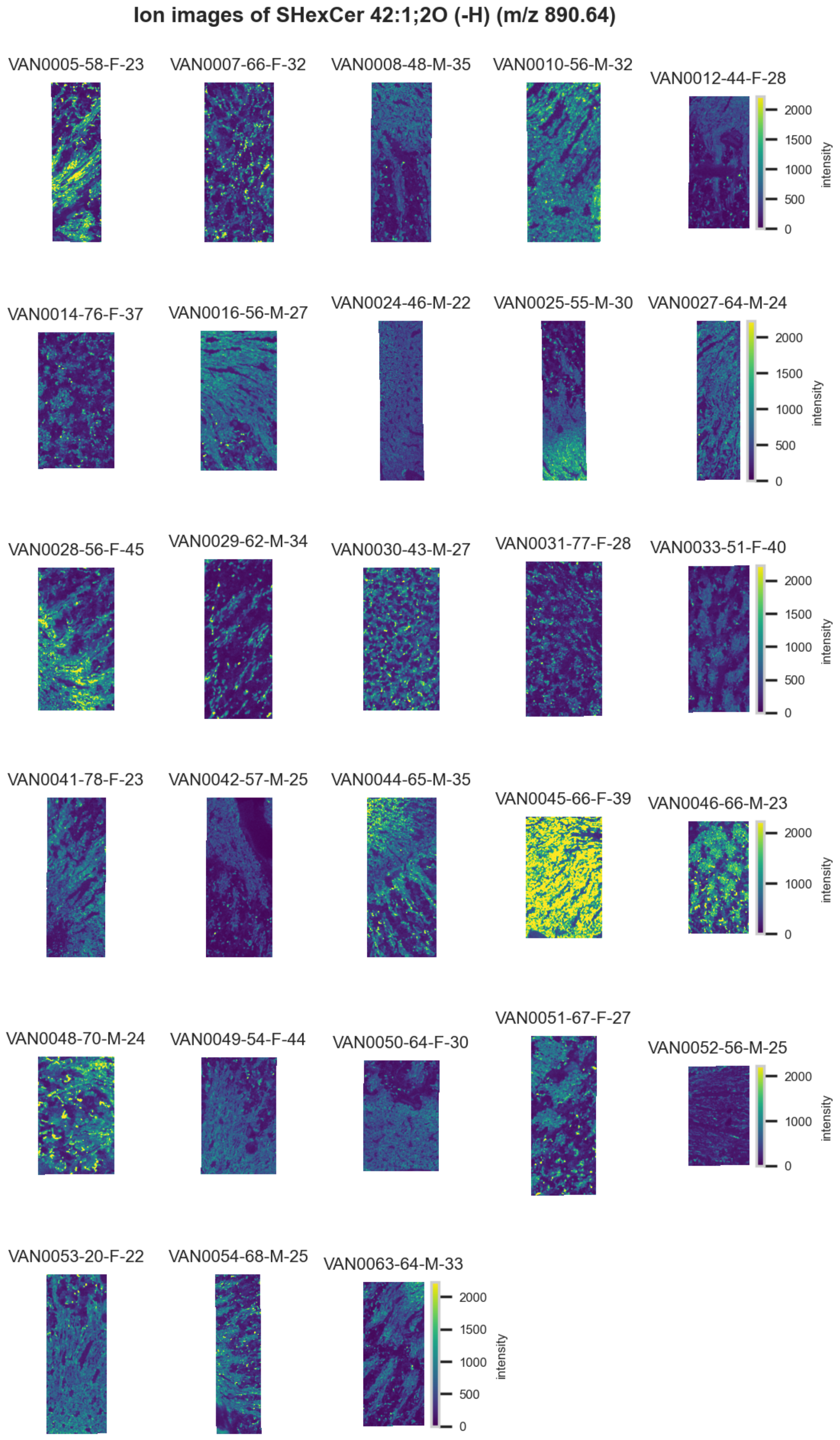

**Figure S113.** 10 µm MALDI IMS images of [SHexCer(42:1,2O)-CH3]$^-$ (*m/z* 890.64) detected in negative ion mode from each donor sample. Multivariate SHAP analysis determined this to be feature 5 for classification of distal tubules.

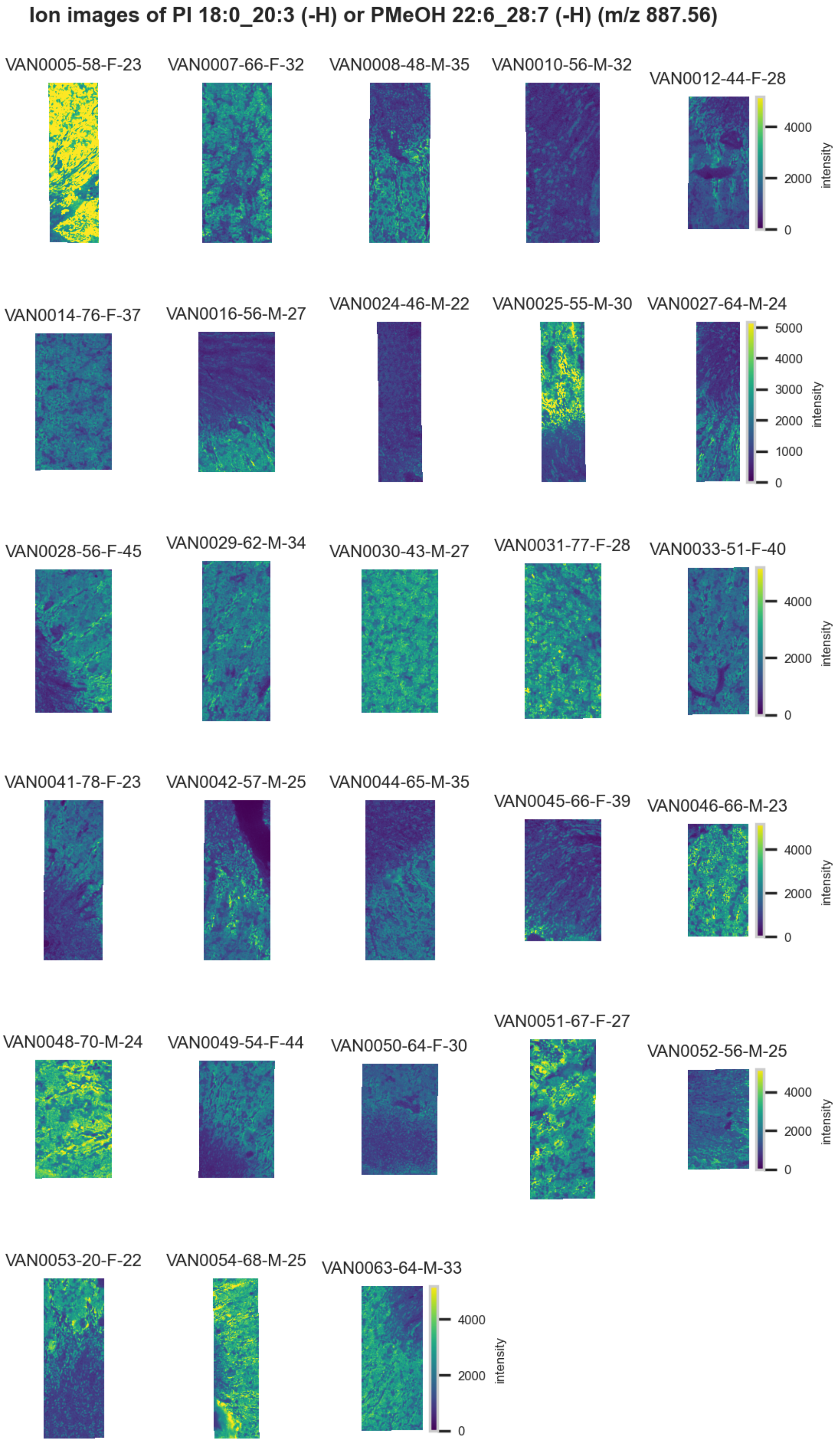

**Figure S114.** 10 µm MALDI IMS images of $[PI(18:0\_20:3)-H]^-$ or $PMeOH(22:6\_28:7)-H^-]^-$ (*m/z* 887.56) detected in <u>negative ion mode</u> from each donor sample. These molecules are isomers and therefore indistinguishable by MS alone. Multivariate SHAP analysis determined this to be feature 6 for classification of distal tubules.

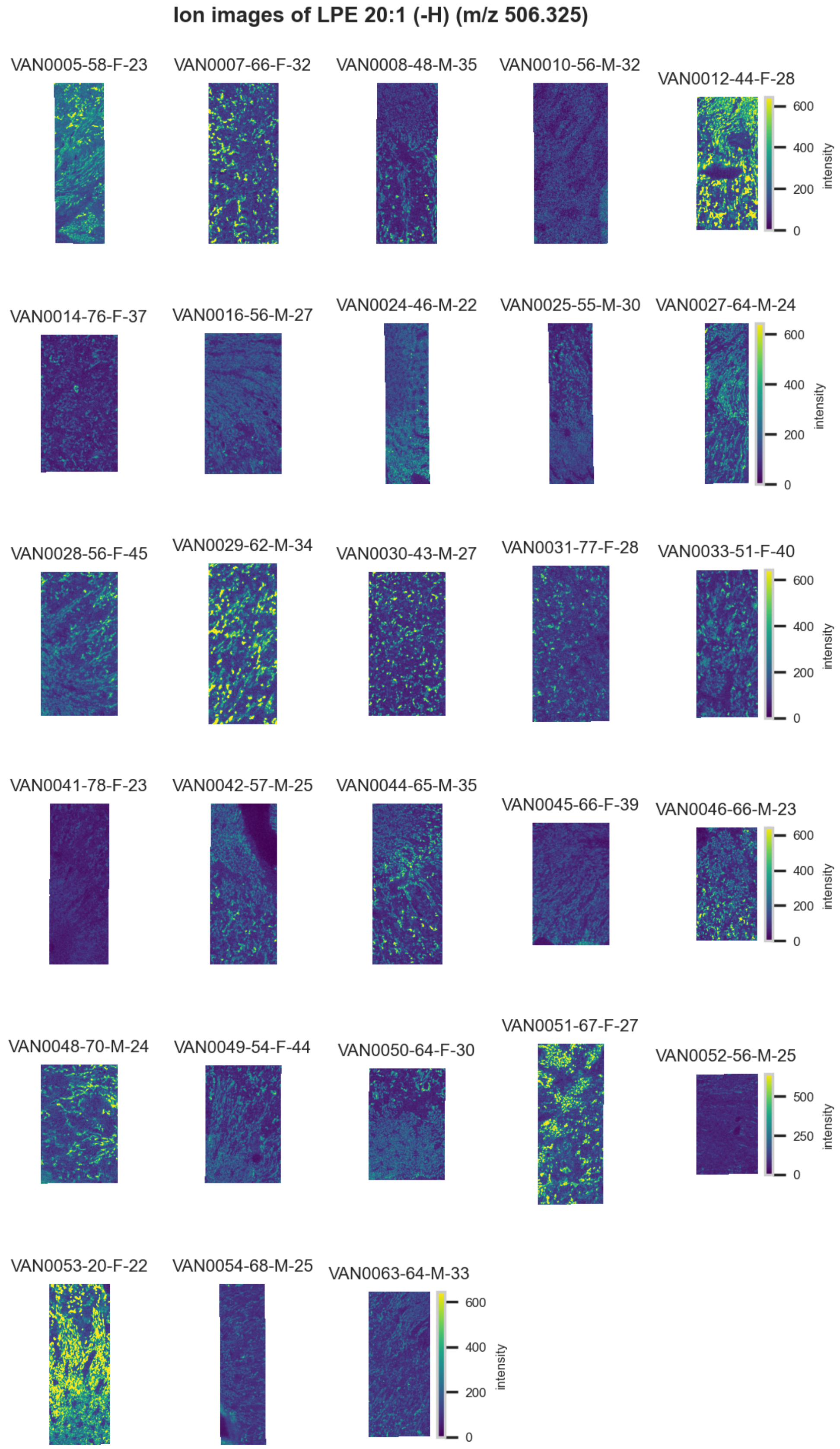

**Figure S115.** 10 µm MALDI IMS images of $[LPE(20:1)-H]^-$ (*m/z* 506.325) detected in <u>negative ion mode</u> from each donor sample. Multivariate SHAP analysis determined this to be feature 7 for classification of distal tubules.

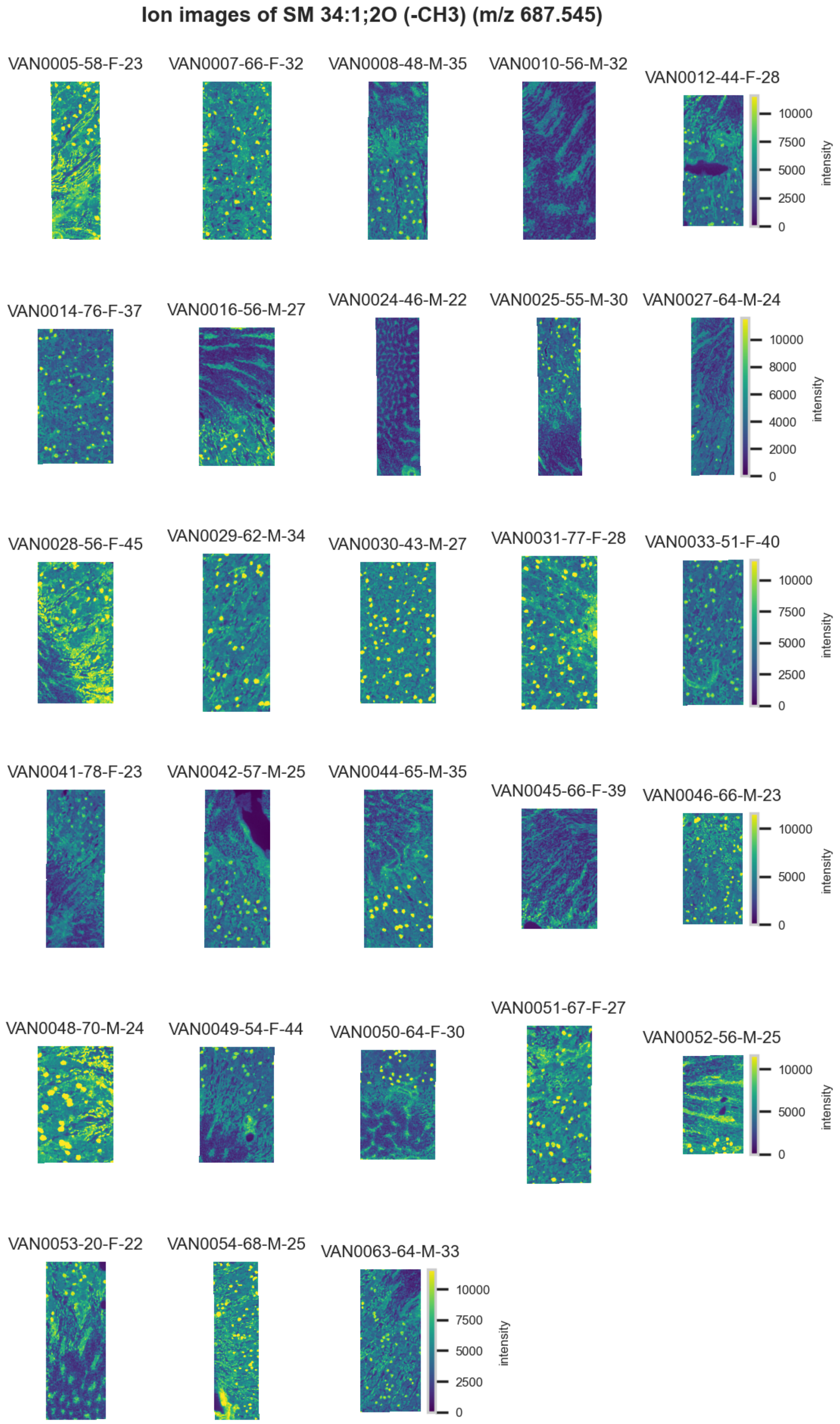

**Figure S116.** 10 µm MALDI IMS images of [SM(34:1,2O)-$CH_3$]$^-$ (*m/z* 687.545) detected in negative ion mode from each donor sample. Multivariate SHAP analysis determined this to be feature 8 for classification of distal tubules.

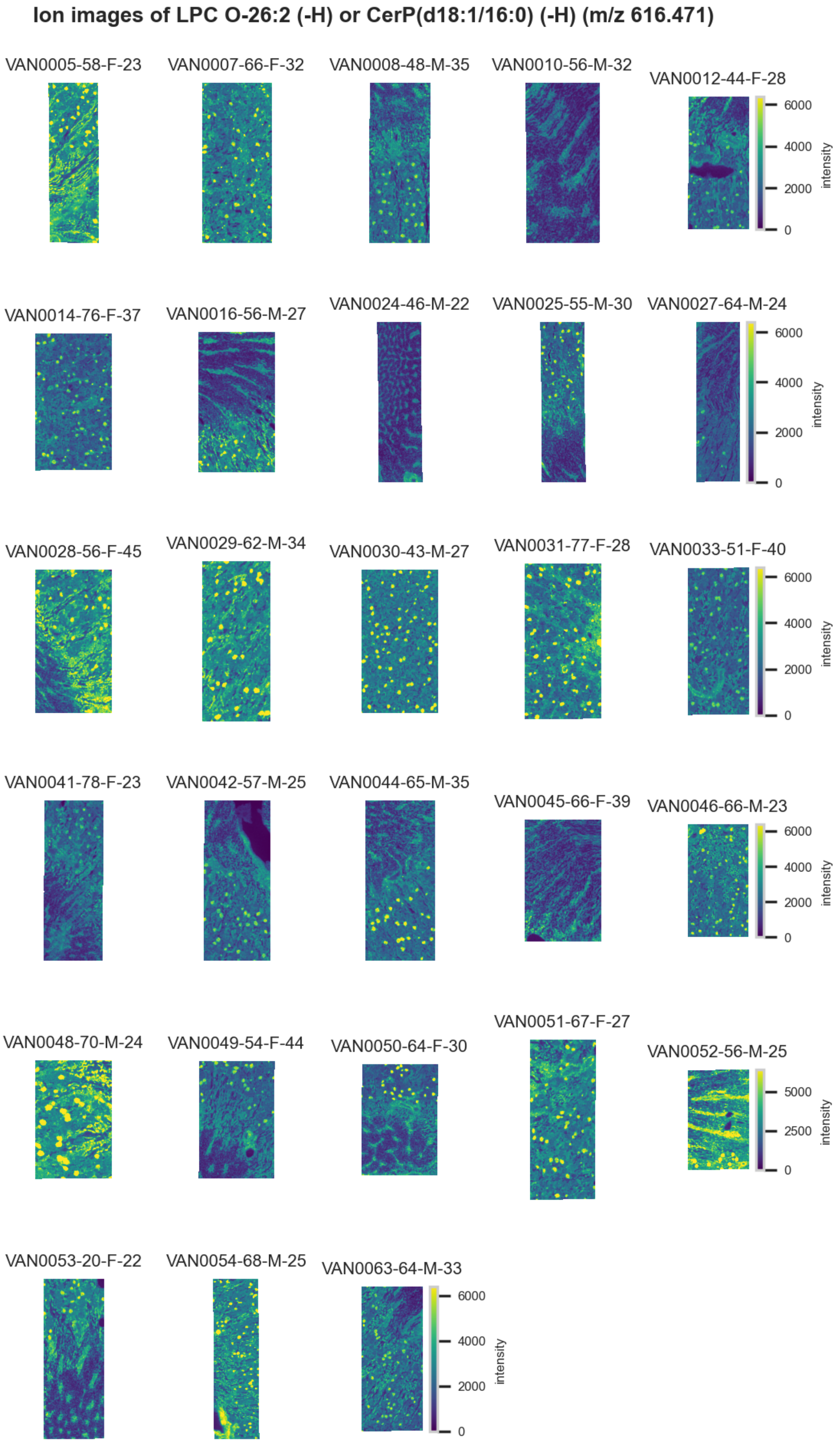

**Figure S117.** 10 µm MALDI IMS images of [LPC(O-26:2)-H]$^-$ or [CerP(d18:1/16:0)-H]$^-$ (*m/z* 616.471) detected in negative ion mode from each donor sample. These molecules are isomers and therefore indistinguishable by MS alone. Multivariate SHAP analysis determined this to be feature 9 for classification of distal tubules.

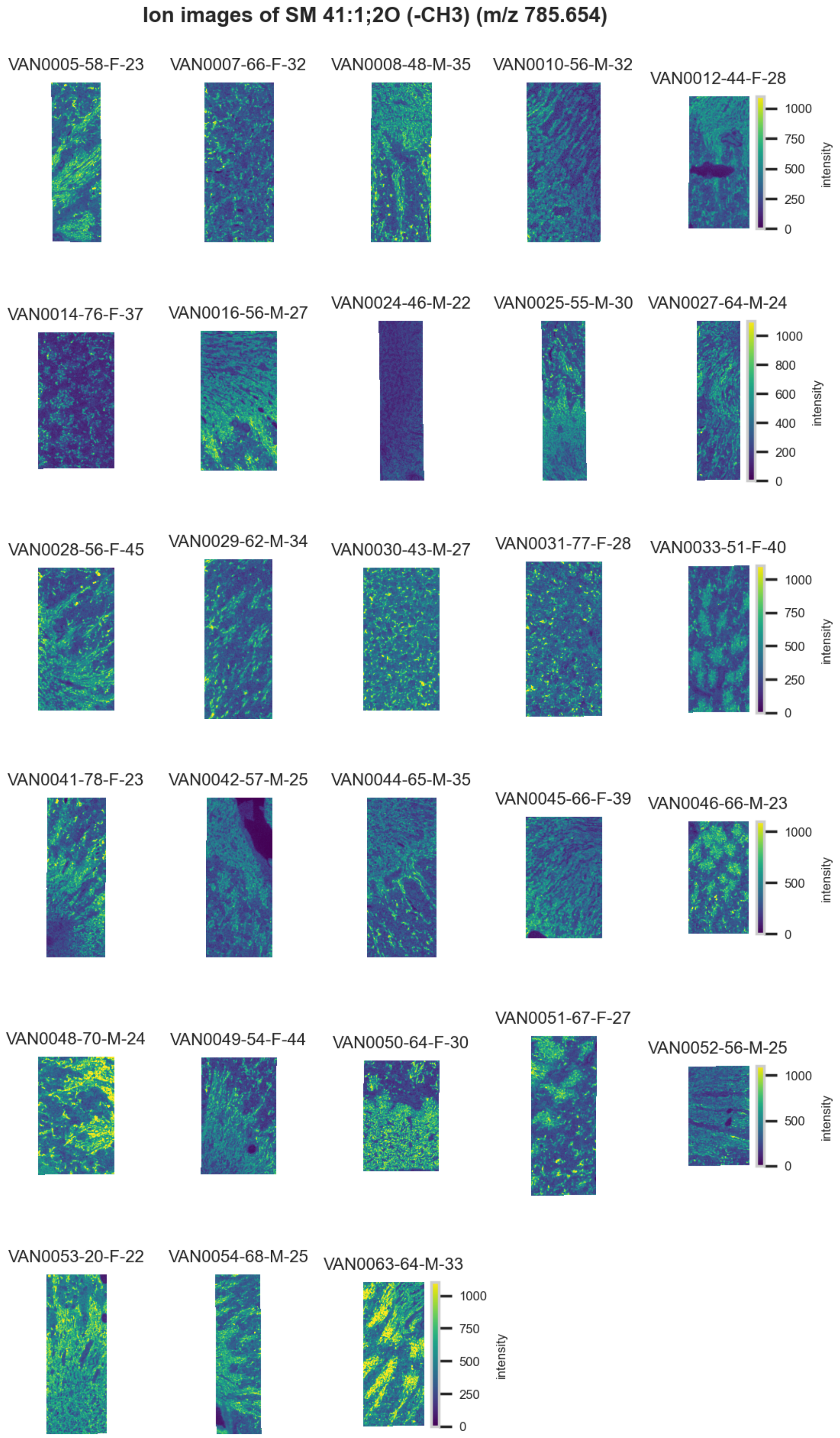

**Figure S118.** 10 µm MALDI IMS images of [SM(41:1,2O)-$CH_3$]$^-$ (*m/z* 785.654) detected in negative ion mode from each donor sample. Multivariate SHAP analysis determined this to be feature 10 for classification of distal tubules.

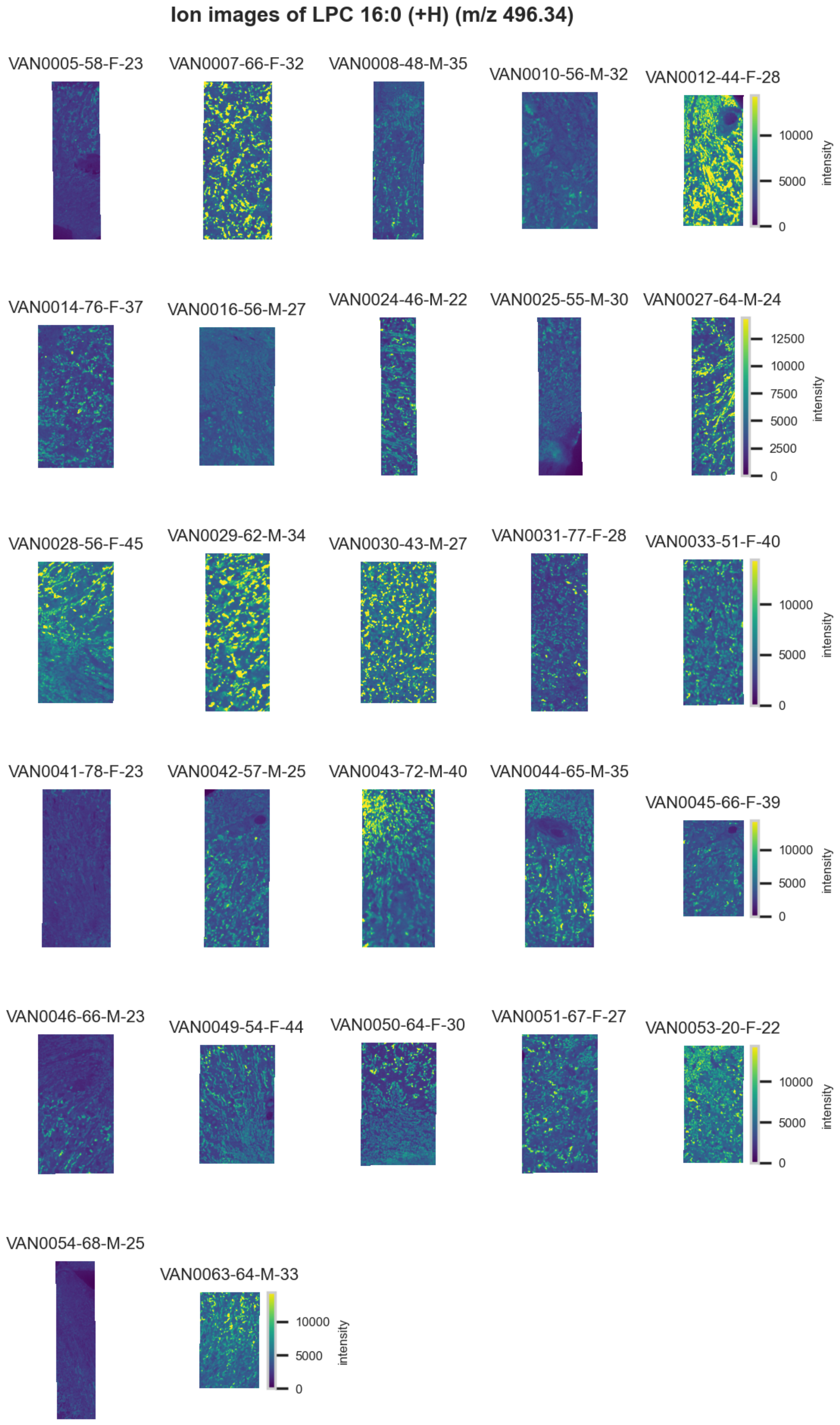

**Figure S119.** 10 µm MALDI IMS images of $[LPC(16:0)+H]^+$ (*m/z* 496.34) detected in positive ion mode from each donor sample. These molecules are isomers and therefore indistinguishable by MS alone. Multivariate SHAP analysis determined this to be feature 1 for classification of distal tubules.

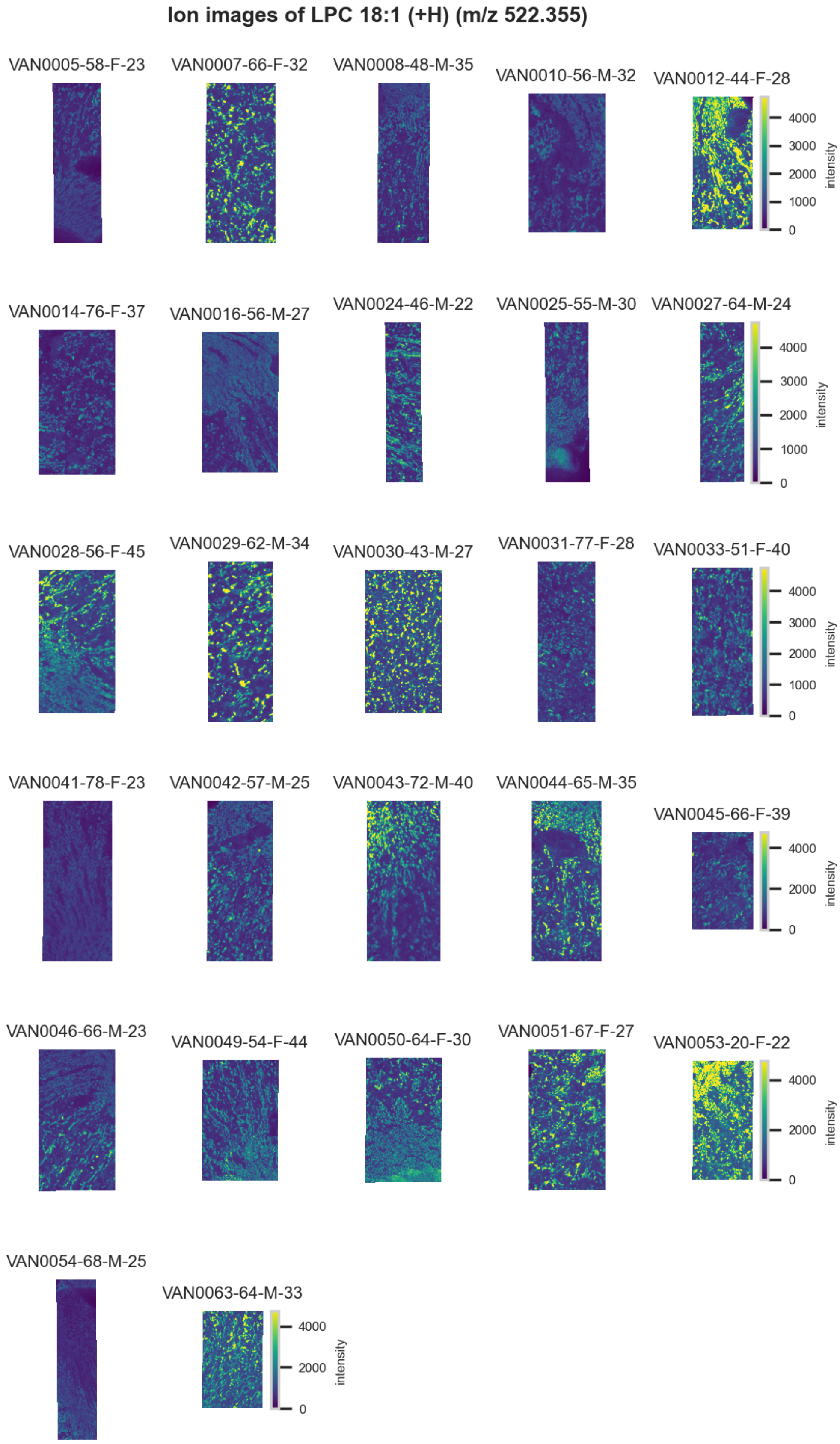

**Figure S120.** 10 μm MALDI IMS images of $[LPC(18:1)+H]^+$ (*m/z* 522.355) detected in <u>positive ion mode</u> from each donor sample. Multivariate SHAP analysis determined this to be feature 2 for classification of distal tubules.

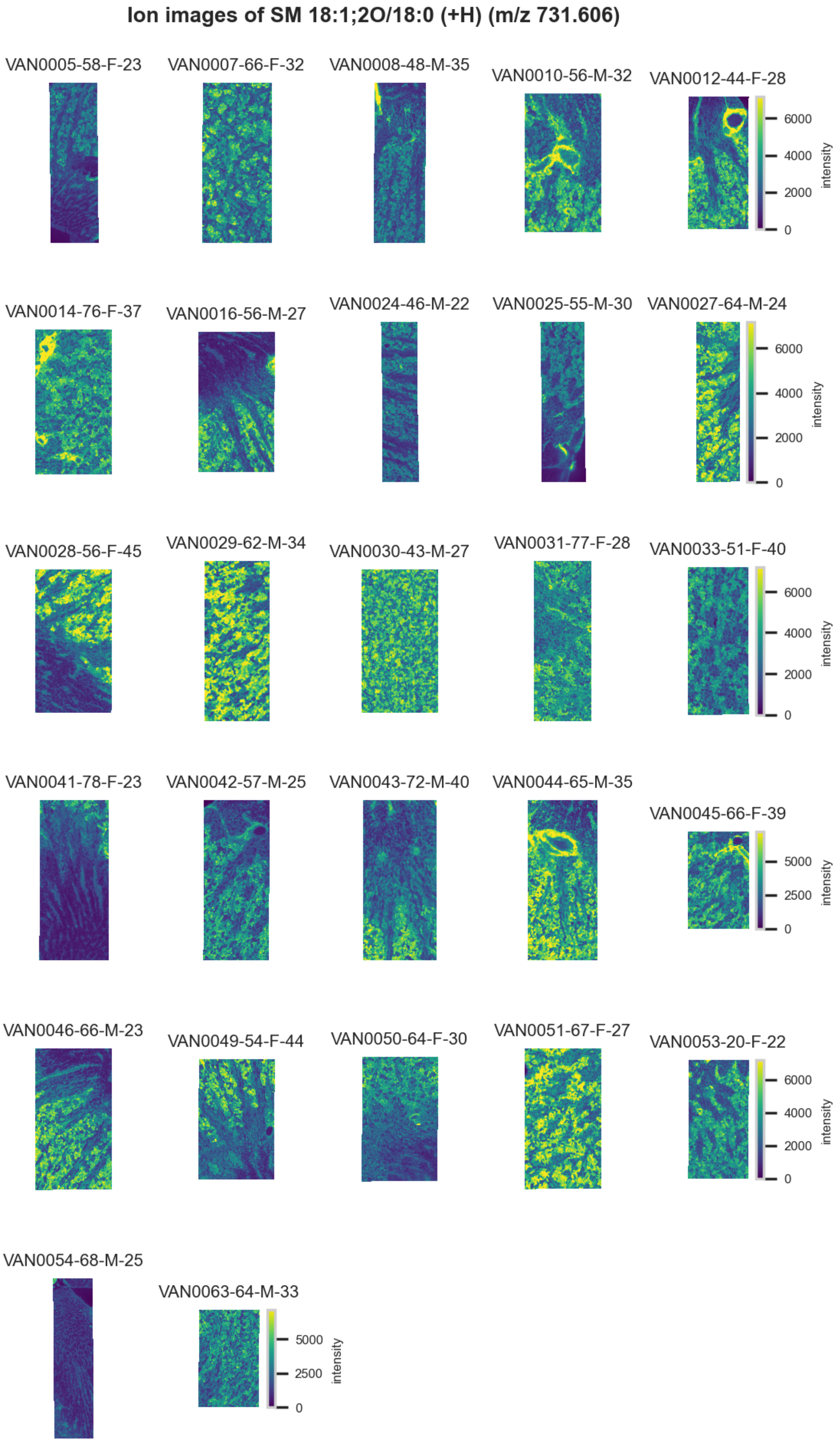

**Figure S121.** 10 µm MALDI IMS images of $[SM(18:1,2O/18:0)+H]^+$ (*m/z* 731.606) detected in positive ion mode from each donor sample. Multivariate SHAP analysis determined this to be feature 3 for classification of distal tubules.

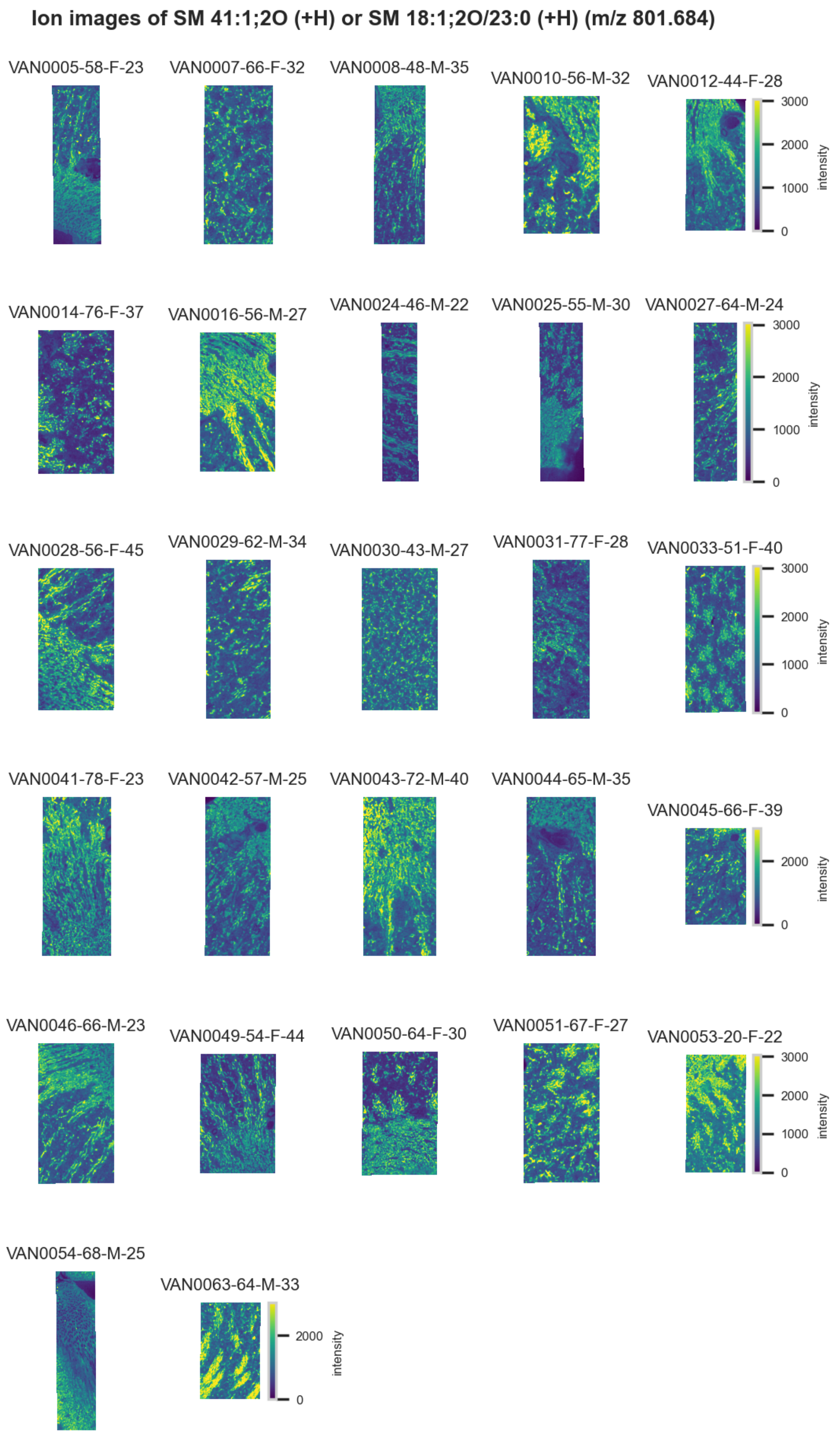

**Figure S122.** 10 µm MALDI IMS images of $[SM(41:1,2O)+H]^+$ or $[SM(18:1,2O/23:0)+H]^+$ (*m/z* 801.684) detected in positive ion mode from each donor sample. These molecules are isomers and therefore indistinguishable by MS alone. Multivariate SHAP analysis determined this to be feature 4 for classification of distal tubules.

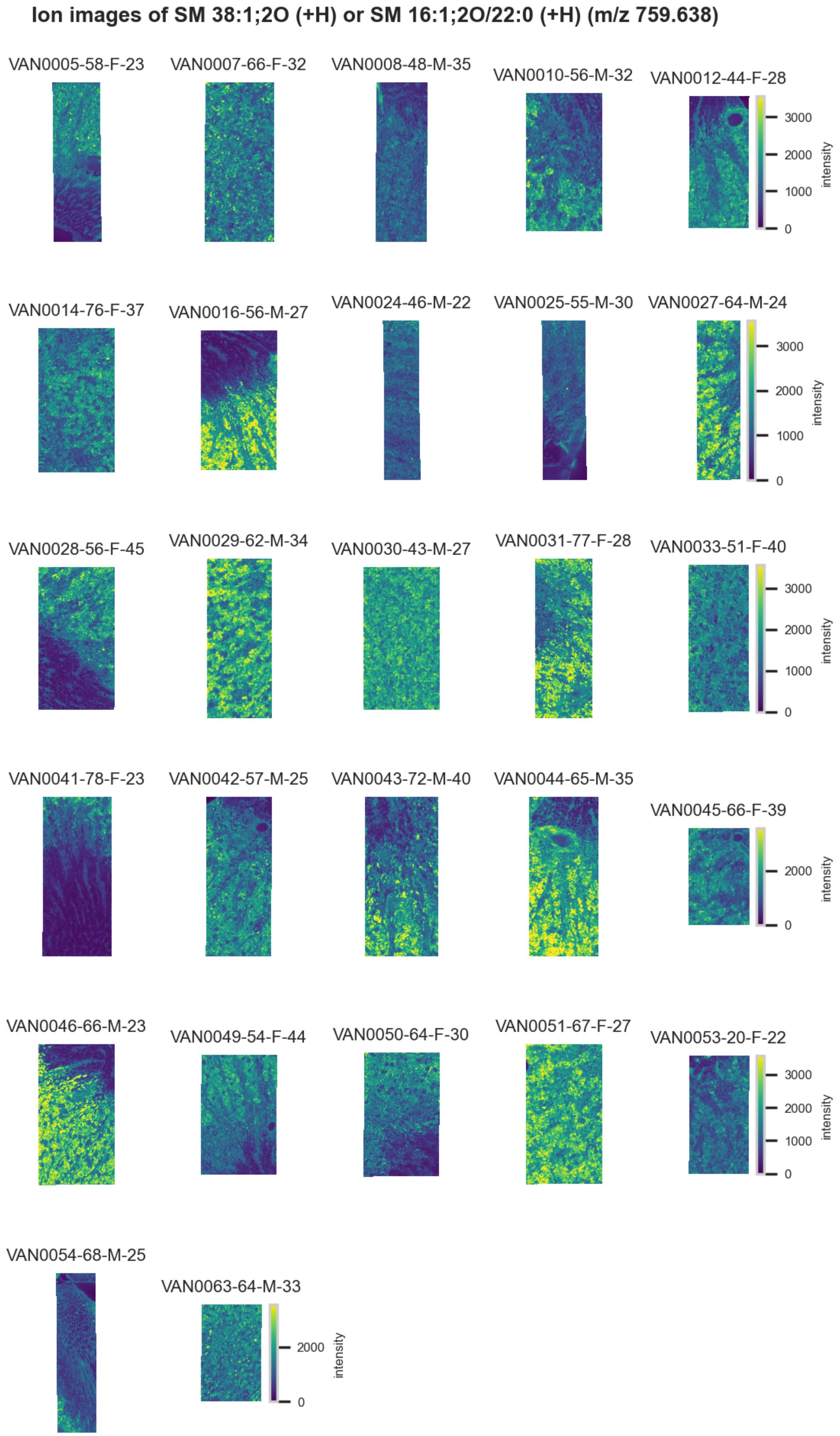

**Figure S123.** 10 µm MALDI IMS images of $[SM(38:1,2O)+H]^+$ or $[SM(16:1,2O/22:0)+H]^+$ (*m/z* 759.638) detected in <u>positive ion mode</u> from each donor sample. Multivariate SHAP analysis determined this to be feature 5 for classification of distal tubules.

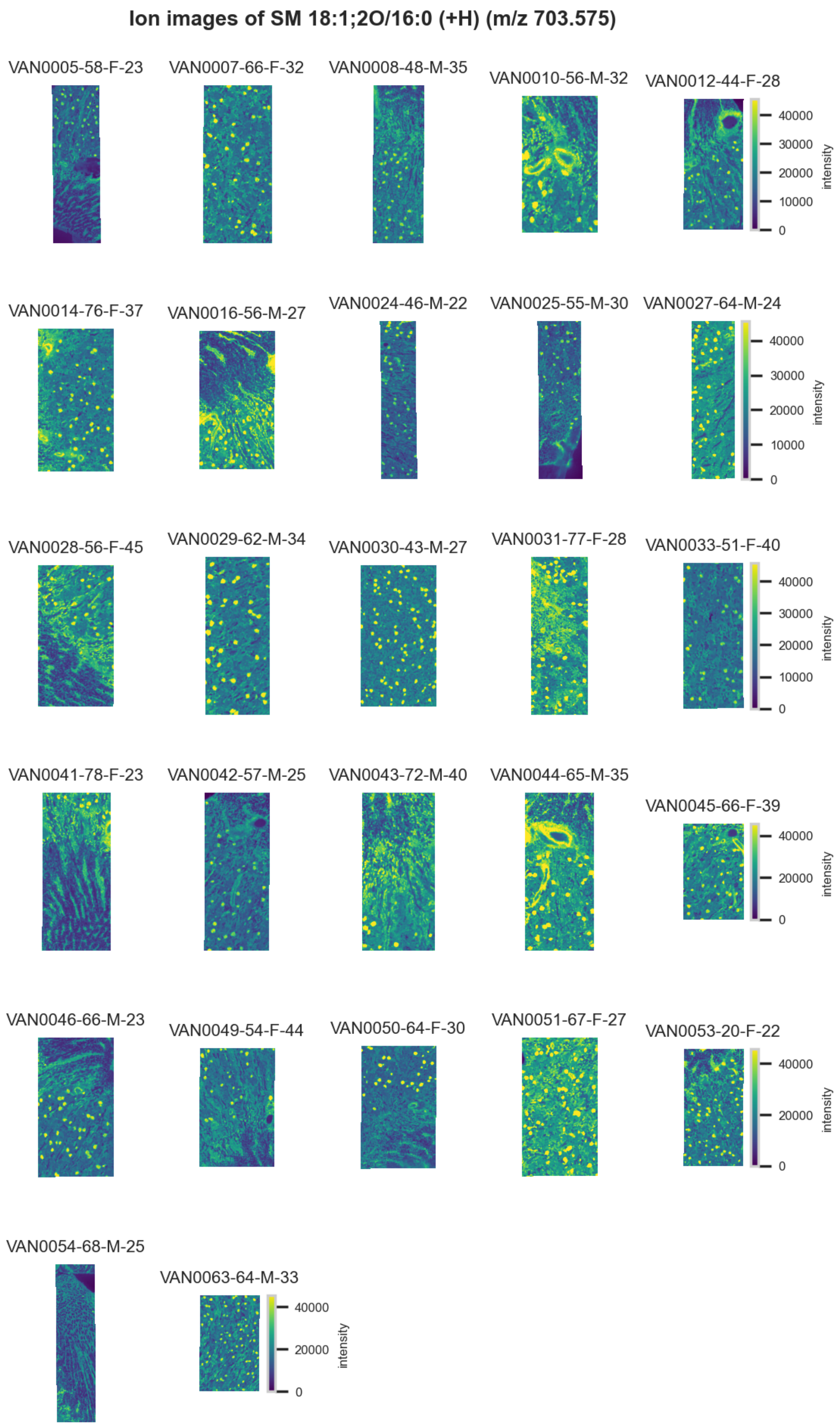

**Figure S124.** 10 µm MALDI IMS images of $[SM(18:1,2O/16:0)+H]^+$ (*m/z* 703.575) detected in positive ion mode from each donor sample. These molecules are isomers and therefore indistinguishable by MS alone. Multivariate SHAP analysis determined this to be feature 6 for classification of distal tubules.

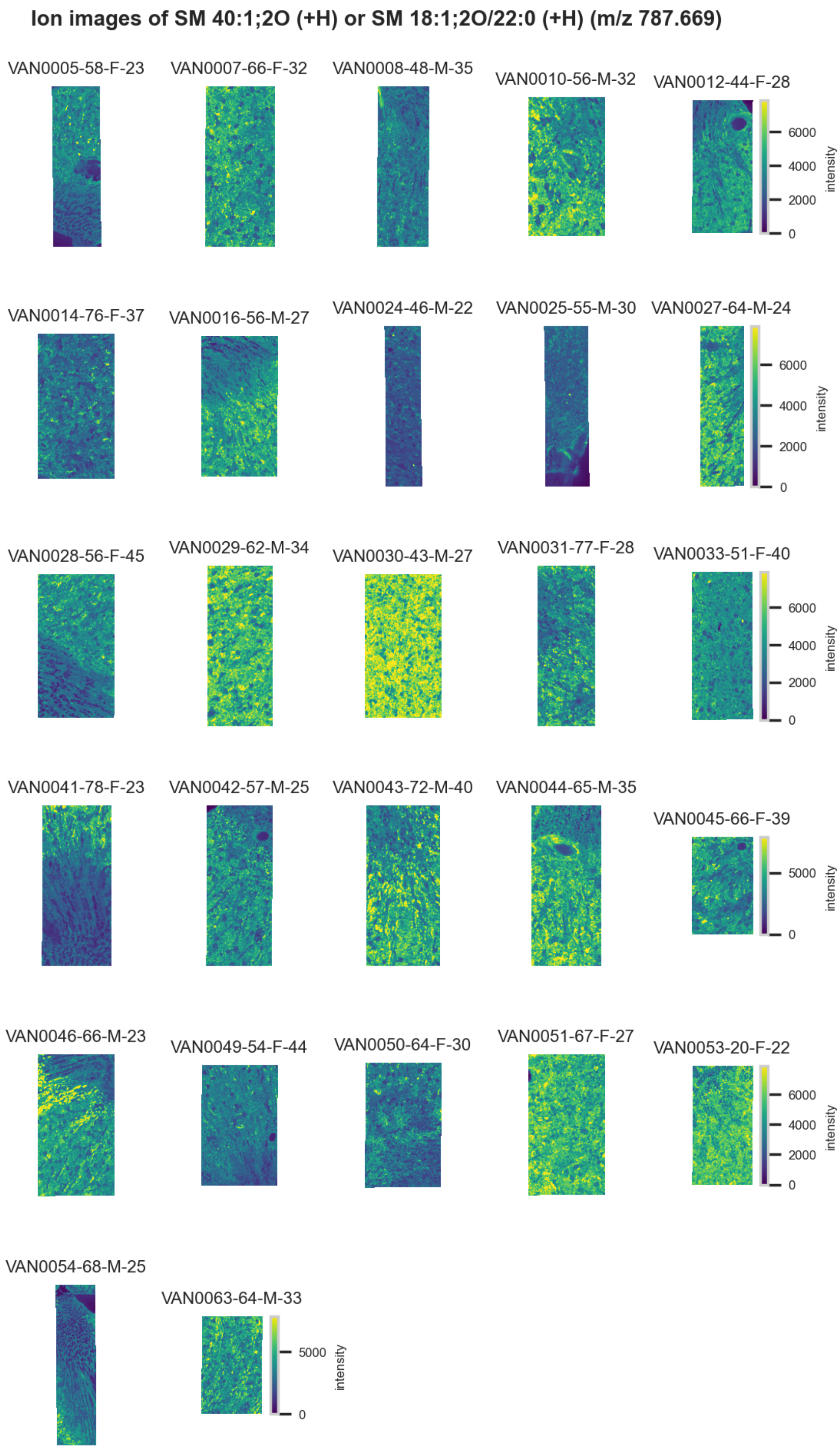

**Figure S125.** 10 µm MALDI IMS images of $[SM(40:1,2O)+H]^+$ or $[SM(18:1,2O/22:0)+H]^+$ (*m/z* 787.669) detected in positive ion mode from each donor sample. These molecules are isomers and therefore indistinguishable by MS alone. Multivariate SHAP analysis determined this to be feature 7 for classification of distal tubules.

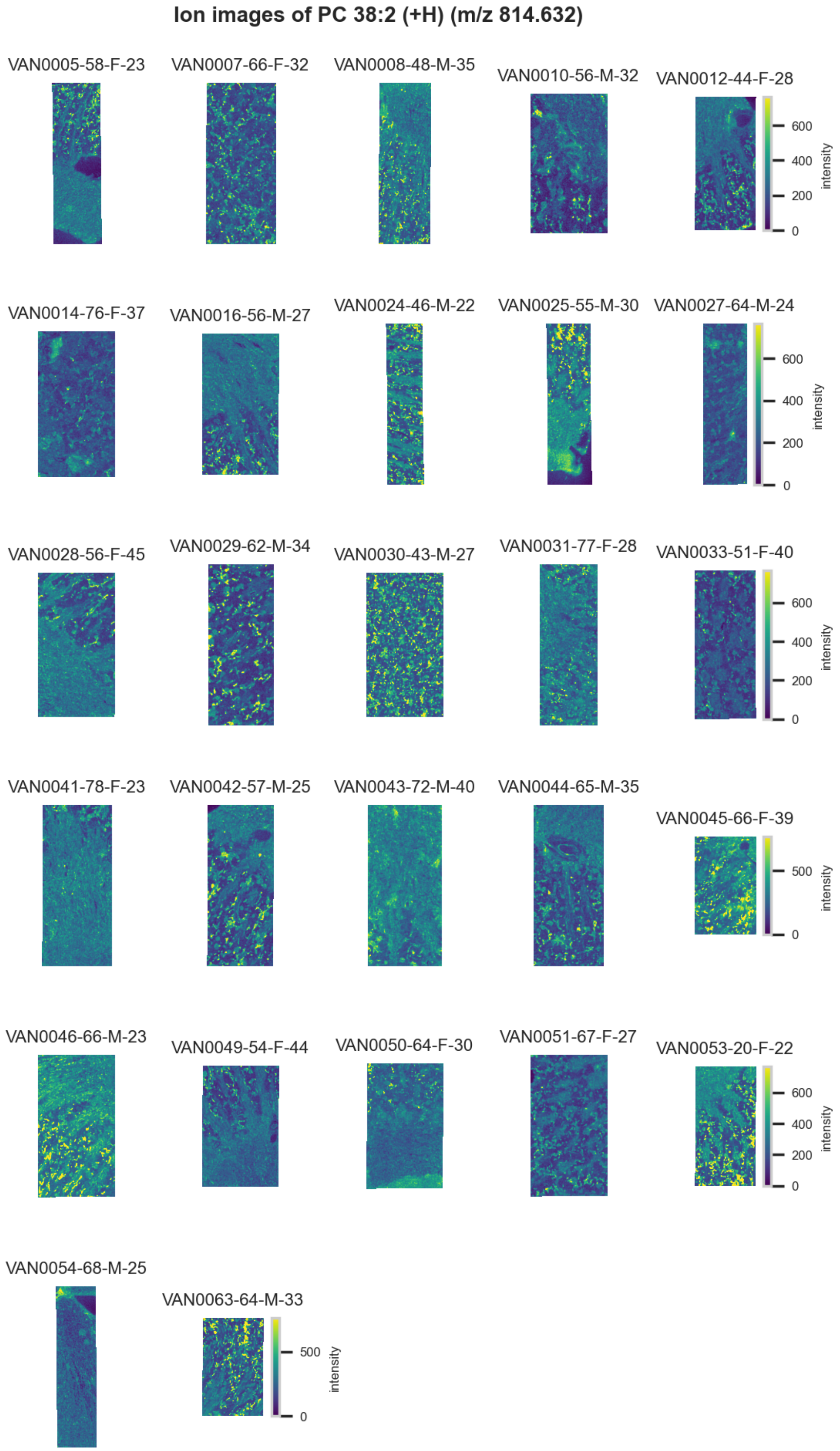

**Figure S126.** 10 µm MALDI IMS images of $[PC(38:2)+H]^+$ (*m/z* 814.632) detected in positive ion mode from each donor sample. Multivariate SHAP analysis determined this to be feature 8 for classification of distal tubules.

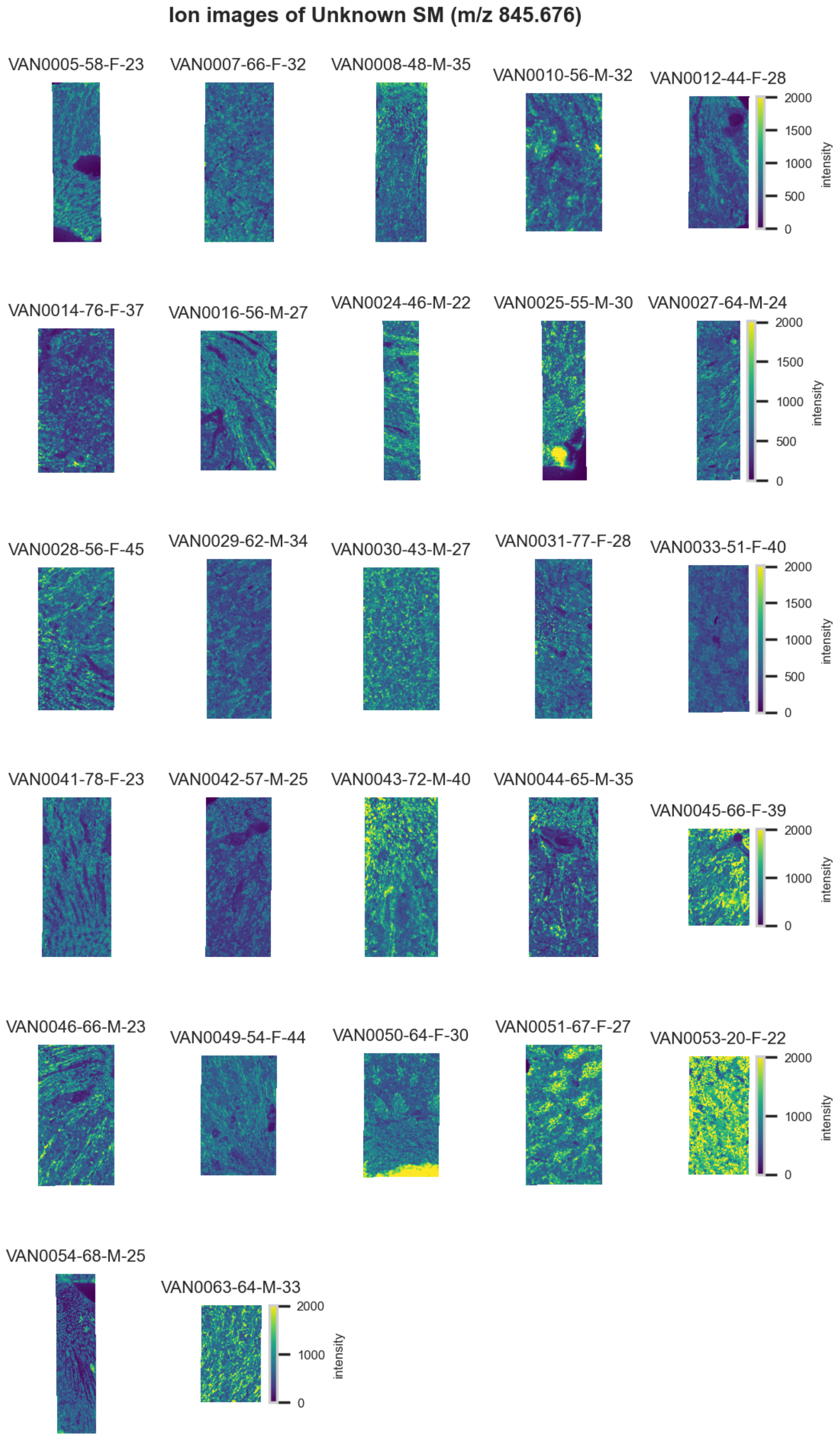

**Figure S127.** 10 µm MALDI IMS images of an unknown SM (*m/z* 845.676) detected in positive ion mode from each donor sample. Multivariate SHAP analysis determined this to be feature 9 for classification of distal tubules.

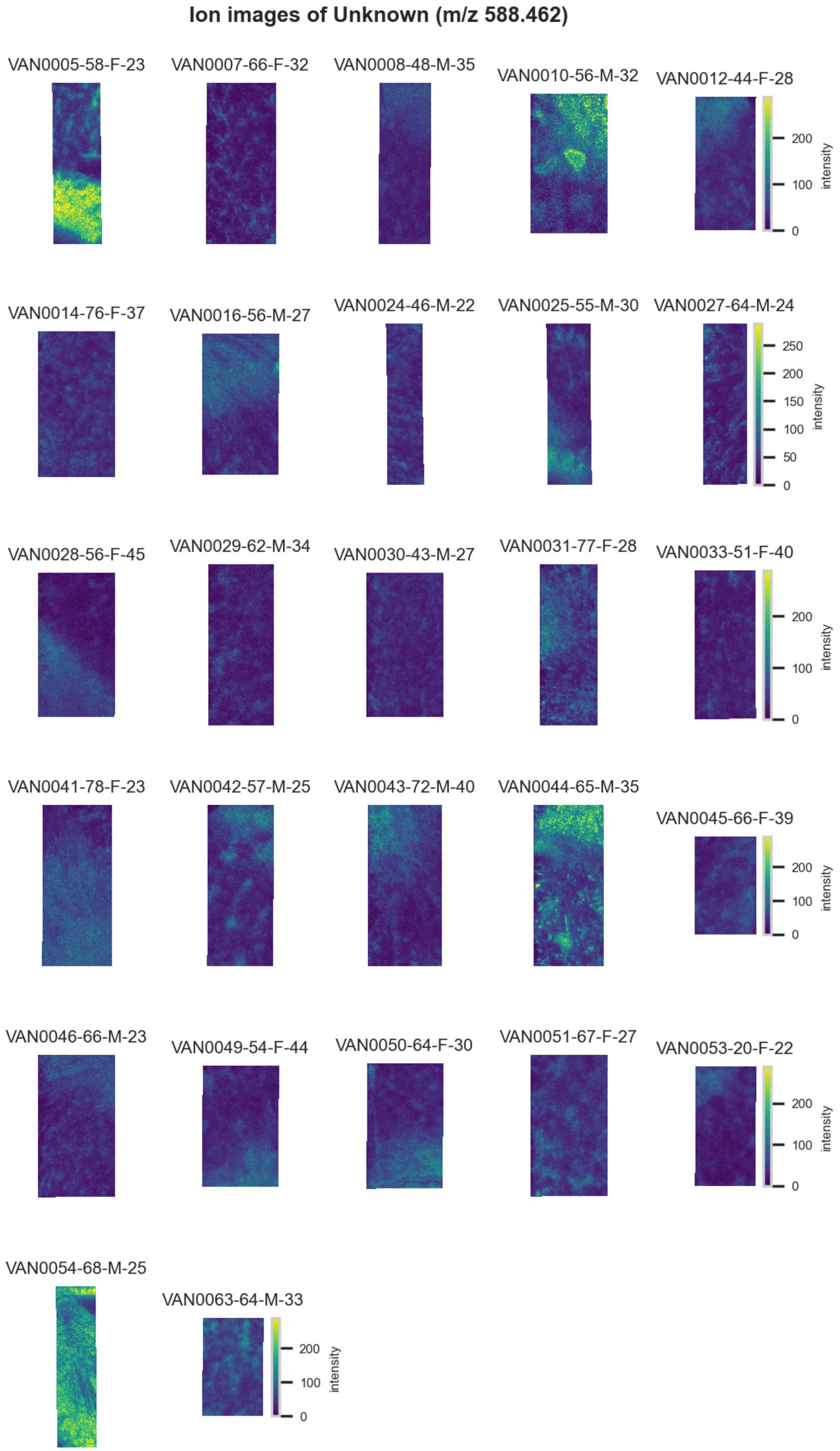

**Figure S128.** 10 µm MALDI IMS images of an unknown (*m/z* 588.462) detected in positive ion mode from each donor sample. These molecules are isomers and therefore indistinguishable by MS alone. Multivariate SHAP analysis determined this to be feature 10 for classification of distal tubules.

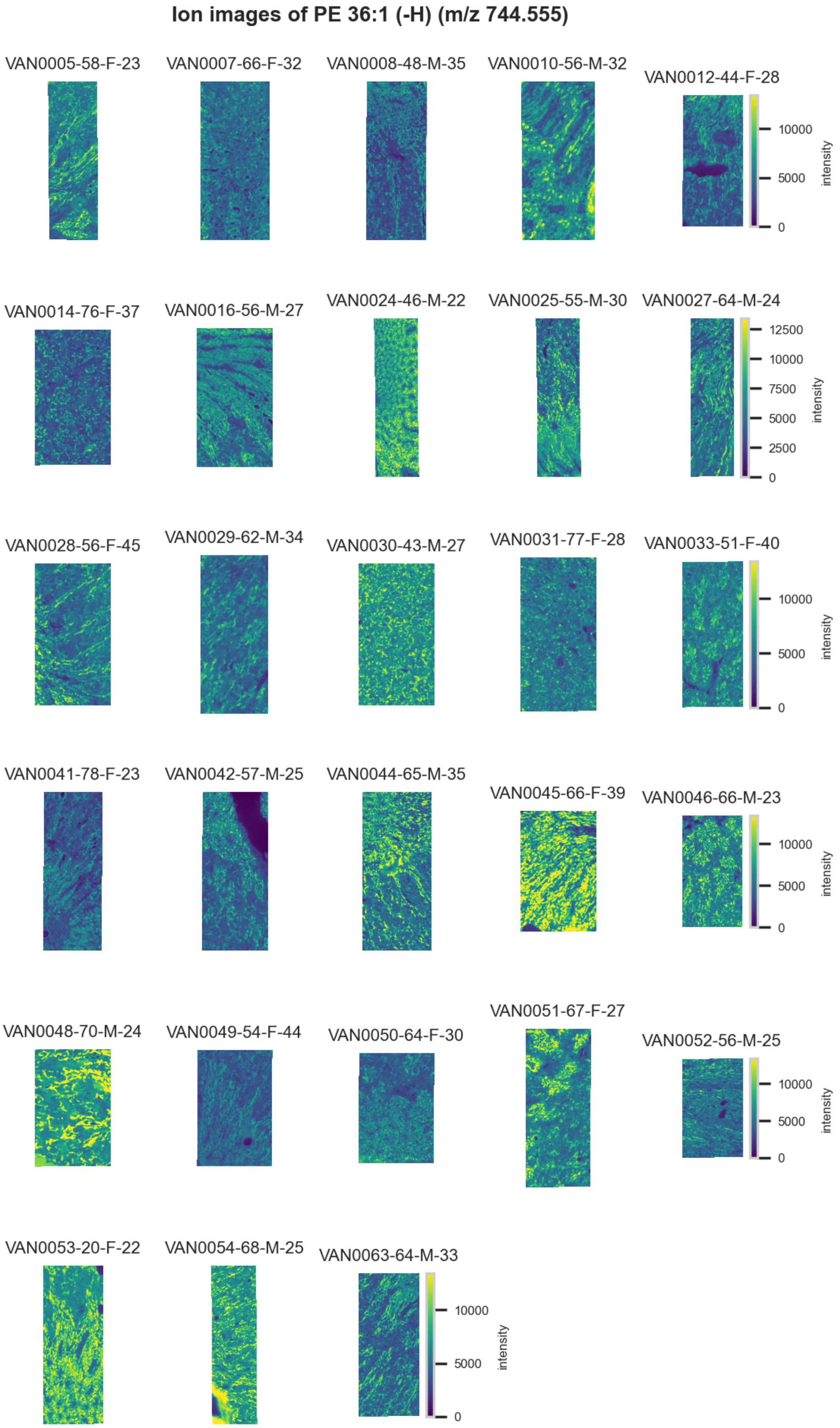

**Figure S129.** 10 µm MALDI IMS images of $[PE(36:1)-H]^-$ (*m/z* 744.555) detected in negative ion mode from each donor sample. Multivariate SHAP analysis determined this to be feature 1 for classification of collecting ducts.

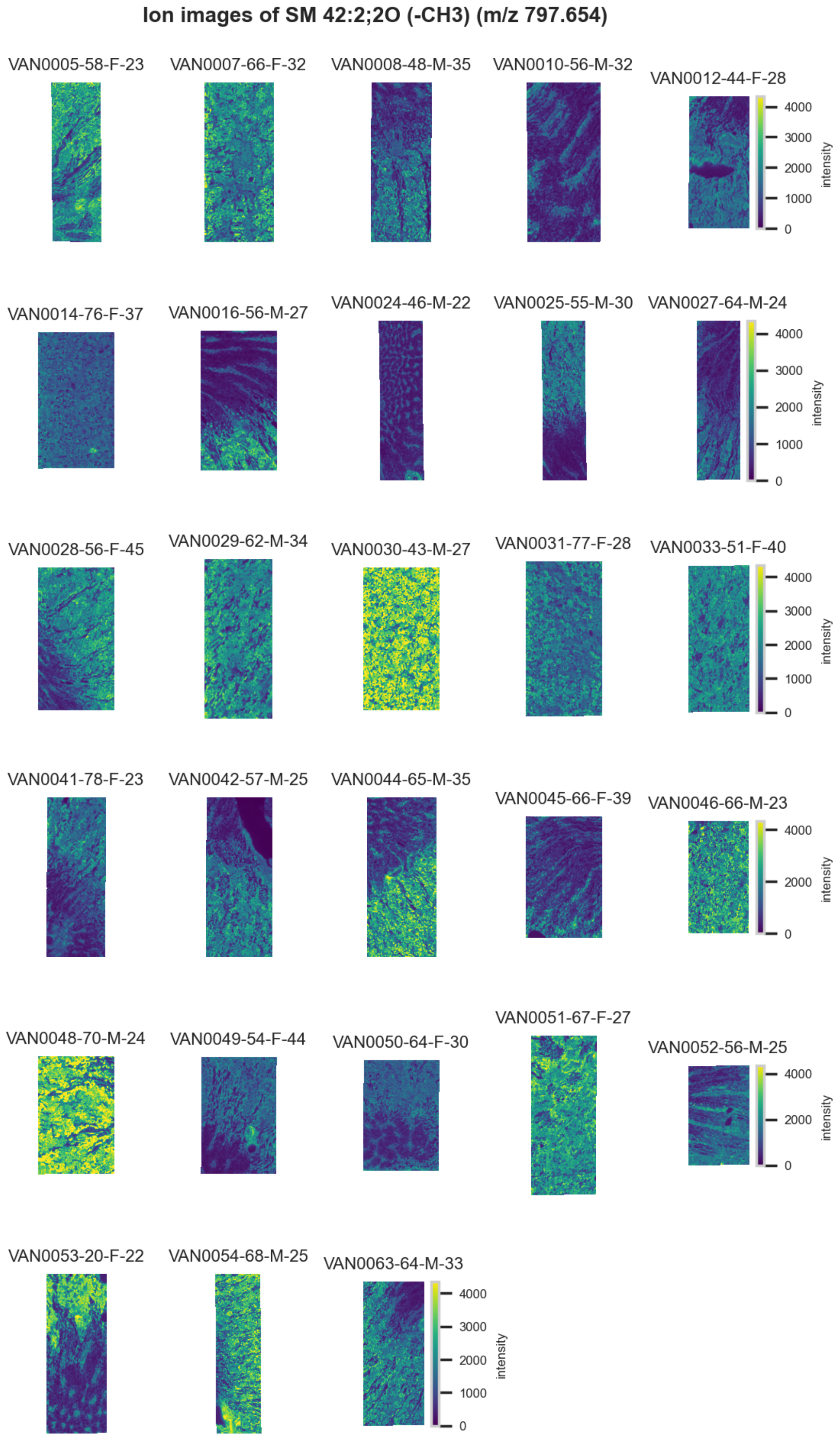

**Figure S130.** 10 μm MALDI IMS images of [SM(42:2,2O)-$CH_3$]$^-$ (*m/z* 797.654) detected in negative ion mode from each donor sample. Multivariate SHAP analysis determined this to be feature 2 for classification of collecting ducts.

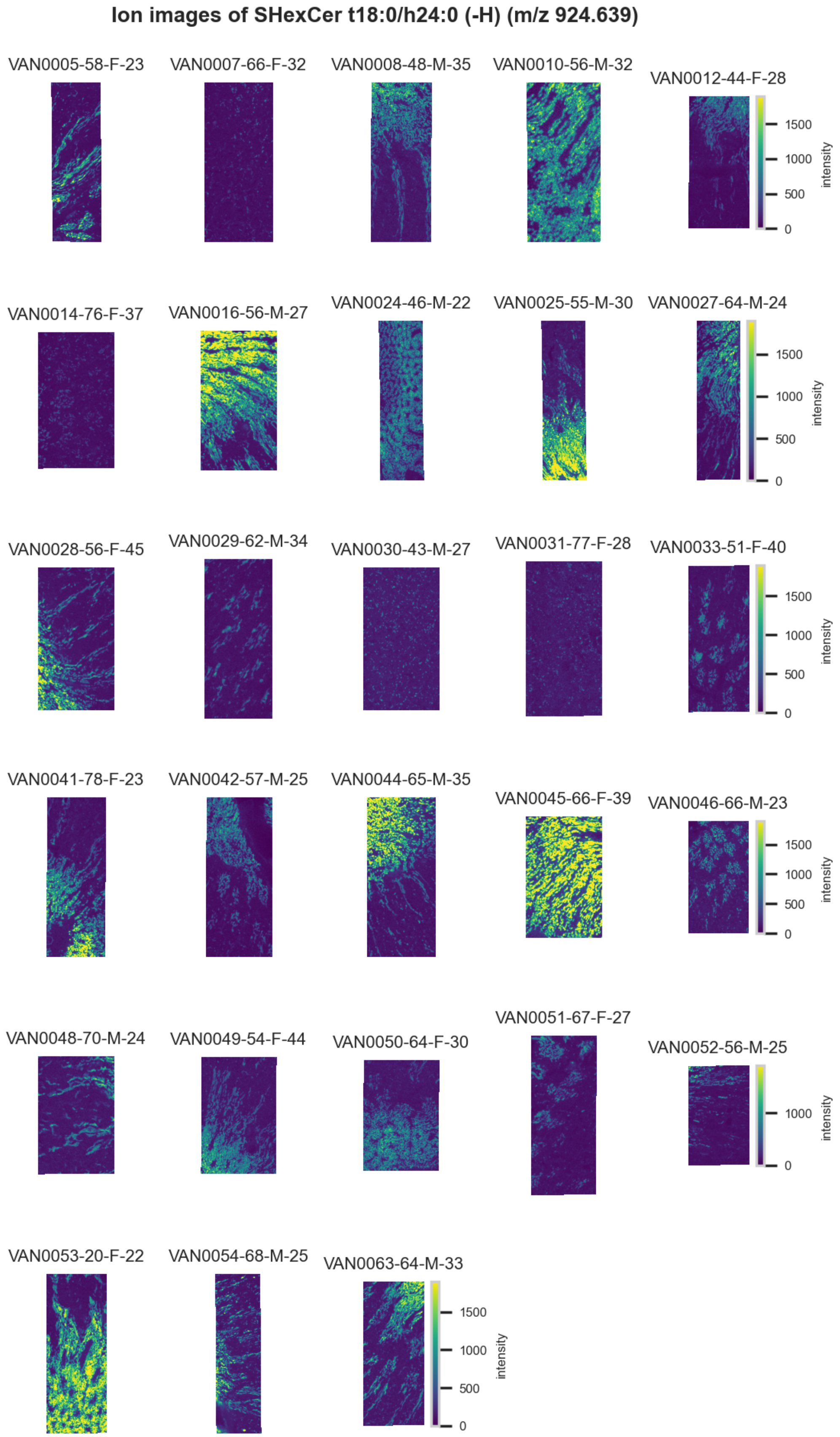

**Figure S131.** 10 µm MALDI IMS images of [SHexCer(t18:0/h24:0)-H]$^-$ (*m/z* 924.639) detected in negative ion mode from each donor sample. Multivariate SHAP analysis determined this to be feature 3 for classification of collecting ducts.

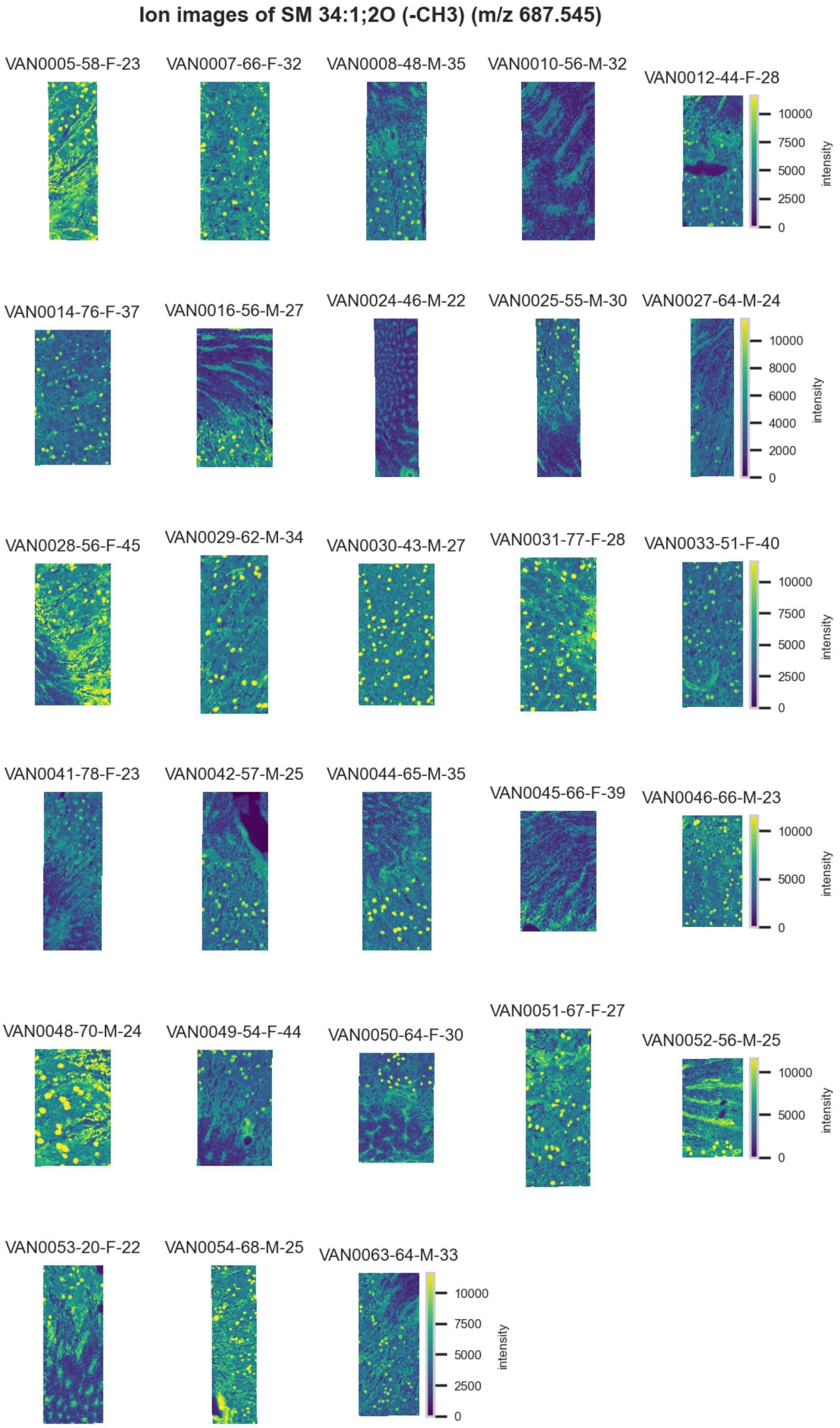

**Figure S132.** 10 µm MALDI IMS images of $[SM(34:1,2O)-CH_3]^-$ (*m/z* 687.545) detected in negative ion mode from each donor sample. Multivariate SHAP analysis determined this to be feature 4 for classification of collecting ducts.

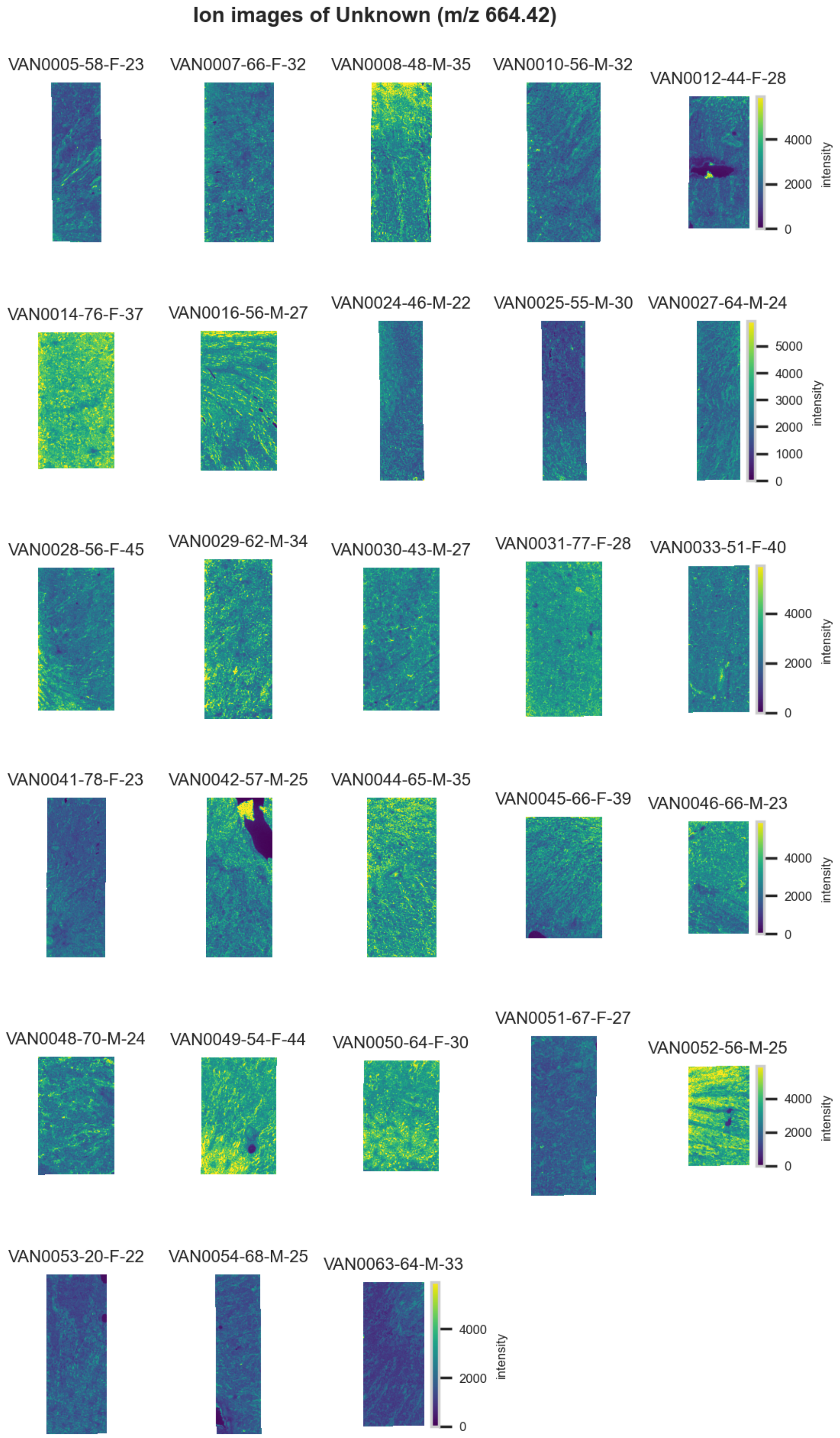

**Figure S133.** 10 µm MALDI IMS images of an unknown (*m/z* 664.42) detected in negative ion mode from each donor sample. Multivariate SHAP analysis determined this to be feature 5 for classification of collecting ducts.

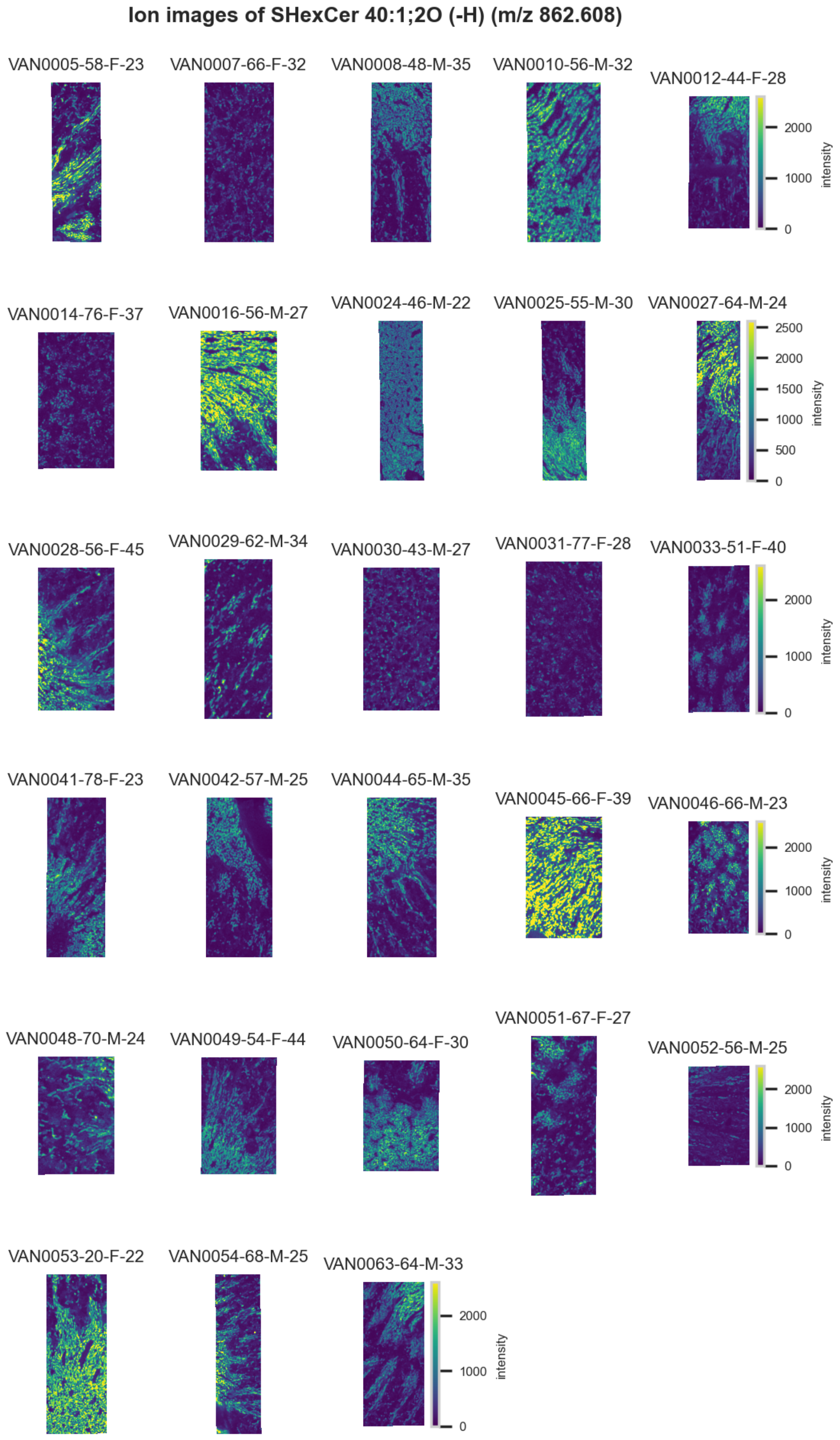

**Figure S134.** 10 μm MALDI IMS images of $[SHexCer(40:1,2O)-H]^-$ (*m/z* 862.608) detected in negative ion mode from each donor sample. Multivariate SHAP analysis determined this to be feature 6 for classification of collecting ducts.

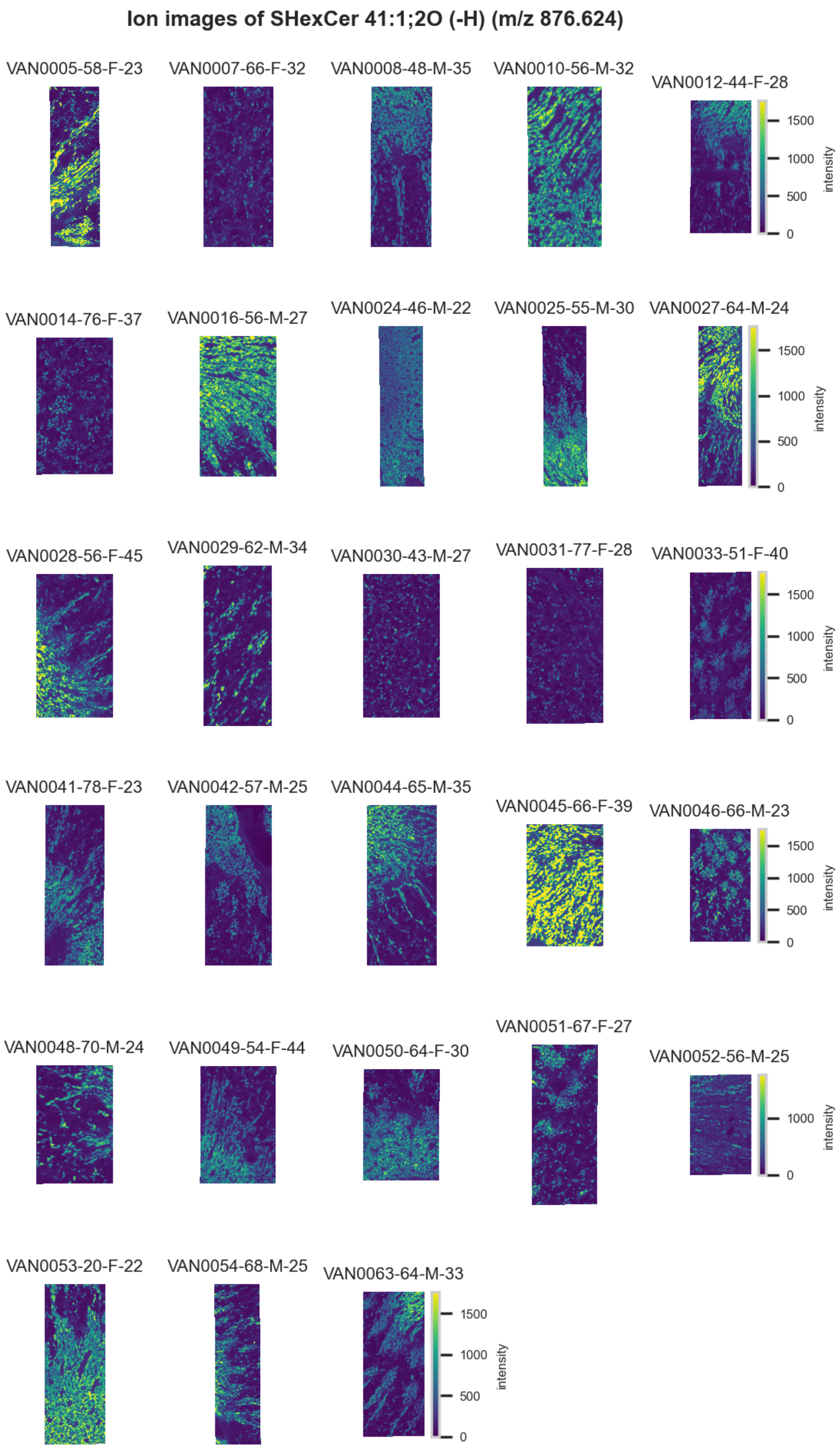

**Figure S135.** 10 µm MALDI IMS images of [SHexCer(41:1,2O)-H]$^-$ (*m/z* 876.624) detected in negative ion mode from each donor sample. Multivariate SHAP analysis determined this to be feature 7 for classification of collecting ducts.

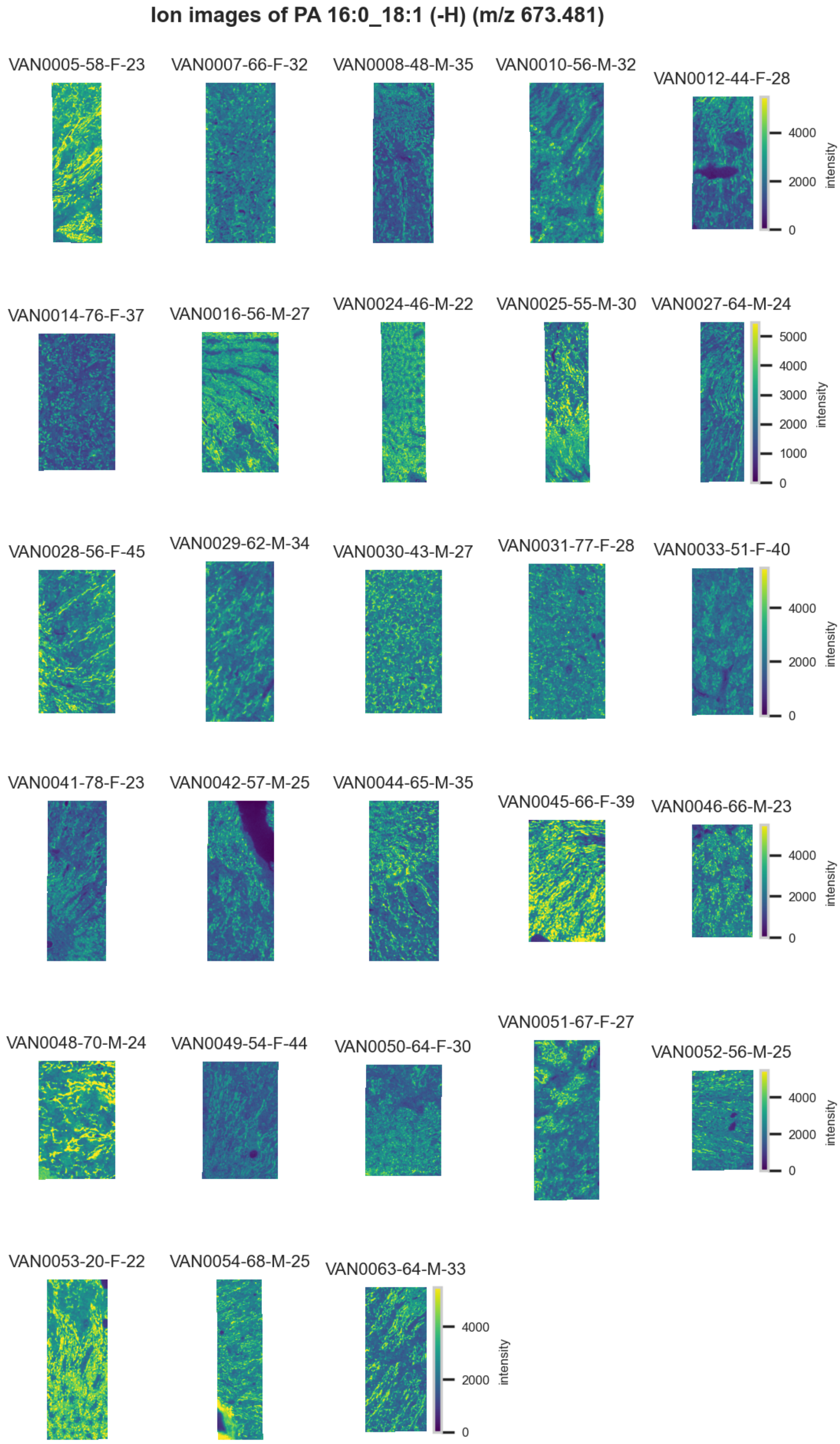

**Figure S136.** 10 μm MALDI IMS images of [PA(16:0_18:1)-H]$^-$ (*m/z* 673.481) detected in negative ion mode from each donor sample. Multivariate SHAP analysis determined this to be feature 8 for classification of collecting ducts.

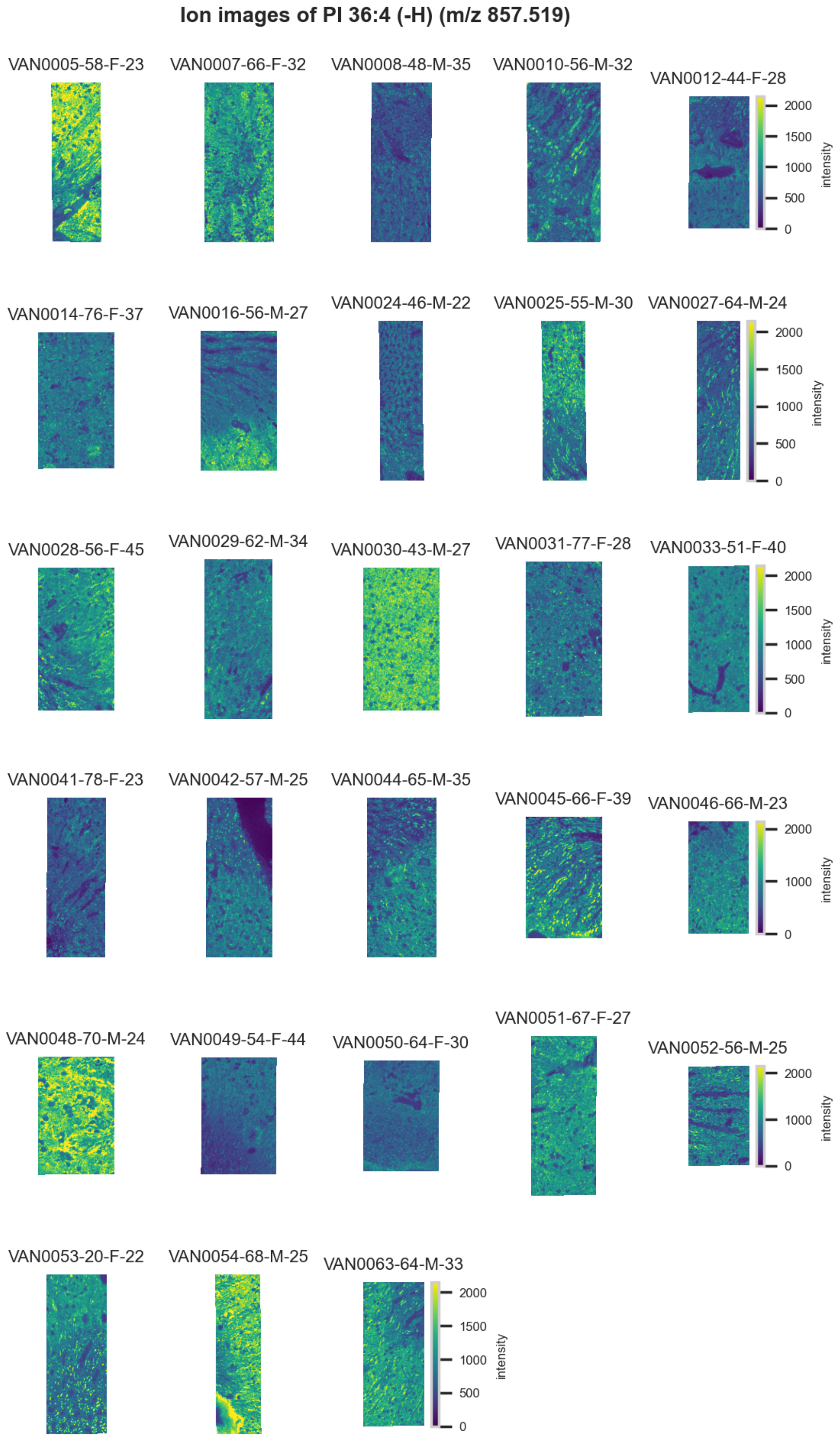

**Figure S137.** 10 µm MALDI IMS images of [PI(36:4)-H]$^-$ (*m/z* 857.519) detected in negative ion mode from each donor sample. Multivariate SHAP analysis determined this to be feature 9 for classification of collecting ducts.

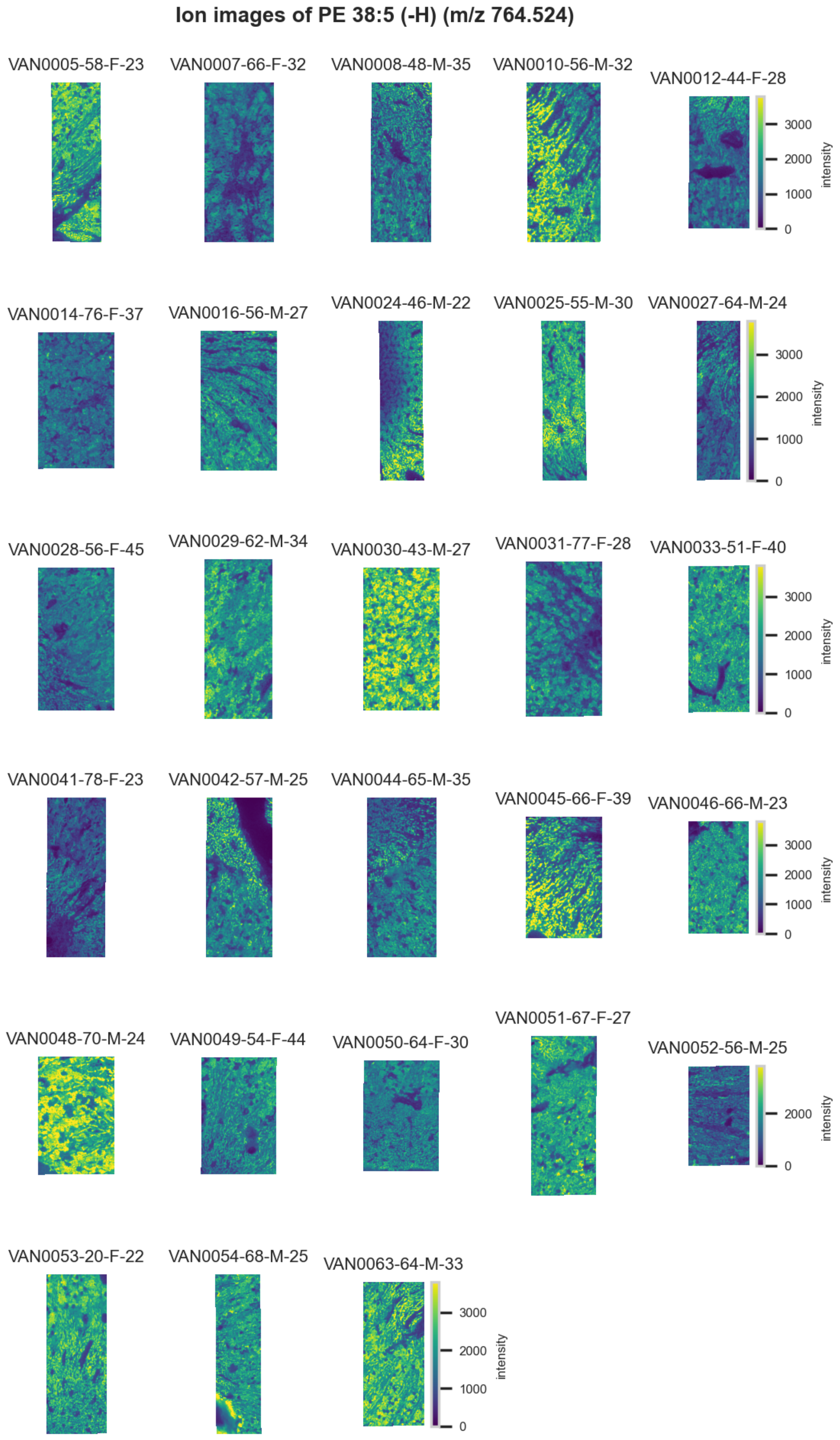

**Figure S138.** 10 µm MALDI IMS images of $[PE(38:5)-H]^-$ (*m/z* 764.524) detected in negative ion mode from each donor sample. Multivariate SHAP analysis determined this to be feature 10 for classification of collecting ducts.

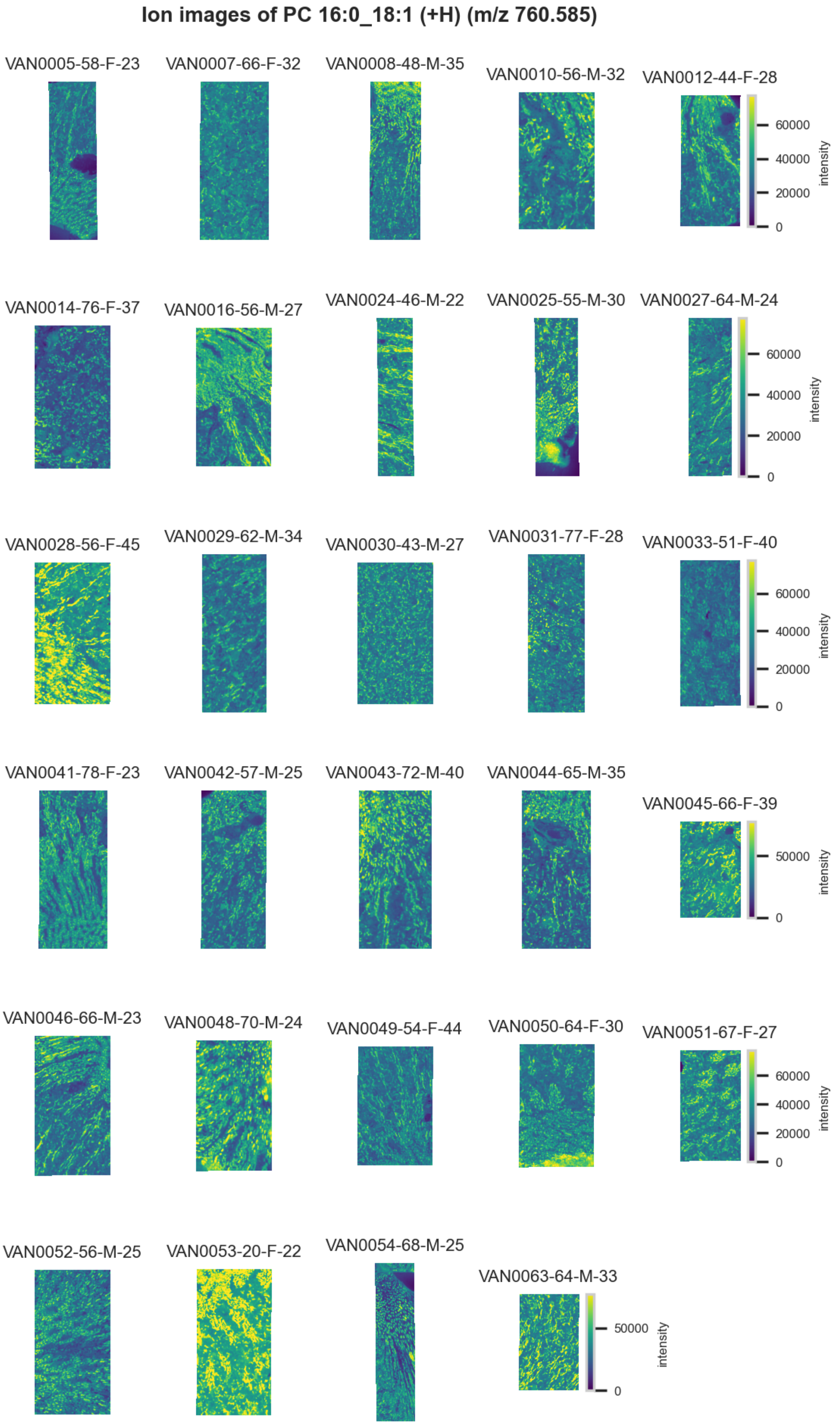

**Figure S139.** 10 μm MALDI IMS images of $[PC(16:0\_18:1)+H]^+$ (*m/z* 760.585) detected in positive ion mode from each donor sample. Multivariate SHAP analysis determined this to be feature 1 for classification of collecting ducts.

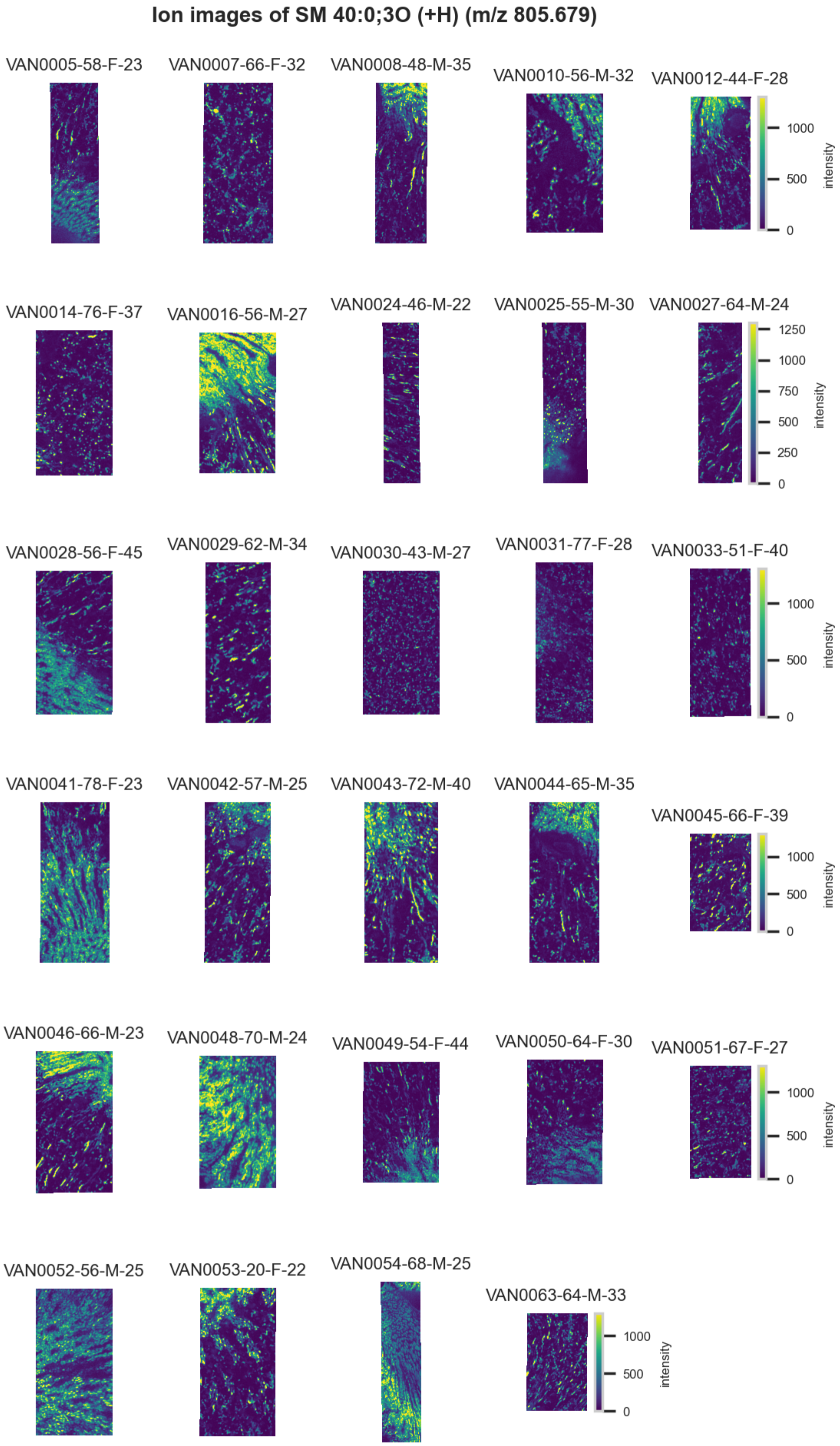

**Figure S140.** 10 µm MALDI IMS images of $[SM(40:0,3O)+H]^+$ (*m/z* 805.679) detected in positive ion mode from each donor sample. Multivariate SHAP analysis determined this to be feature 2 for classification of collecting ducts.

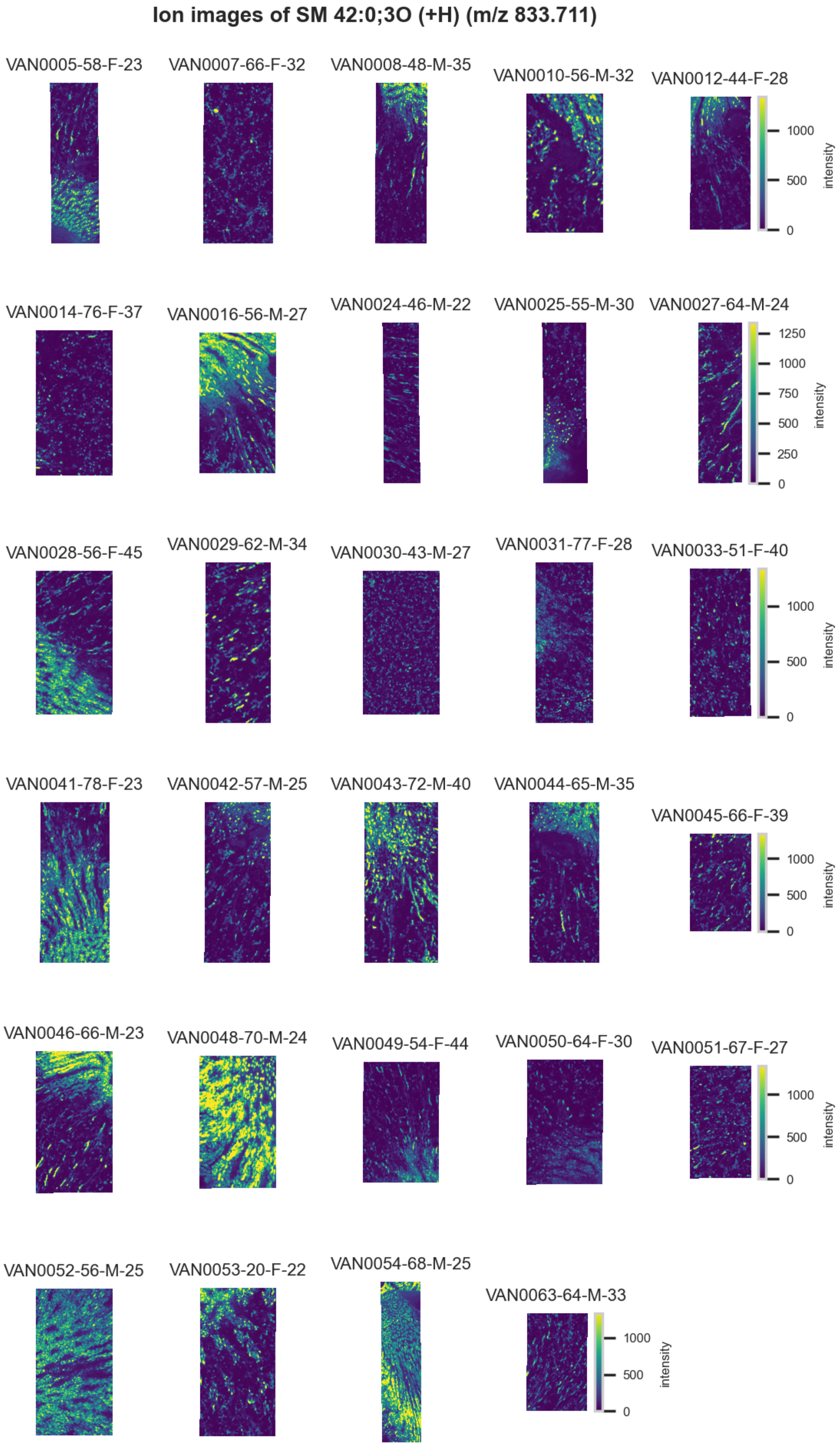

**Figure S141.** 10 μm MALDI IMS images of $[SM(42:0,3O)+H]^+$ (*m/z* 833.711) detected in positive ion mode from each donor sample. Multivariate SHAP analysis determined this to be feature 3 for classification of collecting ducts.

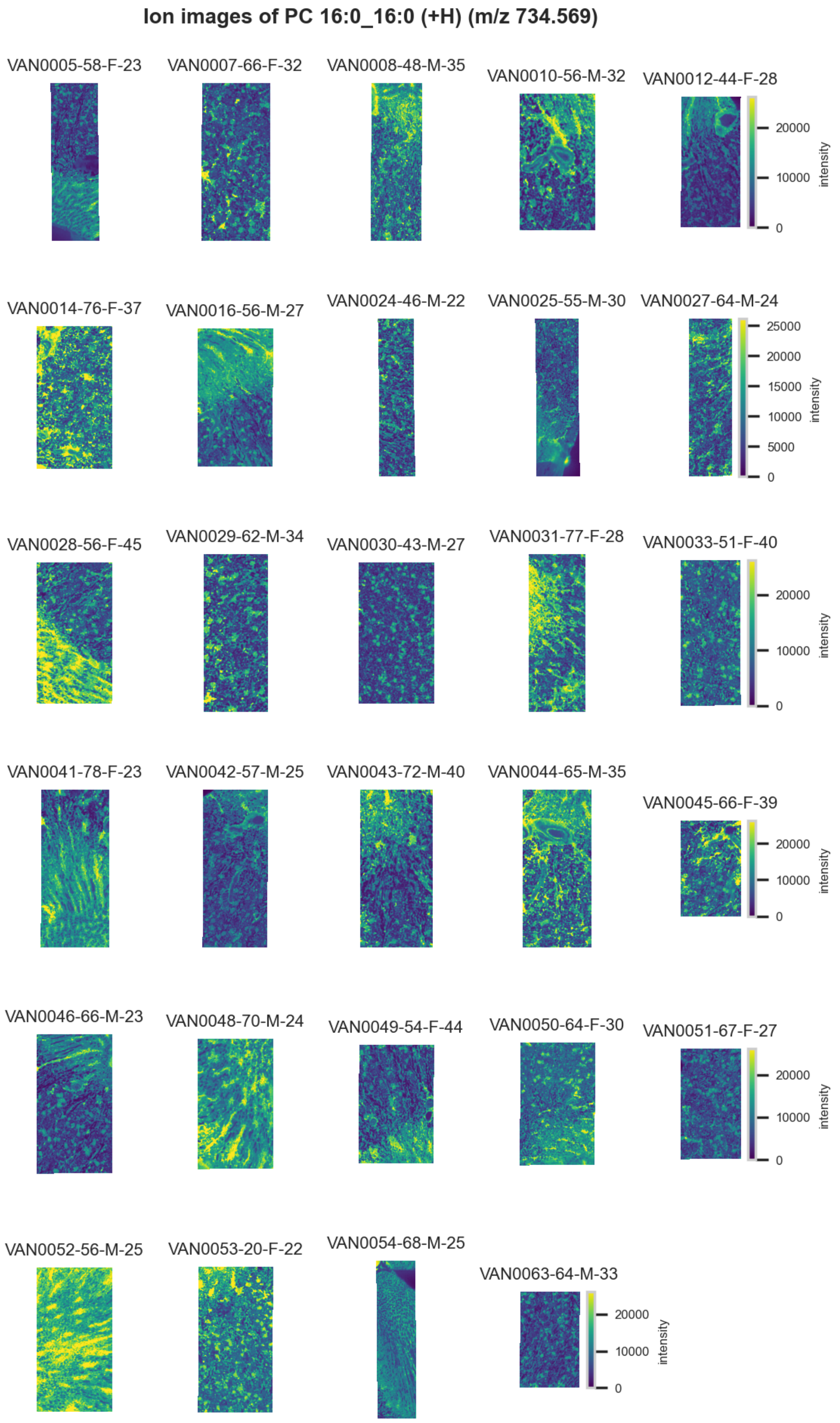

**Figure S142.** 10 μm MALDI IMS images of an unknown $[PC(16:0/16:0)+H]^+$ (*m/z* 734.569) detected in positive ion mode from each donor sample. Multivariate SHAP analysis determined this to be feature 4 for classification of collecting ducts.

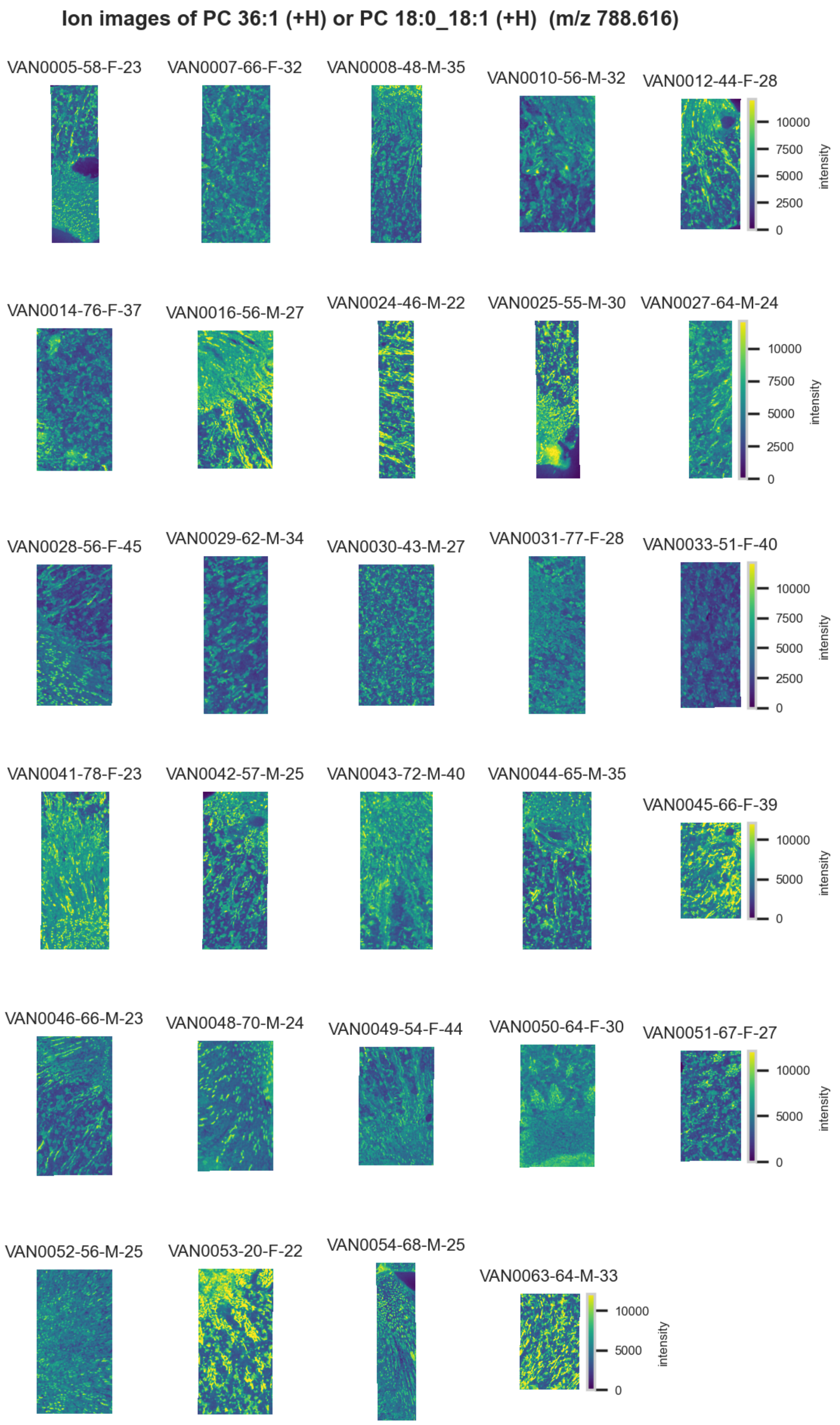

**Figure S143.** 10 µm MALDI IMS images of $[PC(36:1)+H]^+$ or $[PC(18:0\_18:1)+H]^+$ (*m/z* 788.616) detected in positive ion mode from each donor sample. These molecules are isomers and therefore indistinguishable by MS alone. Multivariate SHAP analysis determined this to be feature 5 for classification of collecting ducts.

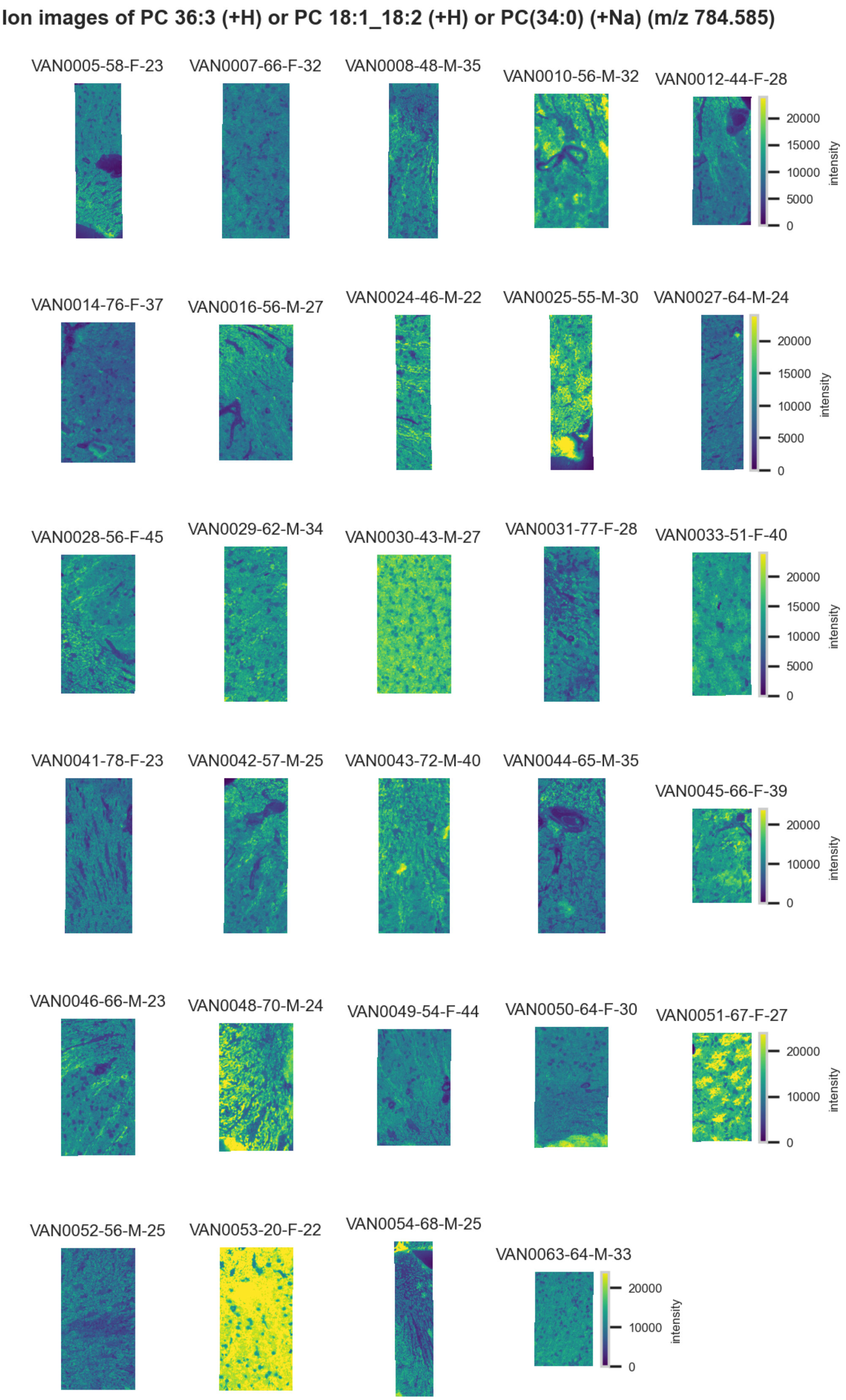

**Figure S144.** 10 µm MALDI IMS images of $[PC(36:3)+H]^+$ or $[PC(18:1\_18:3)+H]^+$ or $[PC(34:0)+Na]^+$ (*m/z* 784.585) detected in positive ion mode from each donor sample. These molecules are isomers and therefore indistinguishable by MS alone. Multivariate SHAP analysis determined this to be feature 6 for classification of collecting ducts.

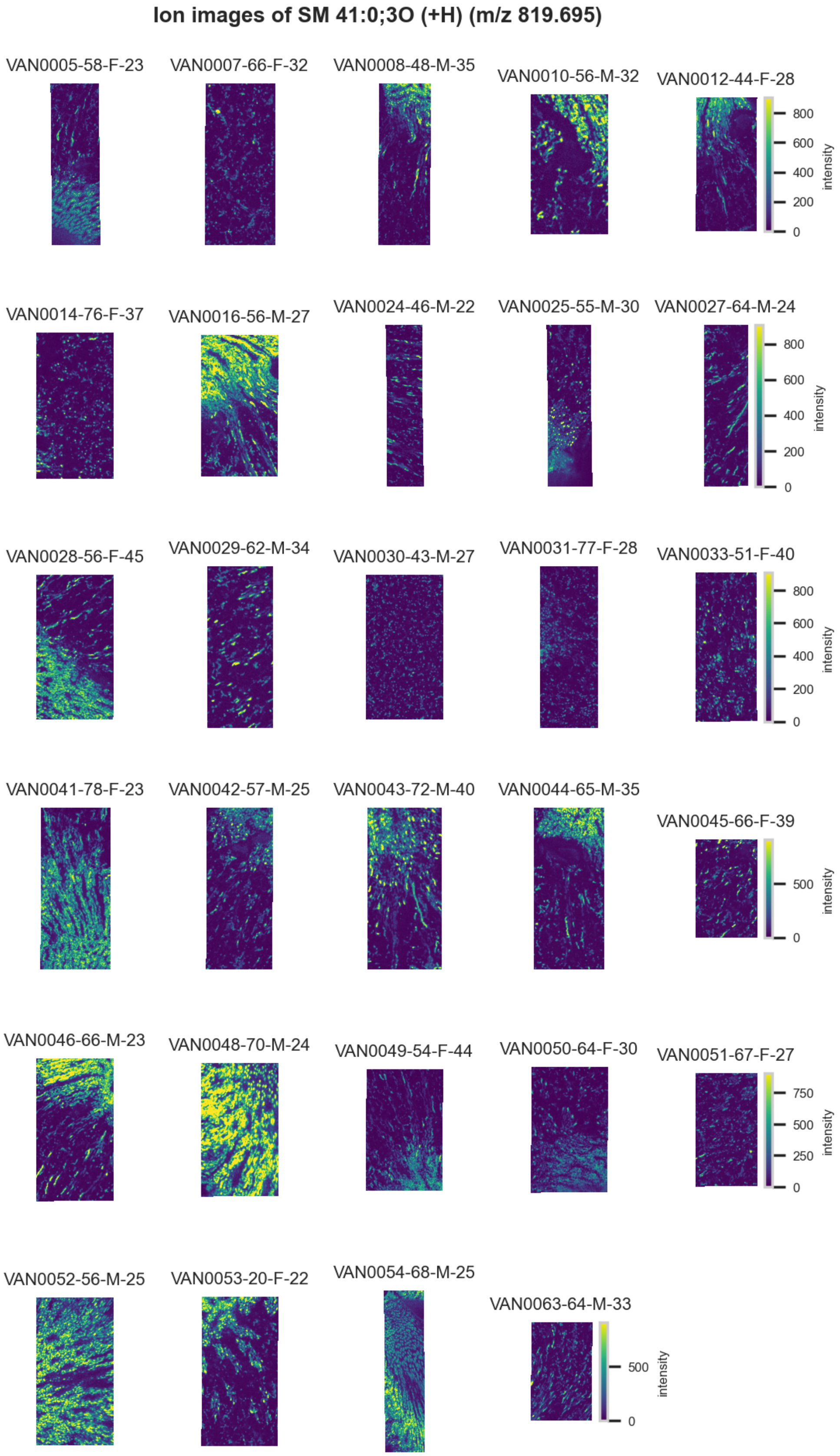

**Figure S145.** 10 µm MALDI IMS images of $[SM(41:0,3O)+H]^+$ (*m/z* 819.695) detected in positive ion mode from each donor sample. Multivariate SHAP analysis determined this to be feature 7 for classification of collecting ducts.

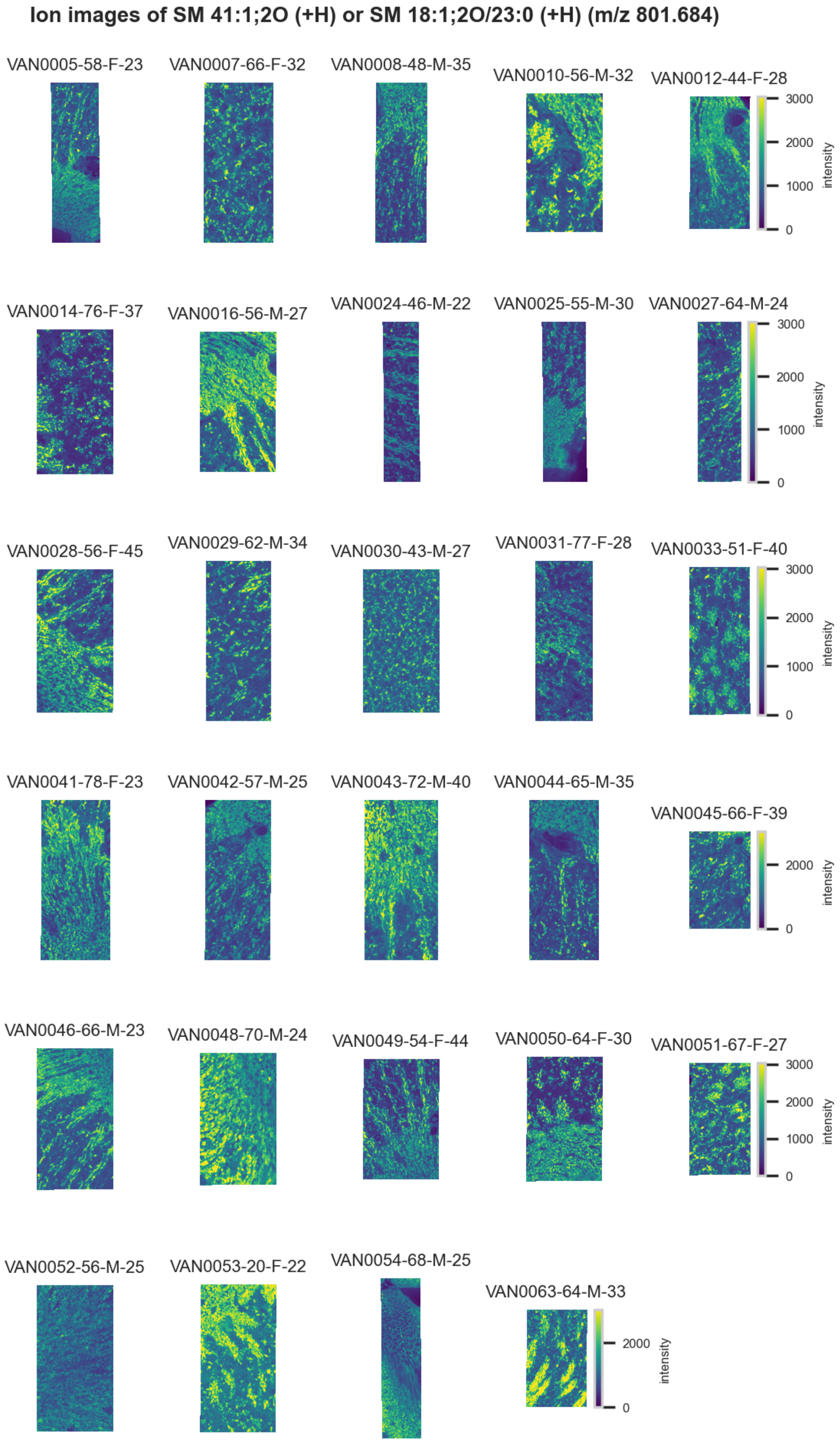

**Figure S146.** 10 µm MALDI IMS images of $[SM(41:1,2O)+H]^+$ or $[SM(18:1,2O/23:0)+H]^+$ (*m/z* 801.684) detected in <u>positive ion mode</u> from each donor sample. These molecules are isomers and therefore indistinguishable by MS alone. Multivariate SHAP analysis determined this to be feature 8 for classification of collecting ducts.

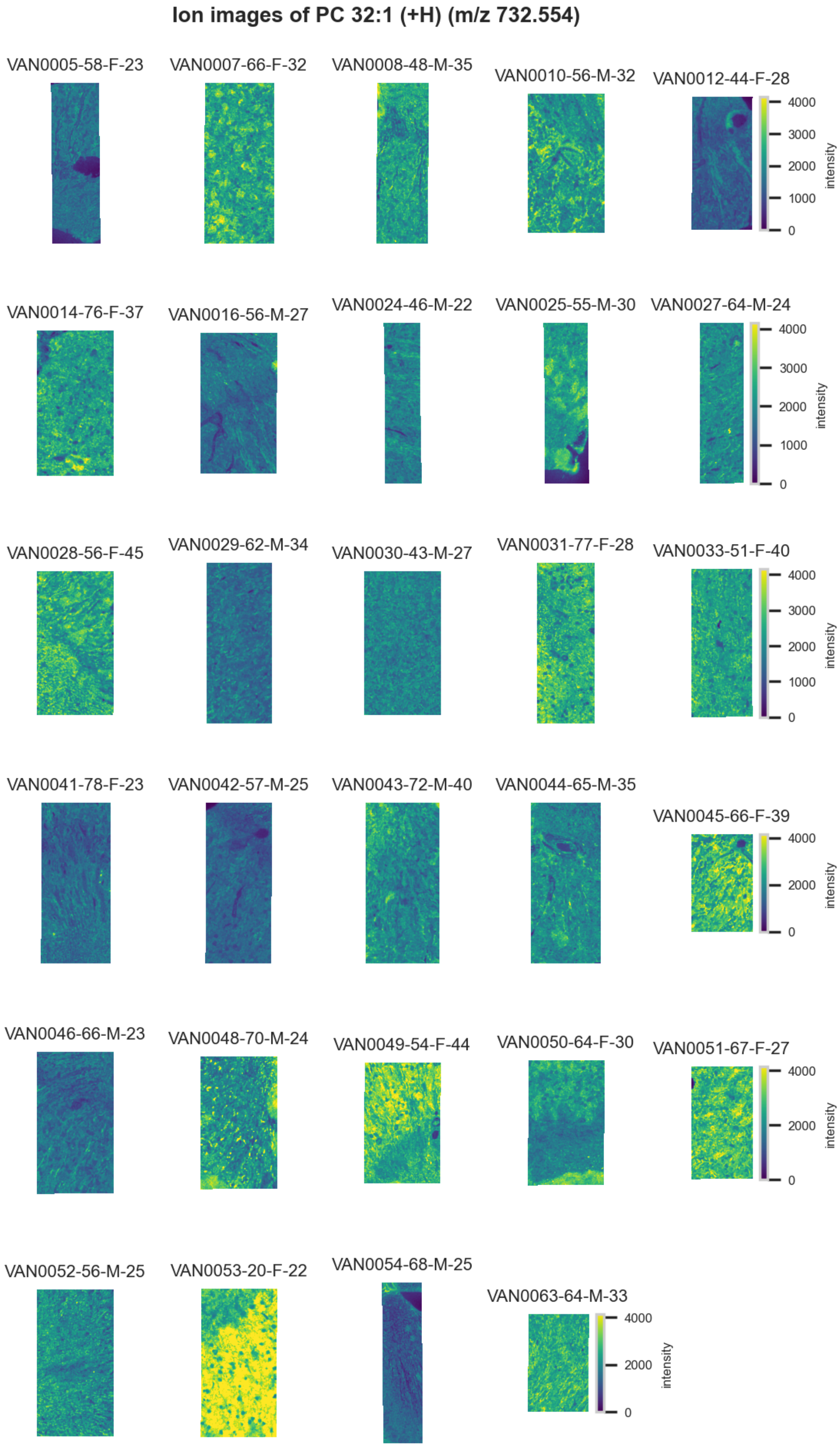

**Figure S147.** 10 µm MALDI IMS images of $[PC(32:1)+H]^+$ (*m/z* 732.554) detected in positive ion mode from each donor sample. Multivariate SHAP analysis determined this to be feature 9 for classification of collecting ducts.

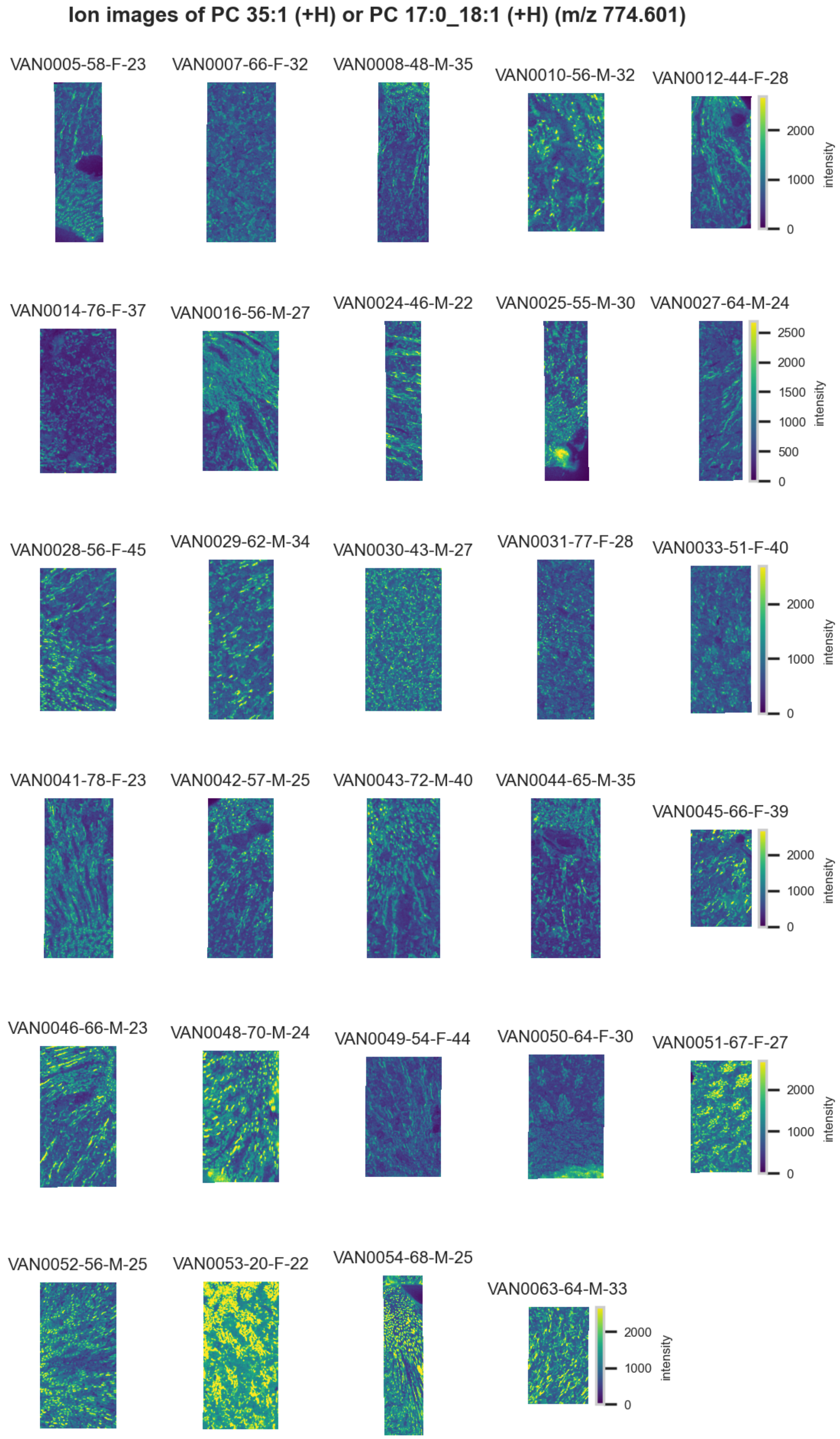

**Figure S148.** 10 µm MALDI IMS images of $[PC(35:1)+H]^+$ or $[PC(17:0\_18:1)+H]^+$ (*m/z* 774.601) detected in positive ion mode from each donor sample. These molecules are isomers and therefore indistinguishable by MS alone. Multivariate SHAP analysis determined this to be feature 10 for classification of collecting ducts.

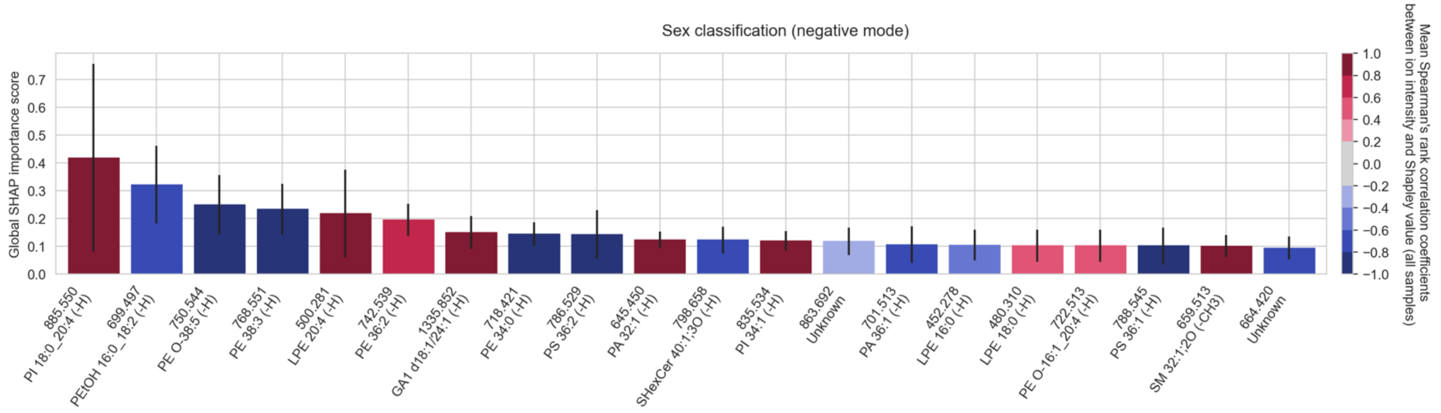

**Figure S149.** The top twenty cohort-wide (absolute) SHAP importance scores reporting the top twenty molecular species among the measured lipid species that enable the classification of **female donor tissue from male donor tissue overall (not specific to an FTU), in negative ion mode**. The error bar is the standard deviation across all donor samples of the tissue sample-wide SHAP importance score for a given molecular species. The color of the bar reports the mean Spearman's rank correlation coefficient (across all donor samples) between the molecular species' mean-centered ion intensity and its Shapley score. Given that the sex classification task is binary (male versus female), we made the female class the target class of our classification model. Therefore, we report positive and negative correlation to female sex. A positive Spearman's rank correlation coefficient indicates that high intensity of the molecular species correlates with female sex. Molecular species that are positively correlated with female sex are negatively correlated with male sex, and vice-versa (refer to Figure 7).

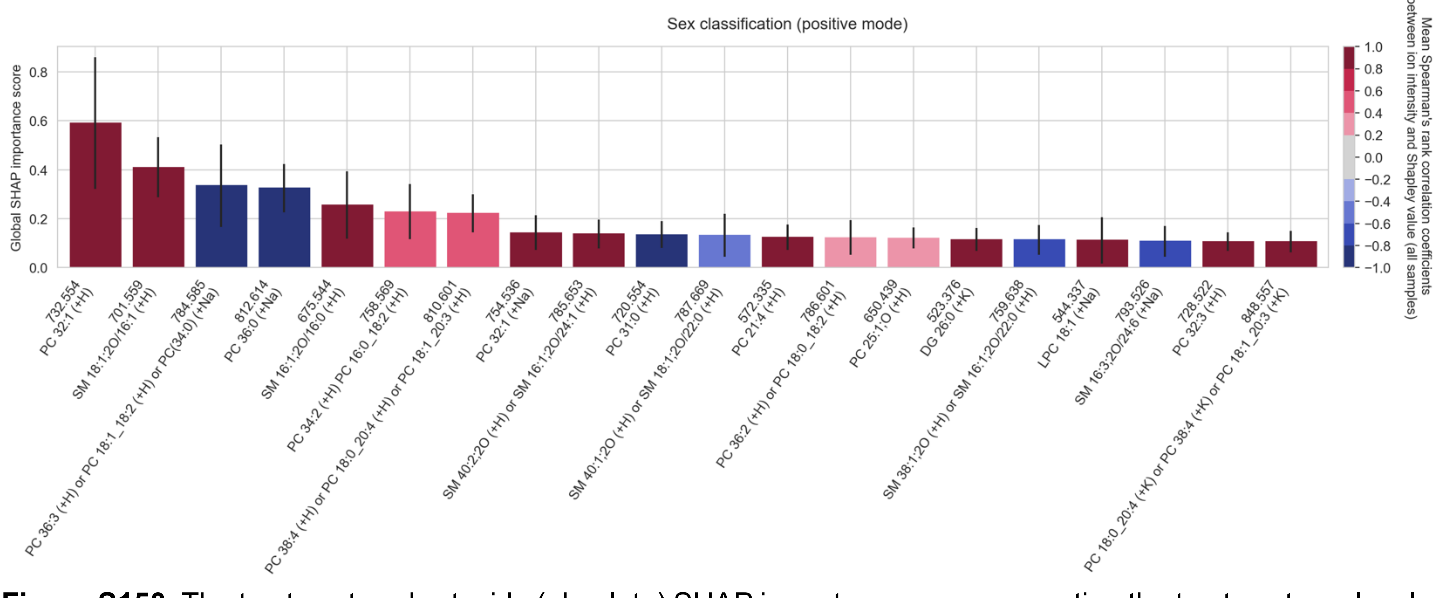

**Figure S150.** The top twenty cohort-wide (absolute) SHAP importance scores reporting the top twenty molecular species among the measured lipid species that enable the classification of **female donor tissue from male donor tissue overall (not specific to an FTU), in positive ion mode**. The error bar is the standard deviation across all donor samples of the tissue sample-wide SHAP importance score for a given molecular species. The color of the bar reports the mean Spearman's rank correlation coefficient (across all donor samples) between the molecular species' mean-centered ion intensity and its Shapley score. Given that the sex classification task is binary (male versus female), we made the female class the target class of our classification model. Therefore, we report positive and negative correlation to female sex. A positive Spearman's rank correlation coefficient indicates that high intensity of the molecular species correlates with female sex. Molecular species that are positively correlated with female sex are negatively correlated with male sex, and vice-versa (refer to Figure 7).

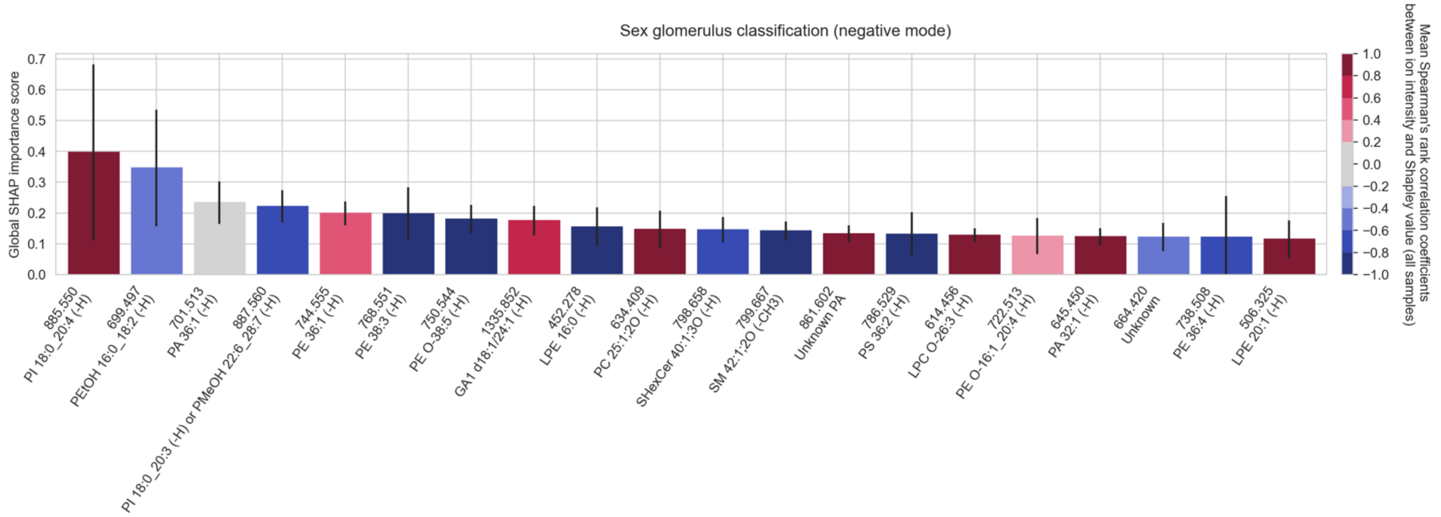

**Figure S151.** The top twenty cohort-wide (absolute) SHAP importance scores reporting the top twenty molecular species among the measured lipid species that enable the classification of **<u>the glomeruli of female donors from the glomeruli of male donors, in negative ion mode</u>**. The error bar is the standard deviation across all donor samples of the tissue sample-wide SHAP importance score for a given molecular species. The color of the bar reports the mean Spearman's rank correlation coefficient (across all donor samples) between the molecular species' mean-centered ion intensity and its Shapley score. A positive Spearman's rank correlation coefficient indicates that high intensity of the molecular species correlates with female sex in glomeruli. Conversely, a negative Spearman's rank correlation coefficient indicates that low intensity of the molecular species correlates with female sex in glomeruli.

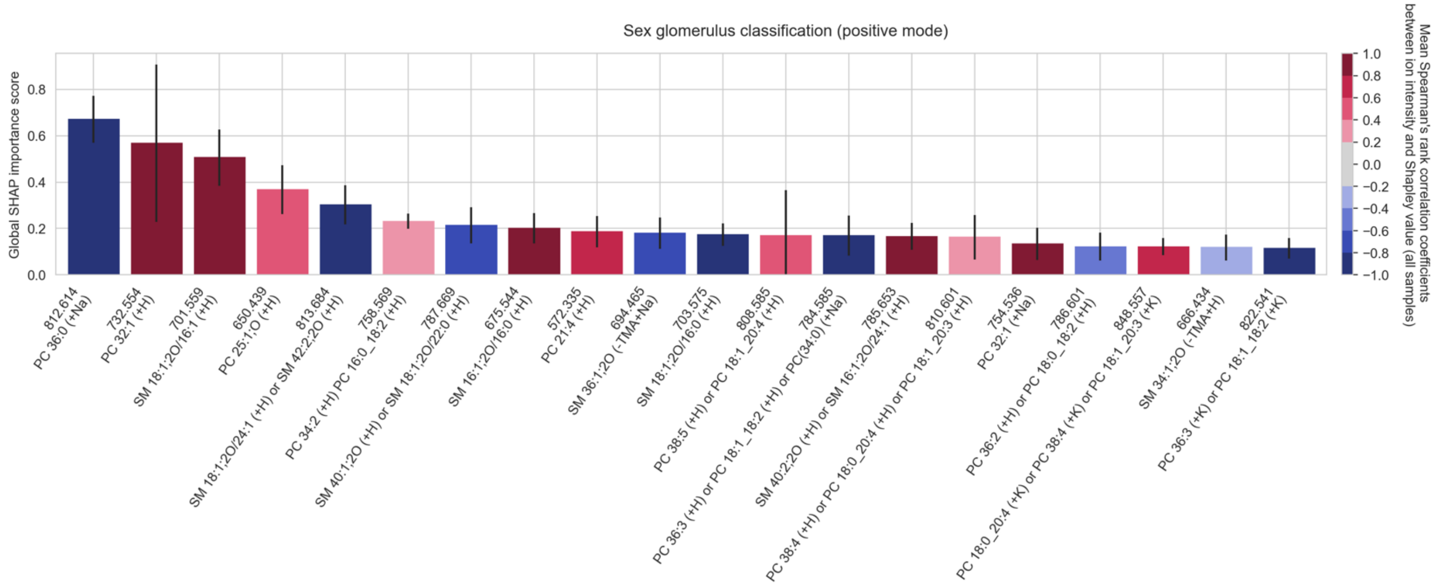

**Figure S152.** The top twenty cohort-wide (absolute) SHAP importance scores reporting the top twenty molecular species among the measured lipid species that enable the classification of **<u>the glomeruli of female donors from the glomeruli of male donors, in positive ion mode</u>**. The error bar is the standard deviation across all donor samples of the tissue sample-wide SHAP importance score for a given molecular species. The color of the bar reports the mean Spearman's rank correlation coefficient (across all donor samples) between the molecular species' mean-centered ion intensity and its Shapley score. A positive Spearman's rank correlation coefficient indicates that high intensity of the molecular species correlates with female sex in glomeruli. Conversely, a negative Spearman's rank correlation coefficient indicates that low intensity of the molecular species correlates with female sex in glomeruli.

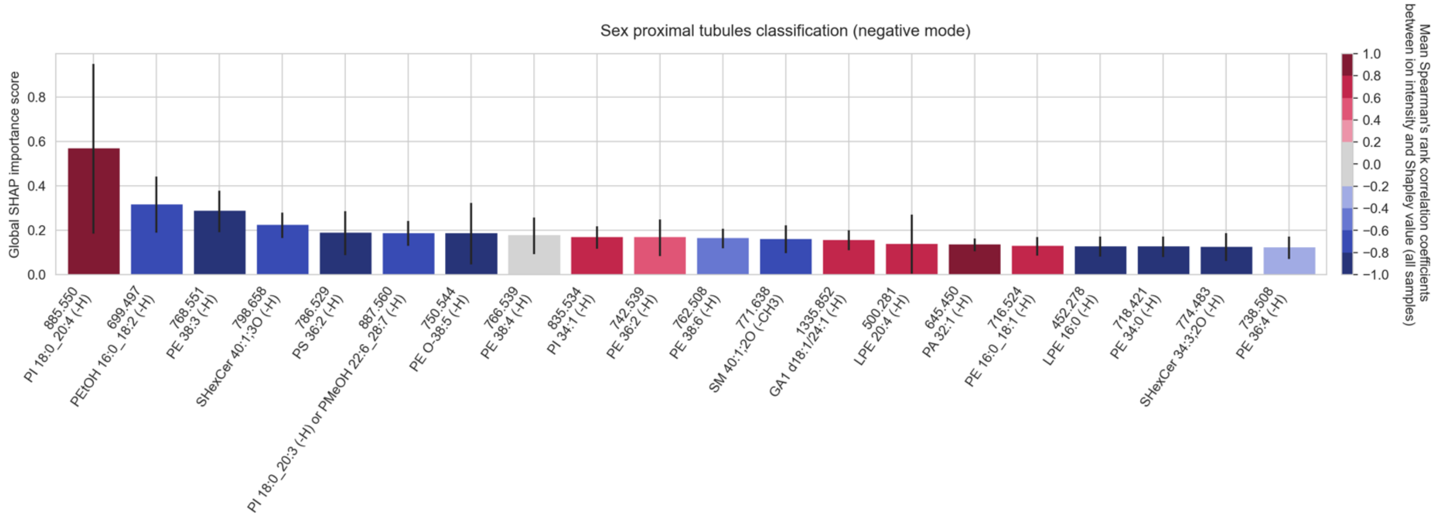

**Figure S153.** The top twenty cohort-wide (absolute) SHAP importance scores reporting the top twenty molecular species among the measured lipid species that enable the classification of **the proximal tubules of female donors from the proximal tubules of male donors, in negative ion mode**. The error bar is the standard deviation across all donor samples of the tissue sample-wide SHAP importance score for a given molecular species. The color of the bar reports the mean Spearman's rank correlation coefficient (across all donor samples) between the molecular species' mean-centered ion intensity and its Shapley score. A positive Spearman's rank correlation coefficient indicates that high intensity of the molecular species correlates with female sex in proximal tubules. Conversely, a negative Spearman's rank correlation coefficient indicates that low intensity of the molecular species correlates with female sex in proximal tubules.

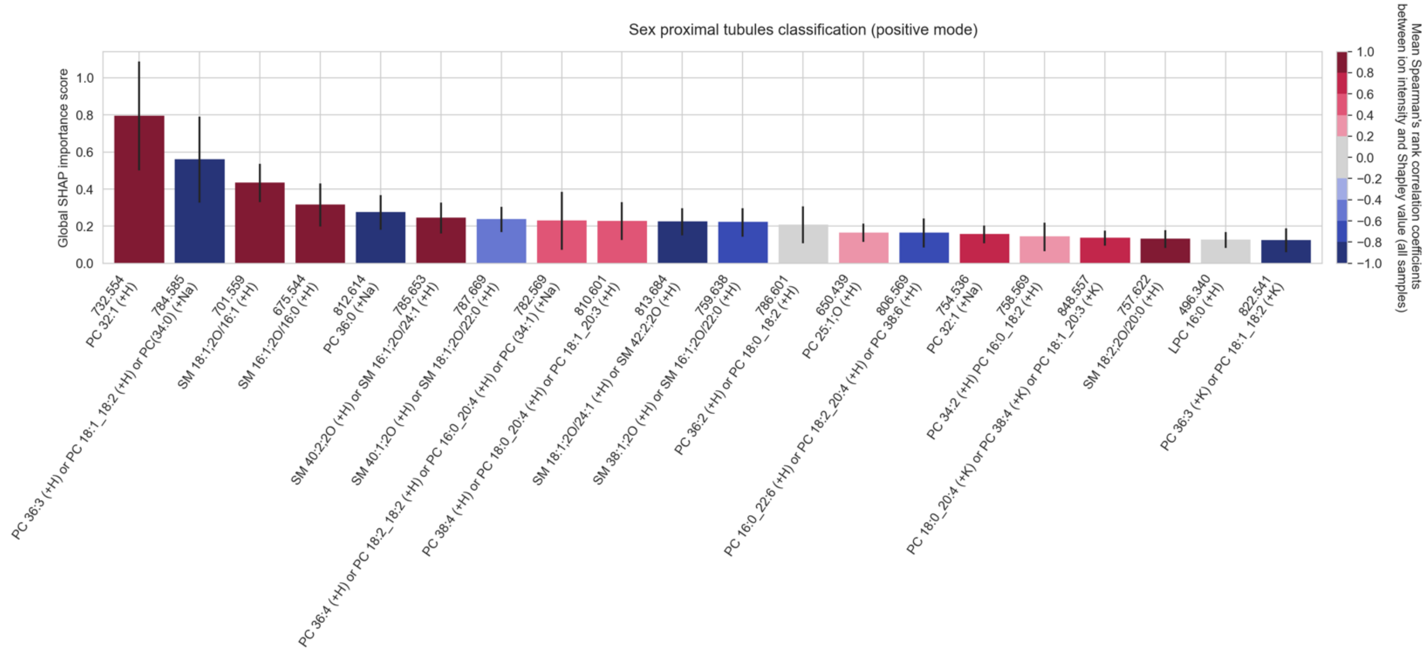

**Figure S154.** The top twenty cohort-wide (absolute) SHAP importance scores reporting the top twenty molecular species among the measured lipid species that enable the classification of **the proximal tubules of female donors from the proximal tubules of male donors, in positive ion mode**. The error bar is the standard deviation across all donor samples of the tissue sample-wide SHAP importance score for a given molecular species. The color of the bar reports the mean Spearman's rank correlation coefficient (across all donor samples) between the molecular species' mean-centered ion intensity and its Shapley score. A positive Spearman's rank correlation coefficient indicates that high intensity of the molecular species correlates with female sex in proximal tubules. Conversely, a negative Spearman's rank correlation coefficient indicates that low intensity of the molecular species correlates with female sex in proximal tubules.

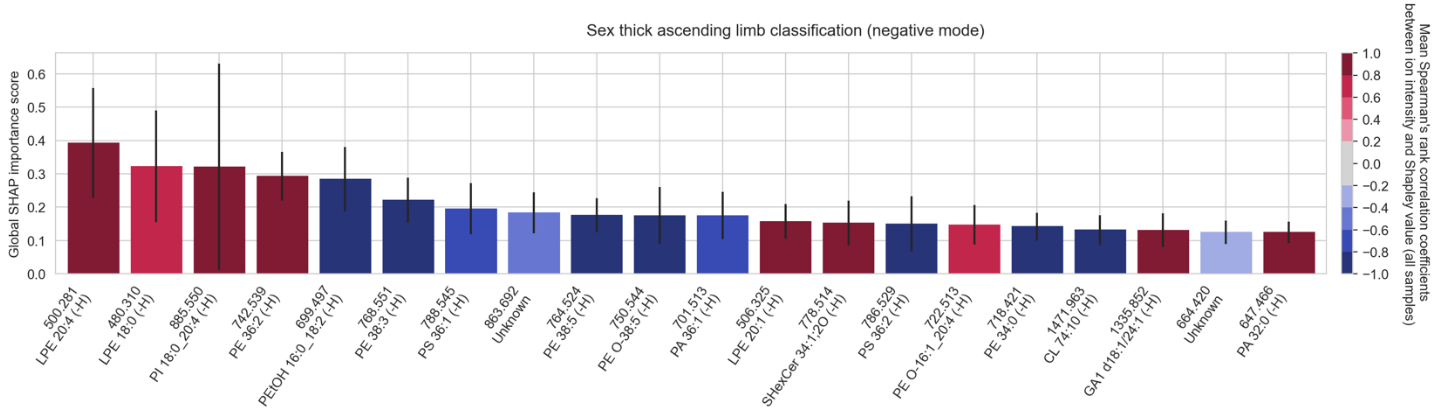

**Figure S155.** The top twenty cohort-wide (absolute) SHAP importance scores reporting the top twenty molecular species among the measured lipid species that enable the classification of **the thick ascending limb of female donors from the thick ascending limb of male donors, in negative ion mode**. The error bar is the standard deviation across all donor samples of the tissue sample-wide SHAP importance score for a given molecular species. The color of the bar reports the mean Spearman's rank correlation coefficient (across all donor samples) between the molecular species' mean-centered ion intensity and its Shapley score. A positive Spearman's rank correlation coefficient indicates that high intensity of the molecular species correlates with female sex in thick ascending limb. Conversely, a negative Spearman's rank correlation coefficient indicates that low intensity of the molecular species correlates with female sex in thick ascending limb.

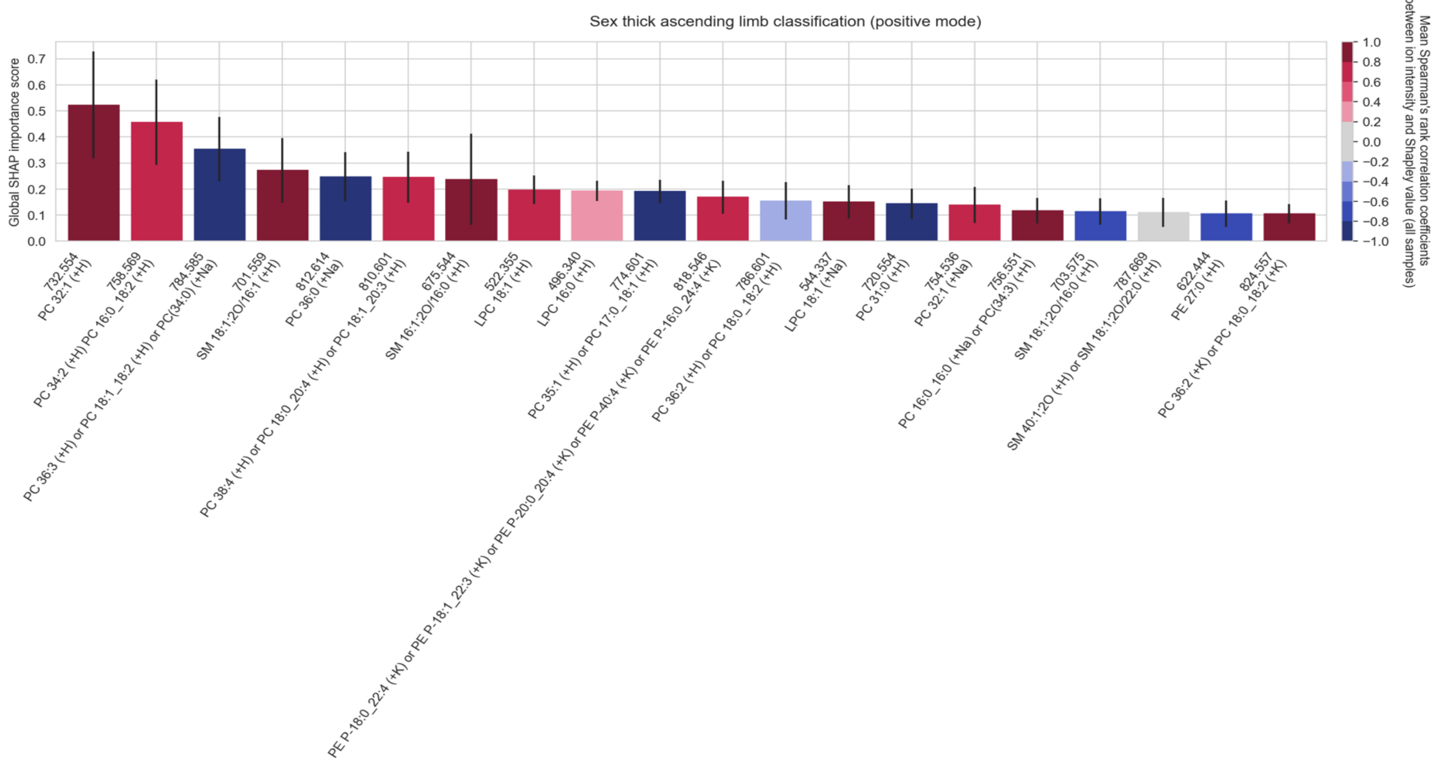

**Figure S156.** The top twenty cohort-wide (absolute) SHAP importance scores reporting the top twenty molecular species among the measured lipid species that enable the classification of **the thick ascending limb of female donors from the thick ascending limb of male donors, in positive ion mode**. The error bar is the standard deviation across all donor samples of the tissue sample-wide SHAP importance score for a given molecular species. The color of the bar reports the mean Spearman's rank correlation coefficient (across all donor samples) between the molecular species' mean-centered ion intensity and its Shapley score. A positive Spearman's rank correlation coefficient indicates that high intensity of the molecular species correlates with female sex in thick ascending limb. Conversely, a negative Spearman's rank correlation coefficient indicates that low intensity of the molecular species correlates with female sex in thick ascending limb.

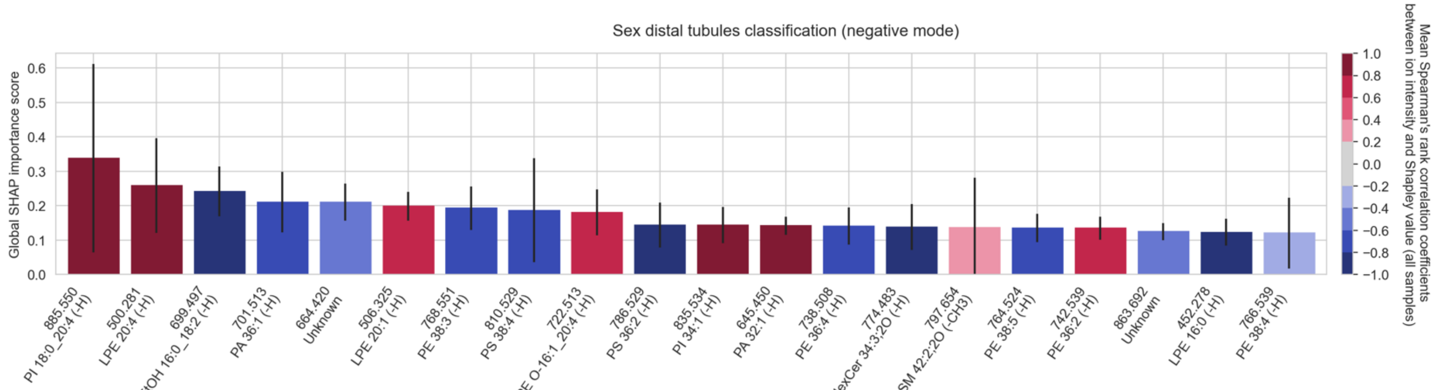

**Figure S157.** The top twenty cohort-wide (absolute) SHAP importance scores reporting the top twenty molecular species among the measured lipid species that enable the classification of **the distal tubules of female donors from the distal tubules of male donors, in negative ion mode**. The error bar is the standard deviation across all donor samples of the tissue sample-wide SHAP importance score for a given molecular species. The color of the bar reports the mean Spearman's rank correlation coefficient (across all donor samples) between the molecular species' mean-centered ion intensity and its Shapley score. A positive Spearman's rank correlation coefficient indicates that high intensity of the molecular species correlates with female sex in distal tubules. Conversely, a negative Spearman's rank correlation coefficient indicates that low intensity of the molecular species correlates with female sex in distal tubules.

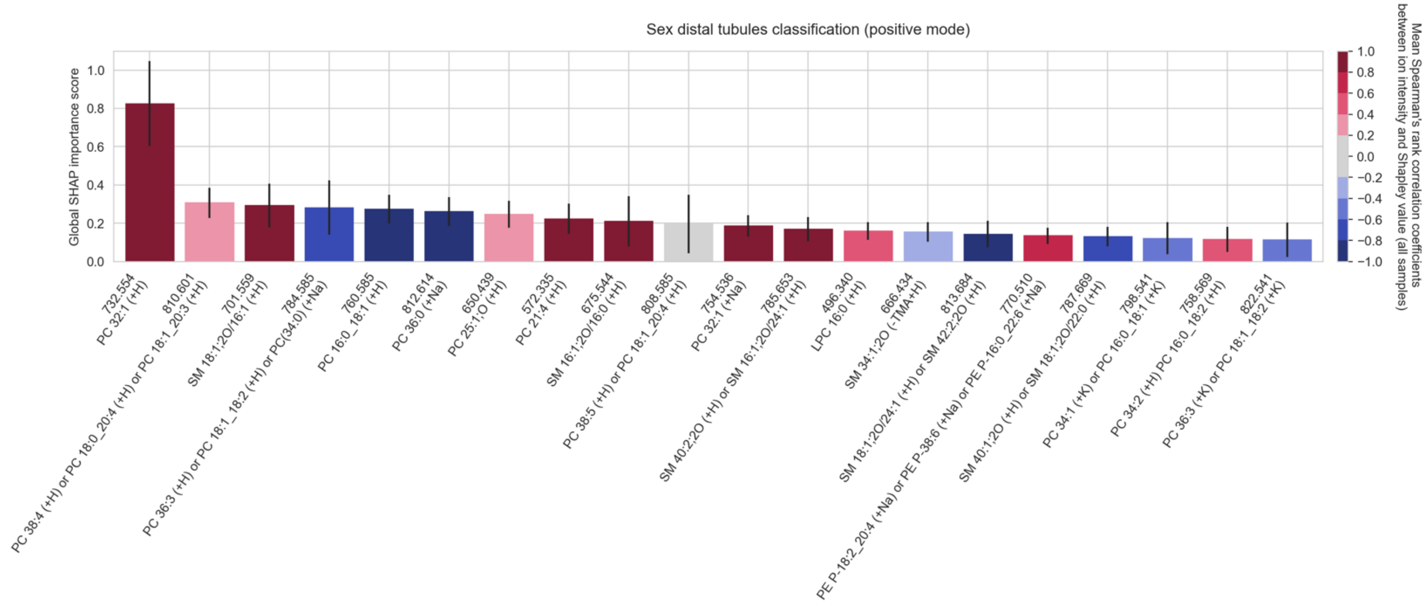

**Figure S158.** The top twenty cohort-wide (absolute) SHAP importance scores reporting the top twenty molecular species among the measured lipid species that enable the classification of **the distal tubules of female donors from the distal tubules of male donors, in positive ion mode**. The error bar is the standard deviation across all donor samples of the tissue sample-wide SHAP importance score for a given molecular species. The color of the bar reports the mean Spearman's rank correlation coefficient (across all donor samples) between the molecular species' mean-centered ion intensity and its Shapley score. A positive Spearman's rank correlation coefficient indicates that high intensity of the molecular species correlates with female sex in distal tubules. Conversely, a negative Spearman's rank correlation coefficient indicates that low intensity of the molecular species correlates with female sex in distal tubules.

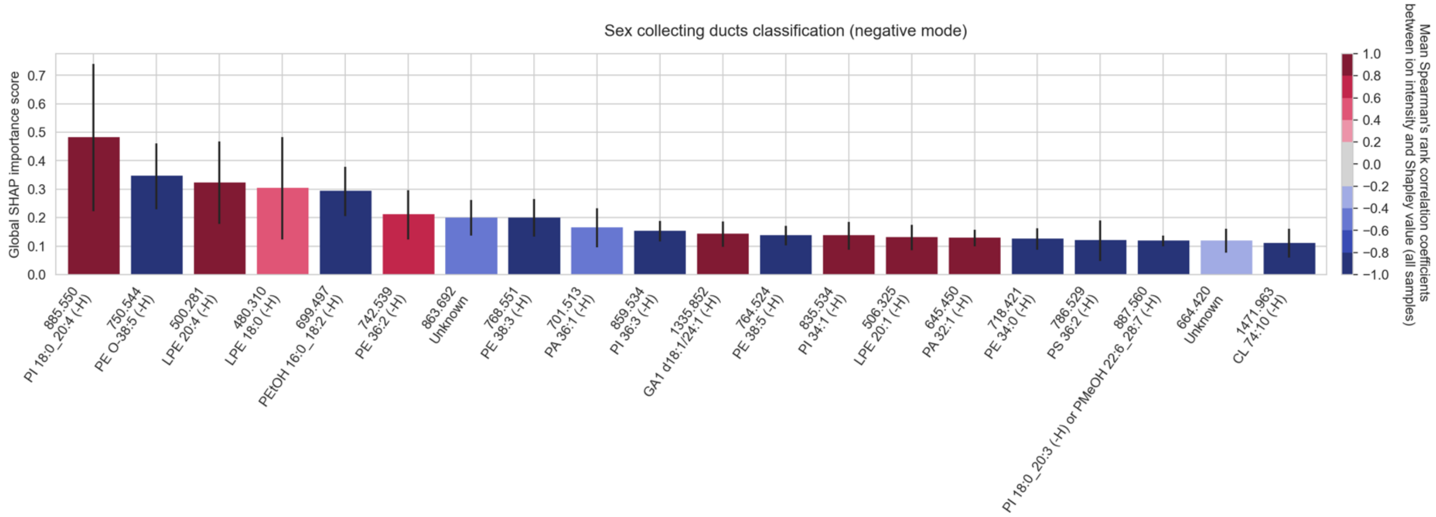

**Figure S159.** The top twenty cohort-wide (absolute) SHAP importance scores reporting the top twenty molecular species among the measured lipid species that enable the classification of **the collecting ducts of female donors from the collecting ducts of male donors, in negative ion mode**. The error bar is the standard deviation across all donor samples of the tissue sample-wide SHAP importance score for a given molecular species. The color of the bar reports the mean Spearman's rank correlation coefficient (across all donor samples) between the molecular species' mean-centered ion intensity and its Shapley score. A positive Spearman's rank correlation coefficient indicates that high intensity of the molecular species correlates with female sex in collecting. Conversely, a negative Spearman's rank correlation coefficient indicates that low intensity of the molecular species correlates with female sex in collecting ducts.

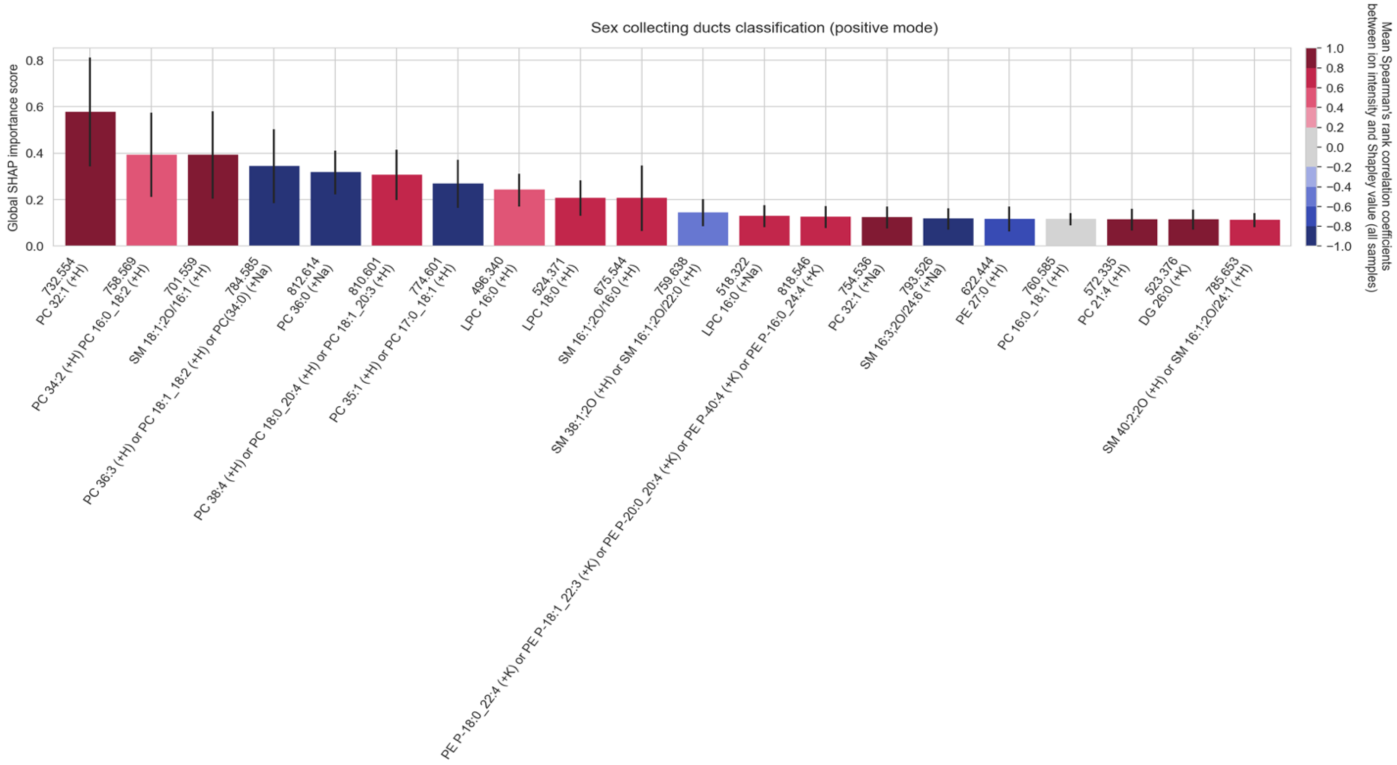

**Figure S160.** The top twenty cohort-wide (absolute) SHAP importance scores reporting the top twenty molecular species among the measured lipid species that enable the classification of **the collecting ducts of female donors from the collecting ducts of male donors, in positive ion mode**. The error bar is the standard deviation across all donor samples of the tissue sample-wide SHAP importance score for a given molecular species. The color of the bar reports the mean Spearman's rank correlation coefficient (across all donor samples) between the molecular species' mean-centered ion intensity and its Shapley score. A positive Spearman's rank correlation coefficient indicates that high intensity of the molecular species correlates with female sex in collecting. Conversely, a negative Spearman's rank correlation coefficient indicates that low intensity of the molecular species correlates with female sex in collecting ducts.

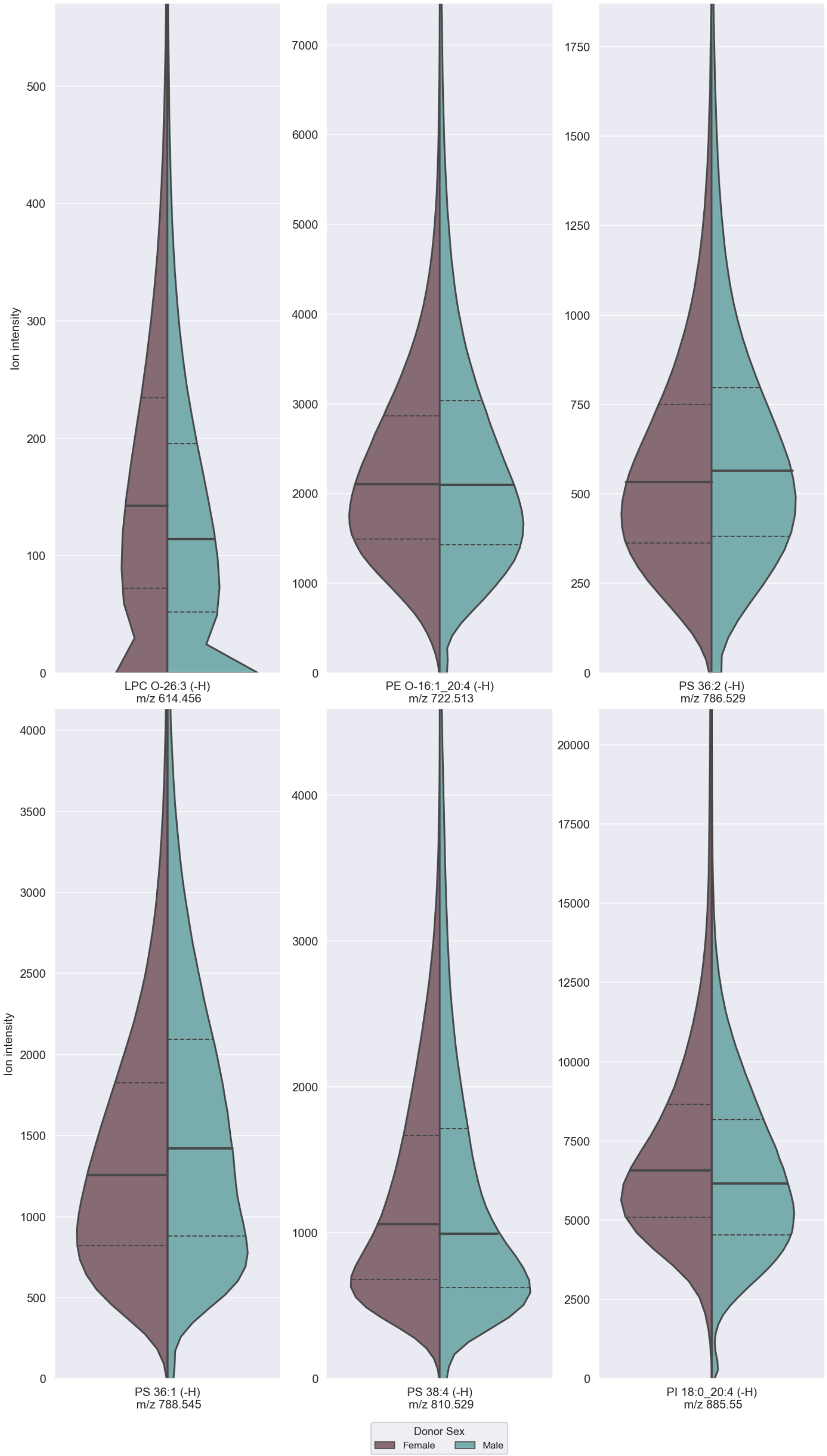

**Figure S161.** Split violin plots of the ion intensity distributions of **a selection of sex biomarker candidates in female (left) and male (right) donor data, in negative ion mode**, approximated using kernel density estimation. The violin plots are cropped at the 99th percentile of the distribution of one of the two classes (whichever is larger) to facilitate visual comparison. The full line indicates the median of each class' distribution, whereas the dashed lines indicate its interquartile range.

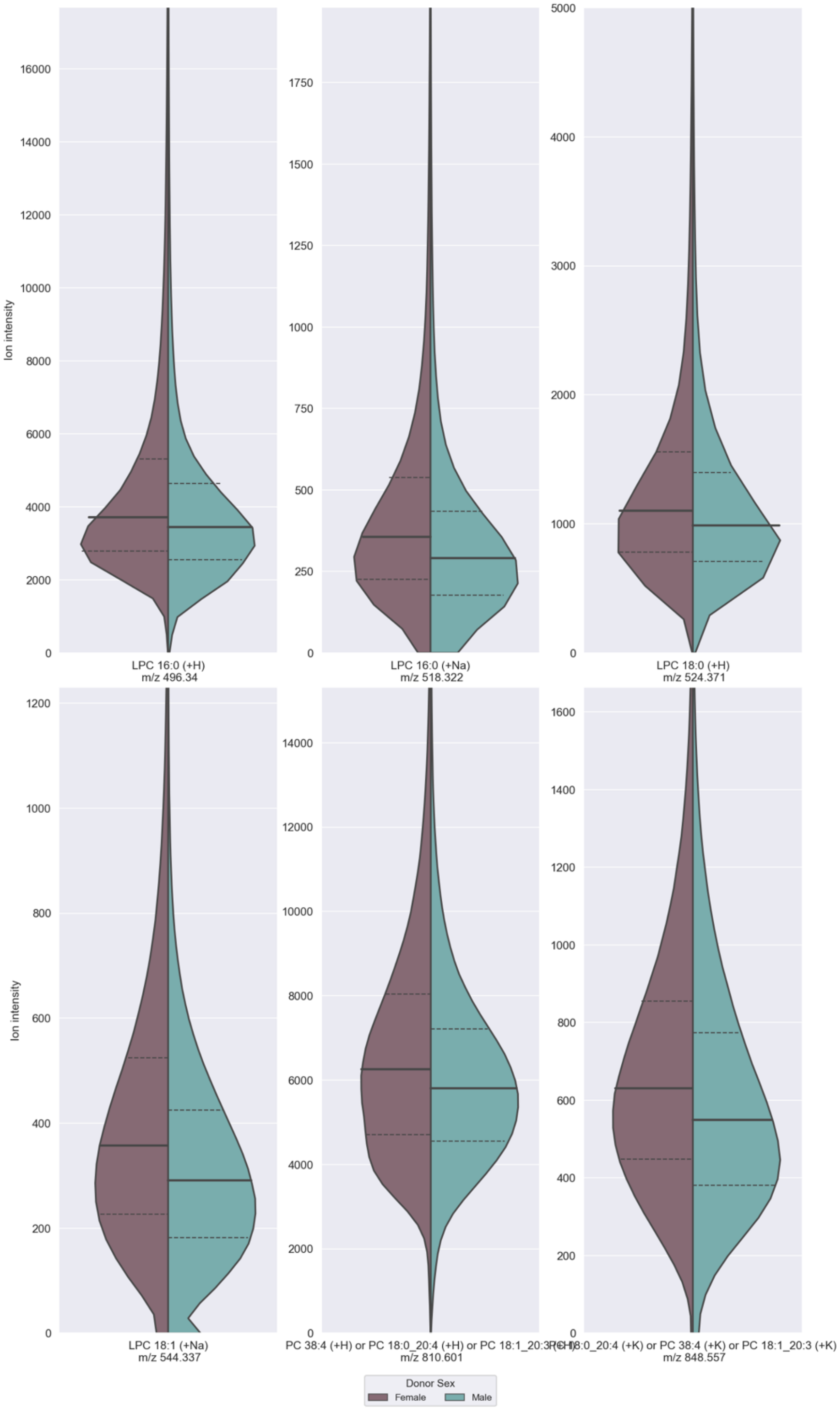

**Figure S162.** Split violin plots of the ion intensity distributions of **a selection of sex biomarker candidates in female (left) and male (right) donor data, in positive ion mode**, approximated using kernel density estimation. The violin plots are cropped at the 99th percentile of the distribution of one of the two classes (whichever is larger) to facilitate visual comparison. The full line indicates the median of each class' distribution, whereas the dashed lines indicate its interquartile range.

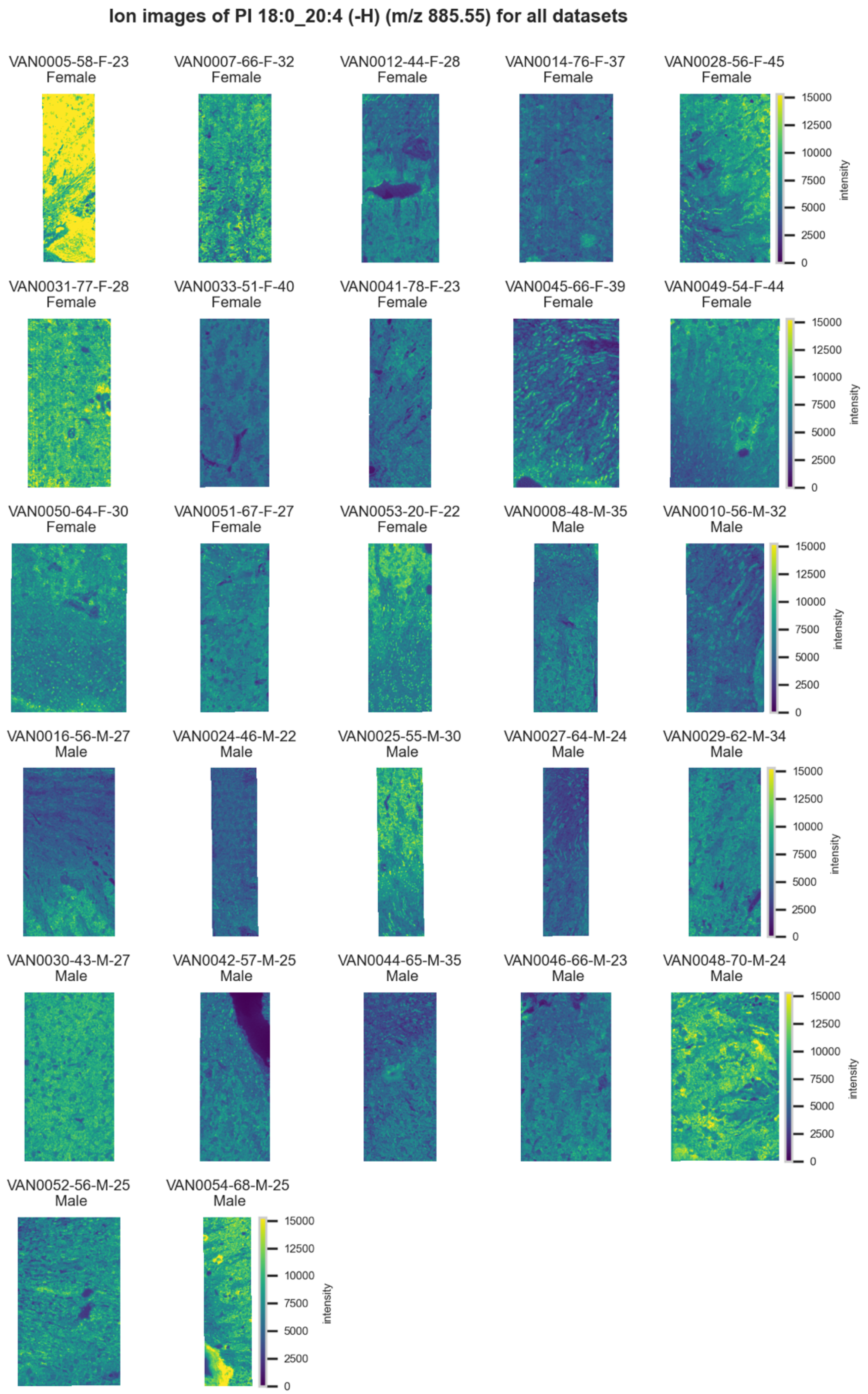

**Figure S163.** 10 µm MALDI IMS images of [PI(18:0_20:4)-H]$^-$ (*m/z* 885.55) detected in negative ion mode from each donor sample. Multivariate SHAP analysis determined this to be feature 1 for classification of sex.

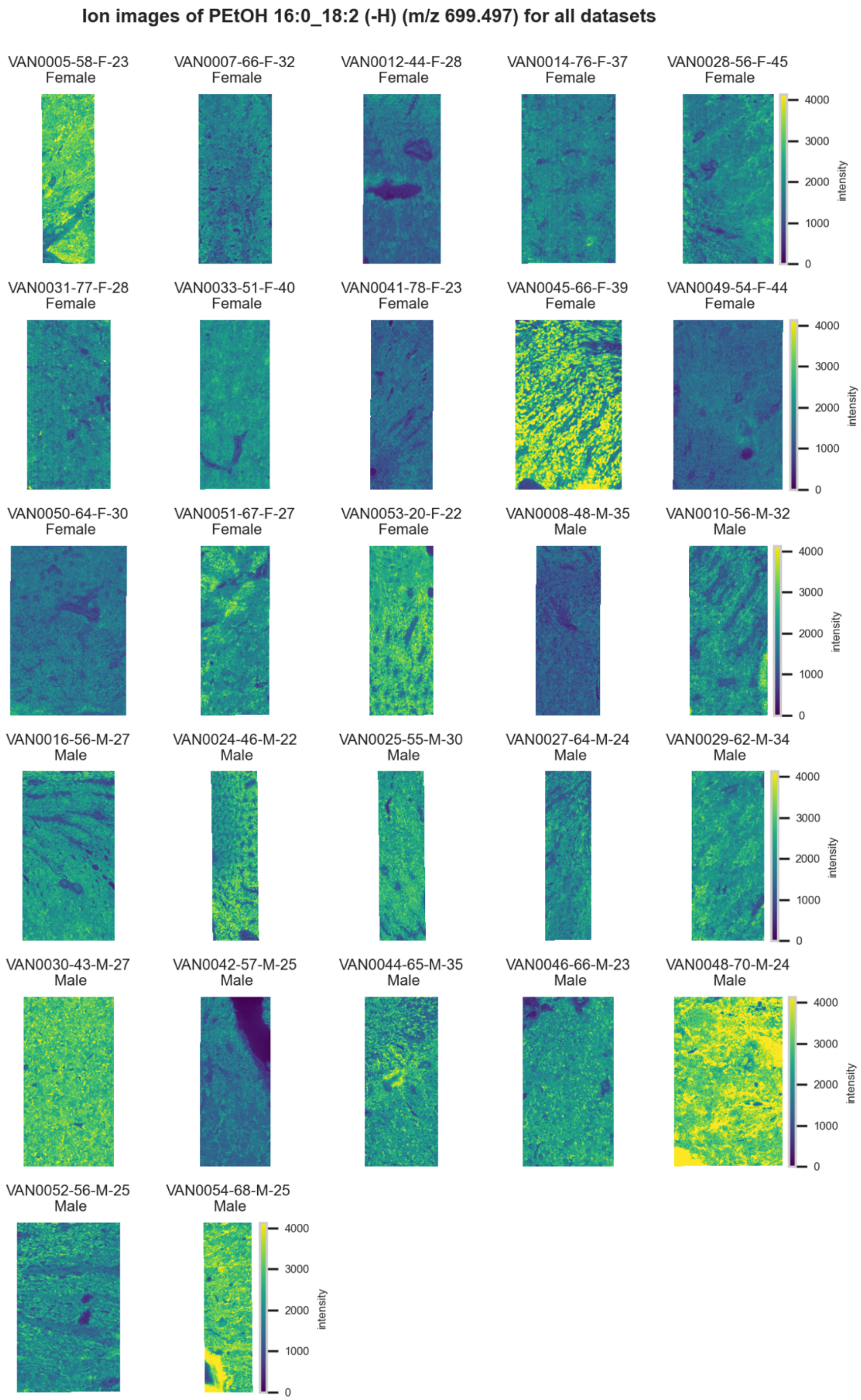

**Figure S164.** 10 µm MALDI IMS images of [PEtOH(16:0_18:2)-H]$^-$ (*m/z* 699.497) detected in <u>negative ion mode</u> from each donor sample. Multivariate SHAP analysis determined this to be feature 2 for classification of sex.

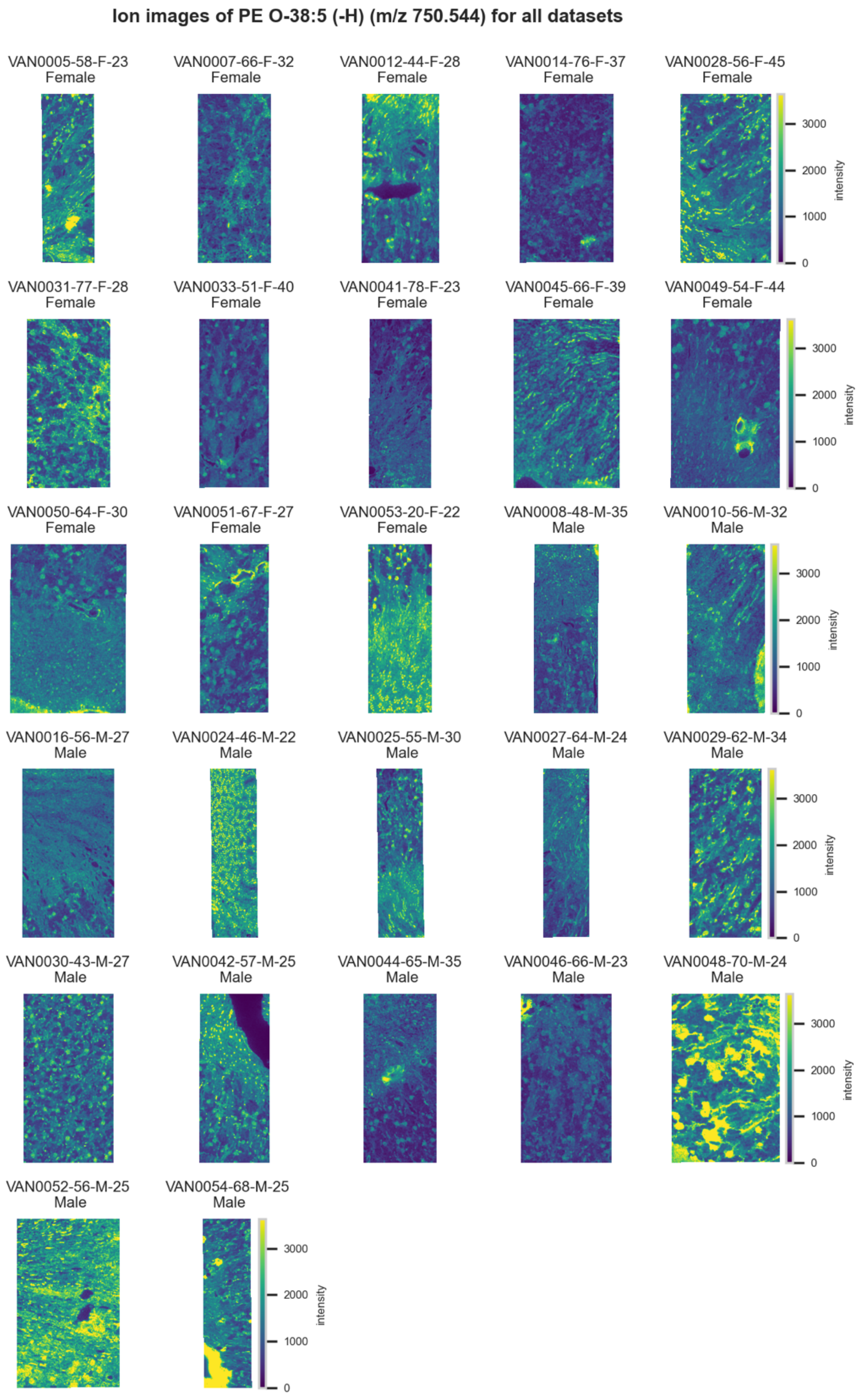

**Figure S165.** 10 µm MALDI IMS images of [PE(O-38:5)-H]$^-$ (*m/z* 750.544) detected in negative ion mode from each donor sample. Multivariate SHAP analysis determined this to be feature 3 for classification of sex.

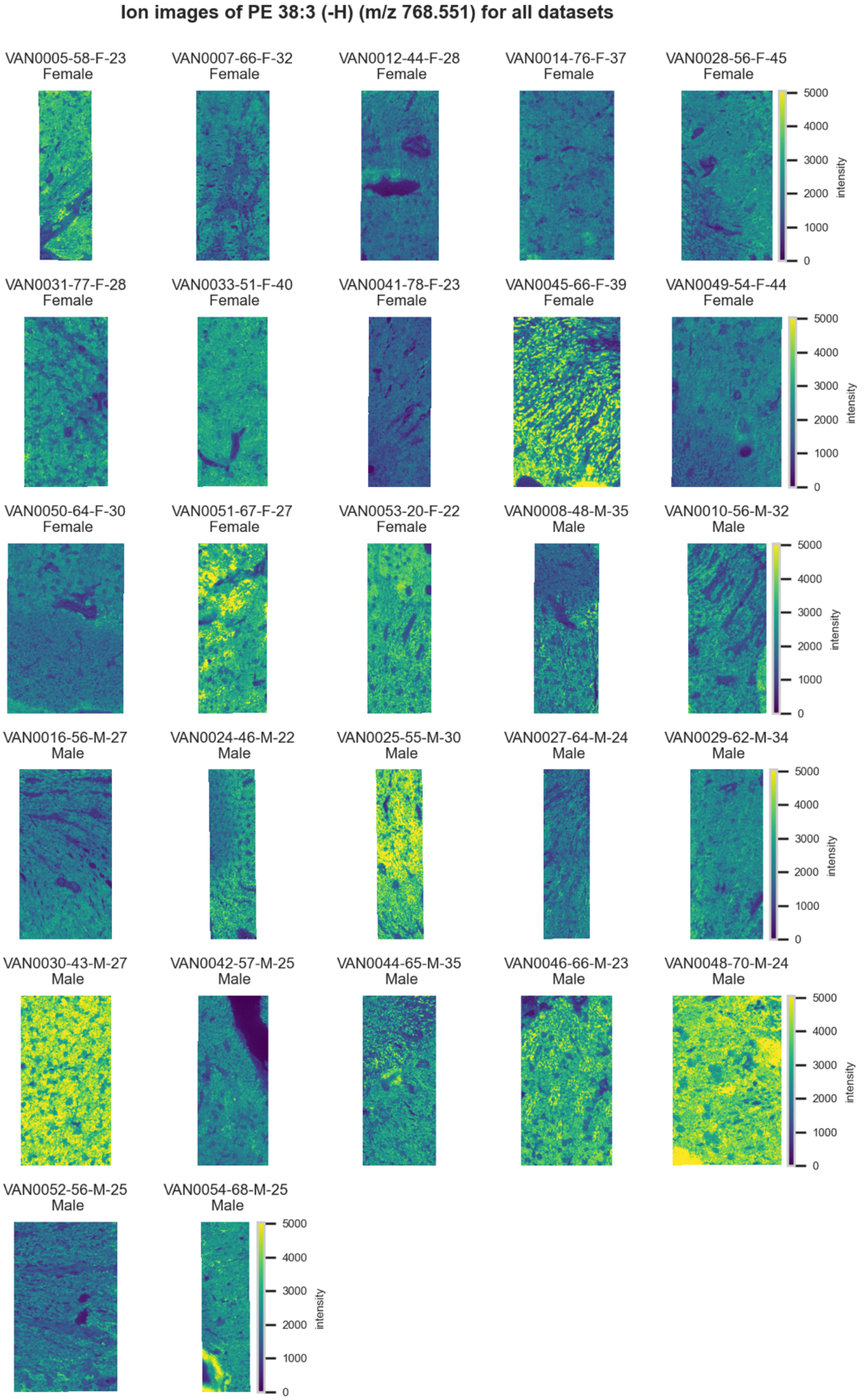

**Figure S166.** 10 µm MALDI IMS images of an unknown [PE(38:3)-H]$^-$ (*m/z* 768.551) detected in negative ion mode from each donor sample. Multivariate SHAP analysis determined this to be feature 4 for classification of sex.

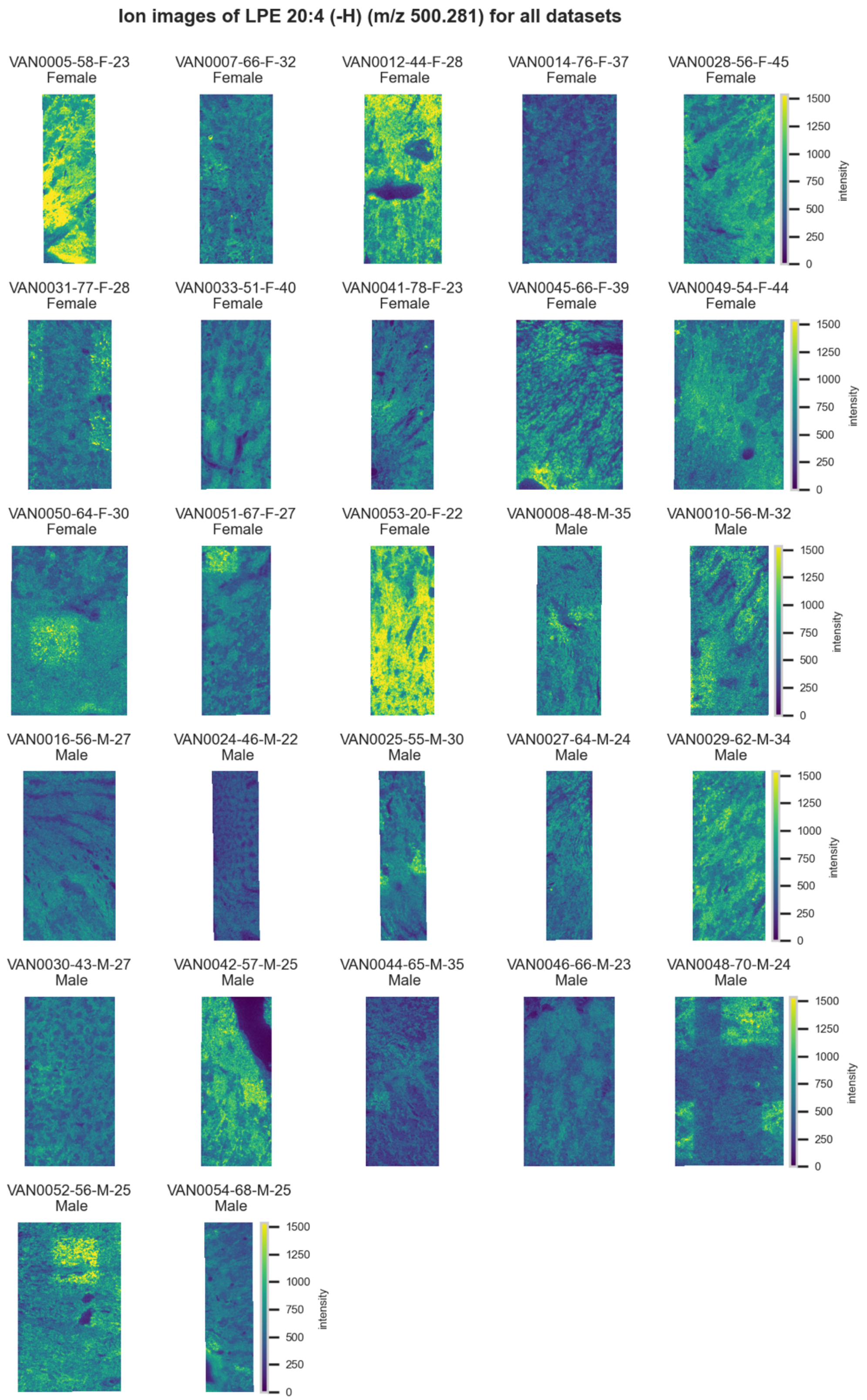

**Figure S167.** 10 µm MALDI IMS images of [LPE(20:4)-H]$^-$ (*m/z* 500.281) detected in <u>negative ion mode</u> from each donor sample. Multivariate SHAP analysis determined this to be feature 5 for classification of sex.

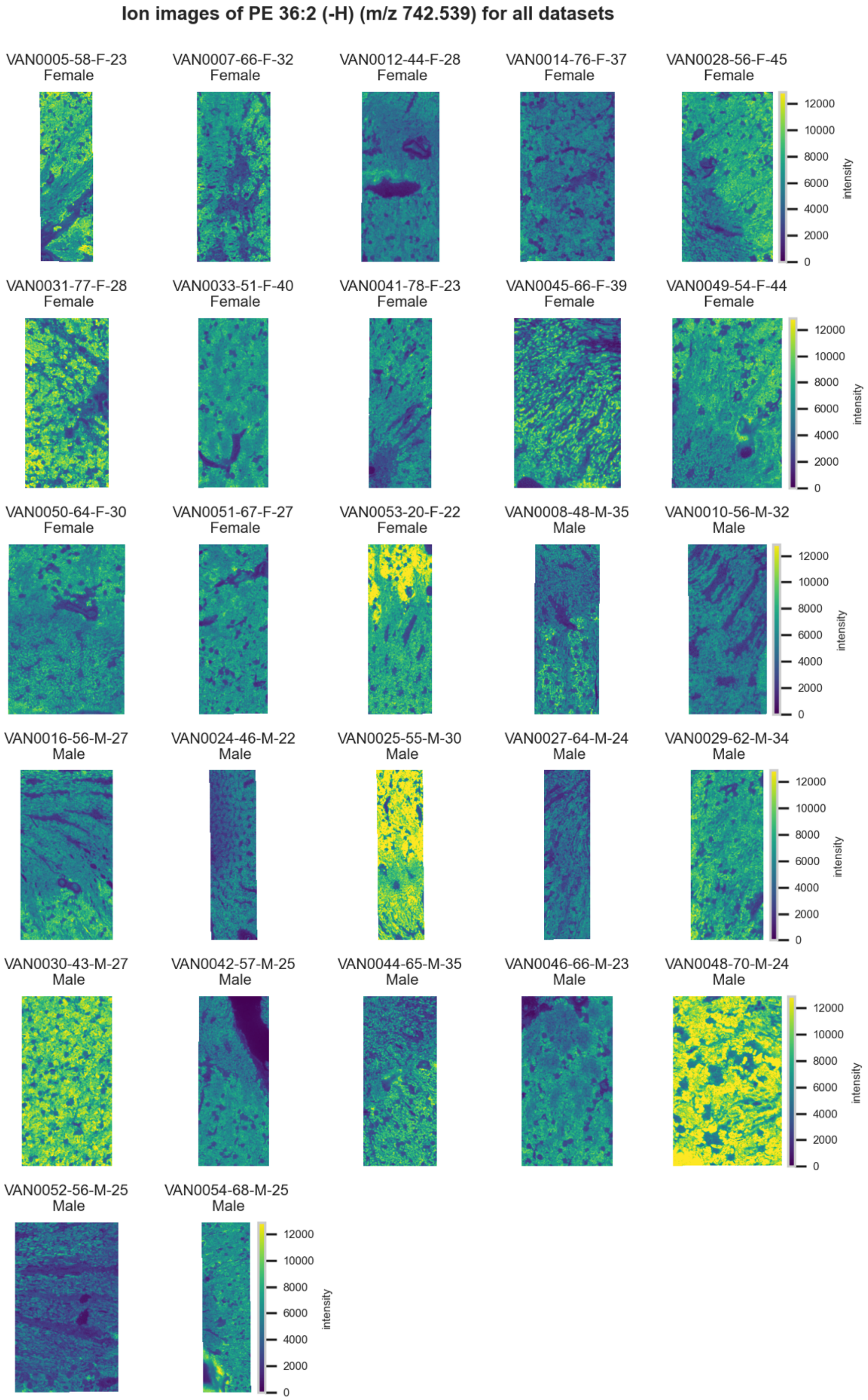

**Figure S168.** 10 µm MALDI IMS images of [PE(36:2)-H]- (*m/z* 742.539) detected in negative ion mode from each donor sample. Multivariate SHAP analysis determined this to be feature 6 for classification of sex.

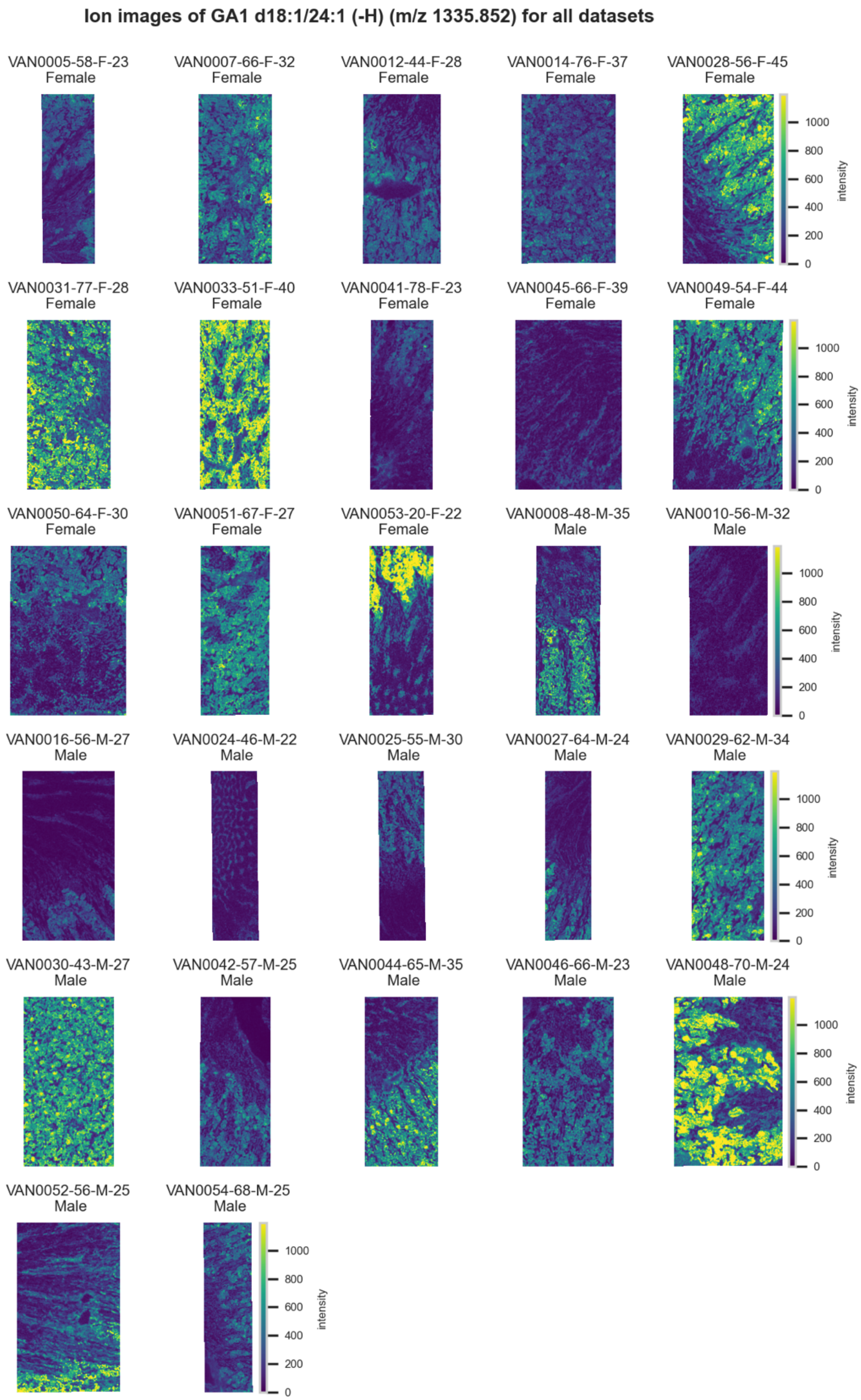

**Figure S169.** 10 µm MALDI IMS images of [GA1(d18:1/24:1)-H]$^-$ (*m/z* 1,335.852) detected in negative ion mode from each donor sample. Multivariate SHAP analysis determined this to be feature 7 for classification of sex.

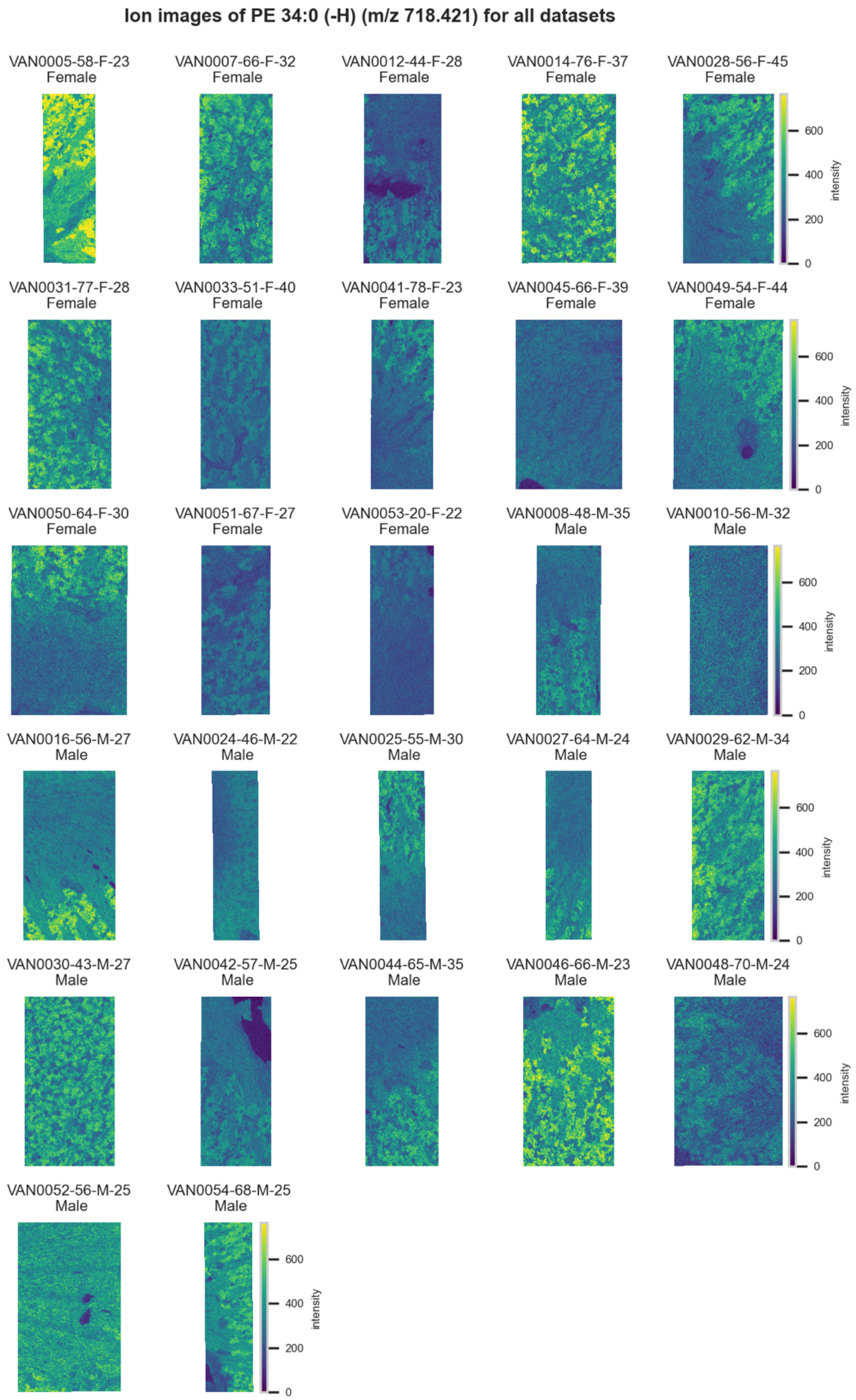

**Figure S170.** 10 μm MALDI IMS images of [PE(34:0)-H]$^-$ (*m/z* 718.421) detected in negative ion mode from each donor sample. Multivariate SHAP analysis determined this to be feature 8 for classification of sex.

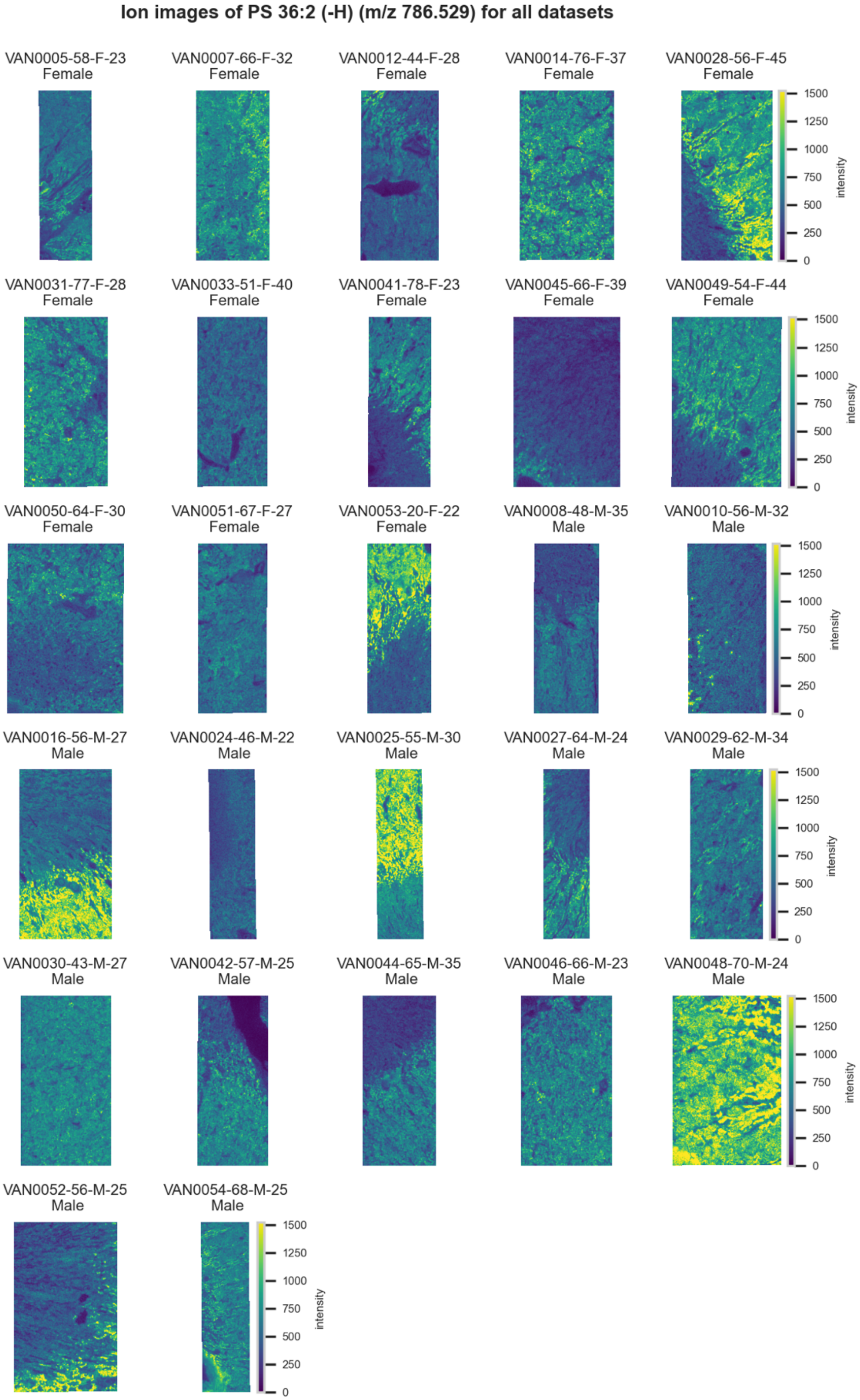

**Figure S171.** 10 µm MALDI IMS images of [PS(36:2)-H]$^-$ (*m/z* 786.529) detected in negative ion mode from each donor sample. Multivariate SHAP analysis determined this to be feature 9 for classification of sex.

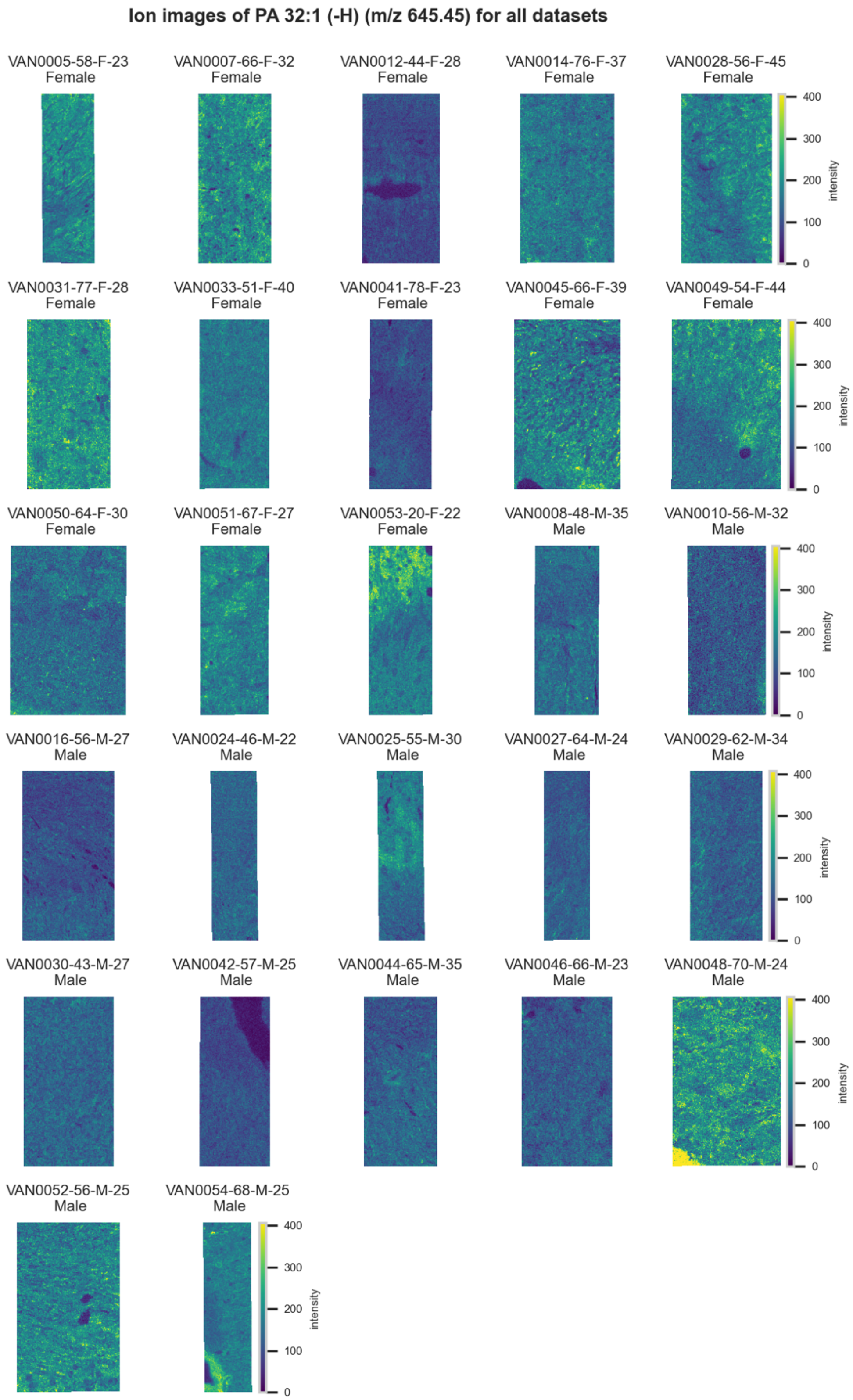

**Figure S172.** 10 µm MALDI IMS images of [PA(32:1)-H]$^-$ (*m/z* 645.45) detected in <u>negative ion mode</u> from each donor sample. Multivariate SHAP analysis determined this to be feature 10 for classification of sex.

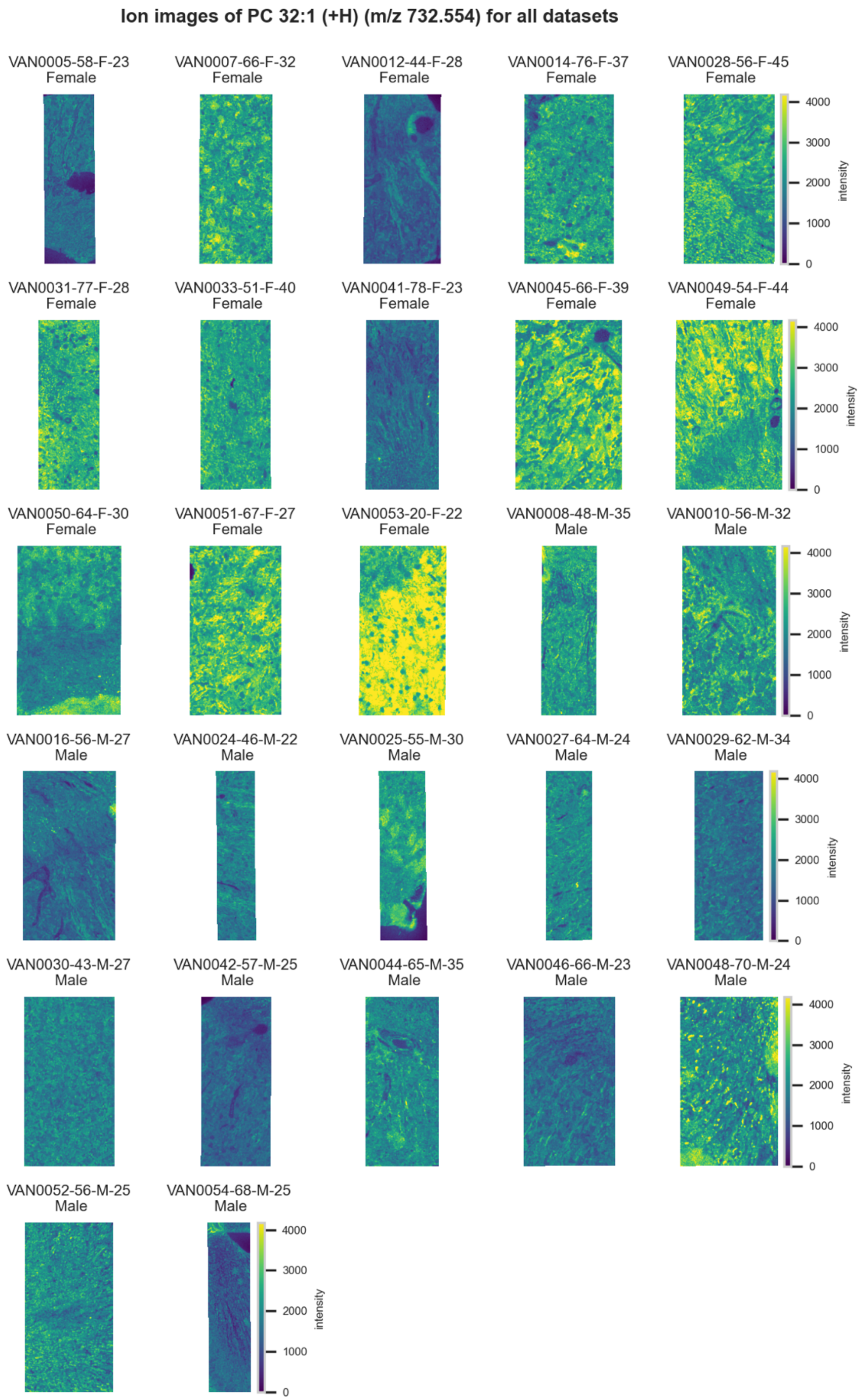

**Figure S173.** 10 µm MALDI IMS images of $[PC(32{:}1)+H]^+$ (*m/z* 732.554) detected in positive ion mode from each donor sample. Multivariate SHAP analysis determined this to be feature 1 for classification of sex.

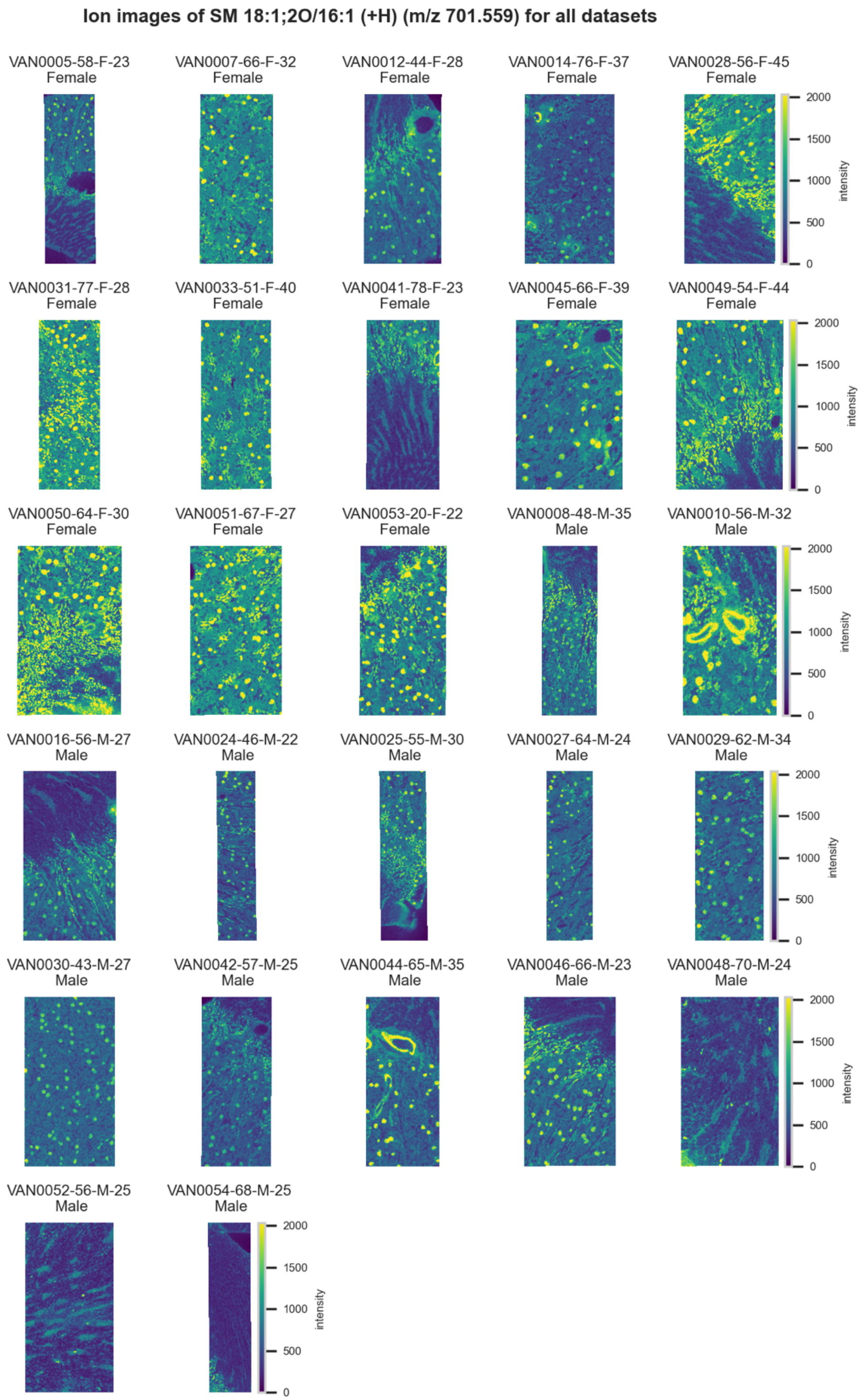

**Figure S174.** 10 µm MALDI IMS images of $[SM(18:1,2O/16:1)+H]^+$ (*m/z* 701.559) detected in positive ion mode from each donor sample. Multivariate SHAP analysis determined this to be feature 2 for classification of sex.

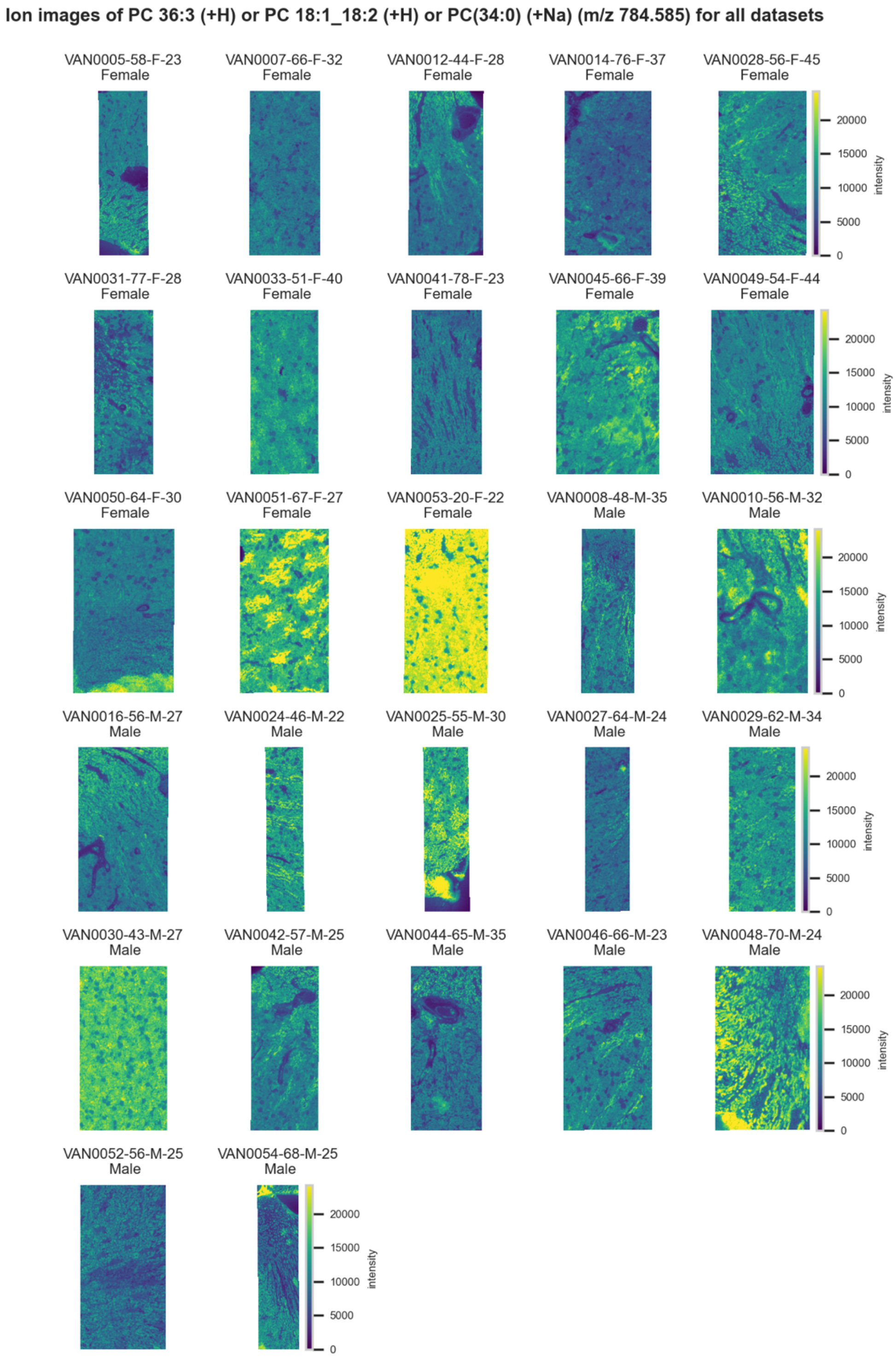

**Figure S175.** 10 µm MALDI IMS images of $[PC(36:3)+H]^+$ or $[PC(18:1\_18:2)+H]^+$ or $[PC(34:0)+Na]^+$ (*m/z* 784.585) detected in positive ion mode from each donor sample. These molecules are isomers or close isobars and therefore indistinguishable by MS alone. Multivariate SHAP analysis determined this to be feature 3 for classification of sex.

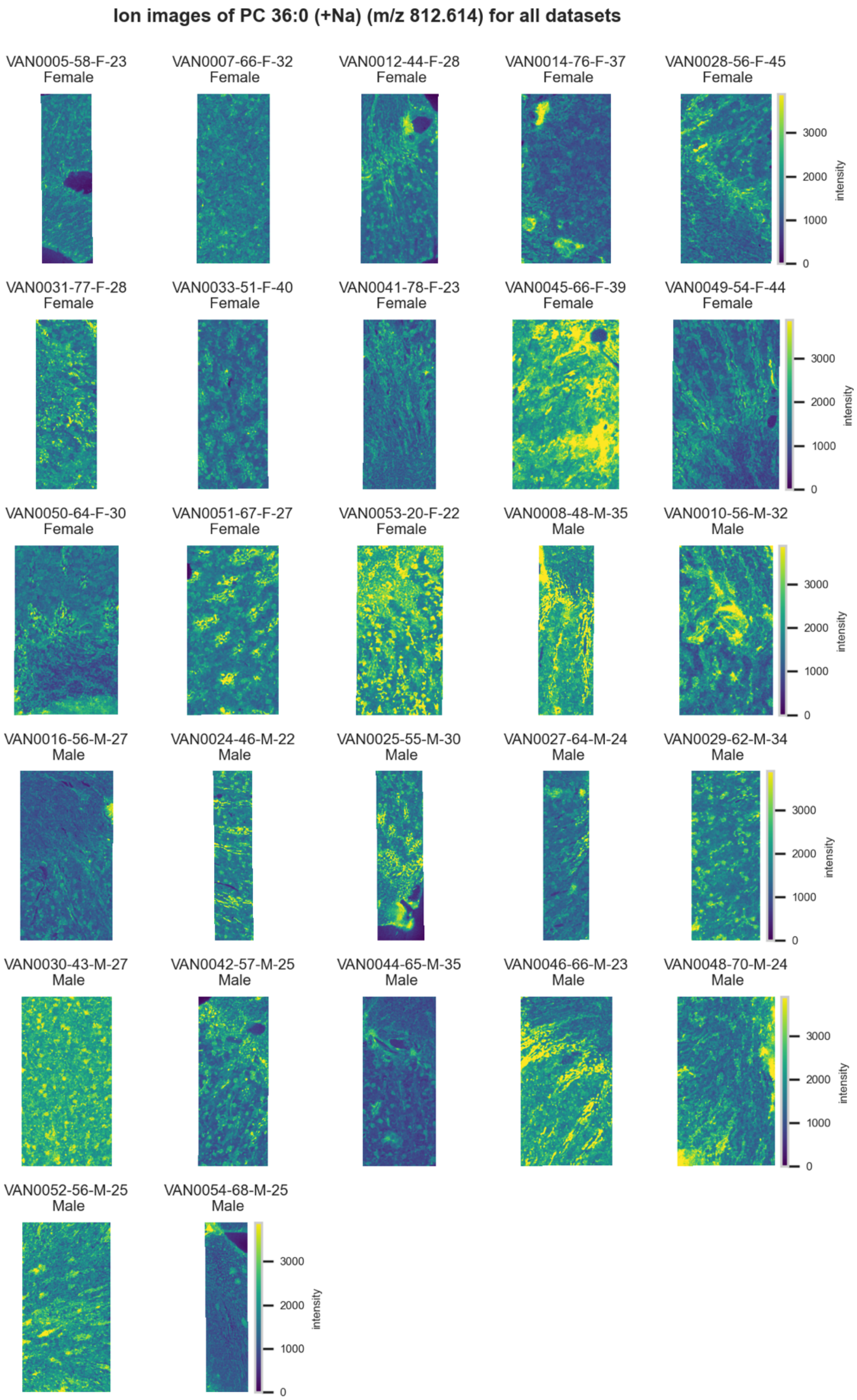

**Figure S176.** 10 μm MALDI IMS images of an unknown $[PE(36:0)+Na]^+$ (*m/z* 812.614) detected in <u>positive ion mode</u> from each donor sample. Multivariate SHAP analysis determined this to be feature 4 for classification of sex.

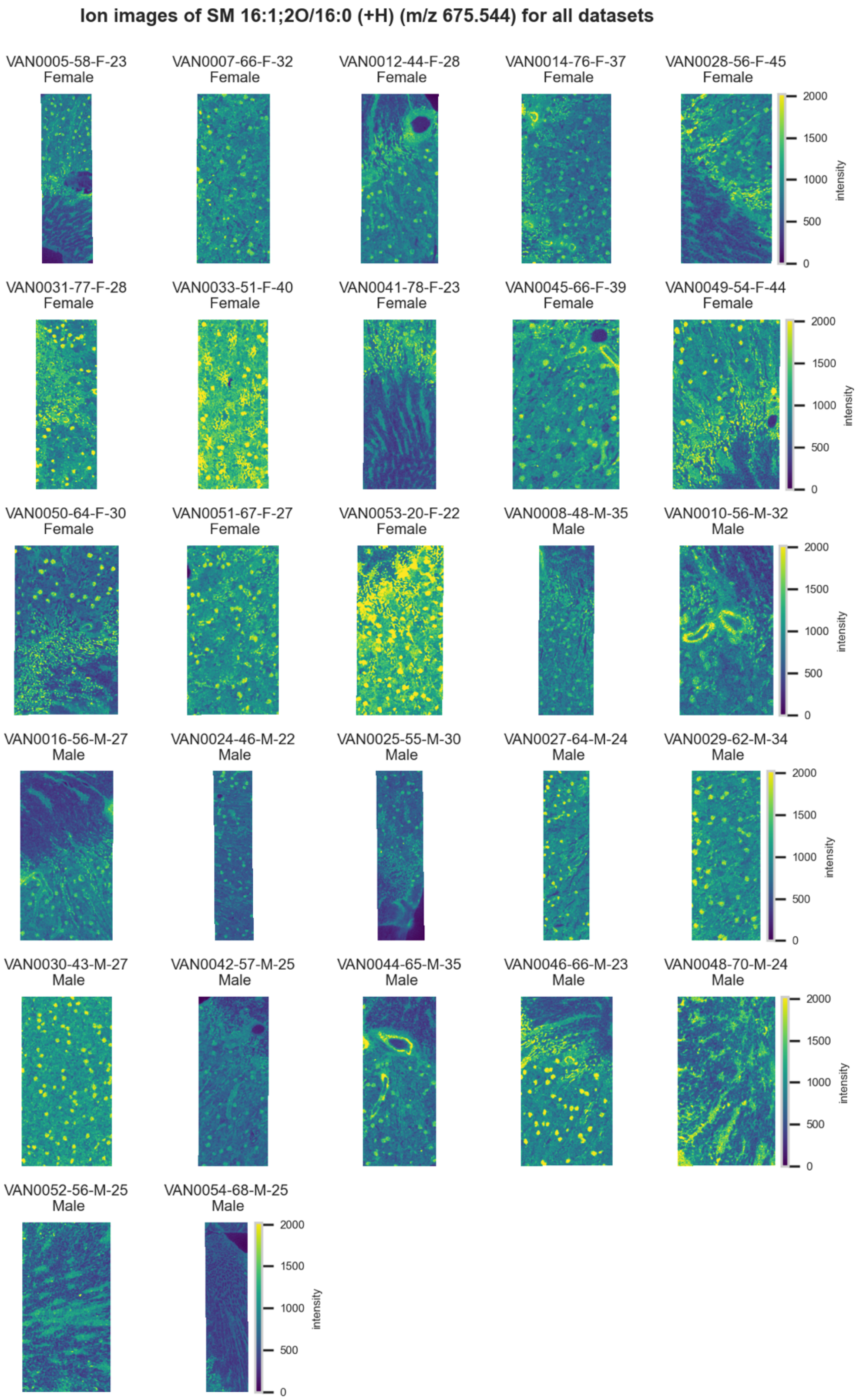

**Figure S177.** 10 µm MALDI IMS images of $[SM(16:1,2O/16:0)+H]^+$ (*m/z* 675.544) detected in positive ion mode from each donor sample. Multivariate SHAP analysis determined this to be feature 5 for classification of sex.

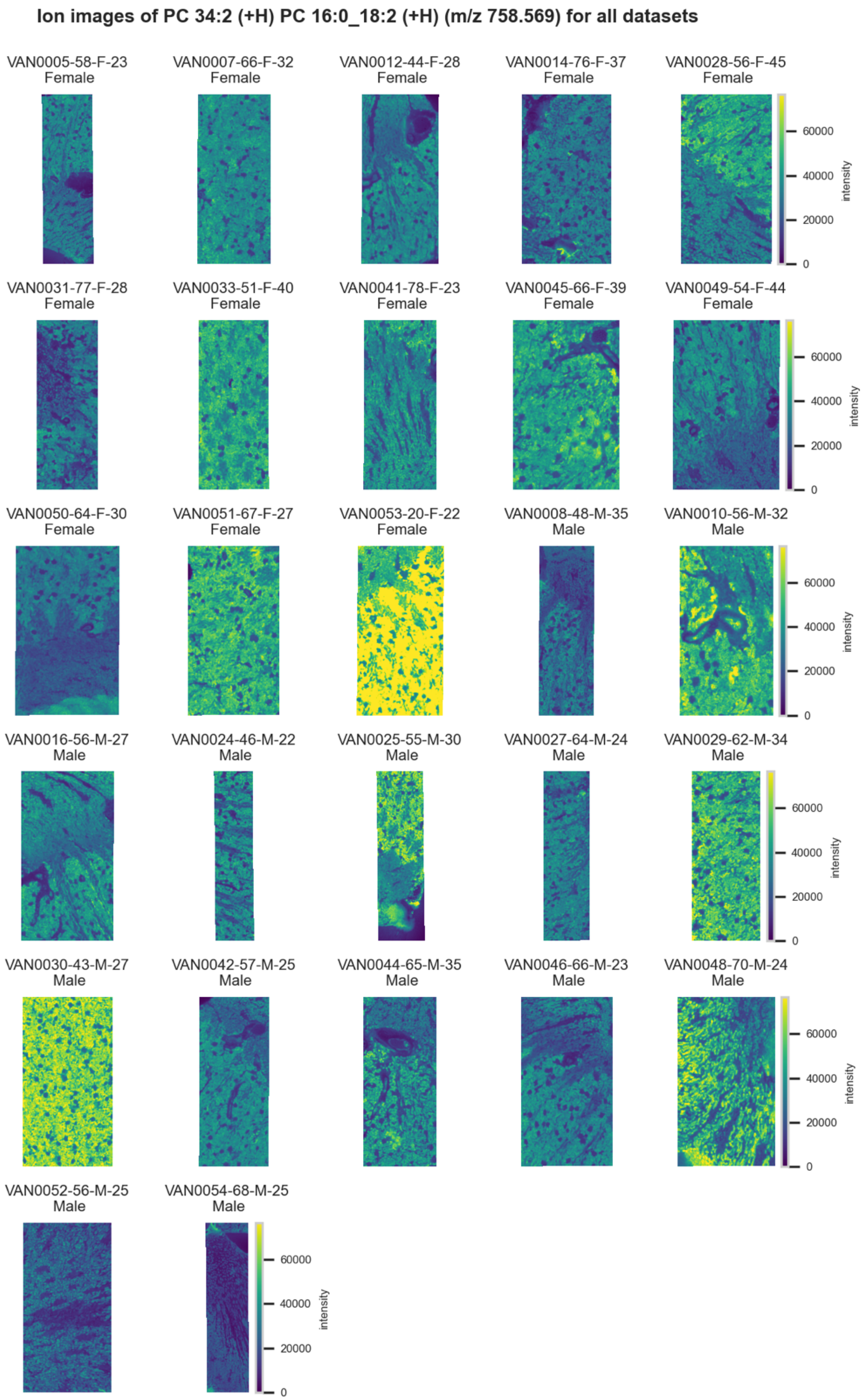

**Figure S178.** 10 µm MALDI IMS images of $[PC(34:2)+H]^+$ or $[PC(16:0\_18:2)+H]^+$ (*m/z* 758.569) detected in positive ion mode from each donor sample. These molecules are isomers and therefore indistinguishable by MS alone. Multivariate SHAP analysis determined this to be feature 6 for classification of sex.

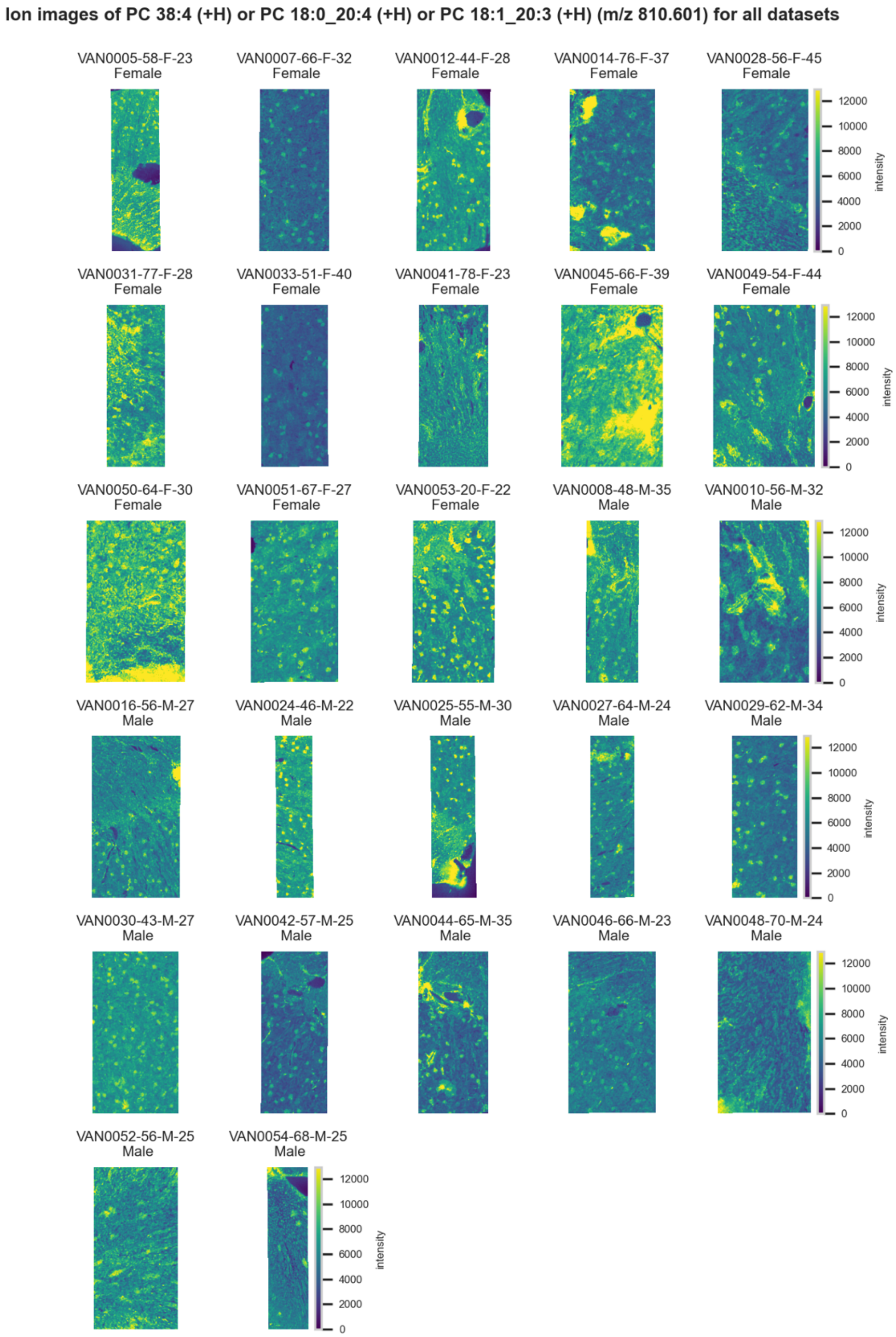

**Figure S179.** 10 µm MALDI IMS images of $[PC(38:4)+H]^+$ or $[PC(18:0\_20:4)+H]^+$ or $[PC(18:1\_20:3)+H]^+$ (*m/z* 810.601) detected in positive ion mode from each donor sample. These molecules are isomers and therefore indistinguishable by MS alone. Multivariate SHAP analysis determined this to be feature 7 for classification of sex.

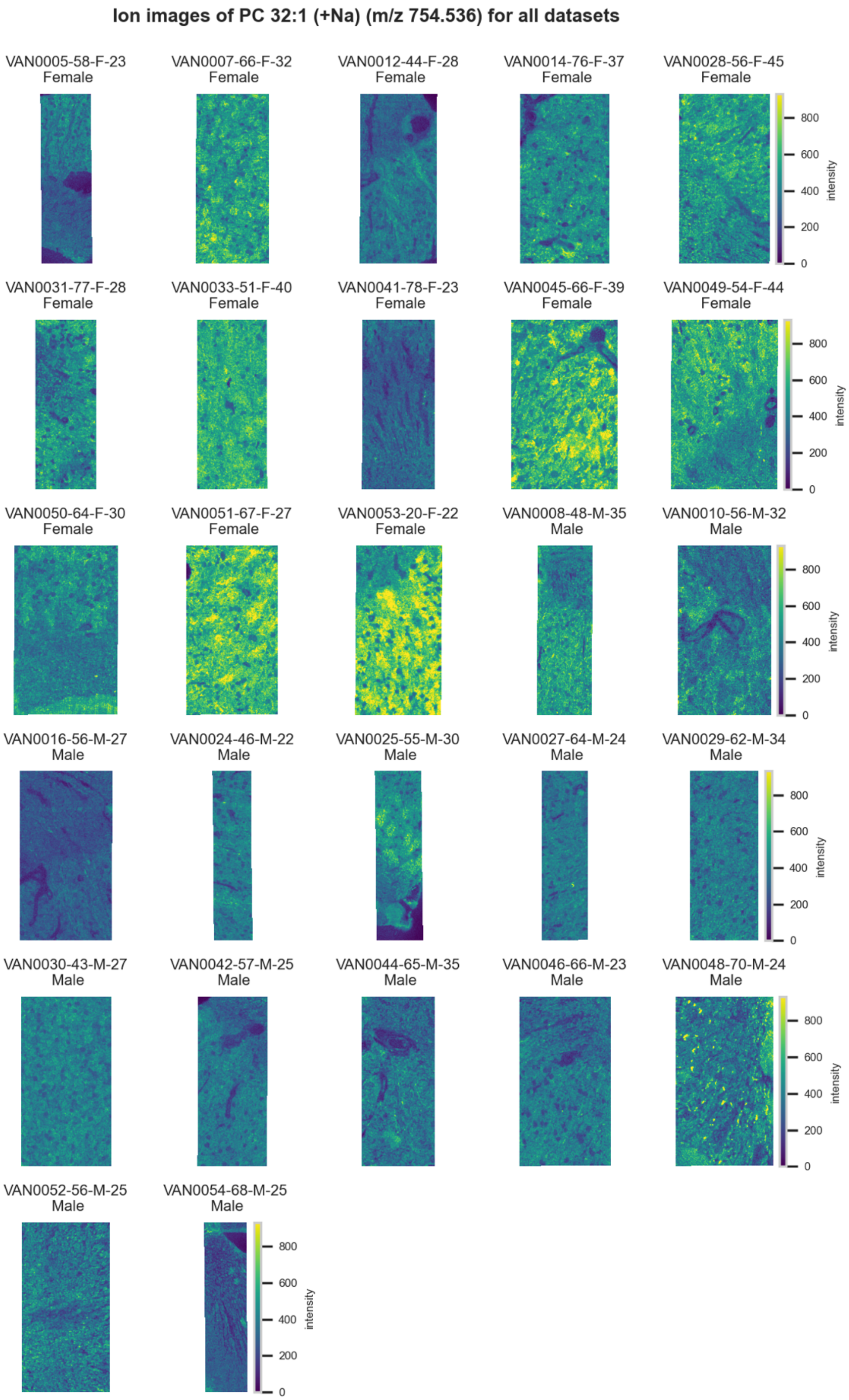

**Figure S180.** 10 μm MALDI IMS images of $[PC(32:1)+Na]^+$ (*m/z* 754.536) detected in positive ion mode from each donor sample. Multivariate SHAP analysis determined this to be feature 8 for classification of sex.

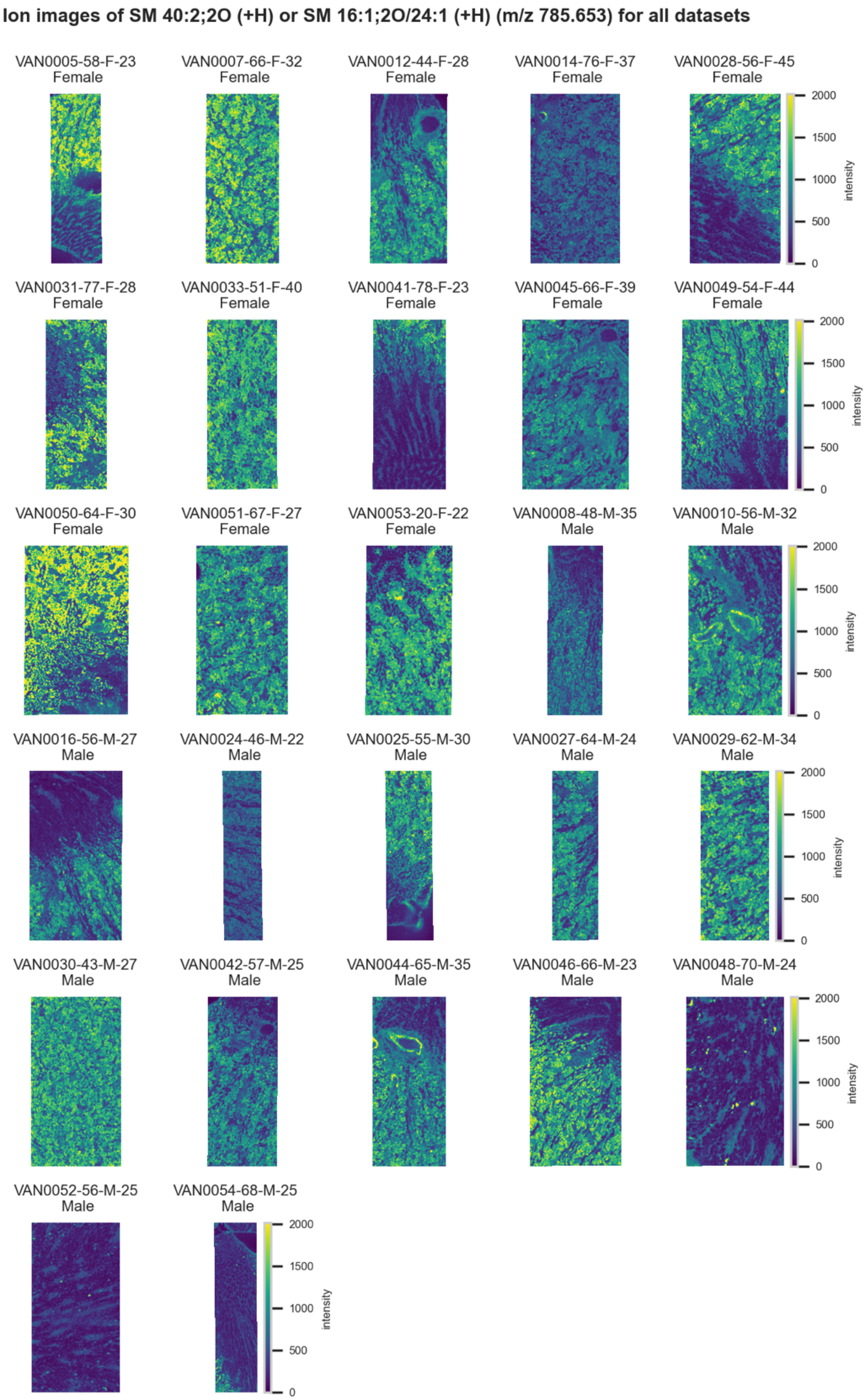

**Figure S181.** 10 µm MALDI IMS images of $[SM(40:2,2O)+H]^+$ or $[SM(16:1,2O/24:1)+H]^+$ (*m/z* 785.653) detected in positive ion mode from each donor sample. These molecules are isomers and therefore indistinguishable by MS alone. Multivariate SHAP analysis determined this to be feature 9 for classification of sex.

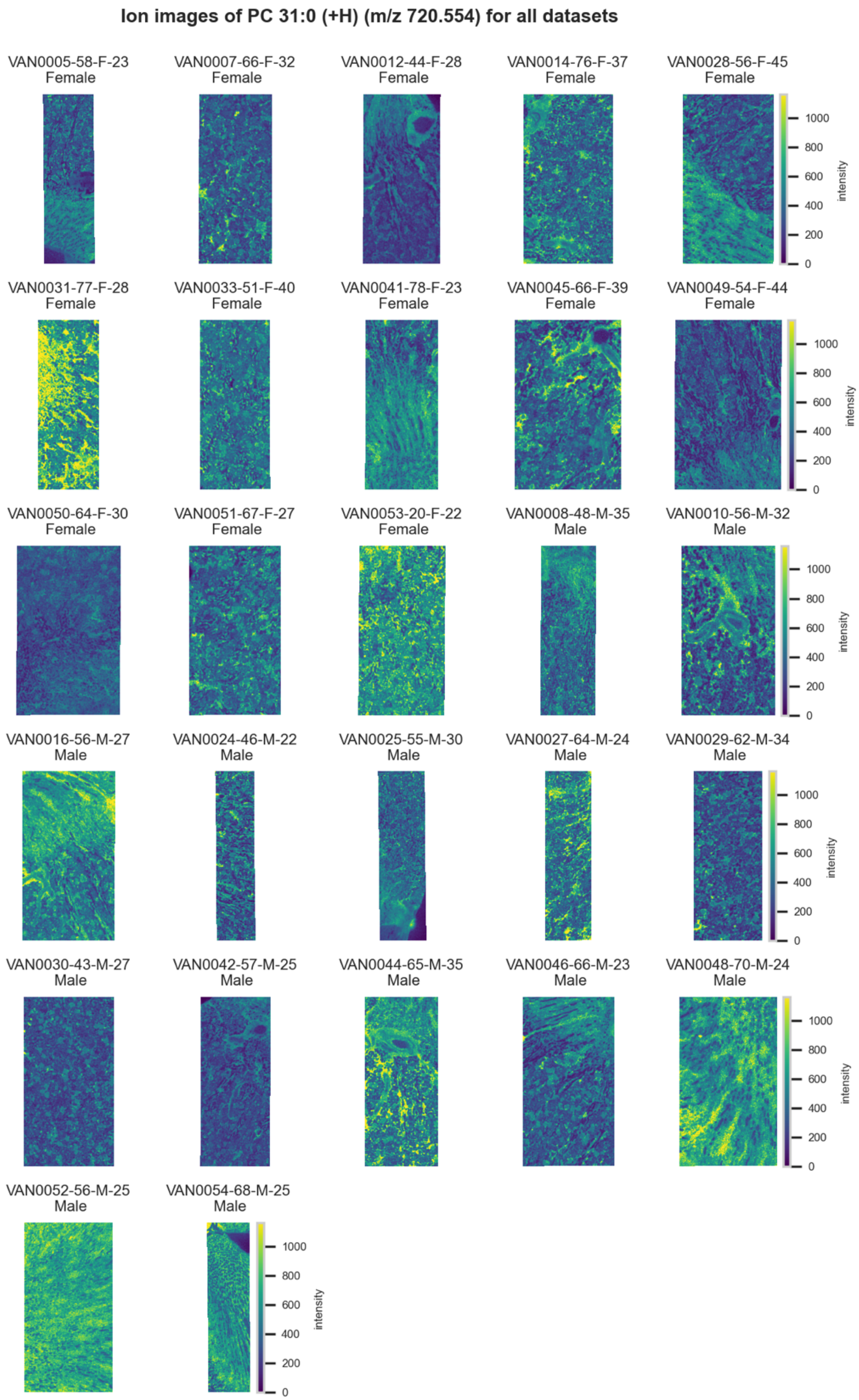

**Figure S182.** 10 µm MALDI IMS images of $[PC(31:0)+H]^+$ (*m/z* 720.554) detected in positive ion mode from each donor sample. Multivariate SHAP analysis determined this to be feature 10 for classification of sex.

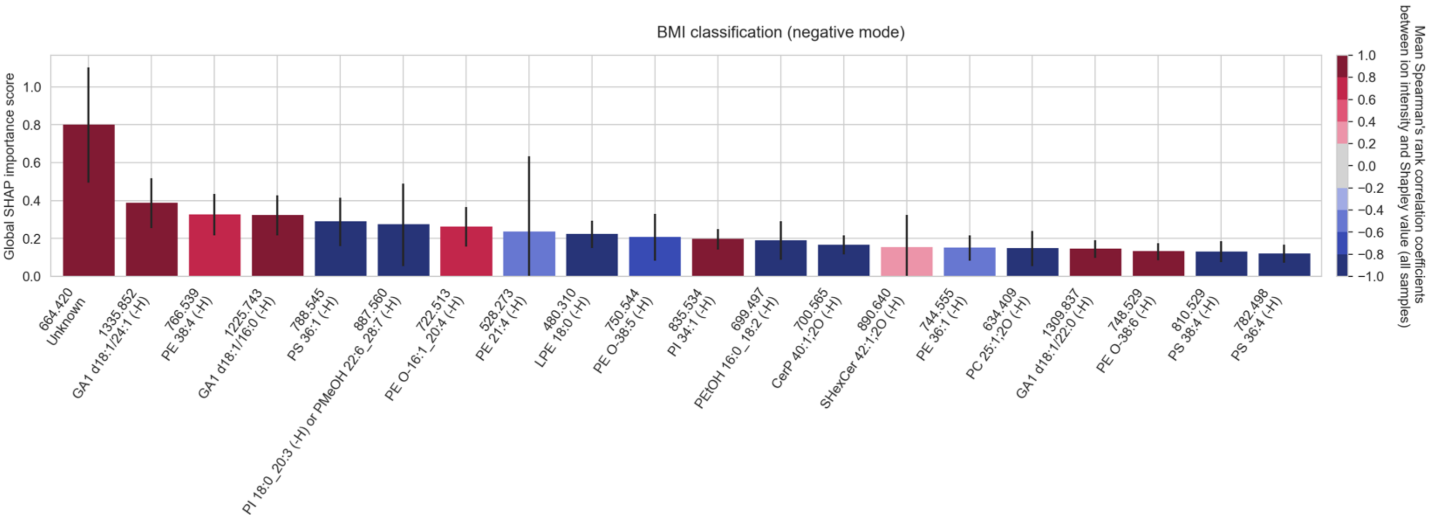

**Figure S183.** The top twenty cohort-wide (absolute) SHAP importance scores reporting the top twenty molecular species among the measured lipid species that enable the classification of **obese donor tissue from normal donor tissue overall (not specific to an FTU), in negative ion mode**. The error bar is the standard deviation across all donor samples of the tissue sample-wide SHAP importance score for a given molecular species. The color of the bar reports the mean Spearman's rank correlation coefficient (across all donor samples) between the molecular species' mean-centered ion intensity and its Shapley score. Given that the BMI classification task is binary (normal-BMI versus high-BMI), we made the high-BMI class the target class of our classification model. A positive Spearman's rank correlation coefficient indicates that high intensity of the molecular species correlates with obesity. Molecular species that are positively correlated with high-BMI are negatively correlated with normal-BMI, and vice-versa (refer to Figure 8).

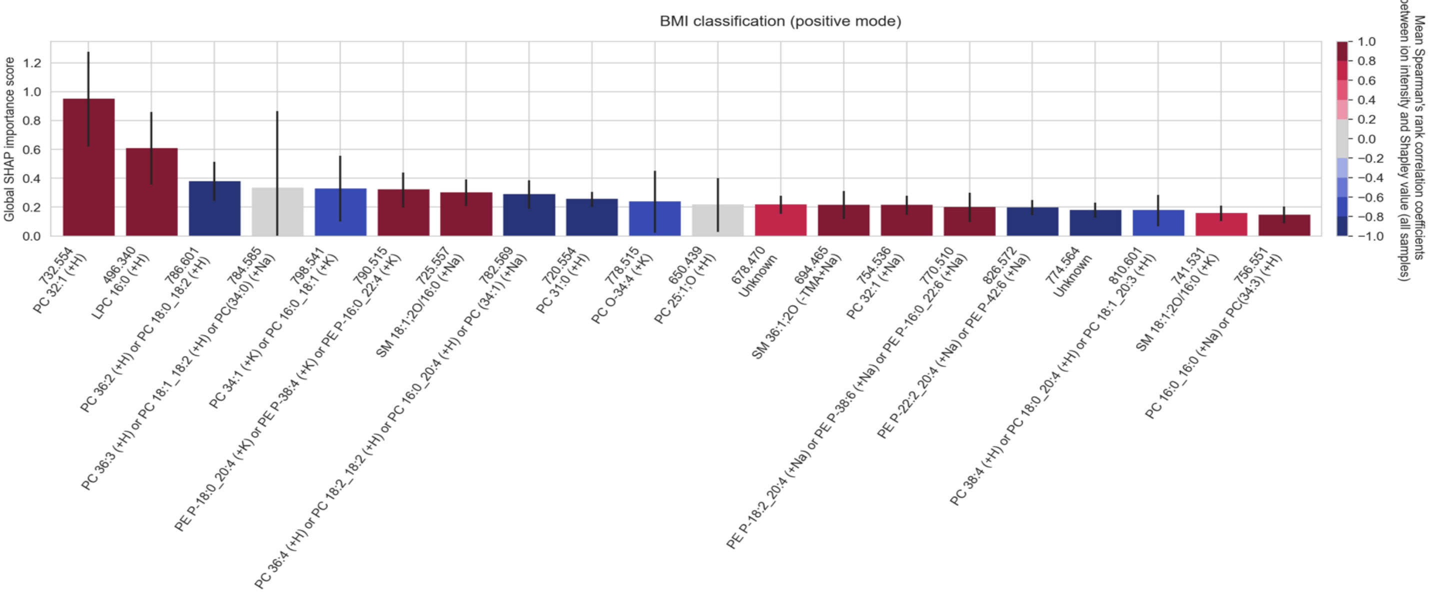

**Figure S184.** The top twenty cohort-wide (absolute) SHAP importance scores reporting the top twenty molecular species among the measured lipid species that enable the classification of **obese donor tissue from normal donor tissue overall (not specific to an FTU), in positive ion mode**. The error bar is the standard deviation across all donor samples of the tissue sample-wide SHAP importance score for a given molecular species. The color of the bar reports the mean Spearman's rank correlation coefficient (across all donor samples) between the molecular species' mean-centered ion intensity and its Shapley score. Given that the BMI classification task is binary (normal-BMI versus high-BMI), we made the high-BMI class the target class of our classification model. A positive Spearman's rank correlation coefficient indicates that high intensity of the molecular species correlates with obesity. Molecular species that are positively correlated with high-BMI are negatively correlated with normal-BMI, and vice-versa (refer to Figure 8).

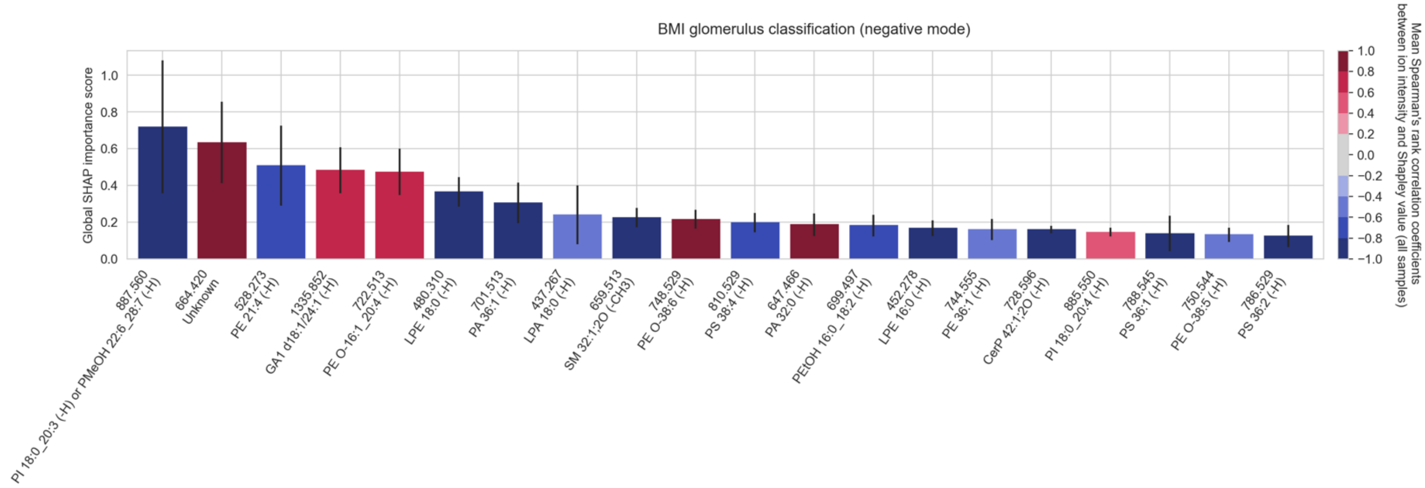

**Figure S185.** The top twenty cohort-wide (absolute) SHAP importance scores reporting the top twenty molecular species among the measured lipid species that enable the classification of **the glomeruli of obese donors from the glomeruli of normal donors, in negative ion mode**. The error bar is the standard deviation across all donor samples of the tissue sample-wide SHAP importance score for a given molecular species. The color of the bar reports the mean Spearman's rank correlation coefficient (across all donor samples) between the molecular species' mean-centered ion intensity and its Shapley score. A positive Spearman's rank correlation coefficient indicates that high intensity of the molecular species correlates with obesity. Conversely, a negative Spearman's rank correlation coefficient indicates that low intensity of the molecular species correlates with obesity.

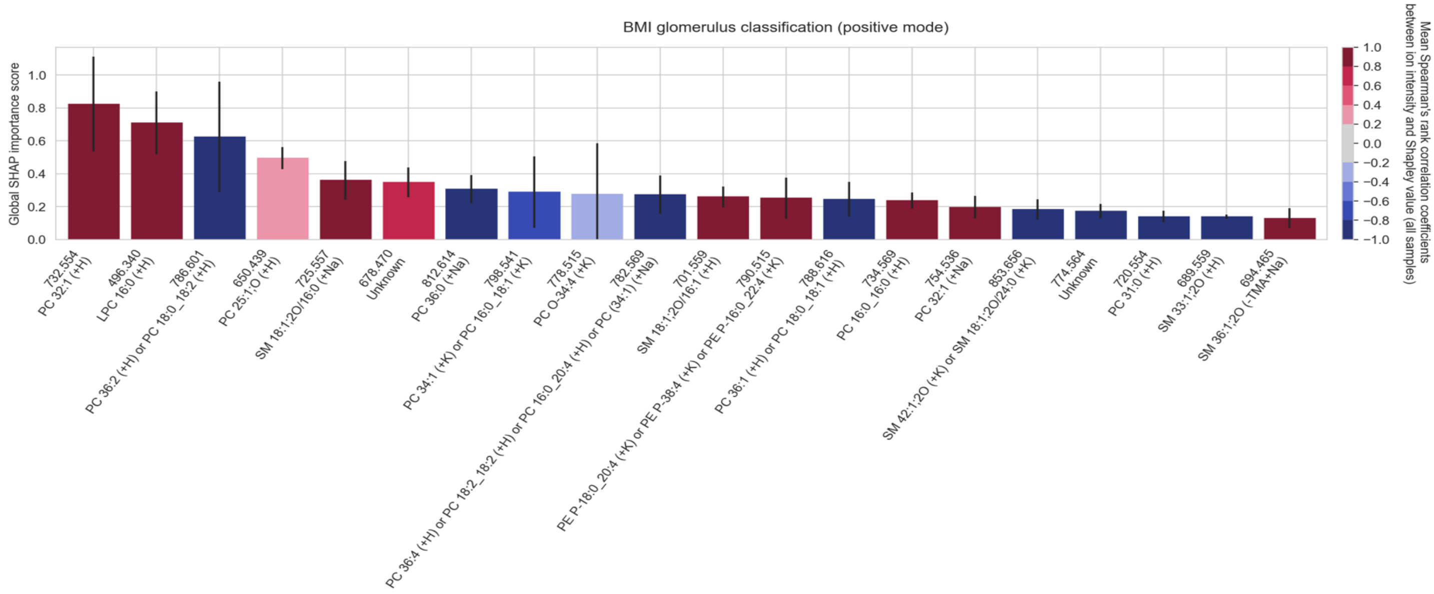

**Figure S186.** The top twenty cohort-wide (absolute) SHAP importance scores reporting the top twenty molecular species among the measured lipid species that enable the classification of **the glomeruli of obese donors from the glomeruli of normal donors, in positive ion mode**. The error bar is the standard deviation across all donor samples of the tissue sample-wide SHAP importance score for a given molecular species. The color of the bar reports the mean Spearman's rank correlation coefficient (across all donor samples) between the molecular species' mean-centered ion intensity and its Shapley score. A positive Spearman's rank correlation coefficient indicates that high intensity of the molecular species correlates with obesity. Conversely, a negative Spearman's rank correlation coefficient indicates that low intensity of the molecular species correlates with obesity.

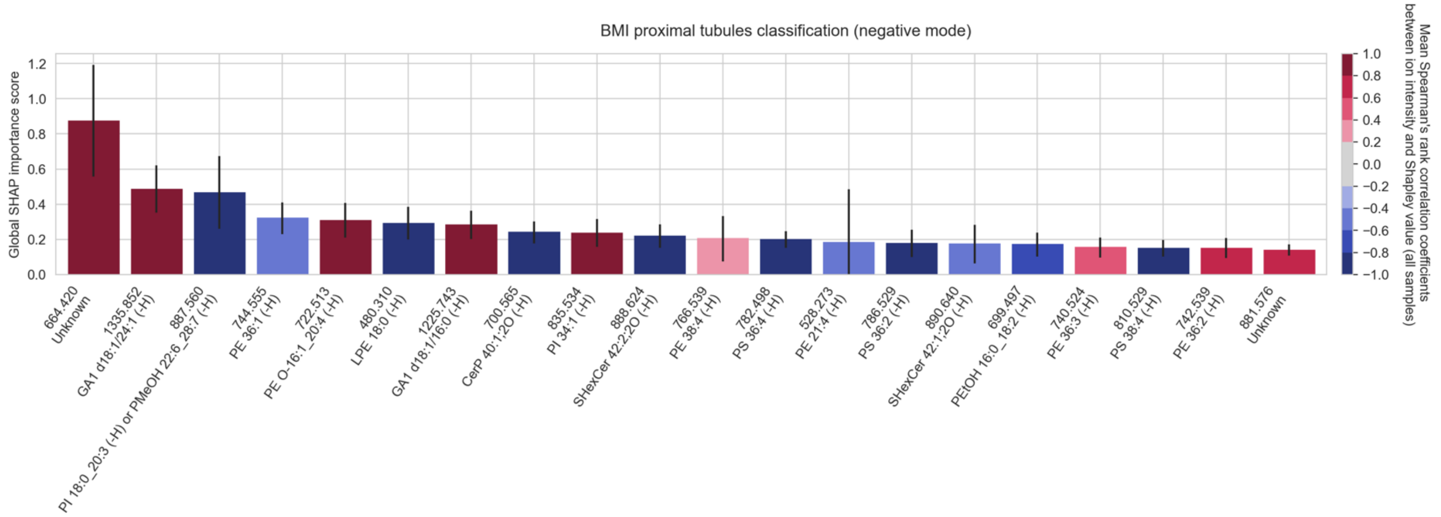

**Figure S187.** The top twenty cohort-wide (absolute) SHAP importance scores reporting the top twenty molecular species among the measured lipid species that enable the classification of **the proximal tubules of obese donors from the proximal tubules of normal donors, in negative ion mode**. The error bar is the standard deviation across all donor samples of the tissue sample-wide SHAP importance score for a given molecular species. The color of the bar reports the mean Spearman's rank correlation coefficient (across all donor samples) between the molecular species' mean-centered ion intensity and its Shapley score. A positive Spearman's rank correlation coefficient indicates that high intensity of the molecular species correlates with obesity. Conversely, a negative Spearman's rank correlation coefficient indicates that low intensity of the molecular species correlates with obesity.

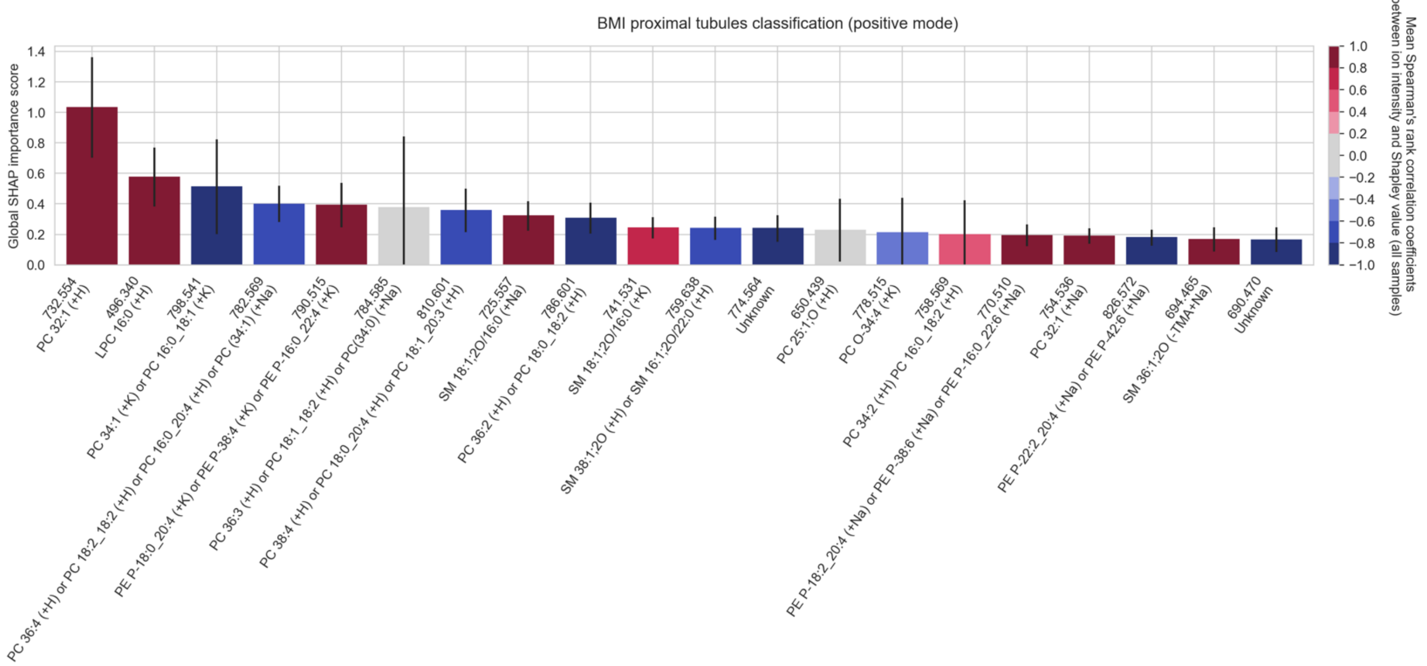

**Figure S188.** The top twenty cohort-wide (absolute) SHAP importance scores reporting the top twenty molecular species among the measured lipid species that enable the classification of **the proximal tubules of obese donors from the proximal tubules of normal donors, in positive ion mode**. The error bar is the standard deviation across all donor samples of the tissue sample-wide SHAP importance score for a given molecular species. The color of the bar reports the mean Spearman's rank correlation coefficient (across all donor samples) between the molecular species' mean-centered ion intensity and its Shapley score. A positive Spearman's rank correlation coefficient indicates that high intensity of the molecular species correlates with obesity. Conversely, a negative Spearman's rank correlation coefficient indicates that low intensity of the molecular species correlates with obesity.

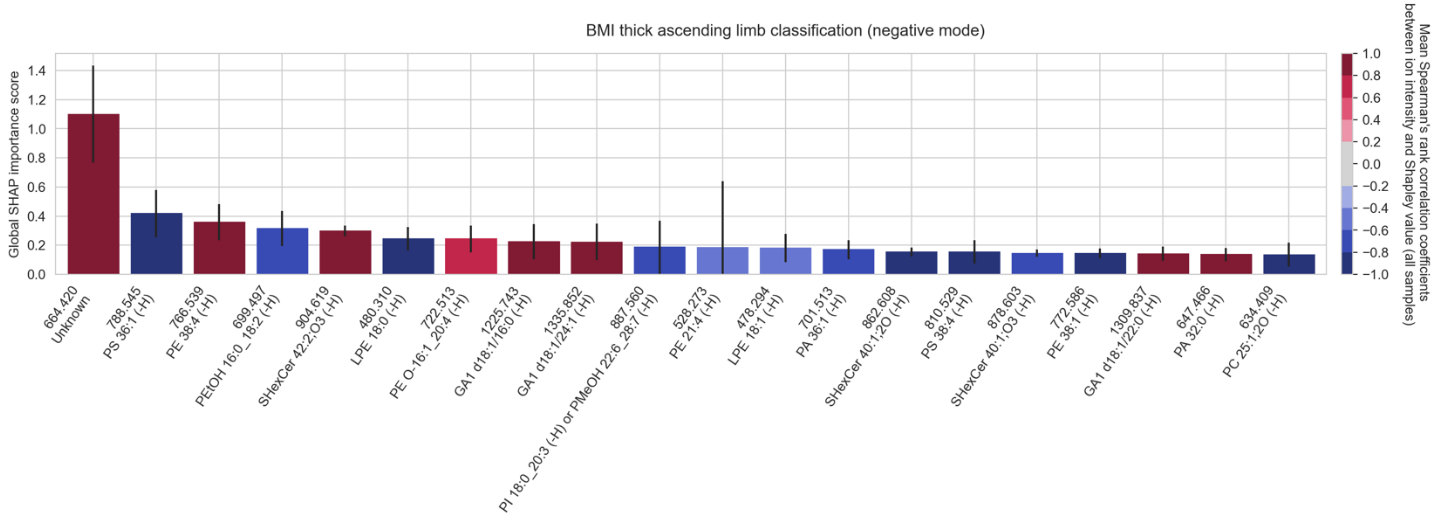

**Figure S189.** The top twenty cohort-wide (absolute) SHAP importance scores reporting the top twenty molecular species among the measured lipid species that enable the classification of **the thick ascending limb of obese donors from the thick ascending limb of normal donors, in negative ion mode**. The error bar is the standard deviation across all donor samples of the tissue sample-wide SHAP importance score for a given molecular species. The color of the bar reports the mean Spearman's rank correlation coefficient (across all donor samples) between the molecular species' mean-centered ion intensity and its Shapley score. A positive Spearman's rank correlation coefficient indicates that high intensity of the molecular species correlates with obesity. Conversely, a negative Spearman's rank correlation coefficient indicates that low intensity of the molecular species correlates with obesity.

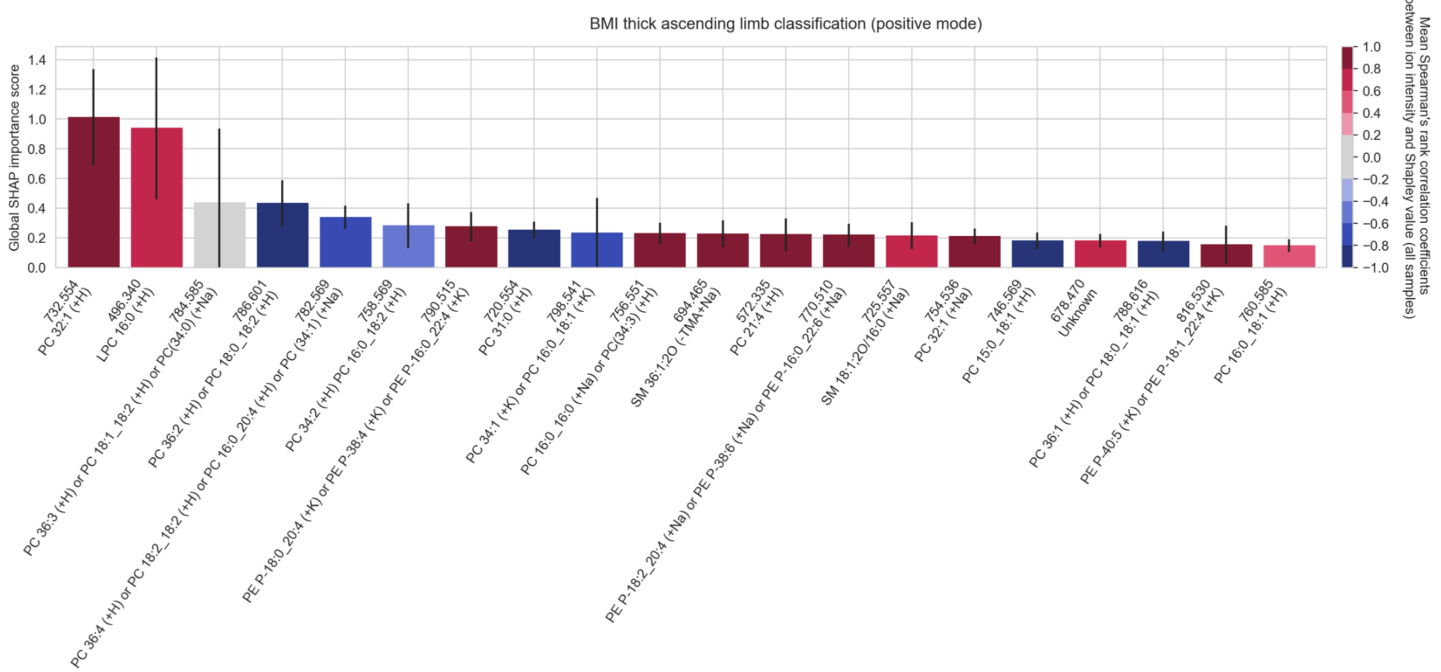

**Figure S190.** The top twenty cohort-wide (absolute) SHAP importance scores reporting the top twenty molecular species among the measured lipid species that enable the classification of **the thick ascending limb of obese donors from the thick ascending limb of normal donors, in positive ion mode**. The error bar is the standard deviation across all donor samples of the tissue sample-wide SHAP importance score for a given molecular species. The color of the bar reports the mean Spearman's rank correlation coefficient (across all donor samples) between the molecular species' mean-centered ion intensity and its Shapley score. A positive Spearman's rank correlation coefficient indicates that high intensity of the molecular species correlates with obesity. Conversely, a negative Spearman's rank correlation coefficient indicates that low intensity of the molecular species correlates with obesity.

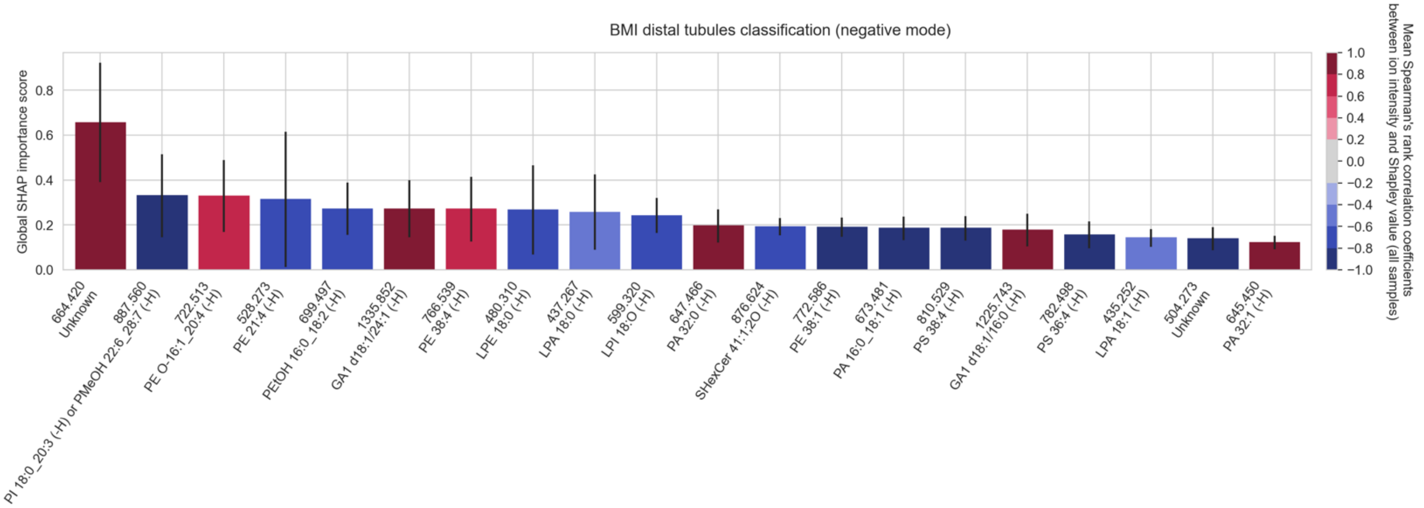

**Figure S191.** The top twenty cohort-wide (absolute) SHAP importance scores reporting the top twenty molecular species among the measured lipid species that enable the classification of **the distal tubules of obese donors from the distal tubules of normal donors, in negative ion mode**. The error bar is the standard deviation across all donor samples of the tissue sample-wide SHAP importance score for a given molecular species. The color of the bar reports the mean Spearman's rank correlation coefficient (across all donor samples) between the molecular species' mean-centered ion intensity and its Shapley score. A positive Spearman's rank correlation coefficient indicates that high intensity of the molecular species correlates with obesity. Conversely, a negative Spearman's rank correlation coefficient indicates that low intensity of the molecular species correlates with obesity.

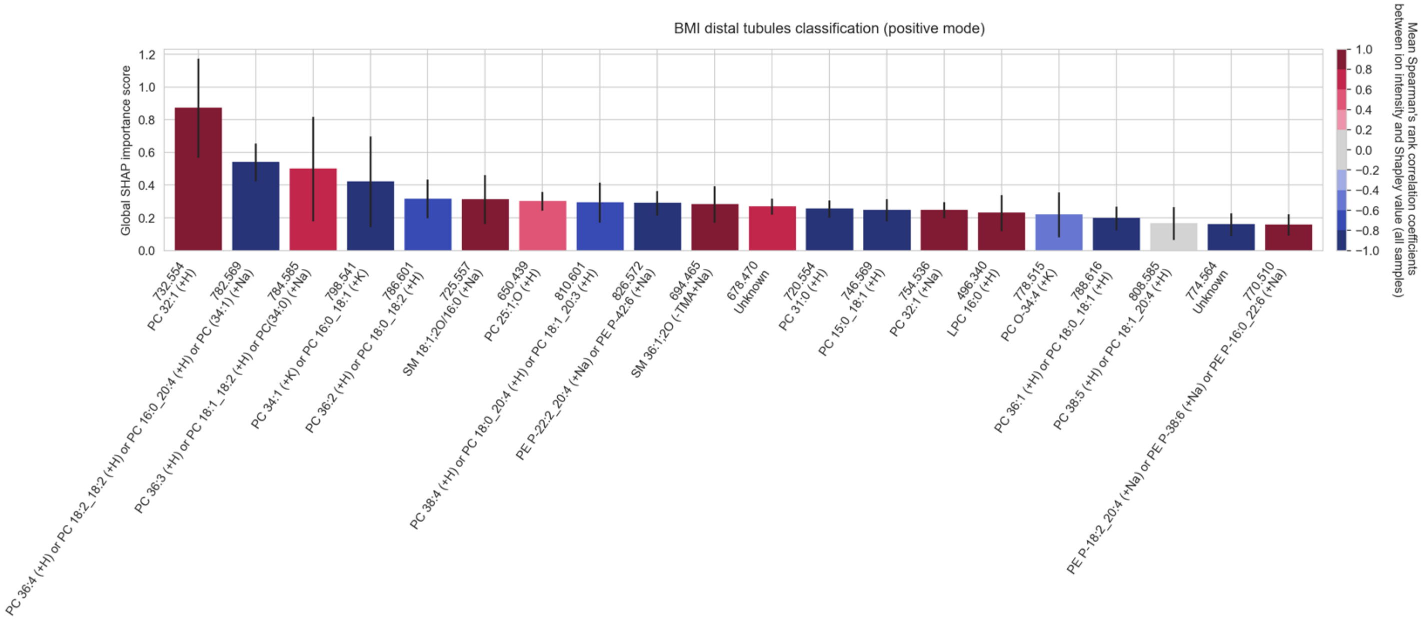

**Figure S192.** The top twenty cohort-wide (absolute) SHAP importance scores reporting the top twenty molecular species among the measured lipid species that enable the classification of **the distal tubules of obese donors from the distal tubules of normal donors, in positive ion mode**. The error bar is the standard deviation across all donor samples of the tissue sample-wide SHAP importance score for a given molecular species. The color of the bar reports the mean Spearman's rank correlation coefficient (across all donor samples) between the molecular species' mean-centered ion intensity and its Shapley score. A positive Spearman's rank correlation coefficient indicates that high intensity of the molecular species correlates with obesity. Conversely, a negative Spearman's rank correlation coefficient indicates that low intensity of the molecular species correlates with obesity.

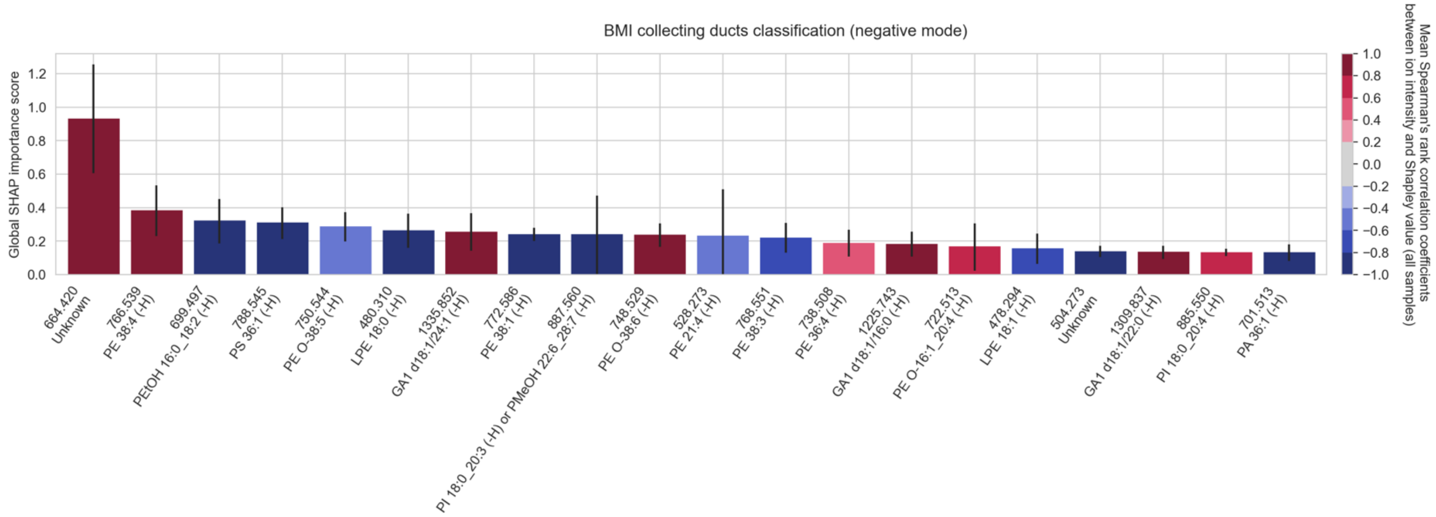

**Figure S193.** The top twenty cohort-wide (absolute) SHAP importance scores reporting the top twenty molecular species among the measured lipid species that enable the classification of **the collecting ducts of obese donors from the collecting ducts of normal donors, in negative ion mode**. The error bar is the standard deviation across all donor samples of the tissue sample-wide SHAP importance score for a given molecular species. The color of the bar reports the mean Spearman's rank correlation coefficient (across all donor samples) between the molecular species' mean-centered ion intensity and its Shapley score. A positive Spearman's rank correlation coefficient indicates that high intensity of the molecular species correlates with obesity. Conversely, a negative Spearman's rank correlation coefficient indicates that low intensity of the molecular species correlates with obesity.

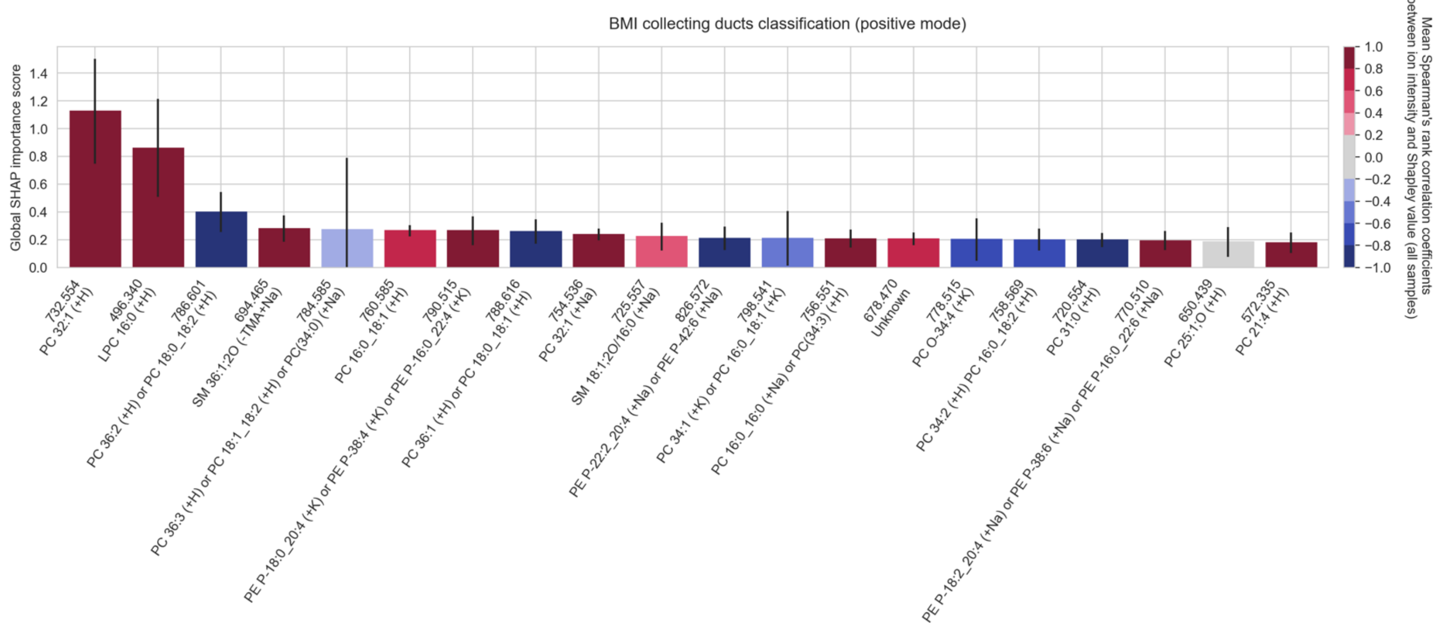

**Figure S194.** The top twenty cohort-wide (absolute) SHAP importance scores reporting the top twenty molecular species among the measured lipid species that enable the classification of **the collecting ducts of obese donors from the collecting ducts of normal donors, in positive ion mode**. The error bar is the standard deviation across all donor samples of the tissue sample-wide SHAP importance score for a given molecular species. The color of the bar reports the mean Spearman's rank correlation coefficient (across all donor samples) between the molecular species' mean-centered ion intensity and its Shapley score. A positive Spearman's rank correlation coefficient indicates that high intensity of the molecular species correlates with obesity. Conversely, a negative Spearman's rank correlation coefficient indicates that low intensity of the molecular species correlates with obesity.

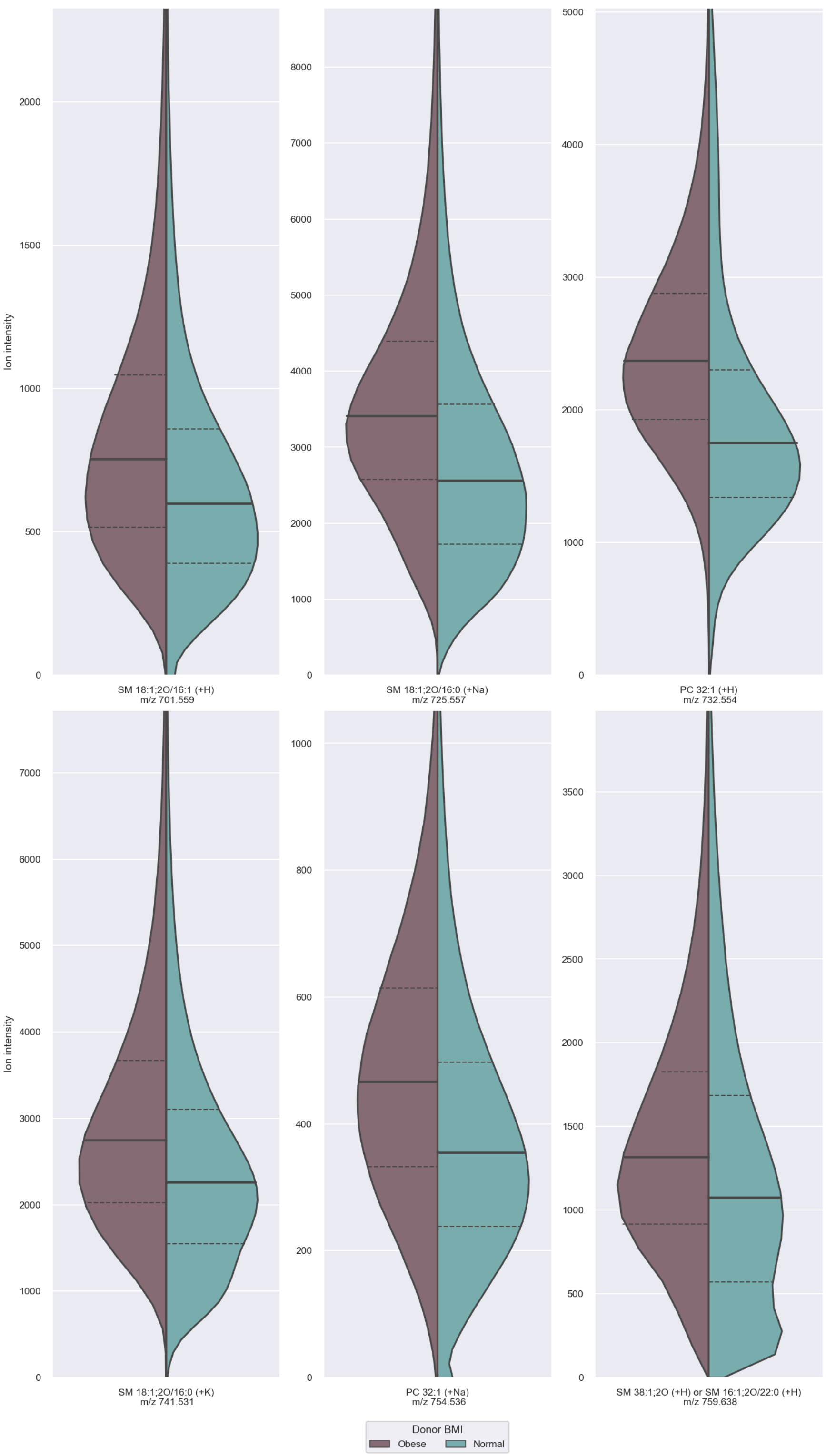

**Figure S195.** Split violin plots of the ion intensity distributions of the **top obesity biomarker candidates in obese (left) and normal (right) donor data, in positive ion mode**, approximated using kernel density estimation. The violin plots are cropped at the 99th percentile of the distribution of one of the two classes (whichever is larger) to facilitate visual comparison. The full line indicates the median of each class' distribution, whereas the dashed lines indicate its interquartile range.

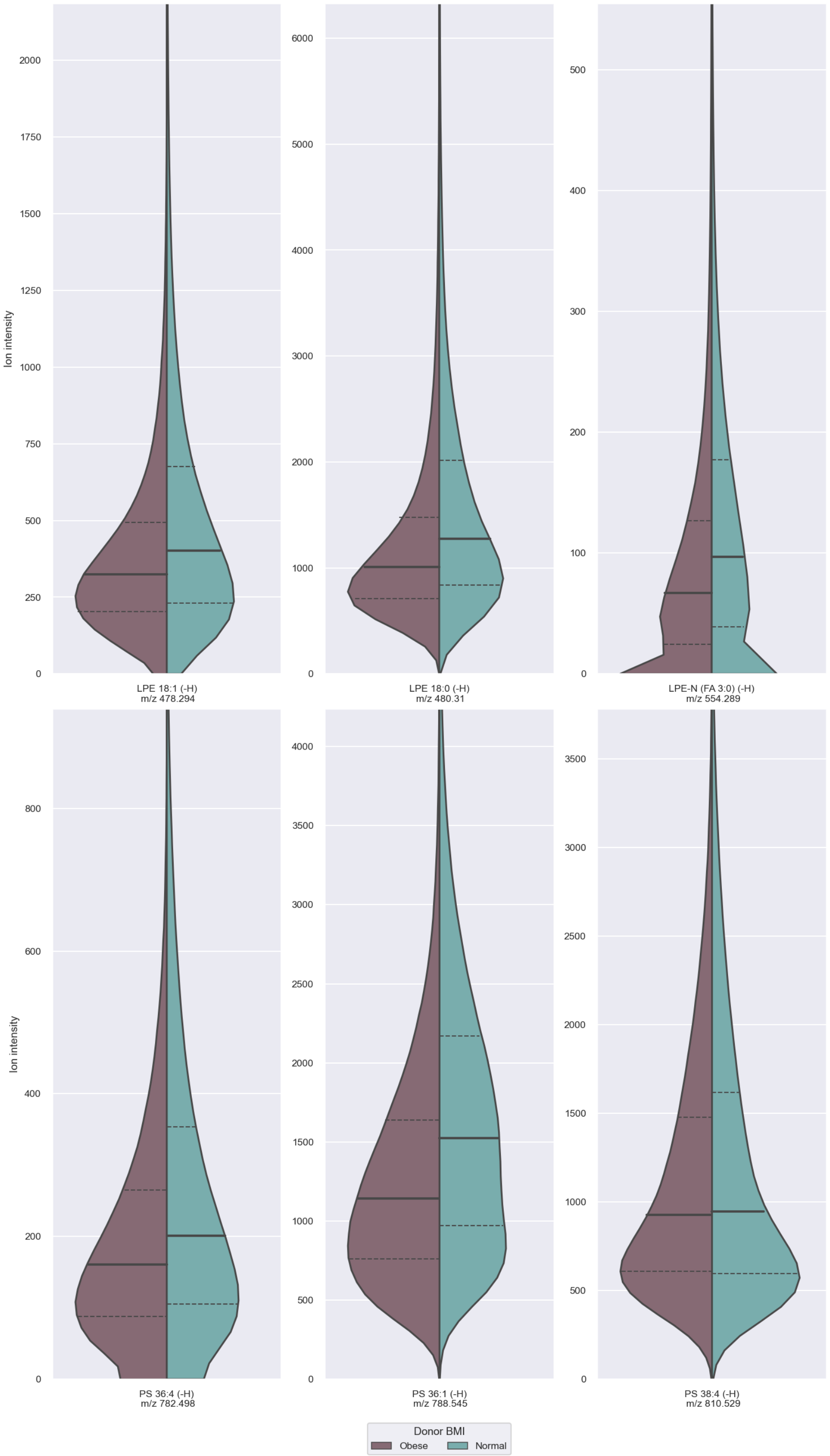

**Figure S196.** Split violin plots of the ion intensity distributions of the **top obesity biomarker candidates in obese (left) and normal (right) donor data, in negative ion mode**, approximated using kernel density estimation. The violin plots are cropped at the 99th percentile of the distribution of one of the two classes (whichever is larger) to facilitate visual comparison. The full line indicates the median of each class' distribution, whereas the dashed lines indicate its interquartile range.

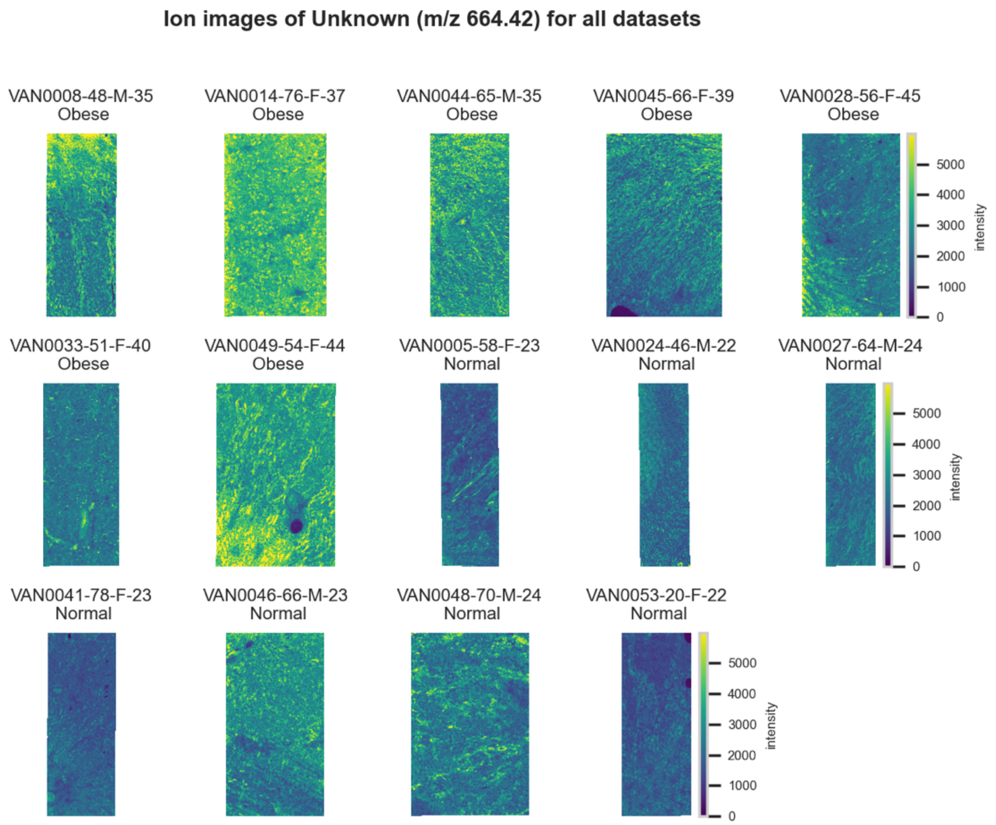

**Figure S197.** 10 µm MALDI IMS images of an unknown (*m/z* 664.42) detected in negative ion mode from each donor sample. Multivariate SHAP analysis determined this to be feature 1 for classification of obesity.

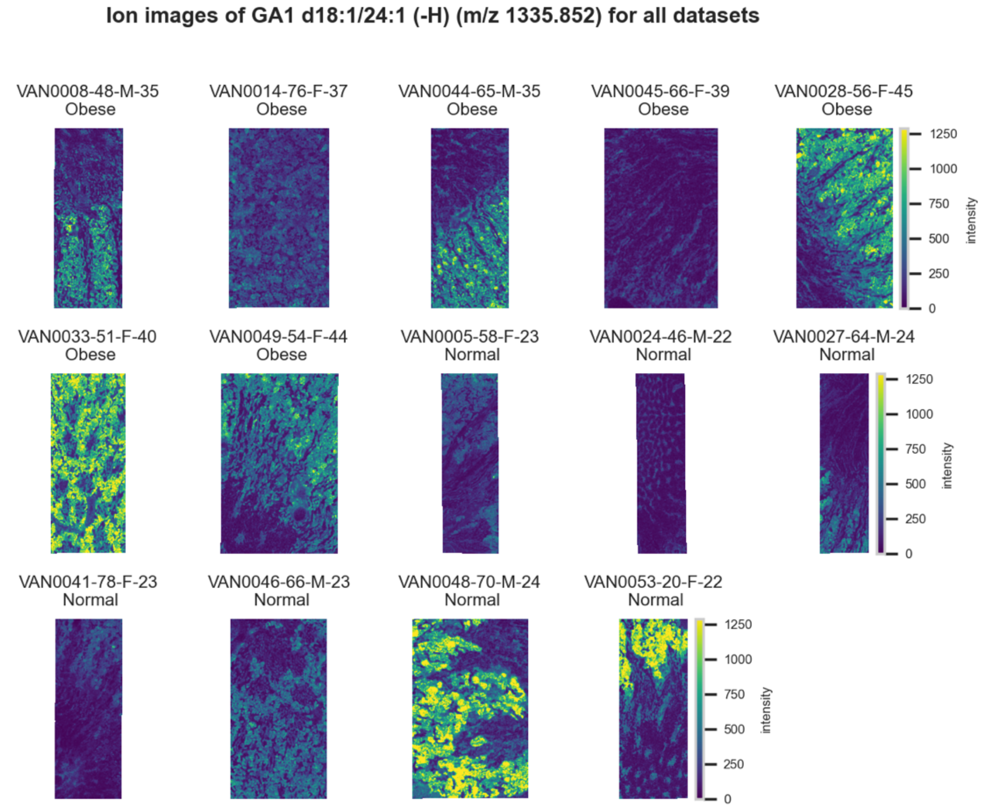

**Figure S198.** 10 µm MALDI IMS images of $[GA1(d18:1/24:1)-H]^-$ (*m/z* 1,335.852) detected in negative ion mode from each donor sample. Multivariate SHAP analysis determined this to be feature 2 for classification of obesity.

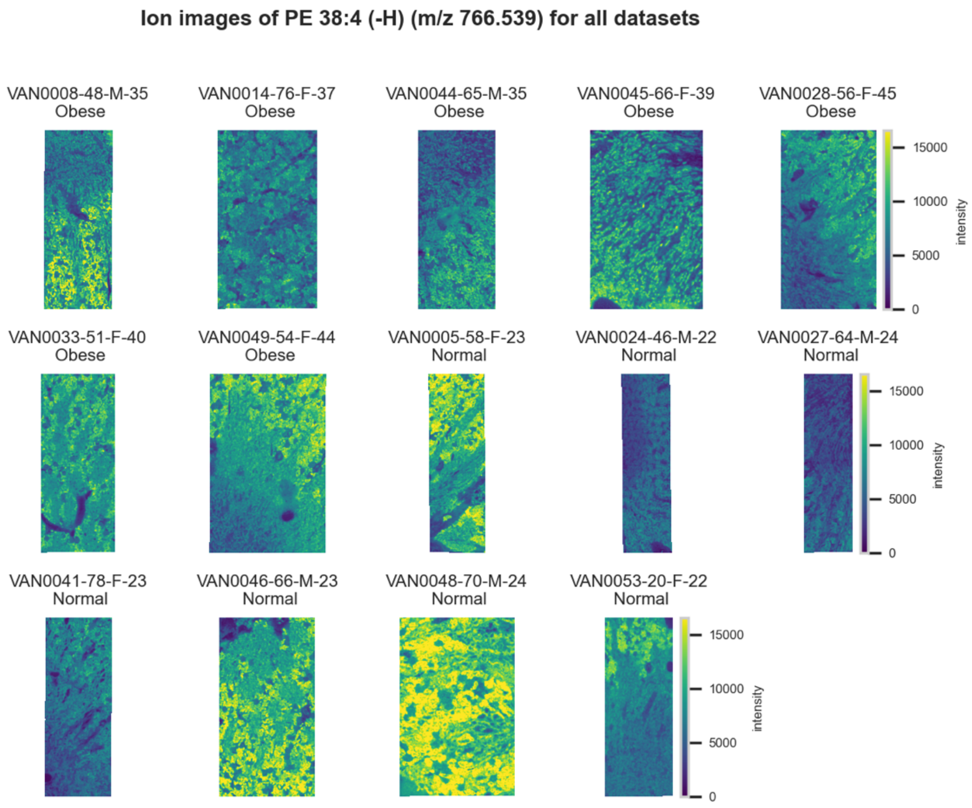

**Figure S199.** 10 µm MALDI IMS images of [PE(38:4)-H]$^-$ (*m/z* 766.539) detected in <u>negative ion mode</u> from each donor sample. Multivariate SHAP analysis determined this to be feature 3 for classification of obesity.

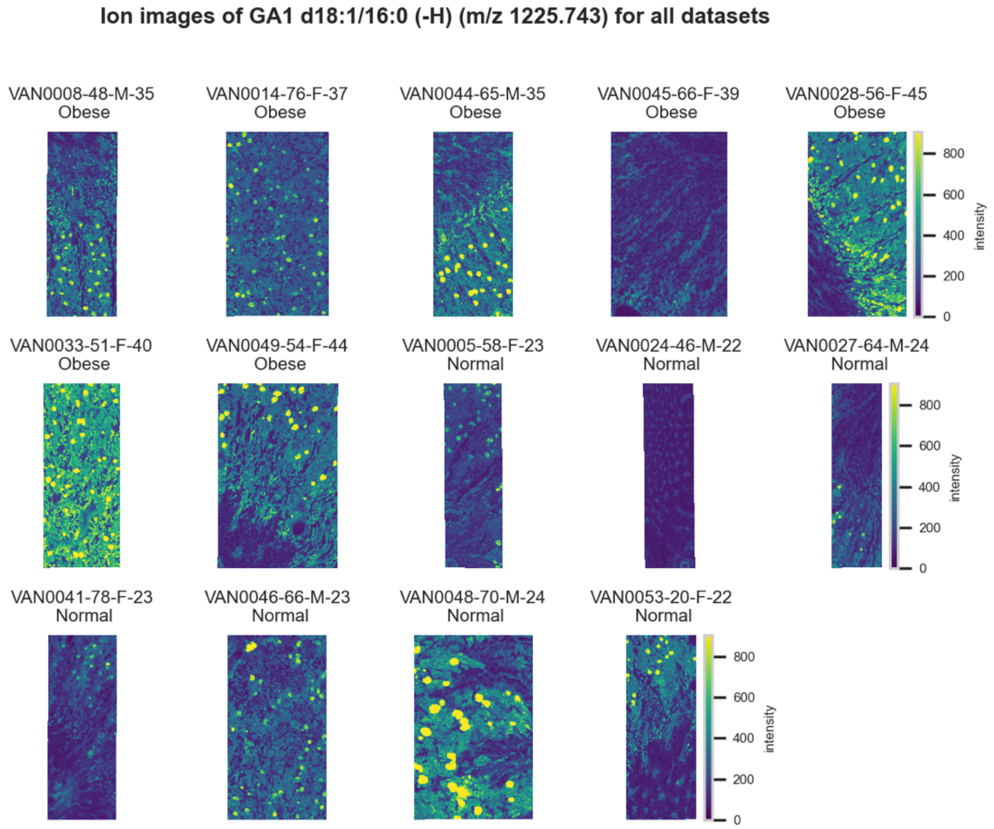

**Figure S200.** 10 µm MALDI IMS images of an unknown [GA1(d18:1/16:0)-H]$^-$ (*m/z* 1,225.743) detected in <u>negative ion mode</u> from each donor sample. Multivariate SHAP analysis determined this to be feature 4 for classification of obesity.

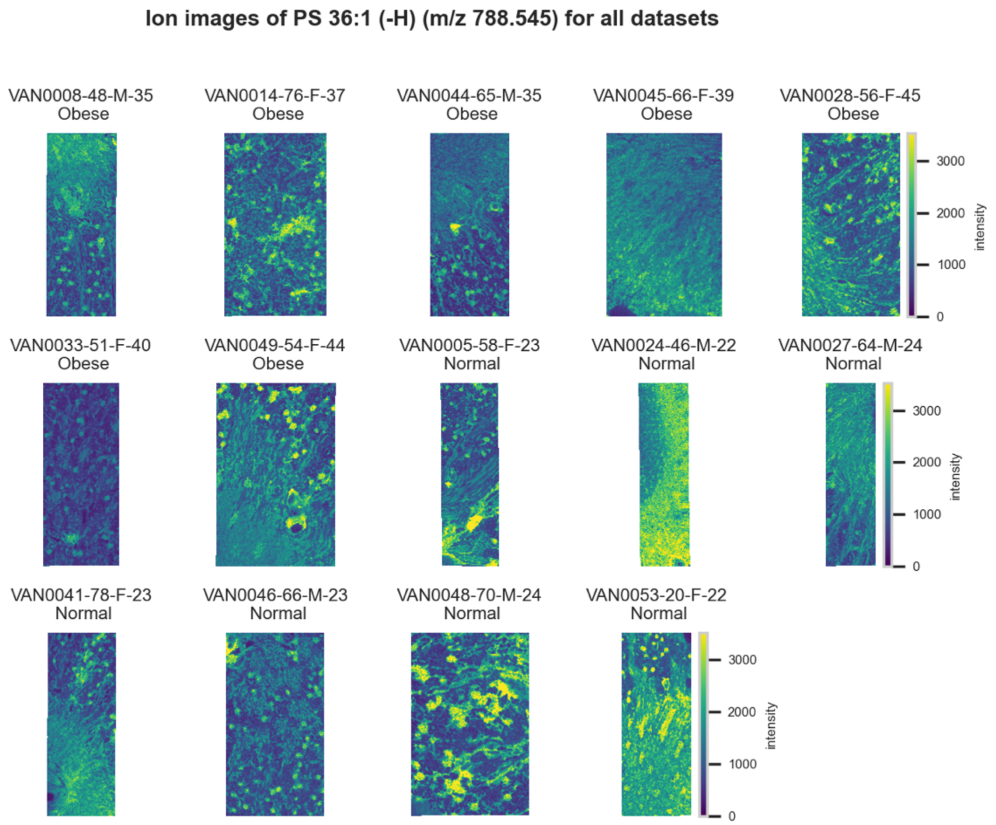

**Figure S201.** 10 µm MALDI IMS images of [PS(36:1)-H]$^-$ (*m/z* 788.545) detected in <u>negative ion mode</u> from each donor sample. Multivariate SHAP analysis determined this to be feature 5 for classification of obesity.

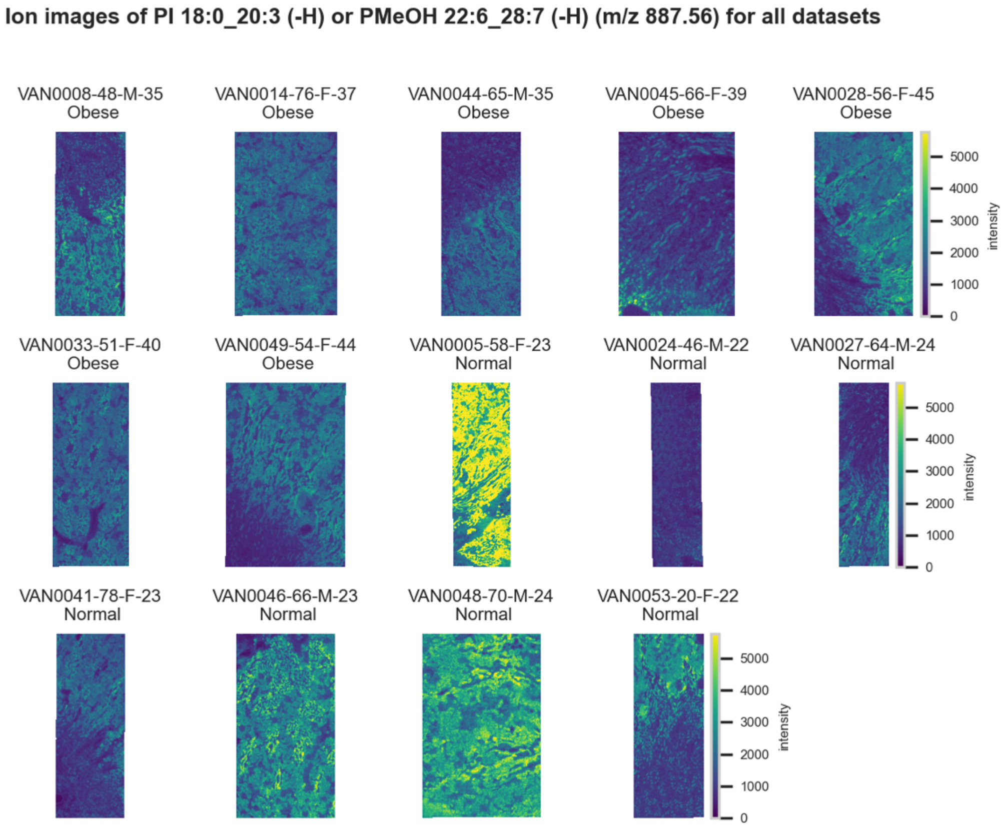

**Figure S202.** 10 µm MALDI IMS images of [PI(18:0_20:3)-H]$^-$ or [PMeOH(22:6_28:7)-H]$^-$ (*m/z* 887.56) detected in <u>negative ion mode</u> from each donor sample. Multivariate SHAP analysis determined this to be feature 6 for classification of obesity.

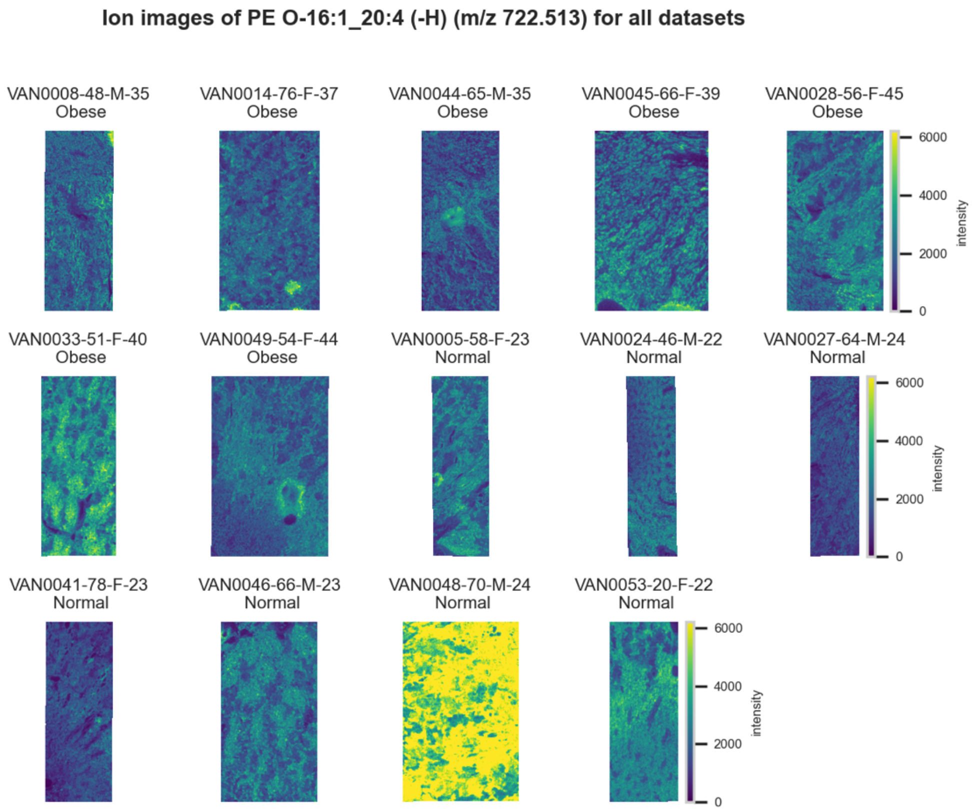

**Figure S203.** 10 µm MALDI IMS images of $[PE(O\text{-}16:1\_20:4)\text{-}H]^-$ (*m/z* 722.513) detected in negative ion mode from each donor sample. Multivariate SHAP analysis determined this to be feature 7 for classification of obesity.

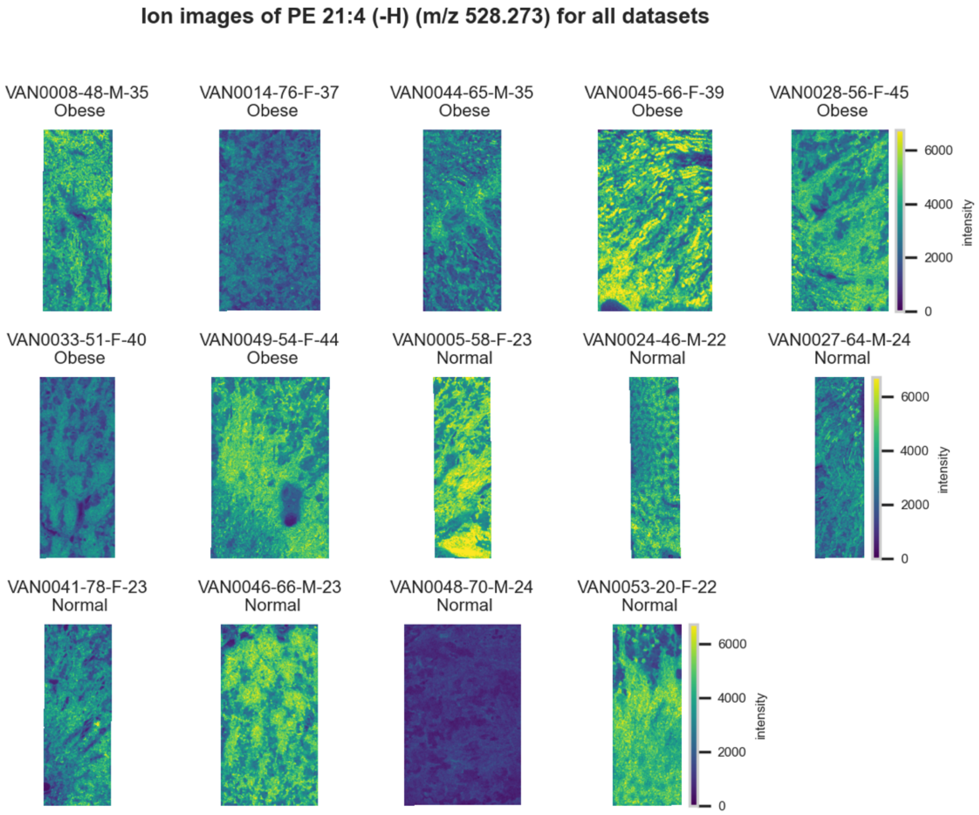

**Figure S204.** 10 µm MALDI IMS images of $[PE(21:4)\text{-}H]^-$ (*m/z* 528.273) detected in negative ion mode from each donor sample. Multivariate SHAP analysis determined this to be feature 8 for classification of obesity.

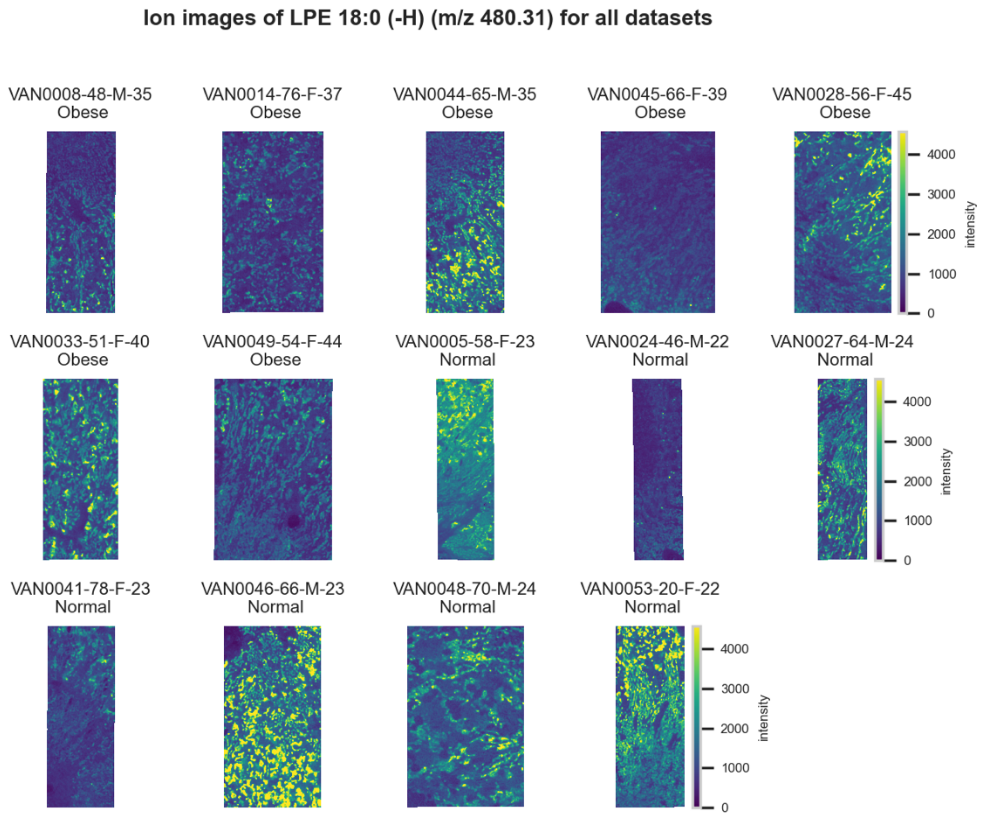

**Figure S205.** 10 µm MALDI IMS images of [LPE(18:0)-H]$^-$ (*m/z* 480.31) detected in negative ion mode from each donor sample. Multivariate SHAP analysis determined this to be feature 9 for classification of obesity.

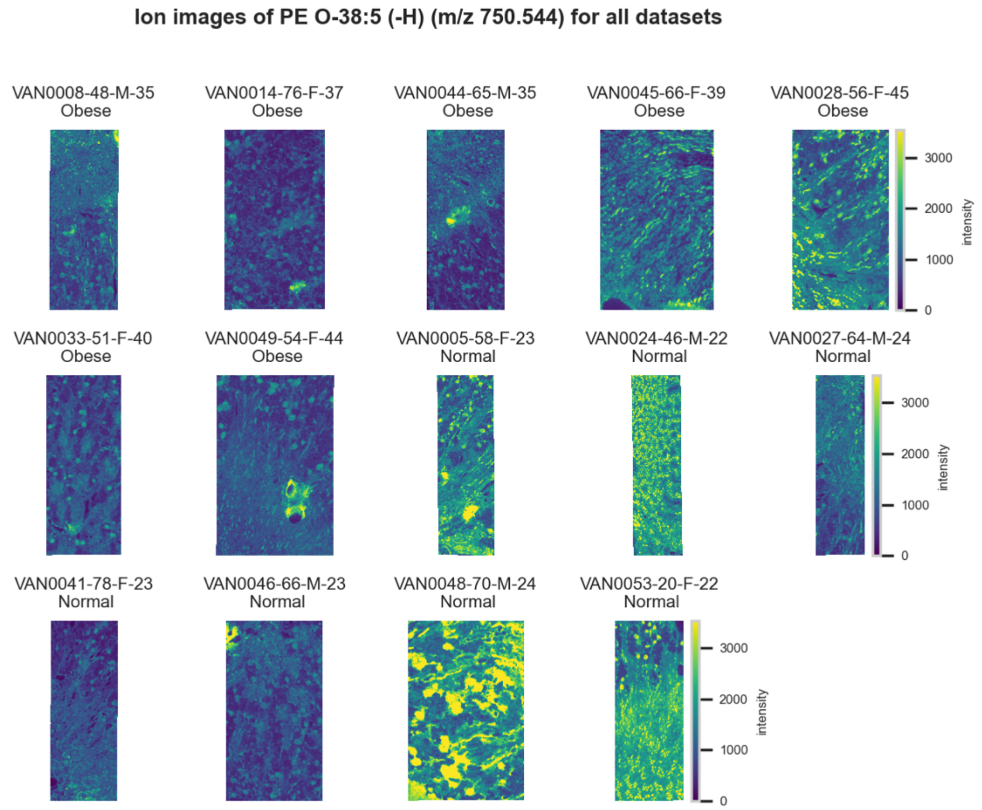

**Figure S206.** 10 µm MALDI IMS images of [PE(O-38:5)-H]$^-$ (*m/z* 750.544) detected in negative ion mode from each donor sample. Multivariate SHAP analysis determined this to be feature 10 for classification of obesity.

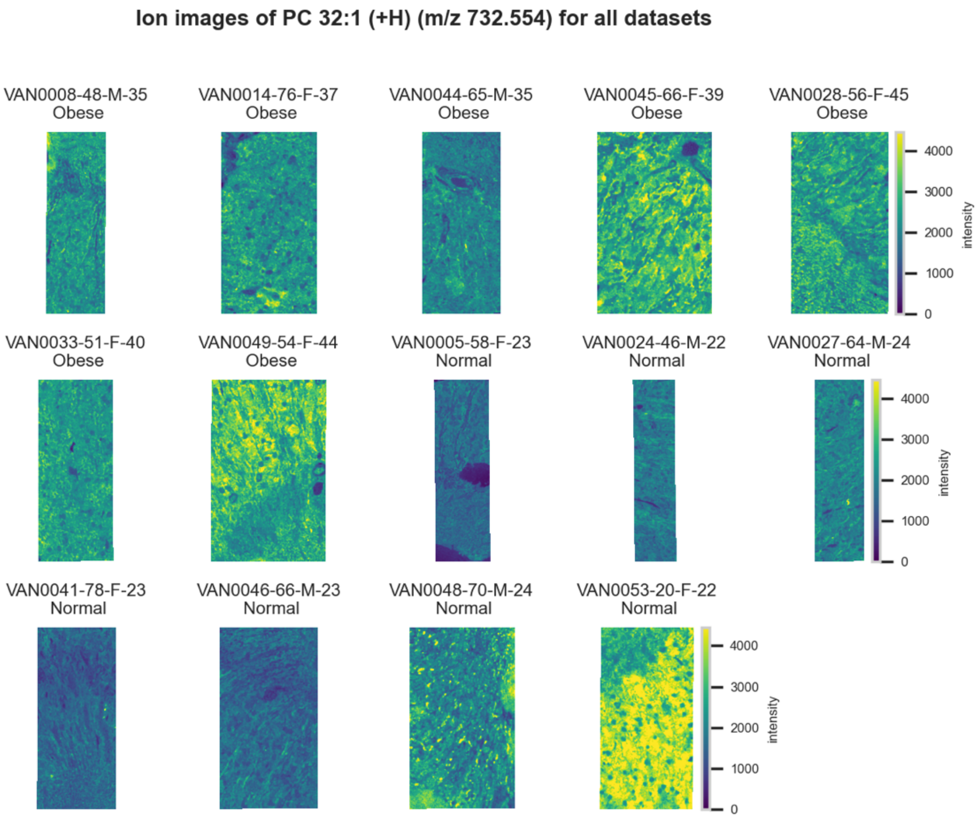

**Figure S207.** 10 µm MALDI IMS images of $[PC(32{:}1)+H]^+$ (*m/z* 732.554) detected in positive ion mode from each donor sample. Multivariate SHAP analysis determined this to be feature 1 for classification of obesity.

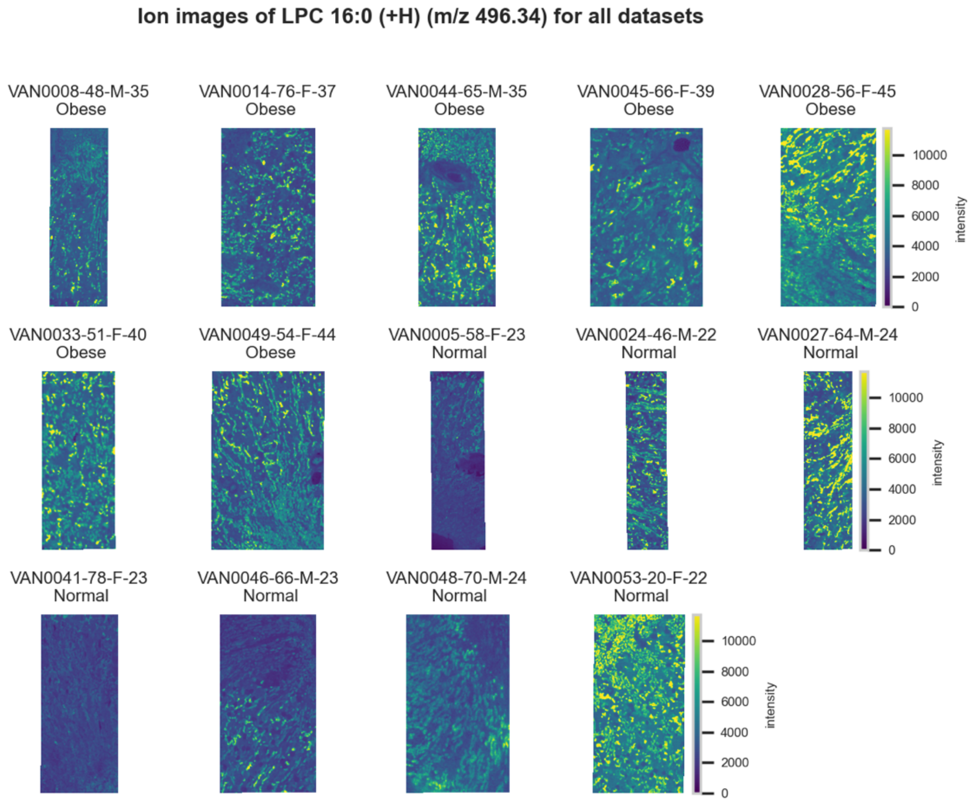

**Figure S208.** 10 µm MALDI IMS images of $[LPC(16{:}0)+H]^+$ (*m/z* 496.34) detected in positive ion mode from each donor sample. Multivariate SHAP analysis determined this to be feature 2 for classification of obesity.

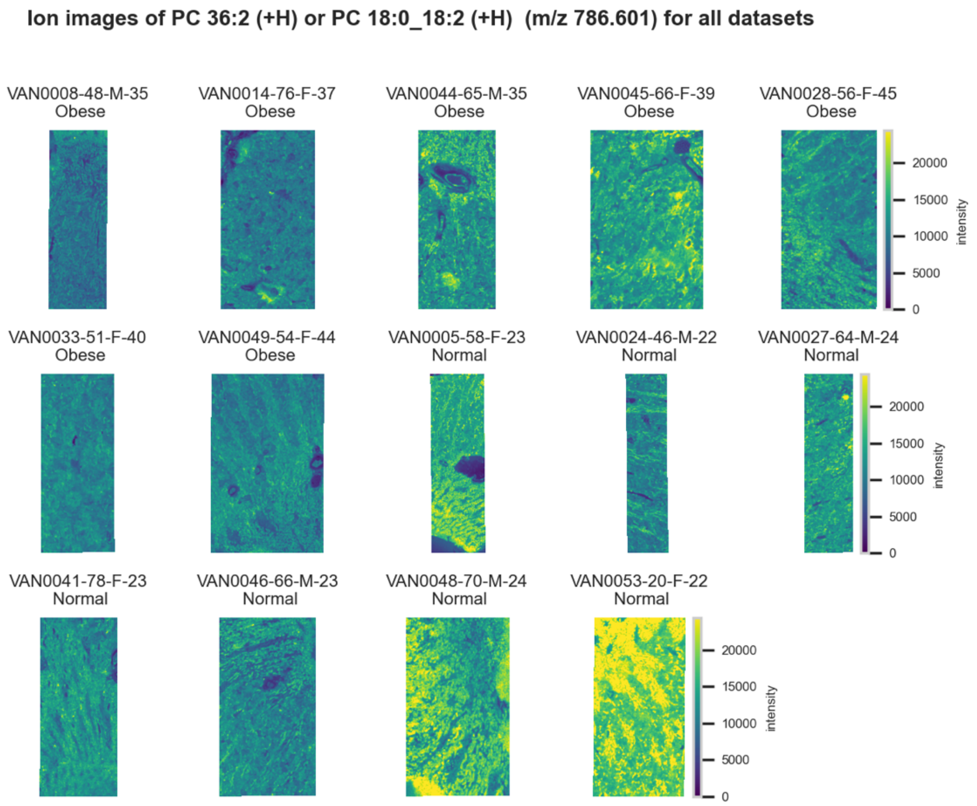

**Figure S209.** 10 µm MALDI IMS images of $[PC(36:2)+H]^+$ or $[PC(18:0\_18:2)+H]^+$ (*m/z* 786.601) detected in positive ion mode from each donor sample. These molecules are isomers and therefore indistinguishable by MS alone. Multivariate SHAP analysis determined this to be feature 3 for classification of obesity.

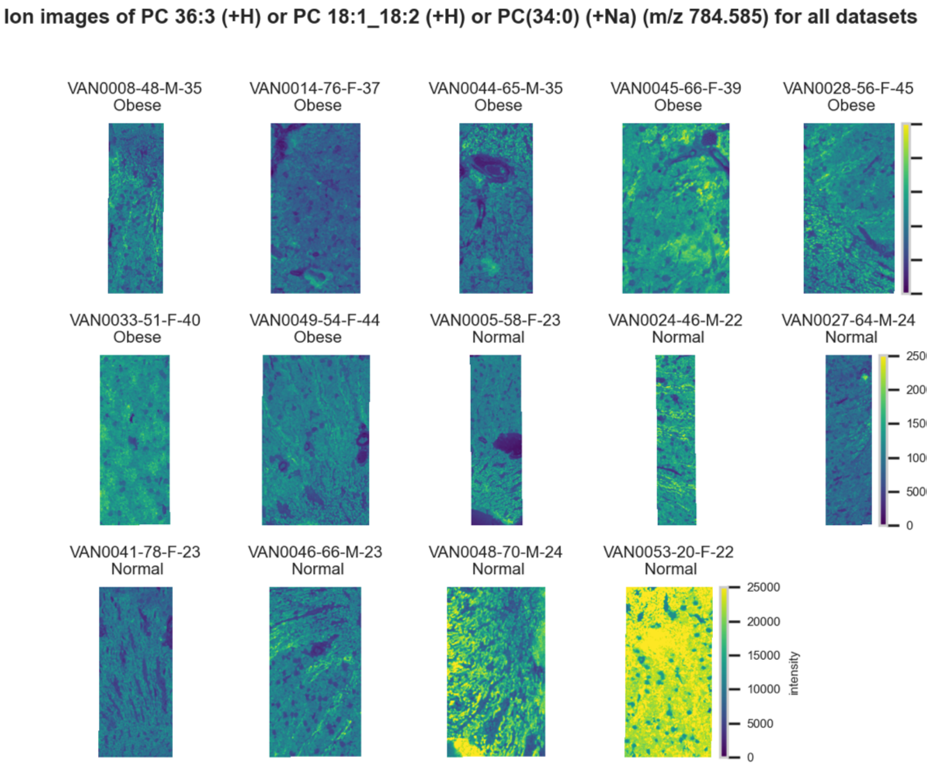

**Figure S210.** 10 µm MALDI IMS images of an unknown $[PC(36:3)+H]^+$ or $[PC(18:1\_18:2)+H]^+$ or $[PC(34:0)+Na]^+$ (*m/z* 784.585) detected in positive ion mode from each donor sample. These molecules are isomers and therefore indistinguishable by MS alone. Multivariate SHAP analysis determined this to be feature 4 for classification of obesity.

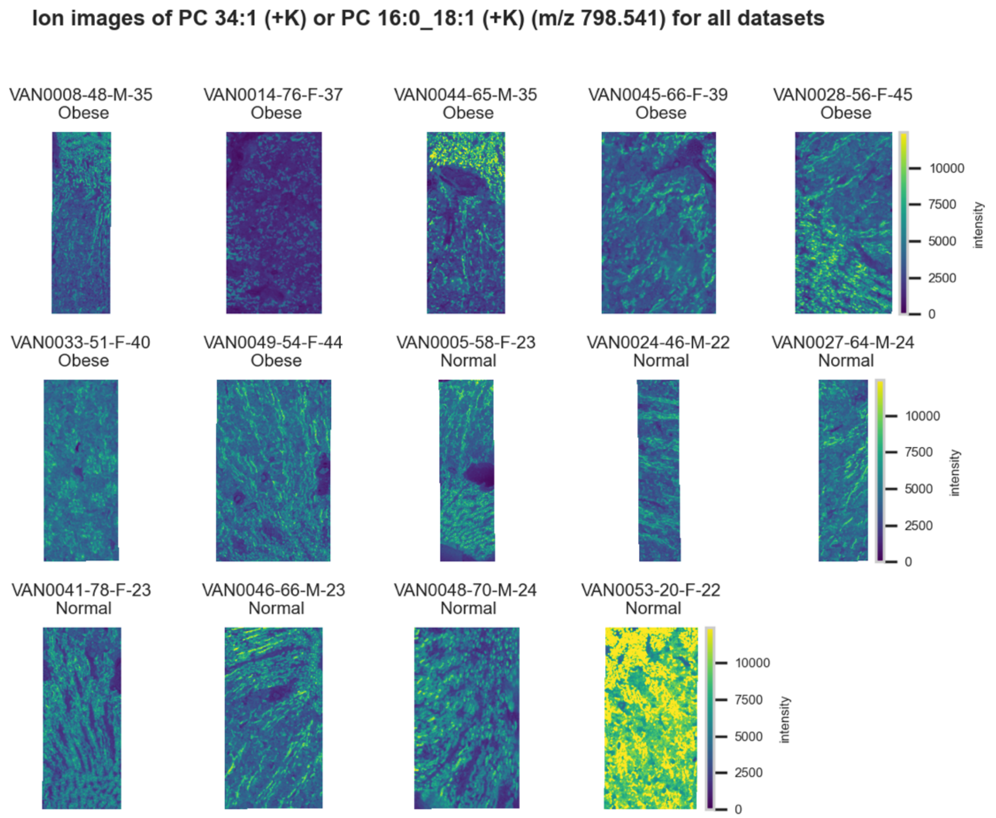

**Figure S211.** 10 µm MALDI IMS images of $[PC(34:1)+K]^+$ or $[PC(16:0\_18:1)+K]^+$ (*m/z* 798.541) detected in <u>positive ion mode</u> from each donor sample. These molecules are isomers and therefore indistinguishable by MS alone. Multivariate SHAP analysis determined this to be feature 5 for classification of obesity.

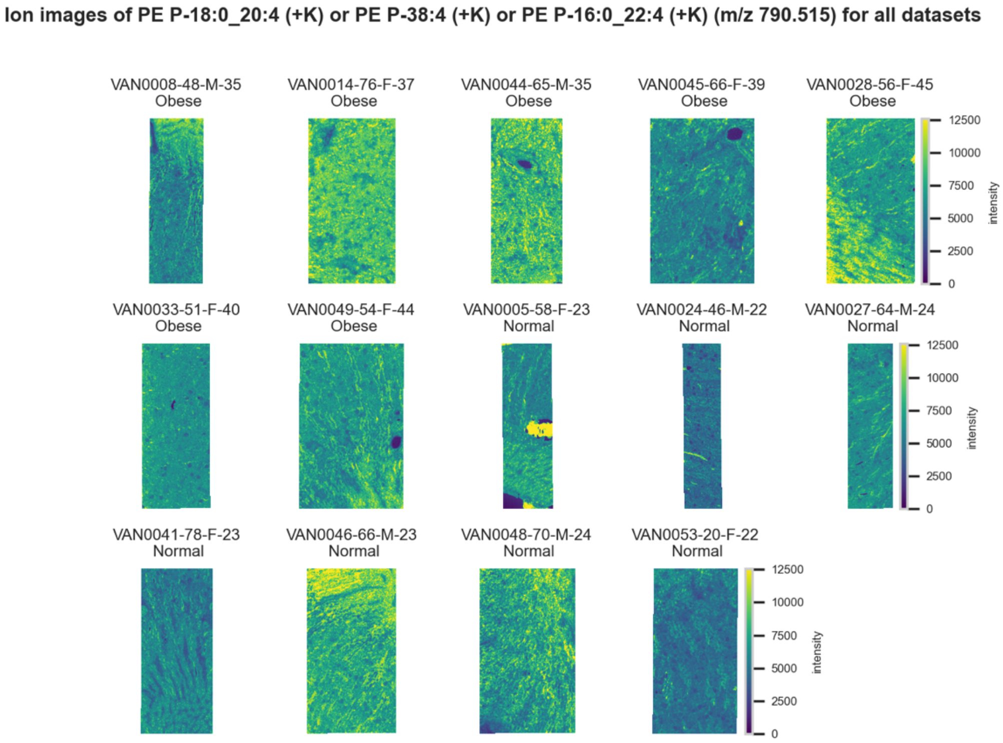

**Figure S212.** 10 µm MALDI IMS images of $[PE(P\text{-}38:4)+K]^+$ or $[PE(P\text{-}18:0\_20:4)+K]^+$ or $[PE(P\text{-}16:0\_22:4)+K]^+$ (*m/z* 790.515) detected in <u>positive ion mode</u> from each donor sample. These molecules are isomers and therefore indistinguishable by MS alone. Multivariate SHAP analysis determined this to be feature 6 for classification of obesity.

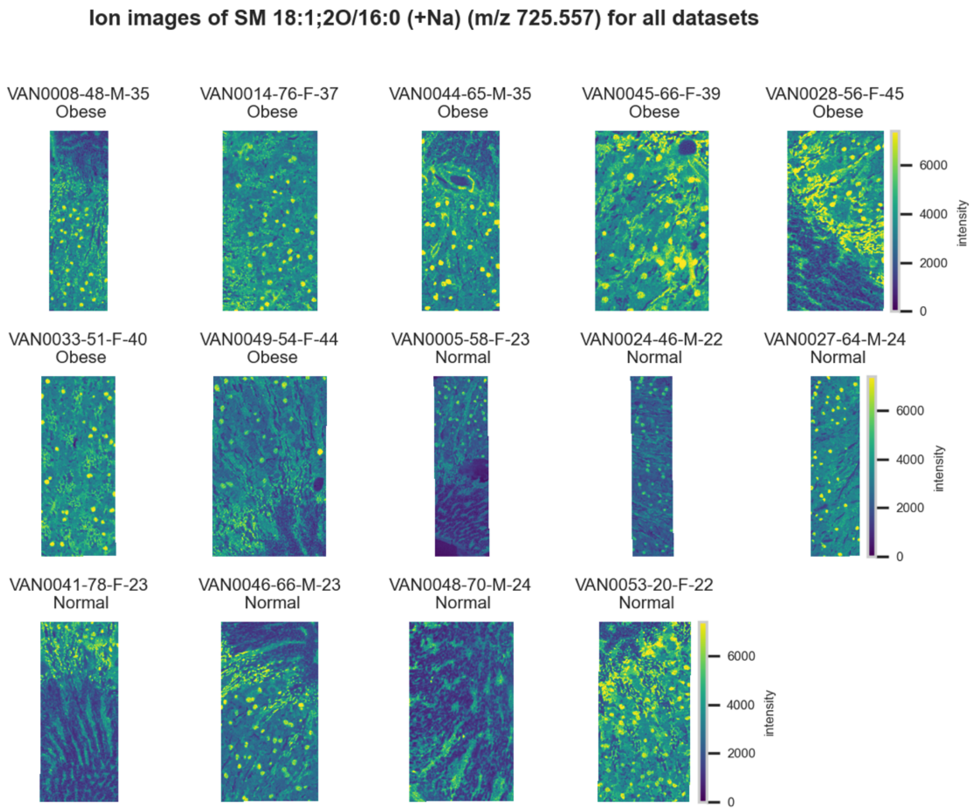

**Figure S213.** 10 µm MALDI IMS images of $[SM(18:1,2O/16:0)+Na]^+$ (*m/z* 725.557) detected in positive ion mode from each donor sample. Multivariate SHAP analysis determined this to be feature 7 for classification of obesity.

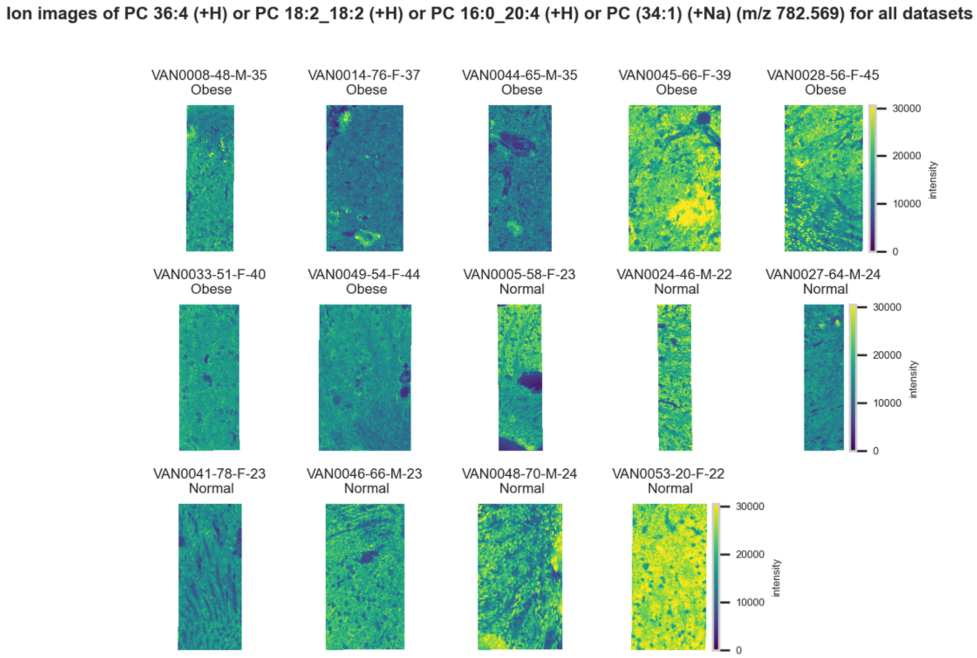

**Figure S214.** 10 µm MALDI IMS images of $[PC(36:4)+H]^+$ or $[PC(16:0\_20:4)+H]^+$ or $[PC(18:2\_18:2)+H]^+$ or $[PC(34:1)+Na]^+$ (*m/z* 782.569) detected in positive ion mode from each donor sample. These molecules are isomers or close isobars and therefore indistinguishable by MS alone. Multivariate SHAP analysis determined this to be feature 8 for classification of obesity.

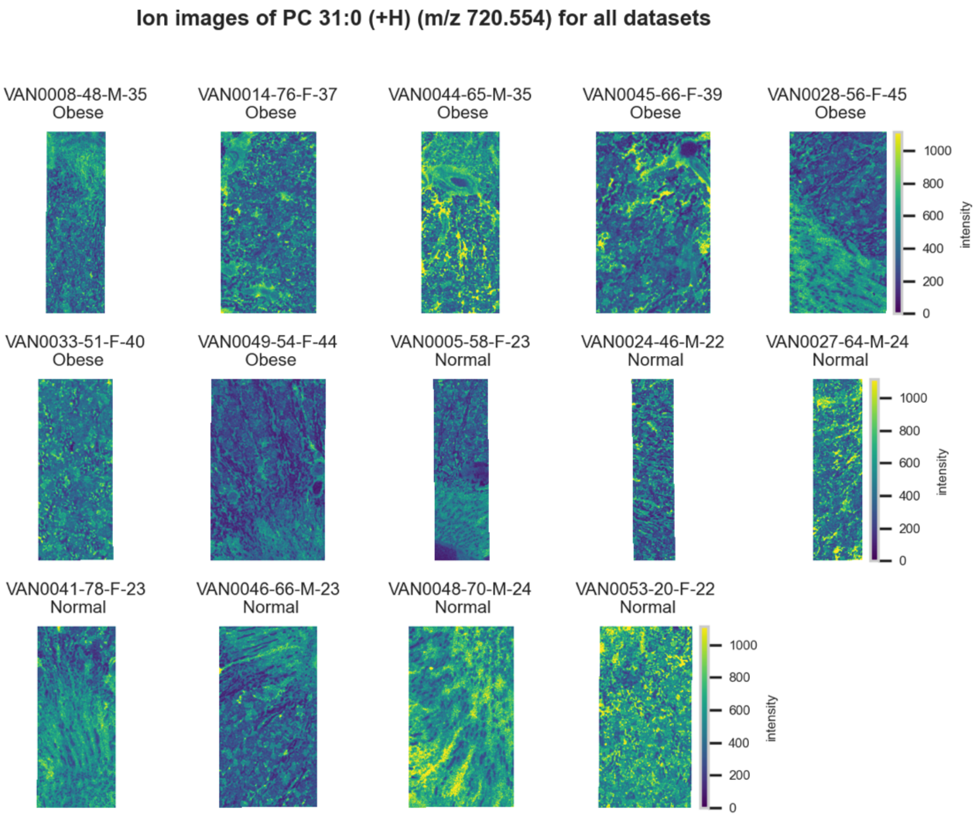

**Figure S215.** 10 µm MALDI IMS images of $[PC(31:0)+H]^+$ (*m/z* 720.554) detected in positive ion mode from each donor sample. These molecules are isomers and therefore indistinguishable by MS alone. Multivariate SHAP analysis determined this to be feature 9 for classification of obesity.

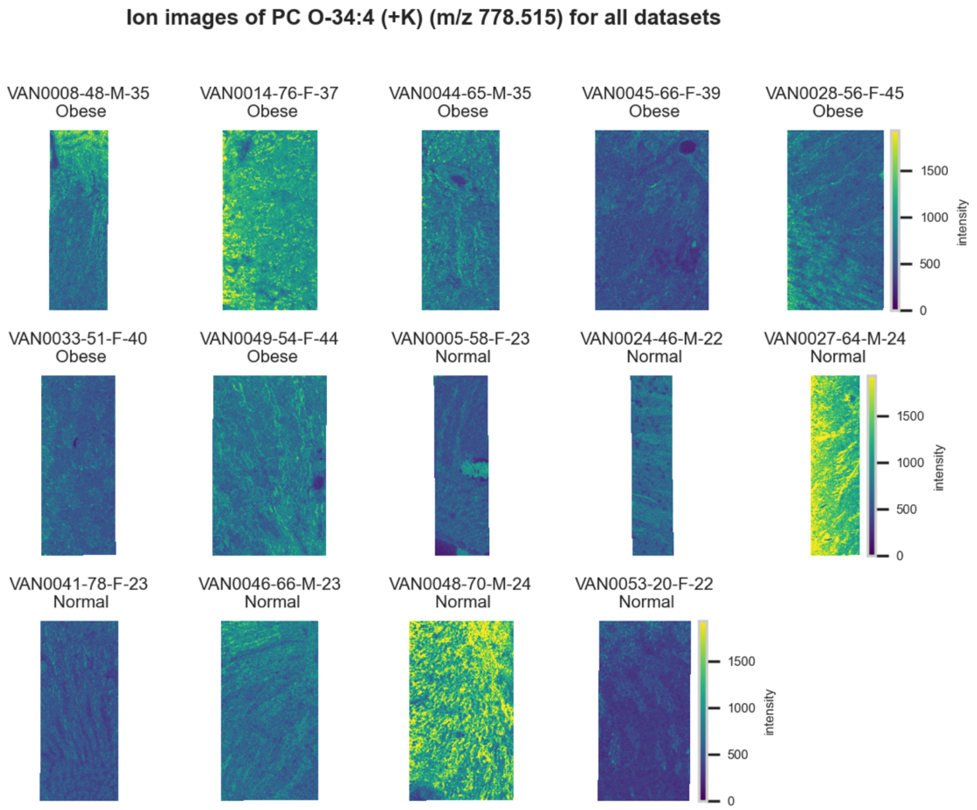

**Figure S216.** 10 µm MALDI IMS images of $[PC(O\text{-}34:4)+K]^+$ (*m/z* 778.515) detected in positive ion mode from each donor sample. Multivariate SHAP analysis determined this to be feature 10 for classification of obesity.
